# The effect of dopamine transporter blockade on optical self-stimulation: behavioral and computational evidence for parallel processing in brain reward circuitry

**DOI:** 10.1101/867481

**Authors:** Ivan Trujillo-Pisanty, Kent Conover, Pavel Solis, Daniel Palacios, Peter Shizgal

## Abstract

The neurobiological study of reward was launched by the discovery of intracranial self-stimulation (ICSS). Subsequent investigation of this phenomenon provided the initial link between reward-seeking behavior and dopaminergic neurotransmission. We re-evaluated this relationship by psychophysical, pharmacological, optogenetic, and computational means. In rats working for direct, optical activation of midbrain dopamine neurons, we varied the strength and opportunity cost of the stimulation and measured time allocation, the proportion of trial time devoted to reward pursuit. We found that the dependence of time allocation on the strength and cost of stimulation was similar formally to that observed when electrical stimulation of the medial forebrain bundle served as the reward. When the stimulation is strong and cheap, the rats devote almost all their time to reward pursuit; time allocation falls off as stimulation strength is decreased and/or its opportunity cost is increased. A 3D plot of time allocation versus stimulation strength and cost produces a surface resembling the corner of a plateau (the “reward mountain”). We show that dopamine-transporter blockade shifts the mountain along both the strength and cost axes in rats working for optical activation of midbrain dopamine neurons. In contrast, the same drug shifted the mountain uniquely along the opportunity-cost axis when rats worked for electrical MFB stimulation in a prior study. Dopamine neurons are an obligatory stage in the dominant model of ICSS, which positions them at a key nexus in the final common path for reward seeking. This model fails to provide a cogent account for the differential effect of dopamine transporter blockade on the reward mountain. Instead, we propose that midbrain dopamine neurons and neurons with non-dopaminergic, MFB axons constitute parallel limbs of brain-reward circuitry that ultimately converge on the final-common path for the evaluation and pursuit of rewards.

**Author summary:** To succeed in the struggle for survival and reproductive success, animals must make wise choices about which goals to pursue and how much to pay to attain them. How does the brain make such decisions and adjust behaviour accordingly? An animal model that has long served to address this question entails delivery of rewarding brain stimulation. When the probe is positioned appropriately in the brain, rats will work indefatigably to trigger such stimulation. Dopamine neurons play a crucial role in this phenomenon. The dominant model of the brain circuitry responsible for the reward-seeking behavior treats these cells as a gateway through which the reward-generating brain signals must pass. Here, we challenge this idea on the basis of an experiment in which the dopamine neurons were activated selectively and directly. Mathematical modeling of the results argues for a new view of the structure of brain reward circuitry. On this view, the pathway(s) in which the dopamine neurons are embedded is one of a set of parallel channels that process reward signals in the brain. To achieve a full understanding of how goals are evaluated, selected and pursued, the full set of channels must be identified and investigated.

## Introduction

We and our cohabitants are fortunate winners. We have enjoyed reproductive success due to a combination of luck and an array of skills among which acumen in cost/benefit decision making is of paramount importance. Rudimentary ability in this domain can be implemented simply, as is evident from the behavior of animals whose nervous systems comprise only hundreds of neurons [1]. In the multi-million-cell nervous systems of mammals, the foundations of more sophisticated cost/benefit decision making are thought to have been heavily conserved [2, 3]. If so, the rodent species so widely studied in neurobiological laboratories are equipped with variants of decision-making circuitry that continues to shape our own choices and actions.

A seminal moment in the study of the neural foundations of cost/benefit decision making was the discovery that rats would work vigorously and indefatigably for focal electrical stimulation of sites in the basal forebrain and midbrain [4]. Despite the artificial spatiotemporal distribution of the evoked neural activity, the rats behaved as procuring a highly valuable, natural goal object, such as energy-rich food. This striking phenomenon, dubbed “intracranial self-stimulation” (ICSS), has been investigated subsequently by means of perturbational, pharmacological, correlational, computational, and behavioral methods that have seen dramatic recent improvements in precision, specificity, and power. For example, the electrical stimulation employed originally activates neurons near the electrode tip with relatively little specificity, whereas contemporary optogenetic methods restrict activation to genetically defined neural populations. Currently employed pharmacological agents are far more selective than the drugs employed in the early studies. Whereas the crude response counts used initially to measure behavioral output are confounded by inherent non-linearity as well as by disruptive side-effects of drugs and motoric activation, contemporary psychophysical methods support inference of the strength and subjective cost of the induced reward as well as the form and parameters of the functions that map observable inputs and outputs into the variables that determine behavioral-allocation decisions. In the current study, we combine, for the first time, direct, specific optical activation of midbrain dopamine neurons, modulation of dopaminergic neurotransmission with a highly selective dopamine-reuptake blocker, psychophysical inference of benefits and costs, computational modeling of the processes that intervene between the optical activation of the dopamine neurons and the consequent behavioral output, and simulation of model output.

### The role of midbrain dopamine neurons in ICSS

Performance for rewarding brain stimulation has long been known to depend on dopaminergic neurotransmission. Drugs that boost dopamine signaling decrease the strength of the stimulation required to support a given level of ICSS, whereas drugs that decrease dopamine signaling necessitate a compensatory increase in stimulation strength at lower doses and eliminate responding at higher ones [5, 6]. It was believed initially that these effects are due to direct activation of dopaminergic neurons by the electrical stimulation. However, dopaminergic fibers are very fine and unmyelinated [7]. Consequently, they have very high thresholds to excitation by extracellular currents. Robust ICSS of sites along the medial forebrain bundle (MFB) is observed using stimulation parameters too weak to produce substantial recruitment of dopaminergic fibers [8]. Moreover, estimates of recovery of refractoriness, conduction velocity, and frequency-following fidelity in the directly activated fibers underlying the rewarding effect implicate fibers that are myelinated and far more excitable than those of dopamine neurons [9–14]. To reconcile these observations with the pharmacological evidence for dopaminergic modulation of ICSS, a “series-circuit” model was proposed [10–12, 15, 16]. According to this model, myelinated fibers of non-dopaminergic neurons transsynaptically activate midbrain dopamine neurons, thus generating the rewarding effect. This model of brain reward circuitry, which has remained virtually unchallenged for nearly forty years, fails to provide a cogent account for the new data and simulations we report here. We thus propose a fundamental revision.

### The roots of the current study

The roots of the current experiment on the rewarding effect produced by optical activation of midbrain dopamine neurons lie in earlier work in which electrical stimulation of the MFB served as the reward. We use the acronyms, eICSS and oICSS, to refer to operant performance for electrical and optical brain stimulation, respectively (Tab 1). Four lines of work on eICSS gave rise to the current oICSS study:

**Table 1.**
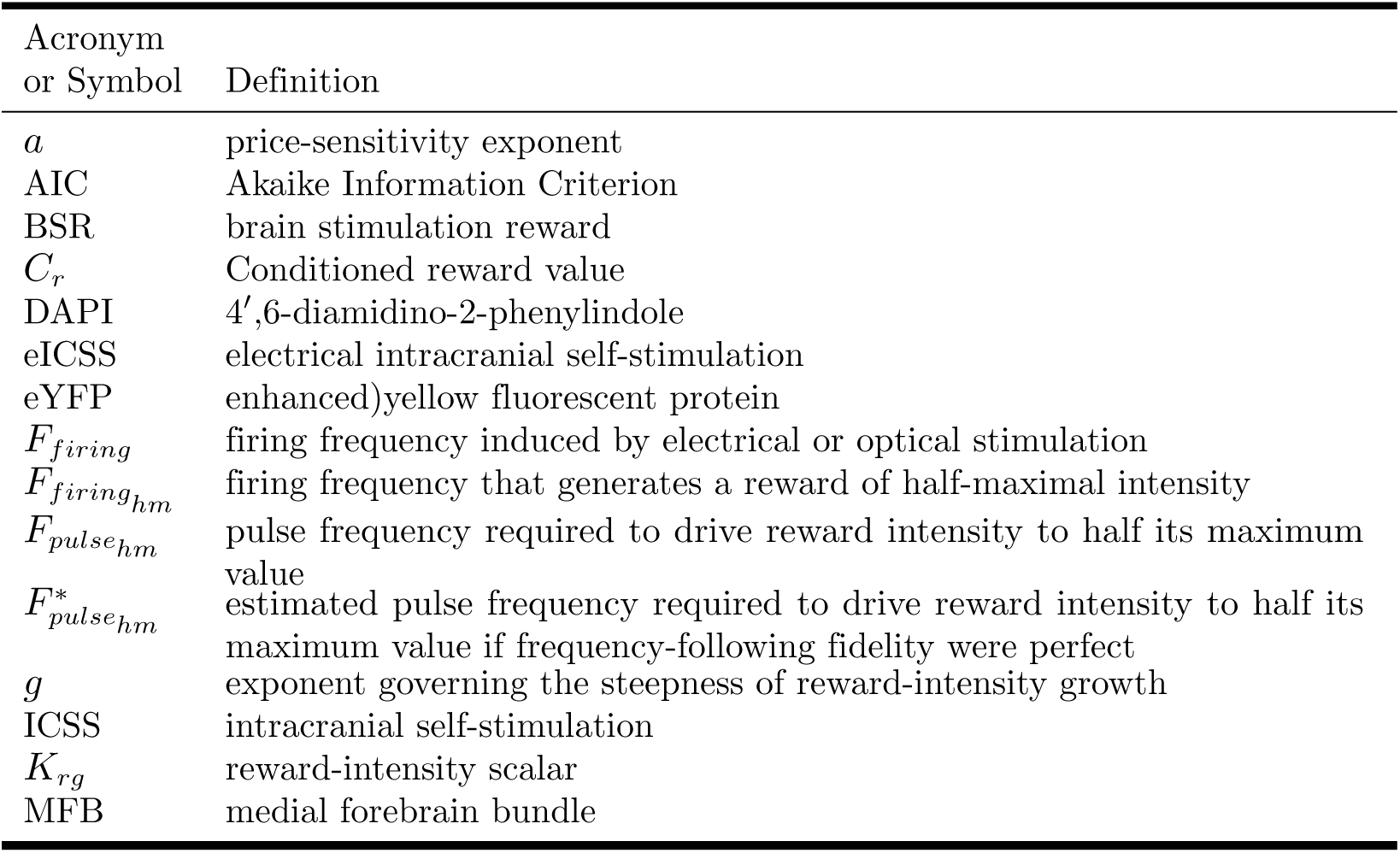
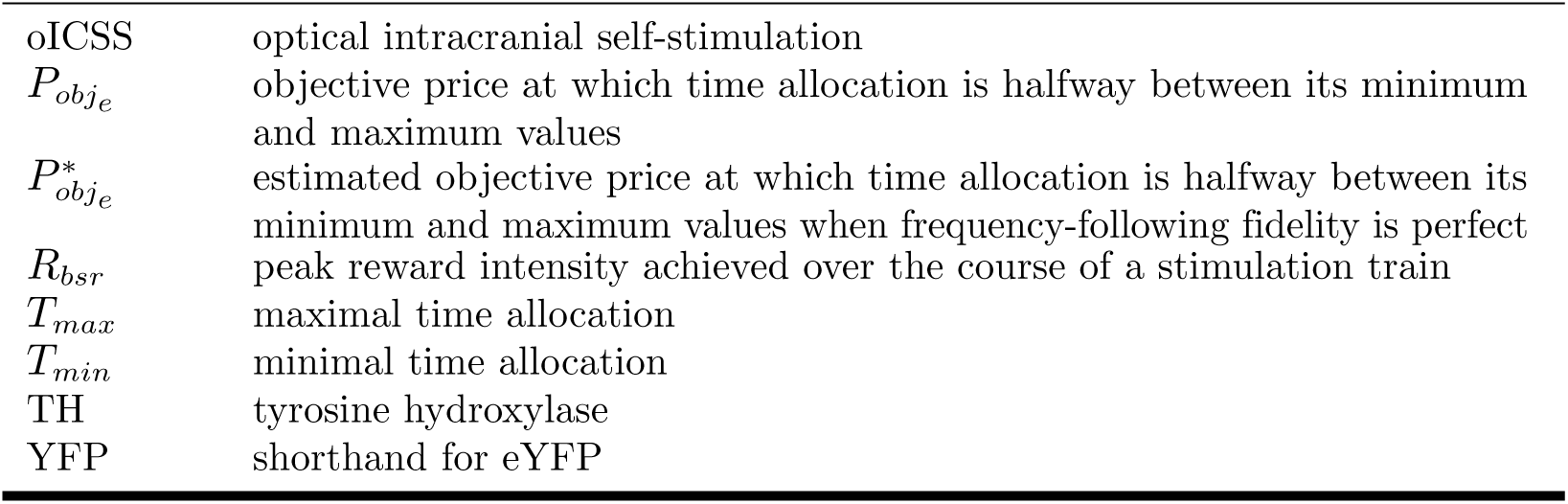
Definition of acronyms and symbols

1. characterization of spatiotemporal integration in the underlying neural circuitry,
2. measurement of how the intensity of the rewarding effect grows as a function of the aggregate rate of induced firing in the directly activated neurons,
3. measurement and modeling of how performance for the electrical reward depends on its strength and cost, and
4. determination of the stage of processing at which perturbation of dopaminergic neurotransmission alters eICSS.

#### The counter model

According to the “counter model” of spatiotemporal integration in the neural circuitry underlying eICSS [17–19], the neural signal that gives rise to the rewarding effect reflects the aggregate rate of induced firing produced by a pulse train of a given duration. Neither the number of activated neurons nor the rate at which they fire matter per se; it is their product that determines the intensity of the rewarding effect. The counter model is well supported empirically [20–23]. An analogous coding principle has been proposed by Murasugi, Salzman & Newsome [24] to account for the effect of electrical microstimulation of cortical area V5 on visual-motion perception.

#### The growth of reward intensity as a function of the aggregate rate of firing

An analogy may help convey what we mean by “reward intensity.” Imagine that a rat tastes a sucrose solution. The rat’s gustatory system is thought to provide two different kinds of information: a) sensory data indicating the concentration and identity of the tastant and b) evaluative data indicating what the sucrose is worth to the rat in current physiological and ecological state [25, 26]. These two signals could diverge, for example when an overfed rat encounters increasingly concentrated solutions. In that case, the sensory “sweetness” signal may continue to increase while the evaluative “goodness” signal plateaus or declines. We use the term, “reward intensity,” by analogy to the evaluative signal. Indeed, we have shown that rats can compare the reward-intensity signals produced by MFB stimulation and intraoral sucrose so as to determine which is larger and can combine them such that a compound electrical-gustatory reward is worth more than either of its constituents delivered singly [27].

The reward-growth function translates the aggregate rate of stimulation-induced firing into the intensity of the rewarding effect. Gallistel’s team used operant matching to describe this function [21–23, 28]. According to the matching law [29–32], subjects partition their time between two concurrent variable-interval schedules in proportion to the relative payoffs from the two schedules. If so, when two schedules are each configured to deliver stimulation trains, the relative payoffs can be inferred from the ratios of work times and reward rates.

Simmons and Gallistel [23] brought out a key feature of the reward-growth function: given a sufficiently high current, reward intensity saturates at pulse frequencies well within the frequency-following capabilities [8] of the directly stimulated substrate. In other words, reward intensity levels off as pulse frequency increases even though the output of the directly stimulated neurons continues to grow. The reward-growth function they described is well fit by a logistic [33], a function that is S-shaped when plotted on semi-logarithmic coordinates (reward intensity vs the logarithm of the pulse frequency).

#### The reward-mountain model

The behavioral method employed in this study entails measurement of time-allocation decisions [34] by laboratory rats. The method is based on a model (Fig 1) of how time allocation is determined by the strength and cost of reward. According to Herrnstein’s single-operant matching law [30, 31, 35], subjects performing an operant response, such as lever pressing, to obtain an experimenter-controlled reward partition their time between “work” (performance of the response required to obtain the reward), and “leisure” (performance of alternate activities such as grooming, exploring, and resting). The higher the benefit from the experimenter-controlled reward and the lower its cost, the larger the proportion of time devoted to work.

**Fig 1.**
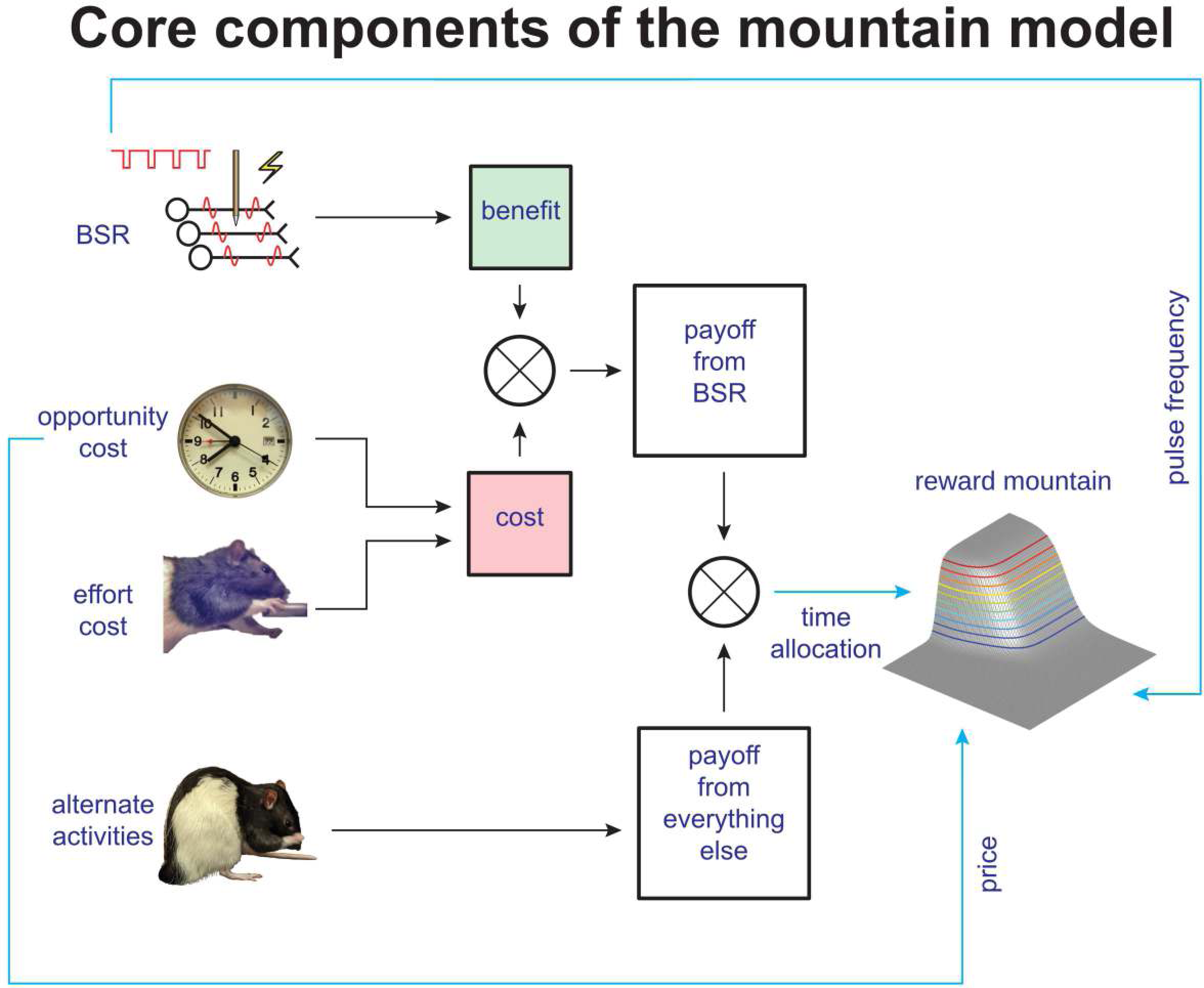
Core components of the reward-mountain model. The self-stimulating rat partitions its time between working for the rewarding stimulation and performing alternate activities, such as grooming, exploring, and resting. The payoff from work depends on the benefit it provides and the cost it entails. The benefit arises from the induced neural activity (shown here to arise from electrical stimulation), whereas the costs are of two different sorts: the intensity of the perceived effort entailed to meet the response requirement and the opportunity cost of the time so expended. The ratio of benefits to costs constitutes the payoff from the experimenter-controlled reward, which is compared to the payoff from alternative activities by means of a behavioral allocation function derived from Herrnstein’s single-operant matching law. The result of this comparison determines allocation of the subject’s time.

The reward-mountain model [36–38] treats the rewarding stimulation as a fictive benefit; although it satisfies no known physiological need, the effect of the stimulation mimics goal objects that do. Two types of costs are incorporated in the model. The first is the intensity of the perceived effort entailed in meeting the response requirement, which consists of holding down a lever for a duration determined by the experimenter. While the rat works to hold down the lever, it cannot groom, rest, or explore. Thus, it pays an opportunity cost [39], which consists of the benefits that would have been obtained from the foregone alternative activities.

In the spirit of the expanded matching law [32], the reward-mountain model equates the payoff from work to the ratio of its benefits and costs. A behavioral-allocation function [37] derived from the generalized matching law [40] compares the payoffs from work and leisure so as to determine the allocation of time to these two sets of activities.

### The effect of perturbing dopamine neurotransmission on the reward mountain

The curve-shift [41–43] or progressive-ratio [44] methods are typically regarded as the “gold standards” for measuring drug-induced changes in the behavioral effectiveness of brain-stimulation reward. These methods assess shifts in the functions relating performance vigor to electrical pulse frequency (in the case of the curve-shift method) or to the number of responses required to earn a reward (in the case of the progressive ratio method). The reward-mountain model shows that these two-dimensional methods yield fundamentally ambiguous results [36, 38]. Performance depends both on the strength of the electrical reward (determined by the pulse frequency) and on response cost (determined, in progressive-ratio testing, by the number of required responses per reward). This dependence is described by a surface in a three-dimensional space (reward-seeking performance versus pulse frequency and response cost (Fig 1)). When the surface is shifted along one of the axes representing the independent variables, its silhouette may also shift along the orthogonal axis [36–38]. An observer using either the curve-shift or progressive-ratio methods views only the silhouette of the surface and thus cannot determine in which way the surface itself (rather than its silhouette) has been displaced. Did administration of a drug shift the reward-growth function, displace the mountain along the cost axis, or both? An observer using either of these convention methods cannot know. In contrast, an observer using the three-dimensional reward-mountain model can answer definitively because the direction of displacement is determined unambiguously [36–38].

Shizgal’s team has used the reward-mountain model to assess the effects of perturbing dopaminergic neurotransmission on performance for rewarding electrical stimulation of the MFB [38, 45, 46]. They found that enhancement of dopaminergic neurotransmission by means of dopamine-transporter blockade [38, 45] or attenuation by means of dopamine-receptor blockade [46] shift the reward mountain almost uniquely along the axis representing response cost. This implies that, contrary to what was long believed [47, 48], the contribution of dopaminergic neurotransmission to brain-stimulation reward is brought to bear downstream (at, or beyond the output) of the reward-growth function.

### The empirical question and its significance

Rats [49] and mice [50, 51] will perform operant responses to obtain direct optical stimulation of midbrain dopamine neurons. The specific empirical question posed in the present study is whether the effect of dopamine-reuptake blockade on performance for rewarding optical stimulation of midbrain dopamine neurons, as measured by shifts in the reward mountain, mimics the effect of this manipulation on performance for rewarding electrical stimulation of the MFB [38, 45]. The outcome matters in at least two ways. First, it bears on the issue of how the output of midbrain dopamine neurons contributes to reward pursuit. Second, it bears on the series-circuit model that, on casual consideration, appears to integrate the findings obtained by means of direct, specific, optical activation of dopamine neurons with the older findings obtained by means of electrical stimulation that activates dopamine neurons indirectly.

Fig 2 illustrates the series-circuit model [10–12, 15, 16], which holds that the rewarding effect produced by electrical stimulation of the MFB arises from the transsynaptic activation of midbrain dopamine neurons. The left half of the figure depicts the generation of the reward-intensity signal subserving eICSS. The observed displacement of the reward mountain by perturbation of dopaminergic neurotransmission [38, 45, 46] requires that the reward-growth function be positioned upstream (to the left) of the input to the midbrain dopamine neurons.

**Fig 2.**
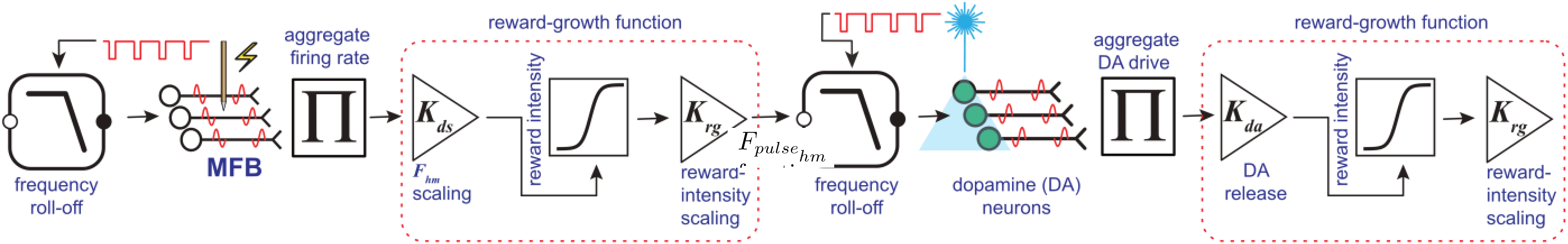
The series-circuit model. According to this model, the rewarding effects produced by both electrical stimulation of medial-forebrain-bundle (MFB) neurons and optical stimulation of midbrain dopamine neurons have a common cause: activation of midbrain dopamine neurons. This activation is due to transsynaptic excitation in the case of eICSS and direct excitation in the case of oICSS. The results of prior eICSS studies [38, 45, 46] require that drugs that perturb dopaminergic neurotransmission alter the computation of reward intensity by actions downstream from the output of a logistic reward-growth function (leftmost box containing an S-shaped curve). The results of the present study require that a similar reward-growth function lies downstream from the dopamine neurons (rightmost box containing an S-shaped curve). The boxed low-pass filter symbols represent the frequency-following functions that map the electrical pulse-frequency into the induced frequency of firing in the directly stimulated neurons subserving eICSS or the optical pulse-frequency into the induced frequency of firing in the dopamine neurons subserving oICSS. *K*_*ds*_ (*ds* stands for “directly stimulated”) scales the input to the reward-growth function for eICSS, whereas *K*_*da*_ scales the input to the reward-growth function for oICSS. *K*_*rg*_ scales the output of the reward-growth functions.

The right half of Fig 2 depicts the generation of the reward-intensity signal subserving oICSS. The four components to the right of the dopamine neurons represent the hypothesis that reward-intensity signals subserving eICSS and oICSS are computed in an analogous fashion. As we will show, this hypothesis is supported by the results of the current study.

Dopamine-transporter blockade increases stimulation-induced dopamine release and thus rescales the *input* to the reward-growth function shown in the right-hand portion of Fig 2 (via the triangular amplifier symbol labeled “*K*_*da*_”). Such an effect shifts the reward-growth function leftward along the logarithmic pulse-frequency axis due to the increased “bang for the buck,” dragging the reward mountain with it. The results confirm this prediction. These shifts along the pulse-frequency axis are orthogonal to those observed in the eICSS studies. We show below that the series-circuit model founders on this discrepancy: it fails to provide a cogent, unified account of both the prior eICSS and present oICSS data. Thus, we advocate abandoning the series-circuit model that has figured so heavily in accounts of eICSS over the past four decades. In its place, we develop a new account that can accommodate both the prior eICSS and the present oICSS findings.

### The issues addressed by the simulations

Multiple non-linear functions intervene between the stimulation we deliver to the brain and the behavioral consequences we observe.

1. The firing frequency of the activated neurons eventually rolls off as pulse frequency is increased sufficiently [8].
2. The intensity of the rewarding effect is an S-shaped function of the aggregate rate of induced impulse flow [22, 23, 28].
3. Subjective opportunity cost is a non-linear function of objective opportunity cost [39].
4. Although the form of the function has yet to be described empirically, subjective effort cost is almost surely a non-linear function of the objective work requirement, rising towards infinity as the physical capabilities of the subject are exceeded.
5. Allocation of behavior to eICSS or oICSS is an increasing sigmoidal function of reward intensity and a decreasing sigmoidal function of subjective cost [30, 31, 36–38].

Intuitive analysis and verbal reasoning rapidly come to grief when confronted with multiple, interconnected non-linearities. To understand how such a set of non-linear, interacting functions produces systematic behavior, it is necessary to capture the known relationships in a quantitative model and to simulate its output in response to the experimental inputs. This is what we have done to complement the empirical experiment. These simulations are summarized here and reported in detail in the accompanying Matlab^®^ Live Script.

The modeling and simulations provide mathematical and logical support for the proposed interpretation of the experimental results and their integration into an account of both oICSS and eICSS. They provide a systematic means for working out the minimal set of “moving parts” that determine reward pursuit, for making testable predictions about the effects of future manipulations, for shedding new light on the role of dopamine neurons in reward pursuit, and for guiding efforts to identify the non-dopaminergic components of the neural circuitry in which the dopamine neurons are embedded.

## Results

Seven rats completed both phases of the study: the power-frequency trade-off and the pharmacological experiment. The data are presented first in two dimensions. The reward-mountain model is then applied to integrate the results of the pharmacological experiment into a three-dimensional structure, to document the effects of dopamine-transporter blockade on the position of the reward mountain, and to interpret these displacements in terms of drug-induced changes in the values of the variables that determine reward pursuit.

### Power-frequency trade-off

The data in Fig 3 were obtained using the FR-1 task. The number of responses emitted per 2-min trial by an exemplar rat (Bechr29) is shown as a function of pulse frequency and optical power; the results for the remaining rats are shown in the supporting-information file (Figs S1-S6). Response rates grew as the pulse frequency was increased, in some cases (Bechr26-29) up to, or well beyond, optical pulse frequencies of 40 pulses s^-1^. In rats Bechr14, 19, 21, and 27, maximum response rats grew as a function of pulse frequency over the lower powers but approached asymptote as the power was further increased. In the remaining rats, maximum response rates grew as a function of pulse frequency over the entire range of tested powers. The position of the rate-frequency curves generally shifted leftwards as optical power was increased.

**Fig 3.**
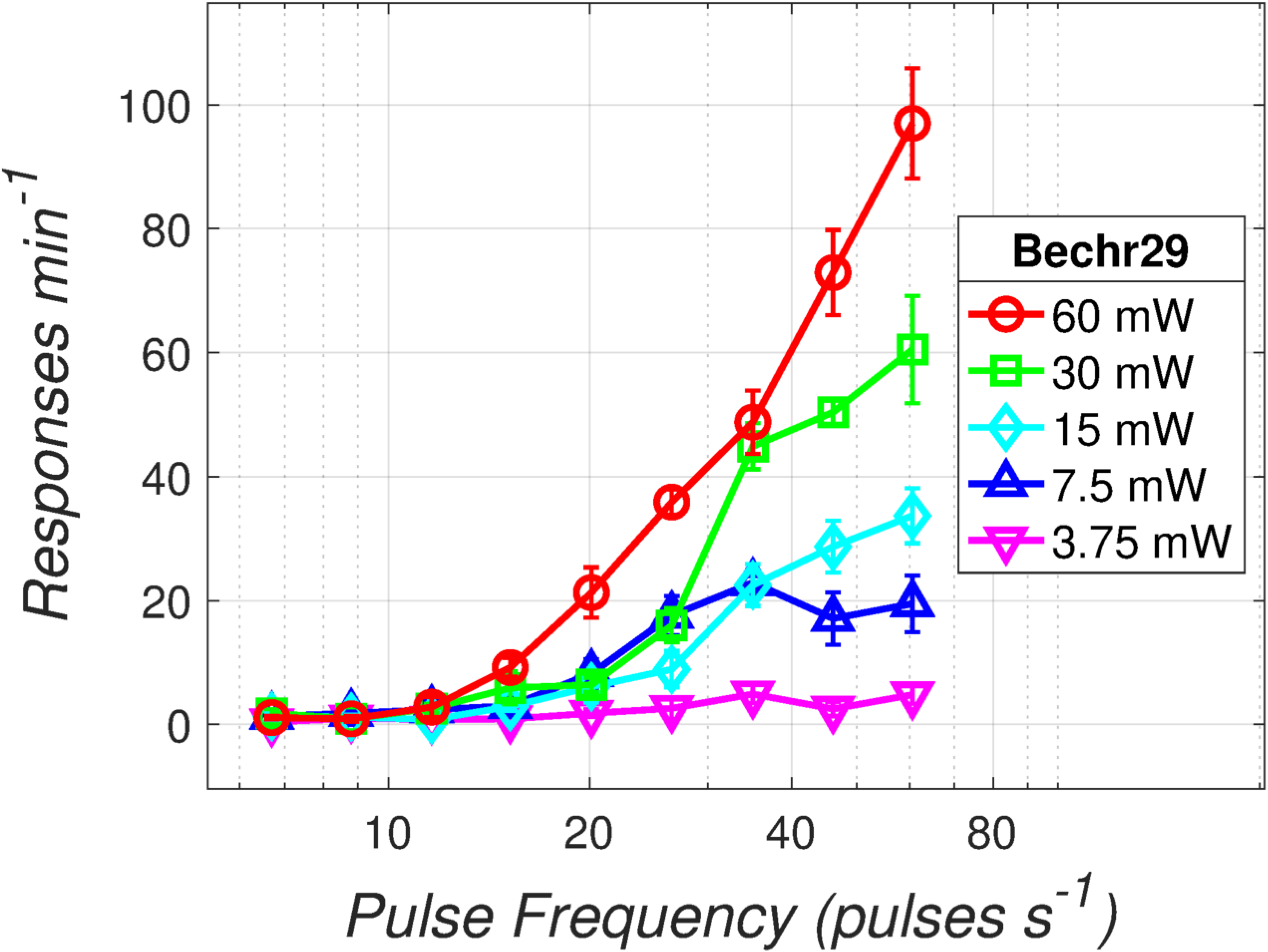
Rate-frequency curves as a function of optical power. The number of responses emitted per 2-min trial by an exemplar rat (Bechr29) is plotted as a function of pulse frequency and optical power.

### Time allocation as a function of reward strength and cost

The data in Fig 4 were obtained using the lever-hold-down task. The two-dimensional views shown here were eventually combined (see below) to generate three-dimensional reward mountains. The curves in Fig 4 plot time allocation as a function of reward strength (“Pulse Frequency”) or opportunity cost (“Price”). The results are from an exemplar rat (Bechr29). Panel A shows frequency-sweep results: Time allocation increased as a function of optical pulse frequency. The time-allocation-versus-pulse-frequency curve obtained following blockade of the dopamine transporter by GBR-12909 is shifted leftwards with respect to the curve obtained in the vehicle condition. Panel **B** shows price-sweep results: Time allocation decreased as a function of increases in opportunity cost (cumulative time that the lever had to be held down to trigger a reward). The time-allocation-versus-price curve obtained following blockade of the dopamine transporter is shifted rightwards with respect to the curve obtained in the vehicle condition. Radial-sweep data were obtained by conjointly decreasing the pulse frequency and increasing the price in stepwise fashion over consecutive trials. Panels **C** and **D** show the same radial-sweep results from two orthogonal vantage points. Time allocation is plotted against pulse frequency in panel **C** and against price in panel **D**. Time allocation increased as a function of pulse frequency and decreased as a function of price. The time-allocation curve obtained following blockade of the dopamine transporter is shifted leftwards along the pulse-frequency axis with respect to the curve obtained in the vehicle condition and rightwards along the price axis. Graphs for the remaining rats are shown in the supporting-information file (Figs S7-S12).

**Fig 4.**
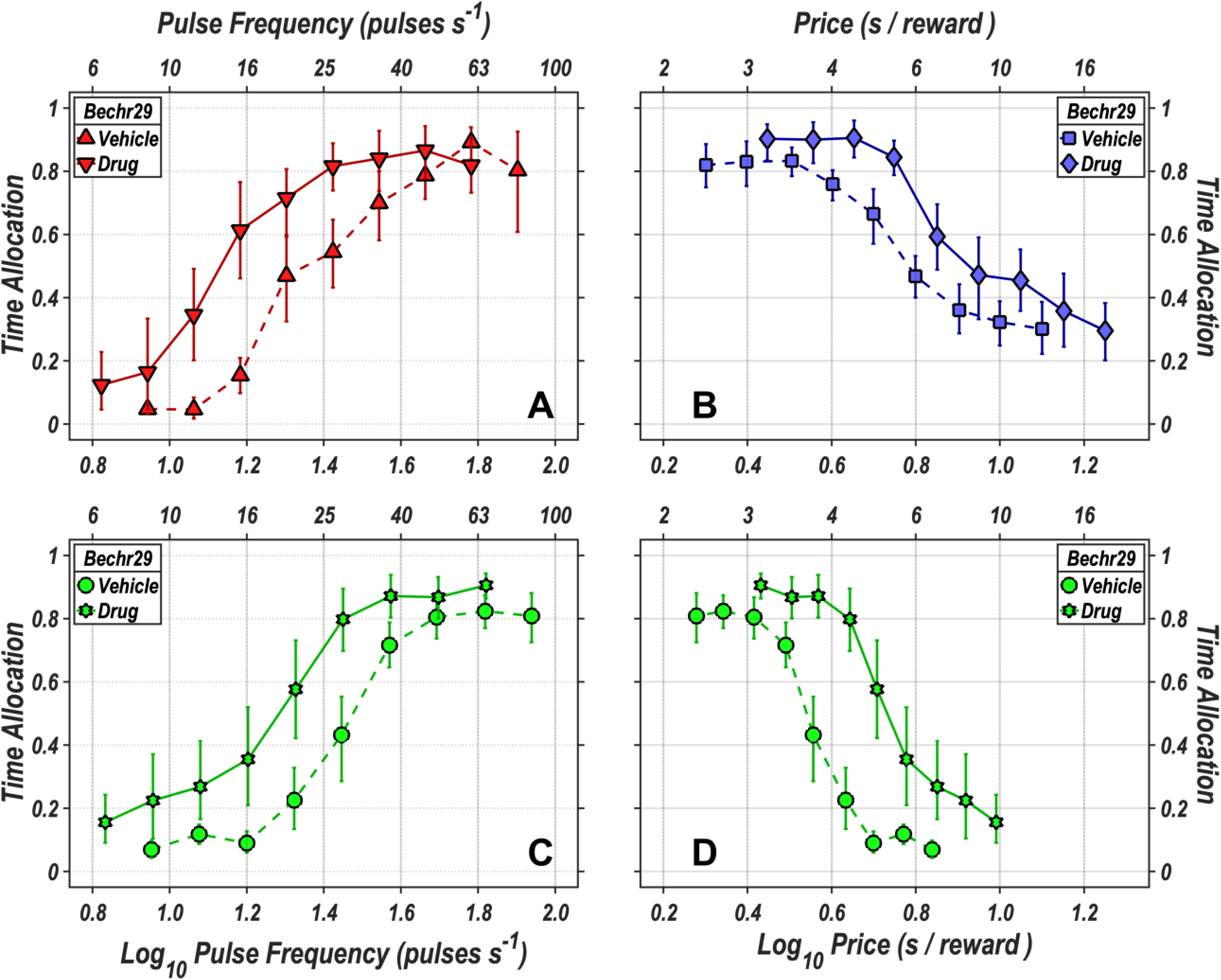
Time allocation as a function of reward strength and cost. **A:** Time allocation as a function of pulse frequency (reward strength) in the vehicle (upright triangles) and drug (inverted triangles) conditions. **B:** Time allocation as a function of price (opportunity cost) in the vehicle (squares) and drug (diamonds) conditions. In the radial-sweep condition, the pulse frequency was decreased and the price decreased concurrently, in stepwise fashion, over consecutive trials. Time allocation is plotted as a function of pulse frequency in panel **C:** and as a function of price in panel **D:**. Data from the vehicle condition are represented by circles, whereas data from the drug condition are represented by Stars of David. The error bars represent 95% confidence intervals. Data are from an exemplar rat (Bechr29).

The drug-induced shifts in the curves shown in Figs 4 and S7-S12 are inherently ambiguous: displacement of these curves along a given axis could be due to any combination of displacements of the underlying reward-mountain structure along the price and/or pulse-frequency axes [37, 38, 52]. This ambiguity is removed by fitting the reward-mountain model, which expresses time allocation as a function of both the price and strength of the rewarding stimulation. In the three-dimensional space of the reward-mountain model, we can determine unambiguously the degree to which dopamine-transporter blockade dispaces the mountain along the price and pulse-frequency axes.

### Model fitting and selection

The standard version of the reward-mountain model has six parameters: 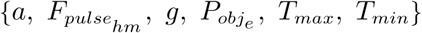. The 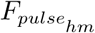 and 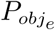 parameters set the location of the mountain along the pulse-frequency and price axes, respectively. 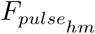 is the pulse frequency at which reward intensity is half maximal, whereas 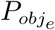 is the price at which time allocation to pursuit of a maximal reward falls midway between its minimal and maximal values, *T*_*min*_ (minimum time allocation) and *T*_*max*_ (maximal time allocation). The slope of the mountain surface along the price axis is set by the price-sensitivity exponent, *a*, whereas the slope along the pulse-frequency axis is determined both by *a* and by the reward-growth exponent, *g*.

In previous studies ([38, 45]), a seven-parameter version of the reward-mountain model sometimes performed better than the standard six-parameter version. The added parameter accommodates cases in which minimum time allocation is higher at low prices than at higher ones. This parameter is called *C*_*r*_ and has been interpreted to represent a reward value assigned to the lever and/or to the act of pressing it [38].

Both the six- and seven-parameter versions of the reward-mountain model are derived in the supporting-information file. (See: Derivation of the reward-mountain model.)

There is a trade-off between the number of parameters fit to a dataset and the precision with which the value of each parameter can be estimated. In order to restrict the number of fitted parameters, we forced common values of *T*_*max*_ and *T*_*min*_ to be fit to the vehicle and drug data. The two location parameters were always free to vary across the vehicle and drug conditions. Four variants of the six-parameter model were fit. The variants are distinguished by whether either, both, or neither of the *a and g* parameters were free to vary across the vehicle and drug condition. Eight variants of the seven-parameter model were fit. These variants are distinguished by the combinations of the *a, C*_*r*_, and *g* parameters that were free to vary across the vehicle and drug conditions.

The 12 candidate models (four variants of the six-parameter version and eight variants of the seven-parameter version) are described in Tab S3. The Akaike Information Criterion (AIC) [53] was used to identify the best-fitting model for each rat. This statistic implements a trade-off between goodness of fit and simplicity. Thus, the AIC penalizes models with large numbers of parameters in comparison to simpler ones. The fits of all of the candidate models to the reward-mountain data from all seven rats converged successfully. Detailed results for rat Bechr29 are shown in Tab S4, and results for all rats are shown in Tabs S5,S6.

### Fitted reward-mountain surfaces

The reward-mountain surfaces fit to the vehicle and drug data from rat Bechr29 are shown in Fig 5, whereas those fit to the data from the remaining rats are shown in the supporting-information file (Figs S17-S22).

**Fig 5.**
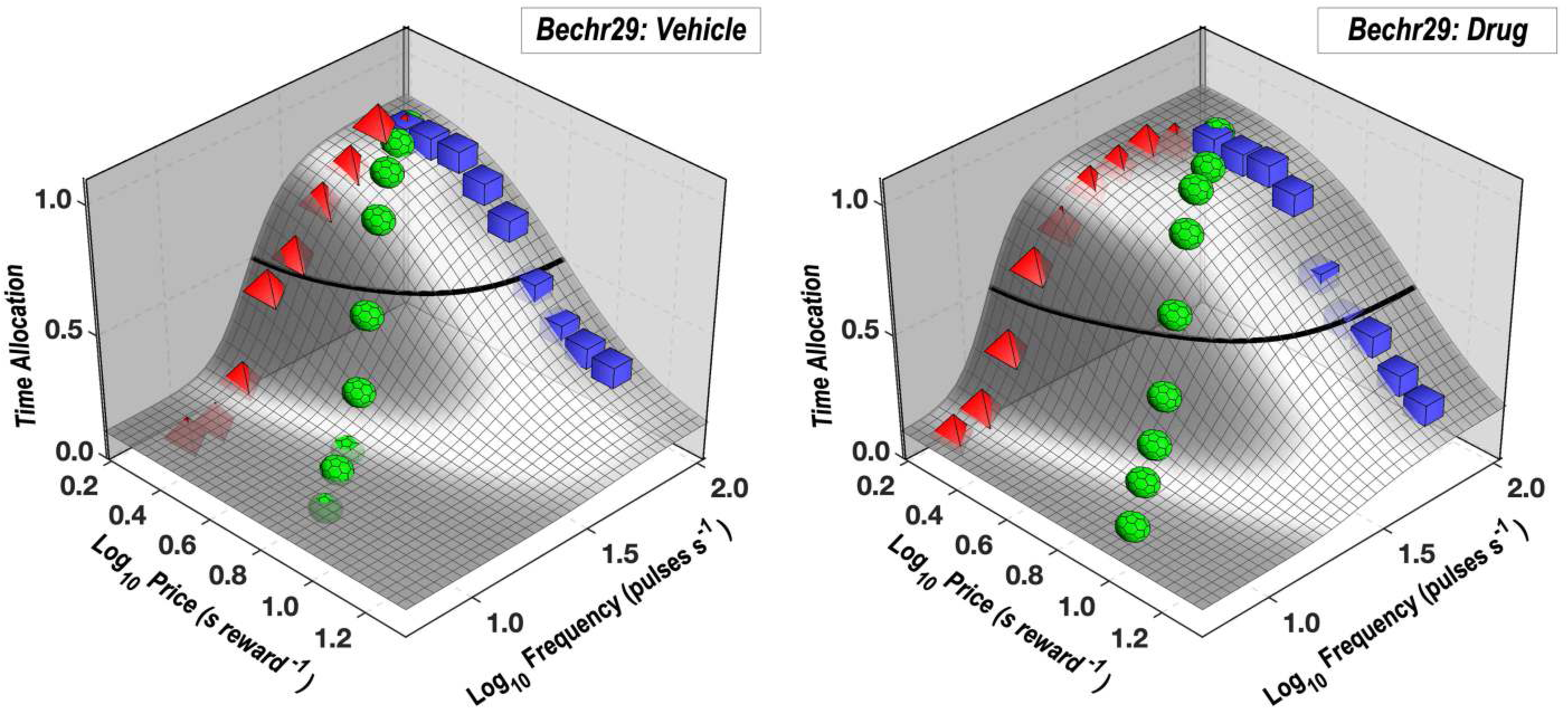
Reward-mountain surfaces fit to the vehicle and drug data from rat Bechr29. The surfaces of the reward-mountain shell are shown in gray. The thick black line represents the contour mid-way between the minimal and maximal estimates of time allocation (the estimated altitudes of the valley floor and summit). Mean time-allocation values for the pulse frequency, price, and radial sweeps are denoted by red pyramids, blue squares, and green polyhedrons, respectively.

To facilitate visualization of the drug-induced shift in the location of the reward mountain in Fig 5, the surfaces have been re-plotted as contour graphs in Fig 6. Horizontal comparison in Fig 6 shows that blockade of the dopamine transporter shifted the mountain downwards along the pulse-frequency axis, whereas vertical comparison shows that the mountain shifted rightwards along the price axis. The shifts are summarized in the bar graph; dot-dash cyan lines show estimates corrected for changes in frequency-following fidelity due to the drug-induced displacement of the mountain along the pulse-frequency axis. The rationale for the correction and the details of its implementation are described in the supporting-information file. (See sections ***Parameters of the frequency-following function for oICSS*** and ***Correction of the location-parameter estimates for changes in frequency-following fidelity***.) Contour and bar graphs for the remaining rats are shown in the supporting-information file (Figs S23-S28).

**Fig 6.**
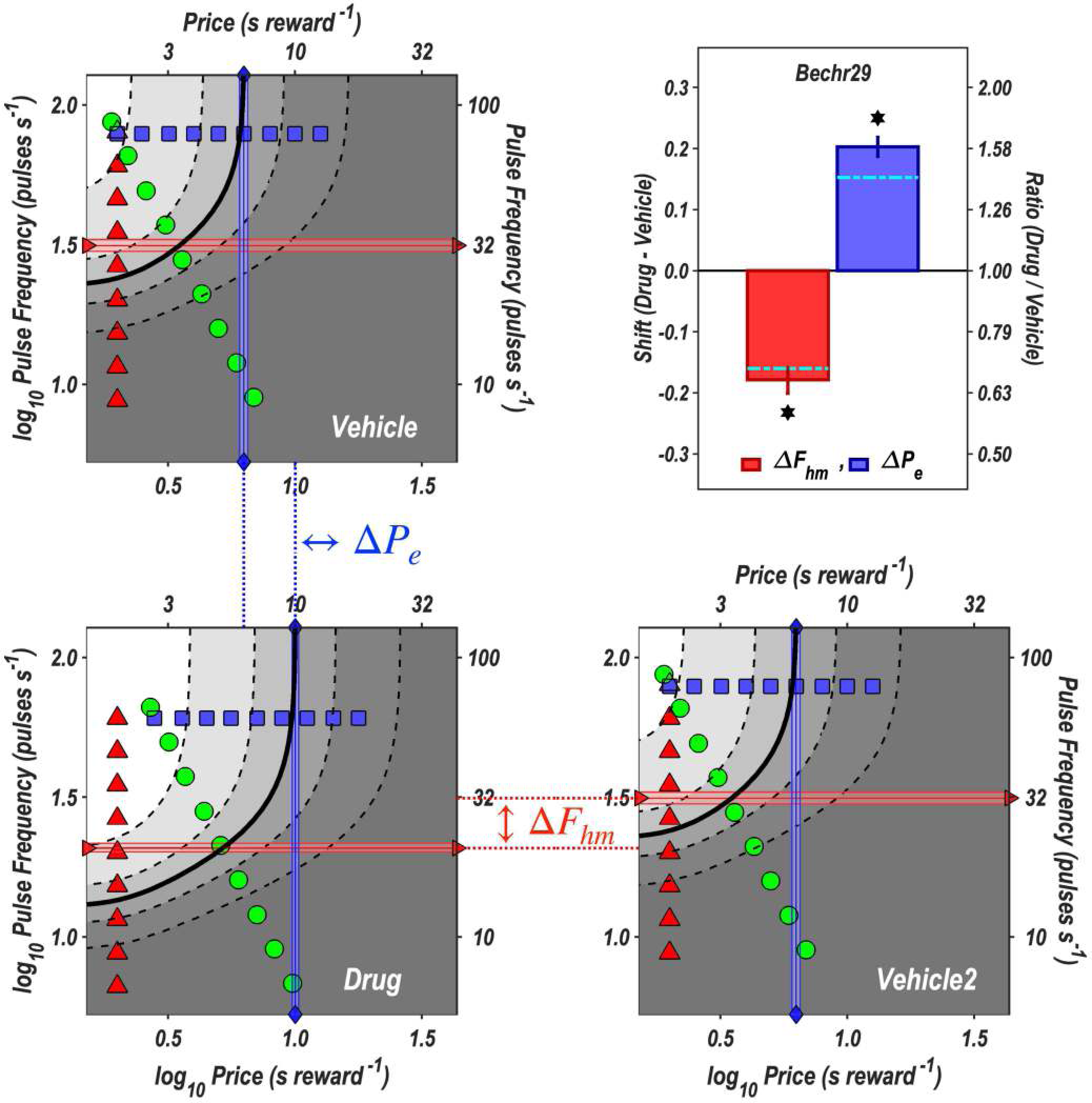
Contour graphs of the reward-mountain surfaces fit to the vehicle and drug data from rat Bechr29. The values of the independent variables along frequency sweeps are designated by red triangles, along price sweeps by blue squares, and along radial sweeps by green circles. The values of the location parameters, 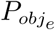 and 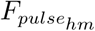, are indicated by blue vertical lines with diamond end points and red horizontal lines with right-facing triangular end points, respectively. The shaded regions surrounding the lines denote 95% confidence intervals. The vehicle data are shown twice, once in the upper-left quadrant and once in the lower right. The dotted lines connecting the panels designate the shifts in the common-logarithmic values of the location parameters of the mountain, which are designated as 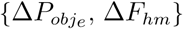 and plotted in the bar graph in the upper-right panel. The dot-dash cyan lines superimposed on the bars show location-parameter estimates corrected for changes in frequency-following fidelity due to the displacement of the mountain along the pulse-frequency axis. (See section ***Correction of the location-parameter estimates for changes in frequency-following fidelity*** in the supporting-information file.) The 95% confidence intervals are shown in the bar graphs as vertical lines.

### Location-parameter estimates

#### F_hm_

The surface-fitting procedure returns the location-parameter value, 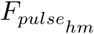 that positions the reward mountain along the ***pulse*-**frequency axis. In studies of eICSS employing the reward-mountain model, it was assumed that this value corresponded to the induced ***firing*** frequency in the directly activated neurons. This firing frequency is the location parameter of the underlying reward-growth function. Given the exceptional frequency-following fidelity of the directly activated neurons subserving eICSS of the MFB [8], it is not unreasonable to assume that each pulse elicits an action potential in most or all of the directly stimulated MFB neurons when the pulse frequency equals 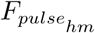. In contrast, the findings reviewed in section ***Parameters of the frequency-following function for oICSS*** of the supporting-information file argue that such an assumption is untenable in the case of oICSS of channelrhodopsin-2 expressing midbrain dopamine neurons. We therefore used the data from the rate-frequency curves obtained here (Figs 3 and S1-S6) and prior studies (e.g., [54]) to estimate the firing frequencies corresponding to the 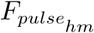 values. We label these as 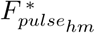 (rather than as 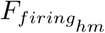) because these values will be plotted along the ***pulse***-frequency axis, and we refer to them as “corrected” estimates of 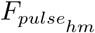.

Tab 2 shows the estimated drug-induced *shifts* in the location of the reward-mountain core along the pulse-frequency axis. The shifts are the differences between the common logarithms of the estimated firing frequencies 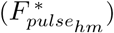 that produced half-maximal reward intensities in the drug and vehicle conditions (Tab S7). Also included are the differences between the estimates for the drug and vehicle conditions and the 95% confidence intervals surrounding these differences. In all cases, the confidence band excludes zero, thus meeting our criterion for a statistically reliable effect. In six of seven cases, the sign of the difference is negative, indicating that the drug shifted the reward mountain downwards along the pulse-frequency axis. Note that the one discrepant shift (for Rat Bechr27) is the smallest. The values in the 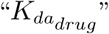 column give the proportional reduction in 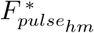 produced by dopamine-transporter blockade.

**Table 2.** Drug-induced shifts of the reward mountain along the pulse-frequency axis. The 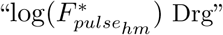 and 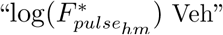 columns list the common logarithms of the values in the 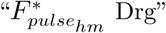 and 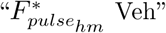 columns of Tab S7. The “diff” column shows the differences between the estimates for the drug and vehicle conditions. The “diff Lo” and “diff Hi” columns designate the lower and upper bounds of the 95% confidence interval surrounding these differences. The “*” character in the “excl 0” column indicates that zero falls outside the 95% confidence interval surrounding the estimates in the”diff” column. Differences so designated meet our criterion for statistical reliability. The rightmost column lists the value of 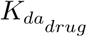 in Eqs S13,S14,S44 implied by the values in the “diff” column.

| Rat | $\log(F_{pulse_{hm}}^*)$<br>Drg | $\log(F_{pulse_{hm}}^*)$<br>Veh | diff | diff<br>Lo | diff<br>Hi | excl 0 | $K_{da_{drug}}$ |
| --- | --- | --- | --- | --- | --- | --- | --- |
| Bechr14 | 1.239 | 1.364 | -0.126 | -0.161 | -0.100 | ★ | 1.335 |
| Bechr19 | 1.015 | 1.160 | -0.145 | -0.206 | -0.088 | ★ | 1.395 |
| Bechr21 | 0.959 | 1.468 | -0.508 | -0.583 | -0.449 | ★ | 3.224 |
| Bechr26 | 1.195 | 1.339 | -0.144 | -0.165 | -0.121 | ★ | 1.393 |
| Bechr27 | 1.307 | 1.221 | 0.086 | 0.048 | 0.124 | ★ | 0.820 |
| Bechr28 | 1.119 | 1.431 | -0.313 | -0.331 | -0.295 | ★ | 2.055 |
| Bechr29 | 1.260 | 1.420 | -0.160 | -0.179 | -0.142 | ★ | 1.446 |

#### P_e_

The estimates that locate the mountain along the price axis have also been corrected so as to remove the contribution of the changes in frequency-following fidelity due to the drug-induced displacement of the mountain along the pulse-frequency axis. (In the supporting-information file, please see sections Correction of the location-parameter estimates for changes in frequency-following fidelity, and Illustration of the correction for imperfect frequency-following fidelity.) Tab 3 gives the common logarithms of the corrected values 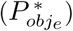 listed in Tab S9 along with the differences between the estimates for the drug and vehicle conditions and the 95% confidence intervals surrounding these differences. In all cases, the confidence band excludes zero, thus meeting our criterion for a statistically reliable effect. The signs of the differences are all positive, indicating that under the influence of dopamine-transporter blockade, a higher price was required to bring time allocated to pursuit of a maximal reward to its middle value. With one exception (Rat Bechr27), the correction reduced the magnitude of the drug-induced shift (as shown by the position of the dot-dash cyan lines in the bar-graph panels of Fig 6 and S23-S28).

**Table 3.**
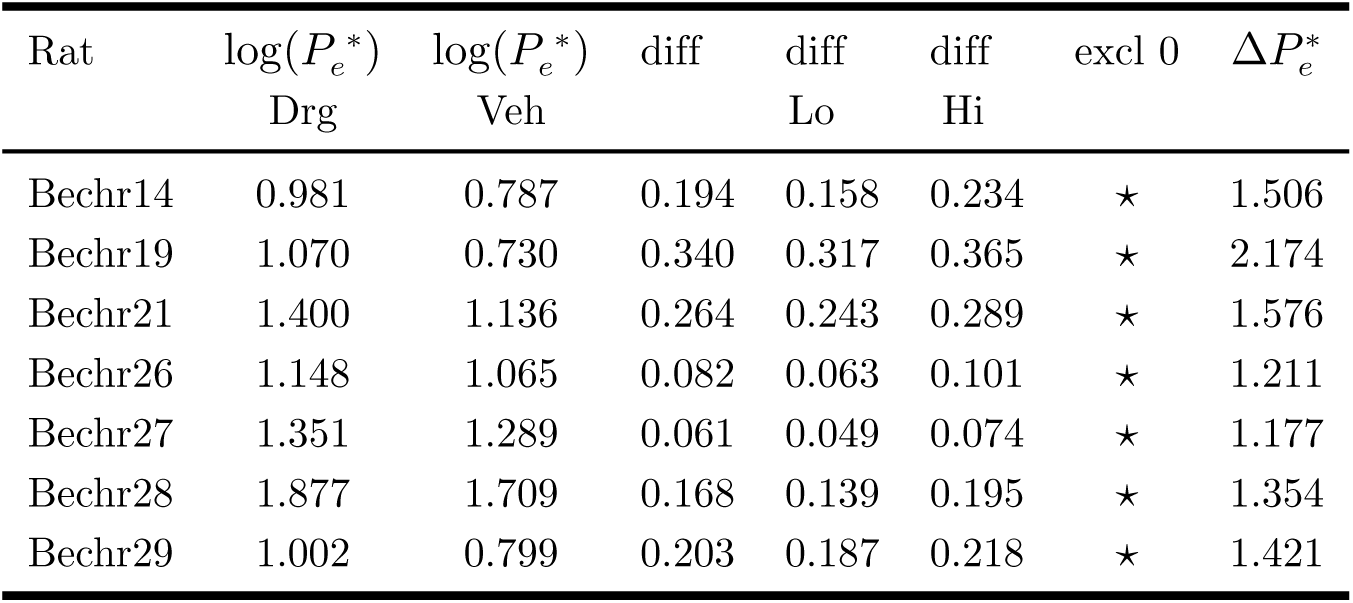
The location of the reward mountain along the price axis. The 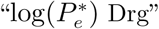 and 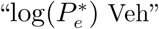 columns list the estimated values that the 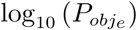 parameter would have attained in the drug and vehicle conditions, respectively, had frequency following been perfect. The “diff” column shows the differences between the estimates for the drug and vehicle conditions, whereas the “diff Lo” and “diff Hi” columns designate the lower and upper bounds of the 95% confidence interval surrounding these differences. The ⋆ character in the “excl 0” column indicates that zero falls outside the 95% confidence interval surrounding the estimates in the”diff” column. Differences so designated meet our criterion for statistical reliability. The rightmost column lists the proportional drug-induced change in the value of 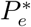 corresponding to the values in the “diff” column. 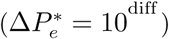

| Rat | $\log(P_e^*)$<br>Drg | $\log(P_e^*)$<br>Veh | diff | diff<br>Lo | diff<br>Hi | excl 0 | $\Delta P_e^*$ |
| --- | --- | --- | --- | --- | --- | --- | --- |
| Bechr14 | 0.981 | 0.787 | 0.194 | 0.158 | 0.234 | $\star$ | 1.506 |
| Bechr19 | 1.070 | 0.730 | 0.340 | 0.317 | 0.365 | $\star$ | 2.174 |
| Bechr21 | 1.400 | 1.136 | 0.264 | 0.243 | 0.289 | $\star$ | 1.576 |
| Bechr26 | 1.148 | 1.065 | 0.082 | 0.063 | 0.101 | $\star$ | 1.211 |
| Bechr27 | 1.351 | 1.289 | 0.061 | 0.049 | 0.074 | $\star$ | 1.177 |
| Bechr28 | 1.877 | 1.709 | 0.168 | 0.139 | 0.195 | $\star$ | 1.354 |
| Bechr29 | 1.002 | 0.799 | 0.203 | 0.187 | 0.218 | $\star$ | 1.421 |

The corrected values of the two location parameters are shown in Fig 7.

**Fig 7.**
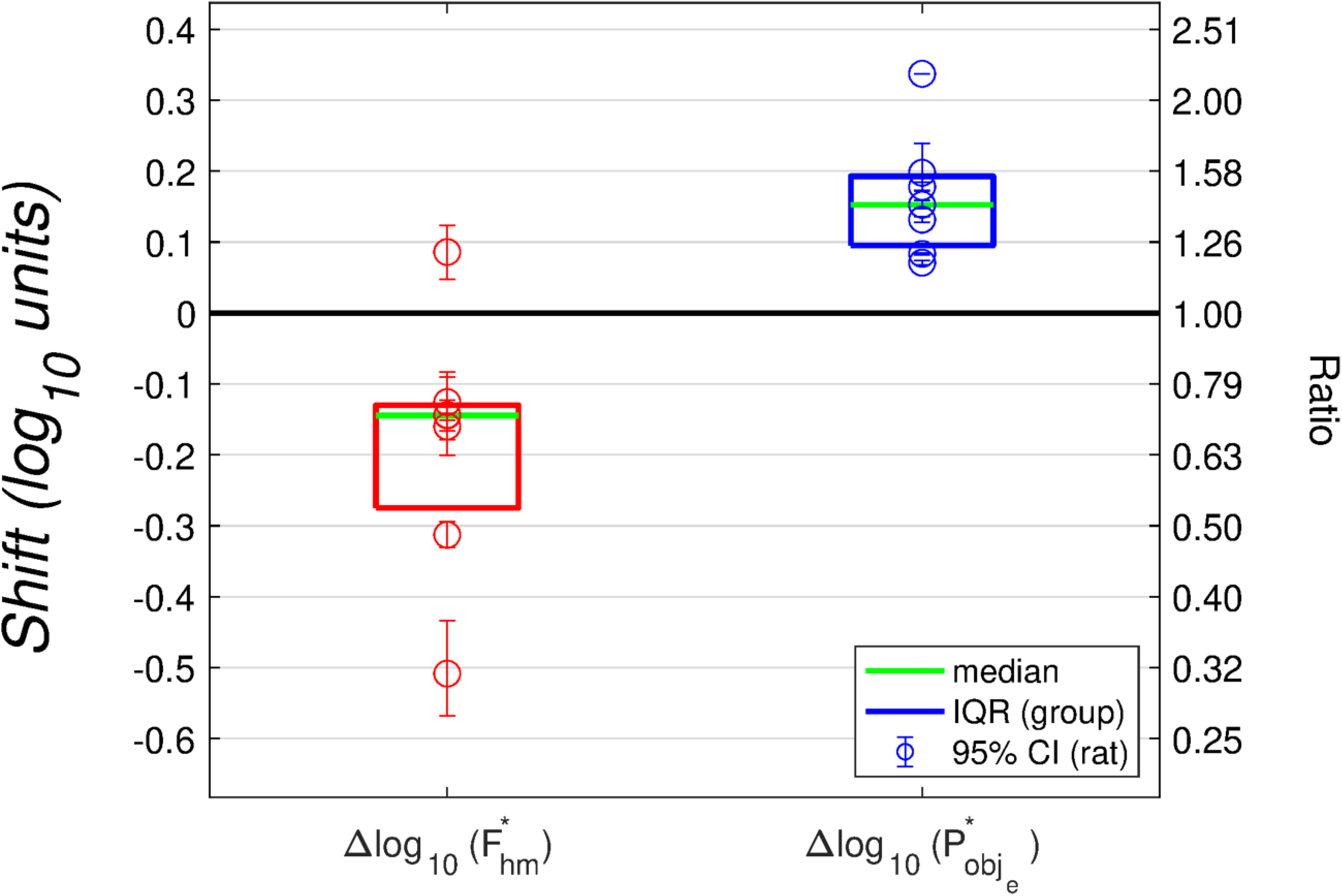
Drug-induced shifts in the location parameters of the reward mountain. The 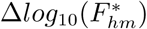 values give the shifts of the reward-growth function and reward mountain along the pulse-frequency axis, whereas the 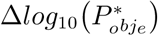 values give the shifts along the price axis. These values have been corrected for the estimated change in frequency-following fidelity due to the drug-induced displacement of the reward mountain along the pulse-frequency axis. According to the reward-mountain model, the 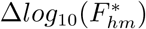 values reflect action of the dopamine transporter blocker prior to the input to the reward-growth function, whereas the 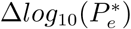 values reflect drug action at, or beyond, the output of the reward-growth function. 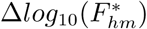 is shorthand for 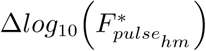.

The corrected values of the location parameters are uncorrelated in both the vehicle (*ρ* = 0.37, *p* = 0.41) and drug (*ρ* = −0.24, *p* = 0.60) conditions), as are the drug-induced ***shifts*** in these values (*ρ* = −0.2830; *p* = 0.5385, Fig S29).

All parameters of the best-fitting models for each rat are shown in Tabs S10-S15 in the supporting information. (The values of the location parameters in these tables are uncorrected.)

### Histology

Immunohistology and confocal microscopy (Fig 8) confirms that the tips of all optical-fiber implants (heavy, angled black lines) were located close to transfected midbrain-dopamine neurons and that ChR2 was selectively expressed in TH-positive neurons.

**Fig 8.**
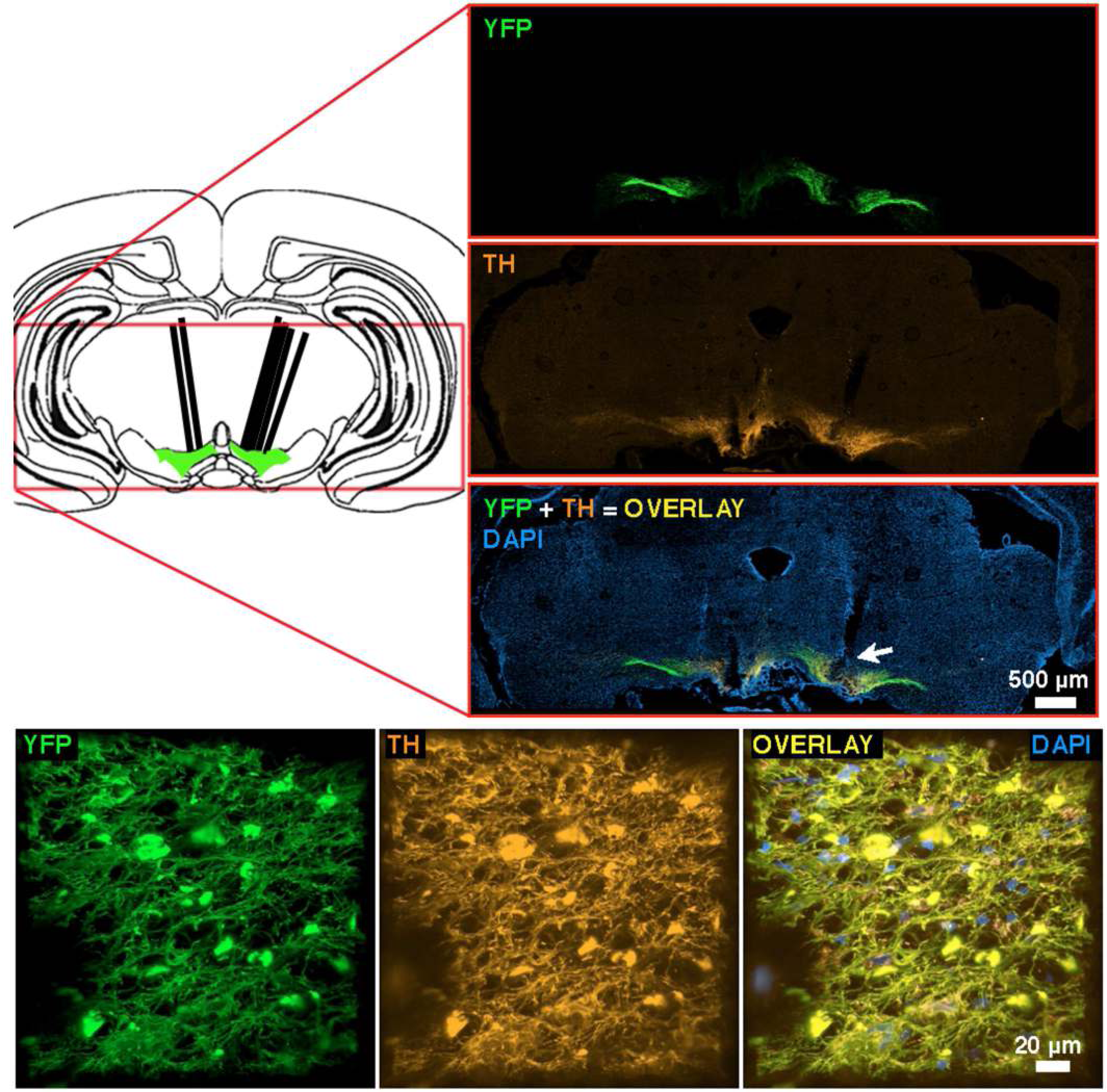
Images of viral construct expression from rat BeChr29. upper left: Schematic representation of a coronal section at the approximate location of the optical-implant tip (modified from [55]). Heavy angled black lines denote the optical-fiber tracks. **upper right:** Representative immunohistochemical images. YFP and TH staining is shown in the top and middle panels respectively. The bottom panel shows the co-expression of YFP and TH (overlay) along with DAPI for anatomical reference. The arrow indicates the approximate location of the tip of the optical implant used for oICSS. **bottom** High magnification 3D reconstruction of the area below the optical-fiber track. Left: YFP-positive neurons; Middle: TH-positive neurons; Right: Overlay of YFP, TH and DAPI staining. Note that YFP is expressed in TH-positive neurons.

**Fig 9.**
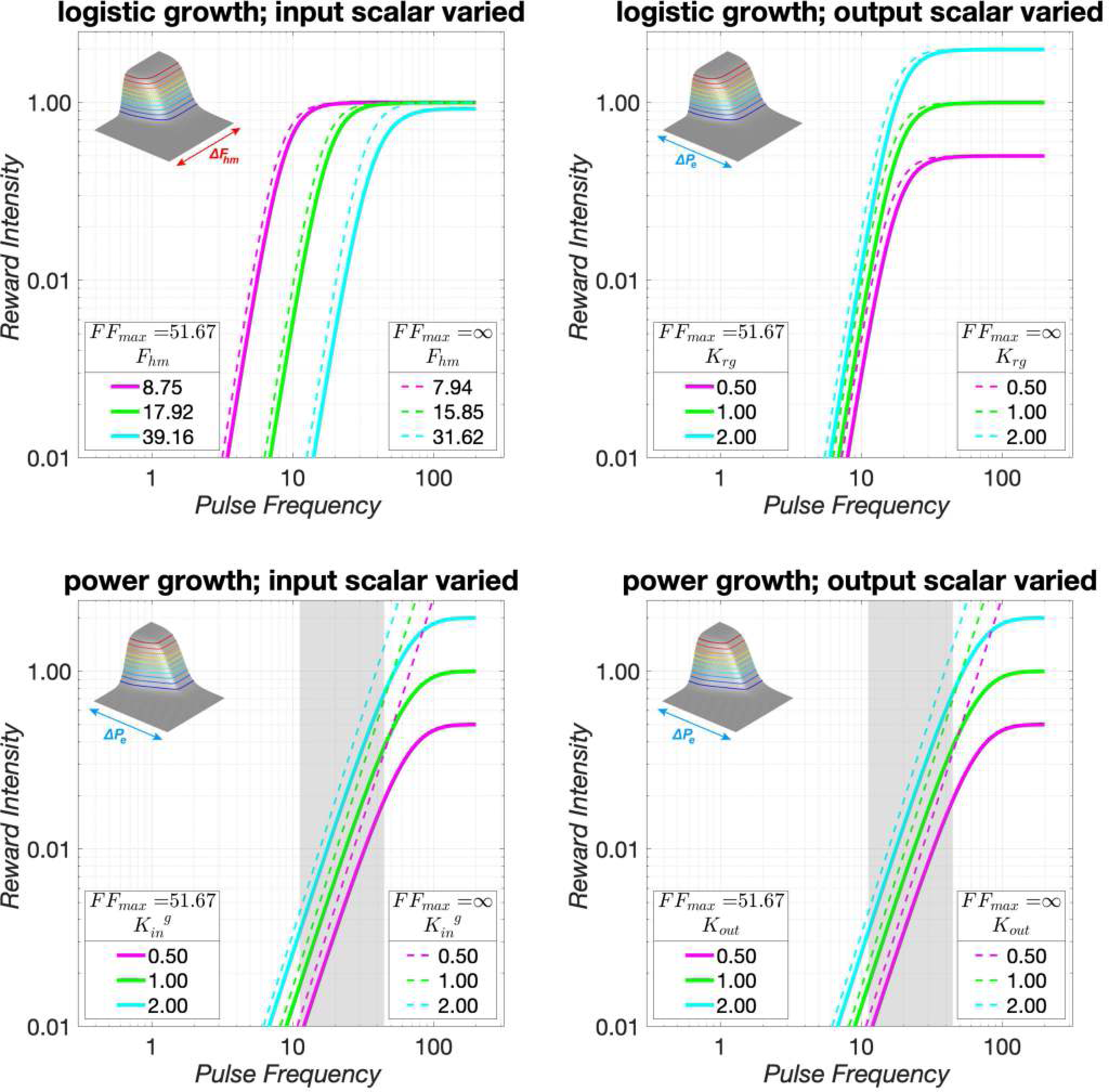
Influence of input- and output-scaling parameters on reward-growth functions. The upper panels show logistic reward-growth functions, whereas the lower panels show power functions. Solid lines map the ***pulse*** frequency into the reward intensity using the assumed frequency-following function (see: Parameters of the frequency-following function for oICSS). *FF*_*max*_ is the maximum firing frequency 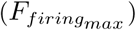 attainable given the form and parameters of the assumed frequency-following function. The dashed lines map the ***firing*** frequency into the reward intensity. (These lines are drawn assuming perfect frequency-following fidelity (*FF*_*max*_ = ∞), and thus pulse frequency and firing frequency are equivalent.) In the case of the logistic reward-growth functions in the upper panels, changing the values of the input- and output-scaling parameters produces independent effects, shifting the reward-growth function in orthogonal directions. In contrast, changing the values of the input- and output-scaling parameters produces identical effects on the power-growth functions plotted in the lower panels.

## Discussion

Decades of psychophysical research on eICSS have provided a detailed portrait of the neuro-computational processes that translate a train of electrical current pulses into subsequent reward-seeking behavior. The reward-mountain model [36–38] integrates these findings so as to predict the allocation of behavior to eICSS from the strength and cost of the rewarding stimulation. The Matlab^®^ live script documented in the supporting-information file simulates several of the principal validations studies, and compares these simulations to empirical results. The simulations and empirical data show that rescaling the input to the reward-growth function shifts the reward mountain along the pulse-frequency axis, whereas rescaling performed at, or beyond, the output of the reward-growth function shifts the reward mountain along the price axis.

Substitution of optical for electrical stimulation [49–51] offers significant advantages: the resulting neural activation is confined to a known, genetically specified, neural population, and the neurons that express the light-sensitive transducer molecule are readily visualized. However, few psychophysical data have been reported concerning the processes that translate the optical input into observable reward-seeking behavior. The results of the present study demonstrate that the dependence of oICSS and eICSS on the strength and cost of rewarding stimulation is formally similar. Nonetheless, boosting dopaminergic neurotransmission by means of dopamine-transporter blockade alters oICSS of midbrain dopamine neurons and eICSS of the MFB in strikingly different ways.

The reward-mountain model was developed to account for data from eICSS experiments. We begin by considering what the current results tell us about generalization of this model to oICSS. We then discuss the significance of the location-parameter values that position the surfaces fitted to the vehicle data within the space defined by the strength and cost of the optical reward. Next, we turn to the main experimental question posed in this study: how does blockade of the dopamine transporter alter the position of the reward mountain? Finally, we discuss the implications of those drug-induced shifts, and we thereby show why a new way of thinking about brain reward circuitry is required to integrate the results of the pharmacological challenge with existing eICSS data.

### The reward-mountain model generalizes successfully to oICSS

The shape of the reward-mountain surface fitted to the oICSS data (Figs 5, S17-S22) resembles that of the surfaces fit to eICSS data reported previously [8, 37–39, 45, 46, 56, 57]. Time allocation falls as pulse-frequency is decreased and/or price is increased. The resemblance between the shapes of the surfaces fitted to prior eICSS and current oICSS data suggests that the reward-mountain model generalized well to the case of oICSS and that pursuit of rewarding optical or electrical stimulation depends similarly on the strength and cost of reward.

Embedded within the reward-mountain model derived in the context of eICSS studies is the notion that reward intensity depends on the aggregate rate of firing induced in the directly stimulated neurons by a pulse train of fixed duration [20–23, 33]. A definitive test of this “counter model” [19] has yet to be reported in the case of oICSS. Nonetheless, there are indications that the counter model holds in this case as well.

Ilango and colleagues trained mice to press a lever to receive optical stimulation of ChR2-expressing midbrain dopamine neurons [58]. They showed that lever-pressing rates increased as a function of both pulse duration and optical power, two variables that conjointly determine the number of neurons activated by the optical stimulation. Lever-pressing also increased when an 8-pulse train was delivered at progressively higher pulse frequencies. This latter result cannot be linked unambiguously to the counter model because the train duration covaried with the pulse frequency; in the case of eICSS, the formal relationships between these two variables and reward intensity differ [33]. The frequency-sweep data reported here were obtained with train duration held constant. Thus, they complement and extend the findings of Ilango and colleagues while avoiding the complication of covariation between the pulse frequency and train duration. Time allocation increased systematically as a function of pulse frequency, as the reward-mountain model and the counter model embedded within it predict.

Ilango and colleagues also measured changes in lever-pressing rates during performance of oICSS on fixed-interval or fixed-ratio schedules of reinforcement [58]. Response rates declined as either the minimum inter-reward interval or the required number of responses per reward was increased.

Fixed-interval schedules set the minimum inter-reward interval, but the subject determines the physical effort expended in harvesting the reward. Although only a single response is required to trigger reward delivery after the fixed interval has elapsed, subjects typically begin responding much earlier and thus expend more effort than is strictly necessary [59]. Fixed-ratio schedules set the number of responses required to trigger reward delivery but cede control of the minimum inter-reward interval to the subject, who can vary response rate so as to bring reward delivery closer (or further) in time, as Ilango and colleagues observed. In contrast to these two classic schedules, the cumulative handling-time schedule employed here fixes both the minimum inter-reward interval and the rate of physical exertion required to secure a reward. This makes it possible to vary the opportunity cost of the reward independently of the required rate of physical exertion, as was done here and in previous experiments carried out in the reward-mountain paradigm. As in the case of the eICSS experiments, increases in the opportunity cost (price) of the reward decreased time allocation systematically (Figs 4, S7-S12, 6, S23-S28).

### Vehicle condition: location parameters and their significance

#### Pulse-frequency axis

In rats working for fixed-duration trains of rewarding, electrical, MFB stimulation, the parameter that positions the reward mountain along the pulse-frequency axis, 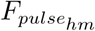, depends on the stimulation current [36]. This variable, (in conjunction with the pulse duration [18, 60]) determines the number of directly stimulated neurons [19] in a manner dependent upon their excitability and spatial distribution with respect to the electrode tip. By analogy, we expect the number of midbrain dopamine neurons recruited by optical pulses and trains of fixed duration to depend on the optical power, the level of ChR2 expression in the dopamine neurons, and the spatial distribution of these neurons with respect to the tip of the optical probe. Given the inevitable variation in ChR2 expression and probe-tip location, it is not surprising that that 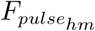 varied over a wide range in the vehicle condition, from 16.3 - 35.8 pulses s^-1^ (Tab S7). Even after correction for differences in frequency-following fidelity, the estimated firing frequencies induced by the 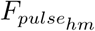 values (i.e., 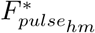) vary more than twofold.

#### Price axis

In principle (and in contrast to 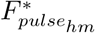), the corrected estimate of the parameter that positions the eICSS reward mountain along the price axis, 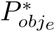, can be compared meaningfully across subjects and electrical stimulation sites. The validity of this comparison rests on the assumption that both the value of alternate activities, such as grooming, resting, and exploring, and the effort cost of performing the lever-depression task do not vary substantially across subjects or stimulation sites. If so, then the price at which a maximally intense, stimulation-generated reward equals the value of alternate activities 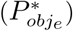 reflects the maximum intensity of the rewarding effect, independent of the value of the stimulation parameters required to drive reward intensity to its upper asymptote. On this view, the rat is willing to sacrifice more leisure in order to obtain strong stimulation of a”good” eICSS site than a poorer one.

Could such a comparison extend meaningfully to different forms and targets of stimulation? The current oICSS experiment was carried out in the same testing chambers as previous eICSS studies. If the value of alternate activities in that environment does not change systematically as a function of whether electrical stimulation of the MFB or optical stimulation of midbrain dopamine neurons serves as the reward, then it is of interest to compare the 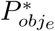 values of the current study to the 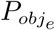 values obtained in prior eICSS experiments. (For reasons discussed in section Correction of the location-parameter estimates for changes in frequency-following fidelity of the supporting-information file, the 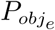 estimates from the eICSS studies do not typically require correction for imperfect frequency-following fidelity). The results of this comparison are intriguing.

The median of the corrected 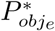 values for the vehicle condition of the present study, 11.8 s, is marginally greater than the median of the 54 values reported in previous eICSS studies [8, 37–39, 45, 46, 56, 57], but the range is much more extreme. Whereas 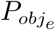 values typically vary over a roughly twofold range in the eICSS studies, the 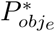 values vary over a more than a tenfold range in the current oICSS dataset. The maximum (56.4) is much greater than any value obtained in the eICSS studies, whereas the minimum (5.4) is smaller.

According to the reward-mountain model, the large magnitude of the maximum 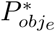 value means that direct optical stimulation of midbrain dopamine neurons can produce a much more potent rewarding effect than electrical stimulation of the MFB, at least in an extreme case. Perhaps the maximal intensity of the rewarding effects reported here varies so profoundly across subjects because the activation of different subsets of dopamine neurons is rewarding for different reasons. If so, the large range of 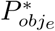 values would add to the accumulating evidence for functional heterogeneity of midbrain dopamine neurons [61–68].

The reward mountain was measured in only seven subjects in the current study, and the dispersion of the tips of the optical probes is modest. Given these limitations and the uncertainly inherent in estimating the location of the optically activated dopamine neurons from the position of the tip, we cannot provide a meaningful assessment of whether the maximum intensity of the rewarding effect, as indexed by the 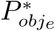 value, is correlated with the location of the optically activated dopamine neurons. It would be interesting to pursue this issue in a larger number of subjects and across a larger region of the midbrain. Mice can be trained to perform oICSS for stimulation of sites arrayed across much of the medio-lateral extent of dopaminergic cell bodies in the ventral midbrain, from the lateral portion of the substantia nigra to the medial boder of the ventral tegmental area [69]. Does 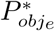 vary systematically as a function of tip location across the full extents of the midbrain region where dopaminergic cell bodies are located?

The reader may be tempted to ascribe the large variation in 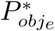 values to variables such as across-subject differences in ChR2 expression and the location of the optical-fiber tip. If, as we argue below, the form of the reward-growth function resembles a logistic, such differences would have contributed to the variation in 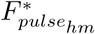 values and not to the 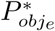 values. That is so because across-subject differences in ChR2 expression and tip location influence the ***input*** to the reward-growth function: the aggregate firing rate induced by the optical stimulation in the midbrain dopamine neurons. In contrast, the scalar that determines the maximum reward intensity (*K*_*rg*_) alters the ***output*** of the reward-growth function (Fig 2), and is one of the components of the 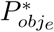 parameter (Eq S30). Thus the reward-mountain model holds that the value of 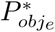 reflects the maximum intensity of the rewarding effect.

According to the argument that the 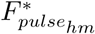 values, but not 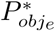 values, should depend on local conditions such as ChR2 expression and the spatial distribution of dopamine neurons with respect to the probe tip, the values of these two location parameters should be uncorrelated. The results are consistent with this expectation.

### Drug-induced shifts in the position of the reward mountain

Tabs 2, S8, S9 show that the dopamine-transporter blocker, GBR-12909, shifted the reward mountain reliably along the pulse-frequency and price axes in all 7 rats. (Only one result is discrepant with those from the rest of the group: the direction of the shift in the data from rat Bechr27, which is also the smallest of the shifts along the pulse-frequency axis.) The magnitudes of the shifts along the two axes are uncorrelated. These results have multiple implications:

1. The reward-growth function responsible for oICSS has independent input-scaling and output-scaling parameters and resembles a logistic.
2. The shifts along the pulse-frequency axis are due to an effect of the drug at, or prior to, the input to the reward-growth function.
3. The shifts along the price axis are due to an independent effect of the drug at, or beyond, the output of the reward-growth function.
4. The shifts along the pulse-frequency axis create grave problems for the hypothesis that the rewarding effect produced by electrical stimulation of the MFB arises solely from transsynaptic activation of midbrain dopamine neurons (the series-circuit model).

#### The reward-growth function for oICSS

The effects of dopamine-transporter blockade on the reward mountain powerfully constrain the underlying reward-growth function, and the functional form implied by these results has far-reaching implications concerning the structure of brain-reward circuitry. We develop this argument by first analyzing the form of the reward-growth function underlying the reward-mountain model, then by demonstrating how the data support this form, and finally by deriving the implications of this functional form for the structure of brain-reward circuity.

The cross-sectional shape of the reward-mountain is determined principally by the form of the reward-growth function. If frequency-following fidelity were perfect and subjective prices were equal to objective ones, then the contour lines would have the same form as the reward-growth function (flipped and rotated, as shown in Fig 10 of [39].)

**Fig 10.**
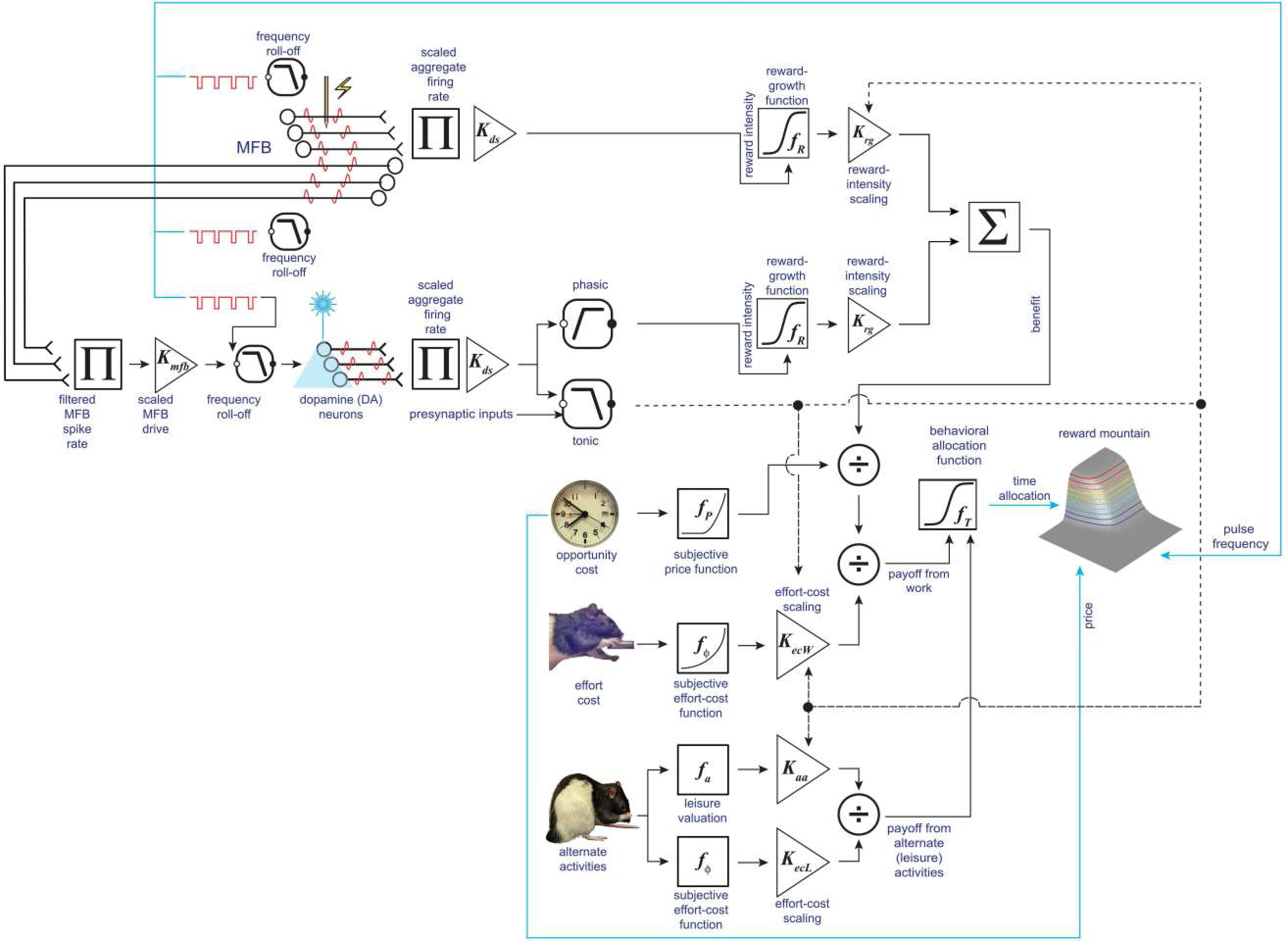
Parallel pathways conveying reward-intensity signals to a final common path. The directly activated neurons subserving eICSS of the MFB are depicted at the left of the top row of symbols. The electrode also activates a second population of fibers, with lower frequency-following fidelity [80], that project directly or indirectly to midbrain dopamine neurons, which have yet lower frequency-following fidelity. Optical stimulation of midbrain dopamine neurons activates only the lower limb of the circuit. The outputs of the reward-growth functions in the two limbs converge (Σ) on the final common path for reward evaluation and pursuit.

The subjective-price function [8] bends the contour lines towards the horizontal at very low prices but has no effect on the location of the reward mountain within the coordinates defined by the independent variables (price and pulse frequency). As estimated by Solomon et al. [39], this function converges on the objective price, and the discrepancy between the subjective and objective values is within 1% once the objective price exceeds 3.18 s. The subjective and objective prices can no longer be distinguished once the objective price approaches the values of the 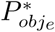 location parameter (Tab S9).

Given the frequency-following function assumed here, frequency-following fidelity is imperfect over much or all of the tested range of pulse frequencies. That said, firing frequency falls only slightly short of the pulse frequency at pulse frequencies below the 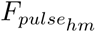 values (Fig S13). The procedure for generating the corrected values of the parameter that locates the reward mountain along the pulse-frequency axis 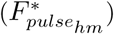 is designed to remove the influence of imperfect frequency-following fidelity. We emphasize that the correction procedure adjusts the estimates of the shifts but cannot manufacture these out of whole cloth. The correction is driven by differences in the location along the pulse-frequency axis of the surfaces fit to the vehicle and drug data. Had the drug failed to displace the mountain surface, the correction would be zero. Although the correction is likely imperfect, the estimated drug-induced displacement of the reward mountain along the pulse-frequency axis (Tab 2) should be due largely to the effect of the drug on the reward-growth function.

Unlike the case for eICSS [21–23, 28], the reward-growth function for oICSS has not yet been measured directly. The fact that a reward-mountain surface based on a logistic reward-growth function fits the current data well provides one hint that this function may indeed be logistic in form, or very similar. But there is a deeper sense in which the results of the present study provide crucial new information about the reward-growth function for oICSS: The displacement of the reward mountain along both the strength and cost axes by dopamine-transporter blockade ***requires*** that the reward-growth function for oICSS have independent location-scaling and output-scaling parameters, like the logistic reward-growth function for eICSS. To explain why this is so, it is helpful to reformat the logistic reward-growth equation (Eq S11), as follows:

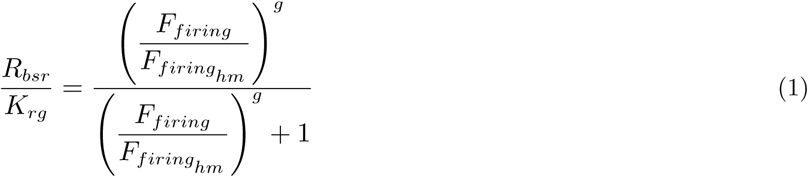

where

*F*_*firing*_ = firing rate produced by a pulse frequency of *F*_*pulse*_ pulses s^-1^

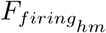 = firing rate required to drive reward intensity to half its maximum value

*g* = exponent that determines the steepness of reward-intensity growth as a function of pulse frequency

*K*_*rg*_ = reward-growth scalar

*R*_*bsr*_ = reward intensity produced by *F*_*firing*_

This format makes clear that *K*_*rg*_ scales the *output* of the logistic reward-growth function (*R*_*bsr*_), whereas 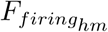 scales the *input* (*F*_*firing*_). These two scalars act independently, as Fig 9 illustrates: changes in 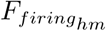 shift the reward-growth function in the log-log plot in the upper-left panel along the X (pulse-frequency) axis, whereas changes in *K*_*rg*_ shift the log-log plot of the reward-growth function in the upper-right panel along the Y (reward-intensity) axis. The inserts in Fig 9 and the contour and bar graphs in Figs S30, S31 show that these orthogonal shifts move the reward mountain along the pulse-frequency and price axes, respectively.

What would happen if frequency-following fidelity remained the same, but the input-scaling and output-scaling parameters of the reward-growth function were no longer independent? We can simulate such a case by replacing the logistic reward-growth function in Eq 1 with a power function:

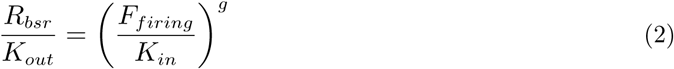

where

*K*_*in*_ = the input-scaling parameter

*K*_*out*_ = the output-scaling parameter

This power-growth function can be rewritten as

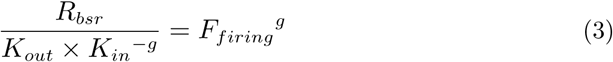

In contrast to the case of logistic growth, the two scaling constants act jointly, in an inseparable manner, and exclusively on the output of the function. Changing either scaling constant shifts the power-growth function along the Y axis but does not change its position along the X axis (lower panels of Fig 9). In contrast to the logistic function (upper panels), the two constants act as one. Fig S33 shows that changing the value of the input-scaling parameter of the power-growth function moves the reward mountain only along the price axis, and Fig S34 shows that changing the value of the output-scaling parameter has the same effect.

The three power-growth functions in the lower panels of Fig 9 designated by solid lines all bend at the same location along the X axis. This is because the frequency-following function that translates pulse-frequency into firing frequency (Eq S1, Fig S13) levels off after that point, truncating the input. However, despite this common landmark, these functions lack a true location parameter: as shown by the dashed lines, the underlying reward-growth function, which maps the firing frequency into reward intensity, rises continuously over the entire domain. In contrast, the logistic curves in the upper-left panel of Fig 9 level off at different locations along the X axis, as determined by their 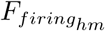 (and *g*) values.

A location parameter analogous to 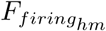 can be defined for any monotonic growth function that has a saturation point, like the logistic reward-growth function for eICSS described by the matching data obtained by Gallistel’s group [21–23, 28]. Such curves have an implicit threshold pulse frequency that just suffices to drive reward intensity out of the baseline noise and a saturating pulse frequency beyond which further increases cannot drive reward intensity higher. The growth of reward intensity is constrained to occur over this interval. The width of the interval between the threshold and saturating pulse frequencies is determined by the reward-growth exponent, *g* (Eq S11), whereas the location of this growth region along the function’s domain is set by the input-scaling parameter, 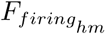. In this study, dopamine-transporter blockade shifted the reward mountain reliably along the pulse-frequency axis (Tab 2, Figs 6, S23-S28). A reward-growth function that lacks independent input- and output-scaling parameters cannot produce such results, as the lower panels of Fig 9) and Figs S33, Fig S34 illustrate. Instead, a logistic-like function is required.

### Shifts along the pulse-frequency axis

Blockade of the dopamine transporter by GBR-12909 increases the amplitude of stimulation-induced dopamine transients recorded in the nucleus-accumbens terminal field [70, 71]. It follows that in the current experiment, a lower pulse frequency will suffice to drive peak dopamine concentration to a given level under the influence of the drug than in the vehicle condition. Transporter blockade would thus rescale upwards the input to the reward-growth function. Such an effect is illustrated in Fig S16. The amplification of the stimulation-induced dopamine release by GBR-12909 is represented by the triangle labelled “*K*_*da*_.” The drug rescales upwards the impact of the phasic stimulation-induced increase in the aggregate firing rate (“DA drive”) on the input to the S-shaped reward-growth function, which is thus shifted leftwards along the pulse-frequency axis (Fig 9), dragging the reward-mountain surface downward along this axis (Fig 6). New optical methods [72, 73] facilitate concurrent measurement of dopamine concentrations and behavioral allocation to reward pursuit and could thus provide a direct test of the proposed mechanism for the observed decreases in 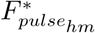.

### Shifts along the price axis

In addition to increasing the amplitude of stimulation-induced dopamine transients, blockade of the dopamine transporter by GBR-12909 also increases the baseline (“tonic”) level of dopamine [45, 70, 71]. The increase in dopamine tone could rescale the output of the reward-growth function upwards, reduce subjective effort costs and/or diminish the value of activities that compete with pursuit of the optical reward. Such effects could arise from increases in *K*_*rg*_ and/or decreases in *K*_*ec*_ or *K*_*aa*_ (Fig S16). Any combination of these effects could account for the observed rightward shifts of the reward mountain along the price axis (Figs 6, S23-S28). This can be seen by reformatting Eq S30 and expressing it verbally as follows:

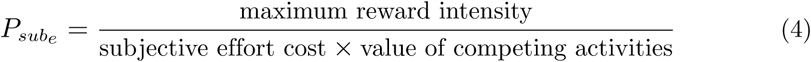

(Reward probability has been omitted from Eq 4 because the probability of reward upon satisfaction of the response requirement was equal to one in the present study.)

Teasing apart the three alternative accounts of the drug-induced change in 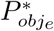 may not be feasible on the basis of behavioral data alone; it will likely require identification of the neural substrates underlying each of these influences and measurement of how dopamine-transporter blockade influences signal flow within each of these circuits.

### Implications for the series-circuit model of brain-reward circuitry

According to the series-circuit model (Fig 2), the rewarding effects produced either by electrical stimulation of the MFB or by optical stimulation of midbrain dopamine neurons arise from a common cause: activation of the dopamine neurons. The optical stimulation excites these neurons directly, whereas the electrical stimulation activates them transsynaptically. We have explained above that a reward-growth function with independent input-scaling and output-scaling parameters lies downstream from the dopamine neurons responsible for oICSS. If so, boosting the peak stimulation-induced dopamine concentration by means of transporter blockade should shift the contour map of the reward mountain downwards along the pulse-frequency axis in both cases. It does not. Whereas the predicted shift is seen in six of seven cases in the current oICSS results, reliable shifts in this direction are absent in all eight cases reported in the analogous eICSS study [45]. Similarly, reliable, downwards shifts were absent in seven of eight eICSS subjects tested under the influence of AM-251, a cannabinoid CB-1 antagonist that attenuates stimulation-induced dopamine release [57] and all six eICSS subjects tested under the influence of the dopamine-receptor blocker, pimozide [46]. This sharp discrepancy is highly problematic for the series-circuit model. If dopamine release in the terminal fields of midbrain dopamine neurons is the sole and common cause of the rewarding effect produced by electrical stimulation of the MFB and optical stimulation of midbrain dopamine neurons, why does the reward mountain respond so differently to perturbation of dopamine neurotransmission in the eICSS and oICSS studies?

Could the different mechanisms by which dopamine neurons are activated explain why dopamine-transporter blockade fails to shift the reward mountain along the pulse-frequency axis in rats working for electrical stimulation of the MFB but succeeds in doing so in rats working for optical stimulation of the ventral midbrain? We doubt that this is a viable explanation. To explain why we must first lay out in some detail this argument for rescuing the series-circuit hypothesis.

Given that ChR2 is found throughout the excitable regions of the cell membrane and that the optical probe is ∼10x larger than the soma of a dopamine neuron, the optically generated spikes likely arise downstream from the somatodendritic region. If dopamine-transporter blockade increased extracellular dopamine concentrations in the somatodendritic region, this would increase binding of dopamine to D2 autoreceptors and potentially decrease the sensitivity of these neurons to synaptic input. Such an effect could well be bypassed in the case of oICSS due to the activation of the ChR2-expressing neurons downstream from the somatodendritic region.

In order for such an explanation to account for the failure of GBR-12909 to shift the reward mountain along the pulse-frequency axis in rats working for electrical stimulation of the MFB, the putative autoreceptor-mediated inhibition would have to exactly counteract the drug-induced enhancement of dopaminergic neurotransmission in the terminal fields of those dopamine neurons that received sufficient excitatory synaptic drive to overcome the influence of the D2 stimulation. (Recall that the reward mountain was not shifted reliably along the pulse-frequency axis in either direction in all eight subjects in the eICSS study.) Such exact counterbalancing seems unlikely.

The impact of the dopamine transporter is very different in the somatodendritic and terminal regions of dopamine neurons. The rate of dopamine reuptake is as much as 200 times higher [74] and dopamine-transporter expression from 3–10 times greater [74, 75] in the terminal region than in the somatodendritic region. In a study carried out in guinea-pig brain slices, GBR-12909 failed to increase electrically induced release of dopamine in the VTA cell-body region [76], in contrast to its potent augmentation of release in terminal regions. This drug greatly boosted the amplitude of dopamine transients recorded voltammetrically in the nucleus accumbens terminal field of rats working for rewarding electrical stimulation of the VTA [70]. A recent study [77] shows very similar release of dopamine in the nucleus accumbens of rats working for electrical stimulation of either the VTA or of the MFB site used in the eICSS study of the effect of GBR on the reward mountain [45], suggesting that GBR-12909 would boost release of dopamine in the nucleus accumbens in response to electrical stimulation of the MFB site employed in the study of the effect of dopamine-transporter blockade on the position of the reward mountain [45]. Moreover, cocaine, which also blocks the dopamine transporter, greatly increased release of dopamine in the nucleus-accumbens shell in response to transsynaptic activation of midbrain dopamine neurons by electrical stimulation of the laterodorsal tegmental area [78, 79]. Taken together, the evidence makes it highly unlikely that in the reward-mountain study by Hernandez et al. [45], autoreceptor-mediated inhibition prevented GBR-12909 from boosting phasic release of dopamine in the terminal fields of midbrain dopamine neurons. Thus, the challenge posed by the present data to the series-circuit model stands, and alternative accounts must be explored.

### Toward a new model of brain-reward circuitry

To account for both the eICSS and oICSS data in the simulations, we propose development and exploration of models in which the reward-intensity signals evoked by electrical stimulation of the MFB and by optical stimulation of midbrain dopamine converge on a final common path. In such models, distinct reward-growth functions translate aggregate firing rate in the MFB neurons and the dopamine neurons into reward intensity. One such model is shown in Fig 10. The two reward-intensity signals (the output of the two reward-growth functions) converge onto the final common path underlying the behaviors entailed in approaching and holding down the lever. Thus, we are proposing that reward-related signals flow in parallel through a portion of the brain reward circuitry subserving ICSS and that the reward-intensity signal in the MFB limb of the circuit bypasses the midbrain dopamine neurons en route to the final common path.

Convergence models face a seemingly daunting challenge: electrical stimulation of the MFB activates midbrain dopamine neurons [71, 80–83]. If so, one would expect such stimulation to drive signaling in both of the hypothesized converging pathways. Wouldn’t this produce at least some displacement of the reward mountain along the pulse-frequency axis in response to dopamine-transporter blockade? The simulations documented in the accompanying Matlab^®^ Live Script show that this is not necessarily the case. Indeed, the simulations show that given reasonable assumptions and values drawn from the current data, a convergence model can replicate the findings reported here. The following paragraphs explain how.

In the eICSS studies entailing measurement of the reward mountain under the influence of drug-induced changes in dopamine neurotransmission, the train duration was 0.5 s, whereas it was 1.0 s in the current oICSS study. Gallistel [18] and Sonnenschein and colleagues [33] showed that the pulse frequency required to sustain half-maximal eICSS performance is a rectangular hyperbolic function of train duration. As shown in the Live Script, we used a rectangular hyperbolic function along with the two sets of chronaxie values from those two papers (which are in remarkable agreement) to estimate the change in 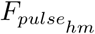 that would be expected from reducing the train duration from 1 s to 0.5 s. The median 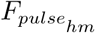 in the vehicle condition of the current study was 27.1 pulses s^-1^ at a train duration of 1.0 s; the estimated and simulated value at a train duration of 0.5 s for the vehicle condition is 37.6 pulses s^-1^. We then set 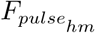 for the upper (MFB) limb of the convergence model (Fig 10) to a value typical of eICSS studies (∼ 77). The graphs in the upper row of Fig 11 were produced by setting the MFB drive to a level equivalent to an optical pulse frequency of 40 pulses s^-1^ (a bit above the estimated 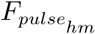 value for an oICSS train duration of 0.5 s). This yields the green reward-growth curve shown in the upper-left graph in Fig 11. Asymptotic reward intensity is quite low. One reason for this is that the lower-limb 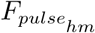 value is well within the roll-off range of the assumed frequency-following function. Another is that frequency-following fidelity in the MFB neurons that generate transsynaptic excitation of the midbrain dopamine neurons is poorer than in neurons that produce the rewarding effect of electrical MFB stimulation [80]. The magenta curve is the sum of the outputs of the upper (cyan curve) and lower (green curve) limbs.

**Fig 11.**
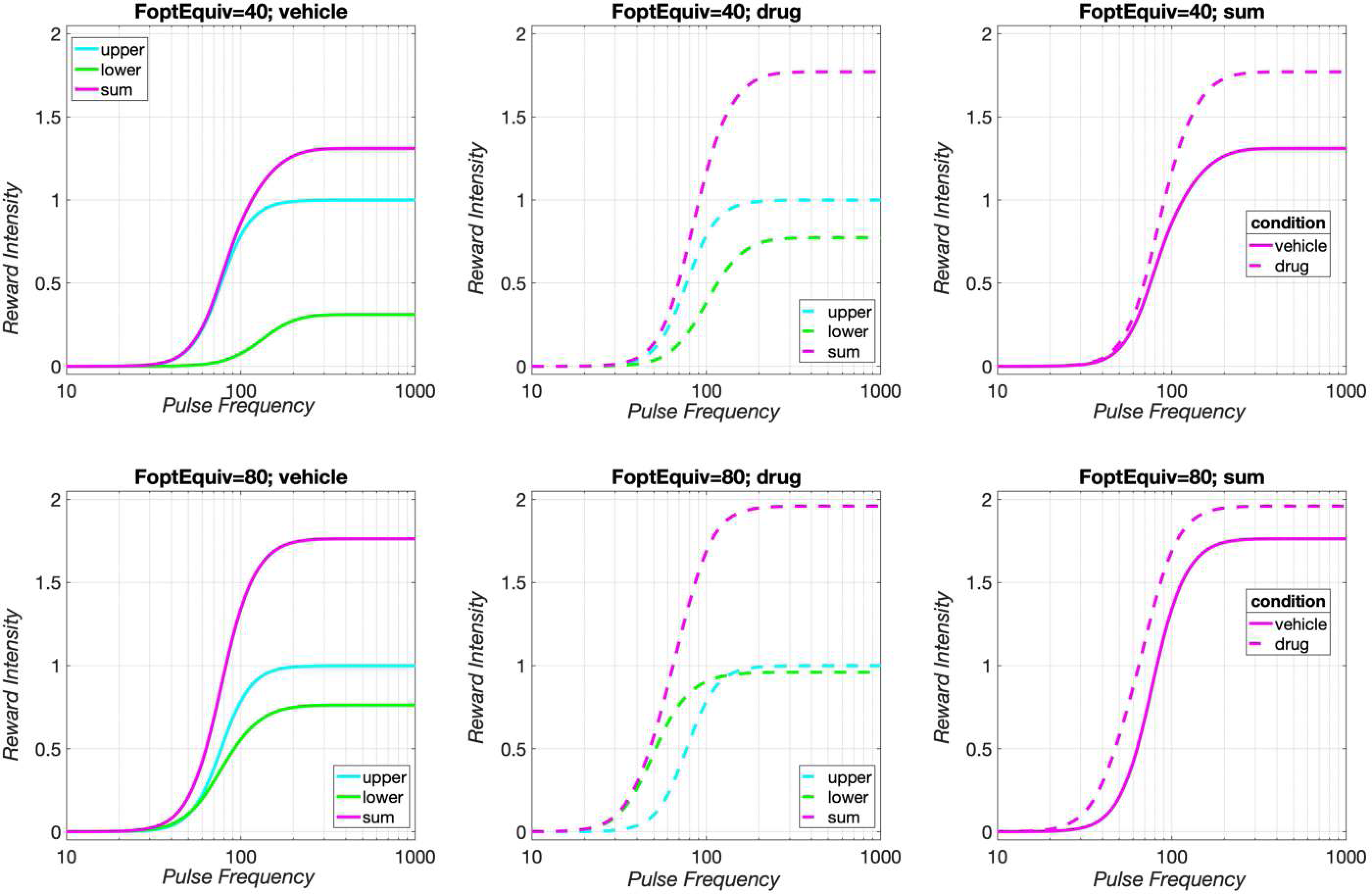
Reward growth in the convergence model. The green curves show the growth of reward intensity in the dopaminergic limb of the convergence model (Fig 10, the cyan curves show the growth of reward intensity in the MFB limb, and the magenta curves show the sum of the reward-intensity values in the two limbs. In the upper row, the strength of the MFB drive is equivalent to optical stimulation more than sufficient to generate half-maximal reward intensity, whereas in the lower row, the MFB drive is equivalent to the strongest optical stimulation employed in the present study. The effect of increased tonic-dopamine signaling is presumed to arise from decreases in subjective effort costs or the value of alternate activities but not from upwards rescaling of reward-intensity. Thus the vertical asymptotes of the cyan curves are not altered by the drug.

**Figure 12.**
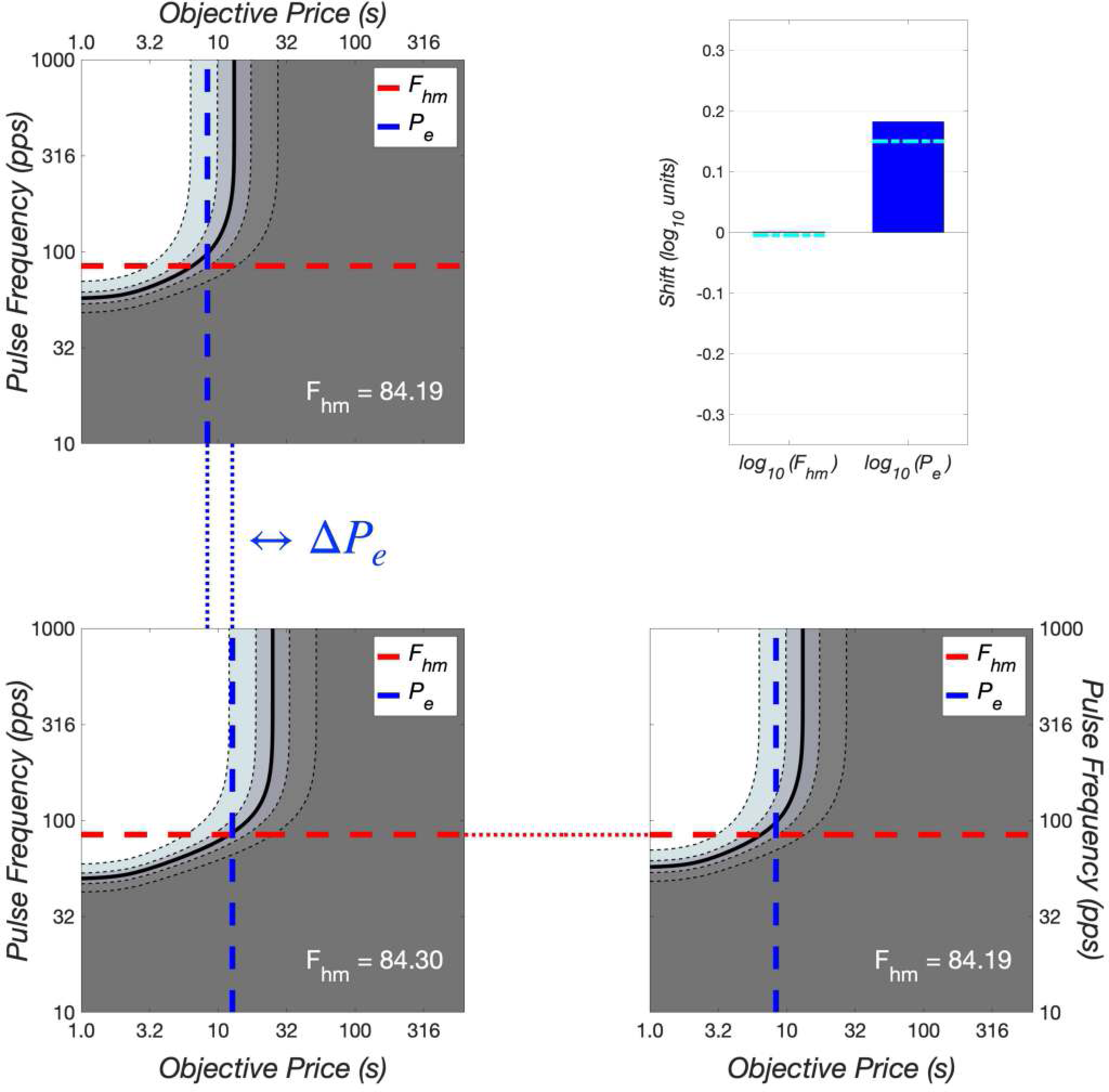

The middle graph in the upper row of Fig 11 shows the simulated effect of dopamine-transporter blockade, with *K*_*da*_ set to 1.4125 in the simulation (estimated value in the current study: 1.3951). The simulated effect of the drug shifts the green reward-intensity curve leftwards. Due to the improved frequency-following fidelity in the drug condition, this boosts the upper asymptote of both the dashed green curve for the lower limb and the dashed magenta curve for the summated output of the two limbs. As a result, the reward mountain shifts rightward along the price axis (Fig S35). (In the simulations, the correction for changes in frequency-following fidelity removes this component of the shift along the price axis, as shown by the dashed cyan line in the bar graph in Fig S35. The remaining component is due to simulated drug-induced reduction in subjective effort costs.)

The left shift along the pulse-frequency axis in Fig 11 is insufficient to overtake the cyan curve for the upper (MFB) limb of the model. Thus, the summated curve for the drug condition (dashed magenta lines in the upper middle and upper right panels) is not shifted laterally with respect to the summated curve for the vehicle condition (solid magenta lines). As a consequence, the simulated reward mountain does not shift along the pulse-frequency axis (Fig S35). Thus, the simulated output obtained using moderate MFB drive on the dopamine neurons replicates the eICSS findings.

What would happen if the MFB drive were much stronger? We repeated the simulations setting the MFB drive to the equivalent of an optical pulse frequency of 80 pulses s^-1^, around the highest value tested in most of the subjects in the current study. The results are shown in the lower row of Fig 11. The reward-growth curve for the lower (dopaminergic) limb of the model (green curve) is now shifted leftwards in the vehicle condition (lower-left panel) and rises to a higher asymptote. From its enhanced starting position along the abscissa, the simulated reward-growth curve for the lower limb in the drug condition is now able to overtake the reward-growth curve for the upper limb, shifting the dashed green curve for the lower limb (middle panel, bottom row) to the left of the dashed cyan curve for the upper limb. As a result, the summated curve for the drug condition (dashed magenta lines, bottom row) is displaced somewhat to the left of the summated curve for the vehicle condition (solid magenta lines, bottom row). This shift drags the reward mountain a short distance down the pulse-frequency axis (Fig S36). Thus, given transsynaptic MFB drive that is sufficiently strong to match extremely intense, direct, optical stimulation, some shift along the pulse-frequency axis is predicted.

This dependence of the output of the convergence model on parameter values is of interest. Although failure to observe shifts along the pulse-frequency axis was the most common result of the experiments in which the eICSS reward-mountain was measured under the influence of drugs that alter dopaminergic neurotransmission (27 of 32 cases reported in [38, 45, 46, 57]), it is not the only result. Reward mountains obtained from three subjects in the cocaine study [38] showed fairly substantial, reliable, shifts along the pulse-frequency axis. (The shift was marginally reliable in a fourth subject when tested initially but disappeared upon re-test.) Although no subject in the GBR-12909 [45] or pimozide [46] studies showed such shifts, one subject in the AM-251 study [57] did. Could variation in the strength of MFB drive on the dopamine neurons explain these apparently discrepant findings?

Taken together, the results of the simulation performed at two levels of MFB drive on the dopamine neurons cast the convergence model in a promising light. With parameter values drawn from the available empirical data (and some as-yet unavoidable generalization from eICSS data to the oICSS case), it can account both for the dominant result of the empirical studies (Fig 11, upper row, Fig S35) as well as for the few systematic deviations from the general pattern. The emergence of such deviations depends on the strength of the MFB drive and the position of the curve for the upper limb along the abscissa as well as on the frequency-following fidelity of both limbs and the fibers that connect them. Thus, a readily testable prediction of the convergence model is that shifts of the eICSS reward mountain in response to perturbation of dopaminergic neurotransmission should be more common when longer train durations are employed. (Lengthening the train duration allows for better frequency-following fidelity in the lower limb as well as in the link that relays MFB drive to the dopamine neurons.) Similarly, higher drug doses should increase the prevalence of such shifts.

The convergence model avoids a puzzling feature of the series-circuit model: the apparent imposition of a processing bottleneck in brain-reward circuitry. Large, myelinated, fast-conducting fibers are expensive metabolically, and they take up valuable neural real-estate. Why funnel the output of such a costly, high-bandwidth pathway exclusively through lower bandwidth dopamine neurons?

In contrast to the convergence model, the series-circuit model does not account for the combined data from the eICSS and oICSS reward-mountain studies entailing pharmacological alteration of dopaminergic neurotransmission. In the series-circuit model, any manipulation that changes phasic dopamine release, or the post-synaptic consequences of this release, should shift the reward mountain along the pulse-frequency axis. Thirty-three of the 38 subjects of the five eICSS and oICSS studies entailing measurement of the reward mountain under pharmacological challenges argue otherwise.

#### A new home for previously orphaned findings?

The convergence models can account for a heretofore unexplained discrepancy between dopamine release and self-stimulation performance. In rats performing eICSS of the MFB, Cossette and colleagues traded off the stimulation current against the pulse frequency [80]. As expected on the basis of the counter model, increases in pulse frequency required that the current be reduced in order to hold behavioral performance constant. In accord with a more detailed study of the frequency response of the MFB neurons subserving eICSS [8], the required current continued to decline as the pulse frequency was increased beyond 250 pulses s^-1^. In contrast, stimulation-evoked dopamine release in the nucleus accumbens, monitored by means of fast-scan cyclic voltammetry, increased very little, or not at all, as the pulse frequency was increased from 120 to 250 pulses s^-1^ and generally declined when the pulse frequency was increased further to 1000 pulses s^-1^. In the series-circuit model, midbrain dopamine neurons relay signals from the directly activated MFB neurons to the behavioral final common path. Thus, the trade-off between the induced firing frequency and the number of directly activated MFB neurons recruited by the current should be manifested faithfully in the stimulation-induced release of dopamine. It was not. The discrepancy between the trade-off functions for eICSS and dopamine release is not explained by the series-circuit model but is readily accommodated by the convergence model.

Huston and Borbély documented rewarding effects produced by electrical stimulation of the lateral hypothalamic level of the MFB in rats that had undergone near-total ablation of telencephalic structures [84, 85]. Although the major telencephalic terminal fields of the midbrain dopamine neurons had been damaged heavily, the rats learned to perform simple movements to trigger delivery of the electrical stimulation. The series-circuit hypothesis leaves these data unexplained, but the convergence model could accommodate them. For example, hypothalamic or thalamic neurons that survived the ablations might relay reward-related signals to brainstem substrates of the behavioral final common path via MFB fibers, as Huston proposed.

Johnson and Stellar [86] made large, bilateral, excitotoxic lesions in the nucleus-accumbens (NAC) and ventral pallidum (VP) of rats that had been trained to perform eICSS of the lateral hypothalamic level of the MFB. The series-circuit model predicts that such extensive damage to a key dopaminergic terminal fields should have greatly reduced the rewarding effectiveness of the electrical stimulation. It did not. The authors concluded that “while the NAC and VP have been shown to be important for various kinds of reward,” … they do “not appear to be critical for the expression of ICSS reward. It may be the case that the ICSS reward signal is processed downstream from the NAC and VP and is therefore unaffected by total destruction of either structure.” That conclusion contradicts the series-circuit hypothesis but it is perfectly compatible with, and indeed anticipates, the convergence model presented here.

### The essential role of modeling and simulation

It is common in behavioral neuroscience to make predictions and assess results on the basis of binary, directional classification. The effect of a manipulation either does or does not meet a statistical criterion. If it does, the value of the output variable is said to go up or go down. The numeric values of the input and output variables typically matter little.

The reward-mountain model is more ambitious. It makes predictions about the form of the relationships between the variables. As is so often the case in biology, the relationships are non-linear. There are ranges of an input variable, such as the pulse frequency, over which an output, such as time allocation, changes little and other ranges over which the output changes a lot. Thus, the snide answer to the question of whether an output will change is “it depends,” and the serious answer entails specifying this dependency in terms of functional forms and their parameters. The numeric values matter.

Albeit in a modest way, the reward-mountain model confronts the problem of convergent causation, the fact that although behavioral output is highly constrained by the relatively small number of muscles, joints and degrees of freedom in an animal body, a very large number of neural controllers have access to the behavioral output. A given level of reward pursuit may be directed at a large, but expensive, reward or at a small, inexpensive one. The reward-mountain model distinguishes such cases by tying reward pursuit to both the strength and costs of reward, and it goes beyond binary and directional classification of effects by specifying the functions that map these physical quantities into the subjective values that determine goal selection and behavioral allocation.

The reward-mountain model is quantitative: given inputs consisting of reward strengths and costs, it outputs time-allocation values. This output changes in specified, lawful ways when internal parameters, such as the scaling of reward intensities, are perturbed by manipulations, such as drug administration. A virtue of such an approach is that it requires explicit statement of assumptions. A shortcoming, perhaps, is that the specification of the numerous, unavoidable assumptions tends to elicit skepticism that may be eluded by verbal formulations that allow assumptions to remain unstated and implicit. In our view, it is best to make our ignorance manifest so as to incite ourselves and our colleagues to reduce it.

Another sense in which we believe the quantitative specification of the model to be important concerns the often-hidden perils of relying on verbal reasoning alone to predict the behavior of systems embodying multiple, interacting, non-linear components [38, 52]. Verbal reasoning is not up to this task. Instead, it requires careful formal, quantitative specification and demonstration of feasibility via simulation, which can reveal lacunae of which the modeler may have been unaware and generate interesting and unexpected results that inspire new experiments. Perhaps most important is that simulation reveals how the model works in a way that verbal descriptions do not capture, and it provides a strong test of whether, in principle, the model actually does what its designers have intended. In the accompanying Matlab^®^ Live Script, we provide the code for performing simulations of the reward-mountain model. We invite the reader to experiment with the code and thereby critique the model.

The simulations lead us to a view rather different than the one attributed to Otto von Bismark concerning the making of democratic laws and the fabrication of sausages. We believe that when modeling decisions about goal selection and pursuit, very close attention should be paid to “how the sausage was made.” For example, it was only by detailed analysis of the internal workings of the convergence model that we came to appreciate the plausibility of its counter-intuitive prediction: stability of the reward-mountain along the pulse-frequency axis under moderate MFB drive in the face of drug-induced enhancement of dopamine signaling.

Diagrams such as Fig 10 include a large number of components and may invite the viewer to imagine William of Ockham turning in his grave. That said, parsimony entails the jettisoning of ***superfluous*** entities. We invite the reader to identify which of the components in the models can be tossed overboard without loss of explanatory power. Can the data from previous eICSS experiments and the present oICSS experiment be explained more simply? If we had found an affirmative answer to this question we would have implemented it. Indeed, we see the models discussed here as over-simplified rather than over-complicated. For example, they say nothing about fundamental matters such as the functional specialization of dopamine subpopulations, how signals from the “milieu interne” modulate the decision variables, or how the subjects learn and update the reward intensities and costs that determine their behavioral allocation. Nonetheless, we argue that the combination of the modeling, simulation and empirical work provides a new perspective on the structure of brain-reward circuitry while challenging a long-established view.

### Finding the MFB substrate

The notion that non-dopaminergic neurons with myelinated axons predominate in the directly stimulated substrate for eICSS of the MFB was proposed in the 1980s on the basis of behaviorally derived estimates of conduction velocity, recovery from refractoriness, and direction of conduction; [10–14, 18]. A 1974 review includes a suggestion that the directly stimulated substrate may be non-dopaminergic [87]. The discrepancy between the behaviorally effective range of pulse frequencies and the frequency-following fidelity of catecholatminergic neurons was also noted during the 1970s [88]. Since that time, little progress has been made toward identifying the directly activated neurons responsible for eICSS of the MFB, although electrophysiological recordings show the rough location of some somata that give rise to fibers with properties compatible with the psychophysically based characterization [89–91]. One likely reason is that the series-circuit model relegates these neurons to a subsidiary role that has thus inspired little empirical investigation: on that view, the MFB fibers merely provide an input, likely one of many [92, 93], to the dopamine neurons ultimately responsible for the rewarding effect.

The convergence model elevates the status of the directly activated neurons subserving the rewarding effect of MFB stimulation. This model asserts that multiple, partially parallel, neural circuits can generate reward and that the dopamine neurons do not constitute an obligatory stage in the final common path for their evaluation and pursuit. From that perspective, it is important to intensify the search for the limb(s) of brain reward circuitry that may parallel the much better characterized dopaminergic pathways. Application of modern tracing methods that integrate approaches from neuroanatomy, physiology, optics, cell biology and molecular biology (e.g., [94]) may well achieve what application of the cruder, older tools failed to accomplish. The detailed psychophysical characterization of the quarry that has already been achieved, particularly the evidence for myelination and axonal trajectory [10–14, 89], can guide the application of such methods.

### Potential implications of parallel channels in brain-reward circuitry

Remarkable success has been achieved in developing tools for specific excitation or silencing of dopaminergic neurons, measuring the activity of these neurons, mapping the circuitry in which they are embedded, and categorizing different functional dopaminergic subpopulations. The neurobiological study of dopamine neurons has been coupled to learning theory, neural computation, and psychiatry, with longstanding application in the study of addictive disorders [6, 16, 95, 96] and emerging linkage to the analysis of depression [97, 98]. Perhaps the brilliance of these successes has obscured the possible roles played by other neurons and circuits in the functions in which dopaminergic neurons have been implicated. We propose that the network in which dopamine neurons are embedded is not the sole source of input to the behavioral final common path for the evaluation and pursuit of rewards. What roles might the alternative sources play in behavioral pathologies and behaviors essential to well being? Addressing that question requires that the existence of such networks be appreciated and addressed, their constituents identified, and their function understood.

## Conclusion

The reward-mountain model was developed to account for data from eICSS experiments in which rats worked for rewarding electrical stimulation of the MFB. At the core of the model is a logistic reward-growth function that translates the aggregate impulse flow induced by the electrode into a neural signal representing the intensity of the reward. On the basis of the direction in which a drug shifts the reward mountain within a space defined by the strength and opportunity cost of reward, the model distinguishes drug actions at, or prior to the input to the reward-growth function from actions at, or beyond, the output. Bidirectional perturbations of dopaminergic neurotransmission have acted selectively in the latter manner, either by rescaling the output of the reward-growth function or by altering other valuation variables, such as subjective reward costs or the attraction of alternate activities: Dopamine-transporter blockers [38, 45] and a dopamine-receptor antagonist [46] shifted the mountain along the axis representing opportunity cost and not along the axis representing pulse-frequency.

The present study demonstrates that the reward-mountain model also provides a good description of how the strength and opportunity cost combine to determine the allocation of behavior to pursuit of direct, optical activation of midbrain dopamine neurons. The results argue that as in the case of eICSS, a logistic-like function with independent input-scaling and output-scaling parameters translates the neural excitation induced by the stimulation into the intensity of the rewarding effect. Unlike the case of eICSS, augmentation of dopaminergic neurotransmission by dopamine-transporter blockade acts as if to rescale the input to the reward-growth function: the reward mountain was shifted along the pulse-frequency axis. In one sense, this is not surprising: by boosting the peak amplitude of stimulation-induced, dopamine-concentration transients in terminal fields, the drug should be expected to reduce the pulse frequency required to generate a reward of a given intensity. However, this result challenges a longstanding model of brain-reward circuitry.

According to the series-circuit model, the rewarding effect of electrical MFB stimulation arises from the activation of highly excitable, non-dopaminergic axons that provide direct or indirect synaptic input to midbrain dopamine neurons; the rewarding effect arises from the transsynaptic activation of these dopamine neurons. The current results show that when these dopamine neurons are activated directly, dopamine-transporter blockade shifts the reward mountain along the pulse-frequency axis. We argue that dopamine-transporter blockade should produce a similar shift in the position of the reward mountain when the dopamine neurons are activated transsynaptically. However, perturbation of dopaminergic neurotransmission has generally failed to shift the reward mountain along the pulse-frequency axis when electrical stimulation of the MFB is substituted for optical stimulation of the midbrain dopamine neurons. Thus, the series-circuit model cannot readily accommodate the results of both the eICSS and oICSS experiments.

We propose that alternatives to the series-circuit model be explored. In the one sketched here, the reward signal carried by the MFB axons runs parallel to the reward signal carried by the midbrain dopamine neurons prior to the ultimate convergence of these two limbs of brain-reward circuitry onto the final common path for the evaluation and pursuit of rewards. This proposal can accommodate findings unexplained by the series-circuit model and suggests a research program that could complement the work that has so powerfully and convincingly implicated dopaminergic neurons in reward.

## Materials and methods

### Subjects

Seven TH::Cre, male, Long-Evans rats weighing 350 g at the time of surgery, served as subjects. Animals were obtained from a TH::Cre rat colony established from three sires generously donated by Drs. Ilana Witten and Karl Deisseroth. Upon reaching sexual maturity, animals were housed in pairs in a 12 h −12 h reverse light-cycle room (lights off at 8:00 AM). The rats were housed singly following surgery.

### Ethics Statement

All experimental procedures were approved by the Concordia University Animal Research Ethics Committee (Protocol #: 30000302) and conform to the requirements of the Canadian Council on Animal Care.

### Surgery

Midbrain dopamine neurons were transfected with the light-sensitive cation channel, *channelrhodopsin 2* (ChR2), fused to the reporter protein, *enhanced yellow fluorescent protein* (eYFP). The construct was delivered by means of a Cre-dependent Adeno-Associated Viral vector (AAV5-DIO-ChR2-EYFP, University of North Carolina Viral Vector Core, Chapel Hill, NC). The virus was injected bilaterally (±0.7 mm ML), at a volume of 0.5 µl, at three different DV coordinates (−8.2, −7.7 and −7.2 mm) and two different AP coordinates (−5.4 and −6.2 mm), to yield a total volume of 3.0 µl per hemisphere. Optical-fiber implants, with a core diameter of 300 µm, were aimed bilaterally at the VTA at a 10° angle (AP: −5.8, DV: 8.02 or 8.12, ML: ± 0.7 mm). Anesthesia was induced by an i.p. injection of a Ketamine-xylaxine mixture (87 mg/kg, 13 mg/kg, Bionicle, Bellville, Ontario and Bayer Inc., Toronto, Ontario, respectably). Atropine sulfate (0.02-0.05 mg/kg, 1 mL/kg, Sandoz Canada Inc., Quebec) was injected s.c. to reduce bronchial secretions, and a 0.3 mL dose of penicillin procaine G (300 000 IU/ml, Bimeda-MTC Animal Health Inc., Cambridge, Ontario) was administered SC, as a preventive antibiotic. “Tear gel” (1% w/v, ‘HypoTears’ Novartis) was applied to the eyes to prevent damage from dryness of the cornea. Anesthesia was maintained throughout surgery by means of isoflurane (1 − 2.5% + O_2_). The head of the rat was fixed to the stereotaxic frame (David Kopf instruments, Tujunga, CA) by means of ear bars inserted into the auditory canal and by hooking the incisors over the tooth bar. Bregma and Lambda were exposed by an incision of the scalp. Three blur holes were drilled in the skull over each hemisphere (AP: −5.4, −5.8, and −6.2 mm; ML:± 0.7,± 2.08,± 0.7, respectively). A 28 gauge injector was loaded with the viral vector. Six 0.5 µL boli of the virus-containing suspension were infused into each brain hemisphere at the following coordinates: AP: −5.4 and −6.2 mm; ML: ± 0.7 mm; DV: −8.2, −7.7 and −7.2 mm. Infusions were performed at a rate of 0.1 µL per minute using a precision pump (Harvard Instruments) and a 10 µL Hamilton syringe (Hamilton Labaoratory prodcuts, Reno, NV). To allow for diffusion, the injector was left in place for ten minutes following each infusion. Optical-fiber implants with a 300 µm, 0.37 numerical-aperture core were constructed following the methods described by Sparta et al. [99]. Optical fibers were aimed bilaterally at the VTA at a 10° angle. The implants were placed at two different DV coordinates to increase the chances of placing the tip of at least one of the optical fibers directly over the neurons that support optical self-stimulation (AP −5.8 mm; ML ± 0.7 mm; DV −8.02 and −8.12 mm). The optical implants were anchored to the skull by means of stainless steel screws and dental acrylic. Gelfoam^™^ (Upjohn Company of Canada, Don Mills, Ontario) was used to fill the holes in the skull and promote healing. Buprenorphine (0.05‘mg/kg SC, 1 mL/kg, RB Pharmaceuticals Ltd., Berkshire, UK) was used as a post-surgery analgesic. The rats were housed singly in the animal care facility for over five weeks to allow for surgical recovery and to achieve appropriate expression and distribution of the protein product of the ChR2-EYFP construct.

### Apparatus

The operant chambers (30 × 21 × 51 cm) had a mesh floor and a clear Plexiglas front equipped with a flashing light located 10 cm above the floor mesh, and a retractable lever (ENV–112B, MED Associates) mounted on a side wall. A 1 cm light was located 2 cm above the lever and was activated when the rat depressed the lever. A blue DPSS laser (473 nm, Shanghai Lasers and Optics Century Co. or Laserglow Technologies, Toronto, ON) was mounted on the roof of each box. The laser was connected to a 1 × 1 FC/M3 optical rotary joint (Doric lenses, Quebec, Canada) by means of a laser coupler (Oz Optics Limited, Ottawa, ON, or Thorlabs, Inc., Newton, New Jersey, USA) and fiber-optic patch cords. Robust, custom-built, optical-fiber patch cords designed for rats [100] were used to attach the implants in the animal’s head to the 1 × 1 FC/M3 optical swivel so as to allow the rat to move without tangling the cable. Experimental control and data acquisition were handled by a personal computer running a custom-written program (“PREF”) developed by Steve Cabilio (Concordia University, Montreal, QC, Canada). The temporal parameters of the electrical stimulation were set by a computer-controlled, digital pulse generator. Stimulation consisted of 1 s trains optical pulses, 5 ms in duration.

### Drug

GBR-12909 was donated generously by the NIMH Chemical Synthesis and Drug Supply Program. It was dissolved in 0.9% saline at a volume of 10 mg/ml. The pH of the solution was adjusted to 5± 0.1 with 0.1M NaOH.

### Self-stimulation screening and initial training

Each animal underwent two to three screening sessions in which only one of the optical implants was attached to the laser. Optical power was measured by means of an optical power meter (PM100D, Thorlabs Inc., Newton, New Jersey, USA, modified to reduce spurious output noise) and adjusted through trial and error for each rat to elicit robust oICSS behavior (30-60 mW, measured at the tip of the patch-cord with the laser operating in continuous-wave mode). Animals were trained by means of the successive approximation procedure to depress the lever to receive optical stimulation (a 1 sec train of 5 ms optical pulses at 80 pulses per second (pps). After the rats had learned to lever press, they were allowed to work for the optical reward, on a Continous Reinforcement (CRF) schedule, during two 15 min trials. The total number of presses was recorded. Both of the implants were tested under these conditions; the implant that yield the larger number of presses was used for the rest of the experiment.

### Training in preparation for measurement of the reward mountain

The rats received further training to prepare them for measurement of the reward mountain (Fig. S37). First, the rats were trained to perform a new reward-procuring response, holding down the lever rather than simply pressing it briefly. This new response was rewarded according to a cumulative-handling-time schedule of reinforcement [101], which delivers a reward when the cumulative time the lever has been depressed reaches an experimenter-defined criterion called the “price” of the reward. In this sense, the price corresponds to what economists call an opportunity cost. To earn a reward, the rats did not have to hold down the lever continuously until the price criterion was met; they could meet the criterion by means several bouts of lever holding separated by pauses. The onset and offset times of each bout of lever depression were recorded.

Next, the values of one or both independent variable (pulse frequency, price) were varied sequentially from trial to trial, thus traversing the independent-variable space along a linear trajectory. Such a traversal is called a “sweep.” Three types of sweeps were carried out. Frequency sweeps were carried out at a fixed price (initially 1-2 s). These sweeps consisted of 10 to 12 trials during which the rat had the opportunity to harvest as many as 60 rewards (except for rat BeChr19, who was allowed to harvest a maximum of 30 rewards per trial due to the unusual effectiveness of optical stimulation in this subject). Each reward was followed by a 2 s *Black Out Delay* (BOD) during which the lever was disarmed and retracted, and timing of the trial-duration was paused. The pulse frequency during the first two trials was set to yield maximal reward-seeking behavior. The first trial was considered a warm-up trial and was excluded from analysis. From the second trial onwards, the rewarding stimulation was decreased systematically from trial to trial by decreasing the pulse frequency in equal proportional steps. The range of tested pulse frequencies was selected as to drive reward-seeking behavior from its maximal to its minimal value in a sigmoidal fashion (S37B). Every trial was preceded by a 10 s *Inter-Trial Interval* (ITI) signaled by a flashing light. During the last 2 s of this period rats received priming stimulation consisting of a non-contingent, 1 s stimulation train, delivered at the maximally rewarding pulse frequency. Rats performed one frequency sweep per session, consisting of ten trials (except for subject BeChr19 who was tested on 12 trials per frequency sweep).

When the rats showed consistent performance across trials and sessions in the frequency-sweep condition, we introduced price sweeps into the training sessions. In the price-sweep condition, the pulse frequency was kept constant at the maximal value for each rat, but the cumulative work time required to harvest the reward (i.e. the price of the reward) was increased systematically across trials. The price was the same on the first two trials of each sweep. As in the frequency-sweep condition, the first trial was considered a warm-up trial and was excluded from analysis. Starting at the second trial of the sweep, prices were increased by equal proportional steps across trials. The prices were set by trial and error so as to yield a sigmoidal transition between maximal and minimal reward-seeking behavior as a function of price (Fig. S37B). Trial duration was set so as to allow the rats to harvest a maximum of 60 rewards per trial (30 in the case of BeChr19). The BOD, ITI, and priming were the same as in the frequency sweep. During price-sweep training sessions, the rats also performed a frequency sweep. The order of the sweeps was randomized across sessions.

Radial sweeps were incorporated when performance on the price sweeps appeared stable. Along radial sweeps, the pulse-frequency was decreased and the price was increased concurrently across trials. Thus, frequency sweeps run parallel to the pulse-frequency axis in the independent-variable space, price sweeps run parallel to the price axis, and radial sweeps run diagonally. The radial sweeps were composed of ten trials; the first trial served as a warm-up and was identical to the second trial. From the second trial onwards, both pulse frequency and price were varied in equal proportional steps so as to yield a sigmoidal decrease in reward-seeking behavior over the course of the sweep (Fig. S37B). The trajectory of the vector defined by the tested pulse frequencies and prices was aimed to pass as close as possible to the point defined by the estimated values of the 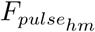 and 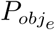 location parameters (see model fitting section). This required fitting the mountain model to preliminary data from each rat and adjusting the pulse frequencies and prices tested along the radial sweep accordingly for the following session. Pulse-frequency and price sweeps were also performed in each session during this phase of testing. The order of presentation of the three different sweeps was random across sessions, and the BOD, ITI, and priming parameters were the same on all trials. Rats were considered ready for drug-test sessions when reward-seeking behavior declined sigmoidally and consistently along all three sweeps and the trajectory of the radial sweep in the independent-variable space passed close to the point defined by the estimated values of 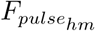 and 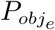.

### Effects of GBR-12909 on the reward mountain

Rats received i.p. injections 90 min prior to behavioral testing. Vehicle (2.0 ml/kg) was administered on Mondays and Thursdays and GBR-12909 (20 mg/kg) on Tuesdays and Fridays. In each session, rats performed a frequency, a price, and a radial sweep in random order. Each sweep consisted of ten trials each (except for the frequency sweep for rat BeChr19, which consisted of 12 trials). The duration of each trial was set so as to allow rats to harvest a maximum of 60 rewards per trial (except for rat BeChr19, who was allowed to harvest a maximum of 30 rewards per trial due to his unusual proclivity to work for very high opportunity costs). Wednesdays and weekends were used as drug elimination days: no testing was conducted on these days, and the rats remained in the animal care facility. Ten vehicle and ten drug sessions were conducted with each rat.

### Calculation of time allocation

The raw data were the durations of “holds” (intervals during which the lever was depressed by the rat) and “release times” (intervals during which the lever was extended but not depressed by the rat). Total work time included 1) the cumulative duration of hold times during a trial, and 2) release times less than 1 s. The latter correction was used because during very brief release intervals, the rat typically stands with its paw over or resting on the lever [101]. Therefore, we treat these brief pauses as work and subtract them from the total release time. Corrected work time for a given trial was defined as the sum of the corrected hold times, and leisure time was defined as the sum of the corrected release times. The dependent measure was time allocation (TA), the ratio of the corrected work and leisure times.

### Model fitting and comparisons

A separate TA calculation was performed for each *reward encounter*: the time between extension of the lever and completion of the response requirement (when cumulative work time equals the set price). The primary datasets thus consisted of the TA values for the reward encounters that occurred during all trials and sessions run following administration of the drug or vehicle. These primary datasets were then resampled 250 times with replacement [38, 102].

#### Fixed parameters

The mountain model and the fitting approach have been described in detail elsewhere [37, 38]. Two versions of the extended reward-mountain model [37] were fit to the present data. Both models include four fixed parameters, two describing the subjective-price function [39] and two describing the frequency-following function [8]. The subjective-price function maps the objective price of the reward into its subjective equivalent. Here, we used the form and parameters of obtained for this function in a study of eICSS of the MFB [39]. The frequency-following function maps the optical pulse frequency into the induced frequency of following in the midbrain dopamine neurons. The function we used here was of the same form as the one described in a study of eICSS of the MFB [8], but different parameter values were required to accommodate the differences between the frequency responses of optically stimulated ChR2-expressing midbrain dopamine neurons and electrically stimulated MFB neurons subserving eICSS. For details, please see the section entitled “Parameters of the frequency-following function for oICSS” in the supporting-information file.

#### Fitted parameters

The first (“standard”) model includes six fitted parameters. The location parameters 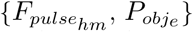 position the mountain along pulse-frequency and price axis, respectively (Fig. 1B), the a and g parameters determine the slope of the mountain surface, and the T_*min*_ and T_*max*_ parameters determine its minimum and maximum altitudes, respectively. The second (“CR”) model includes an additional parameter, CR, that estimates the contribution of conditioned reward [38, 45]. This parameter selectively increases time allocation to pursuit of weak, inexpensive rewards, thus providing a more accurate fit when the lever and/or the act of depressing it become potent secondary reinforcers.

#### Common versus treatment-dependent parameters

Models containing many parameters can prove excessively flexible, and fits employing them may fail to converge. We restricted the flexibility of the models by fitting common values of T_*min*_ and T_*max*_ to the data from the two treatment conditions {vehicle, drug}. The rationale is that the factors causing T_*min*_ to deviate from zero and T_*max*_ to deviate from one tend to be be common across vehicle- and drug-treatment conditions. The main experimental question posed concerns the effect of the drug on the location parameters. Thus, these two parameters 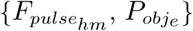 were always free to vary across treatment conditions. Variants of both the standard and CR models were produced in which all, some, or none of the a, CR, and g parameters were common across the two treatment conditions. Thus, twelve models were fit: four variants of the standard model

1. *a* free, *g* common
2. *g* free, *a* common
3. both *a* and *g* free
4. neither *a* nor *g* free

and eight variants of the CR model (the same combinations as for the standard model, but with CR either free or common).

All 12 models were fit to each of the 250 resampled datasets using a procedure developed by Kent Conover, based on the nonlinear least-squares routine in the MATLAB optimization toolbox (the MathWorks, Natick, MA). Mean values for each parameter were obtained by averaging the 250 estimates. Confidence intervals (95%) were estimated by excluding the lowest and the highest 12 values of the 250 estimates. This yields unbiased estimates of the fitted parameters and their dispersions for each subject. The Akaike information criterion [53] was used to select the model that offers the best balance between achieving a good fit and minimizing the number of parameters required to do so. Drug-induced shifts in the location of the 3D structure were considered significant when the 95% confidence interval around the difference between the 250 resampled estimates of the location parameters across drug and vehicle conditions excluded zero (i.e. no difference between conditions).

### Power-frequency trade-off

Following the pharmacological experiment, we explored the effects of systematic changes in optical power and pulse frequency on the number of rewards obtained by each rat. The same subjects were trained to press the lever on an FR-1 schedule to trigger optical stimulation of midbrain dopamine neurons through the same optical implant used in the reward-mountain experiment. We quantified the number of responses emitted in each of a series of nine or ten trials lasting two minutes each. The pulse frequency was decreased systematically across trials: the highest pulse frequency was in effect during the first trial of the series (the “warm-up”) and also on the second trial, and data from the first trial was excluded from analysis. Subsequently, the pulse-frequency decreased in equal logarithmic steps from trial to trial. A single pulse-frequency sweep was run in each of five to six sessions per day. In each session, the optical power (measured at the tip of the patch-cord with the laser operating in continuous-wave mode) was set to one of five values (1.87, 3.75, 7.5, 15 or 30 mW for rat Bechr14; 3.75, 7.5, 15, 30, or 60 mW for rats Bechr21, Bechr28 and Bechr29; and 2.5, 5, 10, 20, and 40 mW for rats Bechr19, Bechr26, and Bechr27). The rats were tested under these conditions for five days. The order of presentation of optical powers was determined pseudorandomly for each rat. Each power was presented in a different sequential order across each testing day (i.e. each power was used 1st, 2nd, 3rd, 4th, or 5th at least once across days, but the preceding and/or subsequent tested powers may have been different across test days). To control for carry-over effects across test sessions, rats were given 30-min breaks between each session: following completion of each of the five to six daily sessions, they were taken out of the operant chambers, brought back to their home cage in the animal-care facility, and had free access to food and water for 30 min before resuming with the following test session. During this 30-min break, the lasers were set to continuous operation mode to minimize fluctuation in optical power due to the cooling of the laser.

### Histology

Rats were sacrificed by means of a lethal injection of pentobarbital i.p. After deep anesthesia had been induced, the rats underwent intracardiac perfusion with phosphate-buffered saline and 4% paraformaldehyde chilled to 4 °C. Upon extraction, the brains were cryoprotected in a solution of 4% paraformaldehyde and 30% sucrose for 48 h at 4 °C and transferred to a −20 °C freezer thereafter. The brains were sliced coronally in 40 µm sections by means of a cryostat, and mounted in electrostatically adhesive slides (Fisherbrand^™^ Superfrost^™^ Plus slides, Fisher Scientific, Pittsburgh, PA). Sections were washed in 0.3% triton in Phosphate-Buffered Saline (PBS) for two minutes, then immersed in 10% donkey serum in PBS for 30 min for blocking.

Anti-Tyrosine hydroxylase antibody (MilliporeSigma, AB152) was diluted to 1:500 and incubated overnight at room temperature. The sections were then washed three times for five min in PBS and incubated with Alexa fluor 594 secondary antibody (Jackson Immuno research laboratories A-11012) for two hours at room temperature. The slides were washed three times for five minutes in PBS and coverslipped using Vectashileld with DAPI. Expression of the construct and restriction to TH-positive sites was confirmed using epifluorescence and confocal microscopy.

### Modeling

A Matlab (version 2019b, The Mathworks, Natick, MA) Live Script was used to simulate the output of the reward-mountain model. Models of the neural circuitry underlying eICSS and oICSS were explored by means of the simulations. The Live Script develops the mountain model from first principles, both for eICSS and oICSS. Key experiments assessing the validity of the model are reviewed, and simulated results are compared to empirical ones. The predictions of the series-circuit model of brain reward circuitry are tested and found to deviate from the empirical results of eICSS studies. An alternate model is proposed and used to simulate the results of eICSS experiments that are not readily explained by the series-circuit model. The text of the Live Script is provided in the Supporting Information along with instructions for downloading and installing the executable code.

## Supporting information

**S1 File:** supp info compressed.pdf.

- Figs S1-S6 Rate-frequency curves as a function of optical power (rats Bechr14-28)..
- Figs S7-S12 Time allocation as a function of reward strength and cost (rats Bechr14-28).
- Tab S1. Definition of acronyms and symbols
- Tab S2. The functions composing the reward-mountain model.
- Derivation of the reward-mountain model
  - Fig S13 Assumed frequency-following function for optical stimulation of midbrain dopamine neurons.
  - Fig S14 Subjective opportunity-cost (“price”) function.
  - Fig S15 Surface and contour plots of two seven-parameter reward-mountain models.
- Adaptation of the reward-mountain model for oICSS
  - Fig S16 The reward-mountain model for oICSS.
  - Parameters of the frequency-following function for oICSS.
  - Displacement of the shell: distinguishing two sources.
  - Correction of the location-parameter estimates for changes in frequency-following fidelity.
- Model fitting and selection
  - Tab S3 The 12 candidate models fit to each dataset.
  - Tab S4 Model-evaluation statistics for the fit of the 12 candidate models to the data from rat Bechr29.
  - Tab S5 Summary statistics for the model that provided the best fit (highest evidence ratio) to the data for each rat.
  - Tab S6 Best-fitting models for all rats.
- Figs S17-S22 Surfaces fit to the vehicle and drug data (rats Bechr14-28).
- Figs S23-S28 Contour graphs of the surfaces fit to the vehicle and drug data and bar graphs of the shifts in the location parameters (rats Bechr14-28).
- Location-parameter estimates
  - Tab S7 The position of the reward mountain along the pulse-frequency axis.
  - Tab S8 Estimates of the maximum normalized reward intensities in the vehicle and drug conditions.
  - Tab S9 The position of the reward mountain along the price axis.
  - Fig S29 Scatter plot of drug-induced shifts in the location parameters.
- Tabs S10-S16 Parameter values from the best-fitting models for rats Bechr14-29.
- The reward-growth function for oICSS
  - Fig S30 Changing the value of the input-scaling parameter, *F*_*hm*_, shifts the mountain along the pulse-frequency axis.
  - Fig S31 Changing the value of the output-scaling parameter, *K*_*rg*_, shifts the mountain along the price axis.
- Illustration of the correction for imperfect frequency-following fidelity
  - Fig S32 Correction for imperfect frequency-following fidelity.
- Comparison of logistic and power growth of reward intensity
  - Fig S33 The input-scaling parameter of the power reward-growth function locates the reward mountain along the price axis.
  - Fig S34 The output-scaling parameter of the power reward-growth function also locates the reward mountain along the price axis.
- Toward a new model of brain-reward circuitry
  - Fig S35 Contour graphs of reward mountains simulated by the convergence model.
  - Fig S36 Contour graphs of reward mountains simulated by the convergence model given very strong MFB input.
- Fig S37 Graphical summary of the experimental procedure.
- Contour lines: the trade-off between pulse frequency and price to hold time allocation constant (Derivation)
- Instructions for downloading and installing the executable Matlab^**®**^ Live Script.
- Text of Matlab^**®**^ Live Script.

## Acknowledgments

Steve Cabilio developed and maintained the experimental-control and data acquisition software used in this study. The experimental-control and data acquisition hardware was designed, built, and maintained by David Munro. Karl Deisseroth and Ilana B. Witten kindly provided TH::Cre^+\-^ rats to establish our breeding colony. Marie-Pierre Cossette assisted Ivan Trujillo-Pisanty in genotyping and breeding-colony maintenance. The NIMH Chemical Synthesis and Drug Supply Program supplied the GBR-12909 used in this study. We received valuable advice on microscopy from Chloë Van Oostende (Concordia Centre for Microscopy and Cellular Imaging). Elizabeth E. Steinberg provided helpful guidance concerning oICSS. Andreas Arvanitogiannis, Peter Dayan and Giovanni Hernandez offered much appreciated comments on the manuscript.

## Supporting information

### Power-frequency trade-off

**Fig S1.**
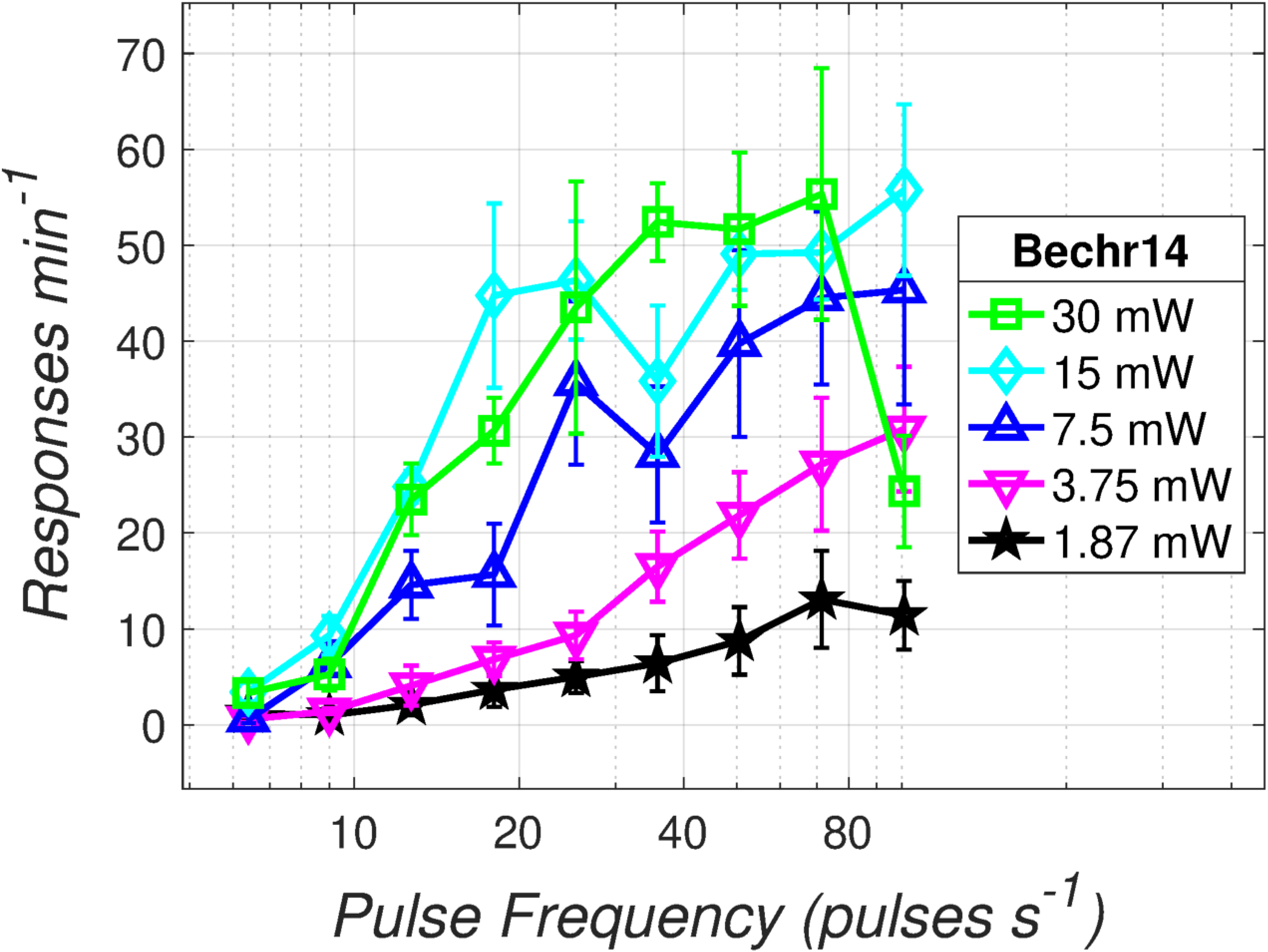
Response-rate versus pulse-frequency graph for rat Bechr14. The number of responses emitted per 2-min trial by an exemplar rat (Bechr29) is plotted as a function of pulse frequency and optical power.

**Fig S2.**
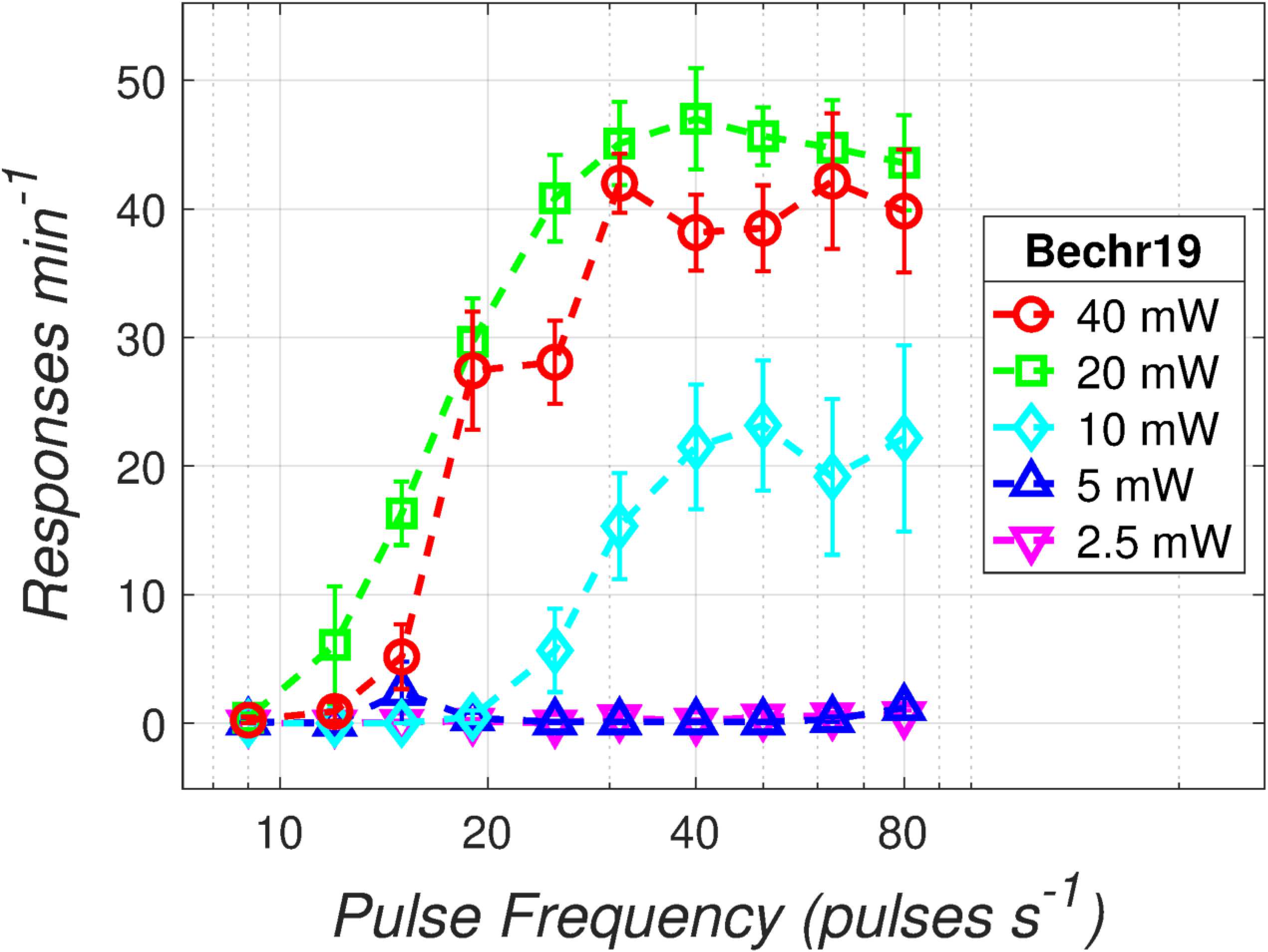
Response-rate versus pulse-frequency graph for rat Bechr19.

**Fig S3.**
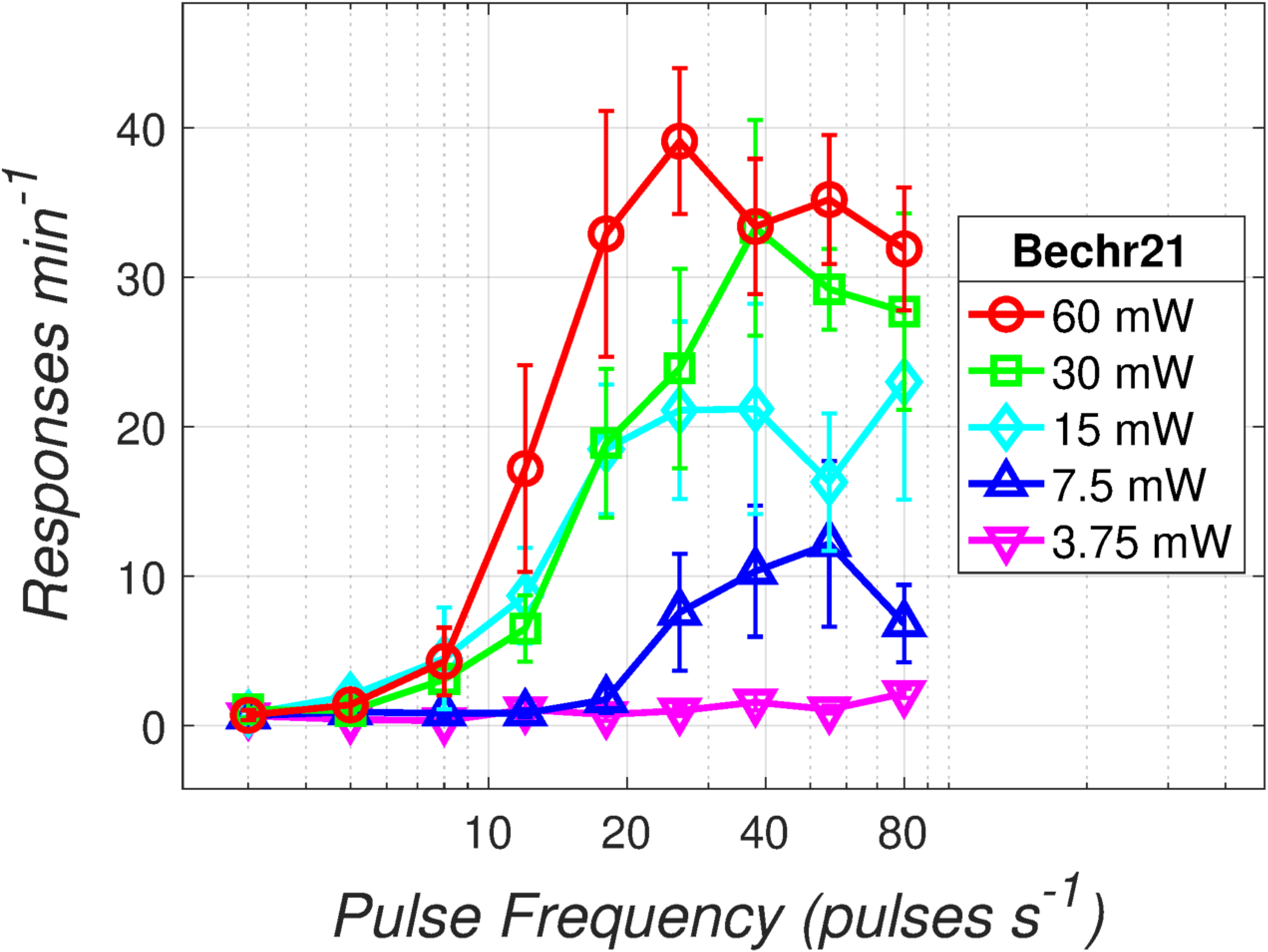
Response-rate versus pulse-frequency graph for rat Bechr21.

**Fig S4.**
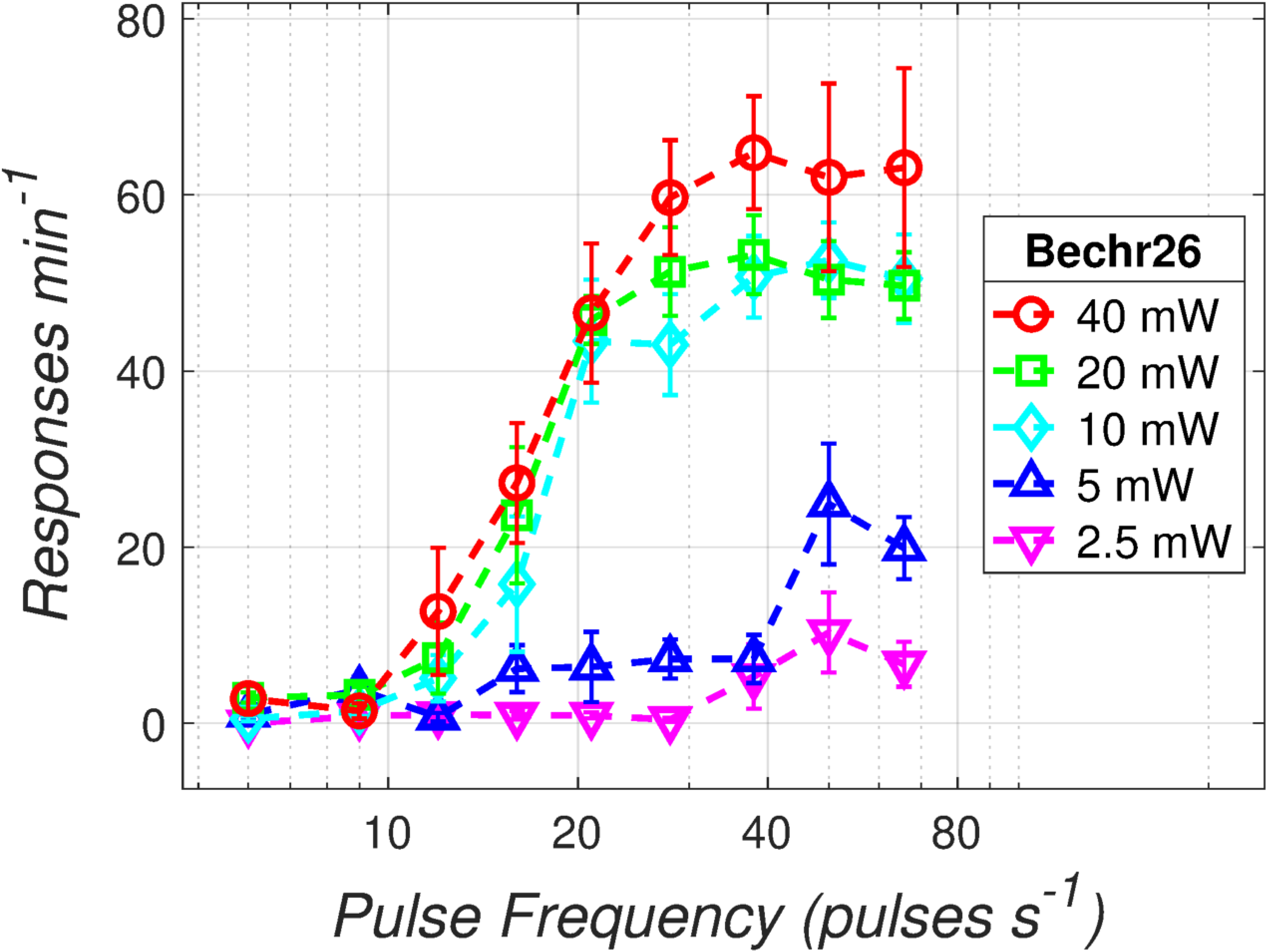
Response-rate versus pulse-frequency graph for rat Bechr26.

**Fig S5.**
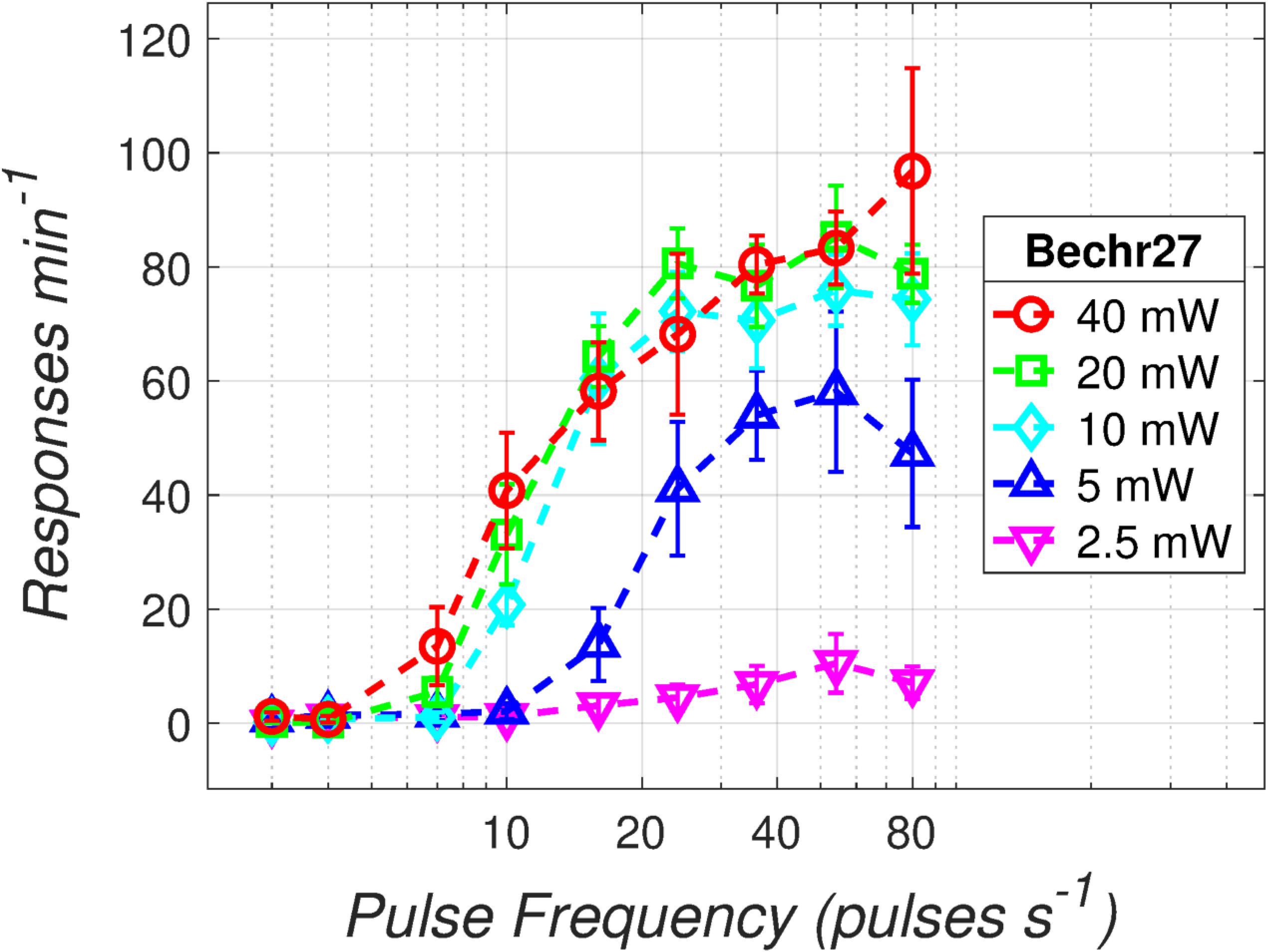
Response-rate versus pulse-frequency graph for rat Bechr27.

**Fig S6.**
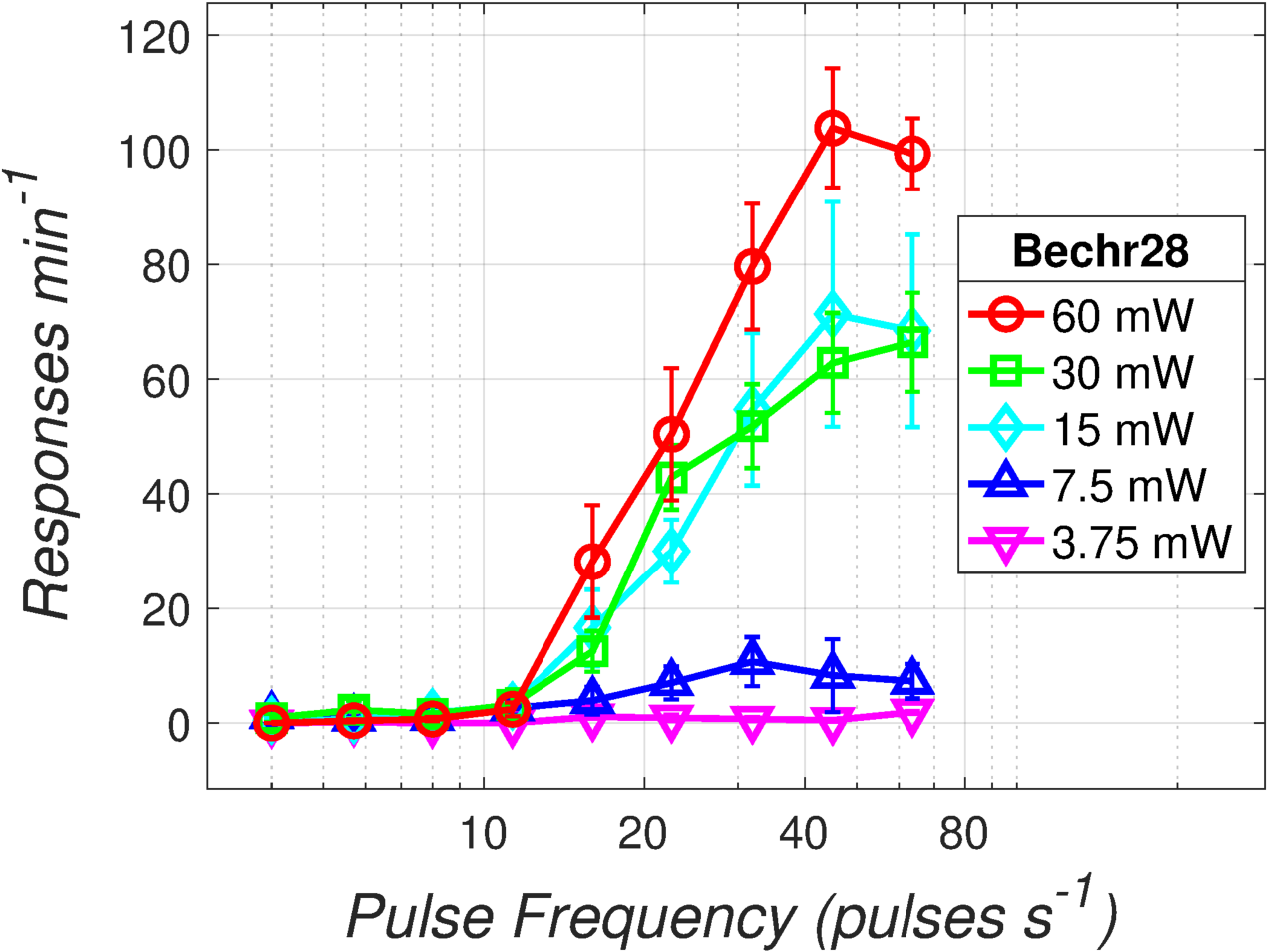
Response-rate versus pulse-frequency graph for rat Bechr28.

The data for rat Bechr29 are shown in Fig 3 in the main text.

### Time allocation as a function of reward strength and cost

**Fig S7.**
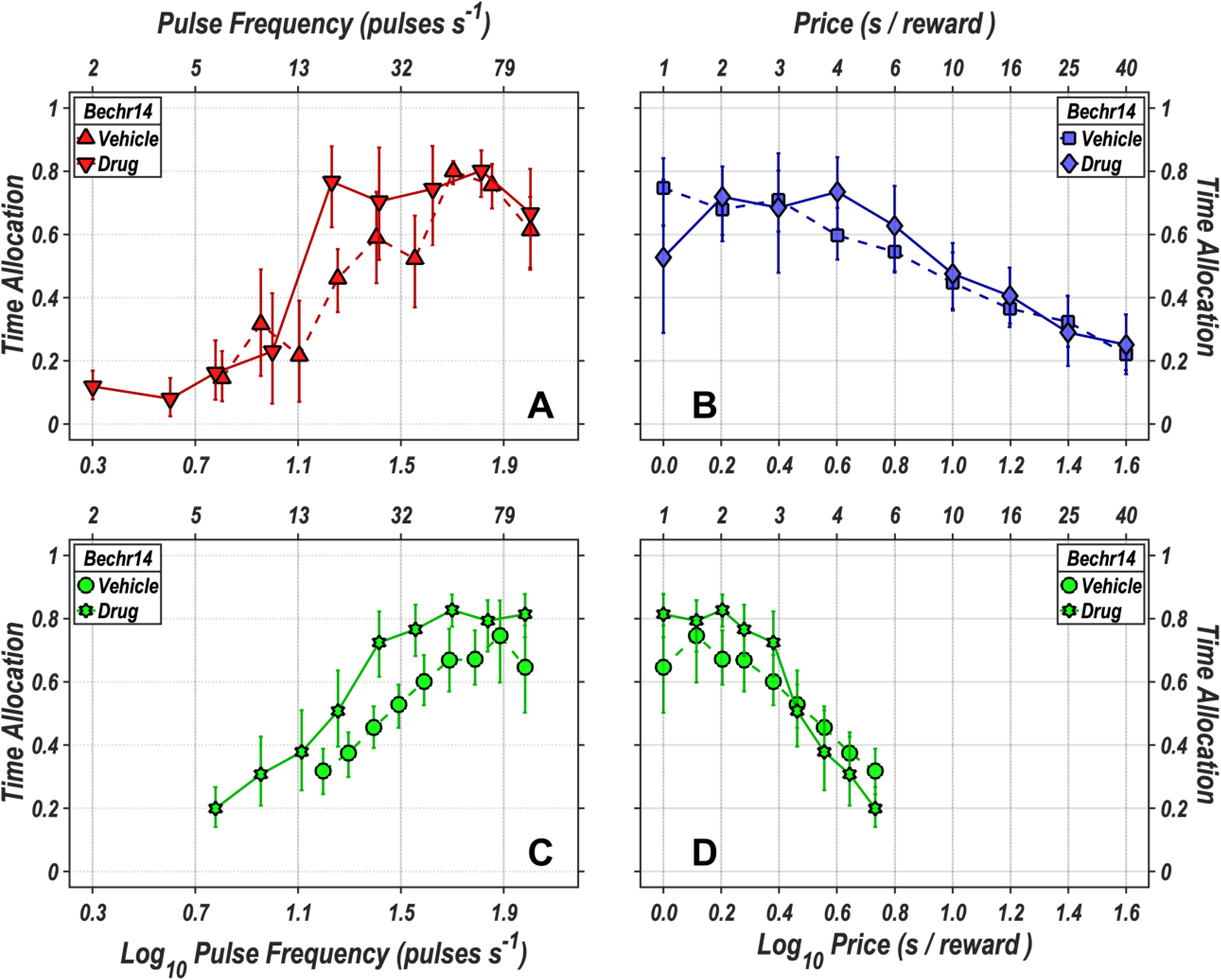
Time allocation as a function of reward strength and cost for rat Bechr14. **A:** Time allocation as a function of pulse frequency (reward strength) in the vehicle (upright triangles) and drug (inverted triangles) conditions. **B:** Time allocation as a function of price (opportunity cost) in the vehicle (squares) and drug (diamonds) conditions. In the radial-sweep condition, the pulse frequency was decreased and the price decreased concurrently, in stepwise fashion, over consecutive trials. Time allocation is plotted as a function of pulse frequency in panel **C**: and as a function of price in panel **D:**. Data from the vehicle condition are represented by circles, whereas data from the drug condition are represented by Stars of David. The error bars represent 95% confidence intervals.

**Fig S8.**
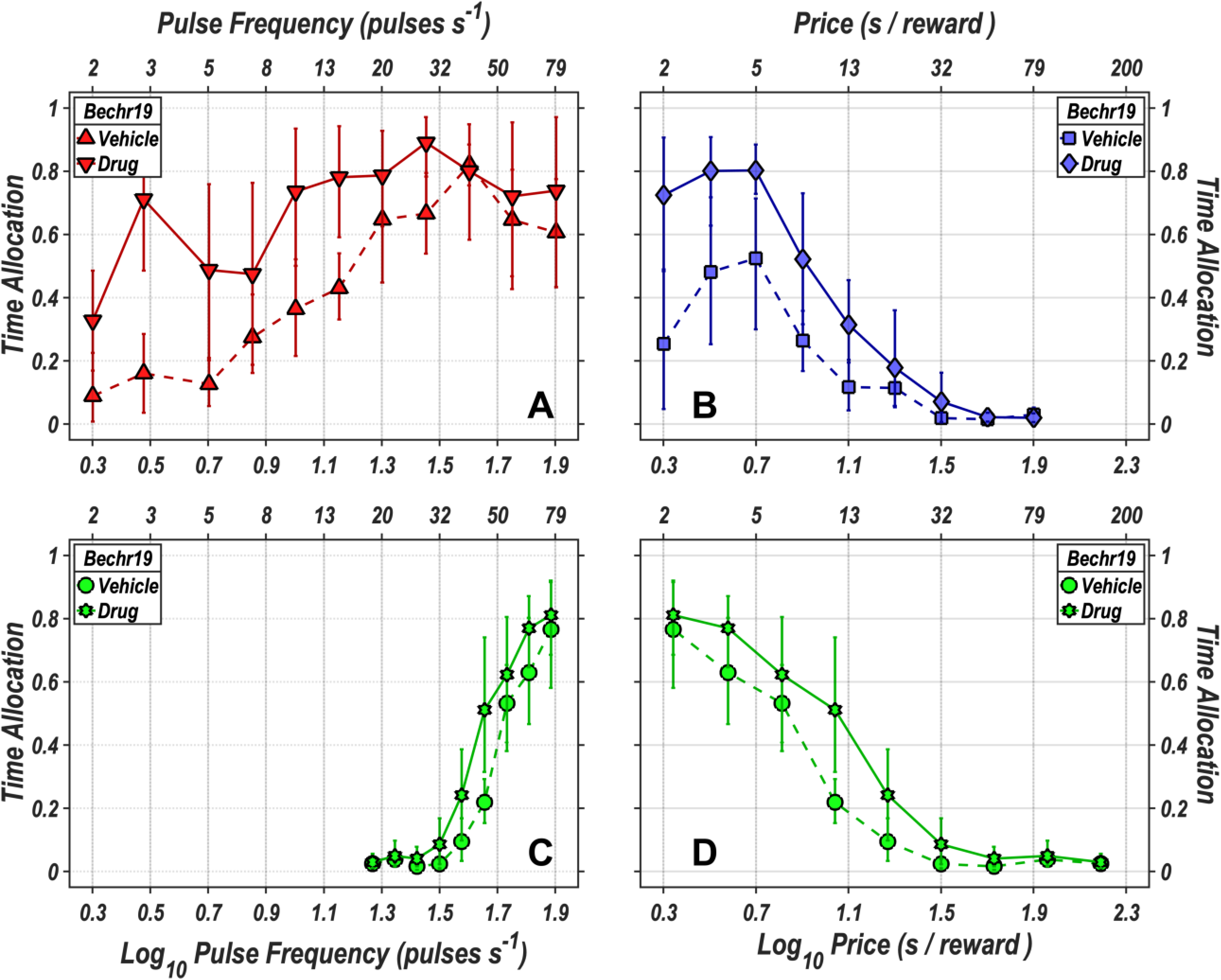
Time allocation as a function of reward strength and cost for rat Bechr19.

**Fig S9.**
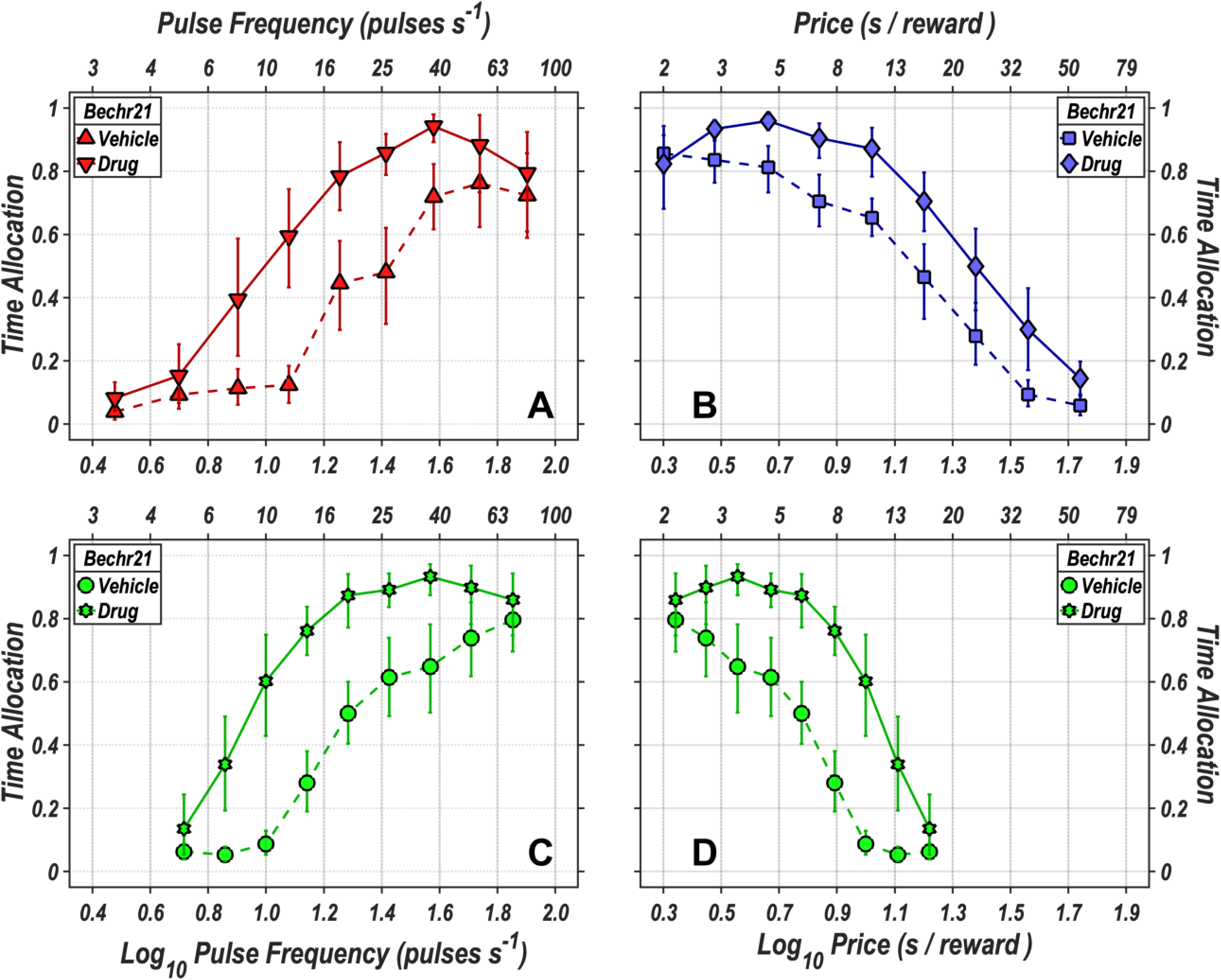
Time allocation as a function of reward strength and cost for rat Bechr21.

**Fig S10.**
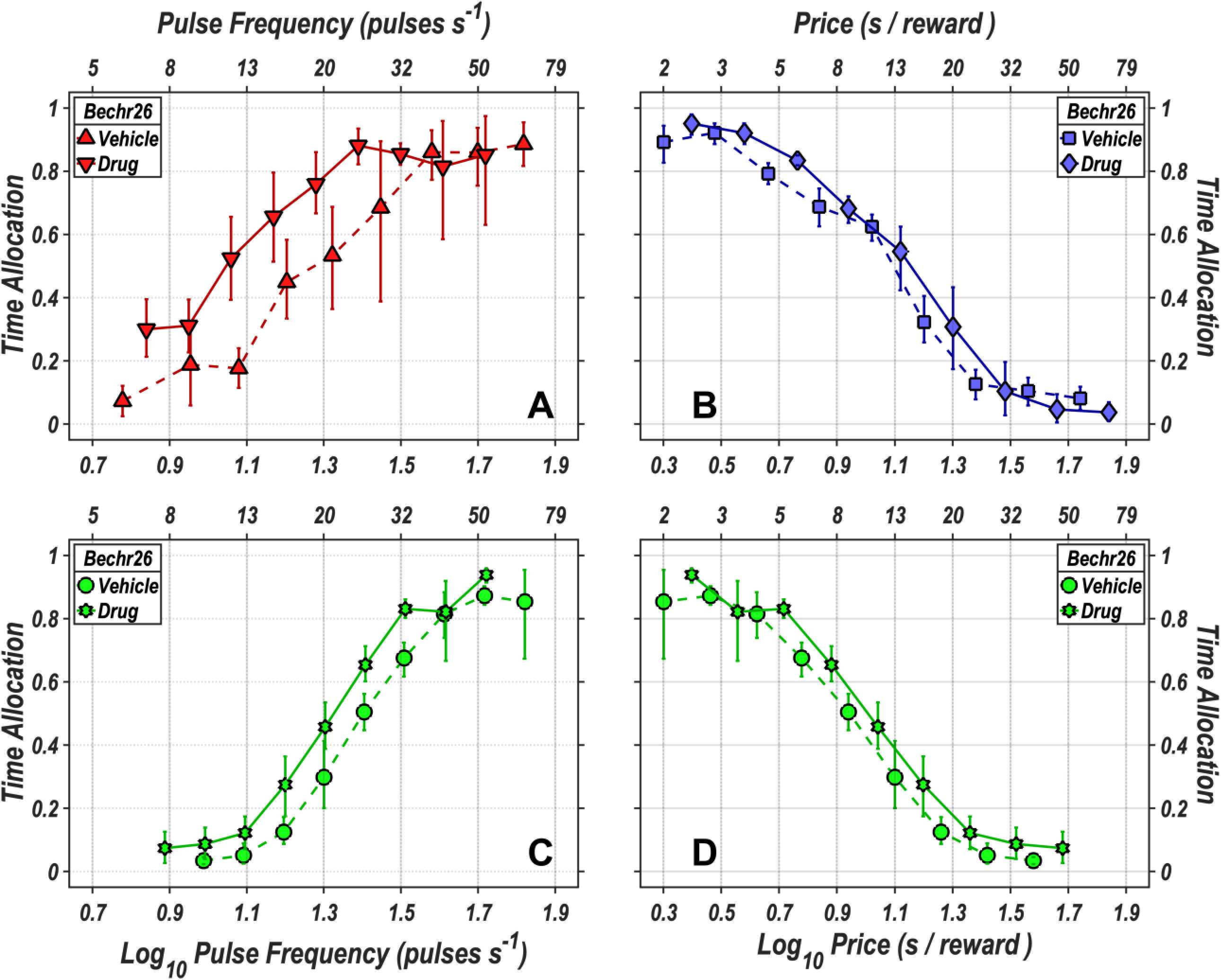
Time allocation as a function of reward strength and cost for rat Bechr26.

**Fig S11.**
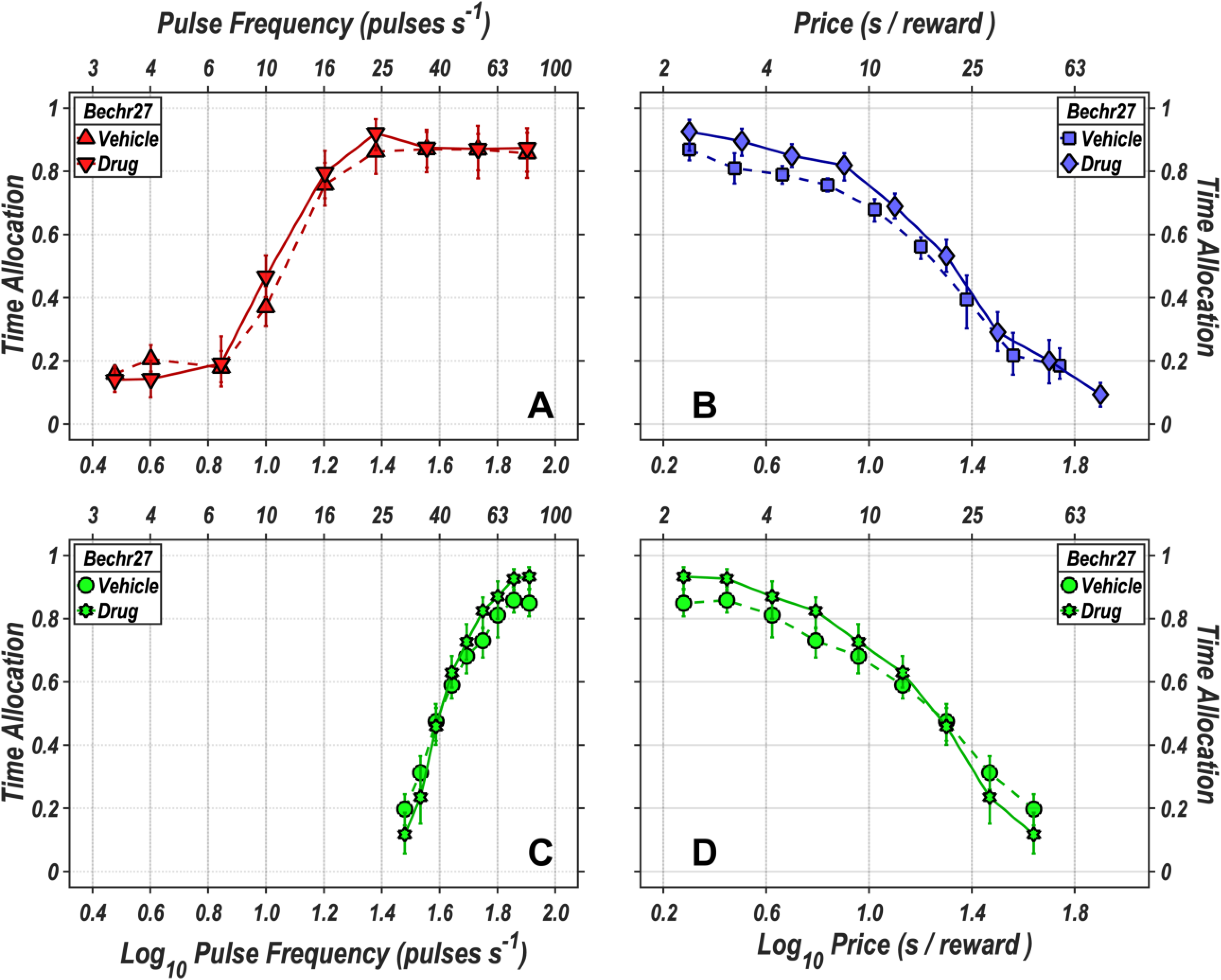
Time allocation as a function of reward strength and cost for rat Bechr27.

**Fig S12.**
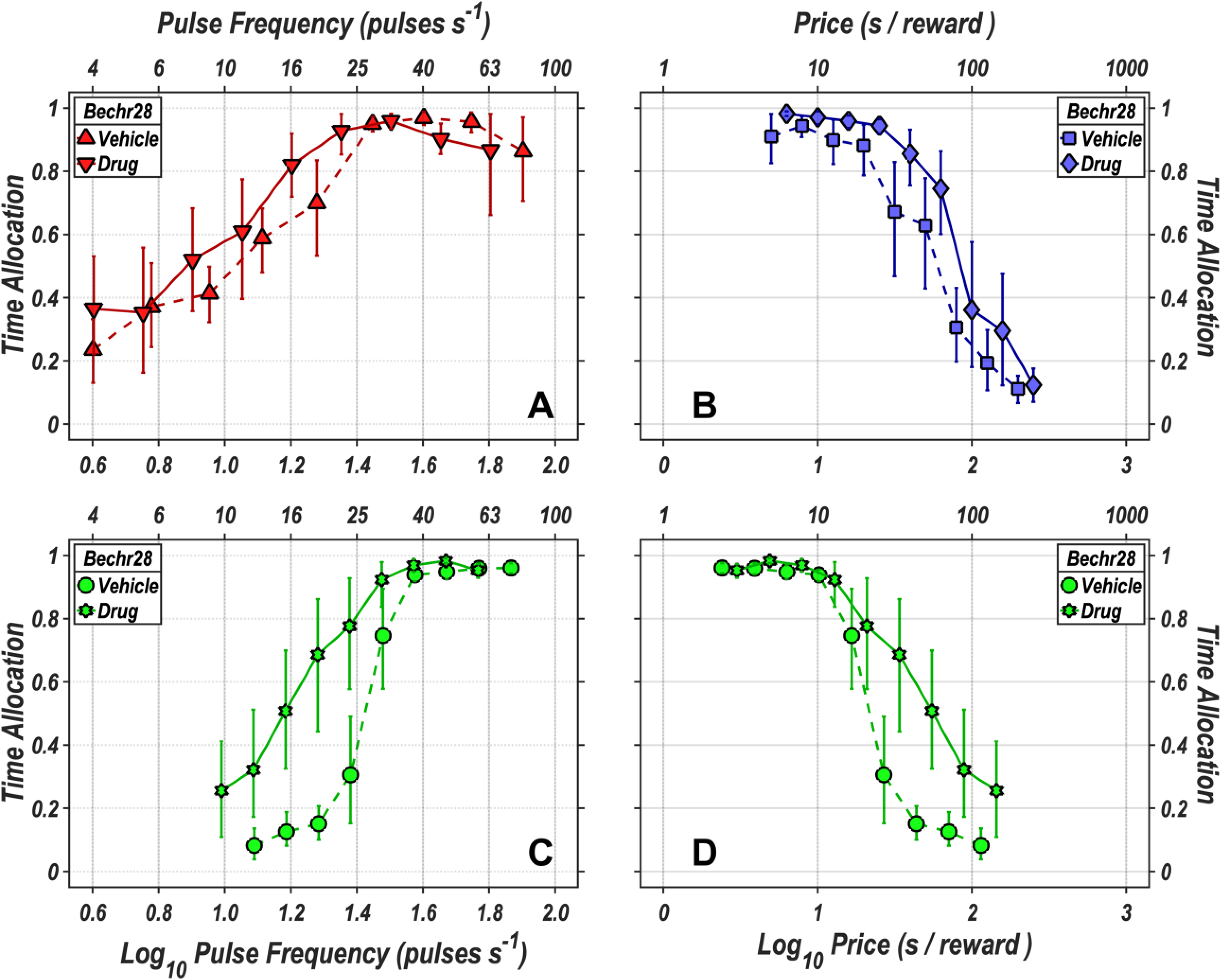
Time allocation as a function of reward strength and cost for rat Bechr28.

The data for rat Bechr29 are shown in Fig 4 in the main text.

### Derivation of the reward-mountain model

Acronyms and symbols employed in this supporting-information file are defined in Tab S1.

**Table S1.** Definition of acronyms and symbols.

| Acronym<br>or Symbol | Definition |
| --- | --- |
| $a$ | price-sensitivity exponent |
| BSR | brain stimulation reward |
| $c$ | chronaxie of the strength-duration function for pulses (pulse duration at which the threshold current is twice rheobase) |
| $C$ | chronaxie of the strength-duration function for trains (train duration at which $F_{firing_{hm}}$ is twice $\rho_{\Pi}$) |
| $C_r$ | Conditioned reward value |
| ChR2 | channelrhodopsin-2 |
| $d$ | pulse duration |
| $D_{burst}$ | duration of stimulation-induced burst of firing in the directly activated neurons |
| $D_{train}$ | train duration (leading edge of first pulse to leading edge of final pulse) |
| DAPI | 4',6-diamidino-2-phenylindole |
| eICSS | electrical intracranial self-stimulation |
| $f_a$ | function that determines the average reward rate derived from performance of alternate activities |
| $f_D$ | function that relates the duration of the stimulation-induced burst of firing to the train duration |
| $f_F$ | frequency-following function that relates the induced firing frequency to the pulse frequency |
| $f_N$ | function that translates the pulse duration and current into the number of electrically recruited, directly stimulated neurons |
| $f_p$ | subjective-probability function |
| $f_P$ | subjective-price function |
| $f_R$ | reward-growth function for eICSS or oICSS |
| $f_T$ | behavioral-allocation function |
| $f_U$ | function that translates the reward rate and subjective rate of exertion into a payoff |
| $f_{\phi}$ | subjective-effort function |
| $F_{bend}$ | parameter governing the abruptness of the roll-off in frequency following |
| $F_{firing}$ | firing frequency induced by electrical or optical stimulation |
| $F_{firing_{hm}}$ | firing frequency that generates a reward of half-maximal intensity |
| $F_{firing_{max}}$ | maximum firing frequency induced by electrical or optical stimulation |
| $F_{hm}$ | notation for $F_{pulse_{hm}}$ used in previous eICSS papers |
| $F_{hm}^*$ | shorthand notation for $F_{pulse_{hm}}^*$ |

Table S1. Definition of acronyms and symbols
| Acronym<br>or Symbol | Definition |
| --- | --- |
| $F_{pulse}$ | pulse frequency in an electrical or optical pulse train |
| $F_{pulse_{hm}}$ | pulse frequency required to drive reward intensity to half its maximum value |
| $F_{pulse_{hm}}^*$ | estimated pulse frequency required to drive reward intensity to half its maximum value if frequency-following fidelity were perfect |
| $F_{ro}$ | pulse frequency in the center of the roll-off region of the frequency-following function |
| $g$ | exponent governing the steepness of reward-intensity growth |
| ICSS | intracranial self-stimulation |
| $I$ | eICSS current |
| $K_{aa}$ | constant scaling the value of alternate activities |
| $K_{da}$ | constant representing the scaling of dopamine release |
| $K_{ec_L}$ | effort-cost scalar for leisure activities |
| $K_{ec_W}$ | effort-cost scalar for work |
| $K_F$ | unit-translation constant to convert pulses $s^{-1}$ into firings $s^{-1}$ neuron$^{-1}$ |
| $K_{IS}$ | current-distance constant |
| $K_{NS}$ | neuron-distance constant |
| $K_{rg}$ | reward-intensity scalar |
| $N$ | number of directly activated neurons |
| MFB | medial forebrain bundle |
| oICSS | optical intracranial self-stimulation |
| $p_{obj}$ | objective probability that a reward will be delivered upon satisfaction of the response requirement |
| $p_{sub}$ | subjective probability that a reward will be delivered upon satisfaction of the response requirement |
| $P_e$ | notation for $P_{obj_e}$ used in previous eICSS papers |
| $P_e^*$ | shorthand notation for $P_{obj_e}^*$ |
| $P_{obj}$ | objective opportunity cost (“price”) of a stimulation train |
| $P_{obj_e}$ | objective price at which $\hat{T} = 0.5$ when $\hat{R} = \hat{R}_{max}$ |
| $P_{obj_e}^*$ | estimated objective price at which $\hat{T} = 0.5$ and $\hat{R} = \hat{R}_{max}$ if frequency-following fidelity were perfect |
| $P_{sub}$ | subjective opportunity cost (“price”) of a stimulation train |
| $\vec{P}_{sub_{\hat{T}=0.5}}$ | vector of subjective prices that hold time allocation midway between its minimum and maximum values |
| $P_{sub_{bend}}$ | parameter controlling the abruptness of the transition from the “blade” to the “handle” of the subjective-price function |
| $P_{sub_{min}}$ | minimal subjective price |
| $\dot{R}_{aa}$ | average rate of reward from performance of alternate (leisure) activities |
| $\hat{\dot{R}}_{aa}$ | $\dot{R}_{aa}$ normalized to vary between 0 and 1 |
| $R_{bsr}$ | peak reward intensity achieved over the course of a stimulation train |
| $\hat{R}_{bsr}$ | $R_{bsr}$ normalized to vary between 0 and 1 as $R_{bsr}$ rises from 0 to $R_{bsr_{max}}$ |
| $\hat{R}_{bsr_{max}}$ | maximum normalized reward intensity |
| $\vec{\hat{R}}_{bsr_{\hat{T}=0.5}}$ | vector of normalized reward intensities that hold time allocation midway between its minimum and maximum values |
| $\dot{R}_{bsr}$ | rate of brain-stimulation reward; reward intensity divided by the subjective price paid to procure the pulse train |
| $T$ | time allocation |

**Table S1. Definition of acronyms and symbols**
| Acronym<br>or Symbol | Definition |
| --- | --- |
| $\hat{T}$ | time allocation normalized to rise between 0 and 1 as $T$ rises from $T_{min}$ to $T_{max}$ |
| $T_{max}$ | maximal time allocation |
| $T_{mid}$ | value of $T$ midway between $T_{min}$ and $T_{max}$; $T_{mid} = \hat{T}_{0.5}$ |
| $T_{min}$ | minimal time allocation |
| TH | tyrosine hydroxylase |
| $U_L$ | payoff from alternate (“leisure”) activities (a.k.a “everything else”) |
| $U_W$ | payoff from brain stimulation reward |
| $\hat{U}_W$ | $U_W$ normalized to vary between 0 and 1 |
| YFP | (enhanced) yellow fluorescent protein |
| $\rho_I$ | rheobase of the strength-duration function for pulses |
| $\rho_{\Pi}$ | rheobase of the strength-duration function for trains |
| $\dot{\phi}_{obj_L}$ | average work rate entailed in performing leisure activities |
| $\dot{\phi}_{sub_L}$ | average rate of subjective exertion entailed in performing leisure activities |
| $\hat{\dot{\phi}}_{sub_L}$ | $\dot{\phi}_{sub_L}$ normalized to vary between 0 and 1 |
| $\dot{\phi}_{obj_W}$ | work rate entailed in holding down the lever |
| $\dot{\phi}_{sub_W}$ | subjective rate of exertion entailed in holding down the lever |

The reward-mountain model provides a framework for integrating the frequency-sweep, pulse-sweep, and radial-sweep data (e.g., Figure 4) in a unified 3D space and for interpreting the drug-induced changes in the position of the resulting 3D structure (the reward mountain). The following derivation of the model first extends earlier depictions that were developed in the context of eICSS studies [1–3] and then adapts the model to accommodate oICSS of midbrain dopamine neurons.

The reward-mountain model predicts time allocation given experimenter-controlled variables that determine the strength and cost of the rewarding stimulation. In most experiments carried out to date, the strength variable is the pulse frequency within a fixed-duration stimulation train, and the cost variable is the work time (opportunity cost, price) required to procure a stimulation train. The functional machinery that generates time-allocation values from these inputs is summarized in Tab S2.

**Table S2.** The functions composing the reward-mountain model. The shell → core functions (left column) map the values of variables that are manipulated or controlled into inputs to the core of the model. The core functions (middle column) map these inputs into the payoffs from work and leisure activities, whereas the core → shell function map the payoffs into the observed dependent variable: time allocation. The accompanying Matlab^®^ Live Script illustrates how the listed functions are implemented. Please see Tab S1 for definitions of the symbols.

| shell → core | core | core → shell |
| --- | --- | --- |
| $F_{firing} = f_F(F_{pulse})$ | | |
| $N = f_N(I, d)$ | | |
| $D_{burst} = f_D(D_{train})$ | $F_{firing_{hm}} = f_H(D_{burst}, N)$ | |
| | $R_{bsr} = f_R(F_{firing}, F_{firing_{hm}})$ | |
| $p_{sub} = f_p(p_{obj})$ | | |
| $P_{sub} = f_P(P_{obj})$ | | |
| $\dot{\phi}_{sub_W} = f_\phi(\dot{\phi}_{obj_W})$ | $U_W = f_U\left(\frac{R_{bsr}}{P_{sub}}, p_{sub}, \dot{\phi}_{sub_W}\right)$ | |
| $\_ = f_a(\_, \_, \dots \_)$ | $\dot{R}_{aa} = f_R(\_)$ | |
| $\dot{\phi}_{sub_L} = f_\phi(\dot{\phi}_{obj_L})$ | $U_L = f_U(\dot{R}_{aa}, \dot{\phi}_{sub_L})$ | $T = f_T(U_W, U_L)$ |

As the table implies, we distinguish between the “shell” and “core” of the model. The shell consists of the variables that are observed (time allocation), manipulated (pulse frequency, price), and controlled (stimulation parameters held constant, physical work required to hold down the lever, affordances of the test environment [4]). The shell is displayed within the space defined by the observed and manipulated variables.

The core (middle column of Tab S2) consists of the functions that compute the intensity of the reward produced by the stimulation train and combine this value with the opportunity and effort costs to generate what we call “payoffs.” Parallel functions in the core compute the value of the alternate activities that compete with pursuit of the stimulation for the rat’s behavior. A set of functions (left column) provides the input to the core by mapping the manipulated and controlled variables into the quantities from which payoffs are derived.

A single function (right column), based on the generalized matching law [5], translates the payoffs generated in the core into the time-allocation values that are manifested in the shell.

Shell variables are objective, whereas core variables are inferred subjective quantities. The core functions are the bridge between the objective inputs that are manipulated or controlled to the observed objective output, time allocation; their form and parameters explain why the manipulated variables cause time allocation to vary in the manner observed in the experiment.

#### Shell → core functions

The arguments of the first three shell → core functions are the four parameters that define a fixed-frequency pulse train: the pulse frequency (*F*_*pulse*_), current (*I*), pulse duration (*d*), and train duration (*D*_*train*_). These functions relay to the core functions the stimulation-induced frequency of firing (*F*_*firing*_), the number of activated neurons (*N*), and the duration of the burst of increased firing (*D*_*burst*_).

##### f_F_

In studies of eICSS, the experimenter typically varies the pulse frequency during a fixed-duration stimulation train in order to control the intensity of the electrically-induced reward. The frequency-following function labeled ***f***_***F***_ maps the manipulated shell variable, *F*_*pulse*_ into the corresponding core variable: the induced frequency of firing in the directly-stimulated substrate, *F*_*firing*_. In the case of eICSS of the MFB, this function has been estimated by psychophysical means and shown to be roughly scalar up to very high pulse frequencies [6], well beyond the typical range of the 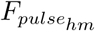 values that locate the reward-mountain shell in the space defined by the two independent variables. Solomon et al. [6] showed that the following function provides a good fit to the frequency-following data for eICSS of the MFB:

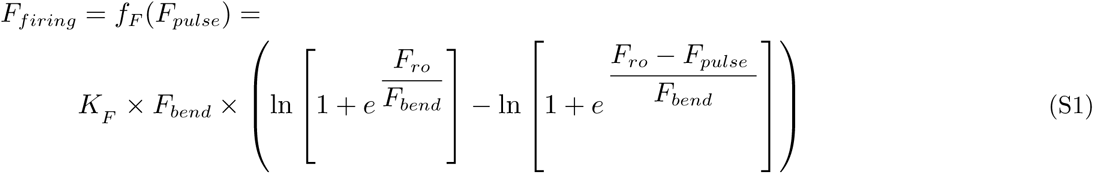

where

***f*_*F*_** = the frequency-following function

*F*_*bend*_ = parameter determining the abruptness of the roll-off in the frequency response; units: *unitless*

*F*_*firing*_ = induced firing rate in the first-stage neurons; units: *firings s* ^− 1^ *neuron* ^− 1^

*F*_*pulse*_ = the pulse frequency; units: *pulses s*^− 1^

*F*_*ro*_ = the pulse frequency in the center of the roll-off region; units: *pulses s*^− 1^

*K*_*F* =_ unit-translation constant; units: *firings pulse*^−1^ *neuron*^− 1^

A plot of the frequency-following function is shown in Fig S13. The form of the function is the same as the one described by Solomon et al. [6]. The choice of parameters is described below in section Parameters of the frequency-following function for oICSS.

**Fig S13.**
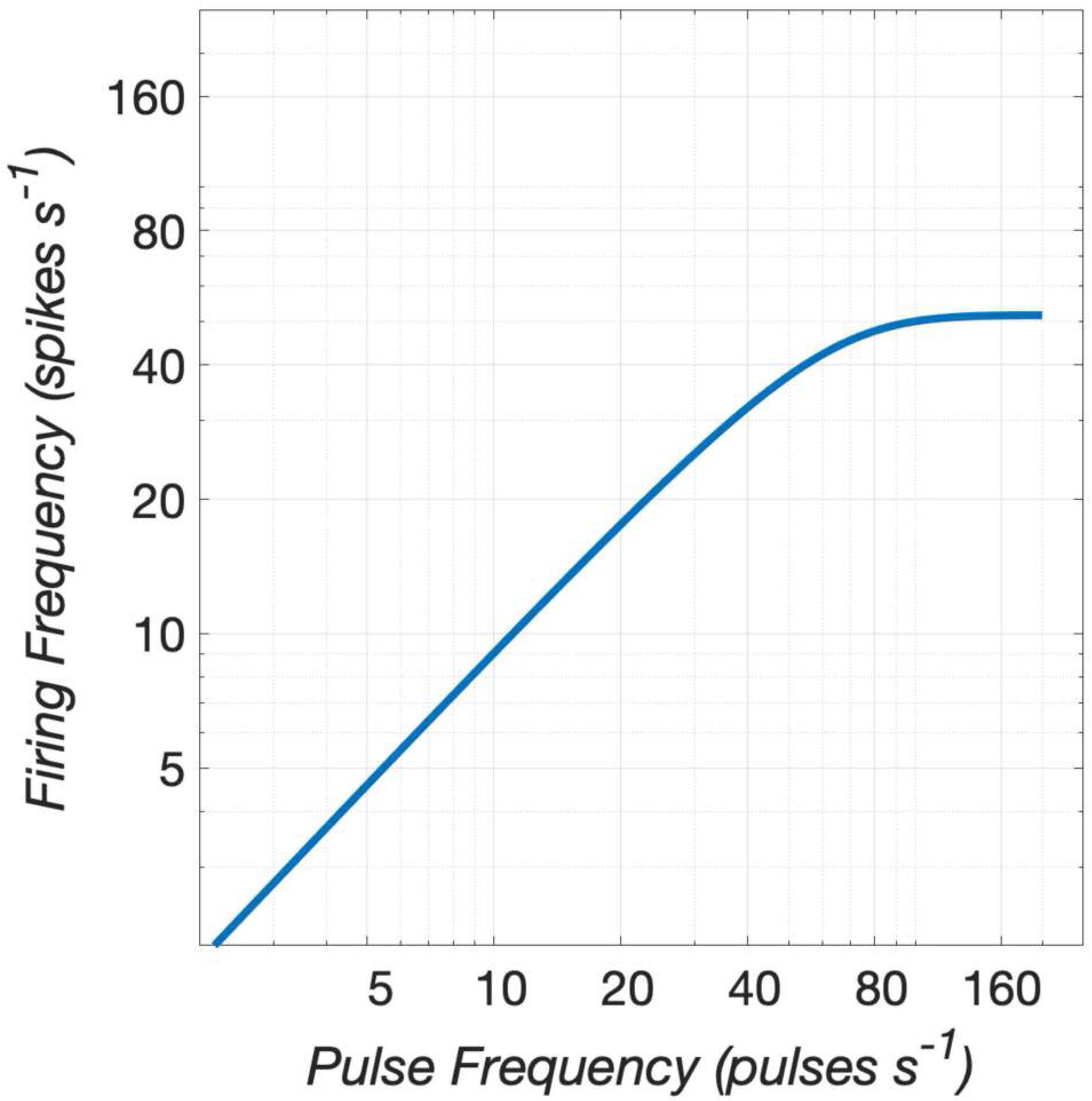
Assumed frequency-following function for optical stimulation of midbrain dopamine neurons. The induced firing frequency is plotted as a function of the optical pulse frequency. Note that frequency-following fidelity is increasingly poor as the pulse frequency increases. At low values, the firing frequency falls only slightly short of the pulse frequency, but by 40 pulses s^−1^, the firing frequency is only ∼80% of the pulse frequency. The maximum induced firing frequency is 51.6 spikes s^−1^

##### f_N_

The *F*_*N*_ function translates the pulse duration and current into the number of electrically excited neurons. These two variables determine conjointly the boundary of the region in which the stimulation excites reward-related neurons. Holding these variables constant, as is the case in most eICSS studies entailing measurement of the reward mountain and in prior studies employing the curve-shift method [7–9], circumvents the need to make assumptions about the spatial distribution of the directly-stimulated neurons subserving the rewarding effect and about their excitability to extracellular stimulation.

Hawkins (cited in [10]) proposed that the number of directly-stimulated (“first-stage”) neurons subserving the rewarding effect is roughly proportional to the current, when pulse duration is held constant. Emprical studies [11, 12] show that the current required to excite a given first-stage neuron varies roughly as a rectangular, hyperbolic function of the pulse duration. Thus,

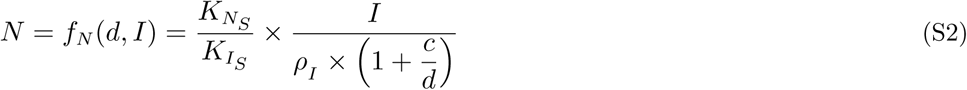

where
*c* = chronxie; pulse duration (units: ms) for which the threshold current is twice *ρ*_*I*_

*d* = pulse duration (units: ms)

*f*_*N*_ = the first-stage recruitment function

*I* = current (units: *μ*A)

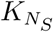 = neuron-distance constant; units: *neurons mm*^−2^

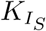 = current-distance constant; units: *μ*A *mm*^−2^

*N* = number of activated first-stage neurons; units: *neurons*

*ρ*_*I*_ = threshold current required to excite a first-stage neuron using a pulse of infinite duration; units: *μ*A

##### f_D_

The third stimulation parameter that has been held constant in most reward-mountain and curve-shift studies is the duration of the stimulation train. The *f*_*D*_ function translates the duration set by the experimenter into the duration of the stimulation-induced increase in the activity of the directly-stimulated substrate. If firing is time-locked to the stimulation pulses, then the duration of this increase, *D*_*burst*_, will equal the train duration, *D*_*train*_. Thus, we assume that

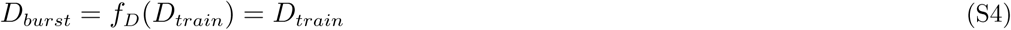

where

*D*_*burst*_ = duration of the stimulation-induced increase in firing above baseline in the directly stimulated neurons subserving the reward effect

*D*_*train*_ = time interval between the leading edges of the first and last pulse in a stimulation train

*f*_*D*_ = the duration-mapping function

##### f_p_

Psychophysical methods have been used to describe the subjective-probability function for BSR, *f*_*p*_ (lower-case subscript) [13]. This function returns the subjective probability that a reward will be delivered upon payment of the price set by the experimenter. Over the range, 0.5 - 1.0, this function was determined to be roughly scalar. In the present study and in all other studies that have entailed measurement of the reward mountain, delivery of the reward upon satisfaction of the response requirement is certain (*p*_*obj*_ = 1).

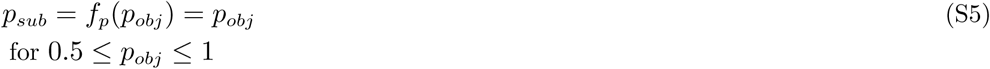

where

*f*_*p*_ = the subjective-probability function

*p*_*obj*_ = objective probability that reward will be delivered upon satisfaction of the response requirement

*p*_*sub*_ = subjective probability that reward will be delivered upon satisfaction of the response requirement

##### f_P_

Past measurements of the reward-mountain in eICSS studies employed the work time required to trigger delivery of a stimulation train (the price) as the cost variable, and that practice is continued here. The function labeled *f*_*P*_ (upper-case subscript) maps the required work time, *P*_*obj*_, into the corresponding subjective variable, *P*_*sub*_. The form and parameters of this function for eICSS of the MFB have been estimated by psychophysical means [14]. We showed that the following psychophysical function more accurately describes the opportunity cost of rewarding brain stimulation than the identity function or functions based either on hyperbolic or exponential discounting:

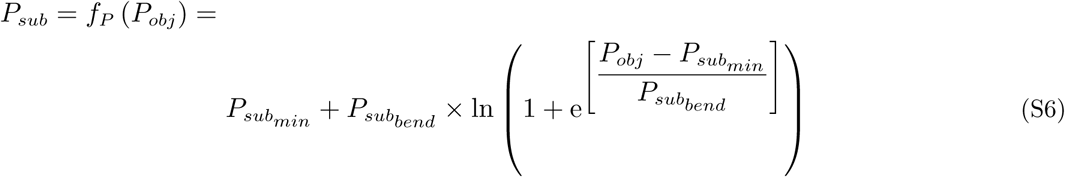

where

*f*_*P*_ = the subjective-price function

*P*_*obj*_ = the “price” of a stimulation train: the cumulative time the lever must be depressed to trigger reward delivery; units: *s*

*P*_*sub*_ = the subjective price of a stimulation train; units: *s*

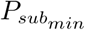 = the minimum subjective price; units: *s*

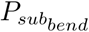 = a constant that controls the abruptness of the transition. from “blade” to “handle;” unitless

For a different view of the subjective-price function, please see [15].

At higher prices, the output of the subjective-price function defined by Eq S6 converges on its input: the subjective price becomes indistinguishable from the objective one. However, as the objective price is reduced below ∼3 s, the subjective price deviates from the objective price and eventually approaches an asymptotic value: 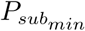. This asymptote has been interpreted to arise from the reduction and eventual disappearance of competition between lever depression and competing activities, such as grooming, resting, and exploring; performance of these competing activities is no longer perceived as beneficial once the available time for their execution becomes sufficiently short.

A plot of the subjective-price function is shown in Fig S14. The parameters employed are the mean values determined by Solomon et al. [14].

**Fig S14.**
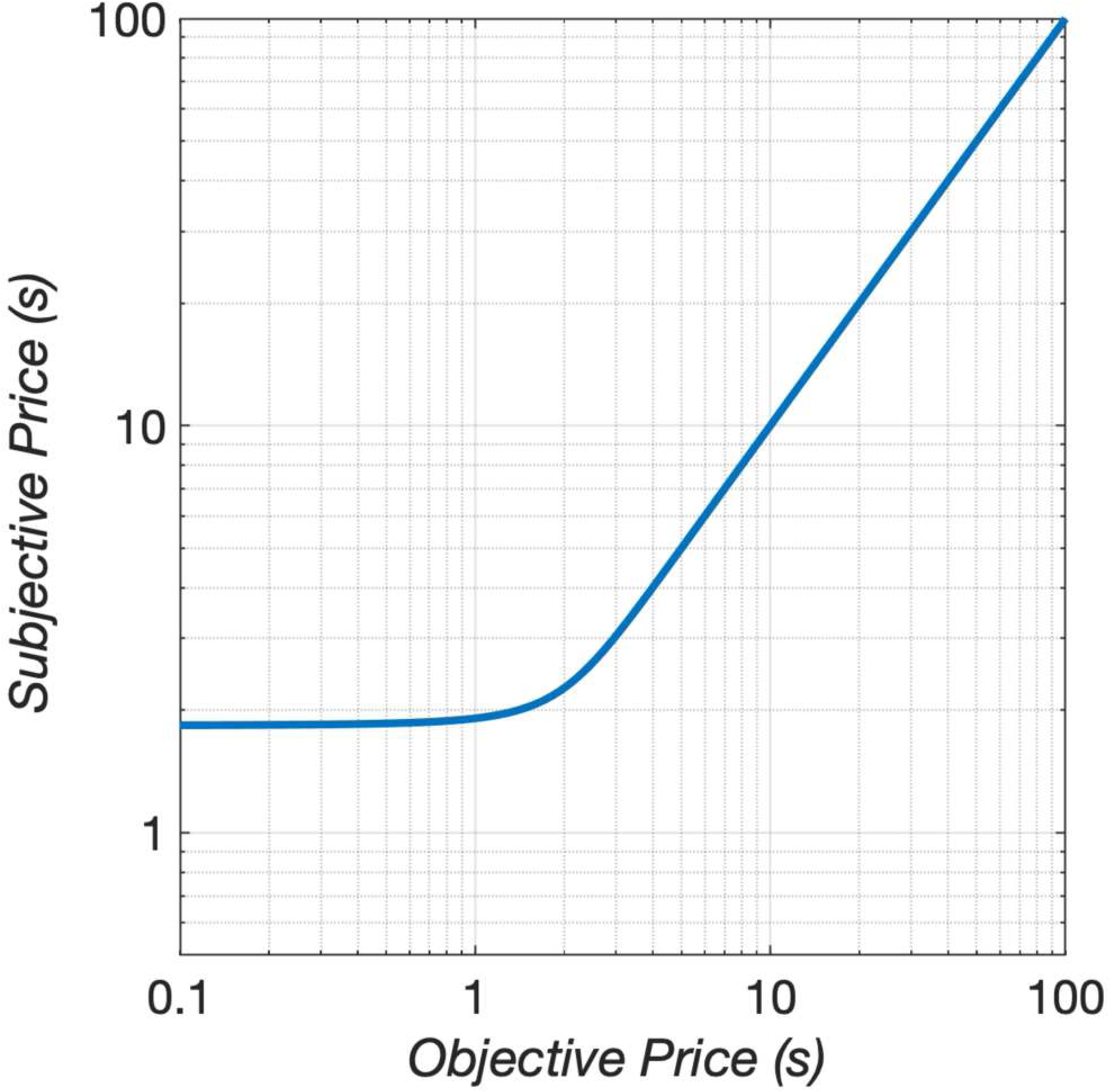
Subjective opportunity-cost (“price”) function. The function maps the objective opportunity cost (cumulative lever-depression time required to trigger reward delivery), *P*_*obj*_),into its subjective equivalent *P*_*sub*_. The form and parameters of this function are based on measurements by Solomon et al. [6].

The arguments of the first five shell → core functions are all variables controlled directly by the experimenter: {*F*_*pulse*_, *d, I, D, p, P*}. These six variables are transformed by the first five shell → core functions into inputs to the core. The arguments of the remaining two shell → core functions listed in Tab S2 are variables arising from features of the test environment that the experimenter attempts to hold constant: the rate of physical work entailed in holding down the lever or in performing activities that compete with pursuit of the rewarding stimulation.

##### f_ϕ_

The effort cost of the reward is the subjective rate of exertion entailed in holding down the lever. We know of no psychophysical studies that reveal the form of this function. That said, we can assume that it accelerates steeply as the effort cost approaches the physical capabilities of the rat. The objective effort cost has been held constant in this and in prior studies carried out in the reward-mountain paradigm. We assume that the subjective effort cost did not co-vary with the manipulated variables. To help ensure this in the current study, the lever was withdrawn for 2 s following the triggering of a stimulation train so as to provide time for any interfering motoric consequences of the stimulation to dissipate.

We treat the subjective rate of exertion required to depress the lever as a constant defined by:

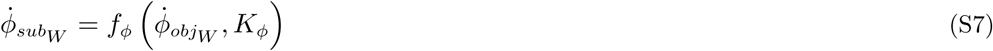

where

*f*_*ϕ*_ = subjective-effort function (form unknown)

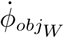 = rate of physical work required to hold down the lever; units: *Js*^−1^

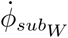 = subjective rate of exertion required to hold down the lever in units we call “*oomphs*” *s*^−1^

*K*_*ϕ*_ = unit-conversion constant; units: *oomphs J*^−1^

The dots over 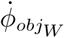 and 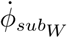 signify that we define these quantities as rates. (No dots are placed over *F*_*firing*_ and *F*_*pulse*_ because doing so would be superfluous and potentially misleading. Frequencies are inherently rates over time (the first time derivative of the pulse or spike number). By omitting the dots, we wish to avoid confusion between these rates and their changes over time (the second derivative of the pulse or spike number).)

The effect of the drug, if any, on the rate of subjective exertion is defined as:

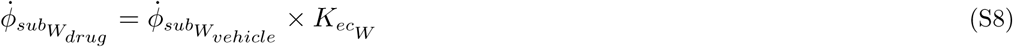

where

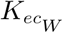 = proportional drug-induced change in the subjective rate of exertion required to hold down the lever; unitless

In the vehicle condition, 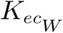 assumes an implicit value of one.

Activities such as grooming and exploring also entail performance of physical work. Thus, *f*_*ϕ*_ is also applied to these activities:

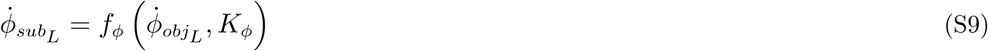

where

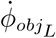 = Average rate of physical work required to perform alternate (“leisure”) activities; units: *Js*^−1^.

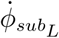 = average subjective rate of exertion entailed in performance of leisure activities in *oomphs s*^−1^.

*K*_*ϕ*_ = unit-conversion constant; units: *oomphs J*^−1^

We assume that 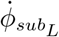 does not covary systematically with the independent variables.

As in the case of the subjective rate of exertion entailed in work, we allow for drug-induced modulation of the subjective rate of exertion entailed in performance of leisure activities.

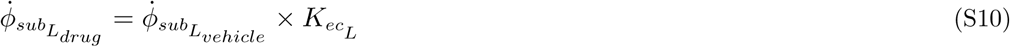

where

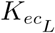 = proportional drug-induced change in the subjective rate of exertion required to hold down the lever; unitless

In the vehicle condition, 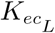 assumes an implicit value of one.

##### f_a_

We do not know which aspects of the leisure activities that compete with pursuit of BSR give rise to reward signals in the brain nor how these signals are encoded. That is why both the arguments and output of the seventh shell → core function (*f*_*a*_) have been left blank. However, we do know that valuation of these activities influences the allocation of time to pursuit of experimenter-controlled rewards: enrichment of the test environment shifts allocation towards alternate activities and away from the experimenter-controlled reward [16]. Thus a function such as *f*_*a*_ must exist. We include *f*_*a*_ in the list of shell → core functions for completeness and in recognition of this requirement.

#### Core functions

Core functions determine the reward rates produced by work (lever depression) and leisure (alternate) activities. These are combined with the associated effort costs to yield a pair of payoffs, which are then passed to the core → shell function for translation into time allocation.

##### f_R_

The core receives a set of spike trains as a result of the combined action of the first three shell → core functions {*f*_*F*_, *f*_*N*_, *f*_*D*_}, one spike train from every activated first-stage neuron. According to the counter model [10, 11, 17], the effects of the spike trains delivered by the individual first-stage neurons are summed, and thus, the intensity of the rewarding effect is determined by the product of the number of activated first-stage neurons and the rate at which they are fired by the stimulation train. This is why a π symbol is used in the flow diagrams to represent the drive produced by a pulse train of fixed duration on the scalar at the input of the reward-growth function (Fig 2).

The logistic form of the reward-growth function was described originally in operant-matching studies carried out by Gallistel’s group [18–20]. Shizgal [21] proposed the following expression for this function:

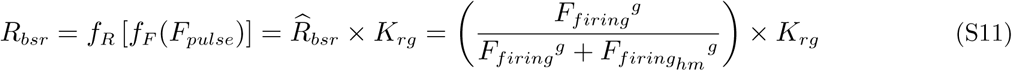

where

*f*_*R*_ = the reward-growth function

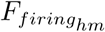 = firing rate required to drive reward intensity to half its maximum value; units: *firings neuron*^−1^ *s*^−1^ 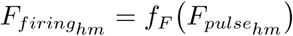

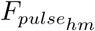 = pulse frequency required to drive reward intensity to half its maximum value; units: *pulses s*^−1^

*g* = the exponent that determines the steepness of reward- intensity growth as a function of pulse frequency

*K*_*rg*_ = reward-growth scalar; units: *hedons*

*R*_*bsr*_ = reward intensity produced by *F*_*firing*_; units: *hedons*

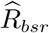 = normalized reward intensity produced by a pulse frequency of *F*_*pulse*_, which, in turn, produces a firing frequency of *F*_*firing*_ in each first-stage neuron; unitless. 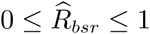

According to Eq S11,

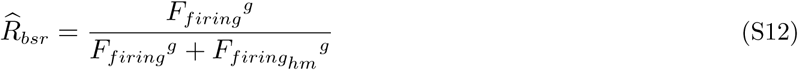

When frequency-following fidelity is sufficiently high, the normalized reward intensity 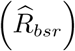 will approach one at high pulse frequencies. The lower the value of the location parameter 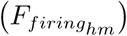 and the higher the value of the reward-growth exponent (*g*), the easier this will be to achieve.

To accommodate the predicted rescaling of the input to the reward-growth function for oICSS by dopamine-transporter blockade, a scalar, *K*_*da*_, is added to Eq S11 in the drug condition of the experiment:

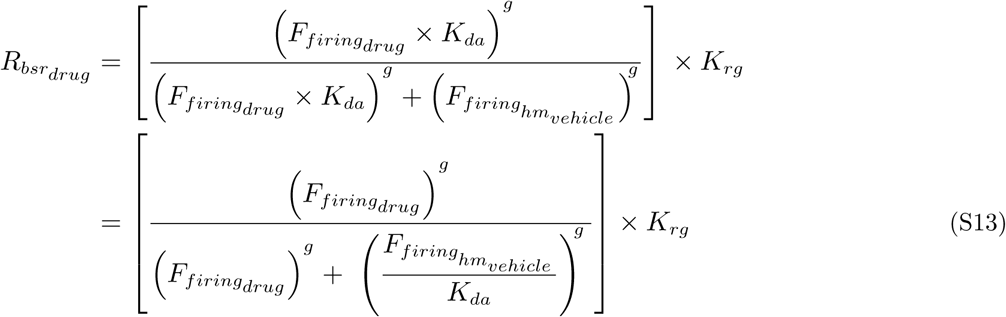

where

*K*_*da*_ = scalar representing the boost in dopamine release due to transporter blockade

Thus,

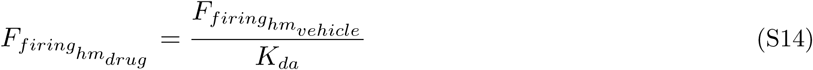

Dopamine-transporter blockade boosts dopamine release, thereby increasing the impact of each firing. The increased value of the scalar (*K*_*da*_) captures this augmented impact, reducing the value of the location parameter of the reward-growth function in the drug condition. Fewer pulses per train are required to produce a reward of a given intensity when *K*_*da*_ increases. Consequently, the reward-growth function shifts leftwards along the pulse-frequency axis. In the simulations of the vehicle condition, *K*_*da*_ is assigned an implicit value of one.

Division by the subjective price transforms the reward intensity into a reward rate:

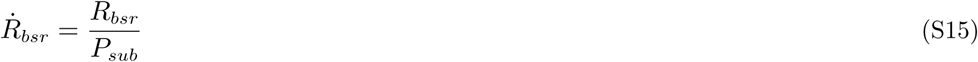

where

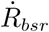 = experienced rate of brain stimulation reward; units: *hedons s*^−1^

##### f_H_

The location parameter of the reward-growth function is the firing rate that drives reward intensity to half its maximal value. The value of this parameter depends on the number of stimulated first-stage neurons, *N* and the interval during which the stimulation train elevates their firing rate, *D*_*burst*_. A prior study of temporal integration in the neural circuitry responsible for eICSS of the MFB [21] implies the following form for the function that determines 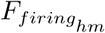:

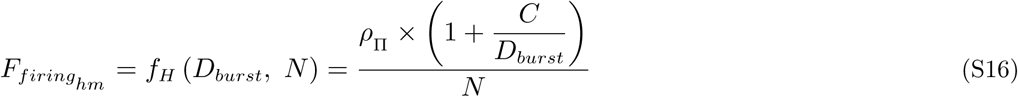

where

*C* = chronaxie: train duration at which 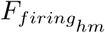 is twice the value of *ρ*_π_; units: *s*

*ρ*_π_ = aggregate rate of firing required to produce a reward of half-maximal intensity when the train duration is infinite; units: *firings s*^−1^

##### f_U_

In keeping with the generalized matching law [22], the benefit from work and its costs are combined in scalar fashion to yield a net payoff. We can format the expression for the payoff as a ratio of two rates: the probability-weighted rate of brain-stimulation reward and the subjective rate of exertion required to hold down the lever (see [23]):

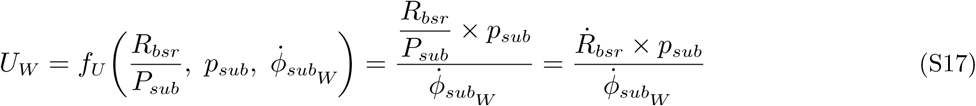

where

*f*_*U*_ = utility function

*U*_*W*_ = payoff from a train of rewarding stimulation; units: *hedons oomph*^−1^

or as a benefit/cost ratio:

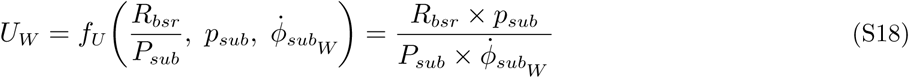

In the case of BSR, there is solid evidence that aggregate impulse flow in the first-stage neurons encodes the signal that will be translated into the intensity of the reward [17, 20]. Although the identity of the first-stage neurons subserving eICSS of the MFB (or any other brain site) remains unknown, directly-driven MFB neurons with properties that match the psychophysically derived portrait of the first-stage fibers have been observed by means of electrophysiological recording [24–26].

The argument of the reward-growth function for leisure activities is left blank, thus signifying our ignorance of how pursuit of these activities is encoded by the brain and translated into a reward rate. That said, the considerable evidence that the value of leisure activities competes effectively with experimenter-controlled rewards [5, 16, 27–29] implies that the payoffs from work and leisure are commensurable. Accordingly, we define a reward rate for the leisure activities that compete with pursuit of BSR:

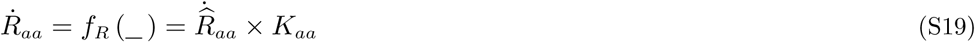

where

*K*_*aa*_ = alternate-activity scalar; units: *hedons s*^−1^

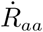 = average rate of reward from alternate (leisure) activities that compete with pursuit of BSR; units: *hedons s*^−1^

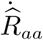 = normalized rate of reward from alternate (leisure) activities; unitless. 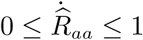

(_) = the unknown variables that give rise to 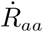

The payoff from these alternate activities is computed in a manner analogous to the computation of the payoff from BSR, as a ratio of reward and subjective-exertion rates:

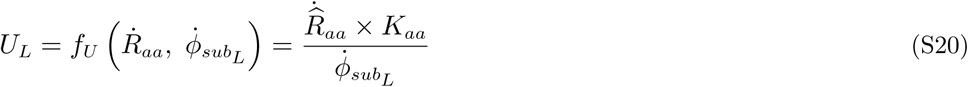

where

*U*_*L*_ = payoff from leisure activities; units: *hedons oomph*^−1^

#### The core → shell function

##### f_T_

The payoffs from pursuit of BSR (*U*_*W*_) and engagement in leisure activities (*U*_*L*_) are used by the sole core → shell function to compute the allocation of time to pursuit of BSR. This behavioral-allocation function is derived from the single-operant matching law [5, 30, 31]:

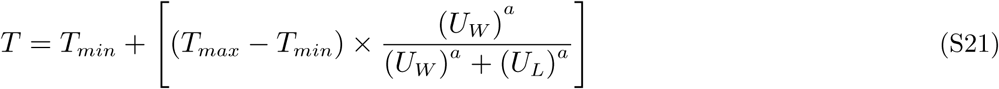

where

*a* = price-sensitivity exponent; unitless

*T* = time allocation; unitless

*T*_*max*_ = maximum time allocation; unitless

*T*_*min*_ = minimum time allocation; unitless

*U*_*W*_ = payoff from pursuit of rewarding brain stimulation (“work”); units: *hedons oomph*^−1^

*U*_*L*_ = payoff from pursuit of alternate (“leisure”) activities; units: *hedons oomph*^−1^

Even when the rat is working maximally, latency to depress the lever is typically greater than zero, and *T*_*max*_ is thus typically less than one. The rat tends to sample the lever at trial onset, even when the payoff from brain stimulation is low. Thus, *T*_*min*_ is typically greater than zero.

The single-operant matching law was formulated initially to account for the behavior of subjects working on variable-interval schedules of reinforcement. The cumulative handling-time schedule in force in the present study [32] is more akin to a fixed-ratio schedule in that the number of rewards earned is strictly proportional to time worked. On such schedules, time allocation shifts more abruptly than on variable-interval schedules as the value of the experimenter-controlled reward is varied. In the original version of the single-operant matching law [30, 31], the payoff terms are not exponentiated. In contrast, the price-sensitivity exponent (*a*) in Eq S21 allows the reward-mountain model to account for time-allocation shifts of varying abruptness.

To simplify the remaining derivation of the reward-mountain model, we define a normalized measure of time allocation:

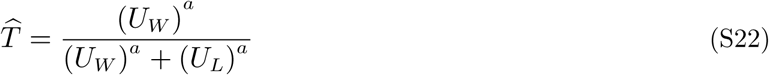

where 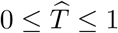

Substituting from Eq S22 in Eq S21, we obtain

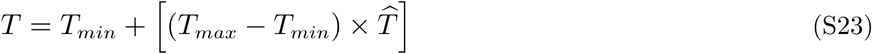

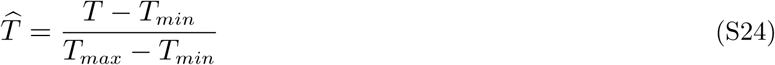

#### Time allocation as a function of reward strength and cost

We are now in a position to tie the dependent variable, time allocation (*T*), to the two independent variables, pulse frequency (*F*_*pulse*_) and price (*P*_*obj*_). Substitution for *U*_*W*_ and *U*_*L*_ from Eqs S17 and S20, respectively, in Eq S22 yields:

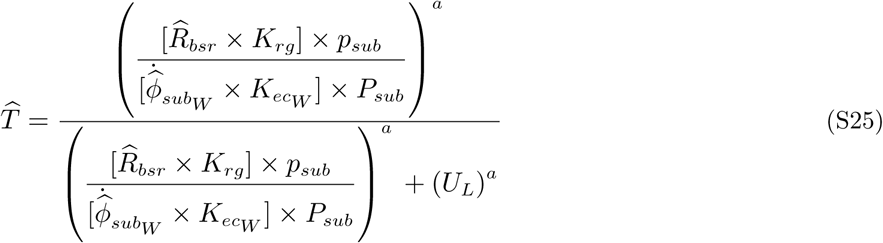

An initial step towards simplifying Eq S25 is to multiply each of the terms on the right by

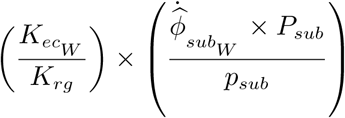

This yields:

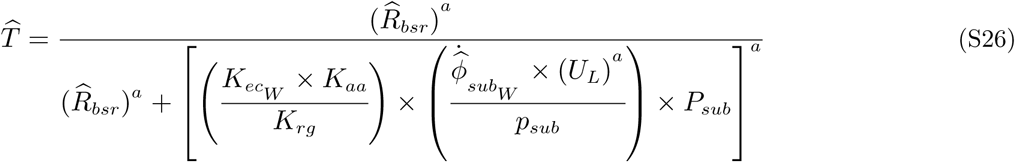

To simplify Eq S25 further, we first define *T*_*mid*_ as the time-allocation value midway between maximal and minimal time allocation:

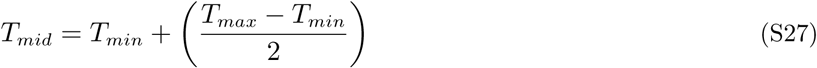

According to Eqs S22 and S23,

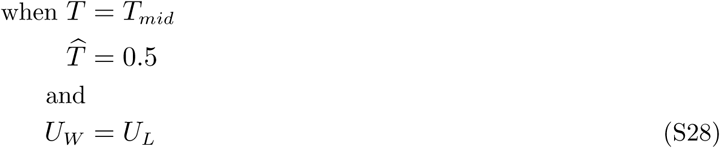

We now hold time allocation at *T*_*mid*_ and drive reward intensity to its maximum value 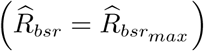. Substituting for *U*_*W*_ from Eq S17 and reversing Eq S28, we obtain

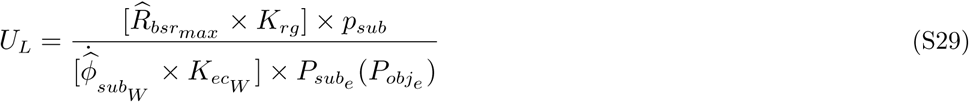

where

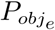 = objective price at which *T* = *T*_*mid*_ when 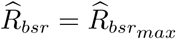

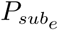 = subective price at which *T* = *T*_*mid*_ when 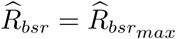

Rearranging the terms yields

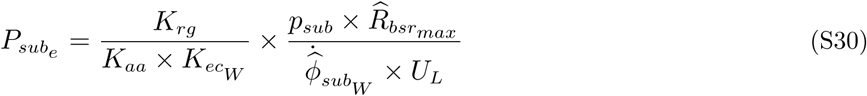

For consistency with previous papers, we use the subscript, “e” in the symbol 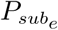 to refer to the fact that the payoff from brain stimulation equals the payoff from alternate activities (“everything else”) when the normalized reward intensity is maximal 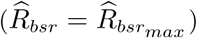 and the subjective price (*P*_*sub*_) equals 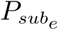. This equivalence between the two competing payoffs is what drives time allocation to the half-way point (*T*_*mid*_) between its minimal (*T*_*min*_) and maximal (*T*_*max*_) values.

Rearranging Eq S30, we obtain:

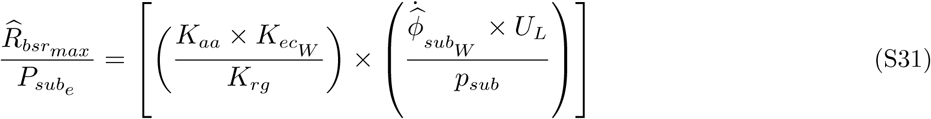

Substituting in Eq S26 from Eqs S11, S24, and S31, we obtain

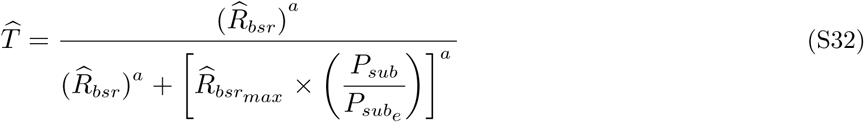

By setting 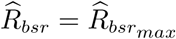 and 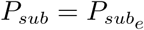 in Eq S32, it can be seen readily that 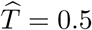, thus satisfying the definition of 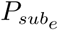 as the price at which time allocation to pursuit of a maximally intense reward is halfway between *T*_*min*_ and *T*_*max*_.

*P*_*sub*_ and 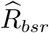 must trade off to hold 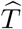 at a given level. The lowest attainable subjective price is the value corresponding to an objective price of zero, which we will call 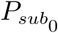. (Negative values of *P*_*obj*_ may be required to drive *P*_*sub*_ to 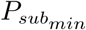.) Consider a vector of subjective prices, 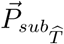, that extends from 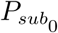 to the highest tested value of *P*_*sub*_ and a corresponding vector of normalized reward intensities, 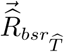, that hold normalized time allocation 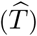 constant over 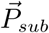. If follows from Eq S32 that

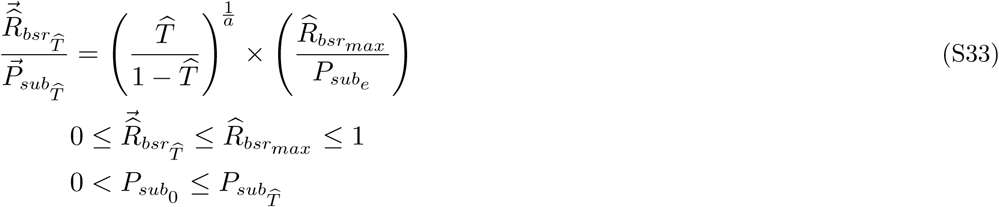

where

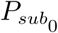 = the subjective price corresponding to an objective price of zero

The higher the subjective price, the higher the normalized reward intensity required to hold time allocation constant. When 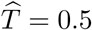, Eq S33 reduces to:

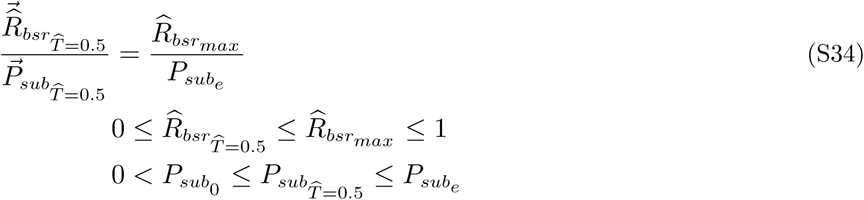

Below (see: Contour lines: the trade-off between pulse frequency and price to hold time allocation constant), we use Eqs S33 and S34 to obtain the equation for the contour lines that provide a two-dimensional description of the reward-mountain surface.

To complete the derivation of the reward-mountain model, we now substitute for 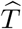 in Eq S23 and expand Eq S32 so that time allocation is expressed in terms of the independent variables: price (*P*_*obj*_) and pulse frequency (*F*_*pulse*_), which appear as the arguments of the subjective-price and frequency-following functions, respectively:

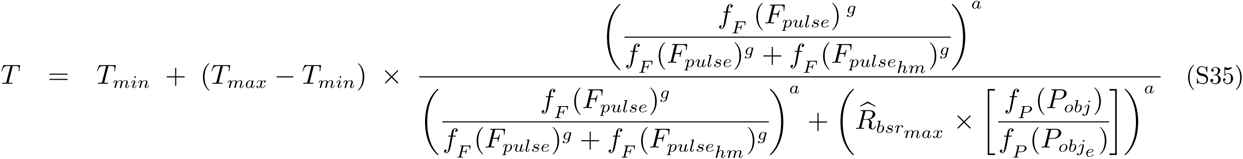

#### The conditioned-reward variant of the reward-mountain model

The six-parameter version of the reward-mountain model incorporates Eqs S1, S6, and S35. The fitted parameters are *a* (the price-sensitivity exponent), 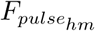 (the pulse frequency at which reward intensity is half maximal), *g* (the reward-growth exponent) 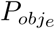 (the price at which time allocation to pursuit of a maximal reward falls midway between its minimal and maximal values), *T*_*min*_ (minimum time allocation), and *T*_*max*_ (maximal time allocation). The seven-parameter version includes an additional parameter, *C*_*r*_, to reflect conditioned reward. This seventh parameter reflects a learned value, above and beyond the payoff from the stimulation train, associated with the lever and/or the act of holding it down. The paper that introduced this parameter [3] incorporated it into the reward-growth function as follows:

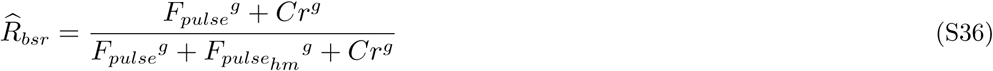

where

*C*_*r*_ = the conditioned reward, expressed in terms of the equivalent pulse frequency

Note that the way the *C*_*r*_ parameter was incorporated into Eq S36 causes this parameter to interact with both the *a* (Eq S35) and *g* (Eq S36) parameters. That form of the model failed to yield consistently converging fits when applied to the current dataset. To address this problem, we altered the way that the *C*_*r*_ parameter is incorporated into the reward-growth function (Eq S11) so as to reduce its interaction with the other parameters:

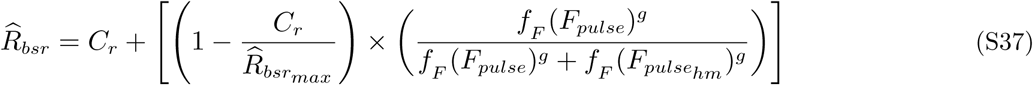

The resulting reward-mountain surface is produced by substituting the expression for 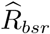 from Eq S37 in Eq S32, as follows:

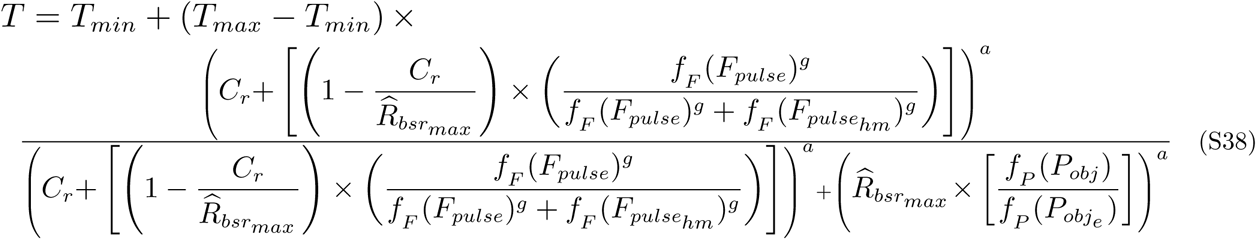

As shown in Fig S15, Eq S38 yields a reward-mountain surface that is all but indistinguishable from the surface generated by the equation in the 2010 paper. Unlike the equation in the 2010 paper, Eq S38 produced well-behaved, converging fits.

**Fig S15.**
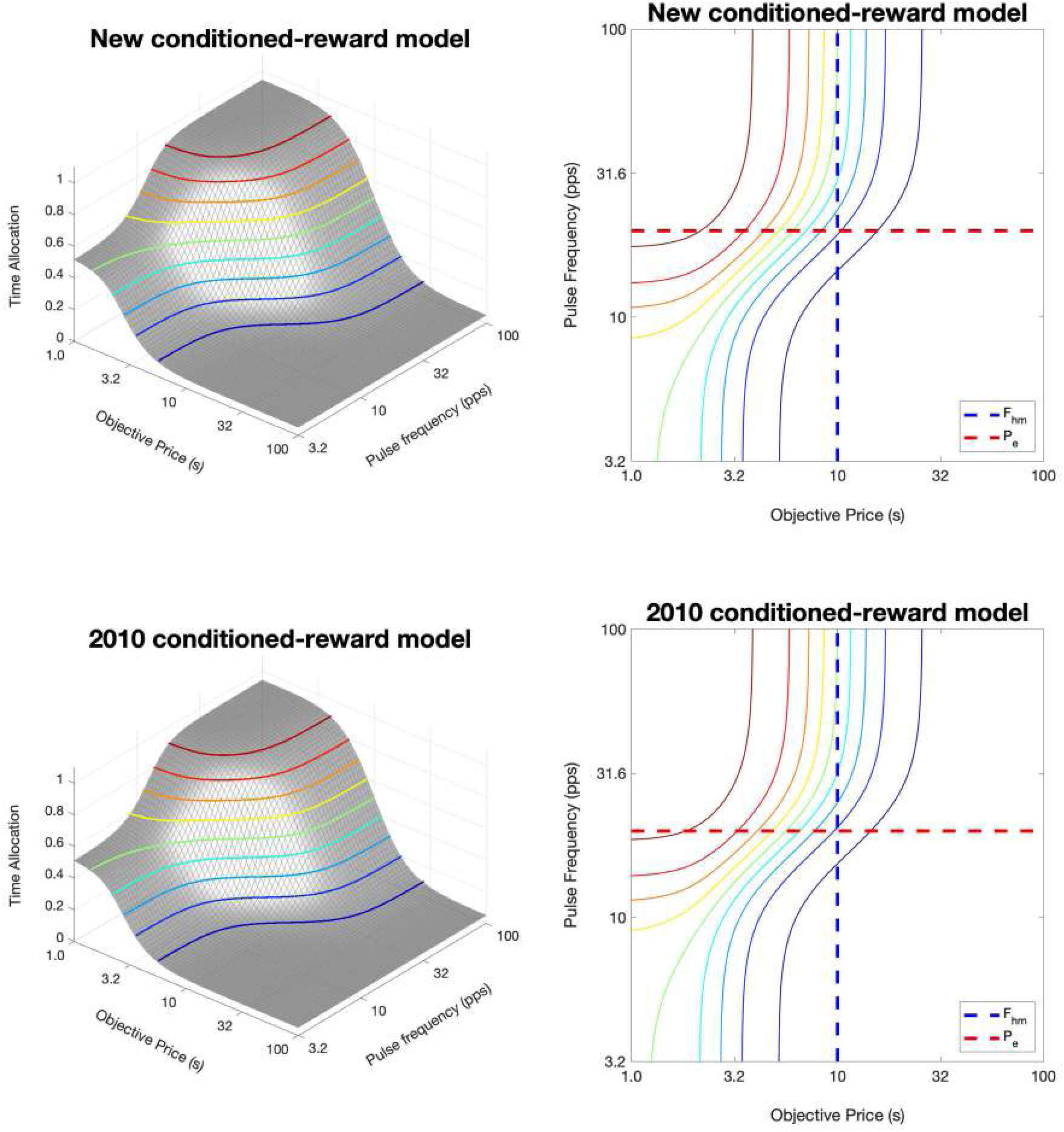
Surface and contour plots of two seven-parameter reward-mountain models. The new version of the model produces a surface that is nearly identical to the one generated by the 2010 version of this model [3]. Fits of the 2010 model to the current datasets failed to converge, whereas fits of the new version of the model converged in all cases.

### Adaptation of the reward-mountain model for oICSS

**Fig S16.**
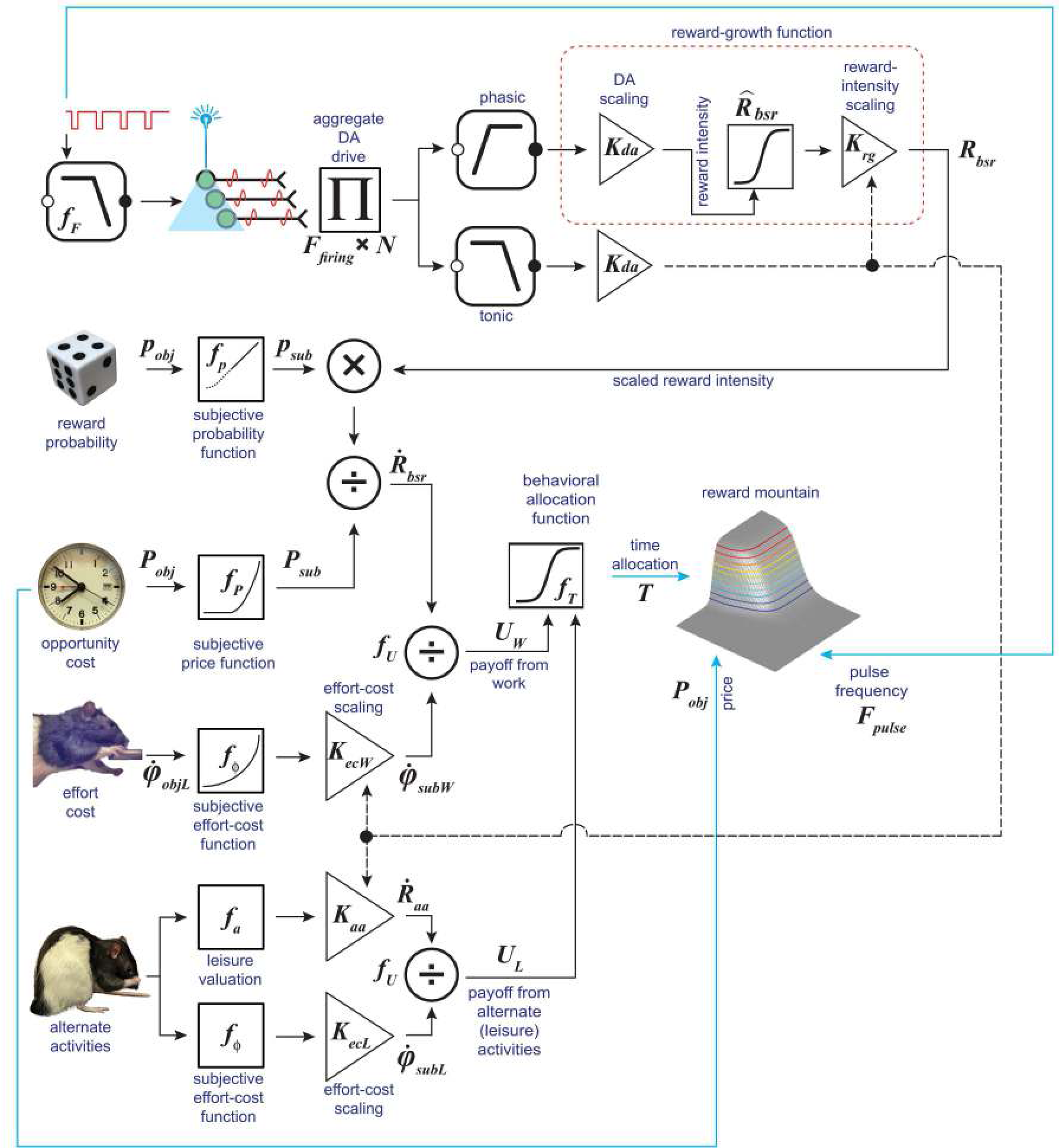
The reward-mountain model for oICSS. A graphical summary of the reward-mountain model, as adapted for oICSS of midbrain dopamine neurons. The symbols are defined in Tab S1.

The reward-mountain surface shows the observed behavioral output (time allocation, *T*) as a function of the two independent variables, the price (*P*_*obj*_) and pulse frequency (*F*_*pulse*_). These three variables constitute the shell of the reward-mountain model, together with the controlled variables: the work rate required to hold down the lever 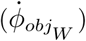, the leisure activities afforded by the test environment, and the average work rate required to perform these activities 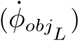. The shell → core functions, {*f*_*F*_, *f*_*p*_, *f*_*P*_, *f*_*ϕ*_, *f*_*a*_}, map the manipulated and controlled variables into the core quantities that determine the payoffs from work and leisure. These mapping functions are shown at the left of the figure. The core functions that compute the reward intensities and corresponding payoffs {*f*_*R*_, *f*_*U*_} are shown in the center. (*f*_*H*_, the function that determines the location parameter of the reward-growth function, is not shown, nor are *f*_*D*_, the function that determines the duration of the stimulation-induced burst of firing and *f*_*N*_, the function that determines the number of recruited neurons (*N*).) The core → shell function, (*f*_*T*_), translates the payoffs from work and leisure into time allocation. It is shown on the right.

Minimal changes were made to adapt the model prior to fitting the reward-mountain surface to the data:

1. New values were used for the parameters of the frequency-following function (Eq S1) so as to accommodate the known properties of midbrain dopamine neurons and the optical-power versus pulse-frequency trade-off data reported here.
2. In eICSS, the current and pulse duration conjointly determine the number of directly stimulated neurons. In oICSS, optical power plays the role assumed by current in eICSS. However, there is insufficient information available to model *f*_*N*_, the function that determines the number of neurons directly activated by a given optical power and pulse duration. Instead, we simply chose values of *N* and *ρ*_π_ that produce simulated reward-mountain surfaces similar to the ones returned by the fits of the model to the data. (See the accompanying Matlab^®^ Live Script.)
3. 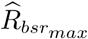 was included explicitly in the reward-mountain model, thus expanding the model to accommodate imperfect frequency-following fidelity.
4. Explicit inclusion of 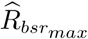 required correction of the location parameters and a more nuanced treatment of drug-induced shifts in the location of the mountain. (See: Displacement of the shell: distinguishing two sources.)
5. The new conditioned-reward variant described above (Eq S38) was used in lieu of the version introduced in our prior studies of the effect of cocaine and GBR-12909 on reward mountains obtained in the Eicss paradigm [3, 33].

#### Parameters of the frequency-following function for oICSS

The form of the frequency-following function (*f*_*F*_) for channelrhodopsin-2-mediated excitation of midbrain dopamine neurons has yet to be determined. Here, we used a function of the same form as the one we had determined previously for eICSS of the MFB [6], but we substituted new parameter values. Dopamine neurons cannot fire nearly as fast as the directly stimulated neurons subserving the rewarding effect of electrical MFB stimulation [6, 34, 35], and the kinetics of channelrhodopsin-2 are slow in comparison to those of the voltage-gated channels responsible for electrically induced neural firing [36]. Thus, frequency-following parameters determined for eICSS of the MFB cannot be used to account for the frequency response of the dopamine neurons subserving oICSS of the ventral midbrain.

Although two studies found that frequency-following fidelity in optically stimulated midbrain dopamine neurons fell to only 40-50% at optical pulse frequencies of 40-50 pulses s^−1^ [34, 35], results of two other electrophysiological studies show very good firing fidelity at 50 pulses s^−1^ [37, 38]. Moreover, results of a recent electrochemical study [39] show that optically induced dopamine release in the nucleus accumbens continued to rise as the pulse frequency was increased from 40-50 pulses s^−1^. Poor frequency-following fidelity was found by Lohani and colleagues in midbrain dopamine neurons optically stimulated at 100 pulses s^−1^ [40], suggesting that the upper limit on the induced firing rate lies at a significantly lower pulse frequency.

Figs 3 and S5 show two cases (data from rats BeChR29 and 27) in which the behavioral effectiveness of the stimulation continues to rise at pulse frequencies up to, or beyond, 60 pulses s^−1^. The curves for the remaining rats approach asymptote earlier, but this does not necessarily reflect failure the higher pulse frequencies to increase firing: saturation of reward-intensity growth [20] could be responsible instead.

In view of the power-frequency trade-off data reported here and the results reported in the studies cited above, we set the middle of the roll-off region of the frequency-following function (*F*_*ro*_) to 50 pulses s^−1^ and the parameter governing the abruptness of the roll-off (*F*_*bend*_) to 20. The resulting frequency-following function is shown in Fig S13. It continues to climb up to optical pulse frequencies of ∼100 pulses s^−1^ to attain a maximum induced firing rate of 51.6 spikes s^−1^.

#### Displacement of the shell: distinguishing two sources

The purpose of the experiment is to draw inferences about brain-reward circuitry from the effect of dopamine-transporter blockade on the location of the shell of the reward mountain within the space defined by the two independent variables, the **objective** opportunity cost of the reward, *P*_*obj*_, and the ***pulse*** frequency, *F*_*pulse*_. However, the core variables about which the inferences are to be drawn operate in spaces defined by the ***subjective*** opportunity cost, *P*_*sub*_, and the induced frequency of ***firing***, *F*_*firing*_ (Tab S2). As we explain below, the core → shell function, *f*_*F*_, which maps the pulse frequency into the evoked frequency of firing, contributes to the estimates of both location parameters of the shell. Its influence must be removed in order to isolate the effects of the drug on 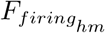 and 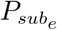, the location parameters of the core. In the depiction and interpretation of the fitted surfaces, we will distinguish between

1. ***primary*** displacement of the shell of the reward mountain due to drug actions on the core components and
2. ***secondary*** (additional) displacement of the shell due to the differential response of the frequency-following function (*f*_*F*_) in the drug and vehicle conditions

In the following section, we explain how to remove the secondary displacements from the location-parameter estimates, thus isolating the primary displacements that are the focus of the study.

#### Correction of the location-parameter estimates for changes in frequency-following fidelity

Fitting the mountain surface to time-allocation data returns the two location parameters of the reward-mountain shell: 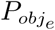, which positions the shell along the price axis, and 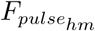, which positions the shell along the pulse-frequency axis. Whereas the value of 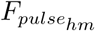 is independent of the value of 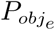, the reverse does not hold when frequency-following fidelity is imperfect. Changes in frequency-following fidelity alter the maximum reward intensity 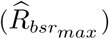, which contributes to the value of both 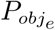 and its subjective equivalent, 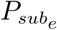. The portion of the shifts in 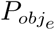 due to changes in frequency-following fidelity must be removed in order decouple estimates of that parameter from shifts along the pulse-frequency axis. Once 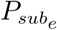 has been suitably corrected, manipulations that act at, or beyond, the output of the reward-growth function (*f*_*R*_, Fig S16) shift the mountain core uniquely along the price axis, whereas manipulations that alter the input to the reward-growth function shift the mountain core uniquely along the frequency axis [2, 3, 41].

Imperfect frequency-following fidelity can also alter the extent to which the surface of the mountain shell shifts along the pulse-frequency axis. This problem will arise if frequency-following fidelity differs in the drug and vehicle conditions. Correction is required in order to estimate the shift of fundamental interest, which is the displacement of the reward-growth function (a core component) along its firing-frequency axis.

##### The frequency-following function

The correction of the location-parameter estimates arises from the form of the frequency-following function (*f*_*F*_). The form and parameters of this function for eICSS of the MFB were described by Solomon et al. [14]. In the absence of analogous data for oICSS, we have assumed the same functional form but have tuned the parameters to accommodate the power-frequency trade-off data reported here and results of prior studies [37–40].

In double-logarithmic coordinates, the frequency-following function (Fig S13) has the form of an inverted hockey stick, with a straight handle that transitions into a flat blade [6]. Pulse frequency is represented along the abscissa of this function, and firing frequency is represented along the ordinate. The induced firing frequency follows the pulse frequency perfectly over the portion of the handle that is truly straight, grows more slowly over a transition zone, and levels off over the blade portion. Thus, within the transition zone, a given drug-induced decrement in firing frequency (a core variable) will correspond to a larger decrement in pulse frequency (a shell variable). Drug-induced displacement of the reward-growth function (*f*_*R*_) towards lower values of 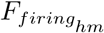 improves frequency-following fidelity (by moving the pulse frequency toward, or onto the straight “handle”). This causes the displacement of the shell to exceed, and thereby overestimate, the underlying displacement of the reward-growth function.

To correct our estimate of how far the reward-growth function has shifted along the pulse-frequency axis, we need to decouple its value from the maximum normalized reward intensity, 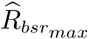. This is achieved by using the assumed frequency-following function (Eq S1) to estimate 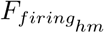 from 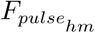.

We define the corrected estimate of the location parameter as follows:

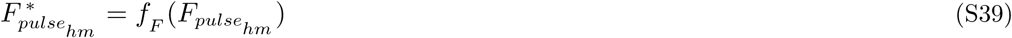

where

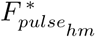 = the estimated value of 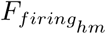, which is the value 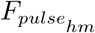 would have attained had frequency-following fidelity been perfect

Eq S39 states that 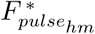 and 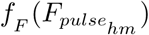 are one and the same. We will plot the value in question in the coordinate space of the shell. There, the pulse frequency, rather than the firing frequency, serves as the ordinate.Thus, in that context, we use 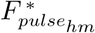 in lieu of 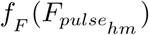 as our notation.

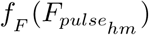 (and thus 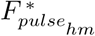)is a value along the ordinate of the frequency-following function (Fig S13), whereas 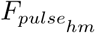 is a value along the abscissa. Given the form of the frequency-following function, 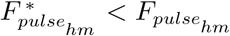 once pulse frequency exceeds the capacity of the neurons to fire reliably to each and every pulse.

Several steps are required to correct the the parameter that locates the reward mountain along the price axis so that it too is decoupled from frequency-following fidelity. The step first is analogous to the estimation of 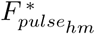 from 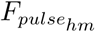: We use the subjective-price equation (Eq S6) to transform 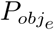 into its subjective equivalent, 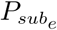.

The next step is to correct 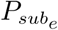 for the effect of imperfect frequency-following fidelity. Eq S30 can be rearranged as follows:

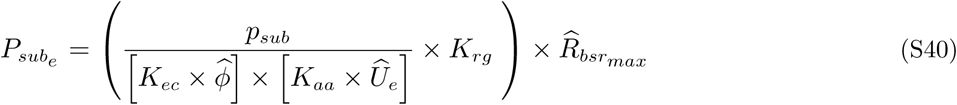

Eq S40 reminds us that 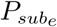 is proportional to the maximum normalized reward intensity that can be attained, 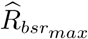. Eq S12 defines 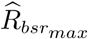 in terms of the maximal attainable firing frequency, 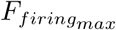, the firing frequency corresponding to 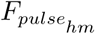, and the reward-growth exponent, *g*. To estimate 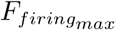, we solve Eq S1 for a pulse frequency more than high enough to drive firing frequency to its maximum (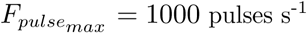) We then use the resulting estimate of 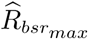 to produce a revised estimate of 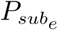:

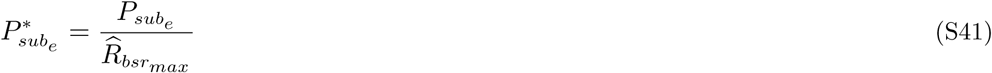

where

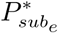 = estimated value that 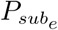 would have attained had frequency-following fidelity been perfect

Last, we transform 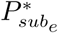 into its objective-price counterpart, 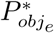 by passing 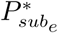 through the back-solution of the subjective-price equation [6]:

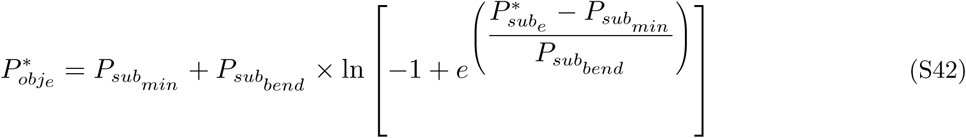

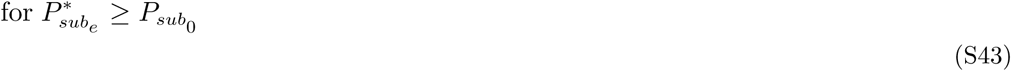

The transformation of 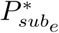 into 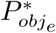 is performed so that the corrected location-parameter estimate can be plotted in the space defined by the independent variables, {*P*_*obj*_, *F*_*pulse*_}. In this space, drug-induced shifts in the position of the reward mountain, corrected for changes in frequency-following fidelity, are depicted as 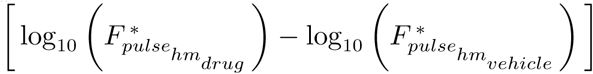 and 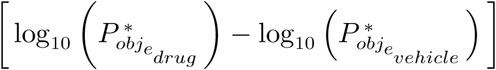.

### Model fitting and selection

#### Model selection

The 12 candidate models fit to the data are described in Tab S3. The models differ in the total number of parameters as well as in the number of parameters free to vary across the vehicle and drug conditions. Models 2, 5, 8, and 11 are based on the six-parameter version of the reward-mountain model (Eq S35), whereas the remaining models are based on the seven-parameter version (Eq S38). The fits of all of the candidate models to the reward-mountain data from all seven rats converged successfully.

**Table S3.** The 12 candidate models fit to each dataset. Values of the “free” parameters were free to differ between the vehicle and drug conditions, whereas a single value was fitted to the data from both conditions in the case of “_com” (common) parameters. Additional columns list the total number of “Common” and “Free” parameters along with their “Totals.” Models 2, 5, 8, and 11 are based on the six-parameter version of the reward-mountain model (Eq S35), whereas the remaining models are based on the seven-parameter version (Eq S38).

| Num | a_free | g_free | CR_free | CR_com | Common | Free | Total |
| --- | --- | --- | --- | --- | --- | --- | --- |
| 1 | 0 | 0 | 1 | 0 | 4 | 6 | 10 |
| 2 | 0 | 0 | 0 | 0 | 4 | 4 | 8 |
| 3 | 0 | 0 | 0 | 1 | 5 | 4 | 9 |
| 4 | 1 | 0 | 1 | 0 | 3 | 8 | 11 |
| 5 | 1 | 0 | 0 | 0 | 3 | 6 | 9 |
| 6 | 1 | 0 | 0 | 1 | 4 | 6 | 10 |
| 7 | 0 | 1 | 1 | 0 | 3 | 8 | 11 |
| 8 | 0 | 1 | 0 | 0 | 3 | 6 | 9 |
| 9 | 0 | 1 | 0 | 1 | 4 | 6 | 10 |
| 10 | 1 | 1 | 1 | 0 | 2 | 10 | 12 |
| 11 | 1 | 1 | 0 | 0 | 2 | 8 | 10 |
| 12 | 1 | 1 | 0 | 1 | 3 | 8 | 11 |

Tab S4 ranks the fits of the 12 candidate models for one rat (Bechr29) by their evidence ratios (the relative likelihood that a candidate model is true in comparison to the best-fitting model). Note that the residual sum of squares for the worst-fitting model (model 10) is slightly lower than in the case of the best-fitting model (model 2). This is not surprising given that the worst-fitting model comprises 12 parameters whereas the best-fitting model comprises only eight. The Akaike Information Criterion (AIC) [42] implements a trade-off between goodness of fit and simplicity. Thus, the AIC penalizes models with large number of parameters in comparison to simpler ones. On the basis of the AIC, model 10 is over 5 × 10^5^ times less likely than model 2 and is ranked accordingly. Summary statistics for the best-fitting model for each rat are listed in Tab S5.

**Table S4.** Model-evaluation statistics for the fit of the 12 candidate models to the data from rat Bechr29. The models are described in Tab S3. “AIC” stands for the Akaike Information Criterion [42]. “Likelihood” refers to the ratio: 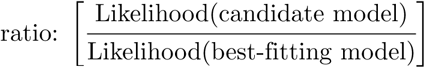, which expresses the relative likelihood that a candidate model is true in comparison to the best-fitting model. The Evidence Ratio (“Ev ratio”) is the inverse of the likelihood ratio. The total number of parameters is listed under “prms,” the residual sum of squares under “RSS,” the total sum of squares under “TSS,” and the adjusted *R*^2^ under “Adj *R*^2^.”

| Model | AIC | Likelihood | Ev_ratio | prms | RSS | TSS | Adj $R^2$ |
| --- | --- | --- | --- | --- | --- | --- | --- |
| 2 | -1,922.58 | 1.000000 | 1.00 | 8 | 13.904 | 12,647.57 | 0.998886 |
| 8 | -1,916.12 | 0.039636 | 25.23 | 9 | 13.897 | 12,647.57 | 0.998885 |
| 5 | -1,915.86 | 0.034860 | 28.69 | 9 | 13.904 | 12,647.57 | 0.998884 |
| 3 | -1,915.86 | 0.034846 | 28.70 | 9 | 13.904 | 12,647.57 | 0.998884 |
| 11 | -1,909.46 | 0.001421 | 703.74 | 10 | 13.896 | 12,647.57 | 0.998883 |
| 9 | -1,909.40 | 0.001377 | 726.48 | 10 | 13.897 | 12,647.57 | 0.998883 |
| 6 | -1,909.14 | 0.001211 | 825.83 | 10 | 13.904 | 12,647.57 | 0.998882 |
| 1 | -1,909.14 | 0.001210 | 826.19 | 10 | 13.904 | 12,647.57 | 0.998882 |
| 12 | -1,902.74 | 0.000049 | 20,330.45 | 11 | 13.896 | 12,647.57 | 0.998881 |
| 7 | -1,902.69 | 0.000048 | 20,826.76 | 11 | 13.897 | 12,647.57 | 0.998880 |
| 4 | -1,902.42 | 0.000042 | 23,852.26 | 11 | 13.904 | 12,647.57 | 0.998880 |
| 10 | -1,896.02 | 0.000002 | 585,023.15 | 12 | 13.895 | 12,647.57 | 0.998878 |

**Table S5.**
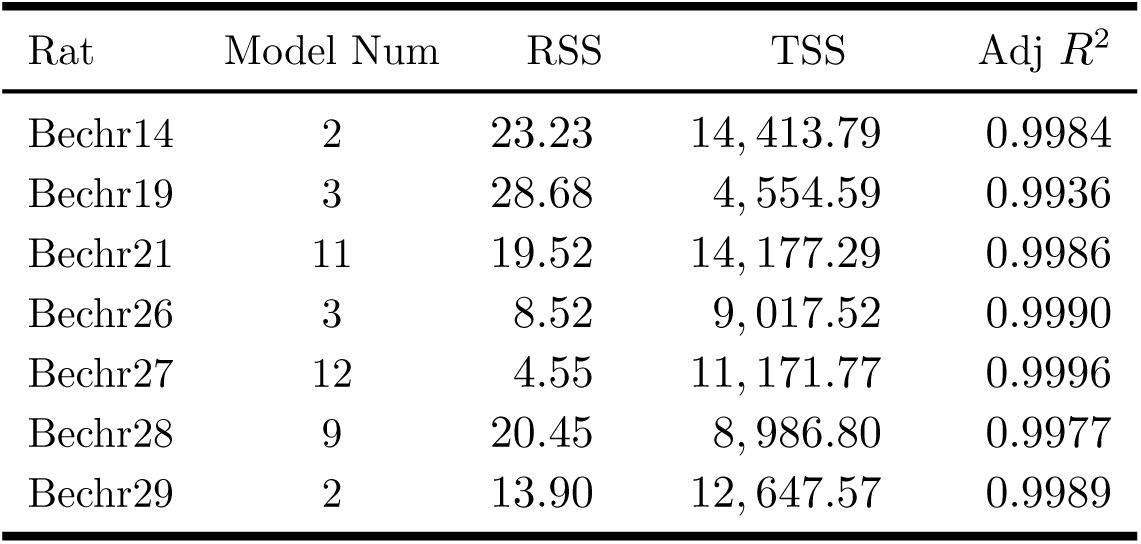
Summary statistics for the model that provided the best fit (highest evidence ratio) to the data for each rat. The models in the “Model Num” column are defined in Tab S3. The residual sum of squares is listed in column “RSS,” the total sum of squares in column “TSS,” and the adjusted R^2^ in column “Adj *R*^2^.”

| Rat | Model Num | RSS | TSS | Adj $R^2$ |
| --- | --- | --- | --- | --- |
| Bechr14 | 2 | 23.23 | 14,413.79 | 0.9984 |
| Bechr19 | 3 | 28.68 | 4,554.59 | 0.9936 |
| Bechr21 | 11 | 19.52 | 14,177.29 | 0.9986 |
| Bechr26 | 3 | 8.52 | 9,017.52 | 0.9990 |
| Bechr27 | 12 | 4.55 | 11,171.77 | 0.9996 |
| Bechr28 | 9 | 20.45 | 8,986.80 | 0.9977 |
| Bechr29 | 2 | 13.90 | 12,647.57 | 0.9989 |

Different variants of the reward-mountain model provided the best fit to the data from different rats. Tab S6 shows the best-fitting model for all rats, as determined by the AIC-based evidence ratio. In four cases (Bechr14,19,26,29), the best-fitting model was one in which common values of the *a* and *g* parameters were fit to the data from both the vehicle and drug conditions; in the remaining three cases, the best-fitting model was one in which the values of *a, g*, or both were free to vary across the vehicle and drug conditions.

In four cases (Bechr19, 26,27,28), the *C*_*r*_ parameter was included in the best-fitting model, whereas in the three remaining cases, it was not. As Fig S15 illustrates, the *C*_*r*_ parameter will be advantageous when time allocation in low-payoff trials is higher along pulse-frequency sweeps than along price or radial sweeps. In no case was the value of this parameter free to vary across the vehicle and drug conditions in the best-fitting model. Thus, there is no evidence that the advantage conferred by addition of the *C*_*r*_ parameter was due to dopamine-transporter blockade.

**Table S6.**
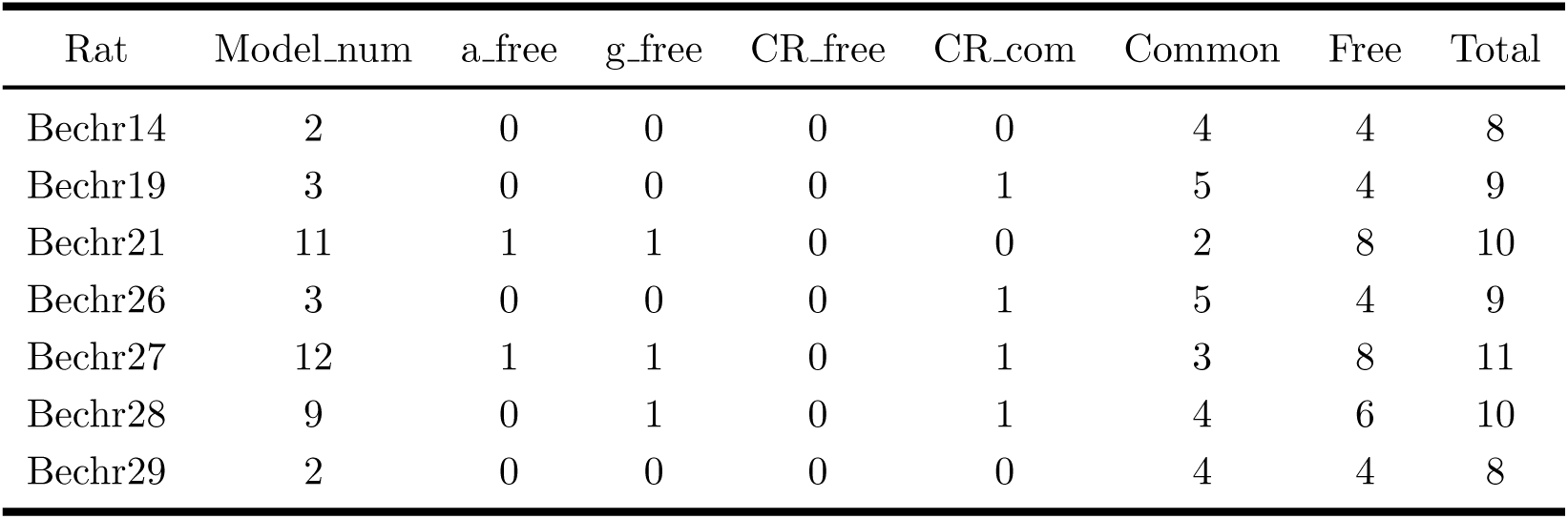
Best-fitting models for all rats. The models are described in Tab S3. Values of the “free” parameters were free to differ between the vehicle and drug conditions, whereas a single value was fitted to the data from both conditions in the case of “com” (common) parameters. Additional columns list the total number of common and free parameters and their totals

### Fitted reward-mountain surfaces

**Fig S17.**
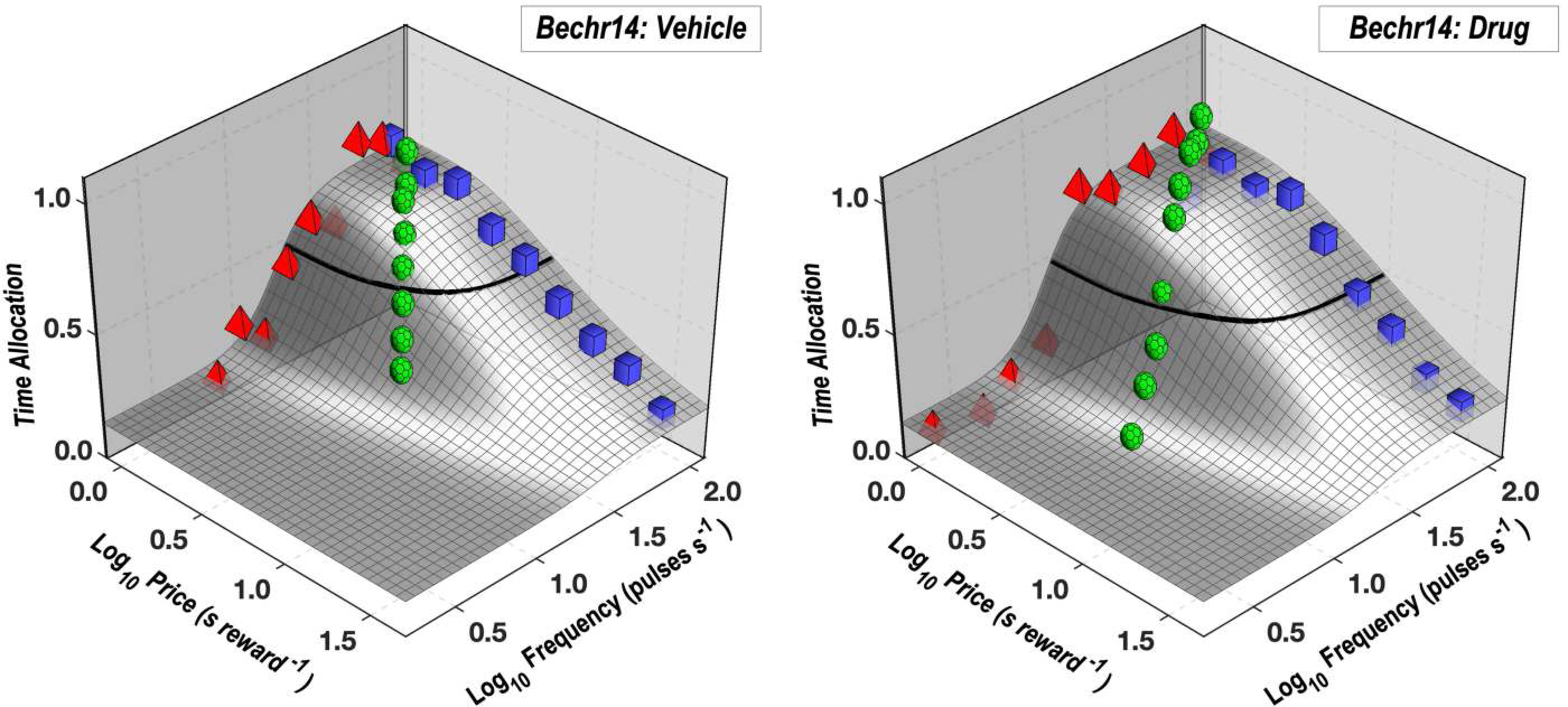
Reward-mountain surfaces fit to the vehicle and drug data from rat Bechr14. The surfaces of the reward-mountain shell are shown in gray. The thick black line represents the contour mid-way between the minimal and maximal estimates of time allocation (the estimated altitudes of the valley floor and summit). Mean time-allocation values for the pulse frequency, price, and radial sweeps are denoted by red pyramids, blue squares, and green polyhedrons, respectively.

**Fig S18.**
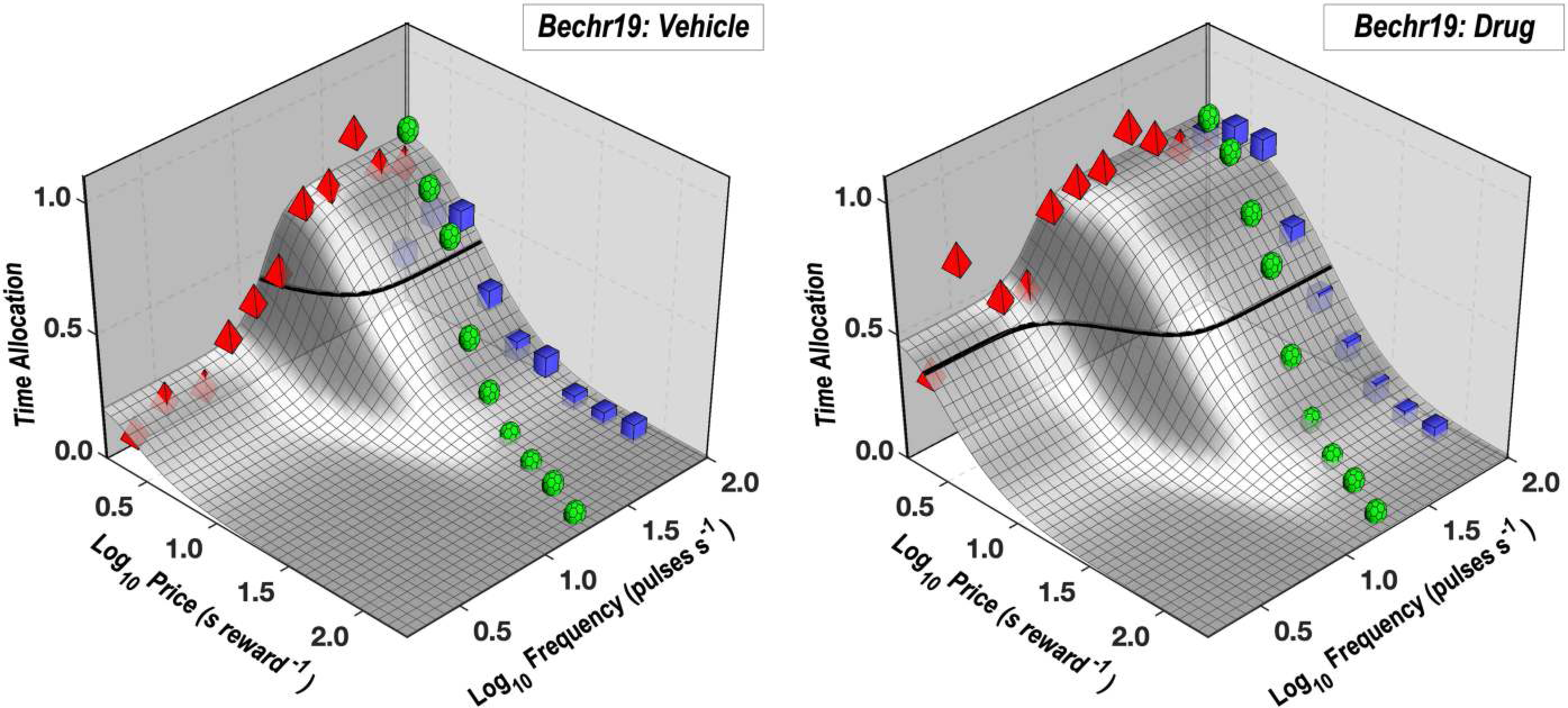
Reward-mountain surfaces fit to the vehicle and drug data from rat Bechr19. See caption for Fig S17.

**Fig S19.**
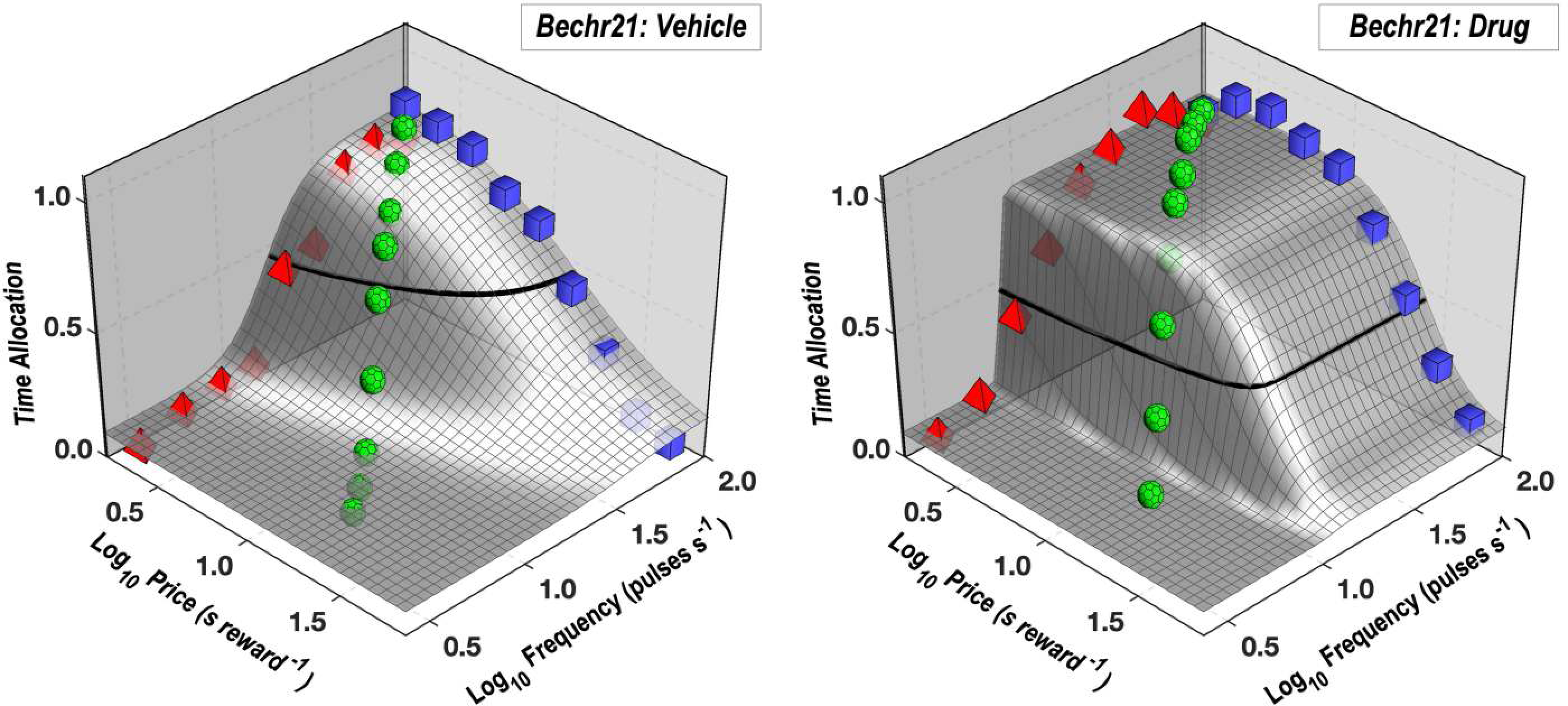
Reward-mountain surfaces fit to the vehicle and drug data from rat Bechr21. See caption for Fig S17.

**Fig S20.**
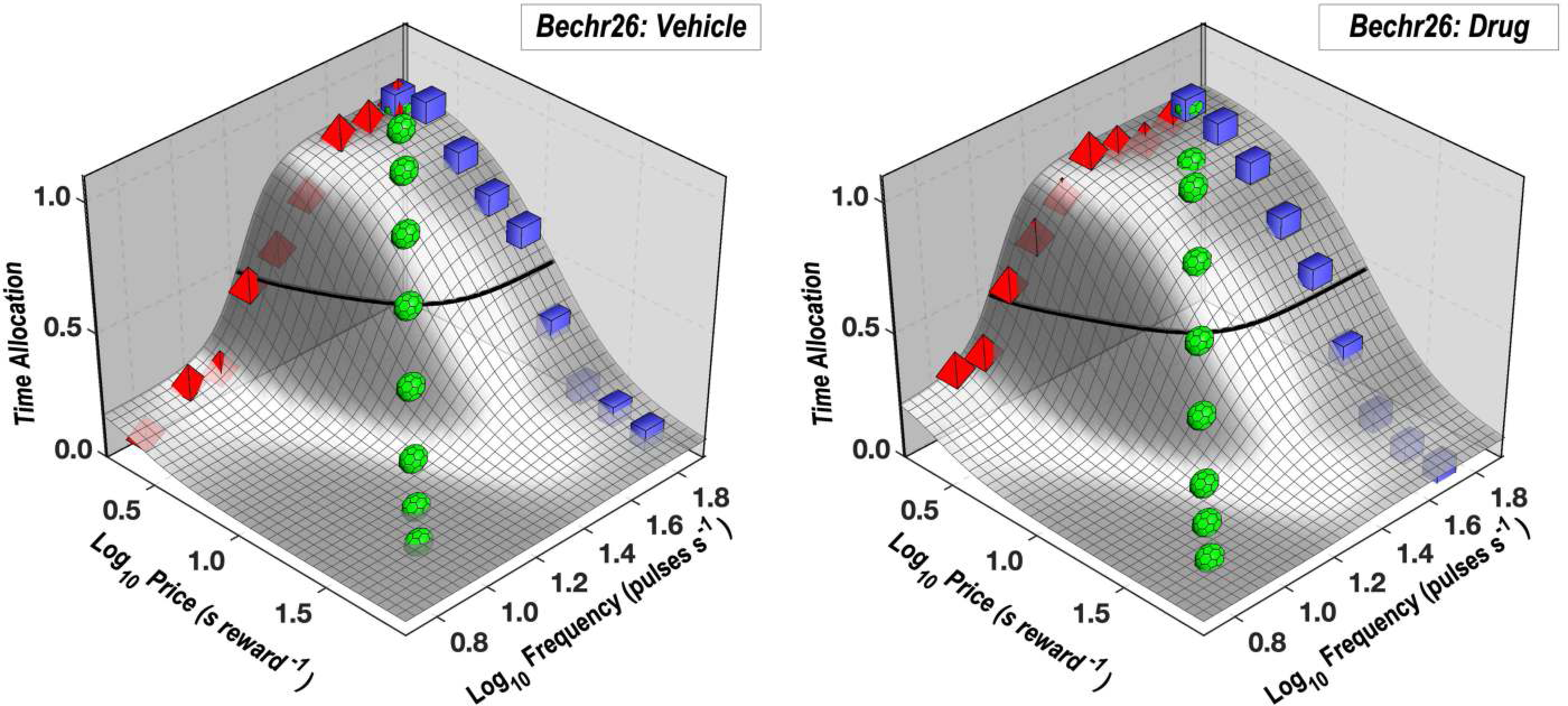
Reward-mountain surfaces fit to the vehicle and drug data from rat Bechr26. See caption for Fig S17.

**Fig S21.**
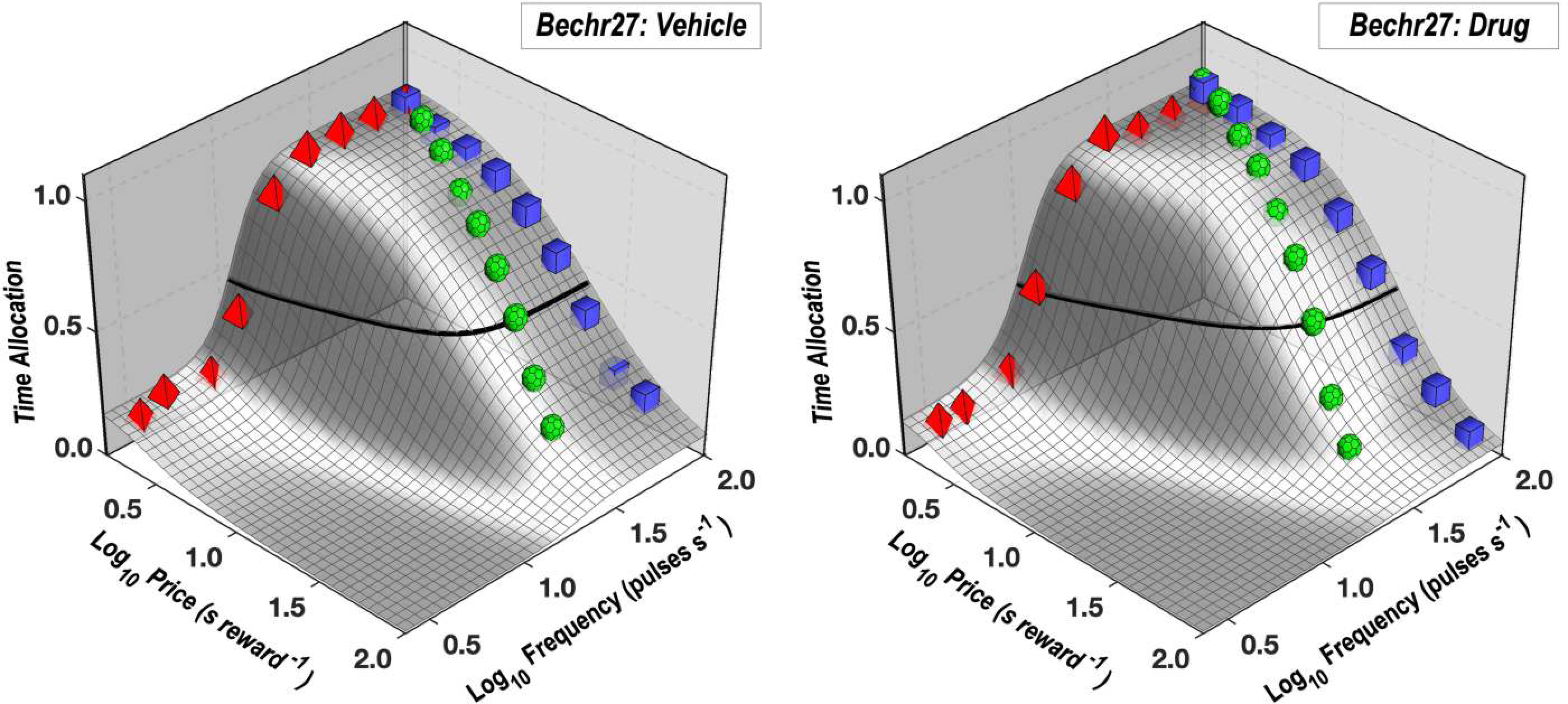
Reward-mountain surfaces fit to the vehicle and drug data from rat Bechr27. See caption for Fig S17.

**Fig S22.**
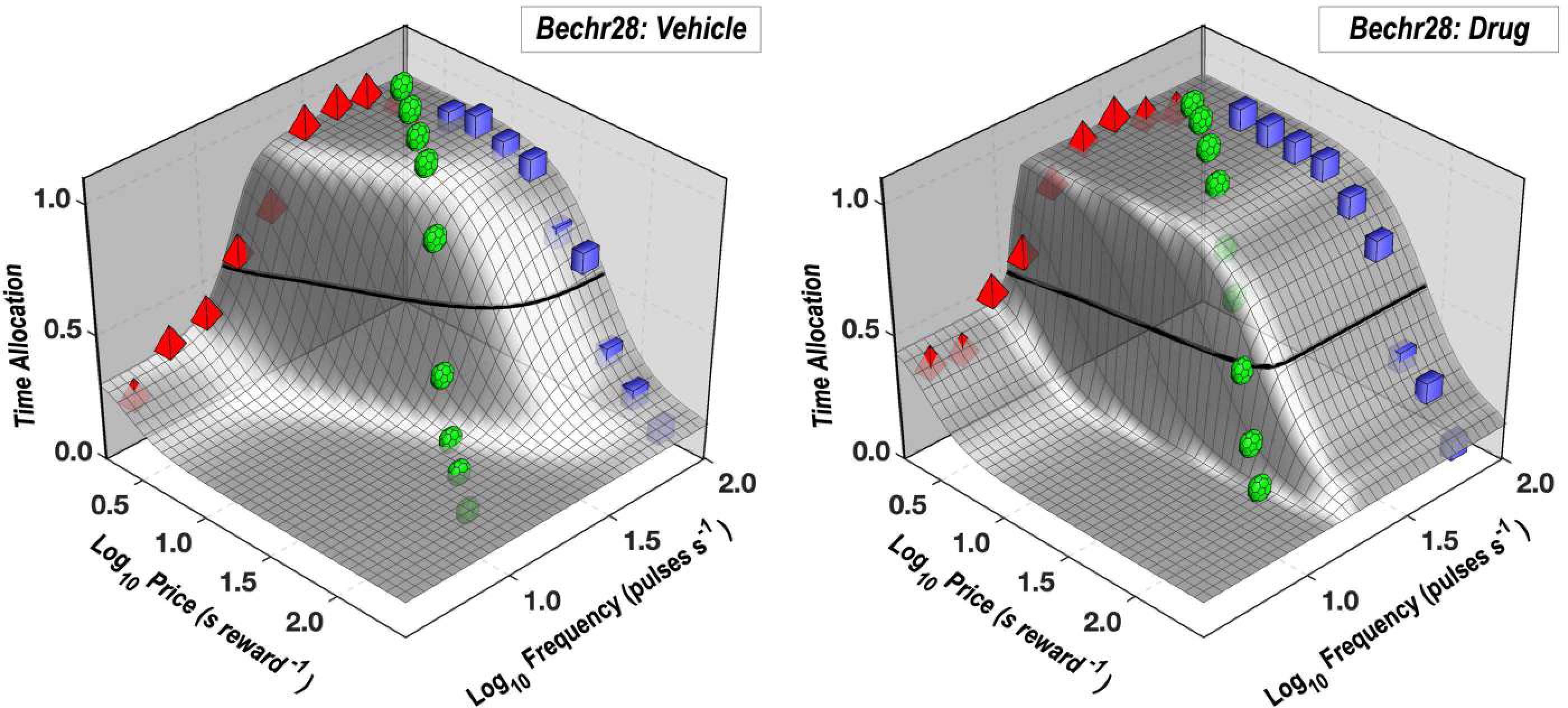
Reward-mountain surfaces fit to the vehicle and drug data from rat Bechr28. See caption for Fig S17.

The corresponding graph for rat Bechr29 is shown in Fig 5 in the main text along with the caption.

### Contour and bar graphs

**Fig S23.**
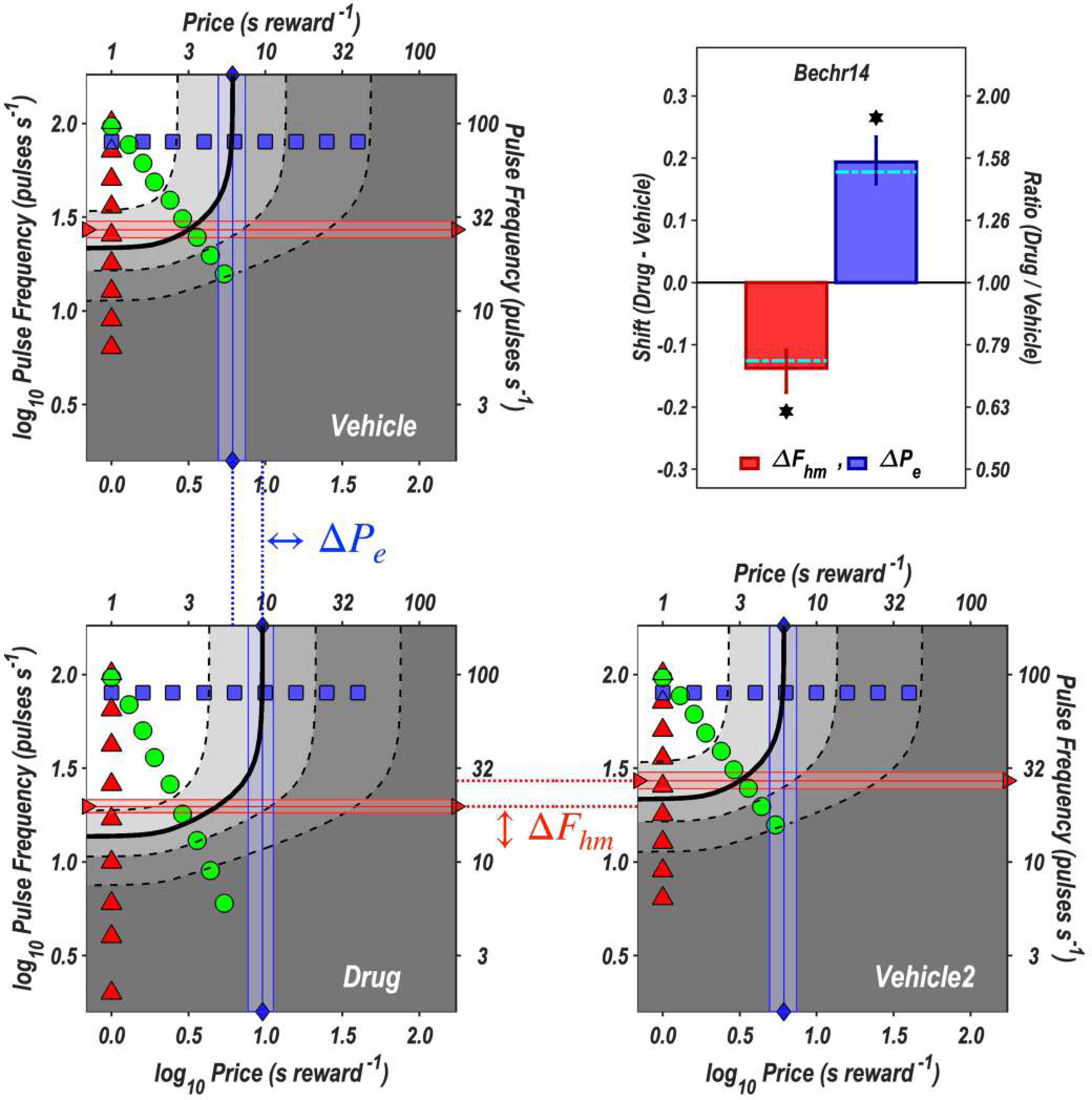
Contour graphs of the surfaces fit to the vehicle and drug data and bar graphs of the shifts in the location parameters for rat Bechr14. The values of the independent variables along frequency sweeps are designated by red triangles, along price sweeps by blue squares, and along radial sweeps by green circles. The values of the location parameters, 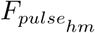 and 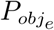, and are indicated by red horizontal lines with right-facing triangular end points and blue vertical lines with diamond end points, respectively. The shaded regions surrounding the lines denote 95% confidence intervals. The vehicle data are shown twice, once in the upper-left quadrant and once in the lower right. The dotted lines connecting the panels designate the shifts in the common-logarithmic values of the location parameters of the mountain, which are designated as 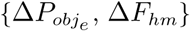 and plotted in the bar graph in the upper-right panel. The dot-dash cyan lines superimposed on the bars show location-parameter estimates corrected for changes in frequency-following fidelity due to the displacement of the mountain along the pulse-frequency axis. (See section ***Correction of the location-parameter estimates for changes in frequency-following fidelity***.) The 95% confidence intervals are shown in the bar graphs as vertical lines.

**Fig S24.**
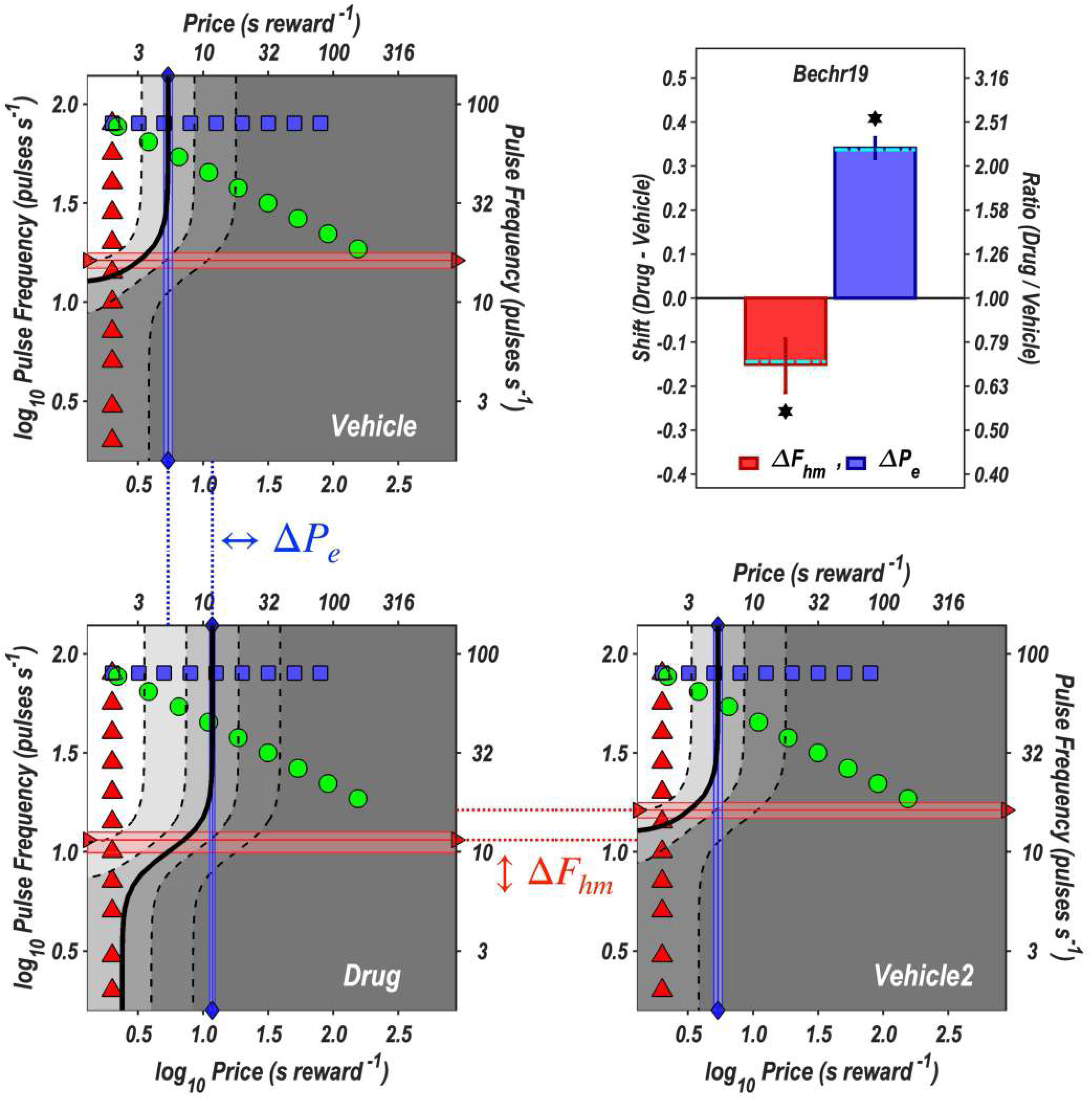
Contour and bar graphs for rat Bechr19. See caption for Fig S23.

**Fig S25.**
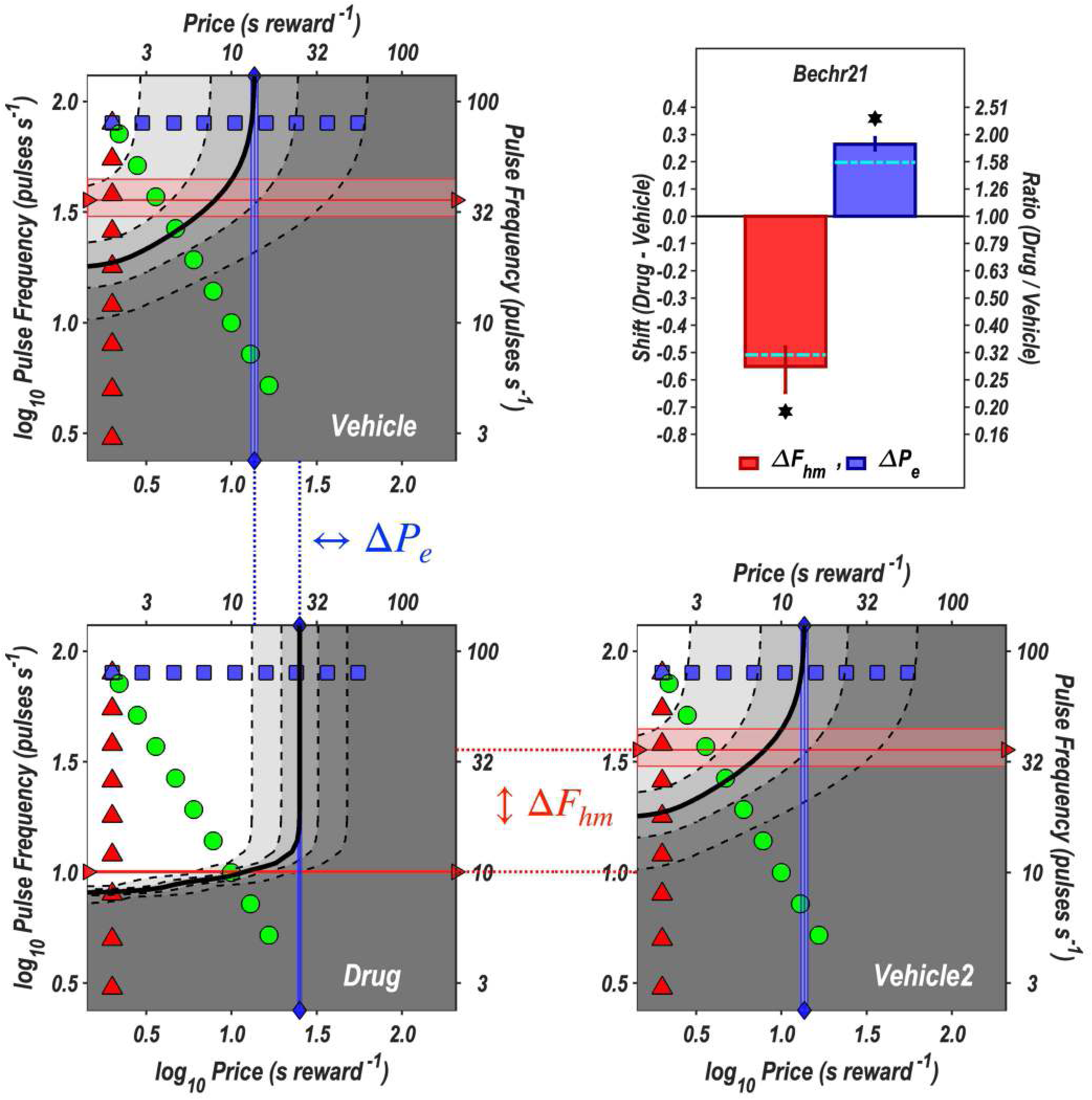
Contour and bar graphs for rat Bechr21. See caption for Fig S23.

**Fig S26.**
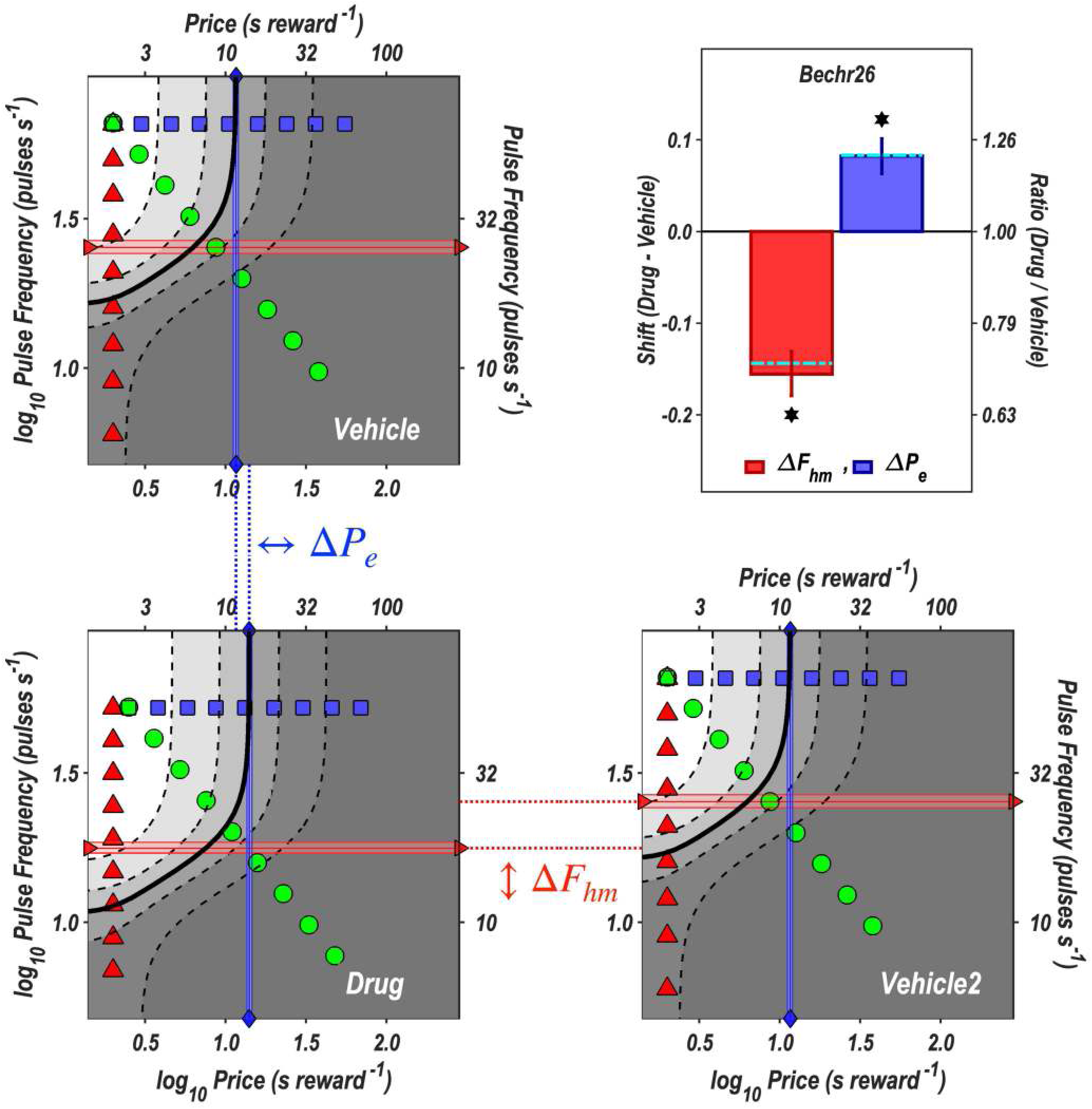
Contour and bar graphs for rat Bechr26. See caption for Fig S23.

**Fig S27.**
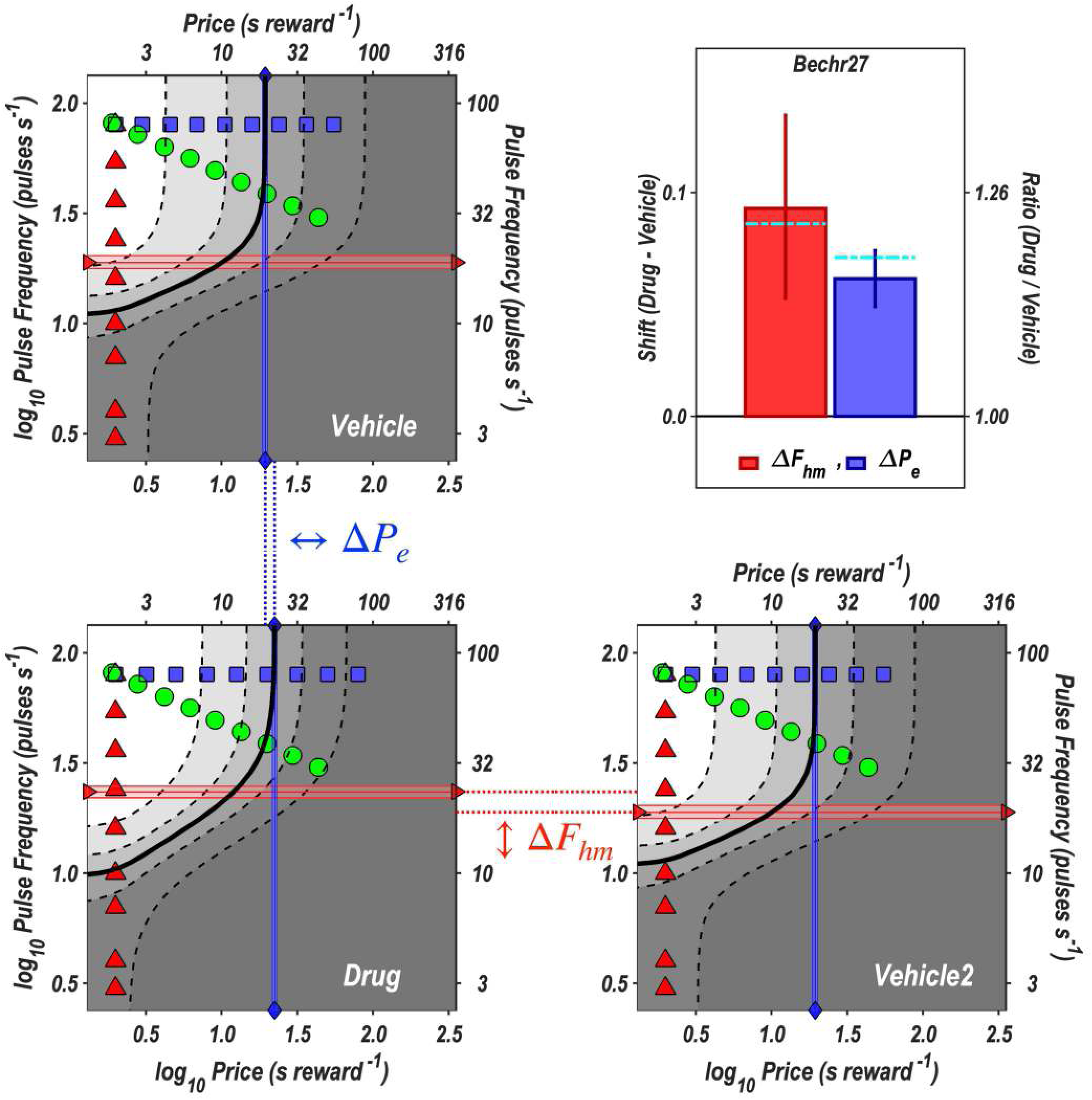
Contour and bar graphs for rat Bechr27. See caption for Fig S23.

**Fig S28.**
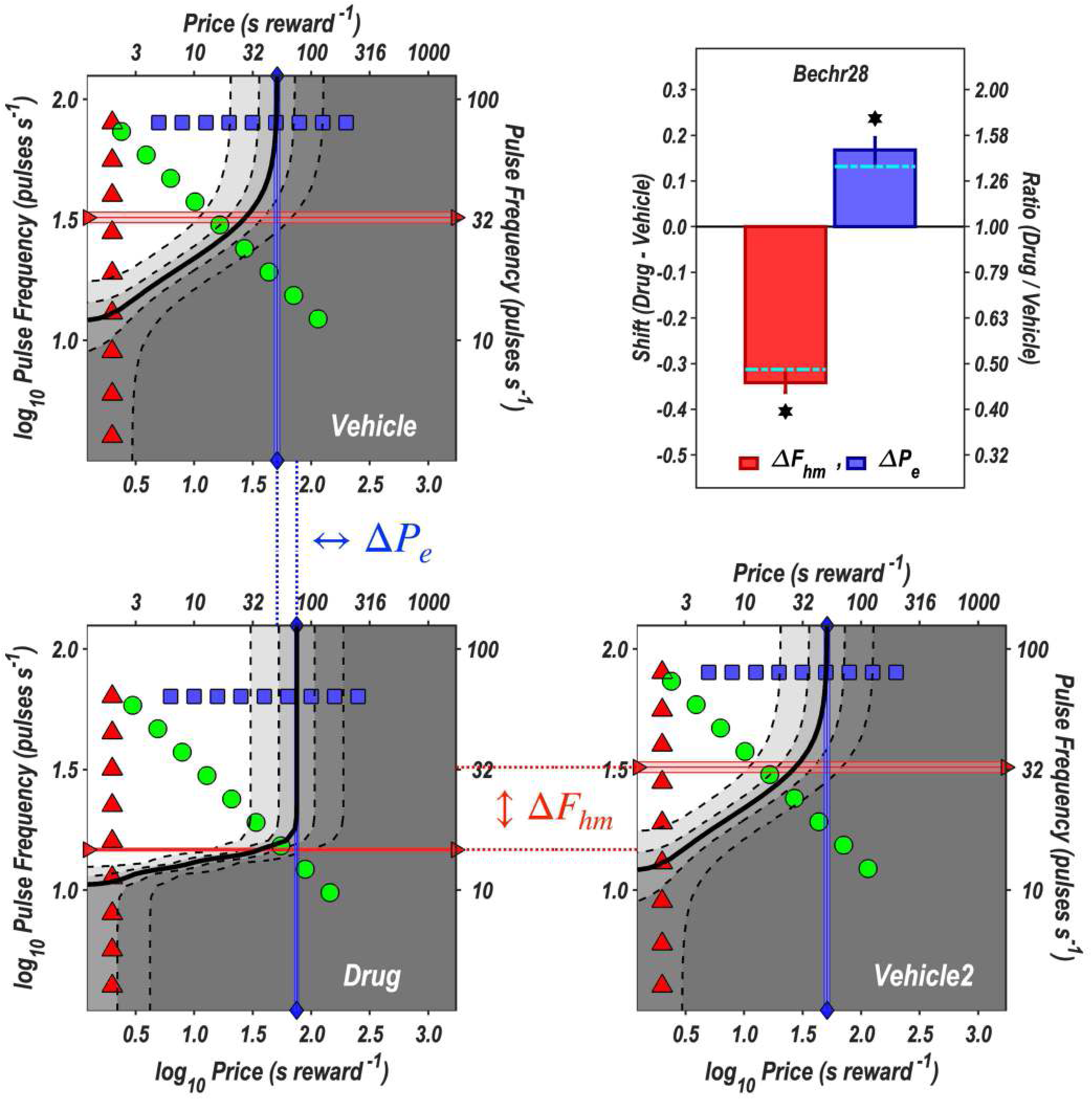
Contour and bar graphs for rat Bechr28. **See** caption for Fig S23.

The corresponding graphs for rat Bechr29 are shown in Fig 6 in the main text.

#### Location-parameter estimates

##### F_hm_

Tab S7 shows the estimates of 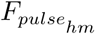 (uncorrected) and 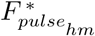 (corrected) for the drug and vehicle conditions. In six of seven cases, a lower pulse frequency sufficed to produce a reward of half-maximal intensity under the influence of dopamine-transporter blockade than in the vehicle condition.

The 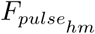 estimates vary over more than a doubling range in both the drug and vehicle conditions. Particularly in the vehicle condition, the higher values fall within a range over which the assumed frequency-following function (??) rolls off, thus preventing the normalized reward-growth function from approaching a value of one at the highest pulse frequencies tested. This is why the uncorrected 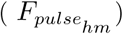 and corrected 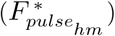 values in Tab S7) differ. For example, the ∼31 pulses s^−1^ that produced a reward of half-maximal intensity in rat Bechr29 in the vehicle condition are estimated to have generated only ∼26 firings s^−1^.

Tab 2 in the main text shows the estimated drug-induced ***shifts*** in the location of the reward-mountain core along the frequency axis.

Eqs S13,S14 express the effect of the drug on the location parameter of the reward-growth function as a divisor: the more the drug boosts dopamine release, the lower the value of the location parameter for the drug condition and thus the farther the reward-growth function is shifted to the left. The rightmost column in Tab 2 lists the values of this divisor implied by the drug-induced shifts in the position of the reward mountain along the pulse-frequency axis. By analogy to Eq S14,

**Table S7.** The position of the reward mountain along the pulse-frequency axis. The 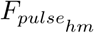 parameter sets the location of the shell of the reward-mountain along the pulse-frequency axis. If frequency-following fidelity were perfect, a pulse frequency of 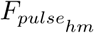 would have induced an identical firing frequency in the optically activated dopamine neurons. Otherwise, the values corrected for imperfect frequency following, which are shown in the 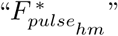 columns, will be lower than the uncorrected values, which are shown in the 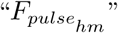 columns. The corrected values in the 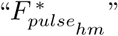 columns are the estimated firing frequencies induced by the pulse frequencies in the 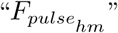 columns, derived from the frequency-following function described in section ??. The values listed in the “Veh” columns are from the fits to the data acquired in the vehicle condition, whereas the values in the “Drg” columns are from the fits to the data acquired under the influence of GBR-12909.

| Rat | $F_{pulse_{hm}}$ Drg | $F_{pulse_{hm}}$ Veh | $F_{pulse_{hm}}^*$ Drg | $F_{pulse_{hm}}^*$ Veh |
| --- | --- | --- | --- | --- |
| Bechr14 | 19.734 | 27.094 | 17.341 | 23.164 |
| Bechr19 | 11.502 | 16.265 | 10.372 | 14.457 |
| Bechr21 | 10.072 | 35.836 | 9.103 | 29.453 |
| Bechr26 | 17.718 | 25.359 | 15.668 | 21.822 |
| Bechr27 | 23.395 | 18.892 | 20.286 | 16.648 |
| Bechr28 | 14.728 | 32.350 | 13.142 | 27.007 |
| Bechr29 | 20.776 | 31.347 | 18.184 | 26.293 |

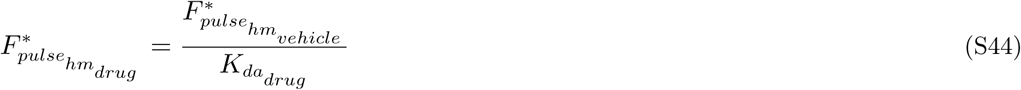

where

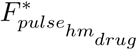 = location parameter of the reward-growth function for the drug condition

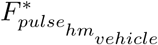 = location parameter of the reward-growth function for the vehicle condition

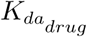 = proportional reduction in the value of the location parameter of the reward-growth function due to dopamine-transporter blockade

It follows that

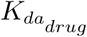 = 10^−diff^

where

diff = 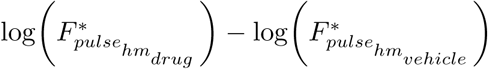

##### P_e_

In previous work employing the reward-mountain model [3, 33, 43, 44], changes in the location of the fitted surface along the price axis have been attributed to variables acting at, or beyond, the ***output*** of the reward-growth function, whereas changes in the location of the fitted surface along the pulse-frequency axis have been attributed to variables acting at, or prior to, the ***input*** to the reward-growth function. The reward-mountain model treats these two sets of changes as independent, a postulate that is largely supported by empirial findings [1, 2, 13]. This interpretation is valid as long as the induced firing frequency can be driven high enough to maximize reward intensity. Tab S8 shows that this assumption does not hold in several of the datasets from the present study: The maximum normalized reward intensity, 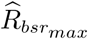, is substantially less than one in these cases.

The deviation of 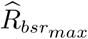 from one is generally greater in the vehicle data than in the drug data. In such cases, a portion of the change in the value of the 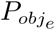 parameter is due to fact that the drug displaced the rising portion of the reward-mountain surface into a range of pulse frequencies over which the fidelity of frequency following is better than in the vehicle condition. That contribution to the change in the value of the 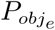 parameter reflects mitigation at the *input* to the reward-growth function (Eq S11), thus undermining the independence of the changes in the parameters that locate the reward mountain along the price and frequency axes. That is why we computed corrected estimates 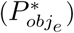, as described in section ***Correction of the location-parameter estimates for changes in frequency-following fidelity*** thus compensating for the differences in the value of 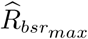 across the vehicle and drug conditions. The estimates of 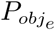 and 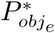 are shown in Tab S9.

**Table S8.** Estimates of the maximum normalized reward intensities in the vehicle and drug conditions. The values in 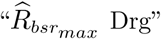 and 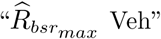 columns are the maximum normalized reward intensities in the vehicle and drug conditions, respectively, based on the frequency-following function described in section ??. The ratios of these values (Drg / Veh) are listed in the “Ratio” column, and the common logarithms of the ratios in the “log(Ratio)” column.

| Rat | $\hat{R}_{bsr_{max}}$ Drg | $\hat{R}_{bsr_{max}}$ Veh | Ratio | log(Ratio) |
| --- | --- | --- | --- | --- |
| Bechr14 | 0.986 | 0.958 | 1.029 | 0.013 |
| Bechr19 | 1.000 | 0.999 | 1.001 | 0.000 |
| Bechr21 | 1.000 | 0.849 | 1.179 | 0.071 |
| Bechr26 | 0.998 | 0.989 | 1.009 | 0.004 |
| Bechr27 | 0.964 | 0.997 | 0.967 | -0.015 |
| Bechr28 | 1.000 | 0.952 | 1.051 | 0.022 |
| Bechr29 | 0.964 | 0.894 | 1.079 | 0.033 |

**Table S9.**
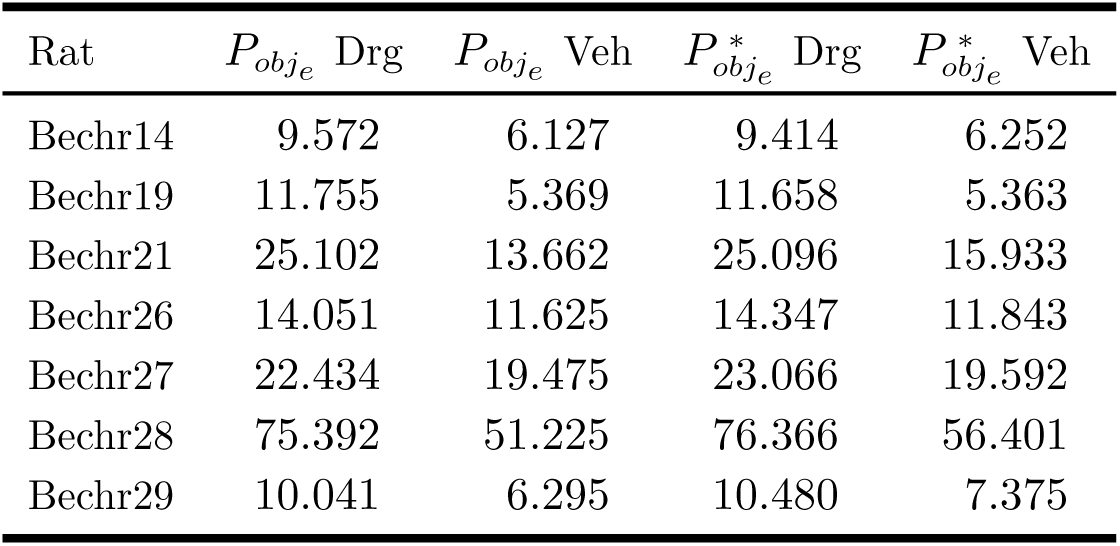
The position of the reward mountain along the price axis. The 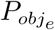 parameter determines the position of the reward mountain along the price axis. When the maximum normalized reward intensity differs between the drug and vehicle conditions due to differences in frequency-following fidelity, the value of the 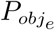 parameter is affected. The values listed in the 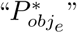 columns have been corrected to remove this effect. They show the estimated value that the 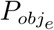 parameter would have attained had frequency-following fidelity been perfect.

| Rat | $P_{obj_e}$ Drg | $P_{obj_e}$ Veh | $P_{obj_e}^*$ Drg | $P_{obj_e}^*$ Veh |
| --- | --- | --- | --- | --- |
| Bechr14 | 9.572 | 6.127 | 9.414 | 6.252 |
| Bechr19 | 11.755 | 5.369 | 11.658 | 5.363 |
| Bechr21 | 25.102 | 13.662 | 25.096 | 15.933 |
| Bechr26 | 14.051 | 11.625 | 14.347 | 11.843 |
| Bechr27 | 22.434 | 19.475 | 23.066 | 19.592 |
| Bechr28 | 75.392 | 51.225 | 76.366 | 56.401 |
| Bechr29 | 10.041 | 6.295 | 10.480 | 7.375 |

Tab 3 in the main text lists the drug-induced shifts in the common logarithms of 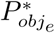.

### The drug-induced shifts in the location-parameter values are uncorrelated

#### Parameter values for best-fitting model for all rats

The values of the location parameters in these tables are uncorrected. Corrected values are listed in Tabs 2 and 3 in the main text.

**Fig S29.**
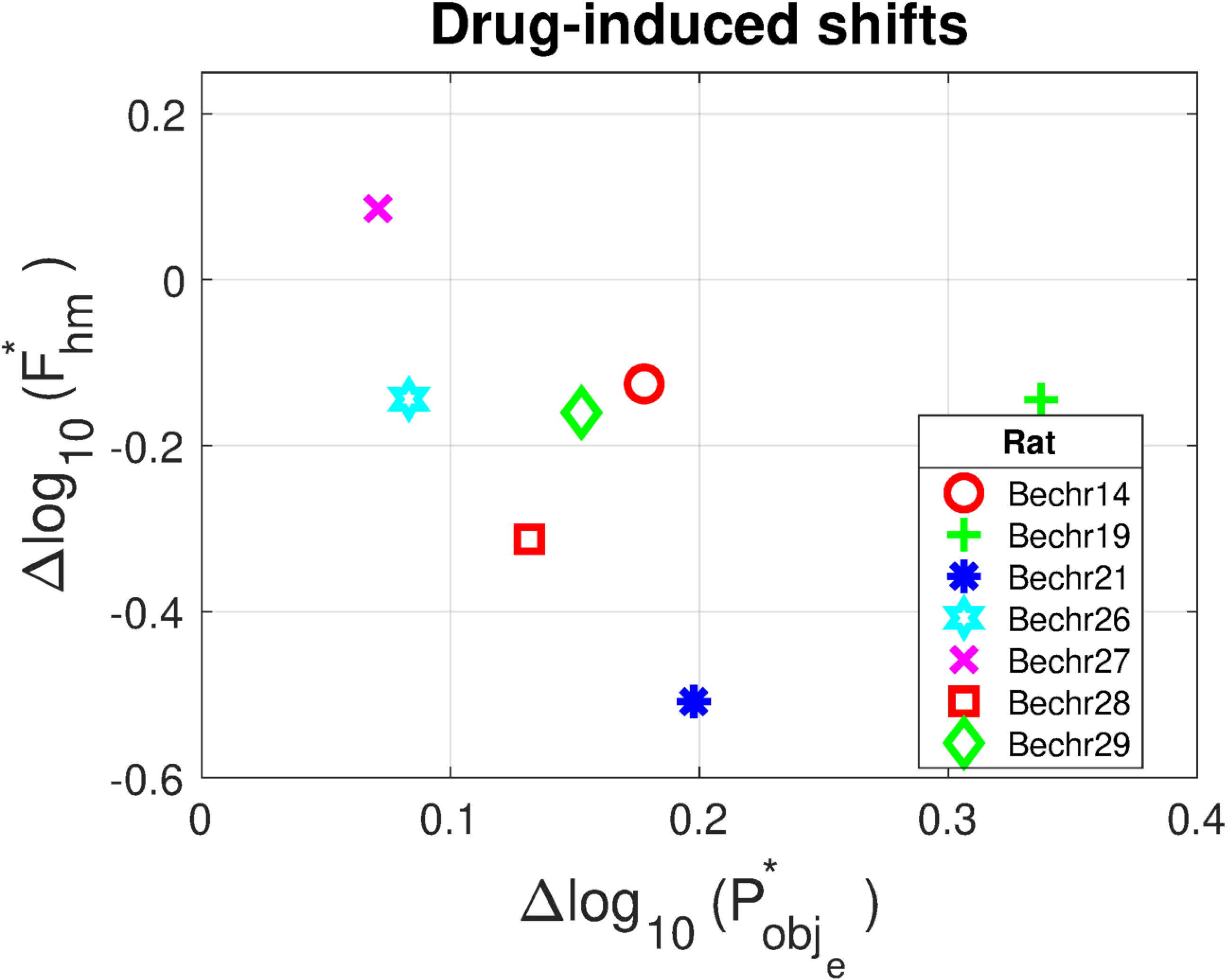
Scatter plot of drug-induced shifts in the location parameters. The corrected estimates of displacement along the price and pulse-frequency axes are shown on the abscissa and ordinate, respectively. 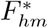 is shorthand for 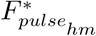

**Table S10.** Parameter values from the best-fitting model for the data from Rat Bechr14. Columns *CB*_*low*_ and *CB*_*high*_ list the upper and lower bounds of the 95.2% confidence intervals.

| Parameter | Vehicle |  |  | Drug |  |  |
| --- | --- | --- | --- | --- | --- | --- |
| | Estimate | $CB_{low}$ | $CB_{high}$ | Estimate | $CB_{low}$ | $CB_{high}$ |
| $a$ | 1.064 | 0.922 | 1.233 | 1.064 | 0.922 | 1.233 |
| $g$ | 3.915 | 3.450 | 4.423 | 3.915 | 3.450 | 4.423 |
| $Log_{10}(F_{hm})$ | 1.433 | 1.390 | 1.478 | 1.295 | 1.262 | 1.331 |
| $Log_{10}(P_e)$ | 0.787 | 0.694 | 0.869 | 0.981 | 0.889 | 1.052 |
| $T_{max}$ | 0.821 | 0.771 | 0.881 | 0.821 | 0.771 | 0.881 |
| $T_{min}$ | 0.144 | 0.127 | 0.159 | 0.144 | 0.127 | 0.159 |

**Table S11.** Parameter values from the best-fitting model for the data from Rat Bechr19. Columns *CB*_*low*_ and *CB*_*high*_ list the upper and lower bounds of the 95.2% confidence intervals.

| Parameter | Vehicle |  |  | Drug |  |  |
| --- | --- | --- | --- | --- | --- | --- |
| | Estimate | $CB_{low}$ | $CB_{high}$ | Estimate | $CB_{low}$ | $CB_{high}$ |
| $a$ | 1.840 | 1.729 | 1.972 | 1.840 | 1.729 | 1.972 |
| $C_r$ | 0.214 | 0.191 | 0.239 | 0.214 | 0.191 | 0.239 |
| $g$ | 5.862 | 3.837 | 13.690 | 5.862 | 3.837 | 13.690 |
| $Log_{10}(F_{hm})$ | 1.211 | 1.172 | 1.247 | 1.061 | 0.996 | 1.100 |
| $Log_{10}(P_e)$ | 0.730 | 0.699 | 0.762 | 1.070 | 1.051 | 1.089 |
| $T_{max}$ | 0.796 | 0.777 | 0.816 | 0.796 | 0.777 | 0.816 |
| $T_{min}$ | 0.005 | 0.000 | 0.009 | 0.005 | 0.000 | 0.009 |

**Table S12.** Parameter values from the best-fitting model for the data from Rat Bechr21. Columns *CB*_*low*_ and *CB*_*high*_ list the upper and lower bounds of the 95.2% confidence intervals.

| Parameter | Vehicle |  |  | Drug |  |  |
| --- | --- | --- | --- | --- | --- | --- |
| | Estimate | $CB_{low}$ | $CB_{high}$ | Estimate | $CB_{low}$ | $CB_{high}$ |
| $a$ | 1.442 | 1.368 | 1.523 | 3.436 | 3.222 | 3.674 |
| $g$ | 3.109 | 2.719 | 3.593 | 10.992 | 10.503 | 11.432 |
| $Log_{10}(F_{hm})$ | 1.554 | 1.480 | 1.649 | 1.003 | 1.000 | 1.007 |
| $Log_{10}(P_e)$ | 1.136 | 1.115 | 1.154 | 1.400 | 1.388 | 1.410 |
| $T_{max}$ | 0.836 | 0.829 | 0.843 | 0.836 | 0.829 | 0.843 |
| $T_{min}$ | 0.097 | 0.091 | 0.103 | 0.097 | 0.091 | 0.103 |

**Table S13.**
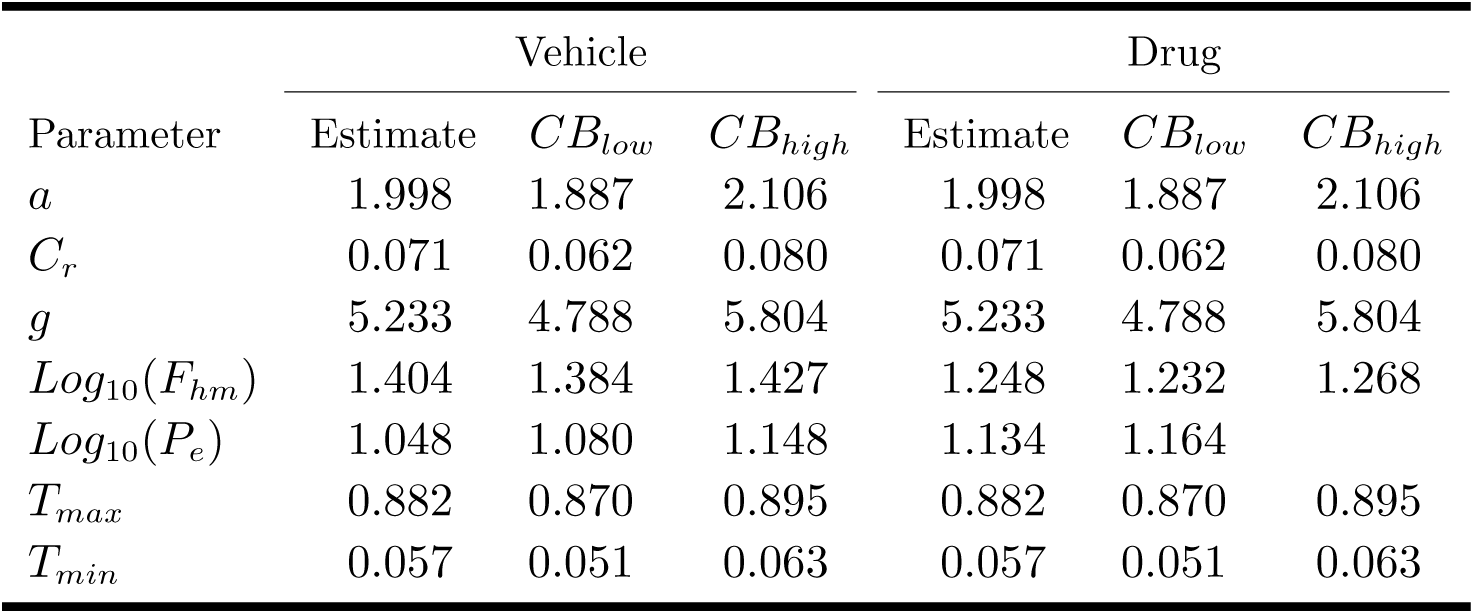
Parameter values from the best-fitting model for the data from Rat Bechr26. Columns *CB*_*low*_ and *CB*_*high*_ list the upper and lower bounds of the 95.2% confidence intervals.

**Table S14.** Parameter values from the best-fitting model for the data from Rat Bechr27. Columns *CB*_*low*_ and *CB*_*high*_ list the upper and lower bounds of the 95.2% confidence intervals.

| Parameter | Vehicle |  |  | Drug |  |  |
| --- | --- | --- | --- | --- | --- | --- |
| | Estimate | $CB_{low}$ | $CB_{high}$ | Estimate | $CB_{low}$ | $CB_{high}$ |
| $a$ | 1.454 | 1.381 | 1.525 | 2.015 | 1.924 | 2.135 |
| $C_r$ | 0.037 | 0.035 | 0.039 | 0.037 | 0.035 | 0.039 |
| $g$ | 5.050 | 4.561 | 5.684 | 3.524 | 3.360 | 3.699 |
| $Log_{10}(F_{hm})$ | 1.276 | 1.248 | 1.307 | 1.369 | 1.342 | 1.394 |
| $Log_{10}(P_e)$ | 1.289 | 1.272 | 1.303 | 1.351 | 1.337 | 1.364 |
| $T_{max}$ | 0.888 | 0.881 | 0.895 | 0.888 | 0.881 | 0.895 |
| $T_{min}$ | 0.009 | 0.000 | 0.028 | 0.009 | 0.000 | 0.028 |

**Table S15.** Parameter values from the best-fitting model for the data from Rat Bechr28. Columns *CB*_*low*_ and *CB*_*high*_ list the upper and lower bounds of the 95.2% confidence intervals.

| Parameter | Vehicle |  |  | Drug |  |  |
| --- | --- | --- | --- | --- | --- | --- |
| | Estimate | $CB_{low}$ | $CB_{high}$ | Estimate | $CB_{low}$ | $CB_{high}$ |
| $a$ | 2.432 | 2.304 | 2.595 | 2.432 | 2.304 | 2.595 |
| $C_r$ | 0.022 | 0.021 | 0.024 | 0.022 | 0.021 | 0.024 |
| $g$ | 4.609 | 4.281 | 5.077 | 17.305 | 16.182 | 18.765 |
| $Log_{10}(F_{hm})$ | 1.510 | 1.489 | 1.532 | 1.168 | 1.163 | 1.173 |
| $Log_{10}(P_e)$ | 1.709 | 1.687 | 1.730 | 1.877 | 1.860 | 1.893 |
| $T_{max}$ | 0.915 | 0.910 | 0.921 | 0.915 | 0.910 | 0.921 |
| $T_{min}$ | 0.143 | 0.135 | 0.151 | 0.143 | 0.135 | 0.151 |

**Table S16.** Parameter values from the best-fitting model for the data from Rat Bechr29. Columns *CB*_*low*_ and *CB*_*high*_ list the upper and lower bounds of the 95.2% confidence intervals.

| Parameter | Vehicle |  |  | Drug |  |  |
| --- | --- | --- | --- | --- | --- | --- |
| | Estimate | $CB_{low}$ | $CB_{high}$ | Estimate | $CB_{low}$ | $CB_{high}$ |
| $a$ | 2.312 | 2.172 | 2.462 | 2.312 | 2.172 | 2.462 |
| $g$ | 3.161 | 2.980 | 3.354 | 3.161 | 2.980 | 3.354 |
| $Log_{10}(F_{hm})$ | 1.496 | 1.476 | 1.518 | 1.318 | 1.304 | 1.335 |
| $Log_{10}(P_e)$ | 0.799 | 0.783 | 0.814 | 1.002 | 0.990 | 1.015 |
| $T_{max}$ | 0.896 | 0.883 | 0.910 | 0.896 | 0.883 | 0.910 |
| $T_{min}$ | 0.112 | 0.107 | 0.118 | 0.112 | 0.107 | 0.118 |

### The reward-growth function for oICSS

**Fig S30.**
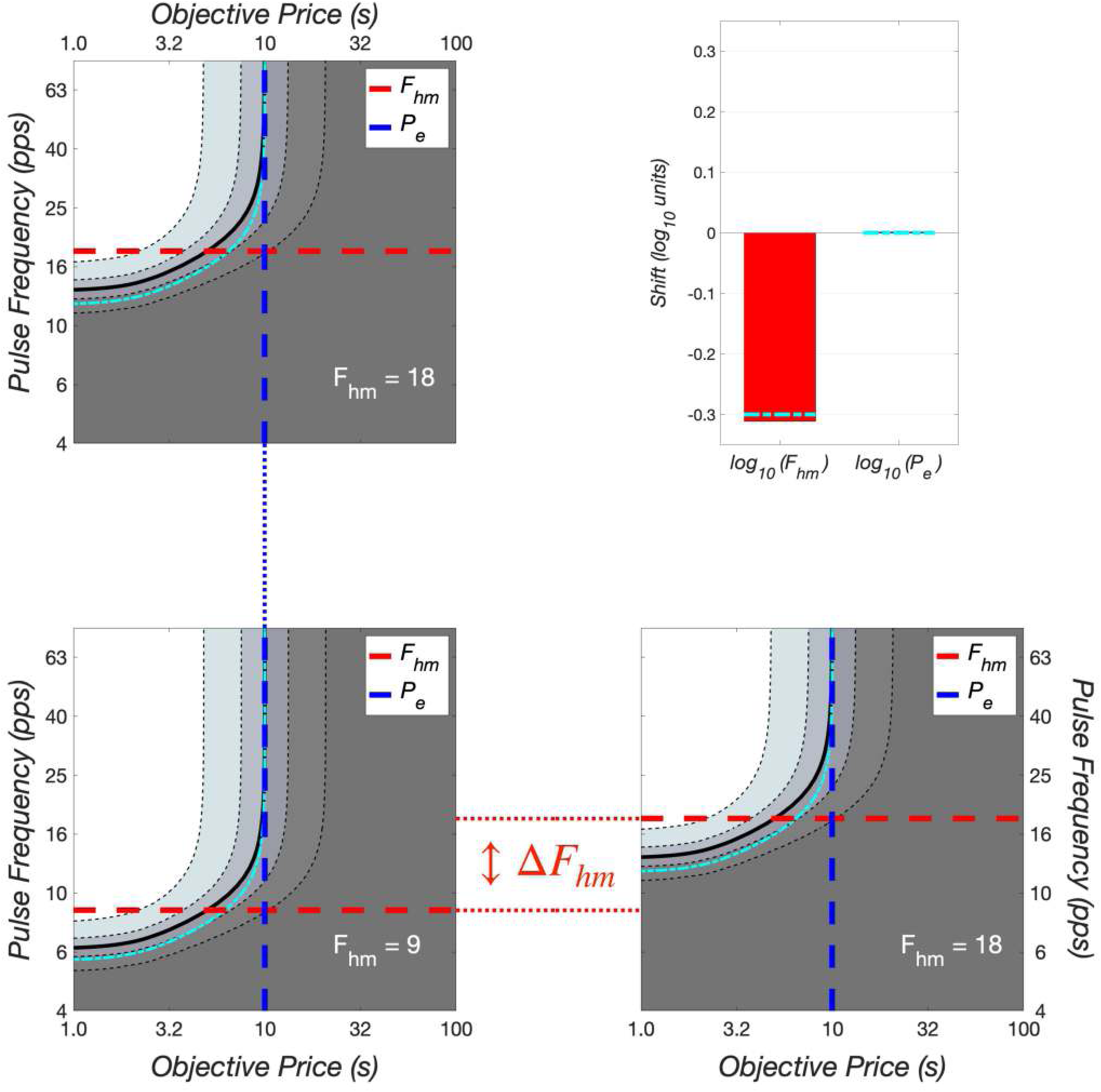
Changing the value of the input-scaling parameter, *F*_*hm*_, shifts the mountain along the pulse-frequency axis. *F*_*hm*_ is shorthand for 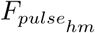

**Fig S31.**
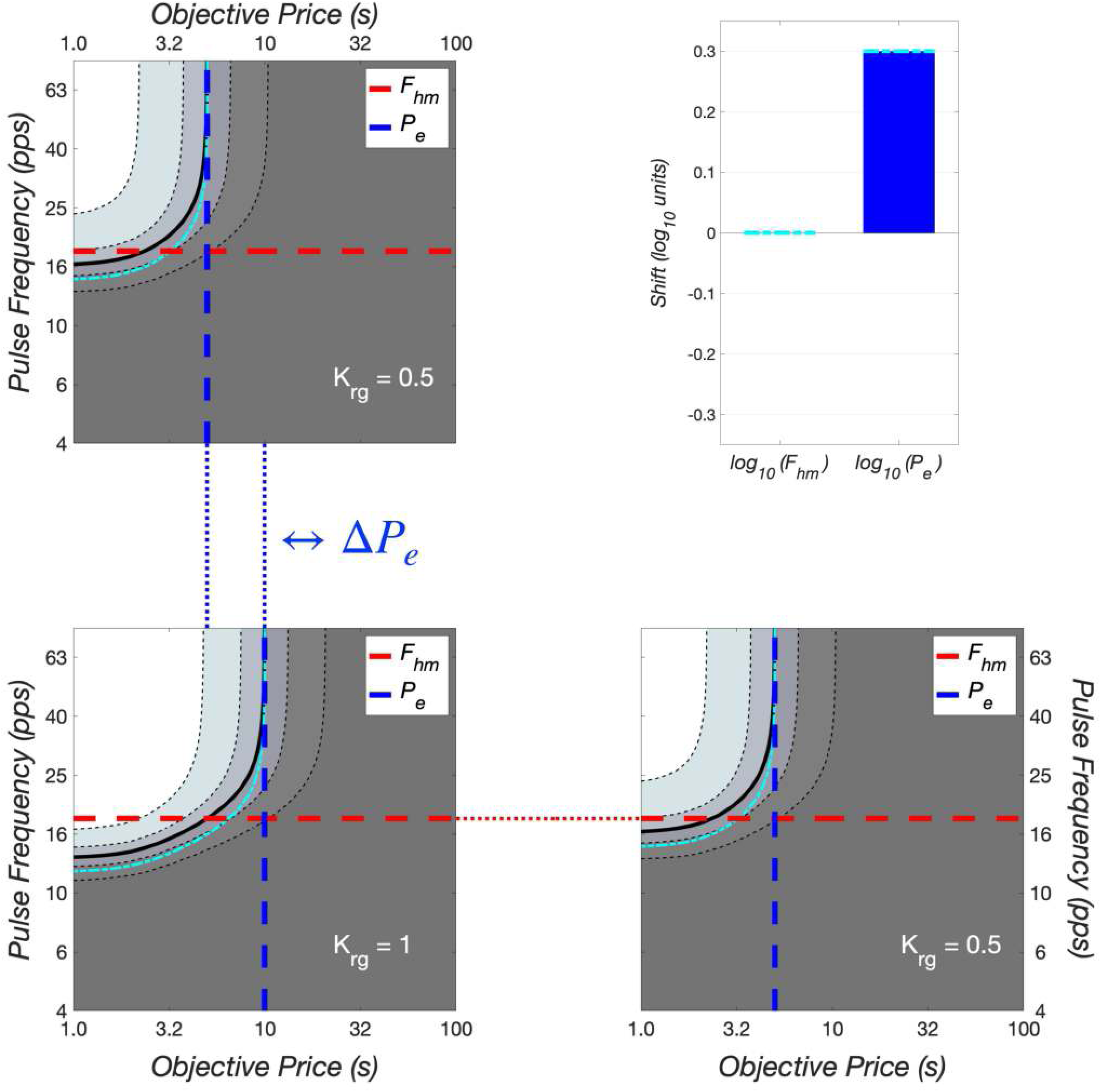
Changing the value of the output-scaling parameter, *K*_*rg*_, shifts the mountain along the price axis. *P*_*e*_ is shorthand for 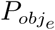

### Illustration of the correction for imperfect frequency-following fidelity

When the parameter that locates the reward mountain along the pulse-frequency axis 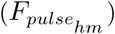 falls within the range over which the induced firing frequency diverges substantially from the pulse frequency, then the magnitude of a drug-induced shift along the pulse-frequency axis is exaggerated, and a fictive shift is produced along the price axis. Fig S32 illustrates this effect and its removal by means of the correction procedure. The dot-dash cyan contour line shows the objective-price and pulse-frequency values that would have driven time allocation halfway between its minimal and maximal values had frequency-following fidelity been perfect. In contrast, the solid, wide, black contour line shows the equivalent objective-price and pulse-frequency values given the assumed frequency-following function. Note that the dot-dash cyan line deviates more from the solid, wide, black line in the simulated vehicle data in the upper left and lower right quadrants than in the simulated drug data in the lower-left quadrant. This is so because the uncorrected location-parameter value (vehicle: ∼ 39 pulses s^−1^; drug: ∼ 18 pulses s^−1^) is much closer to the estimated maximum attainable firing frequency (51.67 firings s^−1^) in the vehicle condition than in the drug condition. As a result, the simulated firing rate falls further below the pulse frequency in the vehicle condition than in the drug condition. The dot-dash lines superimposed on the bar graphs show the result of correcting the location-parameter shifts for this effect.

**Fig S32.**
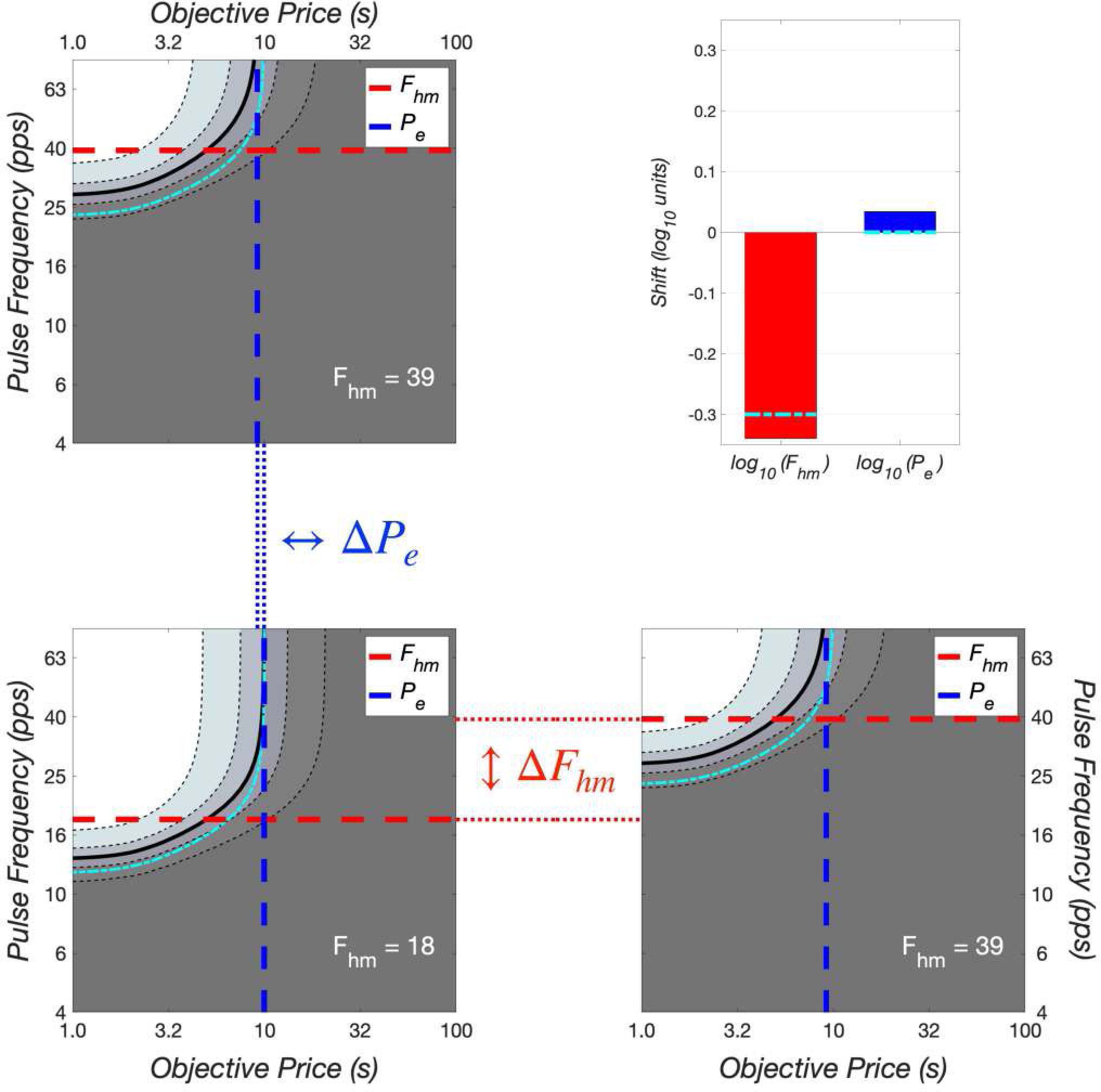
Correction for imperfect frequency-following fidelity. Simulated data, with 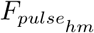 placed well within the region over which frequency-following fidelity falls off substantially. The simualated vehicle data are shown twice, once in the upper-left quadrant and once in the lower right. The dotted lines connecting the panels designate the shifts in the common-logarithmic values of the location parameters of the mountain, which are designated as 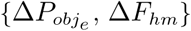 and plotted in the bar graph in the upper-right panel. The dot-dash cyan lines superimposed on the bars show location-parameter estimates corrected for changes in frequency-following fidelity due to the displacement of the mountain along the pulse-frequency axis. *F*_*hm*_ is shorthand for 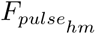

### Comparison of logistic and power growth of reward intensity

**Fig S33.**
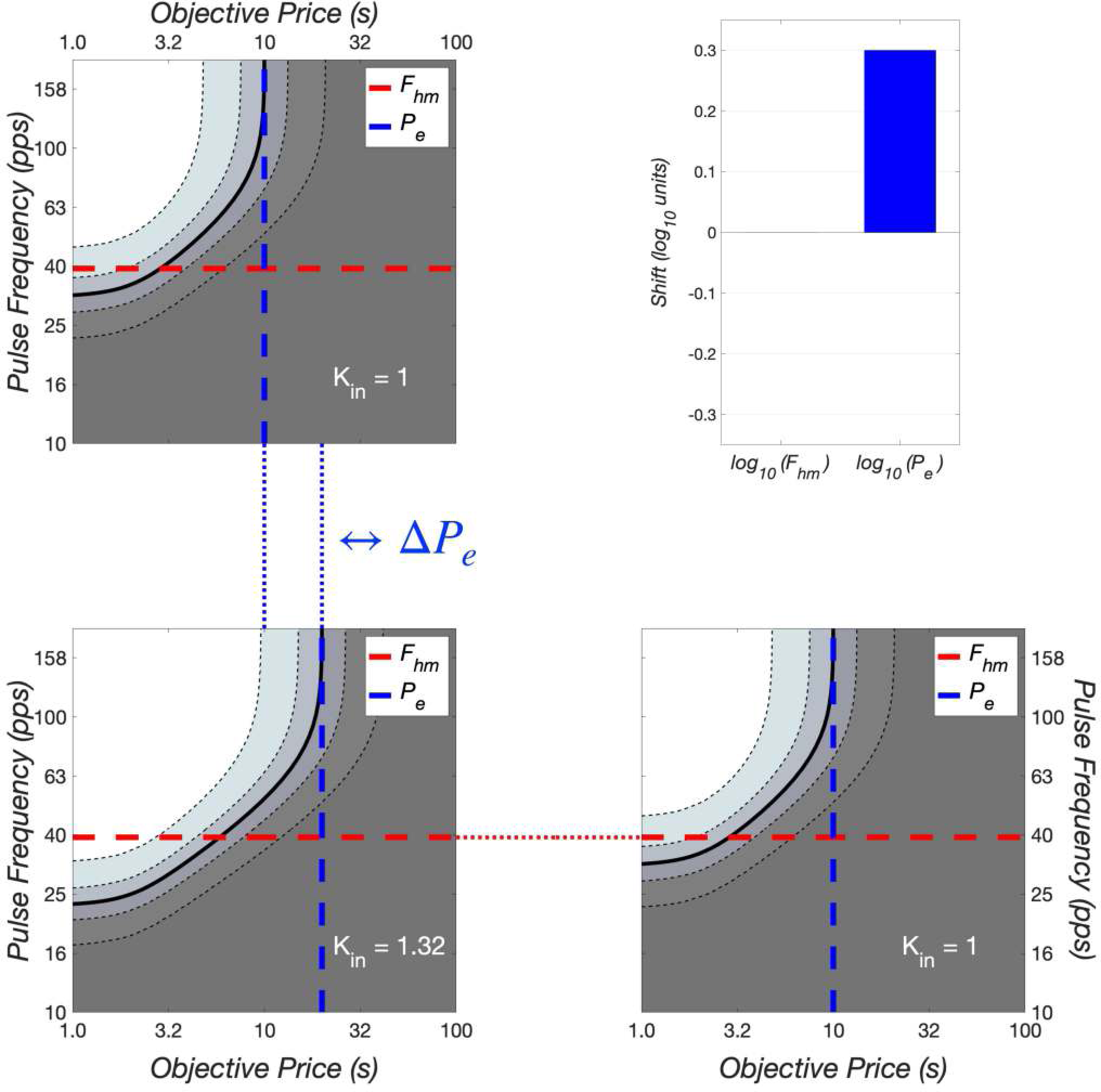
The input-scaling parameter of the power reward-growth function locates the reward mountain along the price axis. Contour- and bar-graph representation of the simulated reward mountains produced by the magenta and green power-reward-growth functions (Eq 2) in the lower-left panel of Fig 9. In contrast to the effect of varying the value of the input-scaling parameter on the location of reward mountains based on logistic reward growth (Figs S30, S31), changing the value of the input-scaling parameter of the power-reward-growth function shifts the mountain along the price axis and not along the pulse-frequency axis.

**Fig S34.**
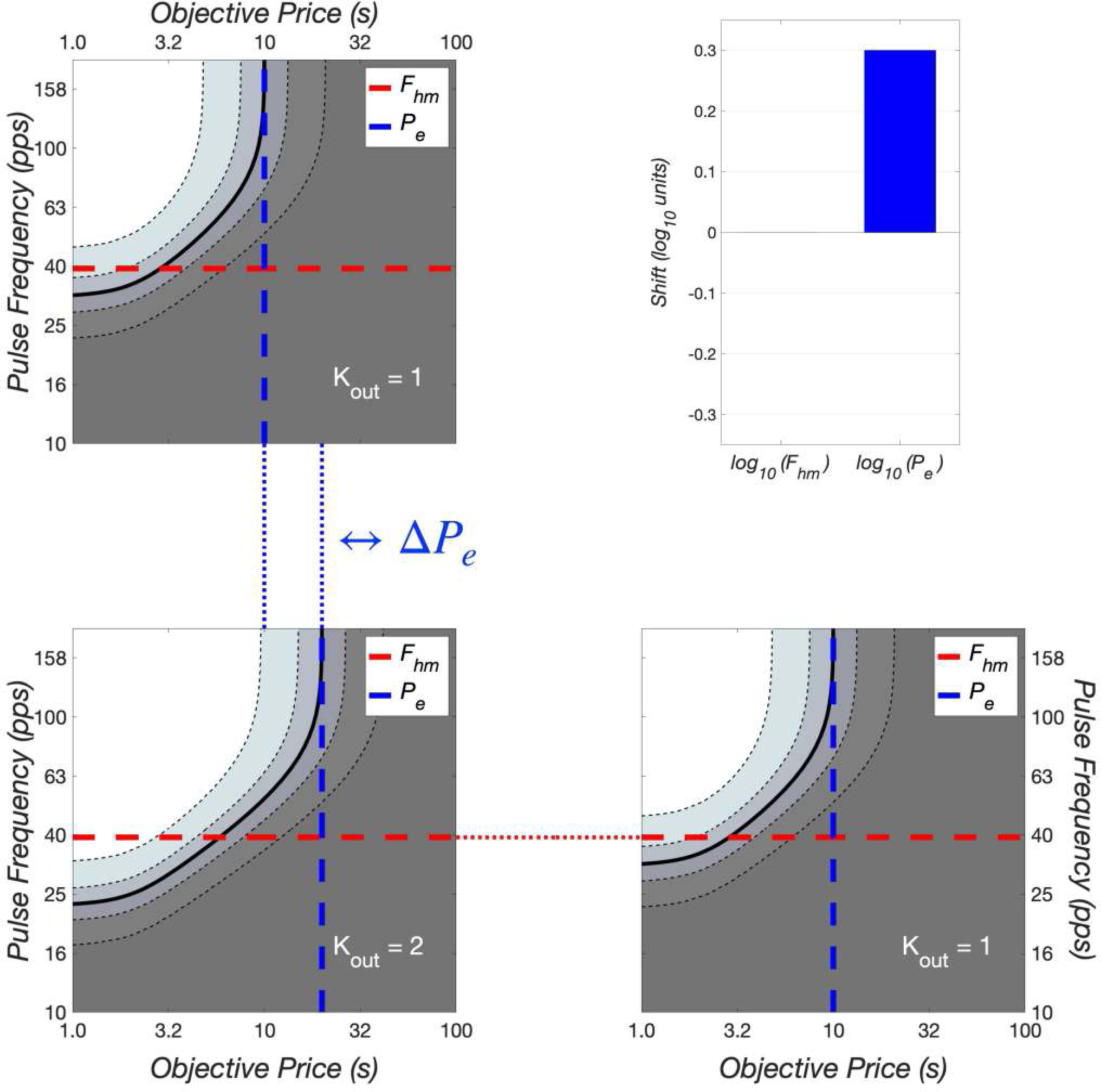
The output-scaling parameter of the power reward-growth function also locates the reward mountain along the price axis. Contour- and bar-graph representation of the simulated reward mountains produced by the magenta and green power-reward-growth functions (Eq 2) in the lower-right panel of Fig 9. Changing the value of the output-scaling parameter of the power-reward-growth function shifts the mountain along the price axis just like the effect of changing the value of the input-scaling parameter shown in Fig S33.

### Toward a new model of brain-reward circuitry

**Fig S35.**
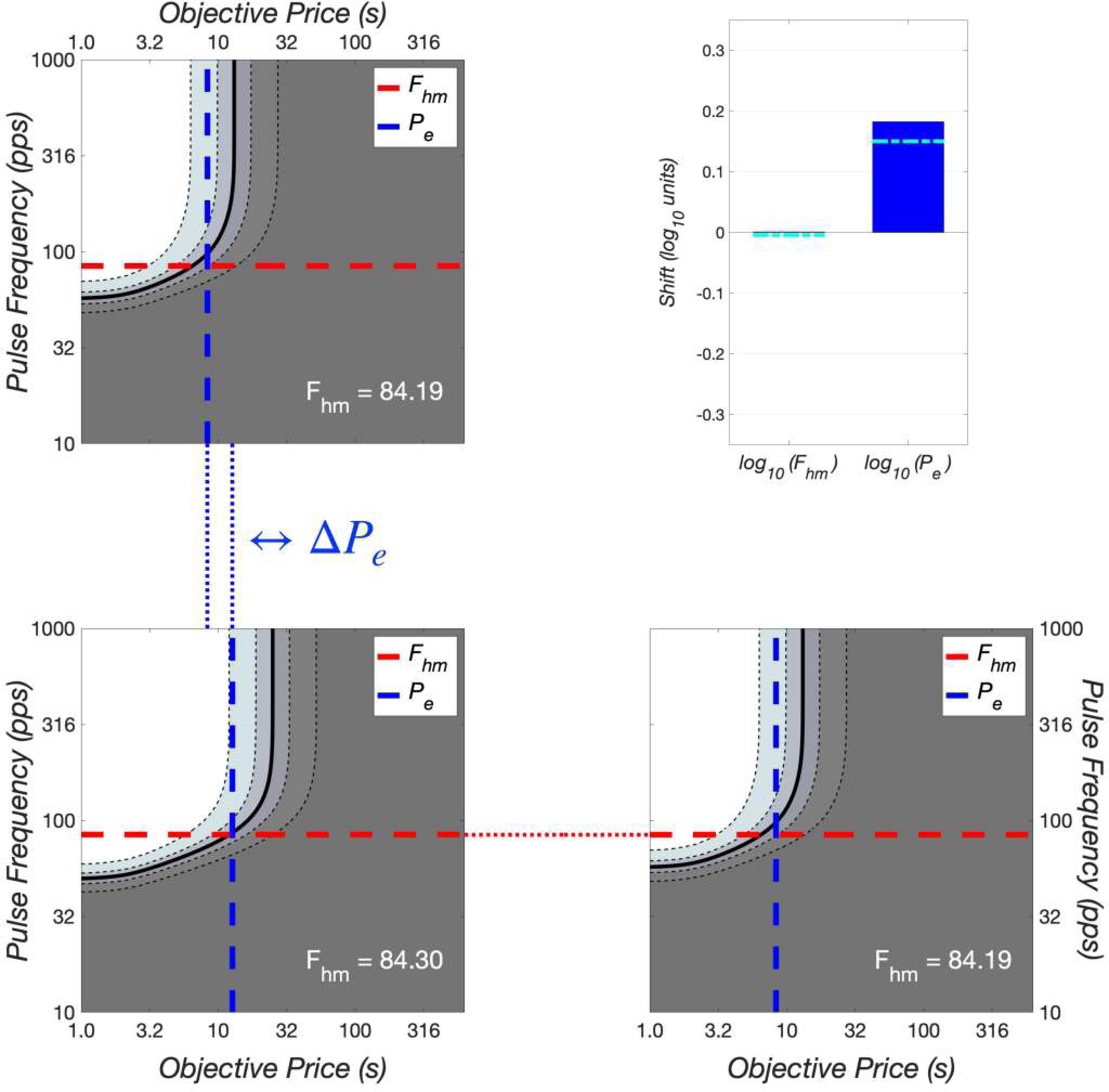
Contour graphs of reward mountains simulated by the convergence model. Dopamine-transporter blockade shifts the simulated reward mountain (almost) exclusively along the pulse-frequency axis, as in the behavioral data. The simulated MFB drive on the dopamine neurons is equivalent to an optical pulse frequency of 40 pulses *s*^−1^.

**Fig S36.**
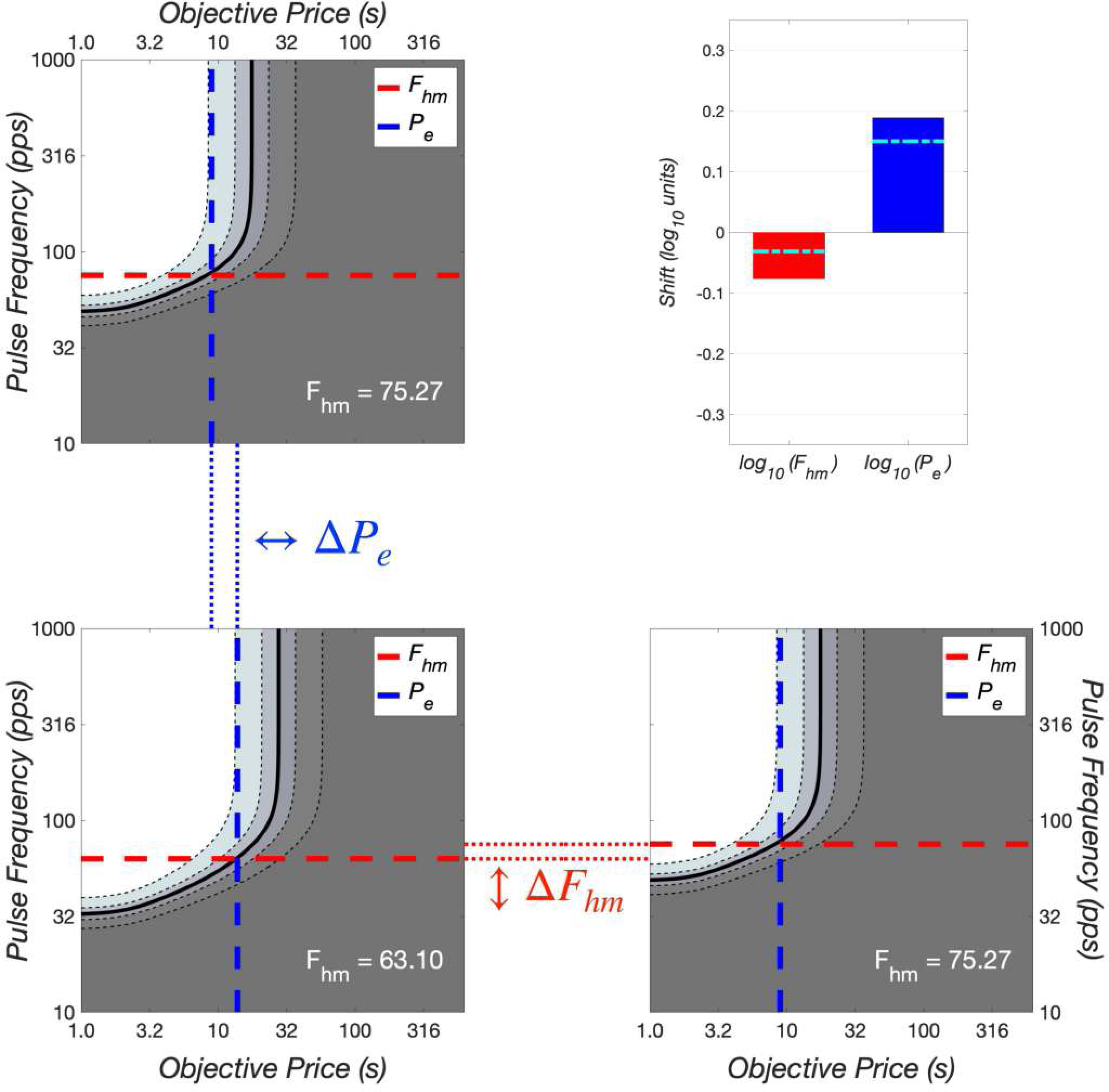
Contour graphs of reward mountains simulated by the convergence model given very strong MFB input. The simulated MFB drive on the dopamine neurons is now equivalent to an optical pulse frequency of 80 pulses *s*^−1^.

### Training in preparation for measurement of the reward mountain

**Fig S37.**
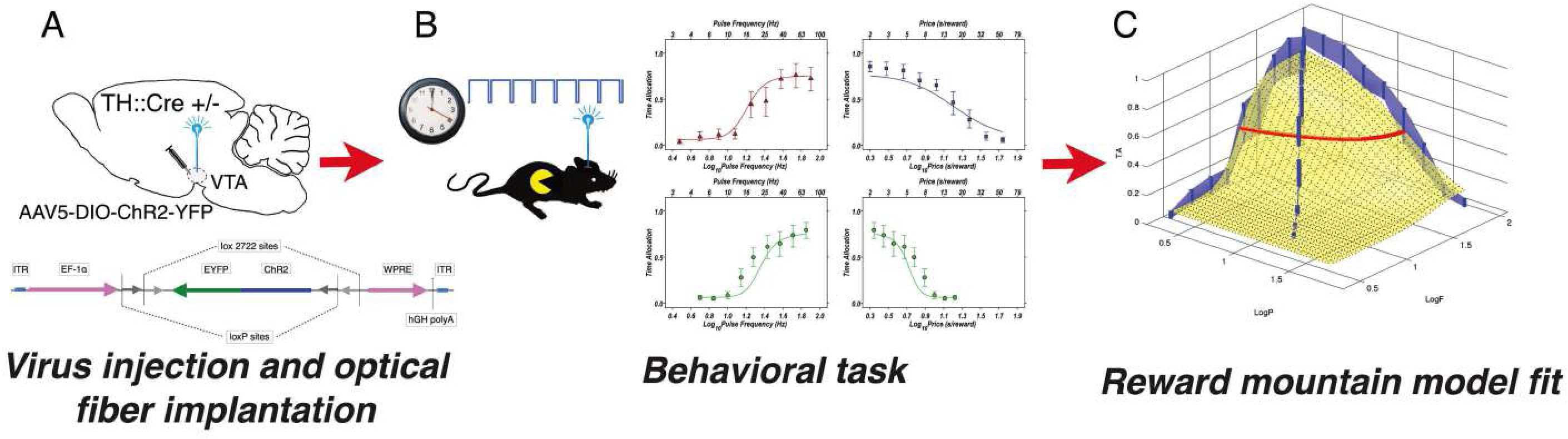
Graphical summary of the experimental procedure. A: TH::Cre +/-rats received bilateral VTA injections of an AAV5 virus bearing a Cre-dependent, ChR2-YFP transcript. Optical fibers were bilaterally aimed at the VTA. B: Rats were trained to hold down a lever for a specified cumulative amount of time to deliver trains of optical stimulation to the VTA. The red curve represents the proportion of trial time the rat spent working for the optical reward as the optical pulse frequency (the reward-strength variable) was systematically manipulated. The blue curve shows the proportion of trial time the rat spent working for a maximal optical reward as the cumulative amount of time required to harvest the reward (“price”) was manipulated systematically. The green curves show proportion of trial time the rat spent working for the optical reward as the strength and price of the reward were simultaneously manipulated. C: The reward-mountain model was fit independently to the data from each rat following injections of GBR-12909 or vehicle. Within subject comparisons were performed.

### Contour lines: the trade-off between pulse frequency and price to hold time allocation constant

Contour graphs provide a compact summary of the reward-mountain surface in a format that facilitates visualization of the direction(s) in which the mountain has been shifted by a manipulation such as administration of a drug. The changes in the values of the location parameters become visually apparent in this format.

Here, we derive the equation for the contour lines, thus updating an earlier derivation [14] in which it had been assumed that the higher pulse frequencies tested drive reward intensity to its maximum attainable value 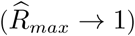. This will indeed be so if the pulse frequencies in question are substantially lower than the frequency-following limit in the directly stimulated neurons. That assumption was usually justified in previous eICSS studies in which the reward mountain was measured [2, 3, 6, 13, 14, 33, 43, 44]. Highly excitably MFB neurons served as the directly activated substrate for the rewarding effect in those studies. In contrast, midbrain dopamine neurons are directly activated substrate in the current study. Not only do these neurons have more limited frequency-following abilities than their MFB counterparts [38, 40], their activation is due to optical excitation of a relatively slow opsin [36] rather than to electrical excitation of voltage-sensitive membrane channels. As we show below, it is likely that the highest pulse frequencies employed in the present study did not always succeed in driving reward intensity to its maximum, particularly in the vehicle condition. To accommodate such cases, we now generalize the previously published expression for the contour lines [14].

We begin by reformatting Eq 33 from the main text as follows:

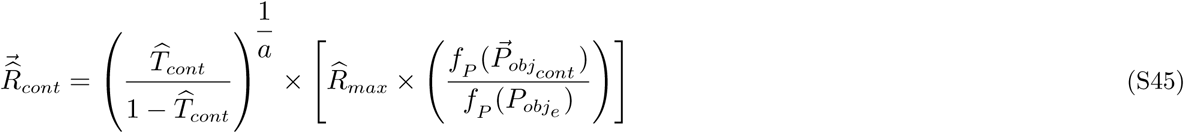

where

*a* = price-sensitivity exponent

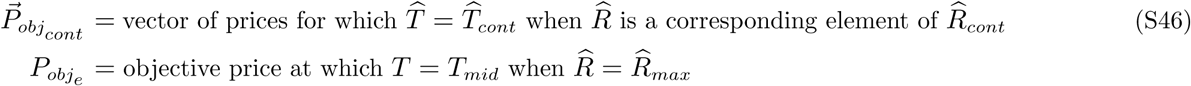

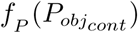 = subjective equivalent of 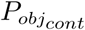

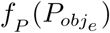 = subective price at which *T* = *T*_*mid*_ when 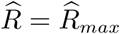

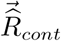 = vector of normalized reward intensities for which 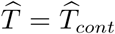 when *P*_*obj*_ is a corresponding element of 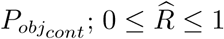

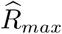 = maximum normalized reward intensity

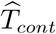 = time allocation represented by the contour line

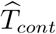 = normalized time allocation represented by the contour line; 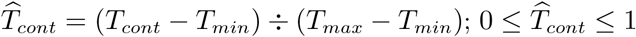;

Substituting for 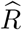 from Eq 12 in the main text, we obtain:

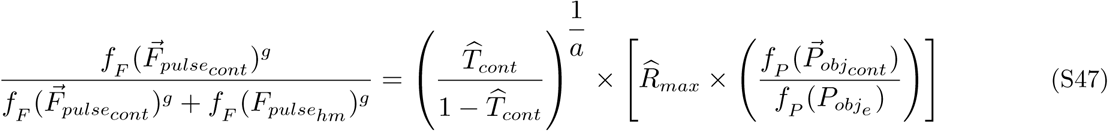

We now multiply both sides by 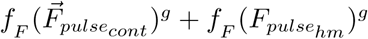, yielding

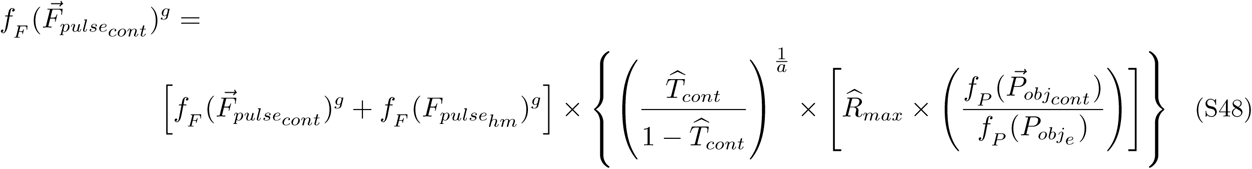

and we then expand the right side to yield:

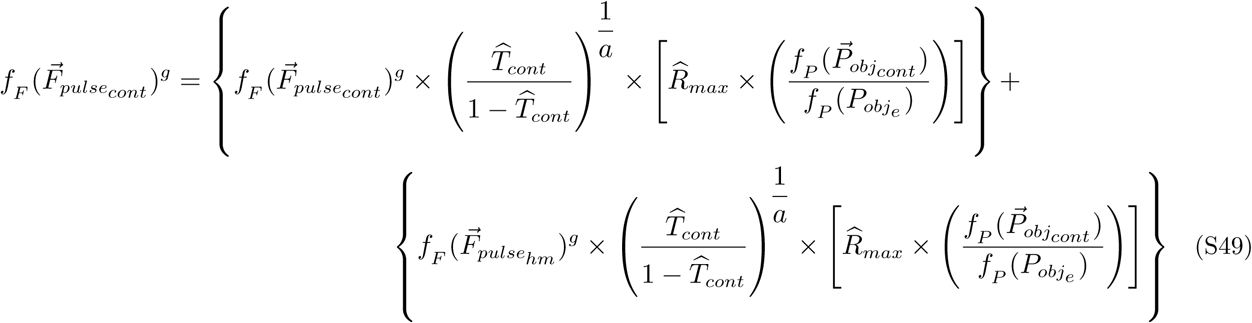

The terms that include 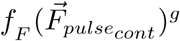 are collected on the left side

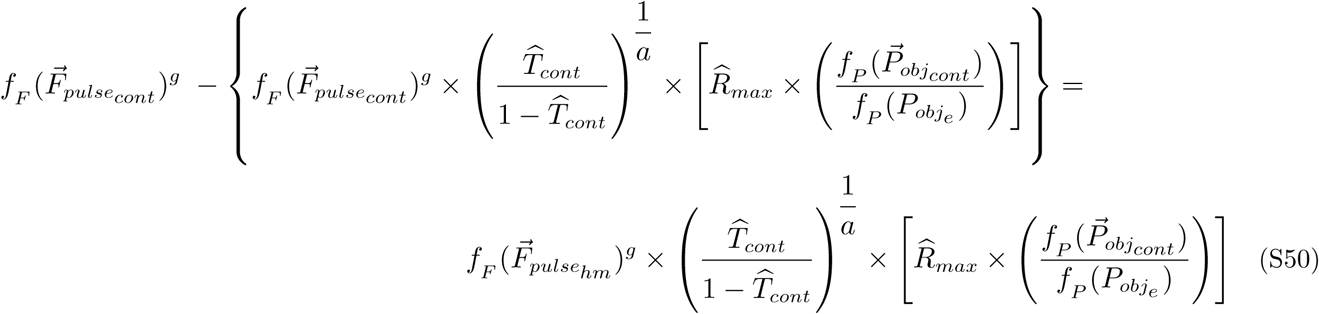

and the left side is factored to yield:

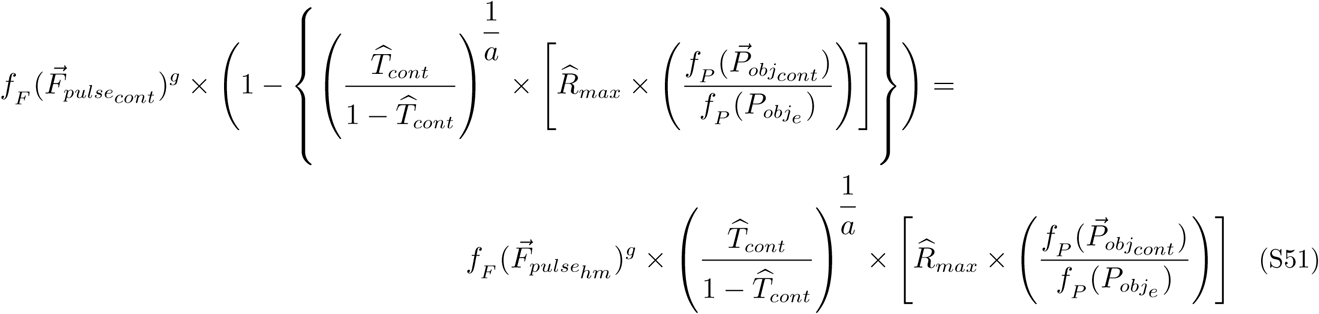

Re-arranging the terms, we obtain:

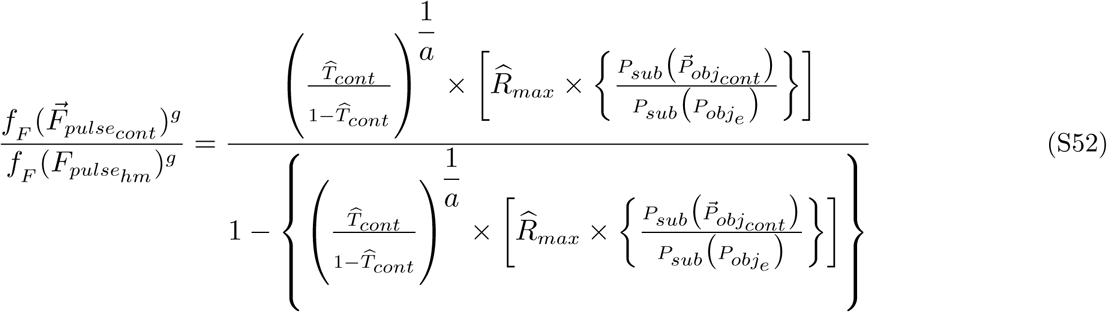

The numerator and denominator of the right side are now multiplied by 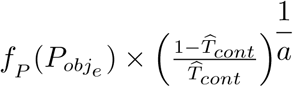 to yield

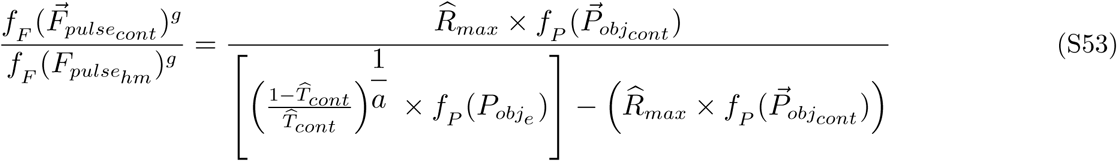

The contour graphs will be plotted in double logarithmic coordinates. In that space, Eq S53 becomes

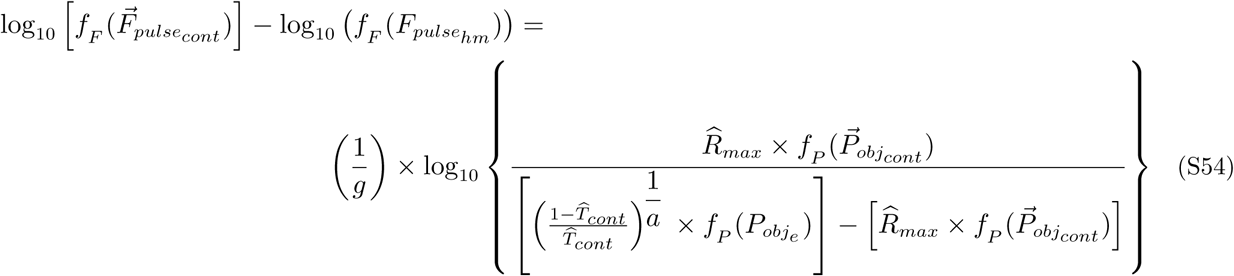

When time allocation falls halfway between *T*_*min*_ and 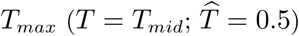, Eq S54 reduces to:

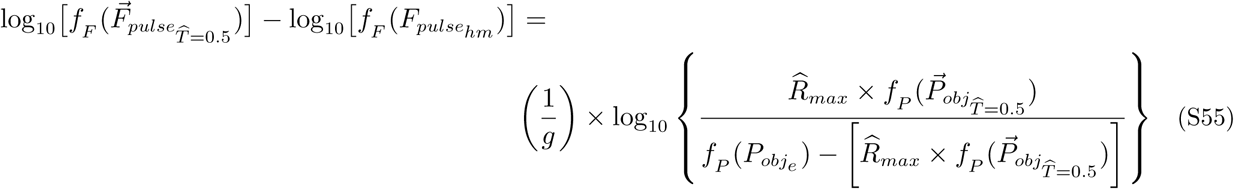

Expanding the right side, we obtain

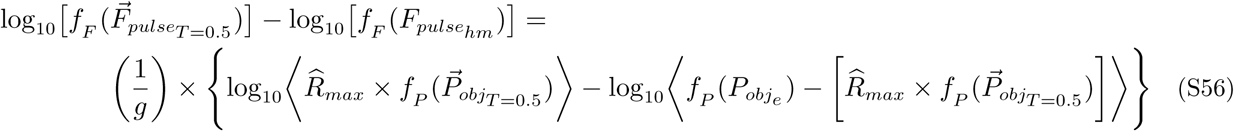

When 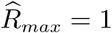, Eq S56 reduces to

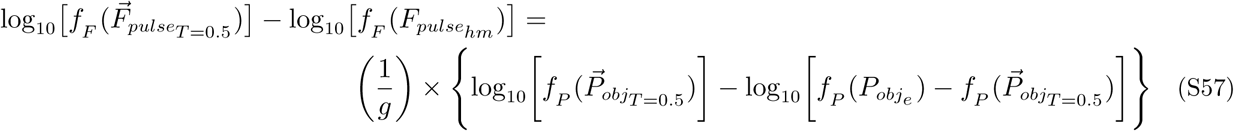

## Instructions for the Matlab Live Script that accompanies the manuscript entitled: The effect of dopamine transporter blockade on optical self-stimulation: behavioral and computational evidence for parallel processing in brain reward circuitry

The Matlab Live Script used in the simulations is available from the following Dropbox folder:

https://www.dropbox.com/sh/75xx1961cmvhxpq/AACuI5RSImSDd8YCig_vos7aa?dl=0

This folder contains three files:

- The Live Script: GBR_eICSS_oICSS_v10.mlx
- A non-executable HTML version of the Live Script that can be viewed in a browser.
- A .zip archive of the 17 graphics files that the Live Script will import: Imported_figures.zip

The setup of the Live Script is described in the section entitled “Preliminaries” and is implemented in lines 4-24.

The Live Script should run on any version of Matlab from R2018a onwards. The script hasn’t been tested on earlier versions, but it may run on versions as far back as R2016a.

Execution of the entire script can take several minutes on a reasonably fast system. It may prove most practical to work through it in sections via the “Run Section” and “Run and Advance” buttons of the Live-Script editor.

Most variables are cleared at the end of each major section. The ones that are required throughout are all defined before line 316 (“Moving the mountain: validation studies”). Once the script has run to that point, the user can navigate directly to sections of interest via the “Go To” button, e.g.

- line 599: “The significance of orthogonal shifts”
- line 870: “At what stage of processing does perturbation of dopaminergic neurotransmission alter reward seeking in eICSS?”
- line 878: “Optical intracranial self-stimulation of midbrain dopamine neurons”
- line 1030: “Dopaminergic modulation of subjective effort cost”
- line 1122: “The series-circuit model of eICSS and oICSS”
- line 1406: “The convergence model”

## MATLAB Live Script to supplement “The effect of dopamine transporter blockade on optical self-stimulation: behavioral and computational evidence for parallel processing in brain reward circuitry”

**This document was not intended to stand alone; It is assumed that anyone working through this script has already read the manuscript, with particular attention to the derivation of the reward-mountain model in the supporting-information file**.

### Preamble

In the current experiment, rats worked for optical stimulation of midbrain dopamine neurons. Time allocated to reward seeking was measured as a function of the strength and cost of the stimulation. We call the resulting three-dimensional data structure the “reward mountain.” Dopamine neurotransmission was perturbed by administration of a dopamine-transporter blocker, and the consequent displacement of the reward mountain was determined.

The reward-mountain model was developed to account for data from experiments on rats working for rewarding electrical brain stimulation (electrical intracranial self-stimulation: eICSS). This model incorporates information obtained over many decades of research on the neural circuitry that intervenes between the tip of the stimulating electrode and the behavioral effects of the stimulation. Although this work has been dogged by uncertainty about the identity of the directly stimulated neurons, painstaking psychophysical experiments have provided extensive information about the directly activated neurons responsible for the rewarding effect, the operating principles of the neural circuitry in which they are embedded, and the reward-seeking behavior generated by this circuitry. Of particular importance to the current study are experiments characterizing the spatiotemporal integration of the electrically evoked reward signals and the growth of the rewarding effect as a function of stimulation strength.

In contrast to the case of eICSS, the identity of the directly stimulated substrate responsible for optical intracranial self-stimulation (oICSS) of midbrain dopamine neurons is known - the stimulation specifically and directly activates the dopaminergic neurons. However, only rudimentary information is available to date about the growth or spatiotemporal integration of the optically induced reward signal. One purpose of this document is to explore what the current experiment reveals about spatiotemporal integration and reward growth in the neural circuitry underlying oICSS. Another purpose is to explore the implications of the current work for understanding how eICSS and oICSS are related at the level of neural circuitry. We show by means of simulation that the results of the current experiment pose grave difficulties for an intuitively appealing account we and others proposed previously: that eICSS arises from the indirect activation of the directly stimulated dopamine neurons that give rise to oICSS. These difficulties challenge widely held notions about the organization of brain circuitry underlying reward.

## Preliminaries

Setting the ***graph2files*** variable to true will cause graphs to be written to external files. These files are stored in a subdirectory of the current folder called ‘Figures.’ Setting this variable to false will speed execution.

The ***show_graphics*** variable determines whether graphics are stored within this Live Script. This variable must be set to false to enable efficient editing of this Live Script. When this Live Script has been executed

with ***show_graphics*** set to true, subsequent editing and task switching will be slowed unacceptably. Set ***show_graphics*** to true only in preparation for saving updated html and/or pdf copies of this file. To prepare for editing after this script has been stored with ***show_graphics*** set to true, reset this variable to false, re-run the script, and save it.

Setting the ***tabs2files*** variable to true will store information about equations, figures, functions, and symbols in external Excel files. Setting this variable to false may speed execution slightly.

Setting the ***saveWS*** variable to true will cause the Matlab workspace to be stored in a .mat file in the folder from which this live script was run.

Be sure to set the default_dir to the directory you wish to use as the default on your system.

As its name implies, the ‘***Imported_Figures***’ folder contains images that this Live Script will import from external files. Those images should be stored in a sub-folder of the folder from which this Live Script will be run. The name of that folder is the argument of the ***set_impfigdir*** command below.

Blocks of redundant code are included in various sections. Although this lengthens the Live Script, it reduces the memory load due to the accumulation of workspace variables. Given that most variables are cleared at the boundaries between major sections, the blocks of redundant code enable debugging by means of “Run Section,” “Run and Advance,” etc. In a future revision, efficiency may be increased by aggregating the variables assigned in these blocks in structures that are saved and re-loaded.

~~~
t_start = tic; % start timer for entire script
tic; % start timer for the current section
global graphs2files show_graphics
graphs2files = true; % save graphics to external files?
tabs2files = true; % save tables of equation #s, figure #s, function #s, and symbol #s?
saveWS = true; % save final workspace?
show_graphics = true; % display graphics in this Live Script?
version = 10; % used in the names of stored files
% define and set the default directory
default_dir = ‘∼/Work/Research/papers/In_Progress/Opto_GBR2/Simulations’;
if ∼exist(default_dir, ‘dir’)
  disp(strcat({‘Default directory ‘}, default_dir, {‘does not exist.’}));
  return
end
cd(default_dir);
% Define the directory that will contain figures generated here & create it if necessary.
global FigDir
FigDir = set_figdir(‘Figures’, graphs2files); % Sub-directory of default directory
% Define the directory containing stored images to be imported.
ImpFigDir = set_impfigdir(‘Imported_figures’); % Sub-directory of default directory
% Define the directory that will house the tables & create if if necessary.
tabdir = set_tabdir(‘Tables’, tabs2files); % Sub-directory of default directory
~~~

The following technical section implements preliminary steps required to set up the simulations.

In order to set up the simulations, some code must be executed to define basic functions and initialize variables.

This document was designed to run on any version of Matlab from R2018a onwards without requiring installation of external function files or scripts. All functions are either built-in (supplied by the Mathworks as part of the standard Matlab installation) or local (i.e., defined at the end of this Live-Script document). This should allow this script to execute on any standard installation of the supported Matlab versions.

The following code initializes the tables that store information about the formatted equations, figures and symbols used in this script.

~~~
%% Initialization
~~~

Initialize the tools that build the tables of equations, figures, functions, and symbols. Load the numbers and descriptions of the pre-defined functions (see below, “***Tools for this live script***” and “***Building blocks for the functions included in the simulations***.”

~~~
global eqn_num eqn_tab fig_num fig_tab fun_num fun_tab sym_num sym_tab init_all;
~~~

~~~
The equation table has been initialized.
The figure table has been initialized.
The function table has been initialized.
The symbol table has been initialized.
This is function #1: (init_all)
This is function #2: (add_eqn)
This is function #3: (add_fig)
This is function #4: (add_fun)
This is function #5: (add_sym)
This is function #6: (FilterFun)
This is function #7: (FilterFunBS)
This is function #8: (LogistNormFun)
This is function #9: (LogistNormBsFun)
This is function #10: (LogistNormBsLocFun)
This is function #11: (PsubFun)
This is function #12: (PsubBsFun)
This is function #13: (ScalarDivFun)
This is function #14: (ScalarDivBsFun)
This is function #15: (ScalarMultFun)
This is function #16: (ScalarMultBsFun)
This is function #17: (PsubEfun)
This is function #18: (PobjEfun)
~~~

Load the numbers and descriptions of the pre-defined functions (see below, “***Tools for this live script***” and “***Building blocks for the functions included in the simulations***.”

Enter symbols defined in the introductory paragraphs above.

~~~
[sym_num, sym_tab] = add_sym(sym_num, sym_tab, “eICSS”, “electrical intracranial self-stimulation”);
[sym_num, sym_tab] = add_sym(sym_num, sym_tab, “oICSS”, “optical intracranial self-stimulation”);
keepVars = who; % Store variables to be retained
toc
~~~

`Elapsed time is 0.110595 seconds.`

## The mountain model

~~~
tic;
~~~

The reward-mountain model was developed to account for operant performance as a function of the cost and strength of an experimenter-controlled reward, within the context of eICSS studies. The task performed to obtain the rewarding stimulation entails putting time on a clock by holding down a lever; a reward is delivered when the cumulative time the lever has been depressed reaches an experimenter-defined criterion (the “***price***” of the reward). The measure of operant performance employed is time allocation: the proportion of trial time that the rat devotes to procurement of the rewarding stimulation (“work”). The model is derived in the main body of the accompanying manuscript and then adapted for application to oICSS. Papers cited here are listed in the reference section of the main body of the manuscript.

In this section, we define the functional building blocks that will be used to simulate the output of the reward-mountain model.

We distinguish between the “shell” and “core” of the model. The shell consists of the variables that are observed (time allocation), manipulated (pulse frequency, price), and controlled (stimulation parameters held constant, physical work required to hold down the lever, affordances of the test environment for leisure activites, such as grooming, resting, and exploring). The core consists of the functions that compute the intensity of the reward produced by the stimulation train and combine this value with the opportunity and effort costs entailed in its procurement, thus generating what we call “payoffs.” A single function based on the generalized matching law translates the payoffs generated in the core into the time-allocation values that are manifest in the shell.

Please see the main body of the text for definitions, explanations, and details.

## Shell→core functions

### The frequency-following function

Solomon et al. (2015) studied frequency-following fidelity in rats working for rewarding electrical stimulation of the medial forebrain bundle (MFB). They showed that the following function provides a good description of the relationship between the induced frequency of firing in the directly stimulated neurons and the pulse frequency:

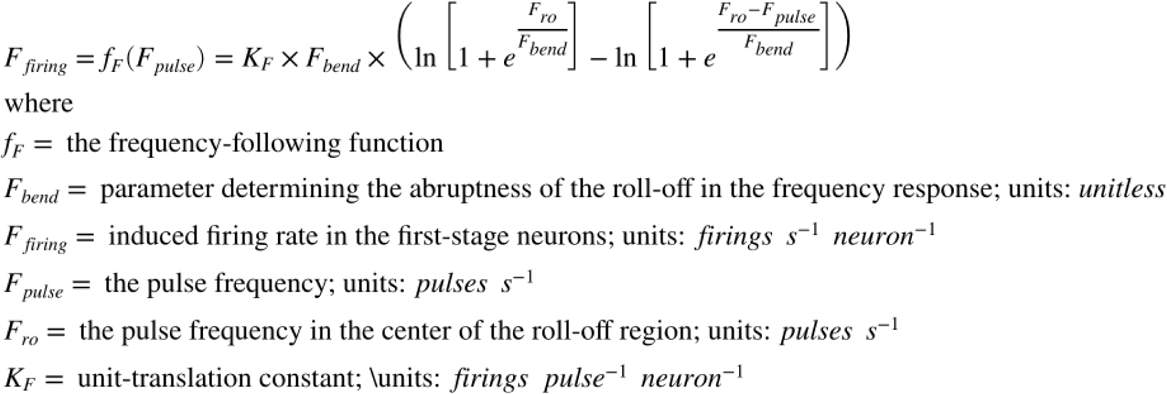

~~~
[eqn_num, eqn_tab] = …
  add_eqn(eqn_num, eqn_tab, “FreqFolElec”, “Frequency of firing as a function of electrical pulse frequency”);
~~~

`This is equation #1: (FreqFolElec)`

~~~
[sym_num, sym_tab] = add_sym(sym_num, sym_tab, “fF”, “frequency-following function”);
[sym_num, sym_tab] = add_sym(sym_num, sym_tab, “Fpulse”, “pulse frequency”);
[sym_num, sym_tab] = add_sym(sym_num, sym_tab, “Fbend”, “parameter determining sharpness of bend in the FreqFol function”);
[sym_num, sym_tab] = add_sym(sym_num, sym_tab, “Fro”, “the pulse frequency in the center of the roll-off region”);
[sym_num, sym_tab] = add_sym(sym_num, sym_tab, “Ffiring”, “induced firing rate of the directly stimuled neurons”);
[sym_num, sym_tab] = add_sym(sym_num, sym_tab, “Kf”, “unit-translation constant for the frequency-following function”);
[fun_num, fun_tab] = add_fun(fun_num, fun_tab, “FilterFun”, “F, Fbend, Fro”,…
  “Frequency-following function”);
~~~

`Function FilterFun has already been entered.`

The following graph relates the induced firing frequency to the electrical pulse frequency:

~~~
logFelecBend = 1.3222; % from Solomon et al., 2015
FelecBend = 10^logFelecBend;
logFelecRO = 2.5587; % from Solomon et al., 2015
FelecRO = 10^logFelecRO;
Felec = logspace(0,3,121);
Fmfb = FilterFun(Felec, FelecBend, FelecRO); % Compute firing rate using the frequency roll-off function
FF_graph = plot_freqFoll(Felec,Fmfb,’mfb’,2,1000,2,1000); % see “Functions that plot graphs and set attributes” below
if show_graphics
  FF_graph.Visible = ‘on’;
end
~~~

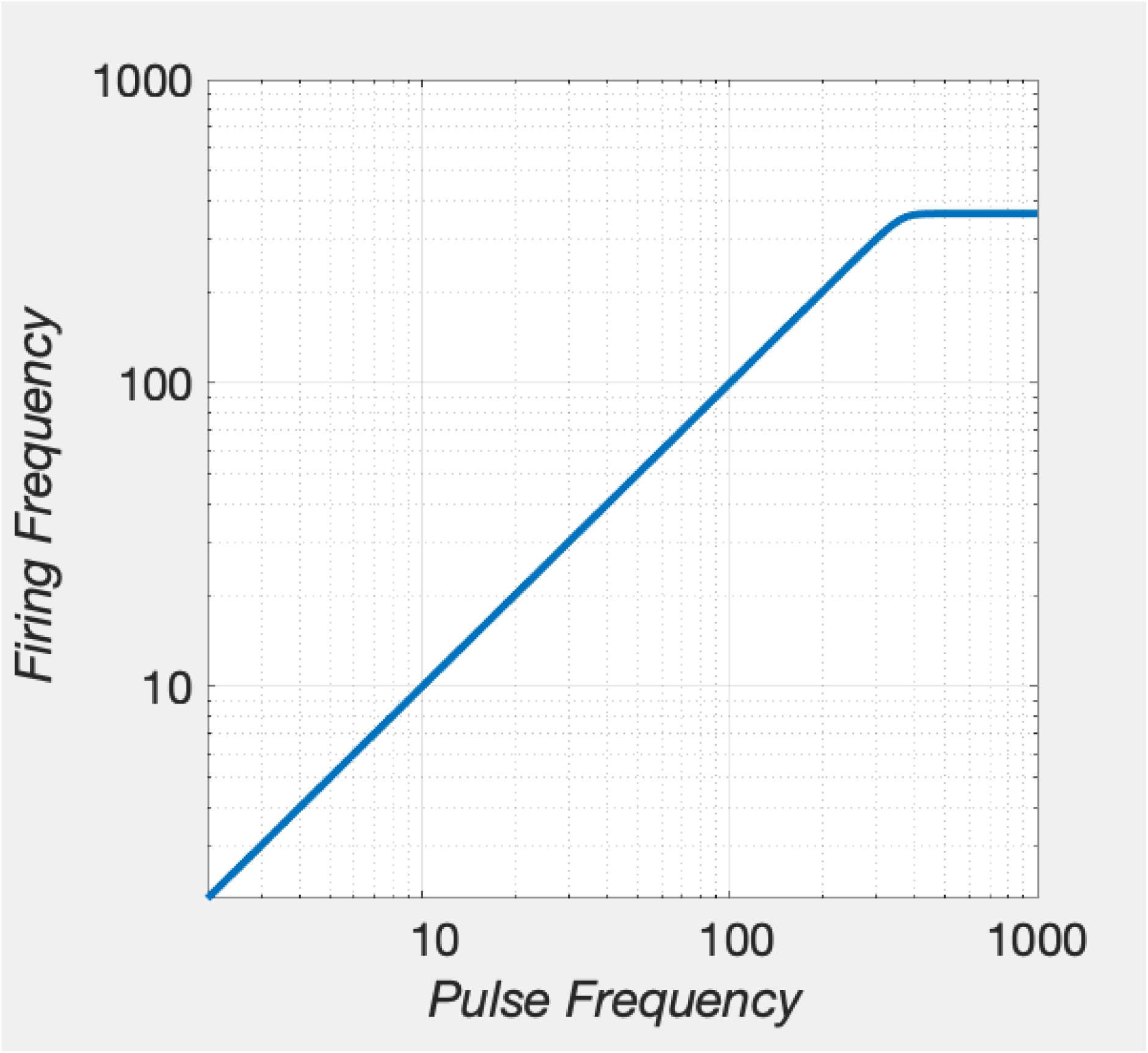

~~~
[fig_num, fig_tab] = add_fig(fig_num, fig_tab, “FreqFolMFB”, …
  “Induced firing frequency in the directly stimulated neurons as a function of pulse frequency”);
~~~

`This is figure #1: (FreqFolMFB)`

This graph shows that the firing of the directly stimulated neurons subserving eICSS of the medial forebrain bundle can follow the pulse frequency faithfully up to very high values (>= 350 pulses per second); the response of the neurons flattens abruptly as pulse frequency is increased further. This high-fidelity frequency following is consistent with the view that the directly stimulated cells subserving the rewarding effect are non-dopaminergic neurons with myelinated axons (Shizgal, 1997; Bielajew & Shizgal, 1986; Gallistel, Yeomans & Shizgal, 1981; Bielajew & Shizgal, 1982; Shizgal et al., 1980).

In addition to finding firing frequencies corresponding to a given pulse frequency, we will also need to do the reverse: finding the pulse frequency required to produce a given firing frequency. Thus, we define a back- solution of the frequency-following function:

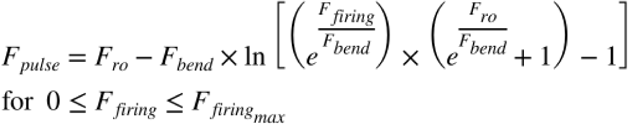

~~~
[eqn_num, eqn_tab] = add_eqn(eqn_num, eqn_tab, “fFbacksolved”, “Back-solution of the frequency-following function”);
~~~

`This is equation #2: (fFbacksolved)`

### The neural-recruitment function

The *f*_*N*_ function translates the pulse duration and current into the number of electrically excited neurons:

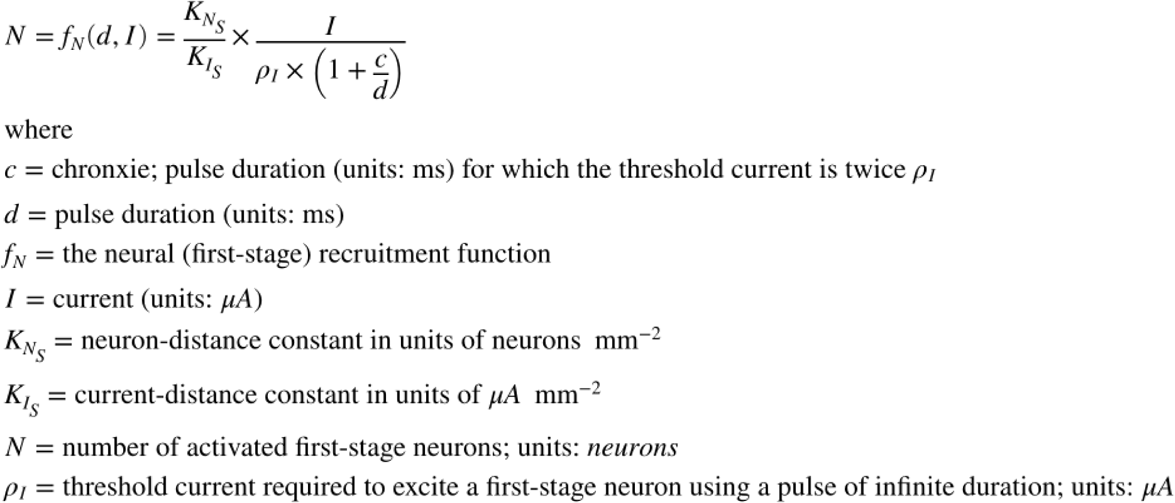

~~~
[eqn_num, eqn_tab] = add_eqn(eqn_num, eqn_tab, “fN”, “Neural-recruitment function”);
~~~

`This is equation #3: (fN)`

~~~
[sym_num, sym_tab] = add_sym(sym_num, sym_tab, “c”, “chronaxie of strength-duration function for pulses”);
[sym_num, sym_tab] = add_sym(sym_num, sym_tab, “d”, “pulse duration”);
[sym_num, sym_tab] = add_sym(sym_num, sym_tab, “fN”, “Neural-recruitment function”);
[sym_num, sym_tab] = add_sym(sym_num, sym_tab, “I”, “current”);
[sym_num, sym_tab] = add_sym(sym_num, sym_tab, “Kns”, “Neuron-recruitment function”);
[sym_num, sym_tab] = add_sym(sym_num, sym_tab, “Kis”, “Current-distance constant”);
[sym_num, sym_tab] = add_sym(sym_num, sym_tab, “N”, “Number of activated first-stage neurons”);
[sym_num, sym_tab] = add_sym(sym_num, sym_tab, “rhoI”, “rheobase: threshold current to excite a first-stage neuron with a pulse of infinite duration”);
~~~

Although we define this function here, we do not implement it or use it in the simulations. Instead, an output value is chosen that, together with a value for the rheobase of the strength-duration function for trains, generates location-parameter values within the range observed in past studies. This function is included in order to summarize the relationships on which existing tests of the counter model have been based (e.g., Simmons & Gallistel, 1994).

### The burst-duration function

For completeness, we include a function that maps the duration of the pulse train into the duration of the evoked increase in the firing of the first-stage neurons. In practice, we assume that the two are equal, as will be case when frequency-following fidelity is high.

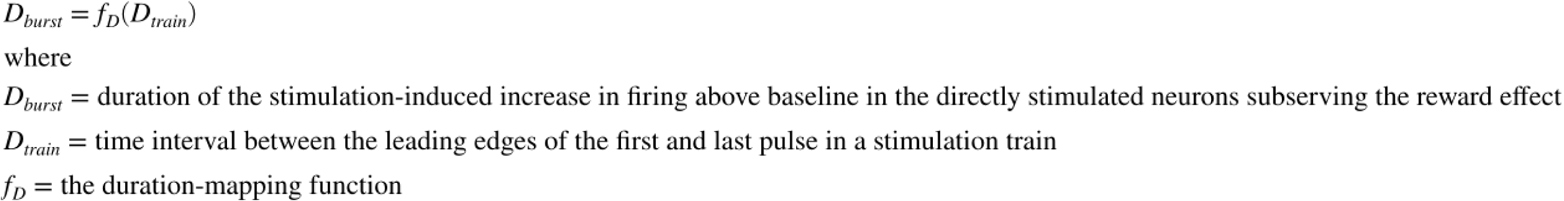

~~~
[eqn_num, eqn_tab] = add_eqn(eqn_num, eqn_tab, “fD”, “Train-burst equation”);
~~~

`This is equation #4: (fD)`

~~~
[sym_num, sym_tab] = add_sym(sym_num, sym_tab, “Dburst”, “Duration of stimulation-induced increase in firing”);
[sym_num, sym_tab] = add_sym(sym_num, sym_tab, “Dtrain”, “train duration”);
[fun_num, fun_tab] = add_fun(fun_num, fun_tab, “fD”, “Dtrain”,…
  “Train-burst function”);
~~~

`This is function #19: (fD)`

### The subjective-probability function

This is another dummy function. As we review below, the subjective probability that a reward will be delivered upon satisfaction of the response requirement appears to equal the objective probability when the latter is 0.5 or higher. In the case of the present oICSS experiment, the objective probability of reward is always one.

*n*.*b*. A lower-case “p” is employed in the symbols.

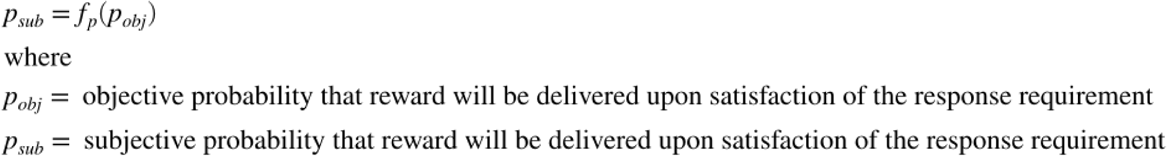

~~~
[eqn_num, eqn_tab] = add_eqn(eqn_num, eqn_tab, “fp”, “Subjective-probability equation”);
~~~

`This is equation #5: (fp)`

~~~
[sym_num, sym_tab] = add_sym(sym_num, sym_tab, “ProbObj”, “Objective probability that reward will be delivered upon satisfaction of the response requirement”);
[sym_num, sym_tab] = add_sym(sym_num, sym_tab, “ProbSub”, “Subjective probability that reward will be delivered upon satisfaction of the response requirement”);
[fun_num, fun_tab] = add_fun(fun_num, fun_tab, “ProbSubFun”, “ProbObj”,…
  “Subjective-probability function”);
~~~

`This is function #20: (ProbSubFun)`

### The subjective-price (opportunity-cost) function

As described in Solomon et al., (2017), the subjective-price function is defined as

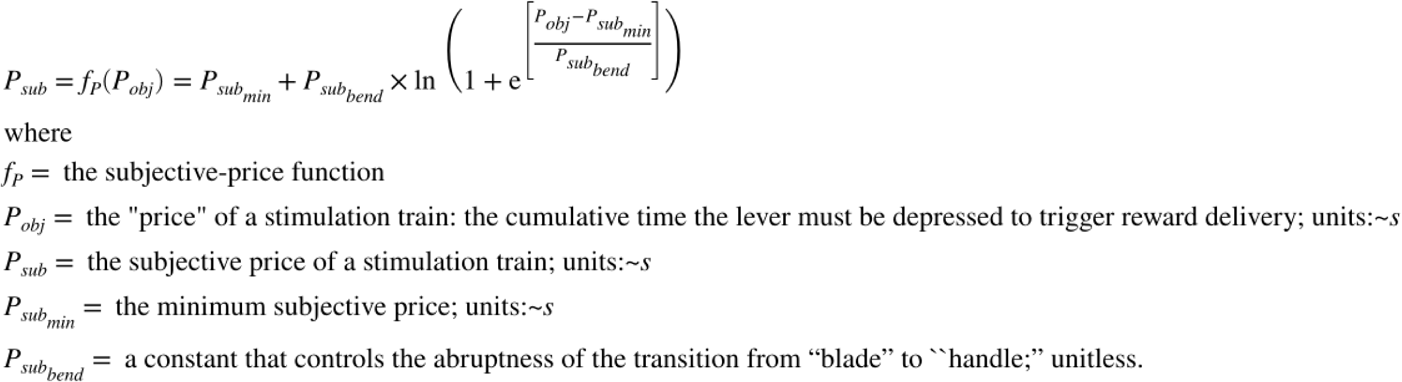

~~~
[eqn_num, eqn_tab] = …
  add_eqn(eqn_num, eqn_tab, “SubjPriceFun”, “Subjective-price function”);
~~~

`This is equation #6: (SubjPriceFun)`

~~~
[sym_num, sym_tab] = add_sym(sym_num, sym_tab, “Pobj”, “objective price (opportunity cost)”);
[sym_num, sym_tab] = add_sym(sym_num, sym_tab, “Psub”, “subjective price”);
[sym_num, sym_tab] = add_sym(sym_num, sym_tab, “PsubBend”, “transition parameter of subjective-price function”);
[sym_num, sym_tab] = add_sym(sym_num, sym_tab, “PsubMin”, “minimum subjective price”);
~~~

and plotted below.

~~~
PsubBend = 0.5;
PsubMin = 1.82;
Pobj = logspace(−1,2,120);
Psub = PsubFun(Pobj, PsubBend, PsubMin);
SP_graph = plot_PsubFun(Pobj,Psub, graphs2files, FigDir);
if show_graphics
  SP_graph.Visible = ‘on’;
end
~~~

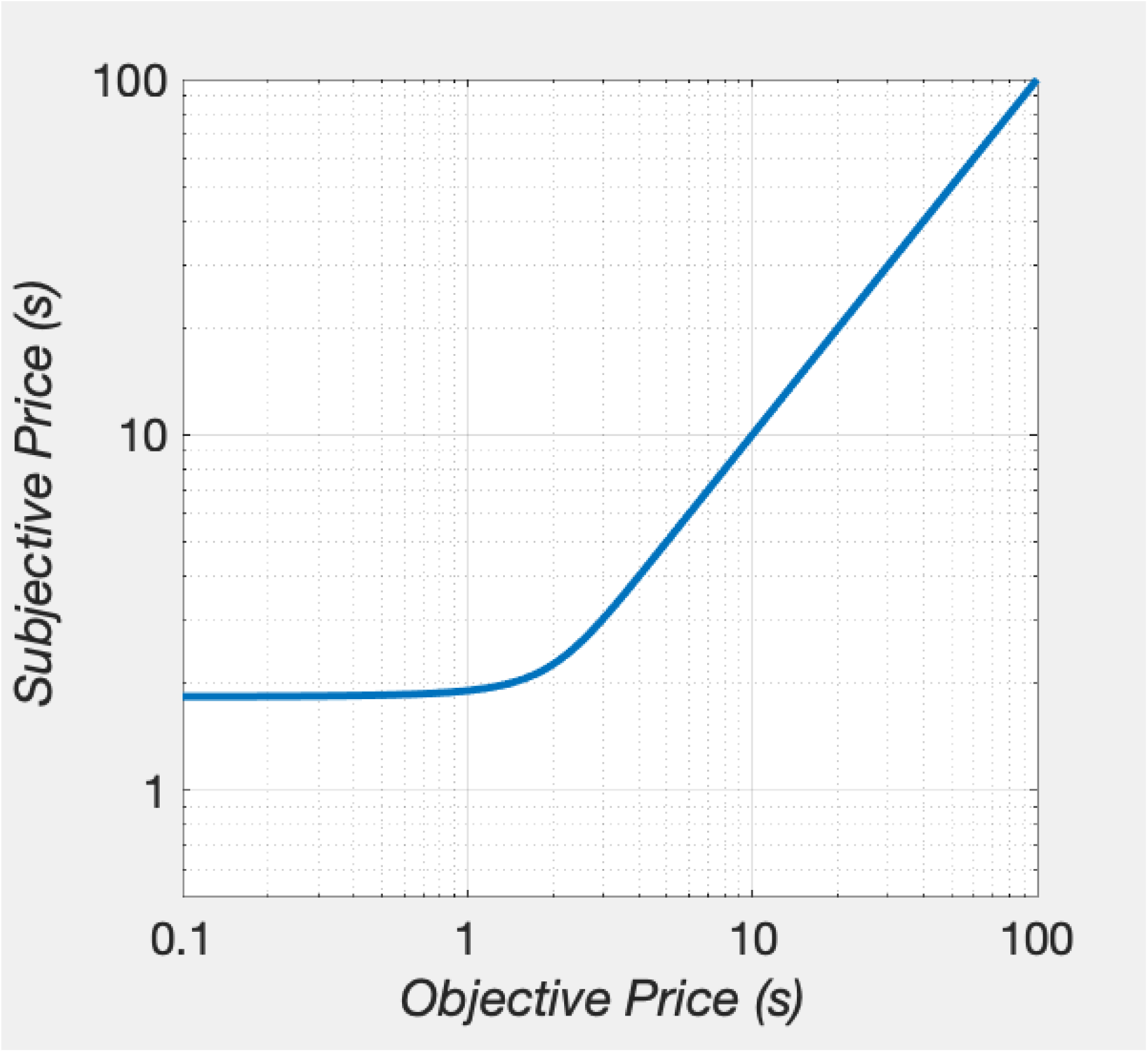

~~~
[fig_num, fig_tab] = add_fig(fig_num, fig_tab, “PsubFun”, … “
  The subjective-price function”);
~~~

`This is figure #2: (PsubFun)`

In addition to finding subjective prices corresponding to objective ones, we will also need to do the reverse. Thus, we define a back-solution of the subjective-price function:

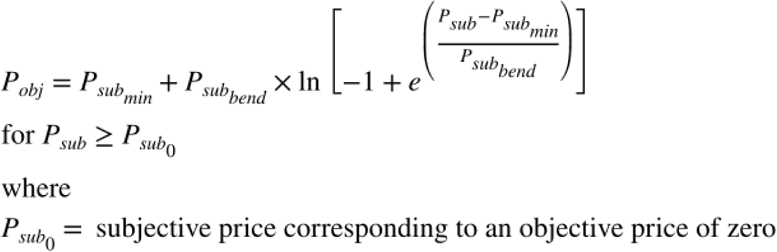

~~~
[eqn_num, eqn_tab] = add_eqn(eqn_num, eqn_tab, “fPbacksolved”, “Back-solution of the subjective-price function”);
~~~

`This is equation #7: (fPbacksolved)`

### The subjective effort-cost function

The physical work entailed in holding down the lever is transformed into the subjective rate of exertion by the following function:

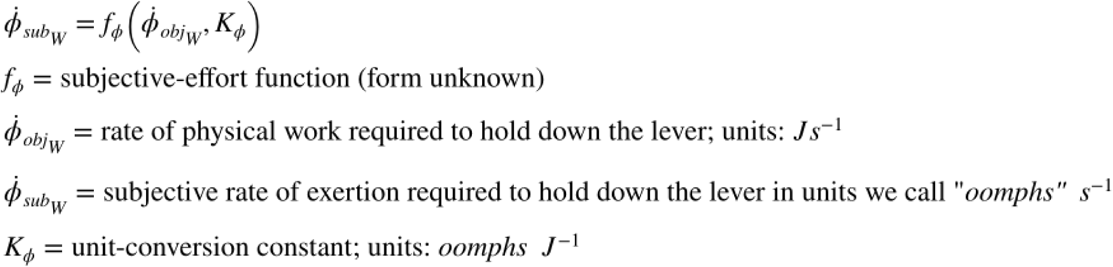

We do not attempt to model the form and parameters of this function. The dots over 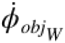 and 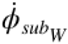 signify that we define these quantities as rates.

~~~
[eqn_num, eqn_tab] = add_eqn(eqn_num, eqn_tab, “fphiW”, “subjective-effort-cost equation for work”);
~~~

`This is equation #8: (fphiW)`

~~~
[sym_num, sym_tab] = add_sym(sym_num, sym_tab, “fphi”, “subjective effort-cost function”);
[sym_num, sym_tab] = add_sym(sym_num, sym_tab, “dotPhiObj”, “objective work rate required to hold down the lever”);
[sym_num, sym_tab] = add_sym(sym_num, sym_tab, “dotPhiSub”, “subjective rate of exertion entailed in holding down the lever”);
[sym_num, sym_tab] = add_sym(sym_num, sym_tab, “Kphi”, “unit-conversion scalar for the effort cost of work”);
[fun_num, fun_tab] = add_fun(fun_num, fun_tab, “fphi”, “dotPhiObj, Kec”,…
   “subjective-effort function”);
~~~

`This is function #21: (fphi)`

The effect of the drug, if any, on the rate of subjective exertion is defined as:

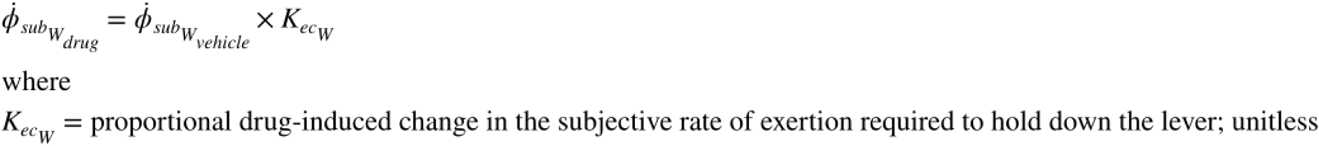

~~~
[eqn_num, eqn_tab] = add_eqn(eqn_num, eqn_tab, “fphiWdrug”, “drug-induced change in the subjective effort cost of work”);
~~~

`This is equation #9: (fphiWdrug)`

~~~
[sym_num, sym_tab] = add_sym(sym_num, sym_tab, “KecW”, “proportional drug-induced change in the subjective rate of exertion required to hold down the lever”);
~~~

In the vehicle condition, 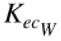 assumes an implicit value of one.

Activities such as grooming and exploring also entail performance of physical work. Thus, the subjective effort-cost function is also applied to these activities:

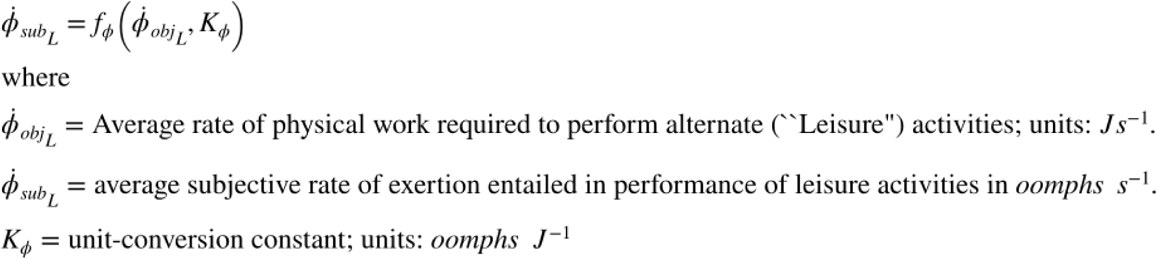

~~~
[eqn_num, eqn_tab] = add_eqn(eqn_num, eqn_tab, “fphiL”, “subjective-effort-cost equation for leisure”);
~~~

`This is equation #10: (fphiL)`

~~~
[sym_num, sym_tab] = add_sym(sym_num, sym_tab, “dotPhiObjL”, “average objective work rate required to perform leisure activities”);
[sym_num, sym_tab] = add_sym(sym_num, sym_tab, “dotPhiSubL”, “average subjective rate of exertion entailed in performing leisure activities”);
~~~

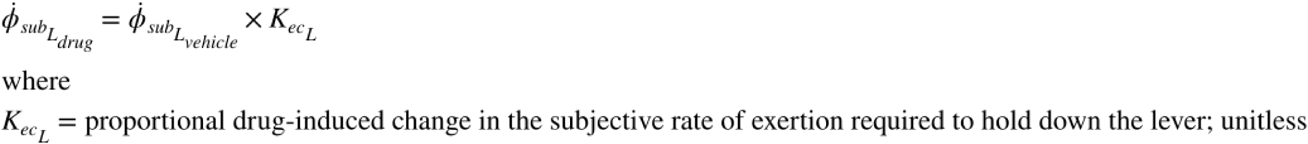

~~~
[eqn_num, eqn_tab] = add_eqn(eqn_num, eqn_tab, “fphiLdrug”, “drug-induced change in the subjective effort cost of leisure activities”);
~~~

`This is equation #11: (fphiLdrug)`

~~~
[sym_num, sym_tab] = add_sym(sym_num, sym_tab, “KecL”, “proportional drug-induced change in the subjective rate of exertion required to perform leisure activities”);
~~~

In the vehicle condition, 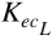 assumes an implicit value of one.

## Core functions

### The reward-growth function for brain stimulation reward (BSR)

The reward-growth function translates the aggregate rate of stimulation-induced firing into the intensity of the rewarding effect. Gallistel’s team used operant matching to describe this function (Gallistel & Leon, 1991, Leon & Gallistel, 1992, Mark & Gallistel, 1993; Simmons & Gallistel, 1994). Shizgal (2003) proposed the following form for this function:

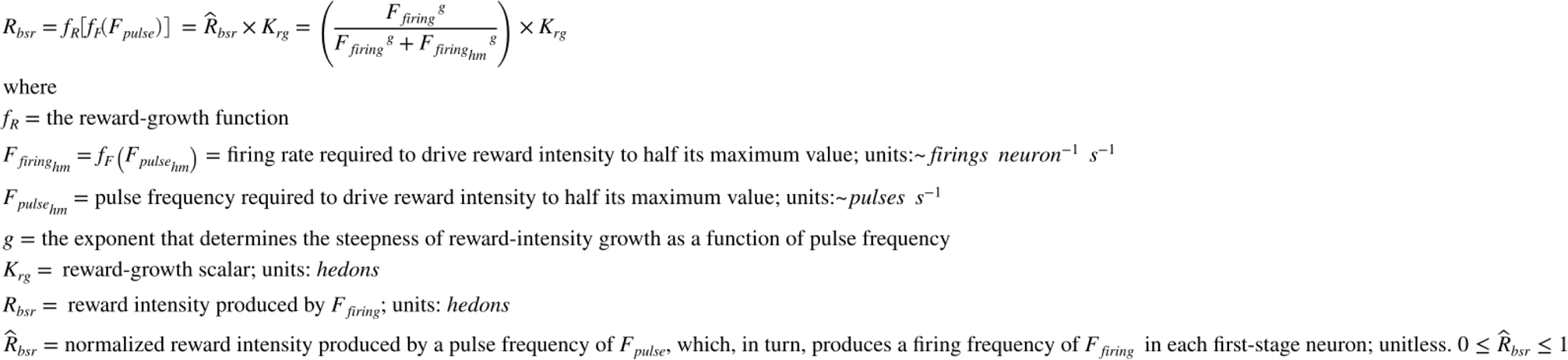

~~~
[eqn_num, eqn_tab] = add_eqn(eqn_num, eqn_tab, “fRbsr”, “reward-growth equation for BSR”);
~~~

`This is equation #12: (fRbsr)`

~~~
[sym_num, sym_tab] = add_sym(sym_num, sym_tab, “fRbsr”, “reward-growth function”);
[sym_num, sym_tab] = add_sym(sym_num, sym_tab, “FfiringHM”, “firing frequency that produces a reward of half-maximal intensity”);
[sym_num, sym_tab] = add_sym(sym_num, sym_tab, “FpulseHM”, “pulse frequency that produces a reward of half-maximal intensity”);
[sym_num, sym_tab] = add_sym(sym_num, sym_tab, “g”, “reward-growth exponent”);
[sym_num, sym_tab] = add_sym(sym_num, sym_tab, “Krg”, “output scalar of the reward-growth function”);
[sym_num, sym_tab] = add_sym(sym_num, sym_tab, “Rbsr”, “reward intensity produced by Ffiring”);
[sym_num, sym_tab] = add_sym(sym_num, sym_tab, “hatRbsr”, “normalized reward intensity produced by a pulse frequency of Fpulse”);
[fun_num, fun_tab] = add_fun(fun_num, fun_tab, “fRbsr”, “F, Fbend, Fhm, Fro, g, Krg”,…
   “Logistic reward-growth function”);
~~~

`This is function #22: (fRbsr)`

It follows from the above that

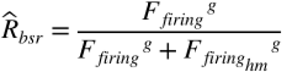

~~~
[eqn_num, eqn_tab] = add_eqn(eqn_num, eqn_tab, “fRbsrNorm”, “normalized reward-growth equation”);
~~~

`This is equation #13: (fRbsrNorm)`

~~~
[fun_num, fun_tab] = add_fun(fun_num, fun_tab, “fRbsrNorm”, “F, Fbend, Fhm, Fro, g”,…
  “Normalized reward-growth function”);
~~~

`This is function #23: (fRbsrNorm)`

To accommodate the predicted rescaling of the input to the reward-growth function for oICSS by dopamine-transporter blockade, a scalar, *K*_*da*_, is added to the reward-growth equation in the drug condition of the experiment:

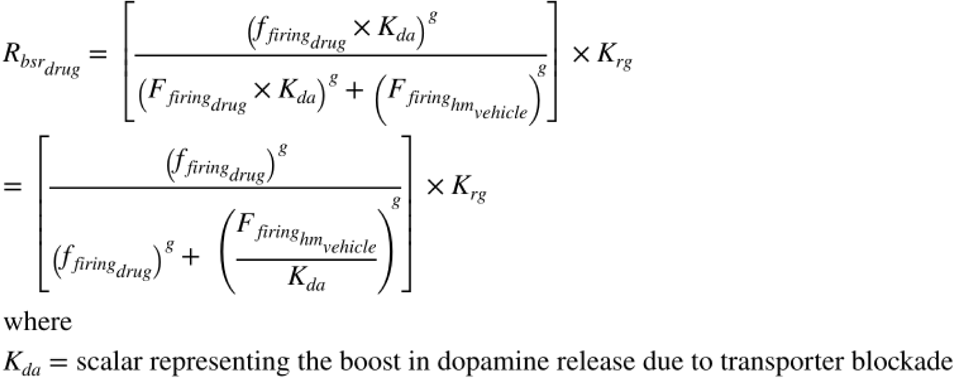

~~~
[eqn_num, eqn_tab] = add_eqn(eqn_num, eqn_tab, “RGfunDrug”, “reward-growth equation for the drug condition”);
~~~

`This is equation #14: (RGfunDrug)`

~~~
[sym_num, sym_tab] = add_sym(sym_num, sym_tab, “Kda”, “scalar representing the boost in dopamine release due to transporter blockade”);
~~~

Thus,

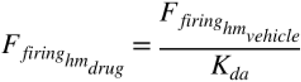

~~~
[eqn_num, eqn_tab] = add_eqn(eqn_num, eqn_tab, “FhmDrug”, “FhmDrug as a function of FhmVehicle”);
~~~

`This is equation #15: (FhmDrug)`

The following graphs plot the reward-growth function for three values of each parameter:

~~~
logFelecBend = 1.3222;
FelecBend = 10^
logFelecBend; logFelecRO = 2.5587;
FelecRO = 10^logFelecRO;
numF = 121; % Number of pulse frequencies in
Felec vector Felec = logspace(0,3,numF);
FelecMat = repmat(logspace(0,3,numF),3,1); % add two rows in preparation for graphing
Fhm1 = 10^1.6; % The number of elements in the parameter vector must equal the # of rows in FelecMat Fhm2 = 10^1.8;
Fhm3 = 10^2.0;
FhmVec = [Fhm1; Fhm2; Fhm3];
FhmMat = repmat(FhmVec,1,numF); % Store the Fhm values in a matrix of the same size as FelecMat
gElec1 = 5; % The number of elements in the parameter vector must equal the # of rows in FelecMat
gElec2 = 10;
gElec3 = 20;
gElecVec = [gElec1; gElec2; gElec3];
gElecMat = repmat(gElecVec,1,numF); % Store the gElec values in a matrix of the same size as FelecMat
Krg1 = 10^-0.2; % The number of elements in the paramer vector must equal the # of rows in FelecMat
Krg2 = 10^0;
Krg3 = 10^0.2;
KrgVec = [Krg1; Krg2; Krg3];
KrgMat = repmat(KrgVec,1,numF); % Store the Krg values in a matrix of the same size as FelecMat
RelecMat_Fhm = fRbsr(FelecMat, FelecBend, FhmMat, FelecRO, gElec1, Krg2);
RelecMat_gElect = fRbsr(FelecMat, FelecBend, Fhm2, FelecRO, gElecMat, Krg2);
RelecMat_Krg = fRbsr(FelecMat, FelecBend, Fhm2, FelecRO, gElec1, KrgMat);
TitleStrSemi = ‘semi-log’;
TitleStrLogLog = ‘log-log’;
pnam = “F_{hm}”;
fnam = “Fhm”;
% The data to be plotted must be in columns. Thus, FelecMat and RelecMat are transposed.
RG_Fhm_semilog = plot_RG(FelecMat’,RelecMat_Fhm’,pnam,FhmVec,fnam,TitleStrSemi,’lin’);
RG_Fhm_loglog = plot_RG(FelecMat’,RelecMat_Fhm’,pnam,FhmVec,fnam,TitleStrLogLog,’log’);
RG_dual_Fhm = dual_subplot(RG_Fhm_semilog, RG_Fhm_loglog, ‘RG_Fhm_semilog_loglog’,…
   graphs2files,FigDir);
if show_graphics
   RG_dual_Fhm.Visible = ‘on’;
end
~~~

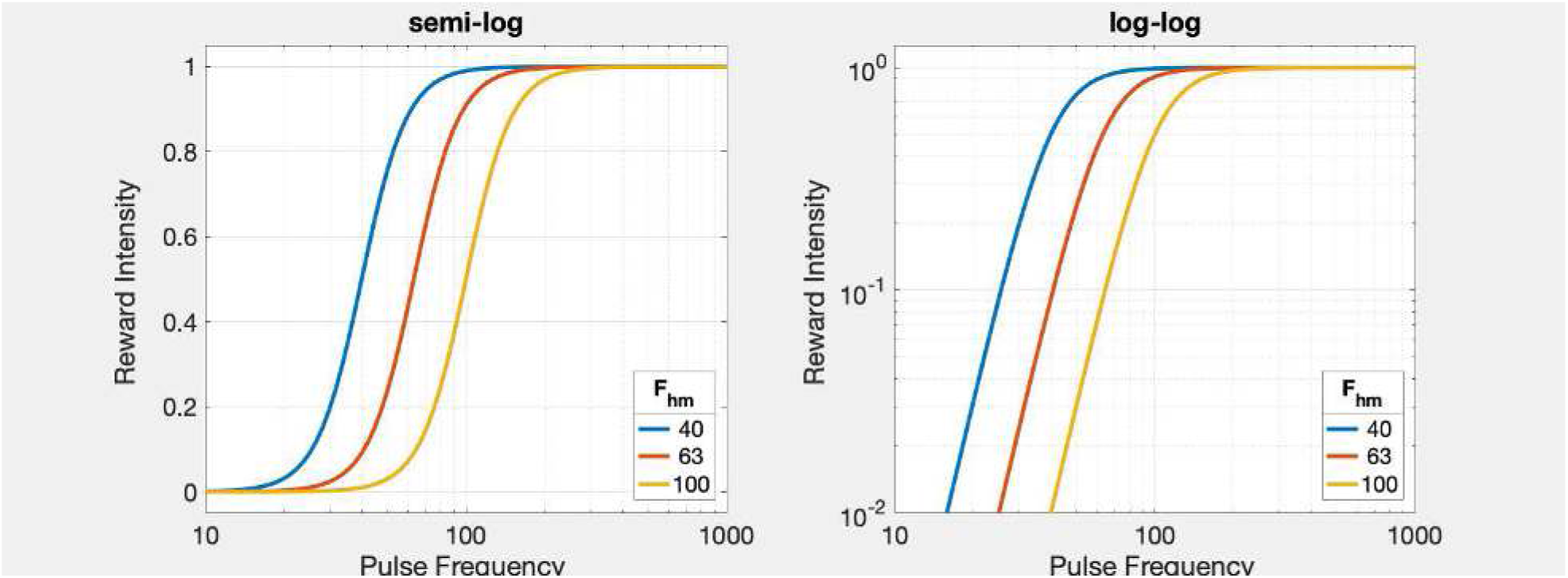

~~~
[fig_num, fig_tab] = add_fig(fig_num, fig_tab, “RGfunsElecFhm”, …
  “Growth of reward intensity at three values of the position parameter”);
~~~

`This is figure #3: (RGfunsElecFhm)`

~~~
[fun_num, fun_tab] = add_fun(fun_num, fun_tab, “plot_RG”, …
  “Fmat,RMat,pnam,pVec,fnam,TitleStr,linlog, varargins”,…
  “function to plot a single reward-growth function”);
~~~

`This is function #24: (plot_RG)`

~~~
[fun_num, fun_tab] = add_fun(fun_num, fun_tab, “dual_subplot”, …
  “g1, g2, dual_sub_out, graphs2files, figdir”,…
  “Function to plot two graphs side-by-side”);
~~~

`This is function #25: (dual_subplot)`

The semi-log plot is on the left, and the log-log plot is on the right.

These graphs shows how the position parameter rescales the ***input*** required to drive reward intensity to a particular level, thus sliding the reward-growth functions laterally.

~~~
pnam = “g”;
fnam = “g”;
RG_gElec_semilog = plot_RG(FelecMat’,RelecMat_gElect’,pnam,gElecVec,fnam,TitleStrSemi,’lin’);
RG_gElec_loglog = plot_RG(FelecMat’,RelecMat_gElect’,pnam,gElecVec,fnam,TitleStrLogLog,’log’);
RG_dual_g = dual_subplot(RG_gElec_semilog, RG_gElec_loglog, ‘RG_gElec_semilog_loglog’,…
  graphs2files, FigDir);
if show_graphics
  RG_dual_g.Visible = ‘on’;
end
~~~

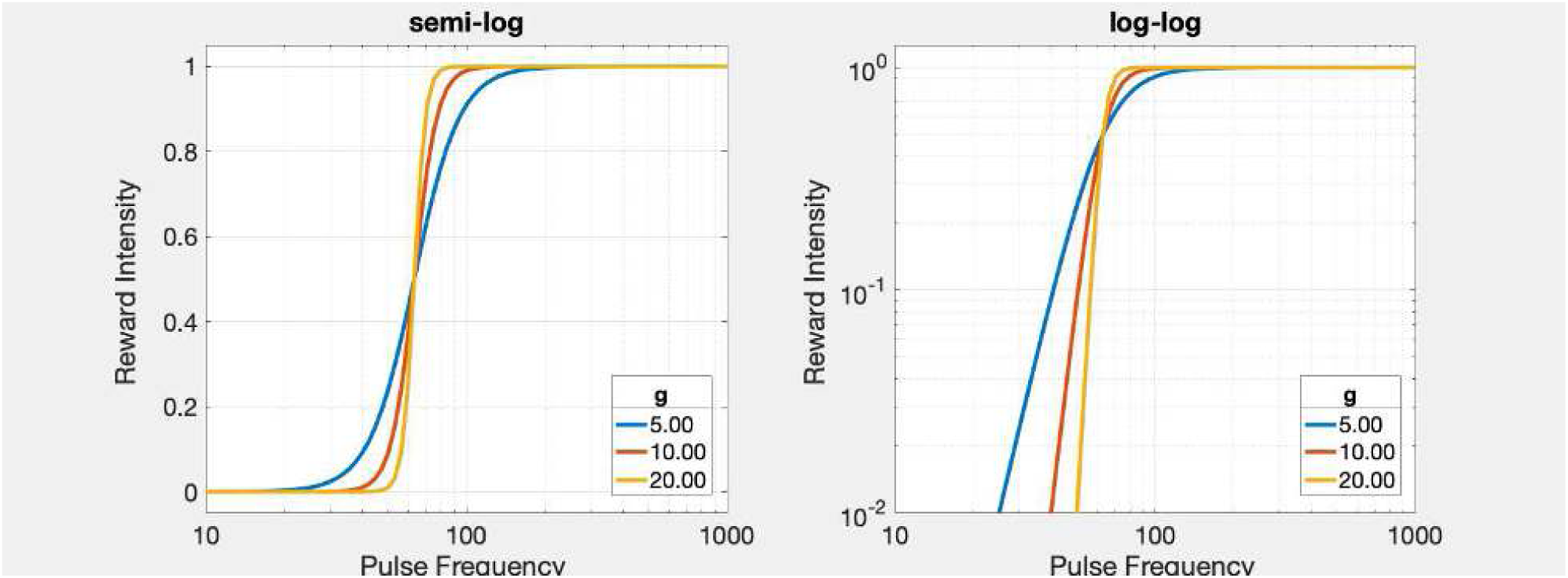

~~~
[fig_num, fig_tab] = add_fig(fig_num, fig_tab, “RGfunsElecG”, …
  “Growth of reward intensity at three values of the steepness parameter”);
~~~

`This is figure #4: (RGfunsElecG)`

The semi-log plot is on the left, and the log-log plot is on the right.

These graphs shows how the steepness parameter rotates the reward-growth functions around their midpoint.

~~~
pnam = “K_{rg}”;
fnam = “Krg”;
RG_Krg_semilog = plot_RG(FelecMat’,RelecMat_Krg’,pnam,KrgVec,fnam,TitleStrSemi,’lin’);
RG_Krg_loglog = plot_RG(FelecMat’,RelecMat_Krg’,pnam,KrgVec,fnam,TitleStrLogLog,’log’);
RG_dual_Krg = dual_subplot(RG_Krg_semilog, RG_Krg_loglog, ‘RG_Krg_semilog_loglog’,…
  graphs2files, FigDir);
if show_graphics
  RG_dual_Krg.Visible = ‘on’;
end
~~~

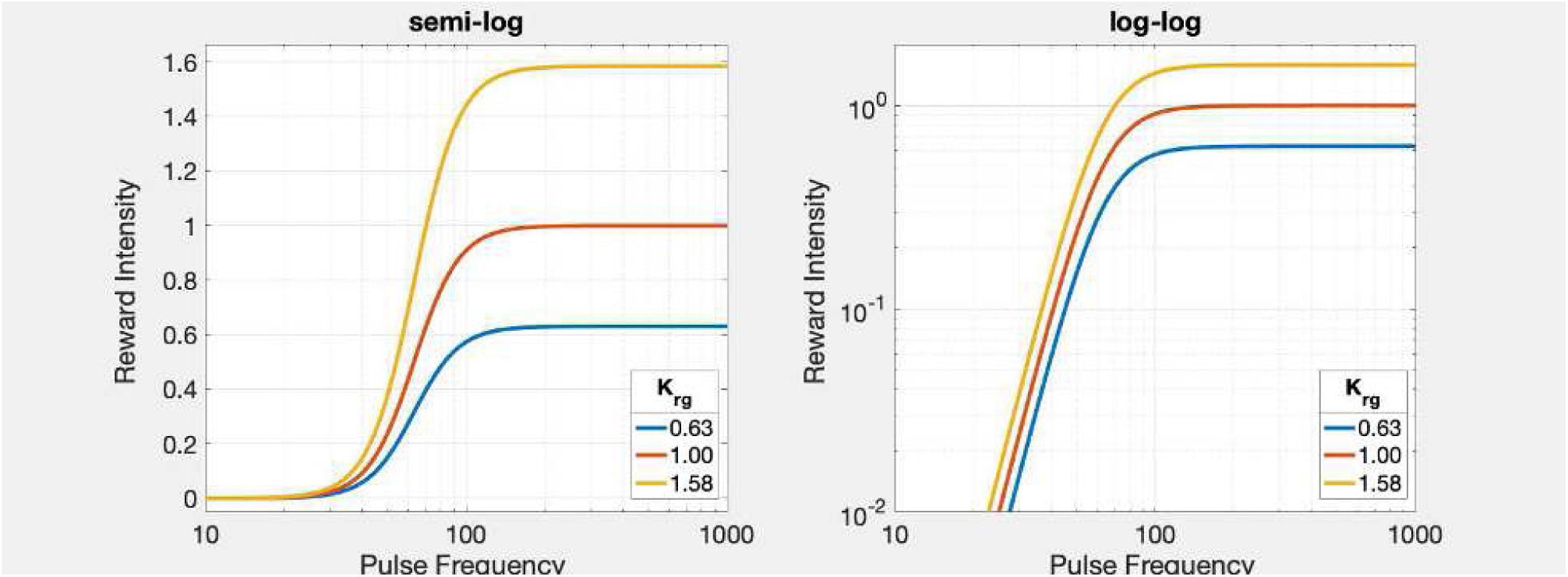

~~~
[fig_num, fig_tab] = add_fig(fig_num, fig_tab, “RGfunsElecKrg”, …
  “Growth of reward intensity at three values of the output-scaling parameter”);
~~~

`This is figure #5: (RGfunsElecKrg)`

These graphs show how the parameter, (*K*_*rg*_), rescales the ***output*** of the reward-growth function vertically, thus determining its asymptotic level.

The preceding graphs show how rescaling of the input to and output from the reward-growth function produce orthogonal changes. This is true as long as as the firing of the directly stimulated neurons keeps pace with the pulse frequency. However, once the value of the position parameter nears the maximum firing frequency of the directly stimulated neurons (e.g., when the current is very low), rightward shifts of the reward-growth function, such as those produced by further decreases in current, are accompanied by decreases in its upper asymptote. The neurons can no longer fire fast enough to drive reward intensity to the same maximum as was achieved when the value of the position parameter was lower (e.g., because the current was higher). This has implications for the interpretation of changes in the location parameters of the reward mountain, as discussed below.

~~~
Fhm4 = 10^2.0;
Fhm5 = 10^2.3;
Fhm6 = 10^2.6
~~~

`Fhm6 = 398.1072`

~~~
FhmVec = [Fhm4; Fhm5; Fhm6];
FhmMat = repmat(FhmVec,1,numF); % Store the Fhm values in a matrix of the same size as FelecMat
RelecMat_FhmHi = fRbsr(FelecMat, FelecBend, FhmMat, FelecRO, gElec1, Krg2);
TitleStrSemi = ‘HiF-semi-log’;
TitleStrLogLog = ‘HiF-log-log’;
pnam = “F_{hm}”;
fnam = “FhmHi”;
% The data to be plotted must be in columns. Thus, FelecMat and RelecMat are transposed.
RG_FhmHi_semilog = plot_RG(FelecMat’,RelecMat_FhmHi’,pnam,FhmVec,fnam,TitleStrSemi,’lin’);
RG_FhmHi_loglog = plot_RG(FelecMat’,RelecMat_FhmHi’,pnam,FhmVec,fnam,TitleStrLogLog,’log’);
RG_dual_FhmHi = dual_subplot(RG_FhmHi_semilog, RG_FhmHi_loglog, ‘RG_FhmHi_semilog_loglog’,…
   graphs2files,FigDir);
if show_graphics
   RG_dual_FhmHi.Visible = ‘on’;
end
~~~

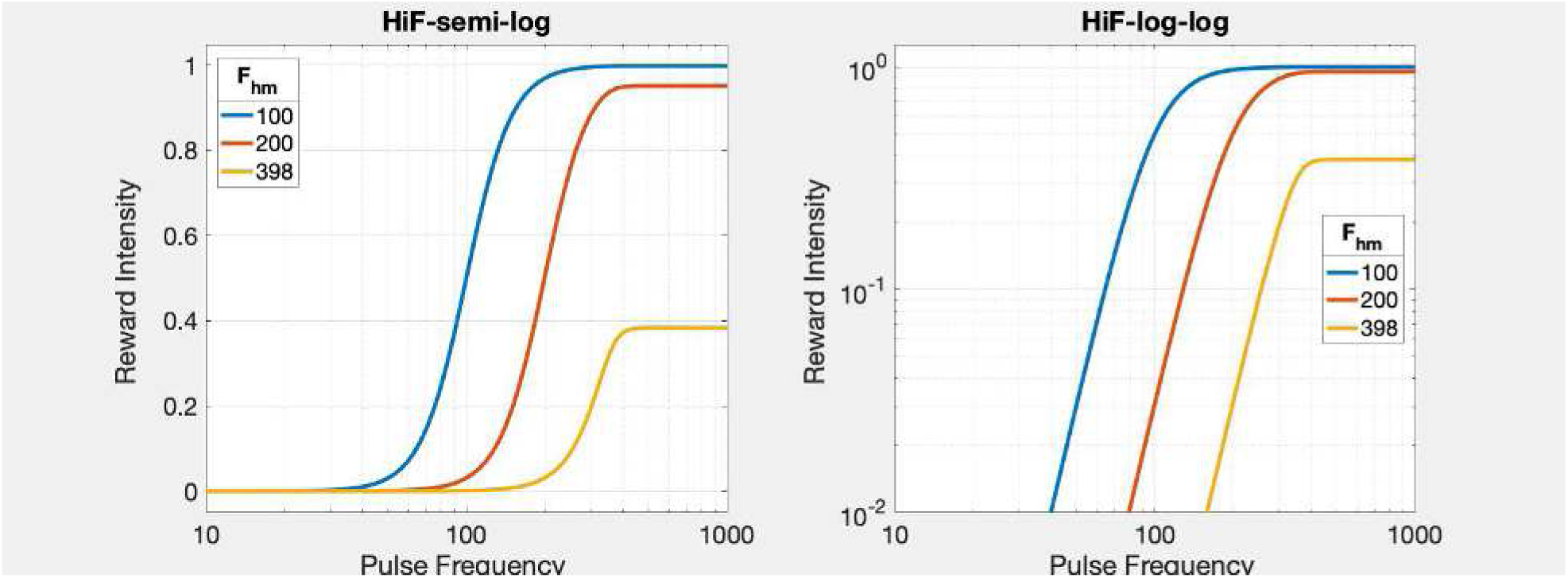

~~~
[fig_num, fig_tab] = add_fig(fig_num, fig_tab, “RGfunsElecFhmHi”, …
  “Growth of reward intensity at three values of the position parameter near FelecRO”);
~~~

`This is figure #6: (RGfunsElecFhmHi)`

When the middle of the frequency roll-off zone 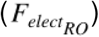 is positioned at 362 pulses *s*^−1^ (the median value reported by Solomon et al., 2015) and *F*_*hm*_ is 100 pulses *s*^−1^, the normalized reward-growth function approaches an upper asymptote of one, as expected. However, doubling *F*_*hm*_ to ∼200 pulses *s*^−1^ moderately decreases the upper asymptote. A further doubling to ∼400 pulses *s*^−1^ pushes *F*_*hm*_ past 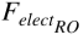, markedly truncating the growth of reward intensity.

~~~
close all;
~~~

### The location parameter of the reward-growth function for BSR

The location parameter of the reward-growth function is the firing rate that drives reward intensity to half its maximal value. The value of this parameter depends on the number of stimulated first-stage neurons, *N*, and the interval during which the stimulation train elevates their firing rate, *D*_*burst*_.

A prior study of temporal integration in the neural circuitry responsible for eICSS of the MFB (Sonnenschein et al., 2003) implies the following form for the function that determines 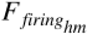;

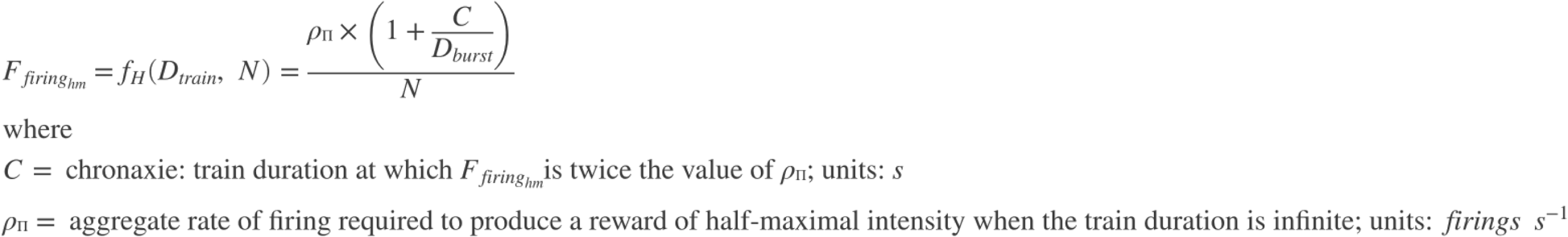

~~~
[eqn_num, eqn_tab] = …
   add_eqn(eqn_num, eqn_tab, “fHfiring”, “strengh-duration function for trains expressed as a firing frequency”);
~~~

`This is equation #16: (fHfiring)`

~~~
[fun_num, fun_tab] = add_fun(fun_num, fun_tab, “fHfiring”, “C, D, N, RhoPi, varargin”,…
   “strengh-duration function for trains expressed as a firing frequency”);
~~~

`This is function #26: (fHfiring)`

~~~
[eqn_num, eqn_tab] = …
  add_eqn(eqn_num, eqn_tab, “fH”, “strengh-duration function for trains expressed as an aggregate firing frequency”);
~~~

`This is equation #17: (fH)`

~~~
[fun_num, fun_tab] = add_fun(fun_num, fun_tab, “fH”, “C, D, RhoPi”,…
  “strengh-duration function for trains expressed as an aggreagate firing frequency”);
~~~

`This is function #27: (fH)`

~~~
[sym_num, sym_tab] = add_sym(sym_num, sym_tab, “Cmfb”, “chronaxie of the strength-duration function for trains”);
[sym_num, sym_tab] = add_sym(sym_num, sym_tab, “Dtrain”, “duration of an electrical pulse train”);
~~~

`Symbol Dtrain has already been entered.`

~~~
[sym_num, sym_tab] = add_sym(sym_num, sym_tab, “RhoPi”, “rheobase of the strength-duration function for trains”);
[fun_num, fun_tab] = add_fun(fun_num, fun_tab, “fH”, “C, D, rhoPi”,…
  “Strength-duration function for trains; generates aggregate firing rate required to produce half-maximal reward intensity”);
~~~

`Function fH has already been entered.`

We assume that *D*_*train*_. = *D*_*burst*_

The counter model is implicit in the equation for 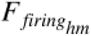. According to this model, the aggregate rate of firing in the directly stimulated neurons determines the intensity of the rewarding effect.

### The payoff from work

In keeping with the generalized matching law (Killeen, 1972), the benefit from work and its costs are combined in scalar fashion to yield a net payoff. We can format the expression for the payoff as a ratio of two rates (see: Gallistel & Gibbon, 2000):

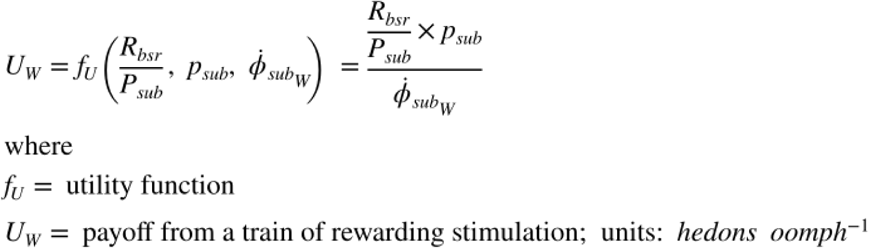

~~~
[eqn_num, eqn_tab] = add_eqn(eqn_num, eqn_tab, “fUW”, “payoff from work”);
~~~

`This is equation #18: (fUW)`

~~~
[sym_num, sym_tab] = add_sym(sym_num, sym_tab, “fU”, “utility function”);
[sym_num, sym_tab] = add_sym(sym_num, sym_tab, “UW”, “payoff from work”);
~~~

or as a benefit/cost ratio:

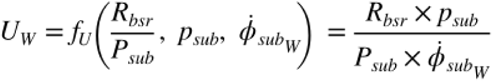

### The reward-growth function for leisure activities

We define a reward rate for the leisure activities that compete with pursuit of BSR as follows:

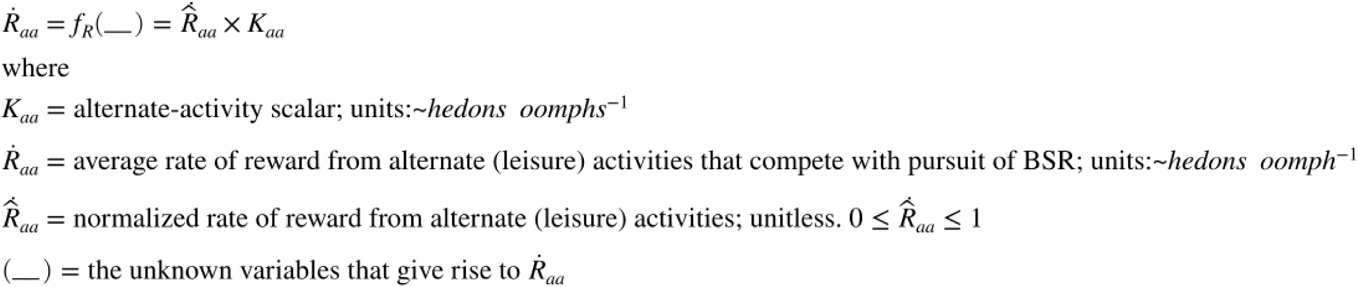

~~~
[eqn_num, eqn_tab] = add_eqn(eqn_num, eqn_tab, “fRaa”, “Leisure reward rate”);
~~~

`This is equation #19: (fRaa)`

~~~
[sym_num, sym_tab] = add_sym(sym_num, sym_tab, “Kaa”, “subjective reward-rate scalar”);
[sym_num, sym_tab] = add_sym(sym_num, sym_tab, “DotRaa”, “Average subjective reward rate from leisure activities”);
[sym_num, sym_tab] = add_sym(sym_num, sym_tab, “DotHatRaa”, “normalized average subjective reward rate from leisure activities”); [fun_num, fun_tab] = add_fun(fun_num, fun_tab, “fRaa”, “Raa,Kaa”,…
  “reward-growth function for leisure activities”);
~~~

`This is function #28: (fRaa)`

### The payoff from leisure activities

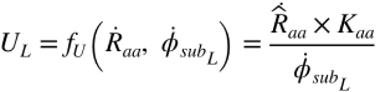

[eqn_num, eqn_tab] = add_eqn(eqn_num, eqn_tab, “fUL”, “payoff from leisure”);

`This is equation #20: (fUL)`

[sym_num, sym_tab] = add_sym(sym_num, sym_tab, “UL”, “payoff from leisure activities”);

### The core → shell function

#### The behavioral-allocation function

The payoffs from pursuit of BSR and engagement in leisure activities are used by the sole core→shell function to compute the allocation of time to pursuit of BSR.

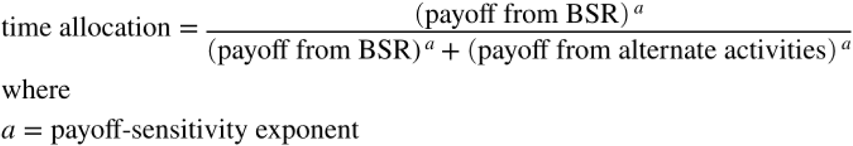

~~~
[sym_num, sym_tab] = add_sym(sym_num, sym_tab, “a”, “payoff-sensitivity exponent”);
~~~

The exponent (*a*) determines how abruptly time allocation changes as a function of changes in payoff. Restating this equation symbolically, we obtain

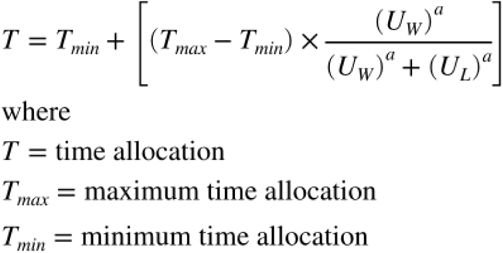

~~~
[eqn_num, eqn_tab] = …
  add_eqn(eqn_num, eqn_tab, “TU”, “Time allocation defined in terms of payoffs”);
~~~

`This is equation #21: (TU)`

~~~
[sym_num, sym_tab] = add_sym(sym_num, sym_tab, “T”, “time allocation”);
[sym_num, sym_tab] = add_sym(sym_num, sym_tab, “Tmax”, “maximum time allocation”);
[sym_num, sym_tab] = add_sym(sym_num, sym_tab, “Tmin”, “minimum time allocation”);
~~~

We also define a normalized measure of time allocation

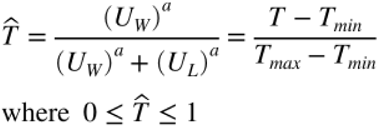

~~~
[eqn_num, eqn_tab] = …
  add_eqn(eqn_num, eqn_tab, “Tnorm”, “Normalized time allocation”);
~~~

`This is equation #22: (Tnorm)`

as well as a value of time allocation, *T*_*mid*_, at which the payoffs from work and leisure are equal, and thus, time allocation falls halfway between its minimal and maximal values:

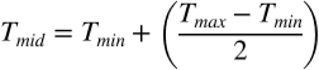

~~~
[eqn_num, eqn_tab] = …
 add_eqn(eqn_num, eqn_tab, “Tmid”, “Mid-range time allocation”);
~~~

`This is equation #23: (Tmid)`

When the reward intensity produced by the stimulation approaches its upper asymptote, the price at which *T* = *T*_*mid*_ is:

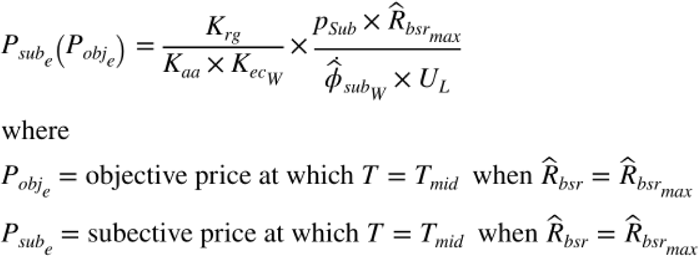

~~~
[eqn_num, eqn_tab] = …
  add_eqn(eqn_num, eqn_tab, “Psub_e”, “Definition of Psub_e”);
~~~

`This is equation #24: (Psub_e)`

~~~
[fun_num, fun_tab] = add_fun(fun_num, fun_tab, “PsubEfun”, …
  “dotPhiObj, Kaa, Kec, Krg, Raa, pObj, RnormMax, varargin”,…
  “function to compute PsubE”);
~~~

`Function PsubEfun has already been entered.`

To obtain an expression for 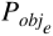, we back-solve the subjective-price equation, as shown above in

~~~
disp(string({strcat({‘Equation ‘},num2str(eqn_tab.Number(eqn_tab.Name==‘fPbacksolved’)))}));
~~~

`Equation 7`

~~~
[fun_num, fun_tab] = add_fun(fun_num, fun_tab, “PobjEfun”, …
  “dotPhiObj, Kaa, Kec, Krg, ObjAA, pObj, PsubBend, PsubMin, RnormMax, varargin”,…
  “Back-solution of the subjective-price function to return PobjE from PsubE”);
~~~

`Function PobjEfun has already been entered.`

We can now define the equation for normalized time allocation as:

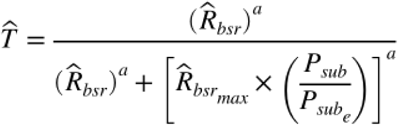

~~~
[eqn_num, eqn_tab] = …
  add_eqn(eqn_num, eqn_tab, “TnormExpand”, “Normalized time allocation expanded”);
~~~

`This is equation #25: (TnormExpand)`

and the full equation for time allocation as:

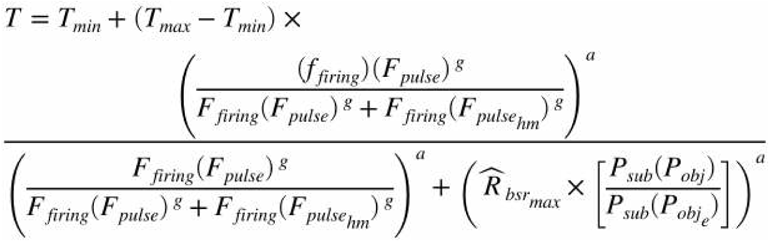

~~~
[eqn_num, eqn_tab] = …
  add_eqn(eqn_num, eqn_tab, “Tfull”, “Full 6-parameter time-allocation equation”);
~~~

`This is equation #26: (Tfull)`

~~~
[fun_num, fun_tab] = add_fun(fun_num, fun_tab, “TAfun”, “a, F, Fbend, FpulseHM, Fro, g, Pobj, PobjE, PsubBend, PsubMin, RnormMax”,… “
6-parameter TA function”);
~~~

`This is function #29: (TAfun)`

### The reward mountain

~~~
disp(string({strcat({‘Equation ‘},num2str(eqn_tab.Number(eqn_tab.Name==‘Tfull’)))}));
~~~

`Equation 26`

defines the surface of the reward-mountain model in a three-dimensional space defined by the two independent variables, the pulse frequency (*F*_*pulse*_) and the price (*P*_*obj*_), and a single dependent variable, time allocation (*T*). Thus, this equation bridges the core of the model by linking three shell variables, two inputs and one output; the bridge is constructed from the shell → core, core, and core → shell functions. The slope of the mountain surface depends on the payoff-sensitivity parameter and the steepness parameter (*a*) of the reward-growth equation, (*g*). The location of the mountain in the plane defined by the independent variables is determined by the two location parameters: whereas *P*_*obj_e*_ positions the mountain along the price axis, 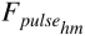 positions the mountain along the pulse-frequency axis. Finally, the altitude of the “valley floor” and “summit” are set by *T*_*min*_ and *T*_*max*_, respectively.

To simulate the mountain model, we must obtain values for the position parameters. 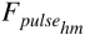. We proceed as follows:

~~~
C = 0.473; % median from Sonnenschein et al., 2003
Dtrain = 0.5; % train duration in eICSS reward-mountain studies
logFelecBend = 1.3222; % from Solomon et al., 2015
FelecBend = 10^logFelecBend;
logFelecRO = 2.5587; % from Solomon et al., 2015 FelecRO = 10^logFelecRO;
FPmax =1000; % for determining RbsrNormMax. This value is sufficiently above FelecRO to maximize the firing rate
N = 100; % arbitrary
RhoPi = 5000; % arbitrary
% RhoPi/N = 50, which is roughly consistent with Sonnenchein et al., 2003
FmfbHM = FpulseHMfun(C, Dtrain, FelecBend, FPmax, FelecRO, N, RhoPi)
~~~

`FmfbHM = 97.3001`

~~~
[fun_num, fun_tab] = add_fun(fun_num, fun_tab, “FpulseHMfun”, …   “C, D, Fbend, FPmax, Fro, N, RhoPi, varargin”,…
  “Function to compute the pulse frequency that produces half-maximal reward intensity”);
~~~

`This is function #30: (FpulseHMfun)`

The parameter that determines the location of the mountain along the price axis is *P*_*obj_e*_.

~~~
dotPhiObj = 1;
Kaa = 1;
Kec = 1;
Krg = 1;
dotRaa = 0.1;
pObj = 1;
PsubBend = 0.5;
PsubMin = 1.82;
~~~

Before computing 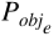, we must determine the maximum value of the normalized reward intensity 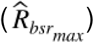.

If the value of the position parameter of the reward-growth function is too high given the frequency roll-off parameters, 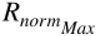 will be less than one.

~~~
[sym_num, sym_tab] = add_sym(sym_num, sym_tab, “RnormMax”, “maximum normalized reward intensity”);
g = 5;
%RnormMax = RnormMaxFun(Fmax, FelecBend, FmfbHM, FelecRO, g)
RnormMax = fRbsrNorm(FPmax, FelecBend, FmfbHM, FelecRO, g)
~~~

`RnormMax = 0.9986`

~~~
PobjE = PobjEfun(dotPhiObj, Kaa, Kec, Krg, dotRaa, pObj, PsubBend, PsubMin, RnormMax)
~~~

`PobjE = 9.9860`

We will now assign a value to the price-sensitivity parameter, *a*, and will simulate the mountain surface. (See ***Functions composing the reward-mountain model*** below.)

~~~
a = 3;
Felec = logspace(0,3,121)’; % column variable
Pobj = logspace(0,3,121); % row variable
Tmfb = TAfun(a, Felec, FelecBend, FmfbHM, FelecRO, g, Pobj, PobjE, PsubBend, PsubMin, RnormMax);
% n.b., numel(Felec) = numel(Pobj). Felec has been transposed. Thus, Tmfb is a square matrix.
MTN = plot_MTN(Felec, Pobj, Tmfb, ‘off’, ‘MTN’, ‘reward mountain’, …
  graphs2files, FigDir); if show_graphics
MTN.Visible = ‘on’;
  end
~~~

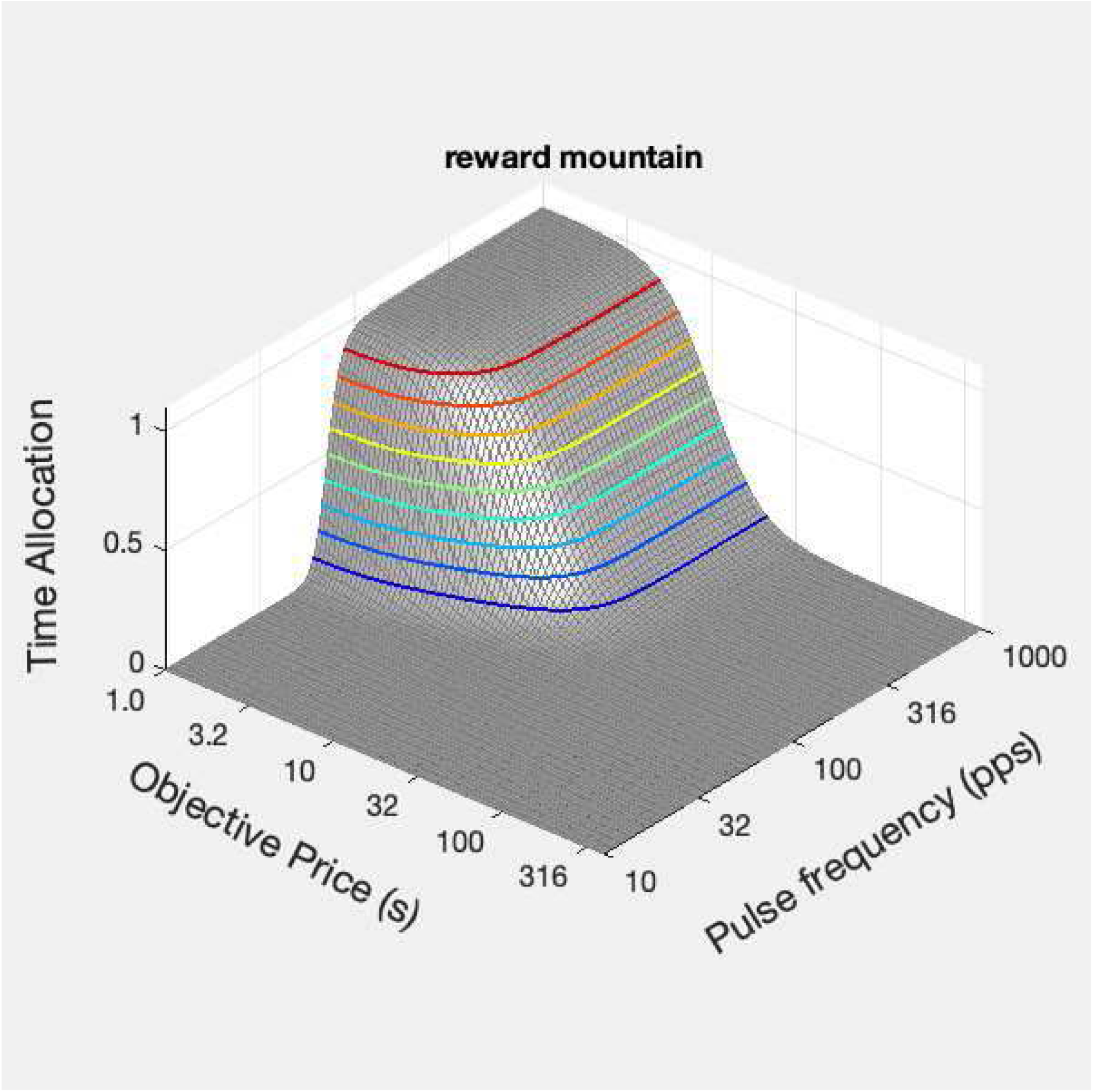

~~~
[fig_num, fig_tab] = add_fig(fig_num, fig_tab, “Mountain”, …
  “3D plot of the reward mountain”);
~~~

`This is figure #7: (Mountain)`

~~~
[fun_num, fun_tab] = add_fun(fun_num, fun_tab, “plot_MTN”, …
  “F, Pobj, T, Visible, mtn_root, title_str, graphs2files, figdir, varargin”,…
  “Function to plot a single reward mountain”);
~~~

`This is function #31: (plot_MTN)`

~~~
clearvars(‘-except’,keepVars{:})
keepVars = who; % Restore cell array containing names of variables to be retained
toc
~~~

`Elapsed time is 25.821254 seconds.`

## Moving the mountain: validation studies

~~~
tic;
close all;
~~~

We have carried out a series of experiments that test the mountain model. Given that there are two independent position parameters 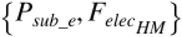, it should be possible to move the mountain independently along either of the axes representing the objective price and the pulse frequency. This prediction has been tested in three ways.

### Effect of varying the stimulation current

Changing the stimulation current alters the number of reward-related axons recruited. According to

~~~
disp(string({strcat({‘Equation ‘},num2str(eqn_tab.Number(eqn_tab.Name==‘FmfbHM’)))}));
~~~

`Equation`

~~~
disp(string({strcat({‘Equation ‘},num2str(eqn_tab.Number(eqn_tab.Name==‘FelecHM’)))}));
~~~

`Equation`

this will change the value of the parameter that positions the mountain along the pulse-frequency axis 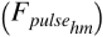, but according to

~~~
disp(string({strcat({‘Equation ‘},num2str(eqn_tab.Number(eqn_tab.Name==‘Psub_e’)))}));
~~~

`Equation 24`

and

`Equation 7`

changing the current will have no effect on the value of the parameter *P*_*obj_e*_ that positions the mountain along the price axis. We will now simulate the effect of changing the current by solving the time-allocation equation for two different values of *N*.

~~~
C = 0.473; % median from Sonnenschein et al., 2003 D = 0.5; % typical train duration
logFelecBend = 1.3222; % from Solomon et al., 2015 FelecBend = 10^logFelecBend;
logFelecRO = 2.5587; % from Solomon et al., 2015 FelecRO = 10^logFelecRO;
FPmax =1000; % for determining RbsrNormMax. This value is sufficiently above FelecRO to maximize the firing rate
N = [79,158]; % Number of neurons recruited by low & hi currents (2-element vector)
RhoPi = 5000; % arbitrary
FmfbHM = FpulseHMfun(C, D, FelecBend, FPmax, FelecRO, N, RhoPi) % FmfbHM is a 2-element vector
~~~

~~~
FmfbHM = 1×2
123.1648 61.5823
dotPhiObj = 1;
Kaa = 1;
Kec = 1;
Krg = 1;
dotRaa = 0.1;
pObj = 1;
PsubBend = 0.5;
PsubMin = 1.82;
FPmax = 1000; % This value is sufficiently above FelecRO to maximize the firing rate g = 5;
RnormMax = fRbsrNorm(FPmax, FelecBend, FmfbHM, FelecRO, g) % RnormMax is a 2-element vector
~~~

~~~
RnormMax = 1×2
0.9955 0.9999
~~~

~~~
PobjE = PobjEfun(dotPhiObj, Kaa, Kec, Krg, dotRaa, pObj, PsubBend, PsubMin, RnormMax)
~~~

~~~
PobjE = 1×2
9.9546 9.9986
~~~

~~~
% PobjE is a 2-element vector
a = 3;
Felec = logspace(0,3,121)’; % column variable Pobj = logspace(0,3,121);% row variable
Tmfb1 = TAfun(a, Felec, FelecBend, FmfbHM(1), FelecRO, g, Pobj, PobjE(1), …
  PsubBend, PsubMin, RnormMax(1));
Tmfb2 = TAfun(a, Felec, FelecBend, FmfbHM(2), FelecRO, g, Pobj, PobjE(2), …
  PsubBend, PsubMin, RnormMax(2));
% n.b., numel(Felec) = numel(Pobj). Felec has been transposed. Thus, Tmfb is a square matrix. MTNloI = plot_MTN(Felec, Pobj, Tmfb1, ‘off’, ‘MTNloI’, ‘low current’, …
  graphs2files, FigDir);
MTNhiI = plot_MTN(Felec, Pobj, Tmfb2, ‘off’, ‘MTNhiI’, ‘high current’, … graphs2files, FigDir);
dual_I_plot = dual_subplot(MTNloI, MTNhiI, ‘MTNloI_hiI’,… graphs2files,FigDir);
if show_graphics
  dual_I_plot.Visible = ‘on’;
end
~~~

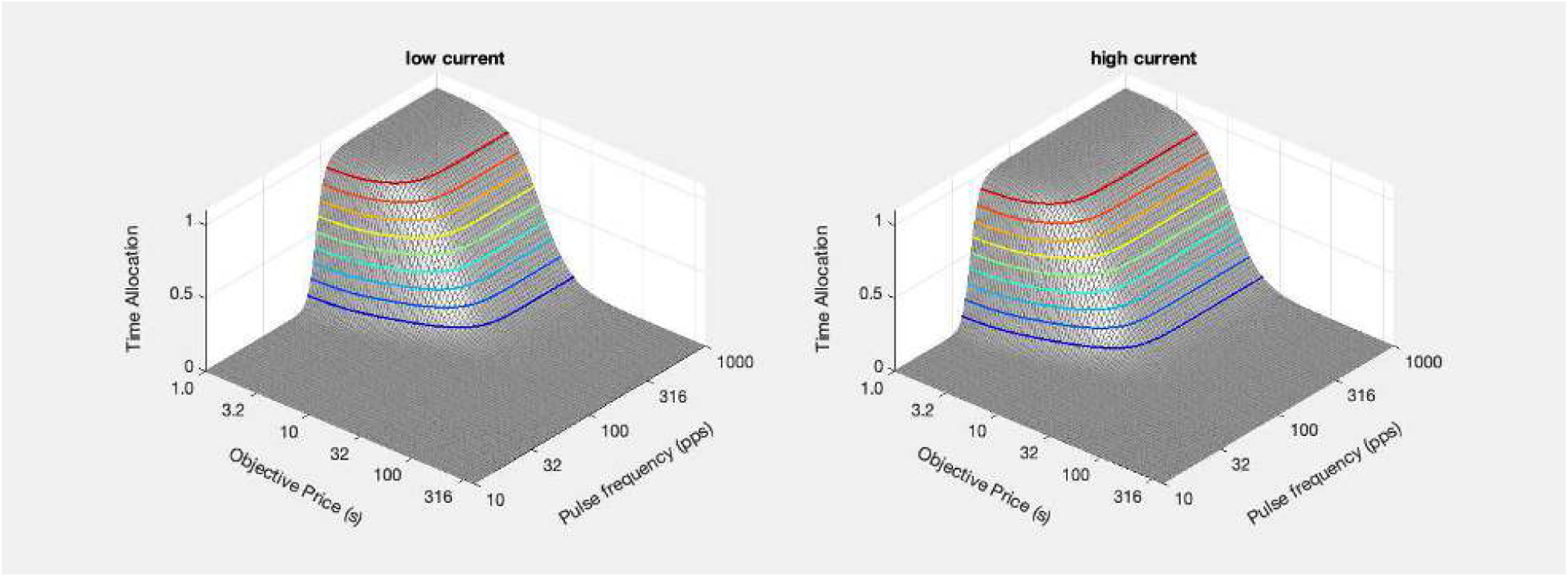

~~~
[fig_num, fig_tab] = add_fig(fig_num, fig_tab, “LoHiI_mtns”, …
  “Effect of changing the current on the reward mountain”);
~~~

`This is figure #8: (LoHiI_mtns)`

(See ***Functions that plot graphs and set attributes*** below.)

Although the change in the position of the mountain following and increase in the number of reward-related neurons recruited can be discerned readily in the 3D plots, the effect is best depicted by means of contour graphs and a bar graph showing the changes in the position paramter. The contour graphs capture in two dimensions all of the spatial information in the 3D plots.

~~~
ContLoI = plot_contour(Felec, Pobj, Tmfb1, PobjE(1), FmfbHM(1), ‘off’, ‘ContLoI’, ‘low current’, …
  strcat({‘N = ‘}, num2str(N(1))), graphs2files, FigDir);
ContHiI = plot_contour(Felec, Pobj, Tmfb2, PobjE(2), FmfbHM(2), ‘off’, ‘ContHiI’, ‘high current’, …
  strcat({‘N = ‘}, num2str(N(2))), graphs2files, FigDir);
bg_LoHiI = plot_bg(FmfbHM(1), FmfbHM(2), PobjE(1), PobjE(2), ‘off’, ‘bg_LoHiI’,…
  graphs2files, FigDir);
bg_root =‘bg_LoHiI’;
quad_I_plot = quad_subplot(ContLoI, ContHiI, bg_LoHiI, ‘quad_LoHiI’, bg_root, …
  graphs2files, FigDir);
if show_graphics
  quad_I_plot.Visible = ‘on’;
end
~~~

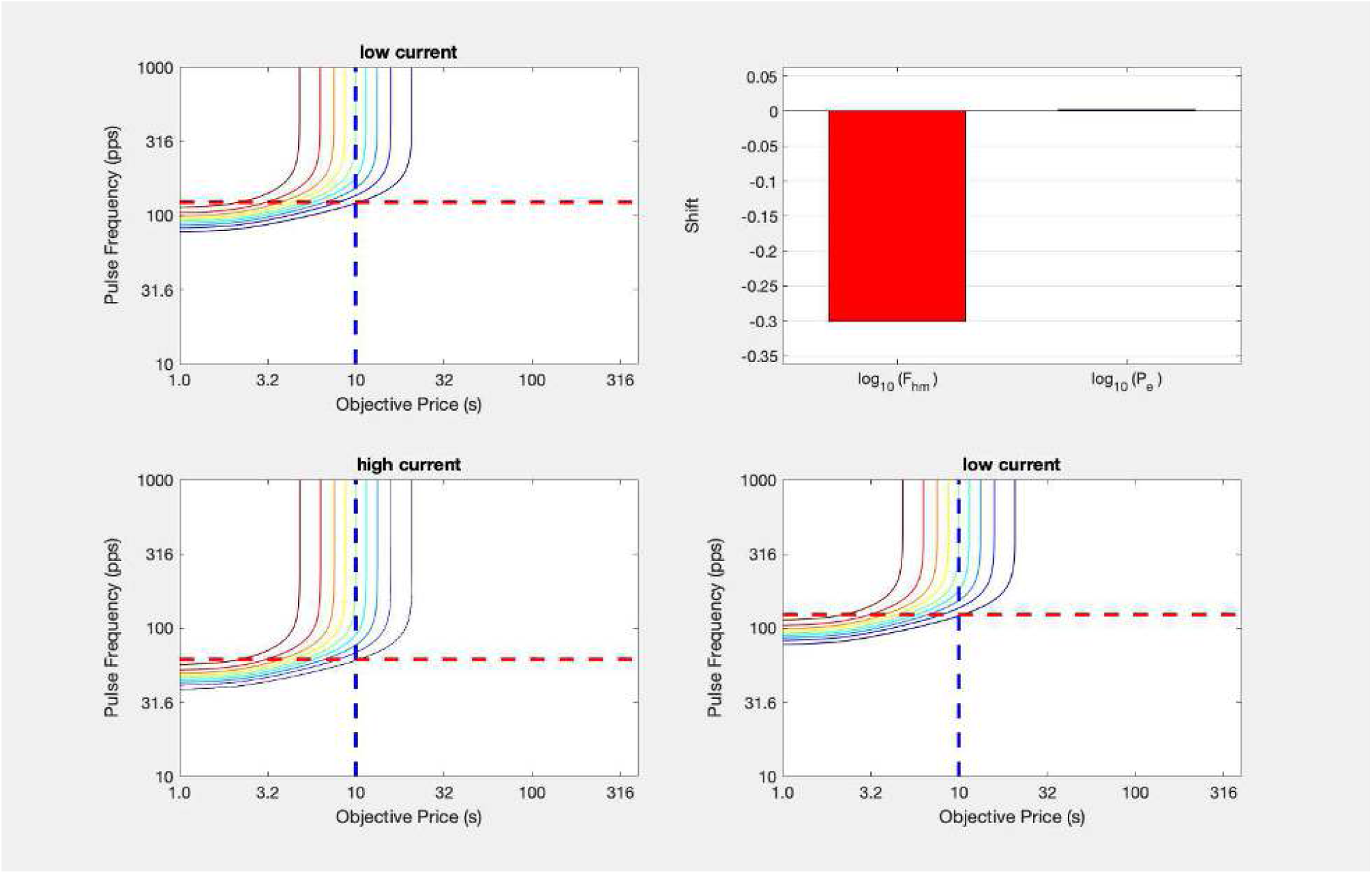

~~~
[fig_num, fig_tab] = add_fig(fig_num, fig_tab, “quad_LoHiI”, …
  “Effect of changing the number of recruited neurons on the position of the reward mountain”);
~~~

`This is figure #9: (quad_LoHiI)`

~~~
[fun_num, fun_tab] = add_fun(fun_num, fun_tab, “plot_contour”, …
  “F, Pobj, T, Pobj_e, Fhm, Visible, mtn_root, title_str, annot_str, graphs2files, figdir, varargin”,…
  “Function to plot the contour graph of a single mountain”);
~~~

`This is function #32: (plot_contour)`

~~~
[fun_num, fun_tab] = add_fun(fun_num, fun_tab, “quad_subplot”, …
  “cont1, cont2, bg, quad_sub_out, bg_root, graphs2files, figdir”,…
  “Function to plot four graphs in a 2 × 2 mosaic”);
~~~

`This is function #33: (quad_subplot)`

The contour graph for the low-current condition is shown twice, once in the upper-left panel and once in the lower-right panel. By comparing the horizontal position of the mountain in the left column, the reader can quickly discern whether there has been a shift along the price axis, and by comparing the vertical position of the mountains in the bottom row, the reader can quickly discern whether there has been a shift along the pulse-frequency axis. As required by the mountain model, the latter shift is observed when the number of neurons recruited is increased due to boost in the stimulation current. The bar graph in the upper right provides a summary of the simulated shifts. The tiny rightward shift along the price axis is due to the fact that is 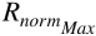 a little closer to one at the higher current than at the lower current (due to the fact that *F*_*hm*_ is lower).

Arvanitogiannis and Shizgal (2008) tested the effect of varying the stimulation current on the position of the reward mountain. In all four rats, the mountain shifted along the pulse-frequency axis as predicted. In two of these rats, there was no corresponding shift along the price axis, but rightward shifts were seen in the remaining two subjects. This deviation from the predictions was attributed to the heterogeneity of the stimulated neurons. Such a deviation would be expected if the stimulation activated two subpopulations of reward-related neurons that project to separate integrators with converging outputs (Arvanitogiannis, Waraczynski & Shizgal, 1996; Arvanitogiannis & Shizgal, 2008). The convergence model described below provides an example of such an arrangement.

~~~
clearvars(‘-except’,keepVars{:})
keepVars = who; % Restore cell array containing names of variables to be retained toc
~~~

`Elapsed time is 10.356255 seconds.`

### Effect of varying the train duration

~~~
tic;
close all;
~~~

In

~~~
disp(string({strcat({‘Figure ‘},num2str(fig_tab.Number(fig_tab.Name==‘quad_LoHiI’)))}));
~~~

Figure 9

the mountain shifts along the pulse-frequency axis because changing the current alters the denominator of the ratio on the right-hand side of

~~~
disp(string({strcat({‘Equation ‘},num2str(eqn_tab.Number(eqn_tab.Name==‘fH’)))}));
~~~

`Equation 17`

Changing the train duration should produce the same qualitative effect as changing the current, but by altering the numerator of the ratio on the right-hand side of

~~~
disp(string({strcat({‘Equation ‘},num2str(eqn_tab.Number(eqn_tab.Name==‘fH’)))}));
~~~

`Equation 17`

instead. By increasing the duration of the train, there is more time for temporal summation and thus, the pulse frequency required to produce a reward of half-maximal intensity decreases.

~~~
C = 0.473; % median from Sonnenschein et al., 2003
D = 0.5; % typical train duration
logFelecBend = 1.3222; % from Solomon et al., 2015
FelecBend = 10^logFelecBend;
logFelecRO = 2.5587; % from Solomon et al., 2015
FelecRO = 10^logFelecRO;
FPmax =1000; % for determining RbsrNormMax. This value is sufficiently above FelecRO to maximize the firing rate
N = 126;
D = [0.25,1.00]; % Short & long train durations (2-element vector)
RhoPi = 5000; % arbitrary
% RhoPi/N = 50, which is roughly consistent with Sonnenchein et al., 2003
29
FmfbHM = FpulseHMfun(C, D, FelecBend, FPmax, FelecRO, N, RhoPi) % FmfbHM is a 2-element vector
~~~

~~~
FmfbHM = 1×2
114.7621 58.4524
~~~

~~~
dotPhiObj = 1;
Kaa = 1;
Kec = 1;
Krg = 1;
dotRaa = 0.1;
pObj = 1;
PsubBend = 0.5;
PsubMin = 1.82;
g = 5;
RnormMax = fRbsrNorm(FPmax, FelecBend, FmfbHM, FelecRO, g) % RnormMax is a 2-element vector
~~~

~~~
RnormMax = 1×2
0.9968 0.9999
~~~

~~~
PobjE = PobjEfun(dotPhiObj, Kaa, Kec, Krg, dotRaa, pObj, PsubBend, PsubMin, RnormMax)
PobjE = 1×2
9.9681 9.9989
~~~

~~~
% PobjE is a 2-element vector
a = 3;
Felec = logspace(0,3,121)’; % column variable
Pobj = logspace(0,3,121); % row variable
Tmfb1 = TAfun(a, Felec, FelecBend, FmfbHM(1), FelecRO, g, Pobj, …
  PobjE(1), PsubBend, PsubMin, RnormMax(1));
Tmfb2 = TAfun(a, Felec, FelecBend, FmfbHM(2), FelecRO, g, Pobj, …
  PobjE(2), PsubBend, PsubMin, RnormMax(2));
MTNloD = plot_MTN(Felec, Pobj, Tmfb1, ‘off’, ‘MTNloD’, ‘short-duration train’, …
  graphs2files, FigDir);
MTNhiD = plot_MTN(Felec, Pobj, Tmfb2, ‘off’, ‘MTNhiD’, ‘long-duation train’, …
  graphs2files, FigDir);
dual_D_plot = dual_subplot(MTNloD, MTNhiD, ‘MTNLoHiD’,…
  graphs2files,FigDir);
if show_graphics
  dual_D_plot.Visible = ‘on’;
end
~~~

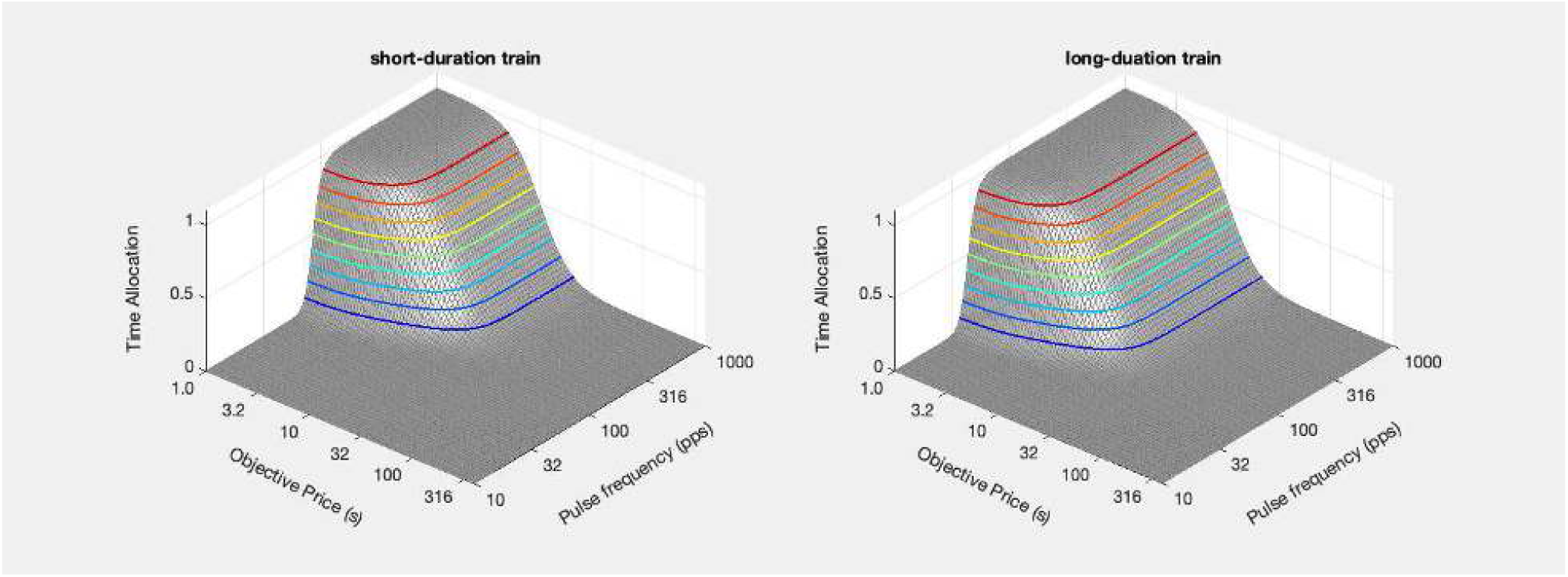

~~~
[fig_num, fig_tab] = add_fig(fig_num, fig_tab, “LoHiD_mtns”, …
  “Effect of changing the train duration on the reward mountain”);
~~~

`This is figure #10: (LoHiD_mtns)`

~~~
ContLoD = plot_contour(Felec, Pobj, Tmfb1, PobjE(1), FmfbHM(1), ‘off’, ‘ContLoD’, ‘short-duration train’, …
  strcat({‘D = ‘}, num2str(D(1))), graphs2files, FigDir);
ContHiD = plot_contour(Felec, Pobj, Tmfb2, PobjE(2), FmfbHM(2), ‘off’, ‘ContHiD’, ‘long-duation train’, …
  strcat({‘D = ‘}, num2str(D(2))), graphs2files, FigDir);
bg_LoHiD = plot_bg(FmfbHM(1), FmfbHM(2), PobjE(1), PobjE(2), ‘off’, ‘bg_LoHiD’,…
  graphs2files, FigDir);
bg_root =‘bg_LoHiD’;
quad_D_plot = quad_subplot(ContLoD, ContHiD, bg_LoHiD, ‘quad_LoHiD’, bg_root, …
  graphs2files, FigDir);
if show_graphics
  quad_D_plot.Visible = ‘on’;
end
~~~

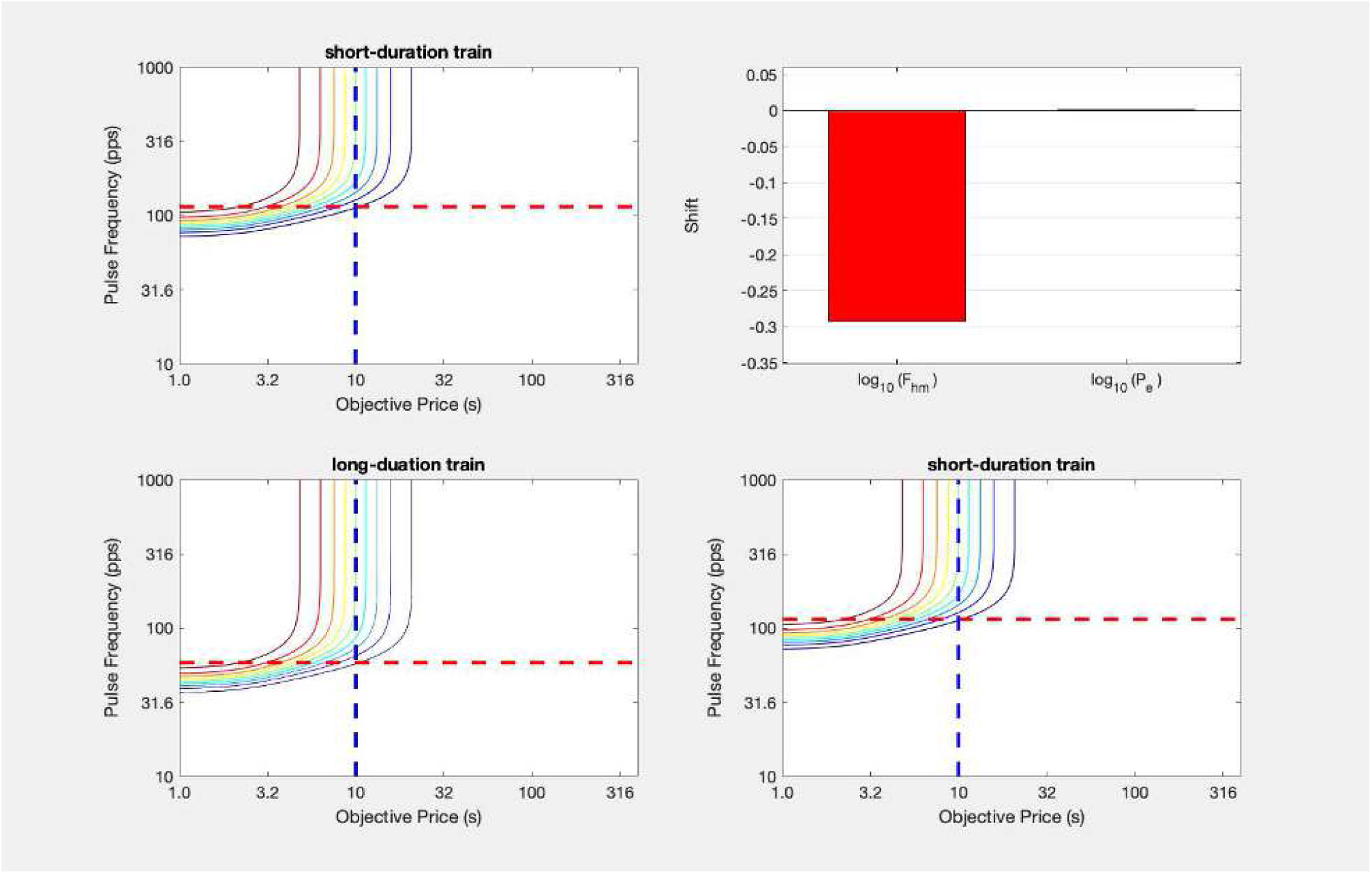

~~~
[fig_num, fig_tab] = add_fig(fig_num, fig_tab, “LoHiD_quad”, …
  “Effect of changing the train duration on the position of the reward mountain”);
~~~

`This is figure #11: (LoHiD_quad)`

~~~
clearvars(‘-except’,keepVars{:})
keepVars = who; % Restore cell array containing names of variables to be retained toc
~~~

`Elapsed time is 13.785165 seconds.`

~~~
tic;
close all;
~~~

Arvanitogiannis and Shizgal (2008) assessed the effect of increasing the train duration from 0.25 to 1.00 s in four rats. The data in all four cases correspond to the prediction: there were statistically reliable shifts of the reward mountain along the pulse-frequency axis but not along the price axis. The effect of increasing the train duration was tested in an additional six rats by Breton et al. (2014). In all six cases, the mountain shifted as predicted along the pulse-frequency axis, and in three of these cases, no reliable shifts along the price axis were observed. In the remaining three cases, increasing the train duration did produce rightward shifts along the price axis. These were again hypothesized to due to the recruitment of reward-related neurons projecting to separate integrators. The convergence model describe below offers a new interpretations of the rightward shifts.

~~~
if show_graphics
  show_imported_graphic(‘TD_quad_Y14.png’, 25, ImpFigDir);
~~~

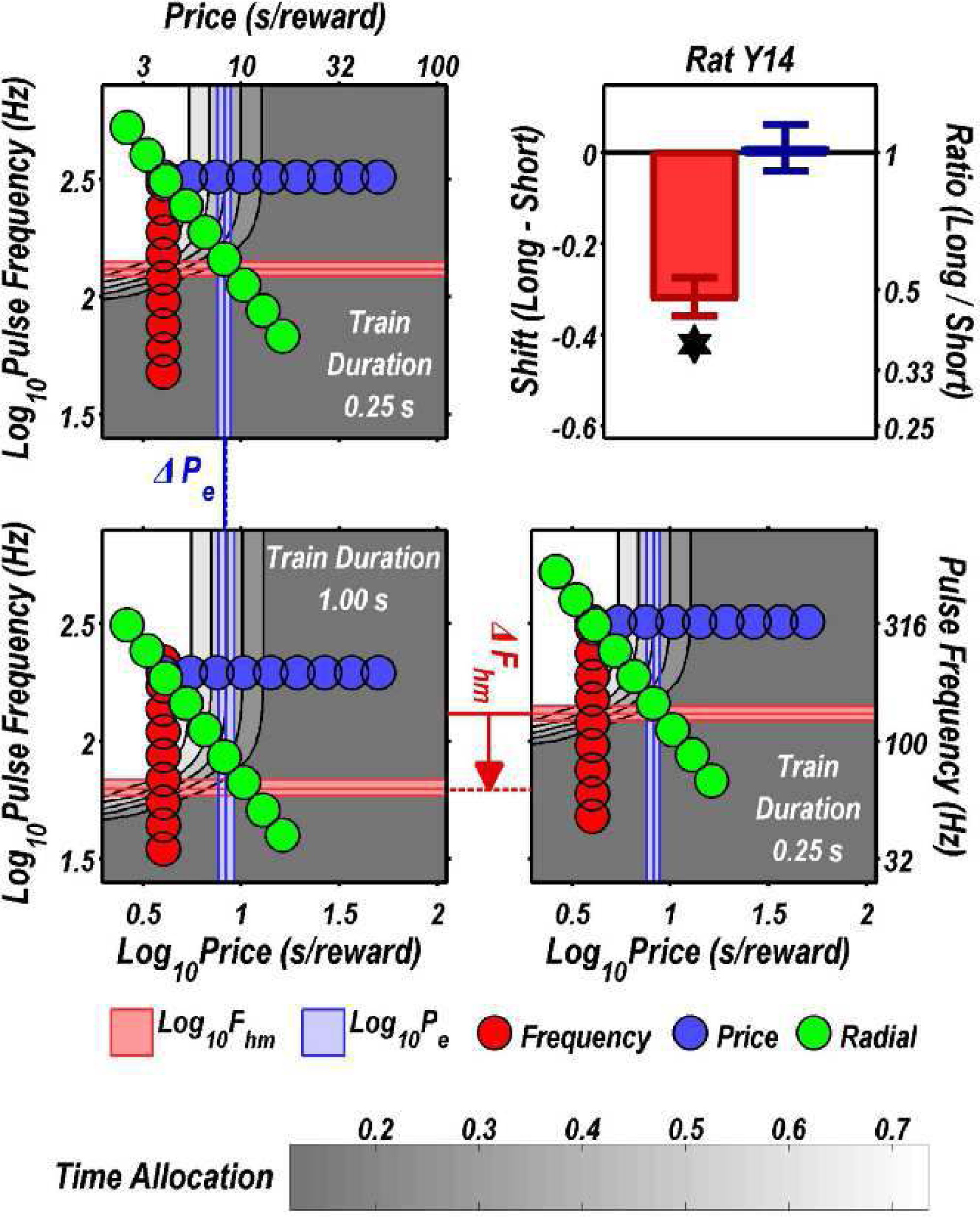

~~~
end
~~~

~~~
[fig_num, fig_tab] = add_fig(fig_num, fig_tab, “LoHiD_quad_Y14”, …
  “Rat Y14: Effect of changing the train duration on the position of the reward mountain”);
~~~

`This is figure #12: (LoHiD_quad_Y14)`

A figure from Breton et al. (2014) showing single-subject data is reproducted above. Note the strong similarity between the simulated results in

~~~
disp(string({strcat({‘Figure ‘},num2str(fig_tab.Number(fig_tab.Name==‘LoHiD_quad’)))}));
~~~

Figure 11

and the empirical results in

~~~
disp(string({strcat({‘Figure ‘},num2str(fig_tab.Number(fig_tab.Name==‘LoHiD_quad_Y14’)))}));
~~~

Figure 12

~~~
toc
~~~

`Elapsed time is 1.508525 seconds.`

### Effect of varying reward probability

~~~
tic;
close all;
~~~

Whereas changing the current or train duration is predicted to shift the reward mountain along the pulse-frequency axis but not along the price axis, changing the probability of delivering a reward upon satisfaction of the response requirement is predicted to produce an orthogonal shift: the mounain should move along the price axis but not along the pulse-frequency axis. This can be seen readily by inspection of

~~~
disp(string({strcat({‘Equation ‘},num2str(eqn_tab.Number(eqn_tab.Name==‘Pobj_e’)))}));
~~~

`Equation`

~~~
disp(string({strcat({‘Equation ‘},num2str(eqn_tab.Number(eqn_tab.Name==‘StrDurFunTr’)))}));
~~~

`Equation`

~~~
and
disp(string({strcat({‘Equation ‘},num2str(eqn_tab.Number(eqn_tab.Name==‘FmfbHM’)))}));
~~~

`Equation`

Intuitively, changing the reward probability should have no effect on the pulse-frequency required to produce a reward of half-maximal intensity, but is should rescale the payoff produced by this reward. Following the change in reward probability, the reward produced by a given pulse frequency is as intense as it was previously, but the ability of this pulse train to compete with alternate sources of reward will depend on the likelihood that the rat gets paid for the work it performs to obtain the electrical stimulation.

We will simulate the effect of changing the reward probability, first from 1.0 to 0.75 and then from 1.0 to 0.5:

C = 0.473; % median from Sonnenschein et al., 2003

D = 0.5; % typical train duration

~~~
logFelecBend = 1.3222; % from Solomon et al., 2015 FelecBend = 10^logFelecBend;
logFelecRO = 2.5587; % from Solomon et al., 2015 FelecRO = 10^logFelecRO;
FPmax =1000; % for determining RbsrNormMax. This value is sufficiently above FelecRO to maximize the firing rate N = 126;
D = 0.5; % train duration RhoPi = 5000; % arbitrary
% RhoPi/N is roughly equal to 50, which is roughly consistent with Sonnenchein et al., 2003
FmfbHM = FpulseHMfun(C, D, FelecBend, FPmax, FelecRO, N, RhoPi)
~~~

`FmfbHM = 77.2222`

~~~
% n.b., FmfbHM1 = FmfbHM2
dotPhiObj = 1;
Kaa = 1;
Kec = 1;
Krg = 1;
dotRaa = 0.1;
pObj = [1, 0.75];
PsubBend = 0.5;
PsubMin = 1.82;
g = 5;
RnormMax = fRbsrNorm(FPmax, FelecBend, FmfbHM, FelecRO, g)
~~~

`RnormMax = 0.9996`

~~~
PobjE = PobjEfun(dotPhiObj, Kaa, Kec, Krg, dotRaa, pObj, PsubBend, PsubMin, RnormMax)
PobjE = 1×2
9.9956 7.4967
~~~

~~~
a = 3;
Felec = logspace(0,3,121)’; % column variable Pobj = logspace(0,3,121); % row variable
Tmfb1 = TAfun(a, Felec, FelecBend, FmfbHM, FelecRO, g, Pobj, …
PobjE(1), PsubBend, PsubMin, RnormMax);
Tmfb2 = TAfun(a, Felec, FelecBend, FmfbHM, FelecRO, g, Pobj, …
PobjE(2), PsubBend, PsubMin, RnormMax);
MTNp1 = plot_MTN(Felec, Pobj, Tmfb1, ‘off’, ‘MTNp1’, ‘p = 1.0’, … graphs2files, FigDir);
MTNp0p75 = plot_MTN(Felec, Pobj, Tmfb2, ‘off’, ‘MTNp0p75’, ‘p = 0.75’, … graphs2files, FigDir);
dual_p0p75_plot = dual_subplot(MTNp1, MTNp0p75, ‘MTNp1vsp0p75’,… graphs2files,FigDir);
if show_graphics dual_p0p75_plot.Visible = ‘on’;
end
~~~

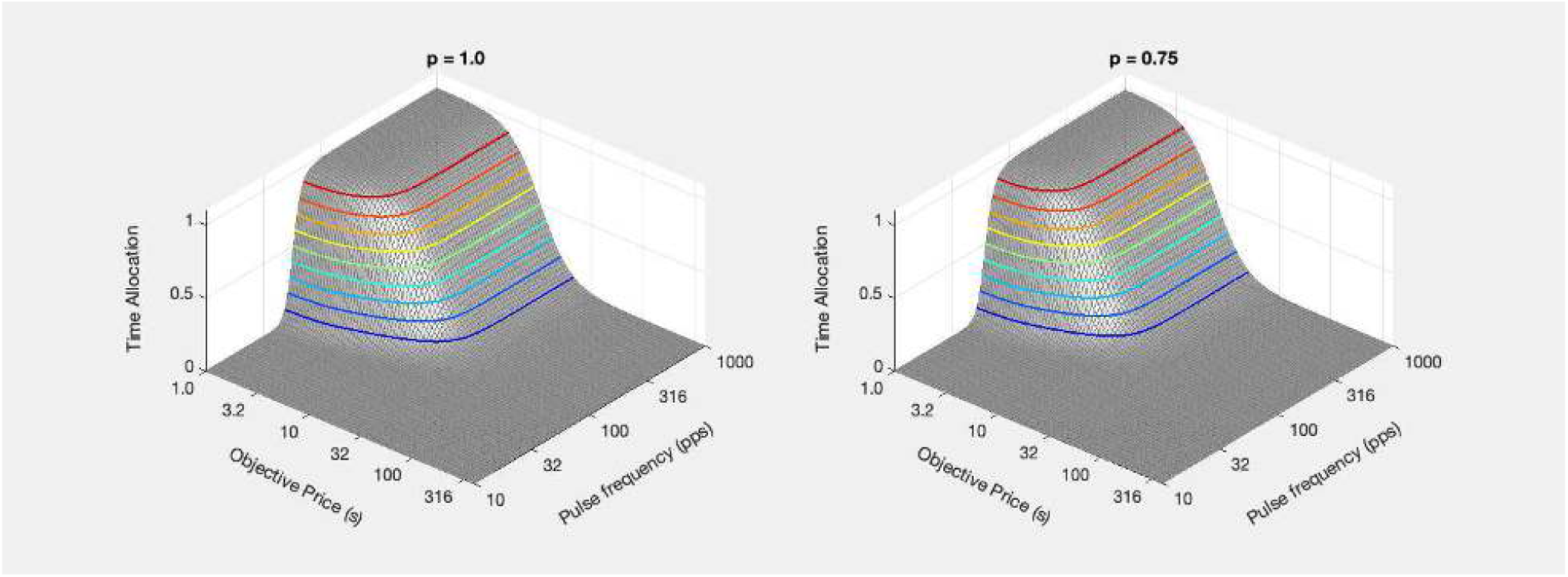

~~~
[fig_num, fig_tab] = add_fig(fig_num, fig_tab, “p1vsp0p75_mtns”, …
  “Effect of changing the reward probability on the reward mountain”);
~~~

`This is figure #13: (p1vsp0p75_mtns)`

~~~
Contp1 = plot_contour(Felec, Pobj, Tmfb1, PobjE(1), FmfbHM, ‘off’, ‘Contp1’, ‘p = 1.0’, …
  strcat({‘p = ‘}, num2str(pObj(1))), graphs2files, FigDir);
Contp0p75 = plot_contour(Felec, Pobj, Tmfb2, PobjE(2), FmfbHM, ‘off’, ‘Contp0p75’, ‘p = 0.75’, …
  strcat({‘p = ‘}, num2str(pObj(2))), graphs2files, FigDir);
bg_p1vsp0p75 = plot_bg(FmfbHM, FmfbHM, PobjE(1), PobjE(2), ‘off’, ‘p1vs0p75_bg’,…
  graphs2files, FigDir, −0.375, 0.075);
bg_root =‘bg_p1vsp0p75’;
quad_p0p75_plot = quad_subplot(Contp1, Contp0p75, bg_p1vsp0p75, ‘quad_p1vsp0p75’, bg_root, … graphs2files, FigDir);
if show_graphics
quad_p0p75_plot.Visible = ‘on’;
end
~~~

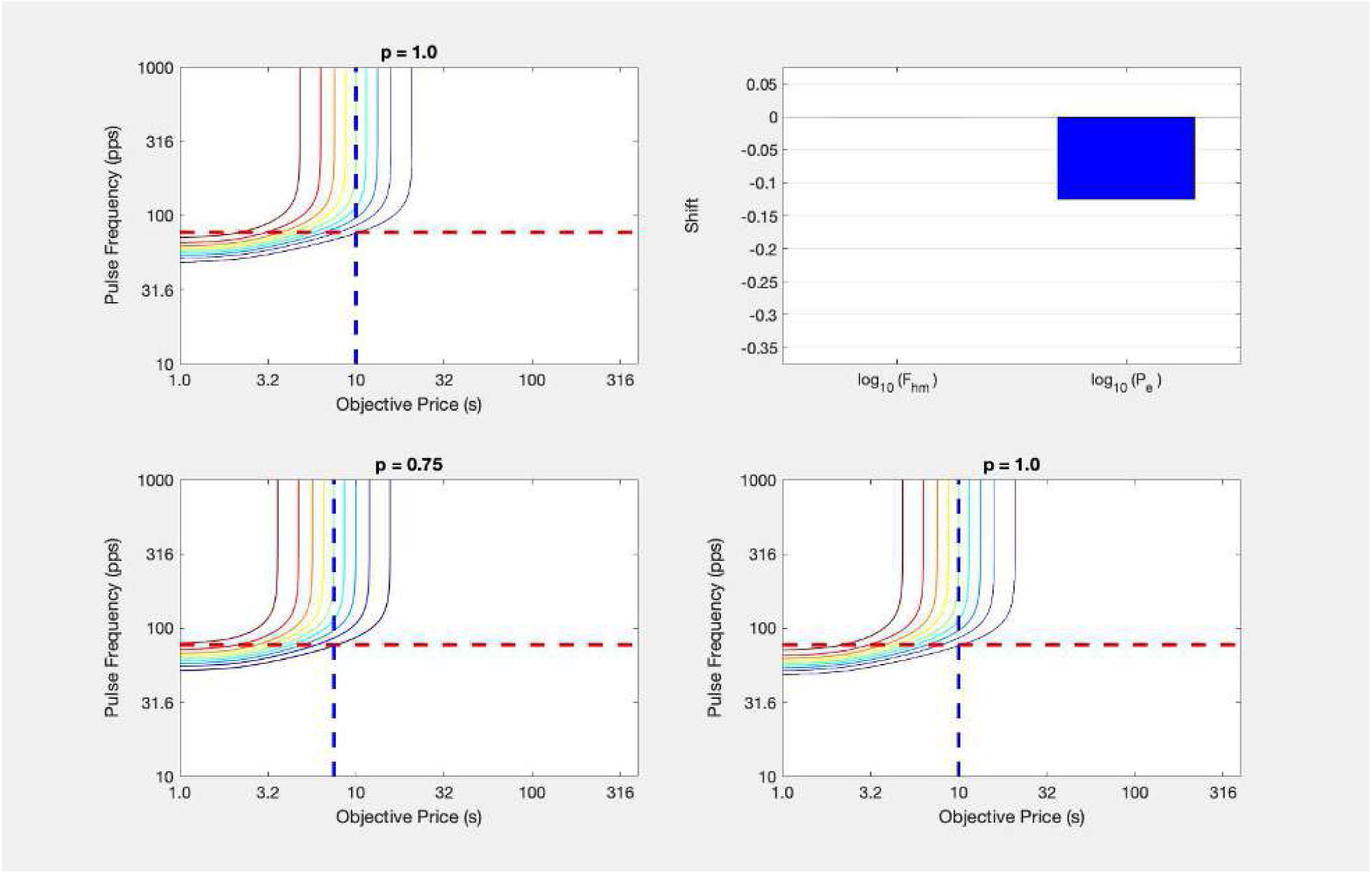

~~~
[fig_num, fig_tab] = add_fig(fig_num, fig_tab, “quad_p1vsp0p75”, …
  “Effect of changing the reward probability on the reward mountain”);
~~~

`This is figure #14: (quad_p1vsp0p75)`

~~~
clear -regexp ^bg ^Cont ^dual ^MTN ^quad; %
Clear only restricted set of variables here toc
~~~

`Elapsed time is 9.850066 seconds.`

~~~
tic;
close all;
~~~

~~~
pObj = [1, 0.5];
PobjE = PobjEfun(dotPhiObj, Kaa, Kec, Krg, dotRaa, pObj, PsubBend, PsubMin, RnormMax)
~~~

~~~
PobjE = 1×2
9.9956 4.9969
~~~

~~~
a = 3;
Felec = logspace(0,3,121)’; % column variable Pobj = logspace(0,3,121); % row variable
Tmfb1 = TAfun(a, Felec, FelecBend, FmfbHM, FelecRO, g, Pobj, …
PobjE(1), PsubBend, PsubMin, RnormMax);
Tmfb2 = TAfun(a, Felec, FelecBend, FmfbHM, FelecRO, g, Pobj, …
  PobjE(2), PsubBend, PsubMin, RnormMax);
MTNp1 = plot_MTN(Felec, Pobj, Tmfb1, ‘off’, ‘MTNp1’, ‘p = 1.0’, … graphs2files, FigDir);
MTNp0p5 = plot_MTN(Felec, Pobj, Tmfb2, ‘off’, ‘MTNp0p5’,   ‘p = 0.5’, … graphs2files, FigDir);
dual_p0p5_plot = dual_subplot(MTNp1, MTNp0p5, ‘MTNp1vsp0p5’,… graphs2files,FigDir);
if show_graphics
dual_p0p5_plot.Visible = ‘on’;
end
~~~

~~~
[fig_num, fig_tab] = add_fig(fig_num, fig_tab, “p1vsp0p5_mtns”, …
  “Effect of changing the reward probability on the reward mountain”);
~~~

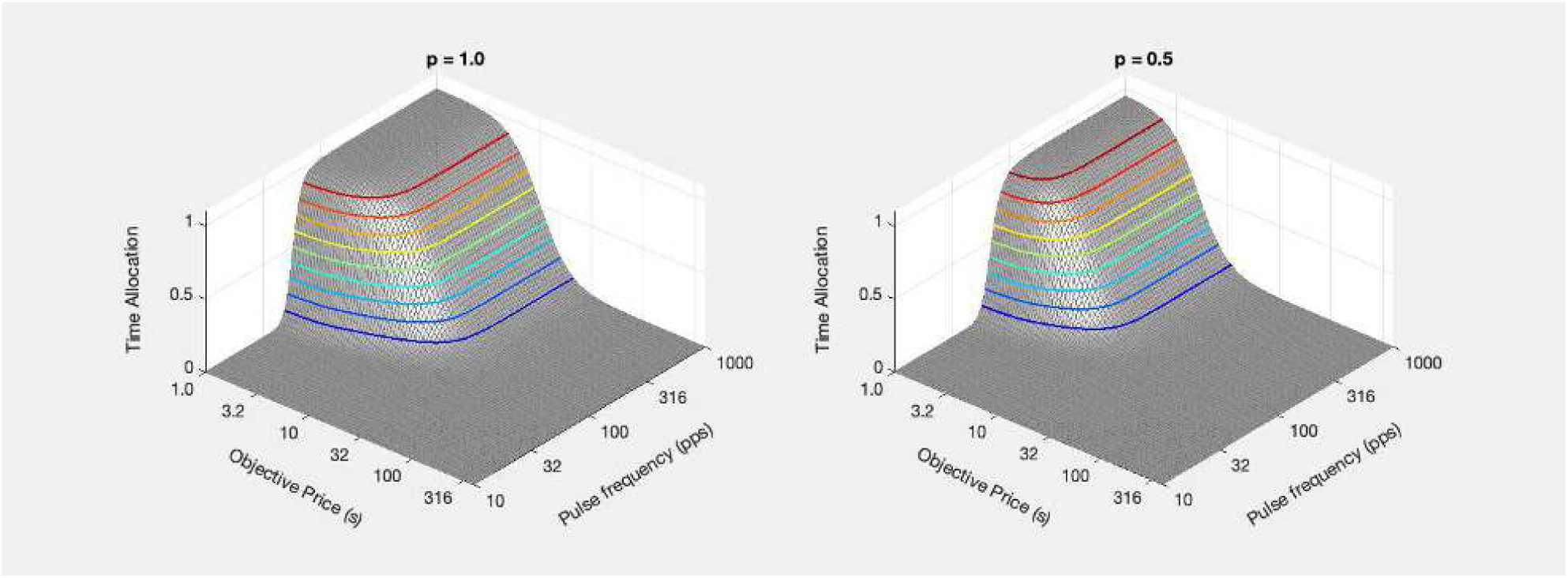

`This is figure #15: (p1vsp0p5_mtns)`

~~~
Contp1 = plot_contour(Felec, Pobj, Tmfb1, PobjE(1), FmfbHM, ‘off’, ‘Contp1’, ‘p = 1.0’, …
strcat({‘p = ‘}, num2str(pObj(1))), graphs2files, FigDir);
Contp0p5 = plot_contour(Felec, Pobj, Tmfb2, PobjE(2), FmfbHM, ‘off’, ‘Contp0p5’, ‘p = 0.5’, …
strcat({‘p = ‘}, num2str(pObj(2))), graphs2files, FigDir);
bg_p1vsp0p5 = plot_bg(FmfbHM, FmfbHM, PobjE(1), PobjE(2), ‘off’, ‘p1vsp0p5_bg’,… graphs2files, FigDir, −0.375, 0.075);
[fig_num, fig_tab] = add_fig(fig_num, fig_tab, “p1vsp0p5_quad”, … “Effect of changing the reward probability on the reward mountain”);
~~~

`This is figure #16: (p1vsp0p5_quad)`

~~~
bg_root =‘bg_p1vsp0p5’;
quad_p0p5_plot = quad_subplot(Contp1, Contp0p5, bg_p1vsp0p5, ‘quad_p1vsp0p5’, bg_root, …
  graphs2files, FigDir);
if show_graphics
  quad_p0p5_plot.Visible = ‘on’;
end
~~~

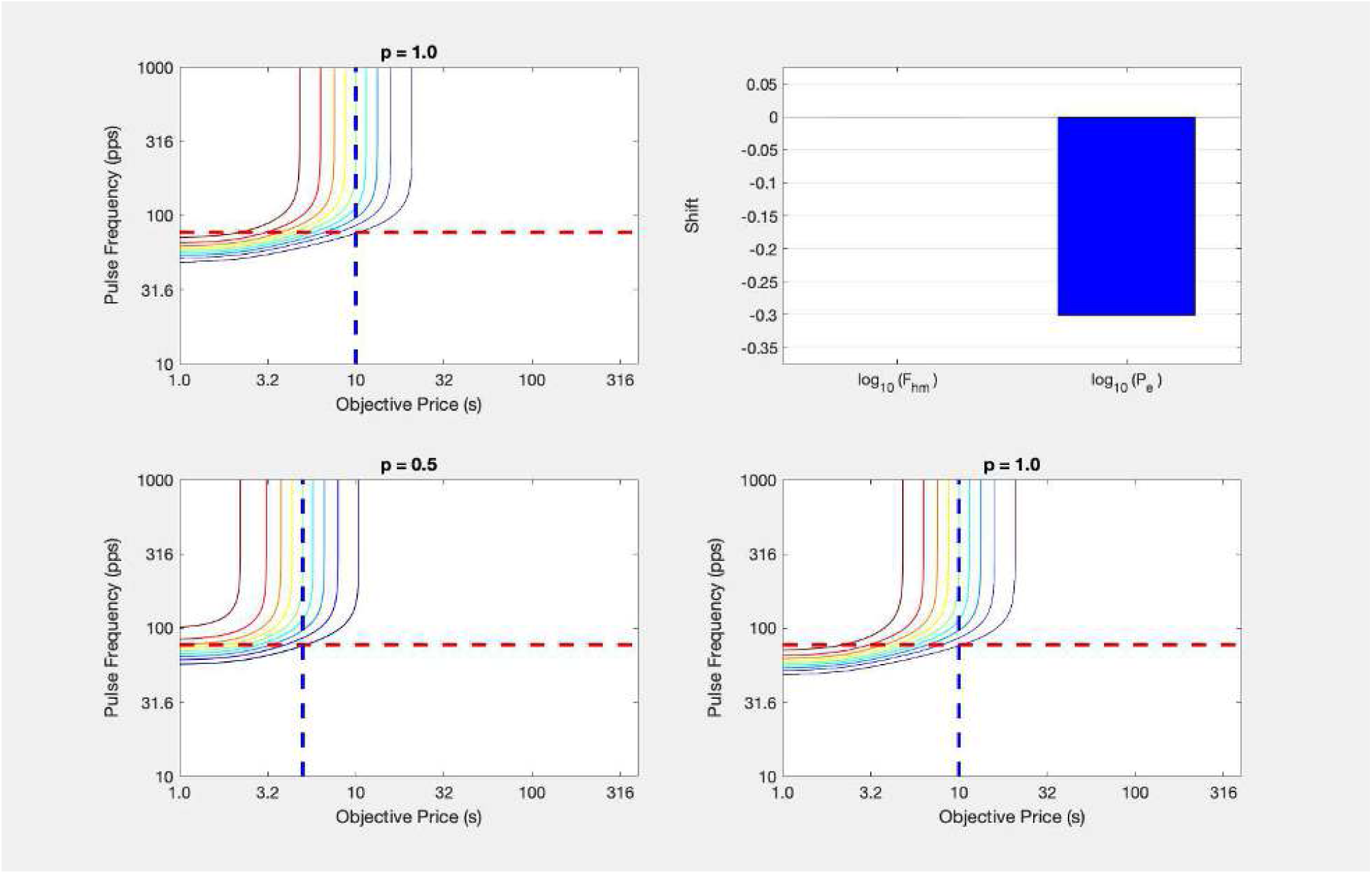

~~~
[fig_num, fig_tab] = add_fig(fig_num, fig_tab, “quad_p1vsp0p5”, …
    “Effect of changing the reward probability on the reward mountain”);
~~~

`This is figure #17: (quad_p1vsp0p5)`

`Elapsed time is 8.880286 seconds.`

~~~
tic;
close all;
~~~

Breton, Conover and Shizgal (2014) tested the effect of decreasing the reward probability from 1.00 to 0.75. The reward mountains obtained from all 10 rats shifted along the price axis as predicted. There was no statistically reliable shift along the pulse-frequency axis in the position of the mountains obtained from six of these rats. In the remaining four cases, the shifts along the pulse-frequency axis were small in comparison to the shifts along the price axis and were inconsistent in direction. Decreasing the probability of reward further to 0.5, increased the size of the shifts along the price axis in all seven rats tested. In five cases, there were no shifts observed along the pulse-frequency axis, and in the two cases in which such shifts were detected, they were inconsistent in direction. Data from one subject showing the effect of reducing the reward probability to 0.5 are shown below

~~~
if show_graphics
    show_imported_graphic(‘prob_shifts_PD8.png’,25,ImpFigDir);
end
~~~

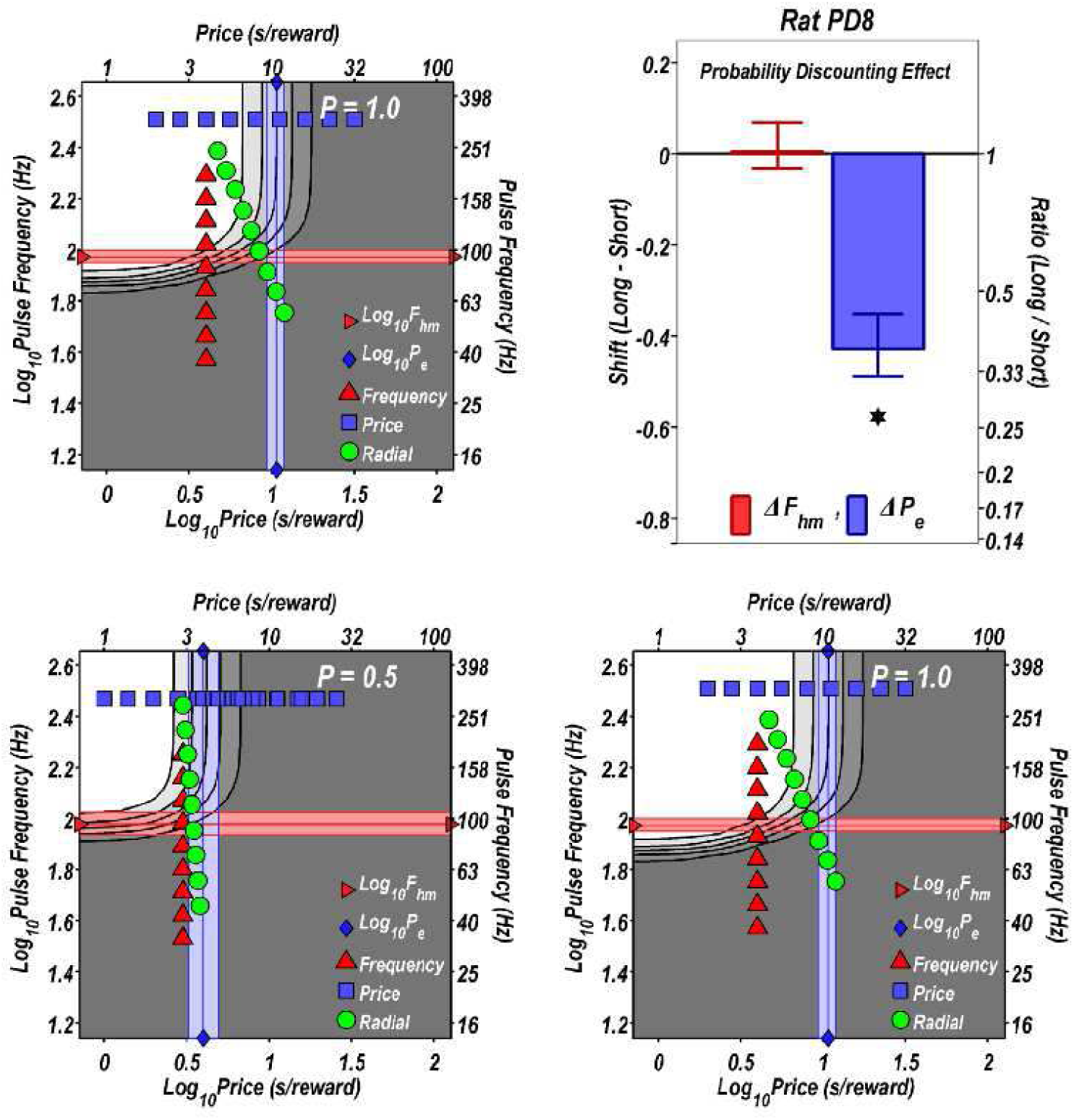

~~~
[fig_num, fig_tab] = add_fig(fig_num, fig_tab, “p1vsp0p5_quad_PD8”, …
     “Rat PD8: Effect of changing the reward probability on the reward mountain”);
~~~

`This is figure #18: (p1vsp0p5_quad_PD8)`

followed by a summary of the entire dataset.

~~~
if show_graphics
    show_imported_graphic(‘prob_disc_all.png’,30,ImpFigDir);
end
~~~

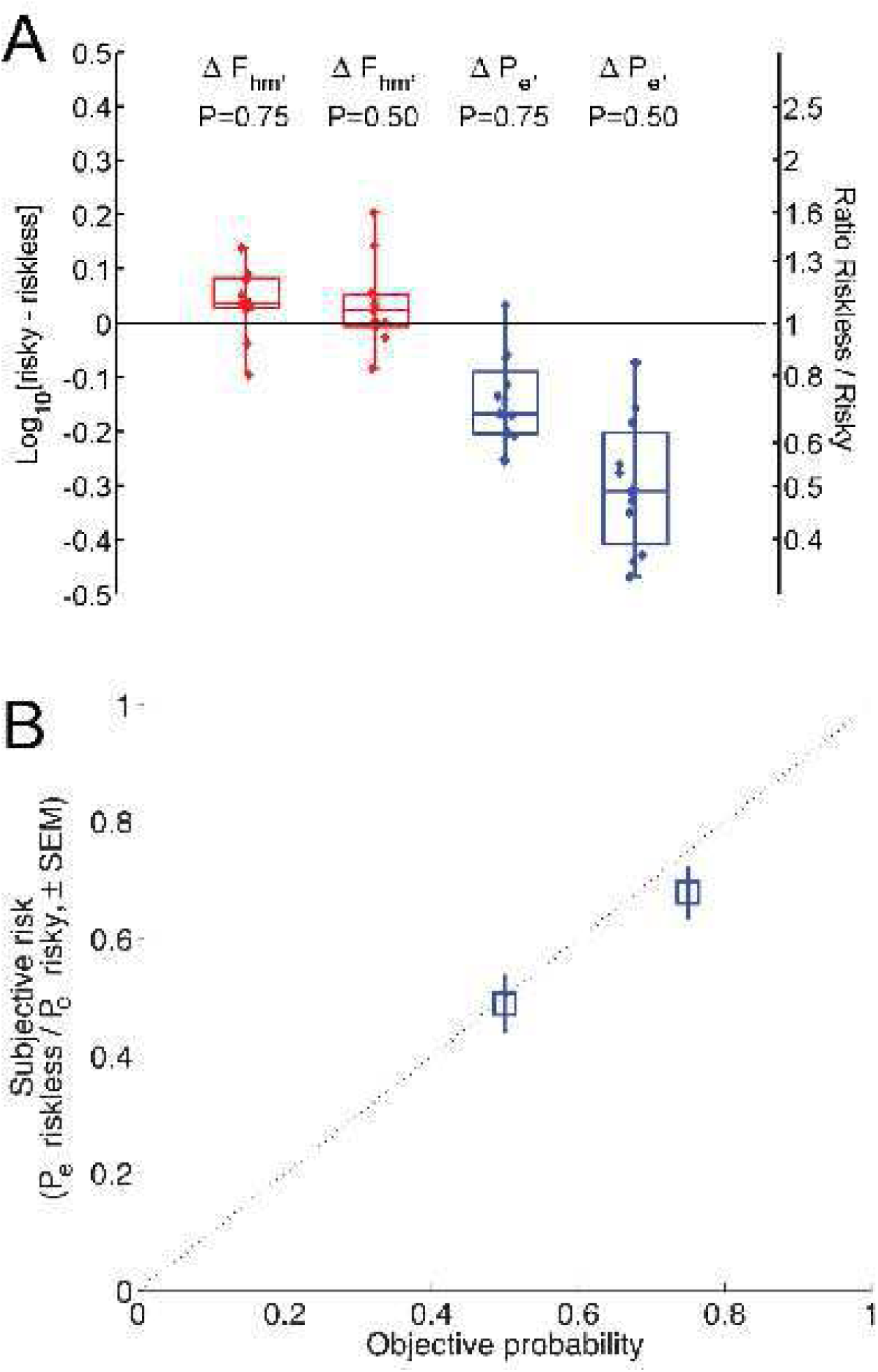

~~~
[fig_num, fig_tab] = add_fig(fig_num, fig_tab, “ProbDiscSummary”, …
      “Summary of the effects of changing the reward probability on the reward mountain”);
~~~

`This is figure #19: (ProbDiscSummary)`

The upper panel of the summary shows that changing reward probability moves the mountain almost exclusively along the price axis and does so in a graded manner; the larger the change in reward probability, the larger the shift along the price axis. On the basis of

~~~
disp(string({strcat({‘Equation ‘},num2str(eqn_tab.Number(eqn_tab.Name==‘PayoffElec’)))}));
~~~

`Equation`

we can estimate the subjective reward probability from the observed shift along the price axis. The lower panel of the summary shows that the subjective reward probabilities estimated in this manner are all but indistinguishable from the objective reward probabilities.

~~~
toc
~~~

`Elapsed time is 1.608664 seconds.`

## The significance of orthogonal shifts

~~~
tic;
close all;
~~~

The validation experiments just described show conclusively that the reward mountain can be displaced along either the pulse-frequency or price axes, as predicted by the underlying model. In most cases, these displacements are orthogonal: the mountain shifts either along one axis or the other.

These observations have important implications for the form of the reward-growth function at the heart of the mountain model. In order for shifts along more than one axis to be possible, the form of the reward-growth function must distinguish changes that rescale its input and output. The logistic form implied by the matching data from the Gallistel lab instantiates such a distinction. Changes in the value of the position parameter, 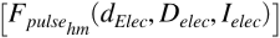, rescale the input, thereby shifting the reward-growth function along the pulse-frequency axis, whereas changes in the value of the maximum attainable reward, (***K***_*rg*_), rescale the output, thereby shifting the maximum value of the reward-growth function vertically. In the figures below, we change the value of the input-rescaling position parameter 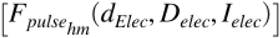 by varying the current (thereby changing *N*, the number of reward-generating neurons recruited), and we rescale the output of the reward- growth function by changing the value of the maximum attainable reward, (***K***_*rg*_).

~~~
logFelecBend = 1.3222; % from Solomon et al., 2015
FelecBend = 10^logFelecBend;
logFelecRO = 2.5587; % from Solomon et al., 2015
FelecRO = 10^logFelecRO;
numF = 121; % Number of pulse frequencies in Felec vector
Felec = logspace(0,3,numF);
FelecMat = repmat(logspace(0,3,numF),3,1); % add two rows in preparation for graphing
C = median([0.453,0.503, 0.268, 0.224, 0.493, 0.64]); % from Sonnenschein et al., 2003
D = 0.5;
RhoPi = (5000 / fHfiring(C, D, 100, 5000)) * 100; % To position the logFhm values near 0.1 log10 units
gElec = 5;
NnLG1 = 10^1.8; % The number of elements in the parameter vector must equal the # of rows in FelecMat
NnLG2 = 10^2;
NnLG3 = 10^2.2;
NnLGvec = [NnLG1;NnLG2;NnLG3];
NnLGmat = repmat(NnLGvec,1,numF); % Store the N values in a matrix of the same size as FelecMat
KrgLG1 = 10^-0.2; % The number of elements in the parameter vector must equal the # of rows in FelecMat
KrgLG2 = 10^0;
KrgLG3 = 10^0.2;
KrgLGvec = [KrgLG1;KrgLG2;KrgLG3];
KrgLGmat = repmat(KrgLGvec,1,numF); % Store the Krg values in a matrix of the same size as FelecMat
FhmLG1 = fHfiring(C, D, NnLG1, RhoPi); % The number of elements in the parameter vector must equal the # of rows in FelecMat
FhmLG2 = fHfiring(C, D, NnLG2, RhoPi);
FhmLG3 = fHfiring(C, D, NnLG3, RhoPi);
FhmLGvec = [FhmLG1;FhmLG2;FhmLG3];
FhmLGmat = repmat(FhmLGvec,1,numF); % Store the Fhm values in a matrix of the same size as FelecMat
RelecLGmat_Nn = fRbsrFull(C,D,FelecMat,FelecBend,FelecRO,gElec,KrgLG2,NnLGmat,RhoPi);
RelecLGmat_Krg = fRbsrFull(C,D,FelecMat,FelecBend,FelecRO,gElec,KrgLGmat,NnLG2,RhoPi);
[fun_num, fun_tab] = add_fun(fun_num, fun_tab, “fRbsrFull”, …
     “C, D, F, Fbend, Fro, g, Krg, N, RhoPi, varargin”,…
     “Full reward-growth function for BSR”);
~~~

`This is function #34: (fRbsrFull)`

~~~
TitleStrSemiLog = ‘logistic growth’;
TitleStrLogLog = ‘logistic growth’;
pnam = “F_{hm}”;
fnam = “Fhm”;
% The data to be plotted must be in columns. Thus, FelecMat and RelecMat are transposed.
RG_NnLG_semilog = plot_RG(FelecMat’,RelecLGmat_Nn’,pnam,FhmLGvec,fnam,TitleStrSemiLog,’lin’);
RG_NnLG_loglog = plot_RG(FelecMat’,RelecLGmat_Nn’,pnam,FhmLGvec,fnam,TitleStrSemiLog,’log’);
pnam = “K_{rg}”;
fnam = “Krg”;
RG_KrgLG_semilog = plot_RG(FelecMat’,RelecLGmat_Krg’,pnam,KrgLGvec,fnam,TitleStrSemiLog,’lin’);
RG_KrgLG_loglog = plot_RG(FelecMat’,RelecLGmat_Krg’,pnam,KrgLGvec,fnam,TitleStrSemiLog,’log’);
dual_subplot(RG_NnLG_loglog, RG_KrgLG_loglog, ‘RG_NnLG_KrgLG_loglog’,…
    graphs2files,FigDir);
if show_graphics
   RG_NnLG_KrgLG_loglog.Visible = ‘on’;
end
[fig_num, fig_tab] = add_fig(fig_num, fig_tab, “LogisticRGLL”, …
    “When reward intensity grows as a logistic function of pulse frequency, shifts are orthogonal”);
~~~

`This is figure #20: (LogisticRGLL)`

~~~
RG_NnLG_KrgLG_semilog = dual_subplot(RG_NnLG_semilog, RG_KrgLG_semilog, ‘RG_NnLG_KrgLG_semilog’,…
    graphs2files,FigDir);
if show_graphics
    RG_NnLG_KrgLG_semilog.Visible = ‘on’;
end
~~~

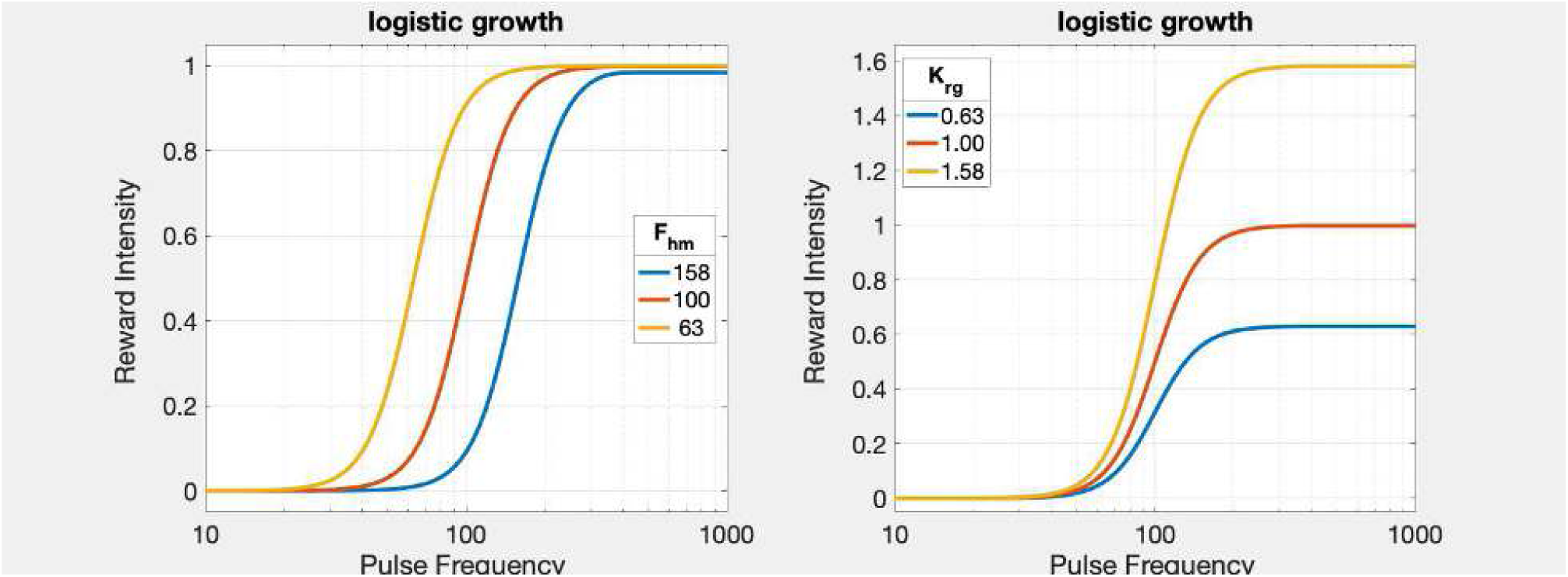

~~~
[fig_num, fig_tab] = add_fig(fig_num, fig_tab, “LogisticRGSL”, …
   “When reward intensity grows as a logistic function of pulse frequency, shifts are orthogonal”);
~~~

`This is figure #21: (LogisticRGSL)`

The horizontal shifts of the reward-growth function are translated into shifts of the reward mountain along the pulse-frequency axis, whereas vertical shifts of the reward-growth function are translated in shifts of the reward mountain along the price axis. Above, we have illustrated such orthogonal shifts by simulating the effects of changing the current or train duration, as shown in

~~~
disp(string({strcat({‘Figure ‘},num2str(fig_tab.Number(fig_tab.Name==‘LoHiI_quad’)))}));
~~~

`Figure`

~~~
disp(string({strcat({‘Figure ‘},num2str(fig_tab.Number(fig_tab.Name==‘LoHiD_quad’)))}));
~~~

Figure 11

Figure 12

and the effects of changing the reward probability, as shown in

~~~
disp(string({strcat({‘Figure ‘},num2str(fig_tab.Number(fig_tab.Name==‘p1vsp0p75_quad’)))}));
~~~

`Figure`

~~~
disp(string({strcat({‘Figure ‘},num2str(fig_tab.Number(fig_tab.Name==‘p1vsp0p5_quad’)))}));
~~~

`Figure 16`

~~~
disp(string({strcat({‘Figure ‘},num2str(fig_tab.Number(fig_tab.Name==‘p1vsp0p5_quad_PD8’)))}));
~~~

`Figure 18`

What would happen if frequency-following fidelity remained the same, but the input-scaling and output-scaling parameters of the reward-growth function were no longer independent? We can simulate such a case by replacing the logistic reward-growth function with a power function:

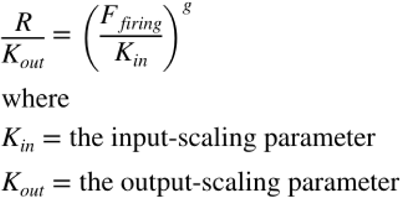

~~~
[eqn_num, eqn_tab] = …
    add_eqn(eqn_num, eqn_tab, “RGpg”, “Power growth of reward intensity”);
~~~

`This is equation #27: (RGpg)`

~~~
[fun_num, fun_tab] = add_fun(fun_num, fun_tab, “fRpg”, “F, Fbend, Fro, g, Kin, Krg”,…
    “Function to compute power growth of reward”);
~~~

`This is function #35: (fRpg)`

This power-growth function can be rewritten as

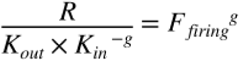

~~~
[eqn_num, eqn_tab] = …
    add_eqn(eqn_num, eqn_tab, “RGpgReformat”, “Power growth of reward intensity”);
~~~

`This is equation #28: (RGpgReformat)`

In contrast to the case of logistic growth, the two scaling constants act jointly, in an inseparable manner, and exclusively on the output of the function. Changing either scaling constant shifts the power-growth function along the y axis but does not change its position along the x axis. As in the case of the logistic reward-growth function, *g* determines the steepness of reward growth (the slope on double-logarithmic coordinates). Although ***K***_*out*_ appears to scale the output, whereas ***K***_*in*_ appears to scale the induced firing frequency,

~~~
disp(string({strcat({‘Equation ‘},num2str(eqn_tab.Number(eqn_tab.Name==‘RGpgReformat’)))}));
~~~

`Equation 28`

shows that the two constants function as one. Thus, reward intensity will grow as a power function of the electrical pulse frequency until the induced firing frequency in the stimulation reward-related neurons asymptotes. Changing the current displaces the ***logistic*** reward-growth function laterally, a shift that is orthogonal to the one produced by changing the scale parameter, ***K***_*rg*_. In contrast, changing the current displaces the ***power*** reward-growth function vertically, the same direction as the shift produced by changing the scale parameter. This is shown graphically below.

In the following simulations, changing the value of *N*serves as a proxy for changes in current. The ***K***_*in*_ parameter is inversely proportional to the number of stimulated neurons (*N*). In the figure legends, the value of the ***K***_*in*_ parameter is normalized to the ***K***_*in*_ value corresponding to the middle value of *N*.

~~~
logFelecBend = 1.3222;
FelecBend = 10^logFelecBend;
logFelecRO = 2.5587;
FelecRO = 10^logFelecRO;
FPmax = 1000;
numF = 121; % Number of pulse frequencies in Felec vector
Felec = logspace(0,3,numF);
FelecMat = repmat(logspace(0,3,numF),3,1); % add two rows in preparation for graphing
gElec = 2.5;
NnPG1 = 10^(2-(0.3/gElec)); % The number of elements in the parameter vector must equal the # of rows in FelecMat
NnPG2 = 10^2;
NnPG3 = 10^(2+(0.3/gElec));
% NnPGvec = [NnPG1;NnPG2;NnPG3];
% NnPGmat = repmat(NnPGvec,1,numF); % Store the N values in a matrix of the same size as FelecMat
FFaggNorm = FilterFun(FPmax,FelecBend, FelecRO) * NnPG2; % Normalization factor
% When Kout = 1 and NnPG = NnPG2, FFaggNorm ensures that R = 1 when FF = FFmax
KinPGnorm = FFaggNorm/NnPG2;
KinPG1 = FFaggNorm/NnPG1;
KinPG2 = FFaggNorm/NnPG2;
KinPG3 = FFaggNorm/NnPG3;
KinPGvec = [KinPG1;KinPG2;KinPG3];
KinPGmat = repmat(KinPGvec,1,numF); % Store the Kin values in a matrix of the same size as FelecMat
% The number of elements in the parameter vector must equal the # of rows in FelecMat
KoutPG1 = 10^-0.3;
KoutPG2 = 10^0;
KoutPG3 = 10^0.3; % RpgMax = 10^0.15 = 1.4125
KoutPGvec = [KoutPG1;KoutPG2;KoutPG3];
KoutPGmat = repmat(KoutPGvec,1,numF); % Store the Kout values in a matrix of the same size as FelecMat
RelecPGmat_Kin = fRpg(FelecMat,FelecBend,FelecRO,gElec,KinPGmat,KoutPG2);
RelecPGmat_Kout = fRpg(FelecMat,FelecBend,FelecRO,gElec,KinPG2,KoutPGmat);
TitleStrSemiLog = ‘power growth’;
TitleStrLogLog = ‘power growth’;
pnam = “K_{in} / K_{in_{norm}}”;
fnam = “Kin”;
KinLgndVec = KinPGvec ./ KinPGnorm;
RG_KinPG_semilog = plot_RG(FelecMat’,RelecPGmat_Kin’,pnam,KinLgndVec,fnam,TitleStrSemiLog,’lin’);
RG_KinPG_loglog = plot_RG(FelecMat’,RelecPGmat_Kin’,pnam,KinLgndVec,fnam,TitleStrLogLog,’log’);
pnam = “K_{out}”;
fnam = “Kout”;
RG_KoutPG_semilog = plot_RG(FelecMat’,RelecPGmat_Kout’,pnam,KoutPGvec,fnam,TitleStrSemiLog,’lin’);
RG_KoutPG_loglog = plot_RG(FelecMat’,RelecPGmat_Kout’,pnam,KoutPGvec,fnam,TitleStrLogLog,’log’);
RG_KinPG_KoutPG_semilog = dual_subplot(RG_KinPG_semilog, RG_KoutPG_semilog, ‘RG_KinPG_KoutPG_semilog’,…
    graphs2files,FigDir);
if show_graphics
    RG_KinPG_KoutPG_semilog.Visible = ‘on’;
end
~~~

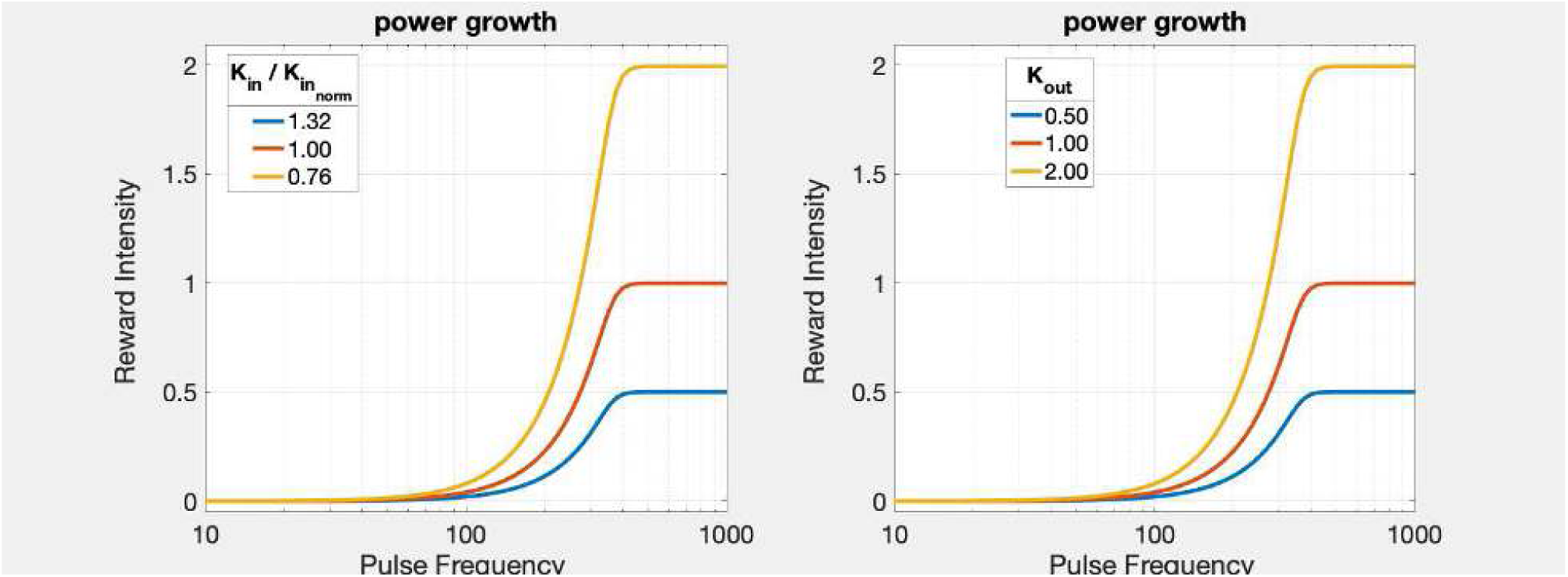

~~~
[fig_num, fig_tab] = add_fig(fig_num, fig_tab, “PowerRGSL”, …
    “When reward intensity grows as a power function, all shifts are vertical”);
~~~

`This is figure #22: (PowerRGSL)`

~~~
dual_subplot(RG_KinPG_loglog, RG_KoutPG_loglog, ‘RG_KinPG_KoutPG_loglog’,…
    graphs2files,FigDir);
if show_graphics
    RG_KinPG_KoutPG_loglog.Visible = ‘on’;
end
%shg
[fig_num, fig_tab] = add_fig(fig_num, fig_tab, “PowerRGLL”, …
    “When reward intensity grows as a power function, all shifts are vertical”);
~~~

`This is figure #23: (PowerRGLL)`

~~~
disp(string({strcat({‘Equation ‘},num2str(eqn_tab.Number(eqn_tab.Name==‘RGpg’)))}));
~~~

`Equation 27`

Unlike the case with logistic growth of reward intensity, the variable that scales the input to the power reward- growth function (***K***_*in*_) and the variable that scales its output (***K***_*out*_) are no longer independent and cannot move the reward-growth curve in orthogonal directions. As a result, the reward mountain can shift only along a single axis when the underlying reward-growth function lacks independent input- and output-scaling parameters. This is demonstated in the following section.

`Elapsed time is 15.340981 seconds.`

~~~
tic;
close all;
~~~

## The reward mountain can shift only along the price axis when the underlying reward-growth function lacks indepedent input- and output-scaling parameters

~~~
dotPhiObj = 1;
Kaa = 1;
Kec = 1;
dotRaa = 0.1;
pObj = 1;
a = 3;
Felec = logspace(0,3,121)’; % column variable
Pobj = logspace(0,3,121); % row variable
logFelecBend = 1.3222;
FelecBend = 10^logFelecBend;
logFelecRO = 2.5587;
FelecRO = 10^logFelecRO;
gElec = 2.5;
N = 100;
PsubBend = 0.5;
PsubMin = 1.82;
FPmax = 1000;
FFmax = FilterFun(FPmax,FelecBend, FelecRO);
Kin = [FFmax, FFmax / 2^(1/gElec)];
Kout = [1,2];
RmaxBase = fRpg(1000,FelecBend,FelecRO,gElec,Kin(1),Kout(1));
RmaxKin = fRpg(1000,FelecBend,FelecRO,gElec,Kin,Kout(1));
RmaxKout = fRpg(1000,FelecBend,FelecRO,gElec,Kin(1),Kout);
PsubEpgBase = PsubEpgFun(dotPhiObj, Kaa, Kec, Kout, dotRaa, pObj, RmaxBase);
PsubEpgKin = PsubEpgFun(dotPhiObj, Kaa, Kec, Kout, dotRaa, pObj, RmaxKin);
PsubEpgKout = PsubEpgFun(dotPhiObj, Kaa, Kec, Kout, dotRaa, pObj, RmaxKout);
[fun_num, fun_tab] = add_fun(fun_num, fun_tab, “PsubEpgFun”, …
    “dotPhiObj, Kaa, Kec, Krg, ObjAA, pObj, Rmax”,…
    “Function to compute PsubE for power reward growth”);
~~~

`This is function #36: (PsubEpgFun)`

~~~
TpgBase = TApgFun(a, Felec, FelecBend, FelecRO, gElec, Kin(1), Kout(1), Pobj, PsubBend, PsubEpgBase, PsubMin);
TpgKin2 = TApgFun(a, Felec, FelecBend, FelecRO, gElec, Kin(2), Kout(1), Pobj, PsubBend, PsubEpgKin(2), PsubMin);
TpgKout2 = TApgFun(a, Felec, FelecBend, FelecRO, gElec, Kin(1), Kout(2), Pobj, PsubBend, PsubEpgKout(2), PsubMin);
[fun_num, fun_tab] = add_fun(fun_num, fun_tab, “TApgFun”, …
   “a, Felec, FelecBend, FelecRO, gElec, Kin, Kout, N, Pobj, PsubBend, PsubEpgBase, PsubMin”,…
   “Time allocation in response to power growth of reward intensity”);
~~~

`This is function #37: (TApgFun)`

~~~
xmin = 0;
xmax = 2.3;
ymin = 1.5;
ymax = 3;
MTNpgKin1 = plot_MTN(Felec, Pobj, TpgBase, ‘off’, ‘MTNpgKin1’, ‘Kin = Kin1’, …
   graphs2files, FigDir, xmin, xmax, ymin, ymax);
MTNpgKin2 = plot_MTN(Felec, Pobj, TpgKin2, ‘off’, ‘MTNpgKin2’, strcat(‘Kin1’,’\div’,’2.0^{1/g}’), …
   graphs2files, FigDir, xmin, xmax, ymin, ymax);
dual_pgKin_plot = dual_subplot(MTNpgKin1, MTNpgKin2, ‘MTNpgKin1Kin2’,…
  graphs2files,FigDir);
if show_graphics
    dual_pgKin_plot.Visible = ‘on’;
end
[fig_num, fig_tab] = add_fig(fig_num, fig_tab, “PG_K1_1vsK1_2_mtns”, …
    “Effect of changing the power-growth input-scaling parameter”);
~~~

`This is figure #24: (PG_K1_1vsK1_2_mtns)`

~~~
MTNpgKout1 = plot_MTN(Felec, Pobj, TpgBase, ‘off’, ‘MTNpgKout1’, ‘Kout = Kout1’, …
    graphs2files, FigDir, xmin, xmax, ymin, ymax);
MTNpgKout2 = plot_MTN(Felec, Pobj, TpgKout2, ‘off’, ‘MTNpgKout2’, ‘Kout = 2*Kout1’, …
    graphs2files, FigDir, xmin, xmax, ymin, ymax);
~~~

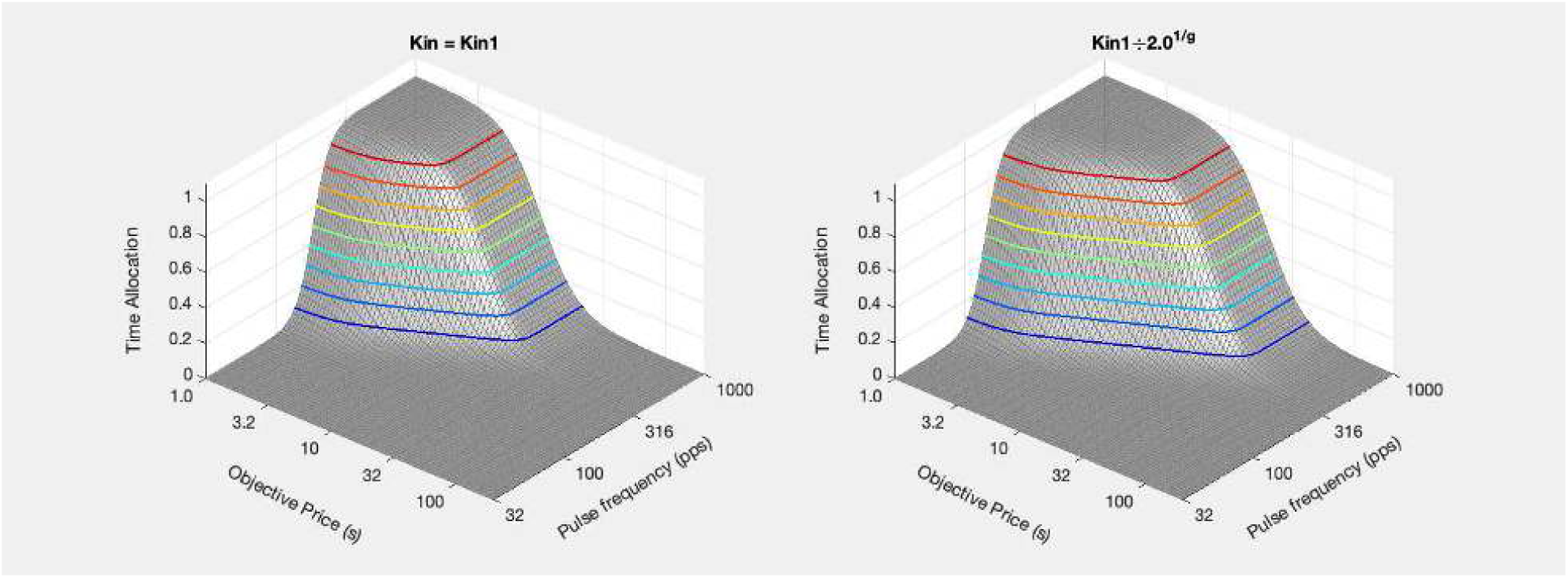

~~~
dual_pgKout_plot = dual_subplot(MTNpgKout1, MTNpgKout2, ‘MTNpgKout1Kout2’,…
    graphs2files,FigDir);
if show_graphics
    dual_pgKout_plot.Visible = ‘on’;
end
~~~

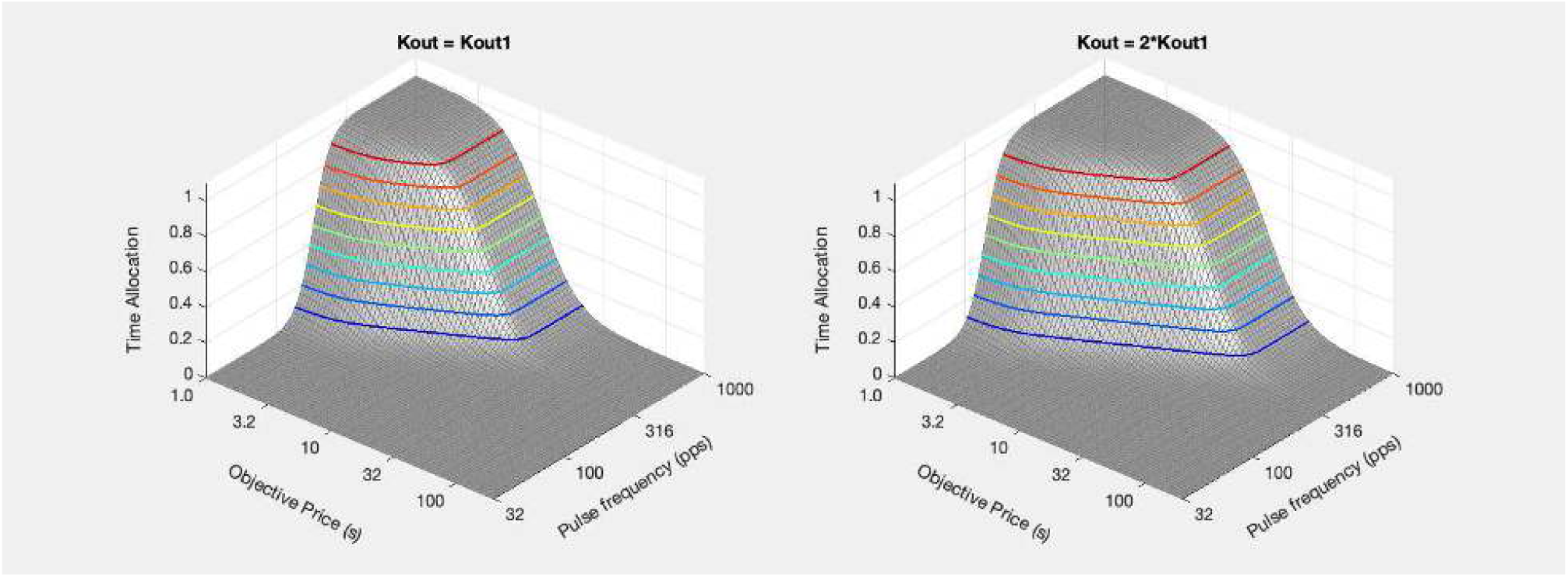

~~~
[fig_num, fig_tab] = add_fig(fig_num, fig_tab, “PG_Kout1vsKout2_mtns”, …
    “Effect of changing the power-growth output-scaling parameter”);
~~~

`This is figure #25: (PG_Kout1vsKout2_mtns)`

~~~
FhmPGKin = find_FhmPG(FPmax, FelecBend, FelecRO, gElec, Kin, Kout(1), RmaxKin);
FhmPGKout = find_FhmPG(FPmax, FelecBend, FelecRO, gElec, Kin(1), Kout, RmaxKout);
[fun_num, fun_tab] = add_fun(fun_num, fun_tab, “find_FhmPG”, …
     “FPmax, Fbend, Fro, g, Kin, Kout, RmaxPG”,…
     “Function to compute the Fhm value for power reward growth”);
~~~

`This is function #38: (find_FhmPG)`

~~~
PobjEpgBase = PsubBsFun(PsubEpgBase, PsubBend, PsubMin);
PobjEpgKin = PsubBsFun(PsubEpgKin, PsubBend, PsubMin);
PobjEpgKout = PsubBsFun(PsubEpgKout, PsubBend, PsubMin);
ContKin1 = plot_contour(Felec, Pobj, TpgBase, PobjEpgKin(1), FhmPGKin(1), ‘off’, ‘ContKin1’, ‘Kin = 1.0’, …
    strcat({‘Kin = ‘}, num2str(round(Kin(1),2))), graphs2files, FigDir,…
    xmin, xmax, ymin, ymax);
ContKin2 = plot_contour(Felec, Pobj, TpgKin2, PobjEpgKin(2), FhmPGKin(2), ‘off’, ‘ContKin2’, ‘Kin = 2.0^{1/g}’, …
    strcat({‘Kin = ‘}, num2str(round(Kin(2),2))), graphs2files, FigDir,…
    xmin, xmax, ymin, ymax);
bg_Kin1vskKin2 = plot_bg(FhmPGKin(1), FhmPGKin(2), PobjEpgKin(1), PobjEpgKin(2), ‘off’, ‘Kin1vsKin2_bg’,…
    graphs2files, FigDir);
bg_root = ‘bg_Kin1vsKin2’;
quad_Kin1vsKin2 = quad_subplot(ContKin1, ContKin2, bg_Kin1vskKin2, ‘quad_Kin1vsKin2’, bg_root, …
    graphs2files, FigDir);
if show_graphics
    quad_Kin1vsKin2.Visible = ‘on’;
end
~~~

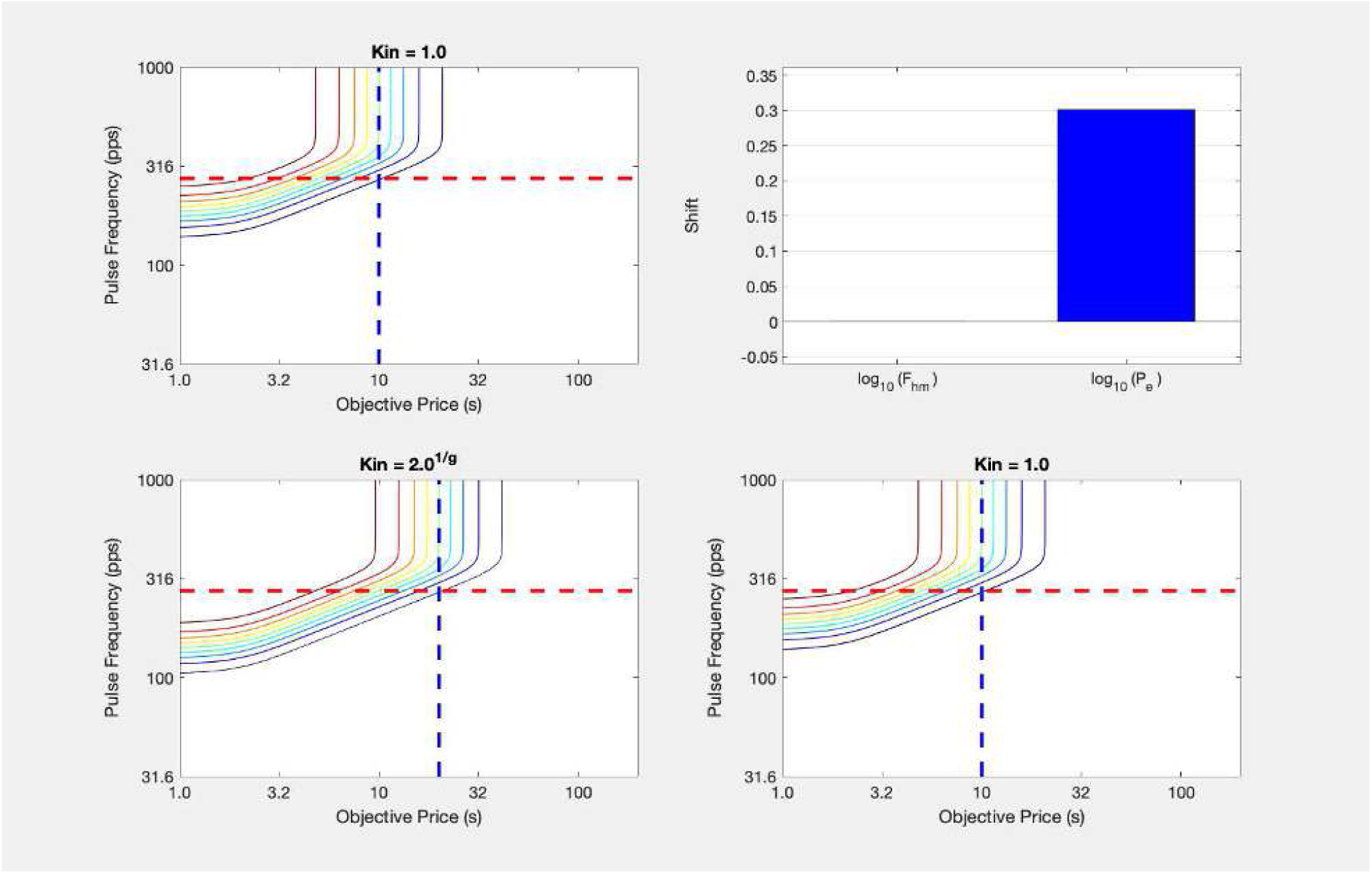

~~~
[fig_num, fig_tab] = add_fig(fig_num, fig_tab, “quad_Kin1vsKin2”, “Effect of increasing the value of the input-scaling paramater”);
~~~

`This is figure #26: (quad_Kin1vsKin2)`

~~~
ContKout1 = plot_contour(Felec, Pobj, TpgBase, PobjEpgKout(1), FhmPGKout(1), ‘off’, ‘ContKout1’, ‘Kout = Kout1’, …
    strcat({‘Kout = ‘}, {‘Kout1’}), graphs2files, FigDir,…
    xmin, xmax, ymin, ymax);
ContKout2 = plot_contour(Felec, Pobj, TpgKout2, PobjEpgKout(2), FhmPGKout(2), ‘off’, ‘ContKout2’, ‘Kout = 2 * Kout1’, …
    strcat({‘Kout = ‘}, {‘2 * Kout1’}), graphs2files, FigDir,…
    xmin, xmax, ymin, ymax);
bg_Kout1vskKout2 = plot_bg(FhmPGKout(1), FhmPGKout(2), PobjEpgKout(1), PobjEpgKout(2), ‘off’, ‘Kout1vsKout2_bg’,…
    graphs2files, FigDir);
bg_root = ‘bg_Kout1vsKout2’;
quad_Kout1vsKout2 = quad_subplot(ContKout1, ContKout2, bg_Kout1vskKout2, ‘quad_Kout1vsKout2’, bg_root, …
    graphs2files, FigDir);
if show_graphics
    quad_Kout1vsKout2.Visible = ‘on’;
end
~~~

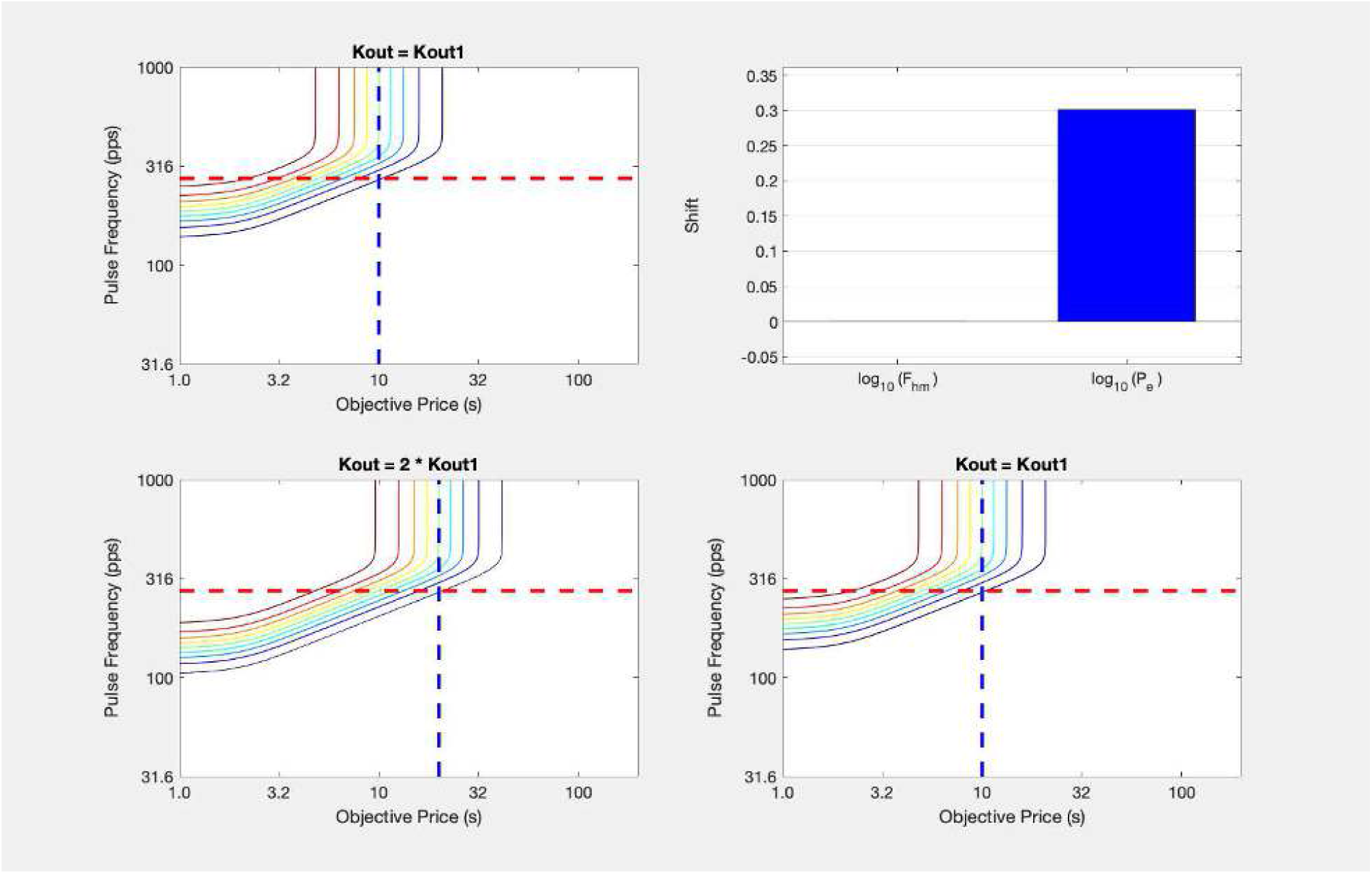

~~~
[fig_num, fig_tab] = add_fig(fig_num, fig_tab, “quad_Kout1vsKout2”, “Effect of increasing the value of the output-scaling paramater”);
~~~

`This is figure #27: (quad_Kout1vsKout2)`

The graphs shown above demonstrate that changes to the input- and output-scaling parameters of the power-growth function produce identical shifts in the position of the reward mountain. Changing the value of either parameter shifts the mountain rightward along the price axis, in sharp contrast to the orthogonal shifts observed when reward intensity grows as a logistic function of pulse frequency.

In the case of eICSS, we know from the experiments of Gallistel’s group that reward intensity grows in a manner similar to a logistic function of pulse frequency. Unlike the case of power growth, logistic growth is characterized by independent parameters that scale the input and output. This characteristic is what causes the effect of varying reward probability to shift the reward mountain in a direction orthogonal to the effects of varying current or train duration. It follows that when such orthogonal shifts are observed, this requires that the underlying reward-growth function incorporates independent parameters that scale the input (the position parameter) and the output (the parameter representing the maximum reward intensity). Thus even in the absence of data from matching experiments or other methods for measuring the growth of reward intensity, we can make inferences about the reward-growth function by observing the way in which various manipulations shift the reward mountain. This will be important for the interpretation of the oICSS data.

`Elapsed time is 18.280454 seconds.`

## At what stage of processing does perturbation of dopaminergic neurotransmission alter reward seeking in eICSS?

~~~
tic;
close all;
~~~

We have described changes in the position parameter 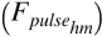 as alterations in the ***sensitivity*** of the reward- growth function. One might best label 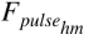 as the “inverse-sensitivity” parameter because the lower its value, the weaker the input required to drive reward intensity to a given level. In contrast, we have described changes in the output-scaling parameter (***K***_*rg*_) as alterations in the ***gain*** of the reward-growth function. Just as turning the volume knob on an audio amplifier changes the loudness produced by weak and strong inputs by the same percentage, changing ***K***_*rg*_ alters the reward intensity produced by both low and high pulse frequencies (within the frequency-following range) by the same percentage. Changes in sensitivity shift the reward mountain along the pulse-frequency axis, whereas changes in gain shift the mountain along the price axis.

Operant peformance in the eICSS paradigm is altered by perturbation of dopaminergic neurotransmission. Boosting dopaminergic signalling typically enhances performance, whereas attenuating dopaminergic signalling typically attenuates performance. These changes have long been attributed to the modulation of reward sensitivity by dopaminergic agents (See: Hernandez et al., 2010). If so, drugs that alter dopaminergic neurotransmission should shift the reward mountain along the pulse-frequency axis and not along the price axis. We have shown that this is not so. In rats working for electrical stimulation of the medial forebrain bundle, we demonstrated that enhancement of dopaminergic signalling by the highly specific reuptake blocker, GBR-12909, shifted the reward mountain rightward along the price axis in 8/10 rats without producing any statistically reliable shifts along the pulse-frequency axis (Hernandez et al., 2012). The D2/D4/5HT7 receptor blocker, pimozide, shifted the reward mountain leftward along the price axis in 5/6 rats without producing any statistically reliable shifts along the pulse-frequency axis (Trujillo-Pisanty, Conover & Shizgal, 2014). To account for these complementary effects, the changes in dopaminergic neurotransmission had to have altered one or more of the variables that determine the position of the reward mountain along the price axis, such as the output-scaling parameter of the reward-growth function (***K***_*rg*_), the subjective effort cost 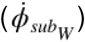, or the payoff from alternate activities (*U*_*L*_).

~~~
if show_graphics
    show_imported_graphic(‘GBR_pimozide_summary.png’, 100, ImpFigDir);
end
~~~

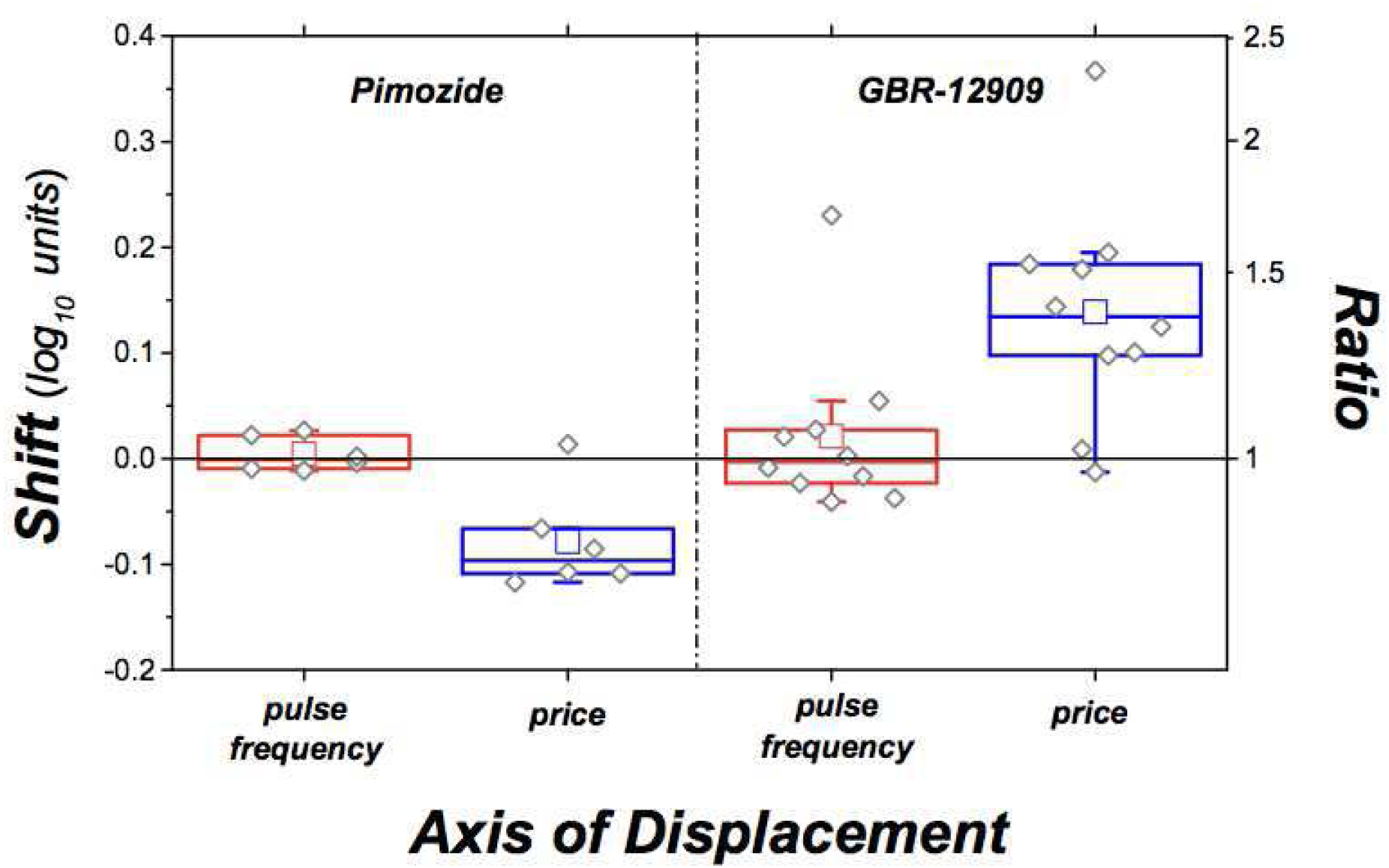

~~~
[fig_num, fig_tab] = add_fig(fig_num, fig_tab, “PimGBRshifts”, …
   “Changes in dopaminergic neurotransmission shift the mountain along the price axis”);
~~~

`This is figure #28: (PimGBRshifts)`

An intuitively appealing interpretation of these data is based on the notion that the neurons subserving eICSS of the medial forebrain bundle project to midbrain dopamine neurons and that it is the transynaptic activation of the dopamine neurons that gives rise to the reward effect of the medial forebrain bundle stimulation (the “series” model of brain-reward circuitry). In the following section, we apply the reward mountain model to the oICSS data from the present paper. We show that the series model ***cannot*** account for the data from both the eICSS and oICSS studies. After demonstrating this, we discuss “convergence” models that show promise for an intergrated account of both eICSS and oICSS, and we propose experiments that put such accounts to empirical test.

~~~
toc
~~~

`Elapsed time is 0.748551 seconds.`

## Optical intracranial self-stimulation of midbrain dopamine neurons

~~~
tic;
close all;
~~~

In the experiments reviewed above, rats worked to trigger trains of electrical current pulses applied to the medial forebrain bundle. In contrast, in the experiment we now discuss, rats worked to trigger trains of optical pulses delivered through an optical fiber positioned over the ventral tegmental area of the midbrain. As a result of viral transcription and Cre-Lox recombination, dopamine neurons with somata in, or adjacent to, the VTA expressed channelrhodopsin-2 (ChR2) and thus could be excited by light delivered at an appropriate wavelength (473 nm). As others have reported previously (Witten et al., 2012), rats expressing ChR2 in midbrain dopamine neurons work vigorously to obtain such optical stimulation.

We show that TH-Cre rats expressing ChR2 in midbrain dopamine neurons learn to perform the cumulative hold-down task to receive optical stimulation of the ventral tegmental area (VTA), where the somata of dopamine neurons projecting to forebrain targets reside. As in the case of eICSS, time allocation varies smoothly as a function of the strength and price of the optical stimulation. The surface defined by the mountain model fits the data well.

~~~
if show_graphics
    show_imported_graphic(‘Bechr29_veh_mtn.png’, 30, ImpFigDir);
end
~~~

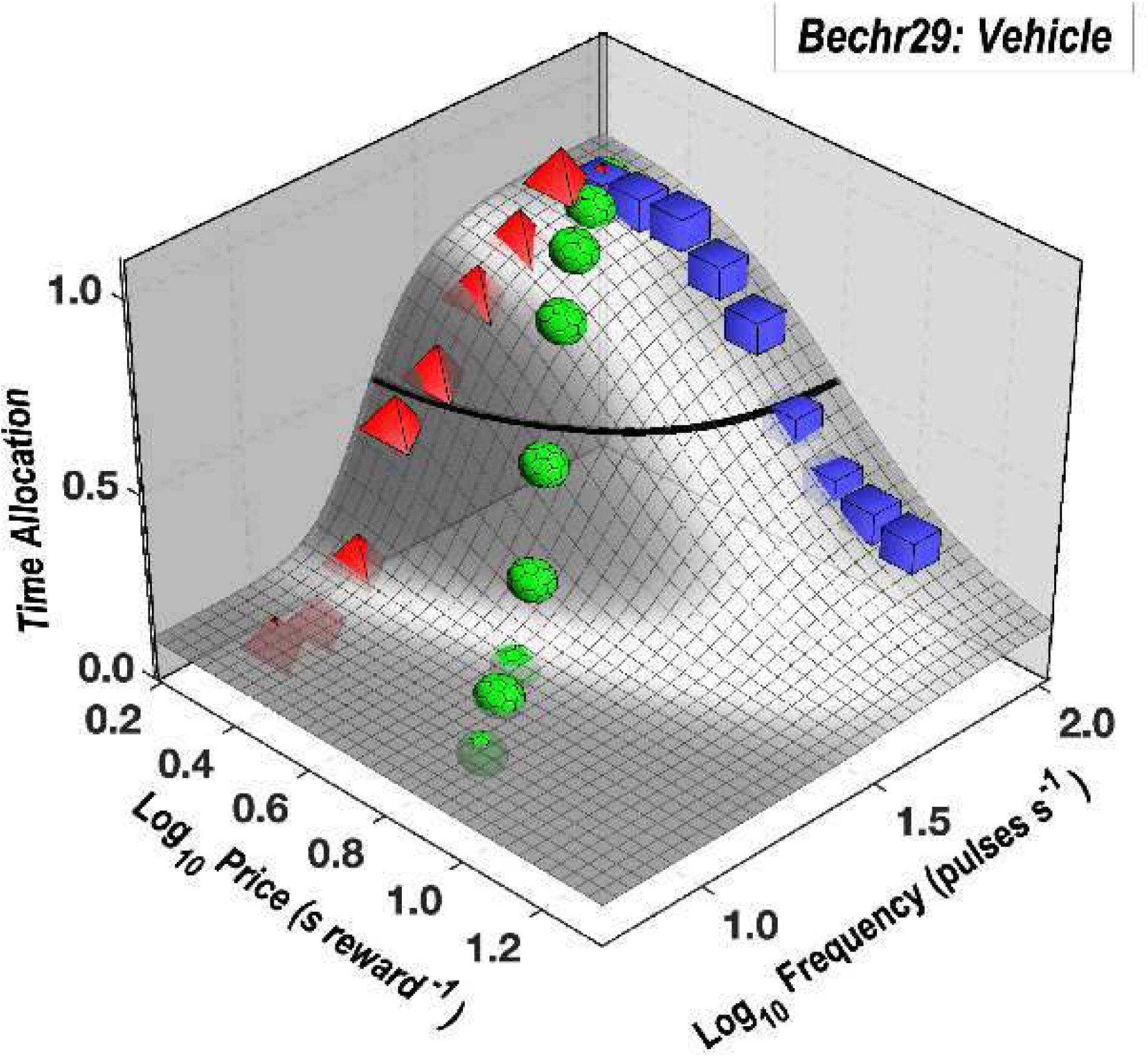

~~~
[fig_num, fig_tab] = add_fig(fig_num, fig_tab, “BeChR29vehMtn”, …
    “The mountain model fitted to vehicle-condition data from rat BeChR29”);
~~~

`This is figure #29: (BeChR29vehMtn)`

### The effect of the dopamine-transporter blocker, GBR-12909

To boost dopaminergic neurotransmission, we administered 20 mg/kg of the dopamine-transporter blocker, GBR-12909. Under the influence of the drug, the mountain fitted to the data from rat BeChR29 shifted leftwards along the pulse-frequency axis and rightwards along the price axis.

~~~
if show_graphics
    show_imported_graphic(‘Bechr29_dual_mountains.png’, 30, ImpFigDir);
end
~~~

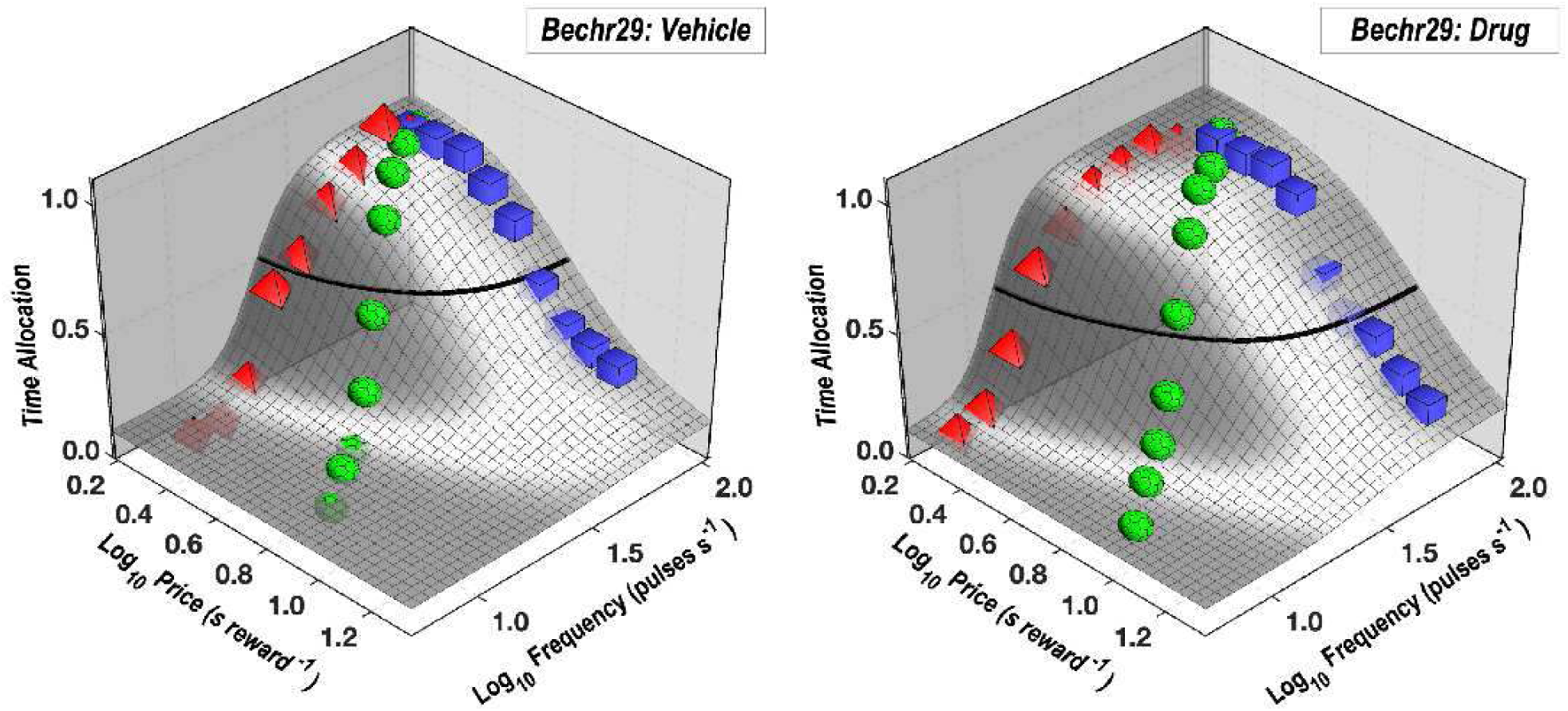

~~~
[fig_num, fig_tab] = add_fig(fig_num, fig_tab, “BeChR29vehdrgMtn”, …
    “Effect of GBR-12909 on the reward-mountain data obtained from rat BeChR29”);
~~~

`This is figure #30: (BeChR29vehdrgMtn)`

These shifts can be most clearly discerned in the contour-plot display and bargraph summary:

~~~
if show_graphics
    show_imported_graphic(‘BeChR29_quad_GBR.png’, 25, ImpFigDir);
end
~~~

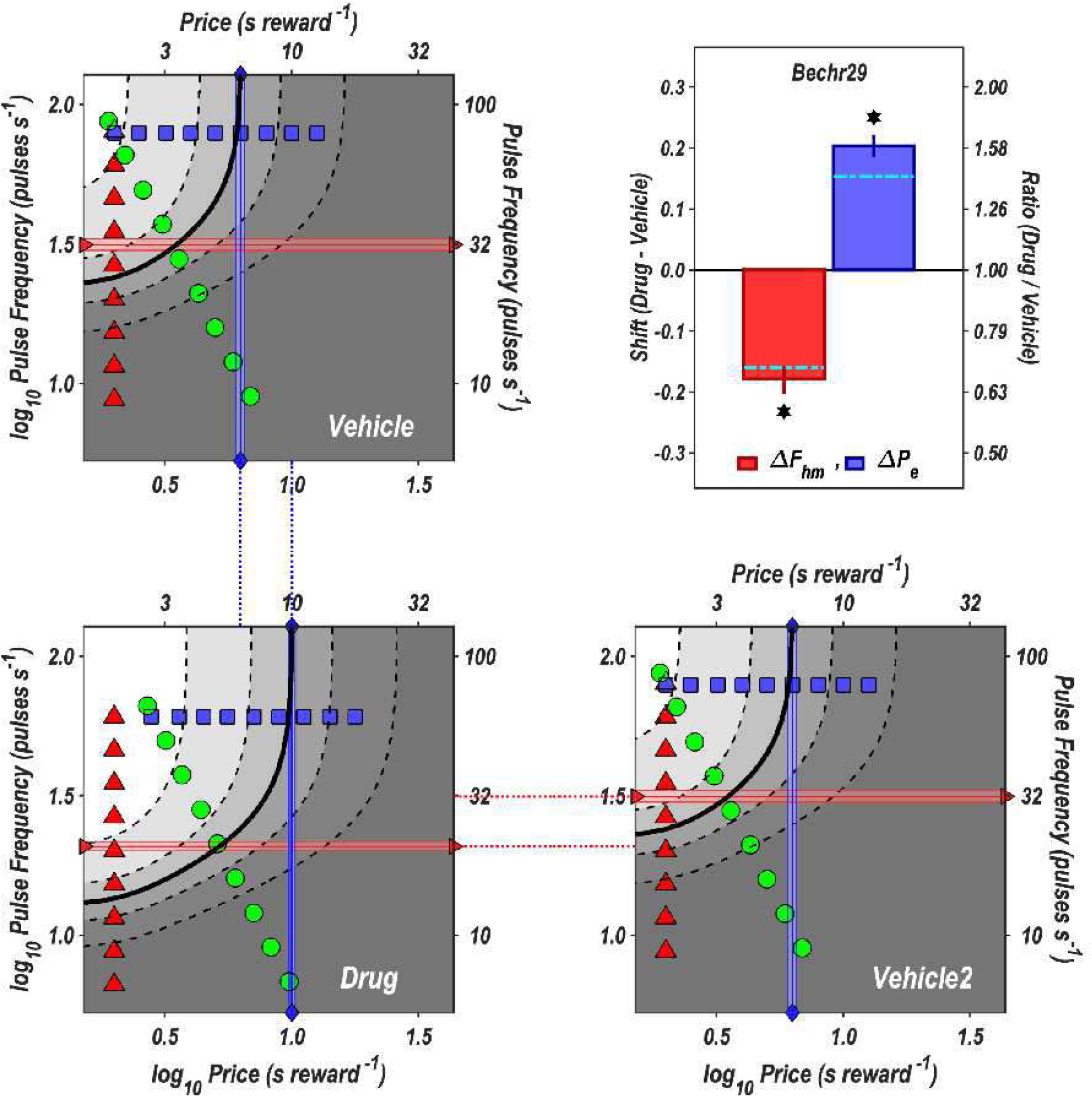

~~~
[fig_num, fig_tab] = add_fig(fig_num, fig_tab, “BeChR29quad”, …
    “Effect of GBR-12909 on the reward-mountain data obtained from rat BeChR29”);
~~~

`This is figure #31: (BeChR29quad)`

This pattern of shifts is seen in the results from 6 of the 7 rats:

~~~
if show_graphics
    show_imported_graphic(‘optoGBR_shift_summary.png’, 100, ImpFigDir);
end
~~~

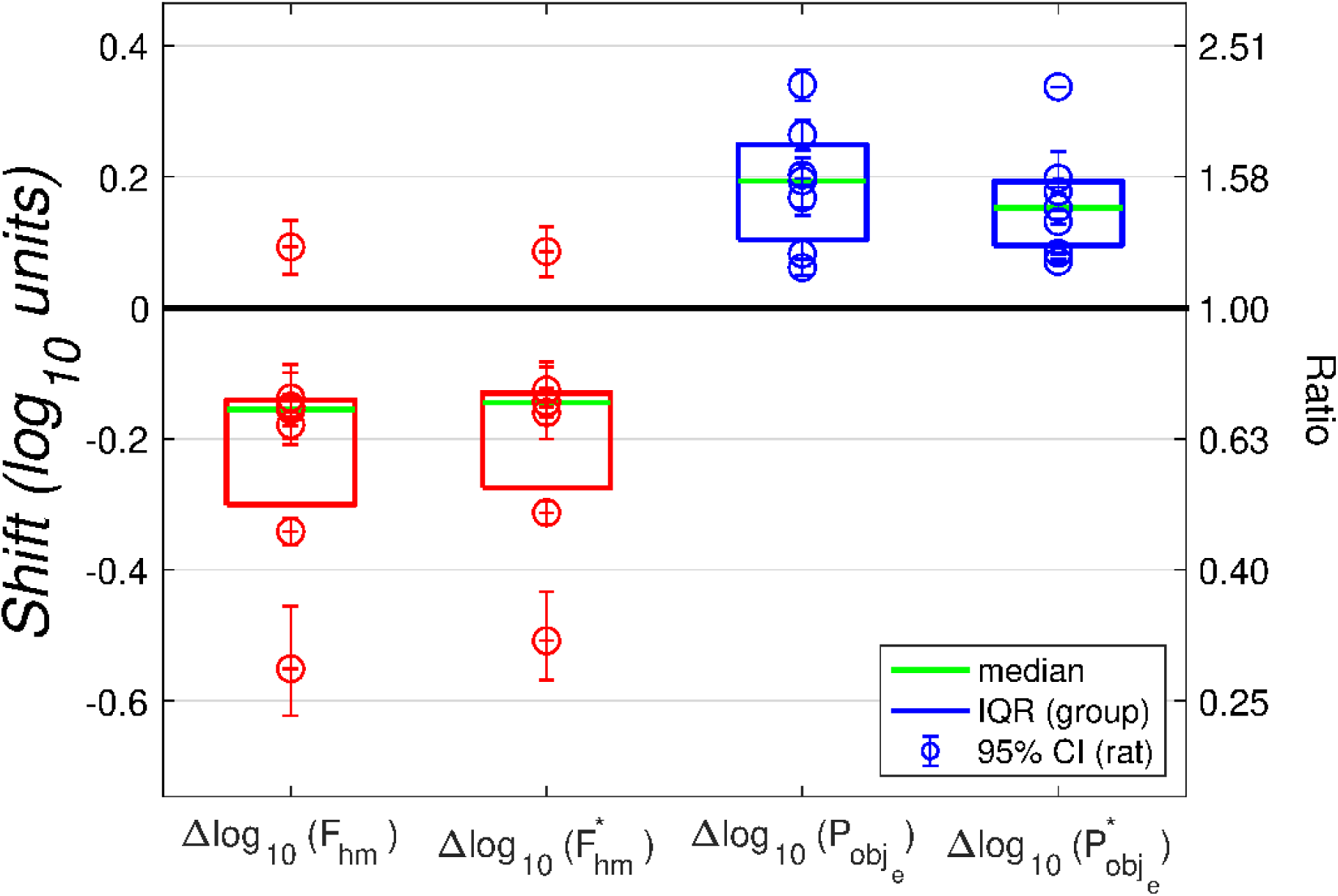

~~~
[fig_num, fig_tab] = add_fig(fig_num, fig_tab, “GBRshiftSummary”, …
    “Effect of GBR-12909 on the reward-mountain data obtained from all 7 rats”);
~~~

`This is figure #32: (GBRshiftSummary)`

The error bars surrounding the data point for each rat represent the bootstrapped 95% confidence interval (CI) for the parameter in question, whereas the boxes denoted by the heavy solid lines represent the inter-quartile range (IQR) of the median parameter estimate for the group of seven rats. The 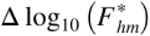 and 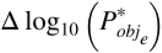 values have been corrected for differential frequency-following fidelity in the vehicle and drug conditions. (See below and main body of the manuscript.)

### Modeling the results

We now undertake modeling to determine how the optical pulse train can be translated into the observed behavioral responses in a manner consistent with the observed effect of dopamine-transporter blockade on the reward mountain.

The first step is to relate the induced frequency of firing in the dopamine neurons to the optical pulse frequency. For this purpose, we adopt a filter function of the form defined by

~~~
disp(string({strcat({‘Function ‘},num2str(fun_tab.Number(fun_tab.Name==‘FilterFun’)))}));
~~~

`Function 6`

, which provides an accurate description of how medial forebrain bundle fibers subserving electrical intracranial self-stimulation respond to the electrical pulse frequency. It is not known how well the functional form describes frequency-following in optically activated, ChR2-expressing, midbrain dopamine neurons. As explained in the main body of the manuscript, parameter values have been chosen on the basis of published electrophysiological, voltammetric, and behavioral data.

The maximum firing frequency in the dopamine neurons is far lower than in the directly stimulated neurons responsible for the rewarding effect of electrical medial forebrain bundle stimulation (Solomon et al., 2015). To reflect this in the equation for the frequency response, we used updated names and values for the parameters:

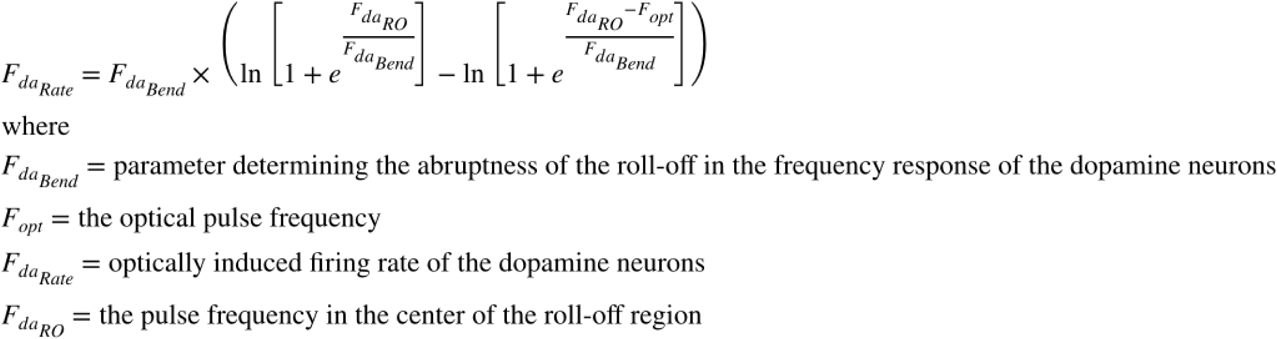

~~~
[eqn_num, eqn_tab] = …
    add_eqn(eqn_num, eqn_tab, “FreqFolDA”, “Frequency of firing as a function of pulse frequency”);
~~~

`This is equation #29: (FreqFolDA)`

~~~
[sym_num, sym_tab] = add_sym(sym_num, sym_tab, “FdaBend”, “parameter determining sharpness of bend in FreqFol function”);
[sym_num, sym_tab] = add_sym(sym_num, sym_tab, “Fopt”, “optical pulse frequency”);
[sym_num, sym_tab] = add_sym(sym_num, sym_tab, “FdaRate”, “optically induced firing rate of the dopamine neurons”);
[sym_num, sym_tab] = add_sym(sym_num, sym_tab, “FdaRO”, “the pulse frequency in the center of the roll-off region”);
~~~

In later stages of the modeling, we use the inverse function, FilterFunBS, to return an optical pulse frequency given the dopamine firing frequency it produces. (see ***Functional building blocks for the simulations)***.

The following graph show the function employed to model frequency-following fidelity in the ChR2-expressing dopamine neurons in response to the optical pulse frequency:

~~~
FdaBend = 20;
FdaRO = 50;
logFopt = 0:0.025:2.4;
Fopt = 10.^logFopt;
FRda = FilterFun(Fopt, FdaBend, FdaRO); % Compute firing rate using the frequency roll-off function
FF_graphDA = plot_freqFoll(Fopt,FRda,’DA’,1,250,1,100);
if show_graphics
    FF_graphDA.Visible = ‘on’;
end
~~~

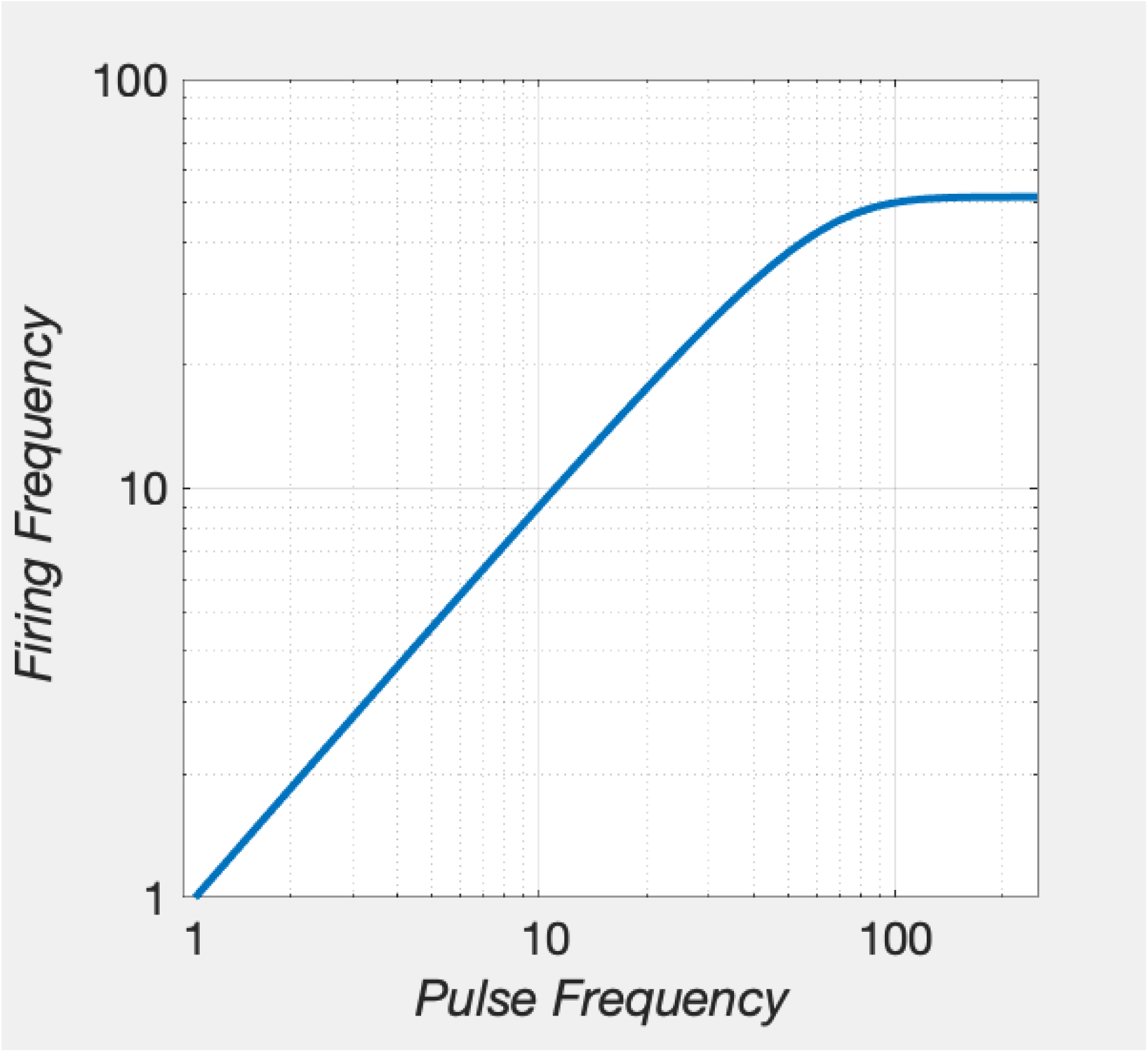

~~~
[fig_num, fig_tab] = add_fig(fig_num, fig_tab, “FreqFolDA”, …
    “Induced firing frequency in dopamine neurons as a function of pulse frequency”);
~~~

`This is figure #33: (FreqFolDA)`

## The spike counter

Performance for optical stimulation of midbrain dopamine neurons depends both on the induced firing frequency and the number of dopamine neurons excited by the optical input (Ilango et al., 2014). The simplest assumption consistent with this finding is that the behavioral effects depend on the aggregate rate of induced firing, as is the case of the behavioral effects produced by rewarding electrical stimulation of the medial forebrain bundle. The aggregate rate of firing induced by the optical stimulation in the population of dopamine neurons is the product of the number of optically activated neurons and the induced firing frequency:

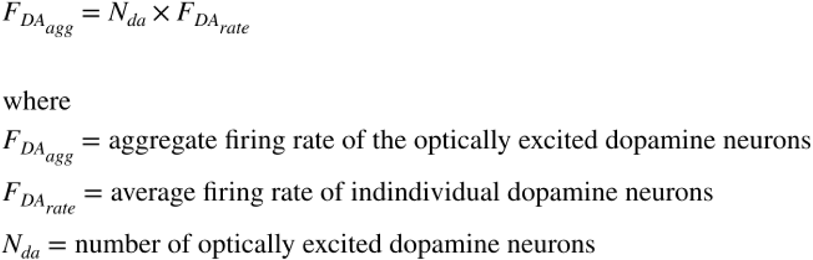

~~~
[eqn_num, eqn_tab] = …
   add_eqn(eqn_num, eqn_tab, “AggDAfr”, “Aggregate firing rate in the dopamine neurons”);
~~~

`This is equation #30: (AggDAfr)`

~~~
[sym_num, sym_tab] = add_sym(sym_num, sym_tab, “FdaAgg”, “aggregate firing rate of the optically excited dopamine neurons”);
[sym_num, sym_tab] = add_sym(sym_num, sym_tab, “Nda”, “number of optically excited dopamine neurons”);
clear a Fopt FRda logFopt;
clear -regexp ^Fda ^FF_graph;
toc
~~~

`Elapsed time is 3.287460 seconds.`

### The effect on the reward mountain produced by modulation of dopaminergic neurotransmission

~~~
tic;
close all;
~~~

The influence of the induced firing of the dopamine neurons can be modulated by drugs, such as transporter blockers, that alter synaptic transmission. To capture such effects, we add a scalar at the output of the dopamine spike count:

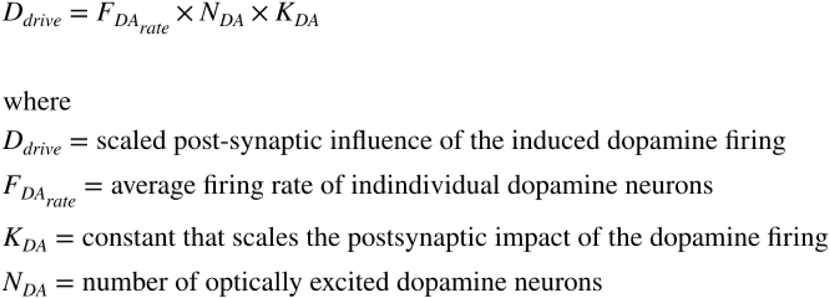

~~~
[eqn_num, eqn_tab] = …
    add_eqn(eqn_num, eqn_tab, “DAdrive”, “scaled post-synaptic influence of the induced dopamine firing”);
~~~

`This is equation #31: (DAdrive)`

~~~
[sym_num, sym_tab] = add_sym(sym_num, sym_tab, “Kda”, “constant that scales the postsynaptic impact of the dopamine firing”);
~~~

`Symbol Kda has already been entered.`

In the following schema, the aggregate rate of firing is represented by the Π symbol and the dopamine-drive scalar by a triangle:

~~~
if show_graphics
    show_imported_graphic(‘counter_opto.png’,60,ImpFigDir);
end
~~~

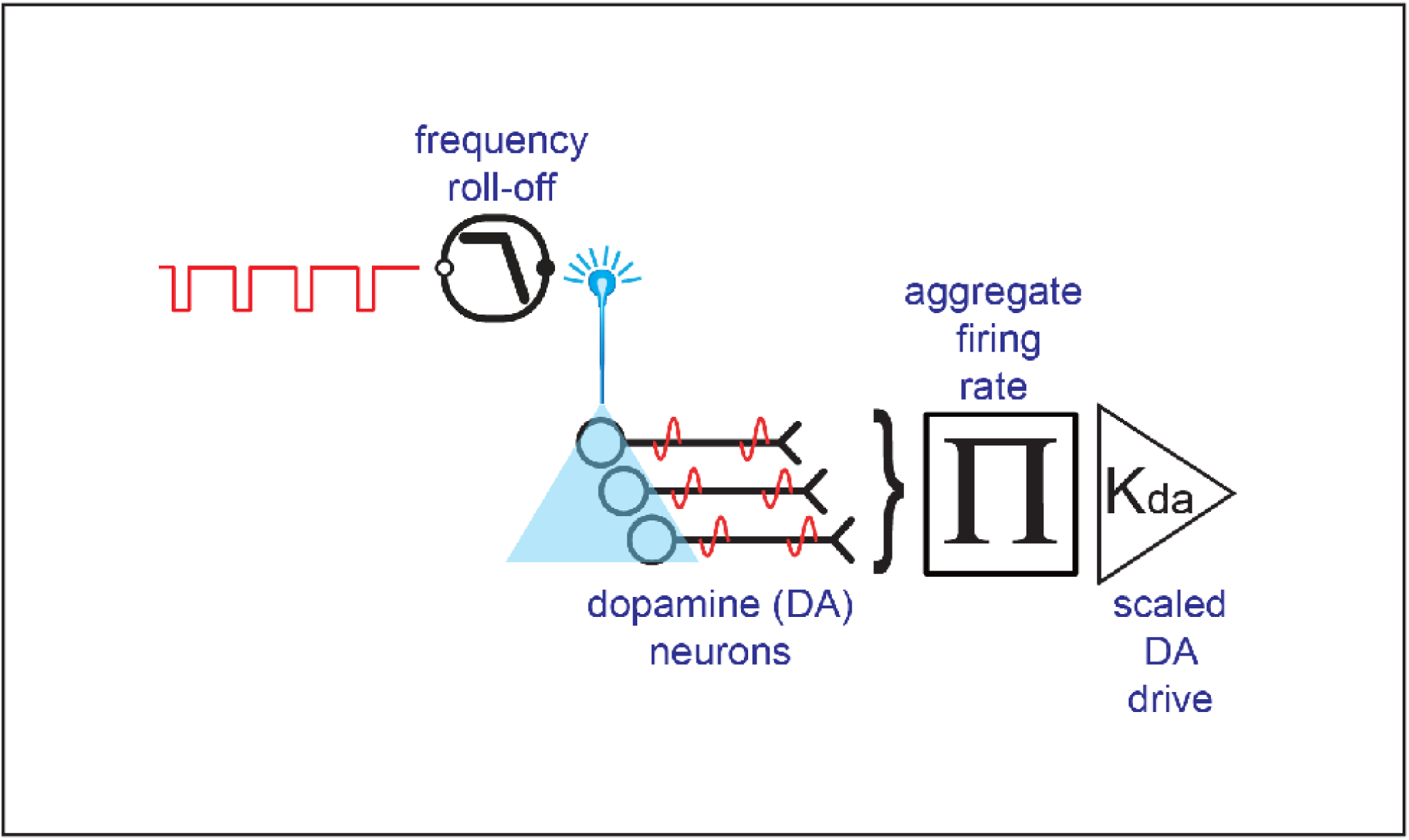

~~~
[fig_num, fig_tab] = add_fig(fig_num, fig_tab, “DAdrive”, …
    “Influence of optically stimulated dopamine release on downstream targets”);
~~~

`This is figure #34: (DAdrive)`

We refer to the scaled output of the activated population of midbrain dopamine neurons as “dopamine drive.”

### The reward-growth function for oICSS

In six of the seven rats, dopamine-transporter blockade shifted the mountain leftward along the pulse-frequency axis. As we demonstrated above, such shifts require that the function that translates dopamine drive into reward intensity (i.e., the reward-growth function) must have a position parameter that is independent of the parameter that sets the maximum reward intensity attainable. The logistic function described by Gallistel’s group in the case of eICSS has this property, and we have therefore added such a function at the output of the counter.

~~~
if show_graphics
    show_imported_graphic(‘DA_logisitc_simplified.png’,50,ImpFigDir);
end
~~~

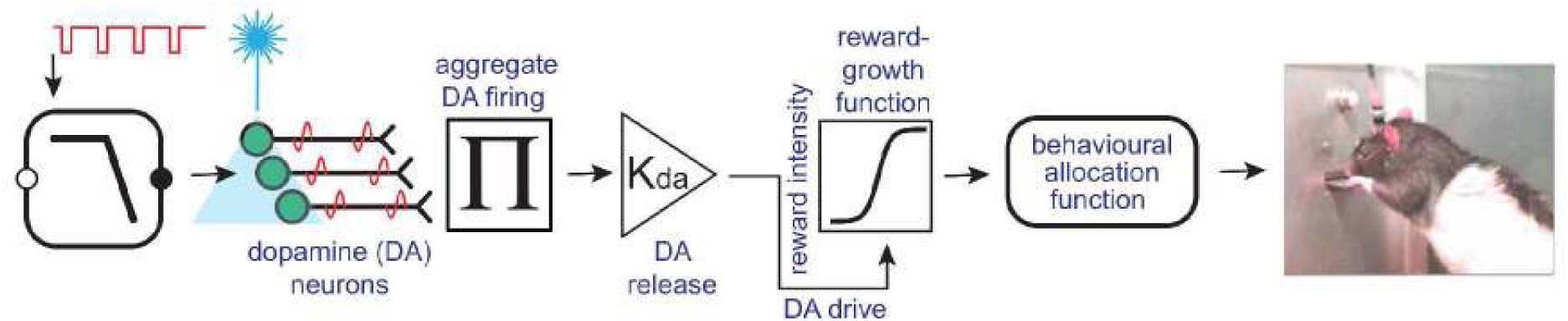

~~~
[fig_num, fig_tab] = add_fig(fig_num, fig_tab, “oICSS_simplified”, …
     “simplied oICSS model with logistic reward growth”);
~~~

`This is figure #35: (oICSS_simplified)`

In the case of eICSS, the form of the two-dimensional reward-growth function employed in the simulations is based on the empirical measurements obtained by Gallistel’s group. No such measurements have yet been obtained for oICSS. Future experiments will be required to determine whether the logistic form or another form with independent input- and output-scaling parameters provides the best description of reward growth in the case of oICSS.

In the case of eICSS, the reward-growth function has been generalized to three dimensions {reward intensity, pulse frequency, train duration} on the basis of additional empirical data (Sonnenschein, Conover, & Shizgal, 2003). Both the pulse frequency and train duration have been manipulated in oICSS experiments (Ilango et al., 2014), but not in a way that makes it possible to describe the joint dependence of reward intensity on these two variables. Future experiments will be required to determine this. For present purposes, we will again adopt the function derived from eICSS data for use in simulating oICSS performance.

We define the position parameter of the reward-growth function for oICSS of midbrain dopamine neurons by modifying

~~~
disp(string({strcat({‘Equation ‘},num2str(eqn_tab.Number(eqn_tab.Name==‘fH’)))}));
~~~

`Equation 17`

as follows:

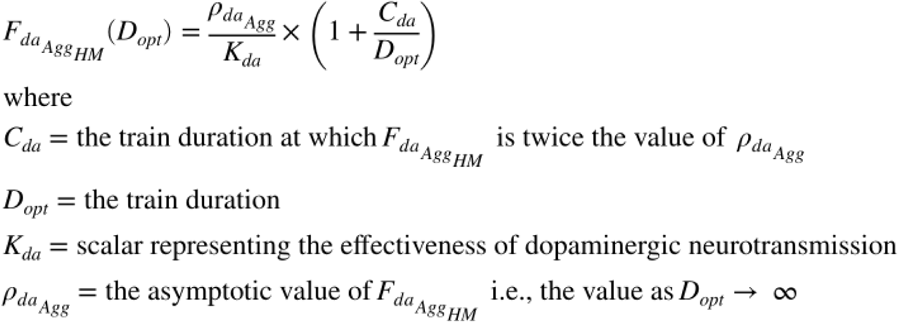

~~~
[eqn_num, eqn_tab] = …
   add_eqn(eqn_num, eqn_tab, “FHda”, “strength-duration function for optical trains delivered to dopamine neurons”);
~~~

`This is equation #32: (FHda)`

~~~
[sym_num, sym_tab] = add_sym(sym_num, sym_tab, “Cda”, “chronaxie of the strength-duration function for trains delivered to dopamine neurons”);
[sym_num, sym_tab] = add_sym(sym_num, sym_tab, “Dopt”, “duration of an optical pulse train”);
[sym_num, sym_tab] = add_sym(sym_num, sym_tab, “RhoDA”, “rheobase of the strength-duration function for trains delivered to dopamine neurons”);
~~~

`The function FpulseHMfun`

~~~
disp(string({strcat({‘Function ‘},num2str(fun_tab.Number(fun_tab.Name==‘FpulseHMfun’)))}));
~~~

`Function 30`

(`See` ***Functions composing the reward-mountain model*** `below`), `accepts an argument list of variable length`. `If` ***K***_*da*_ is appended to the end of the argument list, then it will enter into the calculation of 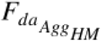 as specified in the above equation. If ***K***_*da*_ is omitted, it assumes an implicit value of one.

The following code produces estimates of 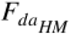 for two values of ***K***_*da*_. The lower value (1) represents the effectiveness of dopaminergic neurotransmission in the vehicle condition, whereas the higher value (2) represents the effectiveness of dopaminergic neurotransmission under the influence of dopamine-transporter blockade.

~~~
C = 0.473; % median from Sonnenschein et al., 2003 D = 1.0; % train duration for oICSS study
FdaBend = 20;
FdaRO = 50;
Kda = [1,10^.2]; NnDA = 100;
RhoPiDA = 1572; % See the calculation in the Series-circuit section below for D = 1 s FPmax = 1000;
FdaHMkDA = FpulseHMfun(C, D, FdaBend, FPmax, FdaRO, NnDA, RhoPiDA, Kda)
~~~

~~~
FdaHMkDA = 1×2
    27.1046 16.4598
~~~

~~~
%FdaHMkDA is a two-element vector
~~~

Note that although Kda(1) is half the value of Kda(2), FdaHMkDA(2) is less than half the value of FdaHMkDA(1). The reason for this is that the frequency-response function for the dopamine neurons has already started to roll off by FdaHMkDA(1) (some dopamine neurons fail to fire once per pulse at this pulse frequency). This will be reflected in a greater discrepancy between RnormMax(1) and one than between RnormMax(2) and one.

To compute time allocation, we also require the value of the parameter that positions the mountain along the price axis, 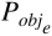. In the initial simulation, we assume that the value of this parameter is unaffected by the blockade of the dopamine transporter.

~~~
dotPhiObj = 1;
64
Kaa = 1;
Kec = 1;
Krg = 1;
dotRaa = 0.1;
pObj = 1;
PsubBend = 0.5;
PsubMin = 1.82;
FPmax = 1000; % This value is sufficiently above FdaRO to maximize the firing rate g = 5;
RnormMax = fRbsrNorm(FPmax, FdaBend, FdaHMkDA, FdaRO, g)
~~~

~~~
RnormMax = 1×2
  0.9615 0.9967
~~~

~~~
% RnormMax is a two-element vector
PobjE = PobjEfun(dotPhiObj, Kaa, Kec, Krg, dotRaa, pObj, PsubBend, PsubMin, RnormMax)
~~~

~~~
PobjE = 1×2
  9.6147 9.9670
~~~

~~~
% PobjE is a two-element vector
~~~

We can now estimate time allocation and simulate the reward mountains for the vehicle and drug conditions using the full logistic reward-growth model shown in the figure below:

~~~
if show_graphics
  show_imported_graphic(‘oICSS_logistic+3Pe_noMasks_v4.png’,25,ImpFigDir);
end
~~~

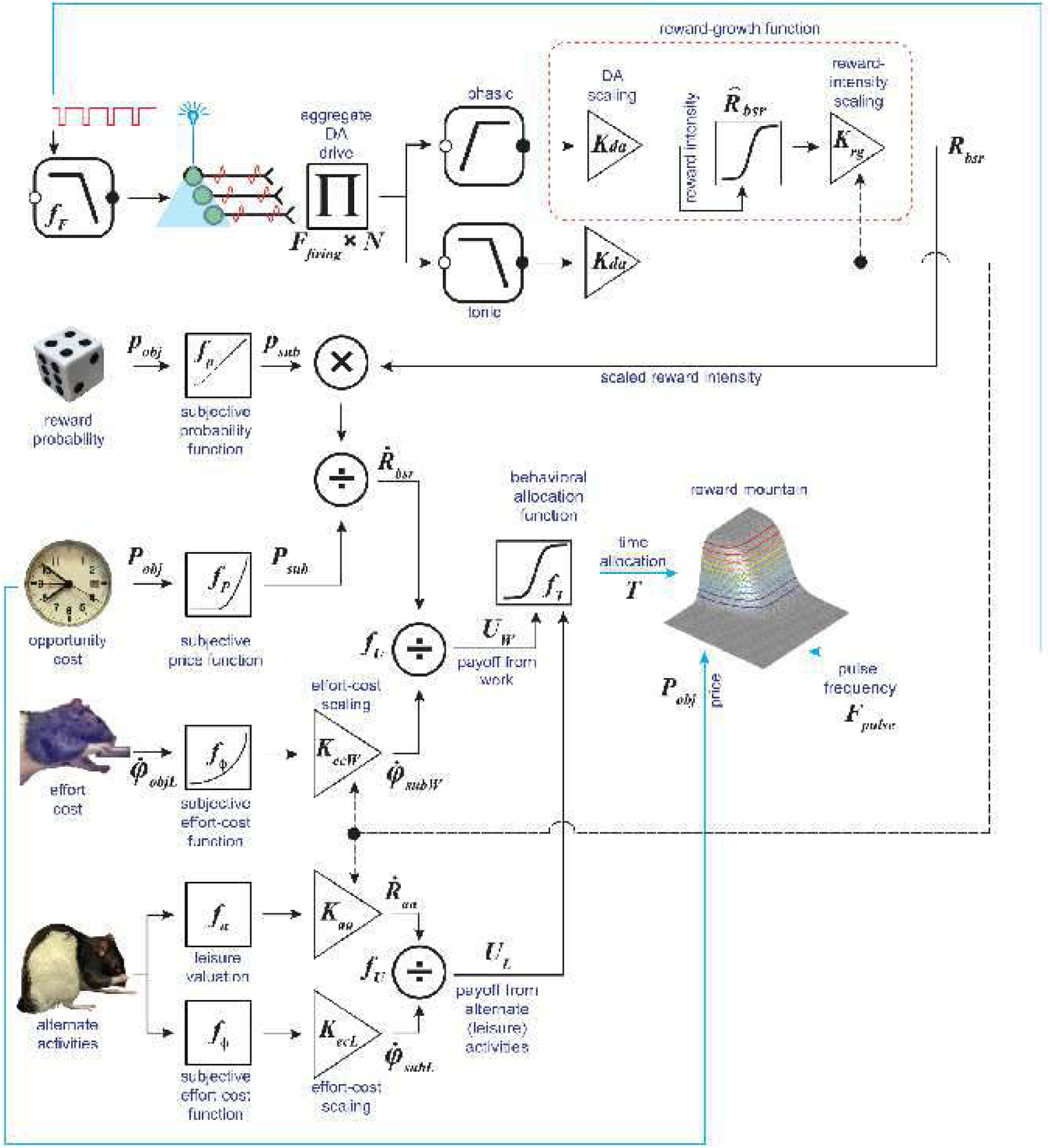

~~~
[fig_num, fig_tab] = add_fig(fig_num, fig_tab,
    “oICSS_logistic”, … “oICSS model with logistic reward growth”);
~~~

`This is figure #36: (oICSS_logistic)`

We will now simulate the output of this model, disregarding, for the time being, any drug-induced changes in dopamine tone.

~~~
a = 3;
Fopt = logspace(0,3,121)’; % column variable
Pobj = logspace(0,3,121); % row variable
Tda1 = TAfun(a, Fopt, FdaBend, FdaHMkDA(1), FdaRO, g, Pobj, PobjE(1), …
  PsubBend, PsubMin, RnormMax(1));
Tda2 = TAfun(a, Fopt, FdaBend, FdaHMkDA(2), FdaRO, g, Pobj, PobjE(2), …
  PsubBend, PsubMin, RnormMax(2));
~~~

Before plotting the bar graphs, we need to correct the estimates of the location parameters for the differential frequency-following fidelity in the drug and vehicle conditions. The mountain is shifted downwards along the pulse-frequency axis in the drug condition and thus, frequency-following fidelity is better than in the vehicle condition.

As we explain in the main body of the manuscript,

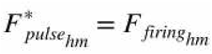

and

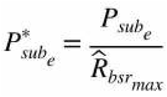

~~~
FdaHMkDAstar = FilterFun(FdaHMkDA,FdaBend,FdaRO);
PsubE = PsubFun(PobjE,PsubBend,PsubMin);
PsubEstar = PsubE ./ RnormMax;
PobjEstar = PsubBsFun(PsubEstar,PsubBend,PsubMin);
title_str1 = strcat({‘Kda = ‘}, sprintf(‘%2.2f’, Kda(1))); xmin = 0;
xmax = 2.3;
ymin = 0.3;
ymax = 2.3;
MTNkDA1 = plot_MTN(Fopt, Pobj, Tda1, ‘off’, ‘MTNkDA1’title_str1, …
  graphs2files, FigDir, xmin, xmax, ymin, ymax);
title_str2 = strcat({‘Kda = ‘}, sprintf(‘%2.2f’, Kda(2)));
MTNkDA2 = plot_MTN(Fopt, Pobj, Tda2, ‘off’, ‘MTNkDA2’, title_str2, …
  graphs2files, FigDir, xmin, xmax, ymin, ymax);
MTNkDA1vskDA2 = dual_subplot(MTNkDA1, MTNkDA2, ‘MTNkDA1vskDA2’,… graphs2files,FigDir);
if show_graphics
    MTNkDA1vskDA2.Visible = ‘on’;
end
~~~

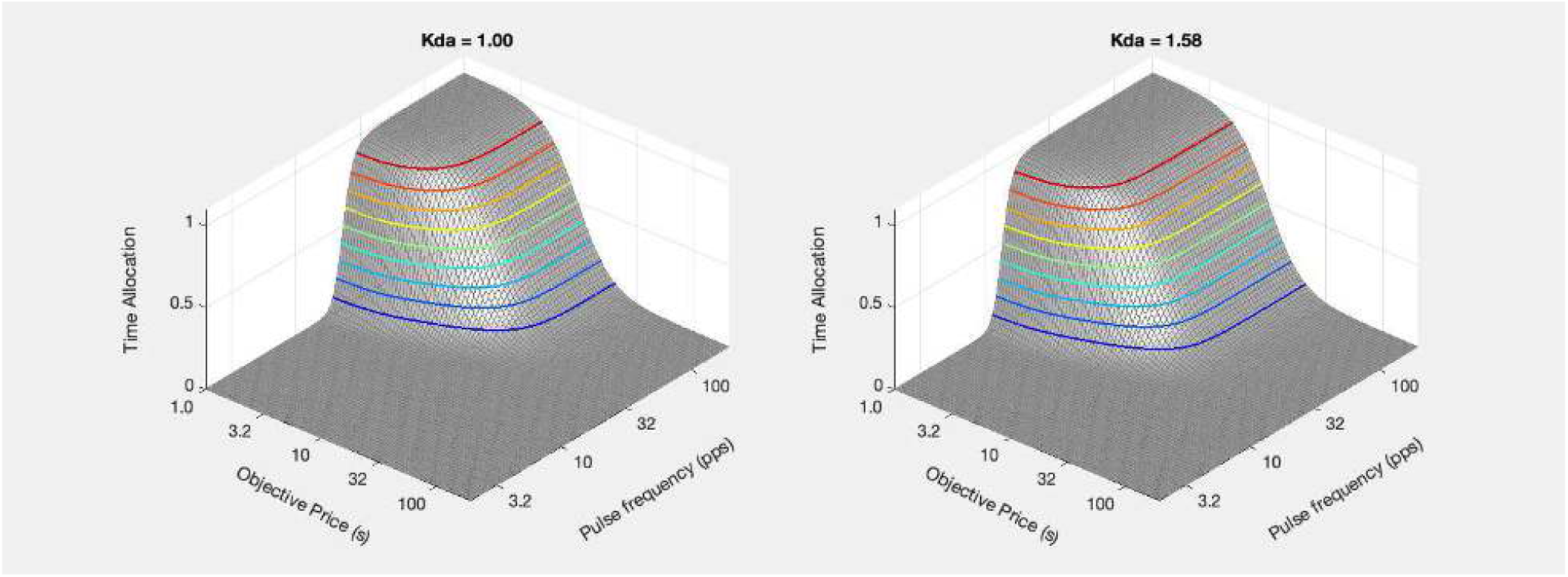

~~~
[fig_num, fig_tab] = add_fig(fig_num, fig_tab, “kDA1vskDA2_mtns”, …
  “Effect of dopamine-transporter blockade on the reward mountain”);
~~~

`This is figure #37: (kDA1vskDA2_mtns)`

~~~
ContkDA1 = plot_contour(Fopt, Pobj, Tda1, PobjE(1), FdaHMkDA(1), ‘off’, ‘ContkDA1’, title_str1, …
   strcat({‘Kda = ‘},num2str(Kda(1))), graphs2files, FigDir,…
   xmin, xmax, ymin, ymax);
ContkDA2 = plot_contour(Fopt, Pobj, Tda2, PobjE(2), FdaHMkDA(2), ‘off’, ‘ContkDA2’, title_str2, …
   strcat({‘Kda = ‘}, num2str(Kda(2))), graphs2files, FigDir,…
   xmin, xmax, ymin, ymax);
bg_kDA1vskDA2 = plot_bgStar(FdaHMkDA(1), FdaHMkDA(2), FdaHMkDAstar(1), FdaHMkDAstar(2),…
   PobjE(1), PobjE(2), PobjEstar(1), PobjEstar(2),…
   ‘off’, ‘kDA1vskDA2_bg’, graphs2files, FigDir);
bg_root = ‘bg_kDA1vskDA2’;
quad_kDA1vskDA2 = quad_subplot(ContkDA1, ContkDA2, bg_kDA1vskDA2, ‘quad_kDA1vskDA2’, bg_root, … graphs2files, FigDir);
if show_graphics
   quad_kDA1vskDA2.Visible = ‘on’;
end
~~~

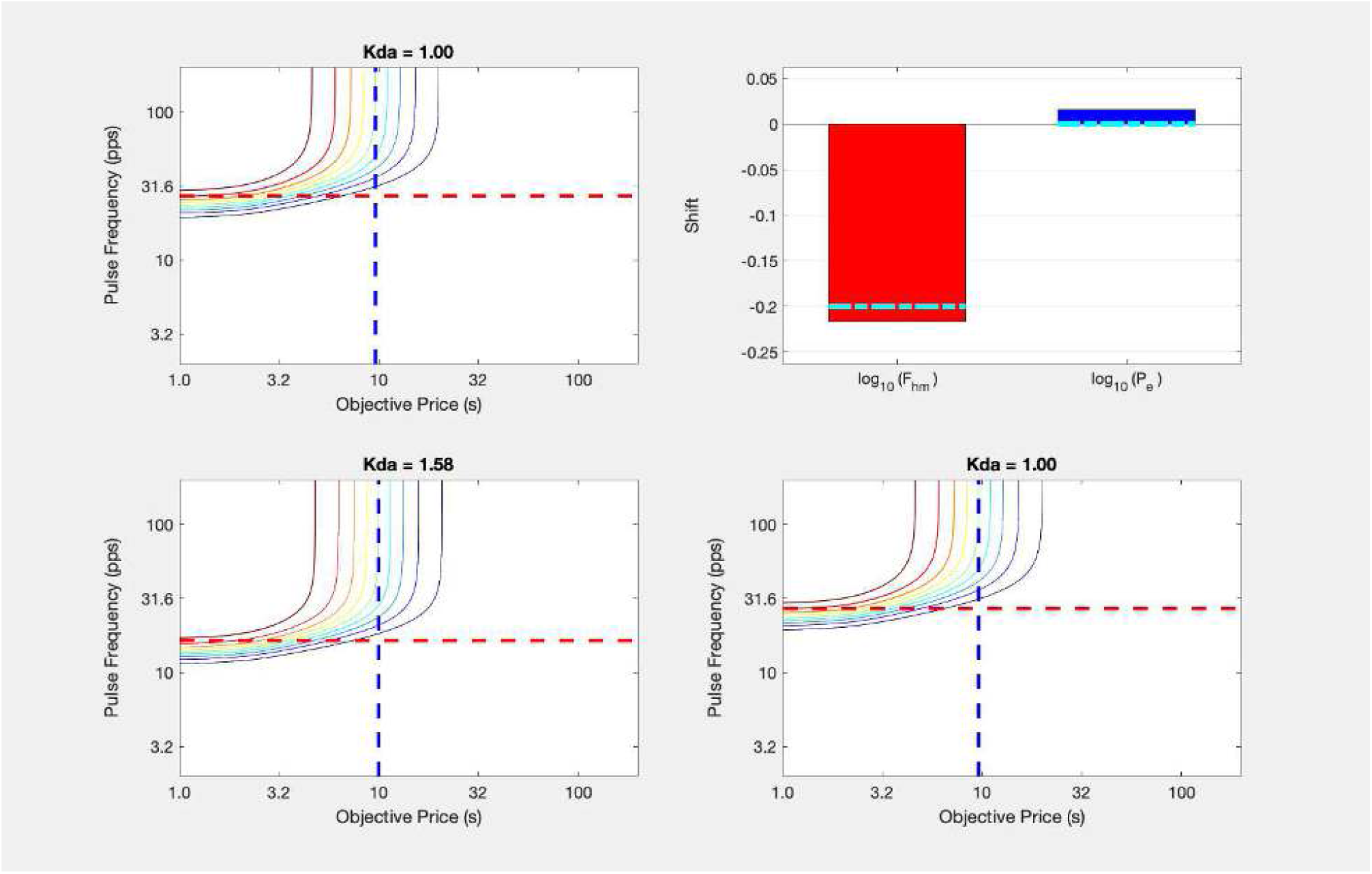

~~~
[fig_num, fig_tab] = add_fig(fig_num, fig_tab, “kDA1vskDA2_quad”, … “
  Effect of dopamine-transporter blockade on the reward mountain”);
~~~

`This is figure #38: (kDA1vskDA2_quad)`

In the above simulation, the drug-induced boost in the effectiveness of dopaminergic neurotransmission has rescaled the input to the reward-growth function. As a result, the mountain shifts leftwards along the pulse-frequency axis. This captures one of the observed effects of GBR-12909.

The small shift along the price axis is due to the fact that 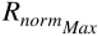 is closer to one in the drug condition than in the vehicle condition. This is due to the fact that ***F***_*hm*_ is lower in the drug condition and thus further from the pulse frequency at which frequency following begins to roll off. The corrected value of log_10_(***F***_*hm*_) is designated by the dot-dash cyan line on the red bar.

The difference in frequency-following fidelity in the drug and vehicle conditions also produces a small shift in log_10_(***P***_*e*_). This shift is removed by the correction (dot-dash cyan line on the blue bar).

The empricially observed rightward shifts along the pulse axis survive the correction for differential frequency-following fidelity. To capture these, a scalar increase must be produced at or beyond the output of the reward- growth function. In the example shown below, we do this by having the drug-induced increase in dopamine tone reduce the subjective effort cost, as proposed by Salomone and colleagues.

`Elapsed time is 10.708417 seconds.`

### Dopaminergic modulation of subjective effort cost

~~~
tic;
close all;
~~~

In

~~~
disp(string({strcat({‘Figure ‘},num2str(fig_tab.Number(fig_tab.Name==‘oICSS_logistic’)))}));
~~~

`Figure 36`

, blockade of the dopamine transporter changes both phasic and tonic dopamine signaling. Given that the lack of correlation between the magnitudes of the resulting shifts along the pulse-frequency and price axes, separate scalars are included for phasic and tonic signaling. The schema indicates three different ways in which increases in tonic signaling could shift the mountain along the price axis, as designated by the black dashed line. The one simulated here is a reduction in the rate of subjective exertion entailed in holding down the lever.

~~~
% Set Fhm to the values obtained from ratBechr29 in the vehicle and drug conditions Kda = [1,1.4458]; % Kda value for Bechr29
FdaHMkDA = [31.34679397, 20.77560124]; % values for Bechr29 FdaHMkDAstar =
[26.29253041, 18.18388501]; % values for Bechr29
PsubBend = 0.5;
PsubMin = 1.8197;
FdaBend = 20;
FdaRO = 50;
FPmax = 200; % This is the value that was employed in the correction of the location parameters
g = 3.160774009; % value for Bechr29; value common for Veh and Drg conditions
RnormMax = [0.893580092, 0.964209143]; % means of resampled values for Bechr29
% PobjE = PobjEfun(dotPhiObj, Kaa, Kec, Krg, dotRaa, pObj, PsubBend, PsubMin, RnormMax, KeffMod)
PobjE = [6.295449067, 10.04096421]; % values for Bechr29
PobjEstar = [7.375024269, 10.47998071]; % values for Bechr29
% We can’t simply perform the corrections of the location parameters on the means.
% The original corrections were performed on the 250 resampled log values.
% The statistics are based on these vectors.
% The log of the mean is not equal to the mean of the logs.
% Similarly, we cannot simply use FilterFun to correct the mean Fhm values and use these to estimate RnormMax
% logFfiringHM (i.e., logFhmStar) and logPobjEstar were estimated from each of the 250 resampled values of
% logFpulseHM and logPobjE. We encounter the same issue.
% The values used above are based on the 250 resampled log values, which were loaded directly from the
% saved workspace that was used in the fits.
~~~

~~~
a = 3;
Fopt = logspace(0,3,121)’; % column variable
Pobj = logspace(0,3,121); % row variable
Tda1 = TAfun(a, Fopt, FdaBend, FdaHMkDA(1), FdaRO, g, Pobj, PobjE(1), …
  PsubBend, PsubMin, RnormMax(1));
Tda2 = TAfun(a, Fopt, FdaBend, FdaHMkDA(2), FdaRO, g, Pobj, PobjE(2), …
  PsubBend, PsubMin, RnormMax(2));
xmin = 0;
xmax = 2.3;
ymin = 0.3;
ymax = 2.3;
title_str1 = strcat({‘Kda = ‘}, sprintf(‘%2.2f’, Kda(1)));
MTNkDAeffMod1 = plot_MTN(Fopt, Pobj, Tda1, ‘off’, ‘MTNoICSSkDAeffMod1’, title_str1, …
  graphs2files, FigDir, xmin, xmax, ymin, ymax);
title_str2 = strcat({‘Kda = ‘}, sprintf(‘%2.2f’, Kda(2)));
MTNkDAeffMod2 = plot_MTN(Fopt, Pobj, Tda2, ‘off’, ‘MTNoICSSkDAeffMod2’, title_str2, …
  graphs2files, FigDir, xmin, xmax, ymin, ymax);
MTNoICSSkDA1vskDA2 = dual_subplot(MTNkDAeffMod1, MTNkDAeffMod2, ‘MTNoICSSkDA1vskDA2’,… graphs2files,FigDir);
if show_graphics
  MTNoICSSkDA1vskDA2.Visible = ‘on’;
end
~~~

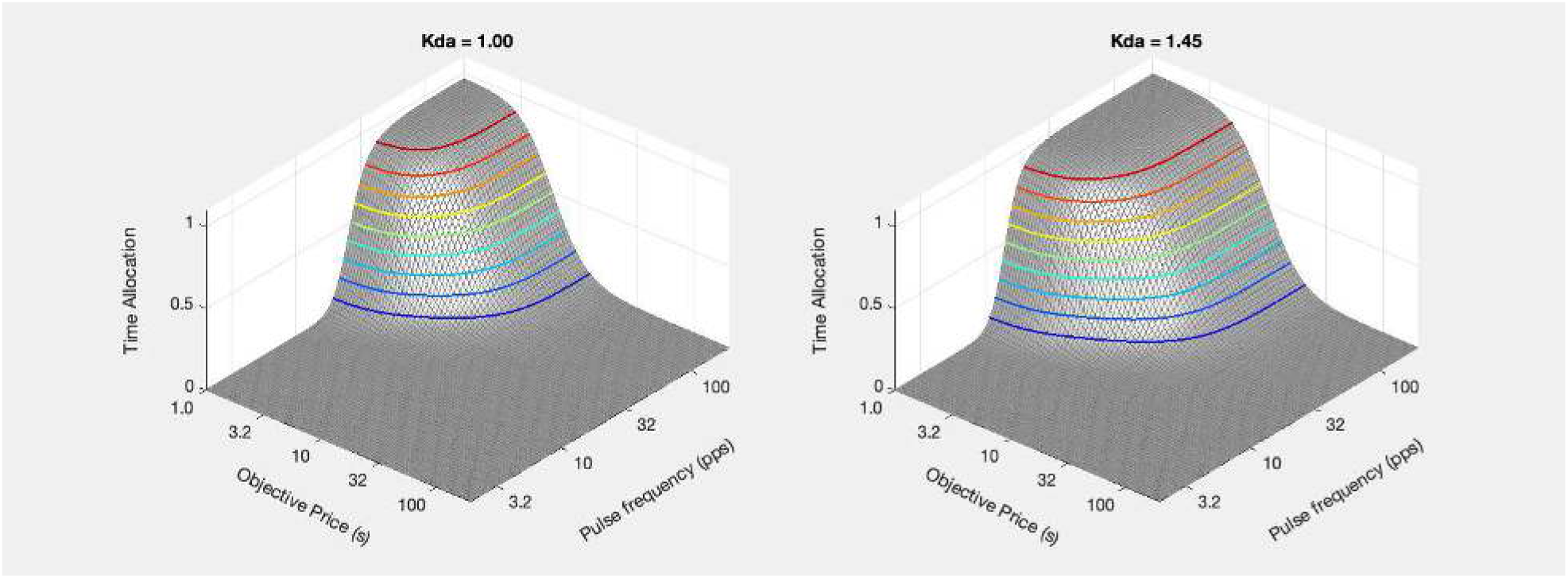

~~~
[fig_num, fig_tab] = add_fig(fig_num, fig_tab, “kDA1vskDAeffMod2_mtns”, …
  “Effect of dopamine-transporter blockade on the reward mountain”);
~~~

`This is figure #39: (kDA1vskDAeffMod2_mtns)`

~~~
contkDAeffMod1 = plot_contour(Fopt, Pobj, Tda1, PobjE(1), FdaHMkDA(1), ‘off’, ‘ContkDAeffMod1’, title_str1’, …
  strcat({‘Kda = ‘}, num2str(Kda(1))), graphs2files, FigDir,…
  xmin, xmax, ymin, ymax);
ContkDAeffMod2 = plot_contour(Fopt, Pobj, Tda2, PobjE(2), FdaHMkDA(2), ‘off’, ‘ContkDAeffMod2’, title_str2, …
  strcat({‘Kda = ‘}, num2str(Kda(2))), graphs2files, FigDir,…
  xmin, xmax, ymin, ymax);
ymin = −0.3581; % from the bargraph for Bechr29
ymax = 0.3652; % from the bargraph for Bechr29
bg_kDA1vskDA2effMod = plot_bgStar(FdaHMkDA(1), FdaHMkDA(2), FdaHMkDAstar(1), FdaHMkDAstar(2),…
PobjE(1), PobjE(2), PobjEstar(1), PobjEstar(2),…
 ‘off’, ‘kDA1vskDA2_bg’, graphs2files, FigDir,…
    ymin, ymax);
bg_root = ‘bg_kDA1vskDAeffMod2’;
quad_kDA1vskDA2effMod = quad_subplot(ContkDAeffMod1, ContkDAeffMod2, bg_kDA1vskDA2effMod,…
     ‘quad_kDA1vskDAeffMod2’, bg_root, graphs2files, FigDir);
if show_graphics
    quad_kDA1vskDA2effMod.Visible = ‘on’;
end
~~~

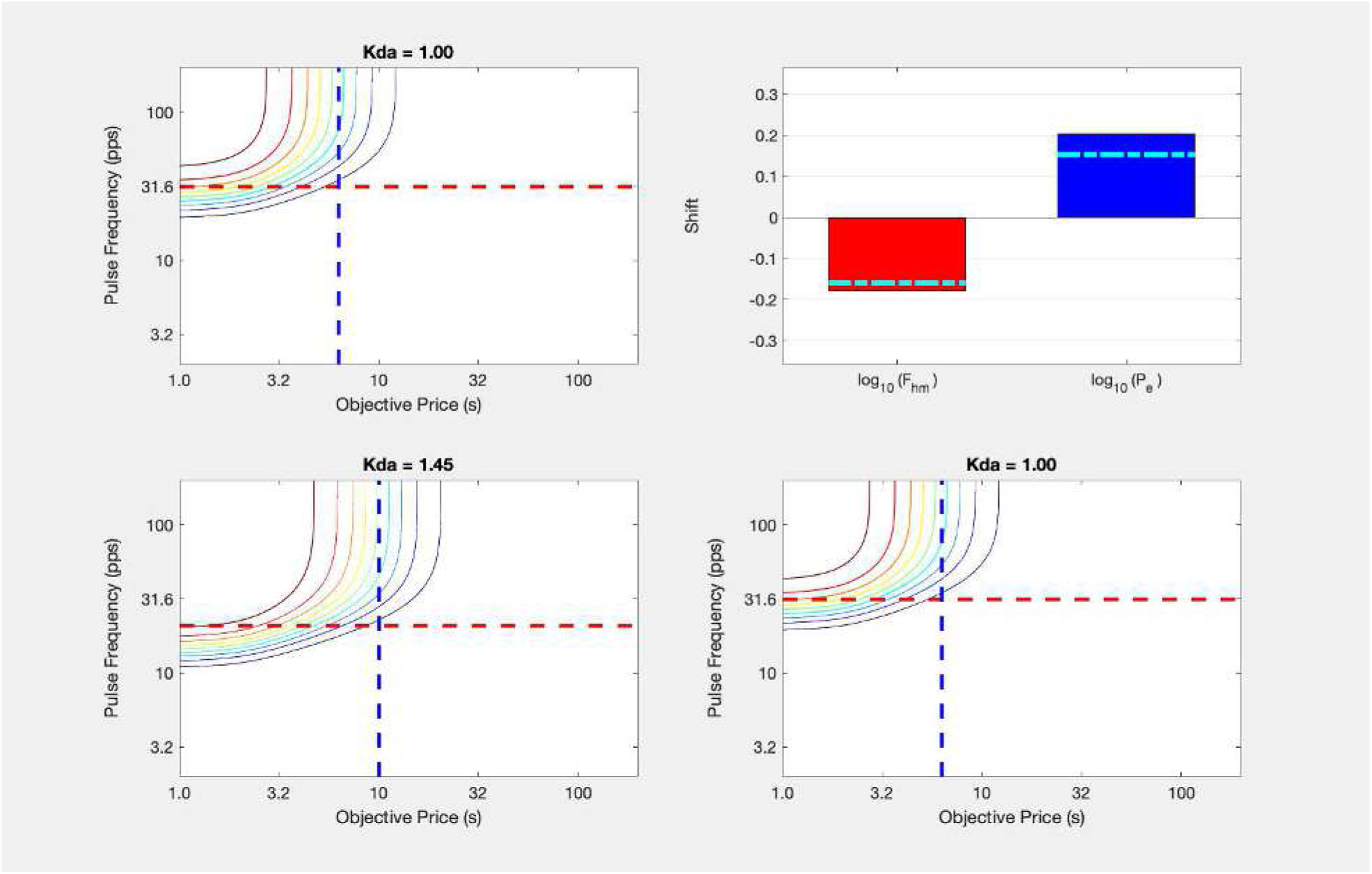

~~~
[fig_num, fig_tab] = add_fig(fig_num, fig_tab, “kDA1vskDA2effMod_quad”, …
   “Effect of dopamine-transporter blockade on the reward mountain”);
~~~

`This is figure #40: (kDA1vskDA2effMod_quad)`

The simulated results are now in qualitative accord with the empirical findings: the mountain shifts leftwards along the pulse-frequency axis and rightwards along the price axis in response to blockade of the dopamine transporter.

Here are the shifts observed in the mountains obtained from rat BeChR29 and the shifts simulated above using the ***K***_*da*_ and 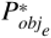 values derived from that rat’s data:

~~~
BeChR29_bg = make_fig_from_png(fullfile(ImpFigDir,’BeChR29_bg.png’), 30);
BeChR29bgVSsimBG = dual_subplot(BeChR29_bg, bg_kDA1vskDA2effMod, ‘BeChR29bgVSsimBG’,…
        graphs2files,FigDir);
% rescale the right panel to 60% width, 80% height & center in right panel
BeChR29bgVSsimBGrs = adjust_right_panel(BeChR29bgVSsimBG, 0.6, 0.825);
if show_graphics
    BeChR29bgVSsimBGrs.Visible = ‘on’;
end
~~~

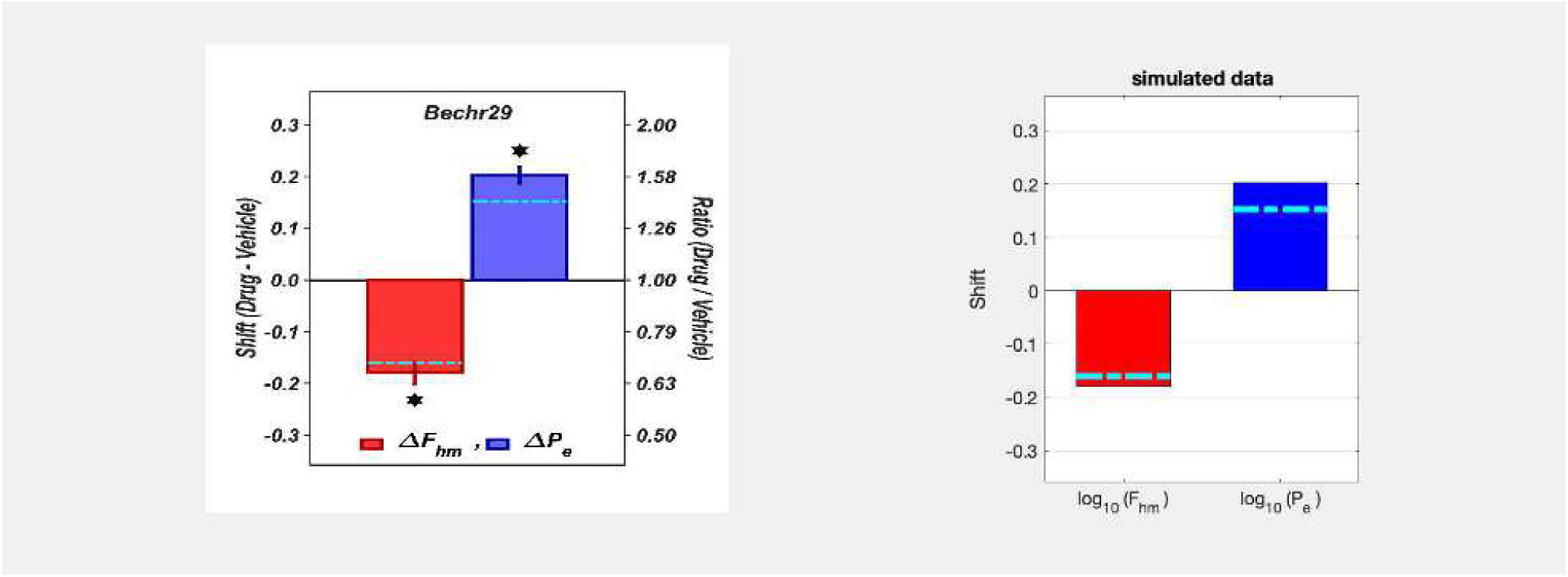

~~~
[fig_num, fig_tab] = add_fig(fig_num, fig_tab, “BeChR29bgVSsimBGrs”, …
    “Simulated and observed effects of dopamine-transporter blockade on the reward mountain”);
~~~

`This is figure #41: (BeChR29bgVSsimBGrs)`

The correspondence is forced by inputting values of 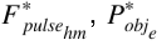, and *g* obtained from the fit of the reward-mountain model to the empirical data. All that this comparison does is to verify the functions for computing time allocation and graphing the results.

Full simulation would yield the same results if we adjusted 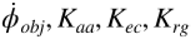, and 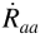 so as to generate the observed values.

`Elapsed time is 11.140447 seconds.`

## The series-circuit model of eICSS and oICSS

~~~
tic;
close all;
~~~

In the previous section, we demonstrate how the effects of dopamine-transporter blockade on oICSS can be explained within the context of the mountain model by a combination of changes in phasic and tonic dopamine signaling. In this section, we pose an additional challenge: can the explanation proposed for the oICSS data also account for the known effects of dopamine-transporter blockade on eICSS? Below, we show that the proposed explanation fails to meet this challenge, we discuss the implications for the “series-circuit” model of eICSS, and we develop a new model that can account for the effects of dopamine-transporter blockade on both oICSS and eICSS.

The series-circuit model treats oICSS and eICSS as behavioral manifestations of the effects produced by injecting signals at two different neural stages of the same pathway. On this view, eICSS arises from electrically induced activation of highly excitable, non-dopaminergic neurons that project directly or indirectly to midbrain dopamine neurons. In other words, the directly activated neurons that give rise to eICSS are in series with the dopamine neurons that render the electrical stimulation rewarding. The midbrain dopamine neurons are excited trans-synaptically in the case of eICSS and directly in the case of oICSS. The consequences of their activation are the same: the subject seeks to re-initiate the stimulation and will pay large effort and opportunity costs when the strength of the stimulation suffices to produce a large increment in the aggregate firing rate of the dopamine neurons.

Here, we show only the portion of the model that generates the reward-intensity signal.

~~~
if show_graphics
    show_imported_graphic(‘series-circuit_DA_double_logisitc_RG_v2.png’,15,ImpFigDir);
end
~~~

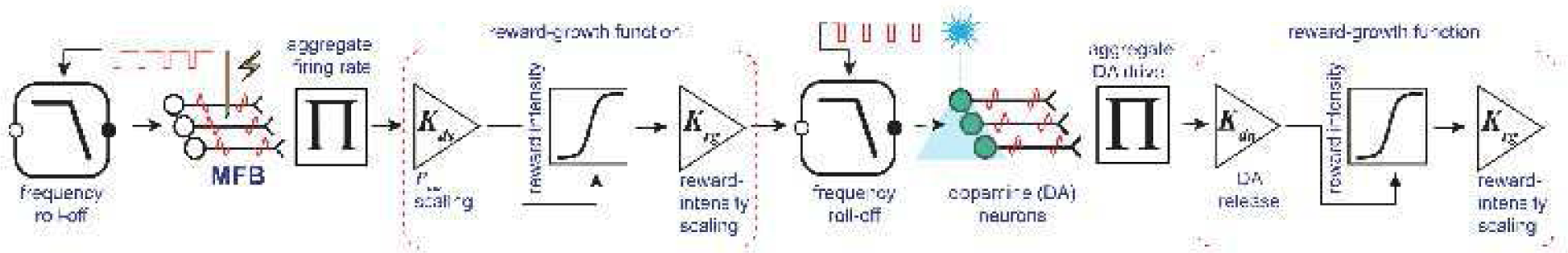

~~~
[fig_num, fig_tab] = add_fig(fig_num, fig_tab, “SeriesCircuit_eICSS_oICSS”, …
    “Series-circuit model of eICSS and oICSS”);
~~~

`This is figure #42: (SeriesCircuit_eICSS_oICSS)`

As the figure above shows, two different reward-growth functions are required, one upstream of the dopamine neurons and another downstream. The upstream reward-growth function is necessary in order to account for the data summarized in

~~~
disp(string({strcat({‘Figure ‘},num2str(fig_tab.Number(fig_tab.Name==‘PimGBRshifts’)))}));
~~~

`Figure 28`

That figure shows that modulation of dopaminergic neurotransmission alters eICSS performance by shifting the reward mountain along the price axis and not along the pulse-frequency axis. This implies that the drug-induced change in dopamine signaling acts at or beyond the output of the reward-growth function for eICSS. For this to occur in the series-circuit model, the reward-growth function for eICSS must lie upstream (to the left) of the dopamine neurons.

A second reward-growth function must lie downstream (to the right) of the dopamine neurons. This is required because GBR-12909 shifted the reward mountain for oICSS along the pulse-frequency axis, as shown in

~~~
disp(string({strcat({‘Figures ‘},num2str(fig_tab.Number(fig_tab.Name==‘BeChR29vehdrgMtn’)))}));
~~~

`Figures 30`

and

~~~
disp(string({strcat({‘‘},num2str(fig_tab.Number(fig_tab.Name==‘GBRshiftSummary’)))}));
~~~

`32`

We explain above (***The significance of orthogonal shifts***) that the mountain will move along the pulse-frequency axis only if the input to the reward-growth function has been rescaled. Such rescaling occurs when the magnitude of dopamine transients is boosted by GBR-12909 and the reward-growth function for oICSS is positioned downstream of the dopamine neurons.

### Series-circuit model: changes in the reward mountain for eICSS in response to dopamine-transporter blockade

To generate a reward mountain for eICSS from the series-circuit model, we proceed in several stages. As was done above (**The reward-growth function for oICSS**), the optical pulse frequency required to produce a half-maximal reward intensity is calculated directly from

~~~
disp(string({strcat({‘Equation ‘},num2str(eqn_tab.Number(eqn_tab.Name==‘fH’)))}));
~~~

`Equation 17`

We next need to compute the electrical pulse frequency (applied to a medial forebrain bundle electrode) that produces excitation in the dopamine neurons equivalent to that produced by a given train of optical pulses delivered to the midbrain dopamine neurons. According the series-circuit model, all inputs that produce the same peak output from the dopamine neurons will produce the same rewarding effect. This will be true regardless of whether the dopamine neurons are excited directly by optical activation or indirectly by trans-synaptic input from medial forebrain bundle neurons activated by electrical stimulation. According to the model, an observer placed downstream from the dopamine neurons and supplied only with information about the aggregate peak output of these neurons cannot know whether optical or electrical stimulation was responsible for a given phasic increase in dopamine release. It is the peak magnitude of this phasic increase in aggregate firing that determines the intensity of the rewarding effect.

(See “**The spike counter”** above.)

To obtain the electrical pulse frequency required to produce a reward of half-maximal intensity, we need to back-solve the equations describing the stages of the model that intervene between the electrode and the dopamine neurons. The back-solutions return the electrical pulse frequency that delivers an input to dopamine neurons equivalent to the optical pulse frequency that produces a half-maximal reward intensity.

The reward-growth functions (S-shaped curves in rectangular boxes) in the flow diagrams are normalized: Their output varies from zero to one and is then scaled by the variable in the triangle to their right. In the case of the upstream reward-growth function in

~~~
disp(string({strcat({‘Figure ‘},num2str(fig_tab.Number(fig_tab.Name==‘SeriesCircuit_eICSS_oICSS’)))}));
~~~

`Figure 42`

(the reward-growth function to the left of the dopamine neurons), that scaling variable is 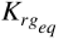. The value of this variable determines the maximum input that the electrode can deliver to the dopamine neurons, scaled in terms of the equivalent optical pulse frequency. In the initial simulation below, 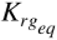 is set to 63 pulses per second, and given the parameters of the frequency-following function of the dopamine neurons 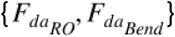, this will drive the dopamine neurons to a near-maximal level (∼43 spikes *s*^−1^). We demonstrate below that the qualitative effect of dopamine transporter blockade is little affected by the value of 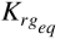.

The function that computes the required electrical pulse frequency in the series-circuit model, *FscHMbs*, comprises three back-solutions. (See ***Functions composing the reward-mountain model*** below.) First, we invert the scaling of the output of the upstream reward-growth function: we divide the optical pulse frequency required to produce a reward of half-maximal intensity by 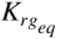. This gives us the output of the upstream normalized reward-growth function (a value between zero and one), the fraction of 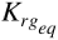 required to produce a half-maximal reward intensity at the output of the dopamine neurons (the FdaHMkDA value). We then backsolve the upstream reward-growth function (by means of the `LogistNormBsFun` function) to obtain the average firing rate of the medial forebrain bundle neurons required to produce the reward intensity in question, given the position parameter of the logistic reward-growth function and the value of its exponent. The position parameter of the logistic is calculated using

~~~
disp(string({strcat({‘Equation ‘},num2str(eqn_tab.Number(eqn_tab.Name==‘fH’)))}));
~~~

`Equation 17`

(the *FFmfbHM* function). Finally, we use the *FilterFunBS* function to obtain the electrical pulse frequency that produces this average firing rate.

The input to *FscHMbs* is the optical pulse frequency required to produce a reward of half-maximal intensity 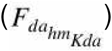 by direct activation of the dopamine neurons, and the output is the equivalent electrical pulse frequency 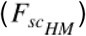.

~~~
[fun_num, fun_tab] = add_fun(fun_num, fun_tab, “FscHMbs”, …
     “FhmDA, FmfbBend, FhmMFB, FmfbRO, gMFB, KrgEq”,…
     “Back solution of the full reward-growth function for the series-circuit (sc) model”);
~~~

`This is function #39: (FscHMbs)`

~~~
[sym_num, sym_tab] = add_sym(sym_num, sym_tab, “FscHM”, “position parameter of RG function in the series-circuit model”);
~~~

We now compute the value of the *FscHMbs* function for the vehicle and drug conditions.

~~~
%% Calculate FscHM for the vehicle and drug conditions
% Calculate the position parameter of the upstream reward-growth function
C = 0.473; % median from Sonnenschein et al., 2003
D = 0.5; % typical train duration
FdaBend = 20;
logFmfbBend = 1.3222; % from Solomon et al., 2015
FmfbBend = 10^logFmfbBend
~~~

`FmfbBend = 20.9991`

~~~
FdaRO = 50;
gMFB = 1.58; % reduce the value due to the embedding within the DA RG function
logFmfbRO = 2.5587; % from Solomon et al., 2015
FmfbRO = 10^logFmfbRO
~~~

`FmfbRO = 361.9929`

~~~
NnMFB = 126;
RhoPiMFB = 5000; % RhoPi/N is within ∼20% of 50, which is roughly consistent with Sonnenchein et al., 2003
FPmax = 1000;
FmfbHM = FpulseHMfun(C, D, FmfbBend, FPmax, FmfbRO, NnMFB, RhoPiMFB)
~~~

`FmfbHM = 77.2222`

In the vehicle condition of the current experiment, the median 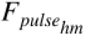 value for oICSS was 27.1. The train duration was 1 s, whereas it was 0.5 s in the corresponding eICSS study. The form and parameters of the temporal-integration function for oICSS of midbrain dopamine neurons are unknown. Faut de mieux, we will use the functional form and parameters obtained from eICSS to compute the Fhm value for a 0.5 s train that corresponds to the value obtained in the current oICSS study for 1 s trains.

In the series-circuit model, the output of the MFB neurons must pass through the midbrain dopamine neurons. If so, the temporal-integration characteristics obtained in eICSS experiments reflect those of both the directly stimulated and dopamine stages of the circuit. According to this model, integration in the dopamine stage cannot be faster than estimated in the eICSS study.

It follows from

~~~
disp(string({strcat({‘Equation ‘},num2str(eqn_tab.Number(eqn_tab.Name==‘fH’)))}));
~~~

`Equation 17`

that the Fhm value for a 0.5 s train that corresponds to an Fhm value for a 1 s train is given by

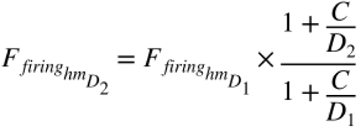

~~~
[eqn_num, eqn_tab] = add_eqn(eqn_num, eqn_tab, …
   “FfiringD1D2”, “calculates equivalently effective pulse frequencies at two different train durations”);
~~~

`This is equation #33: (FfiringD1D2)`

Given the frequency-following function, the median 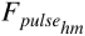 value for the vehicle condition (27.1) corresponds to a firing frequency of 23.1.

~~~
FpulseHMmedVeh = 27.094;
FfiringHMmedVeh = FilterFun(FpulseHMmedVeh, FdaBend, FdaRO)
~~~

`FfiringHMmedVeh = 23.1475`

Given these values

~~~
C = 0.473;
D1 = 1;
D2 = 0.5;
FfiringHmD1 = FfiringHMmedVeh
~~~

`FfiringHmD1 = 23.1475`

~~~
FfiringHmD2 = FfiringHmD1 .* (1+(C./D2)) ./ (1+(C./D1))
~~~

`FfiringHmD2 = 30.5805`

~~~
FPmax = 1000;
FpulseHmD2 = FilterFunBS(FdaBend, FdaRO, FfiringHmD2,FPmax)
~~~

`FpulseHmD2 = 37.6179`

The MFB will have to deliver excitation equivalent to 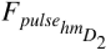 to deliver a half-maximal reward intensity from the dopamine neurons at a train duration of 0.5 s. We will determine the values of 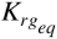 and 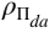 so as to obtain the above 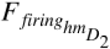 value in the vehicle condition.

We first set the number of activated dopamine neurons arbitrarily to 100.

~~~
NnDA = 100;
~~~

Then, we use

~~~
disp(string({strcat({‘Equation ‘},num2str(eqn_tab.Number(eqn_tab.Name==‘FfiringD1D2’)))}));
~~~

`Equation 33`

to find the required value of 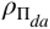, keeping in mind that 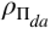 is the aggregate firing rate required to produce a half-maximal reward intensity with a train of infinite duration. Setting D1 to infinity makes the denominator of

~~~
disp(string({strcat({‘Equation ‘},num2str(eqn_tab.Number(eqn_tab.Name==‘FfiringD1D2’)))}));
~~~

`Equation 33`

equal to one, so the equation is simplified to:

~~~
RhoPiDA = NnDA * (FfiringHmD2/(1+(C/D2)))
~~~

`RhoPiDA = 1.5715e+03`

We are now positioned to compute the MFB pulse frequency that will drive the dopamine neurons to produce a reward of half-maximal intensity.

~~~
Kda = [1,10^0.15]; % Effect of dopamine-transporter blockade on the frequency of firing required
% to produce a reward of half-maximal intensity. Median Kda in current oICSS study: 10^0.1446
FPmax = 1000; % This pulse frequency well above FdaRO and can thus serve to estimate their firing rate
FdaHMkDA = FpulseHMfun(C, D2, FdaBend, FPmax, FdaRO, NnDA, RhoPiDA, Kda) % result is a two-element vector
~~~

~~~
FdaHMkDA = 1×2
   37.6179 25.1418
~~~

Now, we work backwards through the MFB input to find the electrical pulse frequency that will drive the dopamine neurons to produce a reward of half-maximal intensity.

~~~
KrgEq = 63;
FFmax = FilterFun(FPmax, FmfbBend, FmfbRO);
if FFmax > LogistNormBsFun(gMFB,FmfbHM,max(FdaHMkDA)/KrgEq)
    for j = 1:length(FdaHMkDA)
        FscHM(j) = FscHMbs(FdaHMkDA(j), FmfbBend, FmfbHM, FmfbRO, FPmax, gMFB, KrgEq) % result is a two-element vector
    end
else
    display(“FscHM could not be calculated because KrgEq is too low.”);
    display(“Set KrgEq such that FFmax > FilterFun(FPmax, Fbend, Fro).”);
    return
end
~~~

~~~
FscHM = 99.0574
FscHM = 1×2
   99.0574 59.5983
~~~

~~~
% Calculate PobjE for the vehicle and drug conditions
dotPhiObj = 1;
Kaa = 1;
Kec = 1;
KeffMod = Kda; % We tie the subjective effort cost to the effect of the drug
Kda
~~~

~~~
Kda = 1×2
    1.0000 1.4125
~~~

~~~
% Median shift in Pe in the current oICSS study: 10^0.1525
Krg = 1;
dotRaa = 0.1;
pObj = 1;
PsubBend = 0.5;
PsubMin = 1.82;
Felec = logspace(0,3,121)’; % column variable
~~~

To solve the time-allocation equation for the series-circuit model, we must first compute the drive on the dopamine neurons that is produced by each value of the electrical pulse frequency (*F*_*elec*_). We express this drive in terms of the optical pulse frequency that produces equivalent firing in the dopamine neurons, 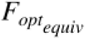. The *FoptEquivFun* function first translates the electrical pulse frequency into the induced rate of firing in the medial forebrain bundle neurons, then translates this firing rate into a normalized reward intensity, and then converts the normalized reward intensity into the equivalent optical pulse frequency. Once this has been done, time allocation can be computed using the same equation and parameter values that were used to generate the reward mountain for optical stimulation. The *FoptEquivFun* function is the inverse of the *FscHMbs* function. (See ***Functions composing the reward-mountain model*** below.)

~~~
[fun_num, fun_tab] = add_fun(fun_num, fun_tab, “FoptEquivFun”, …
    “Felec, FmfbBend, FmfbHM, FmfbRO, gMFB, KrgE”,…
    “Function to compute the optical pulse frequency that produces the same DA output as Felec”);
~~~

`This is function #40: (FoptEquivFun)`

~~~
[sym_num, sym_tab] = add_sym(sym_num, sym_tab, “FoptEquiv”, “optical pulse frequency that applies the same drive on the dopamine neurons as Felec”);
FoptEquiv = FoptEquivFun(Felec, FmfbBend, FmfbHM, FmfbRO, gMFB, KrgEq);
FPmaxEq = max(FoptEquiv) % This is the highest equivalent optical pulse frequency that the DA neurons will “see”
~~~

`FPmaxEq = 57.9538`

~~~
gDA = 5; % Use a value consistent with the oICSS study; median in vehicle condition was 4.61
RnormMax = fRbsrNorm(FPmaxEq, FdaBend, FdaHMkDA, FdaRO, gDA)
~~~

~~~
RnormMax = 1×2
    0.6146 0.9228
~~~

~~~
% RnormMax is a 2-element vector
PobjE = PobjEfun(dotPhiObj, Kaa, Kec, Krg, dotRaa, pObj, PsubBend, PsubMin, RnormMax, KeffMod)
~~~

~~~
PobjE = 1×2
    6.1461 13.0354
~~~

~~~
% PobjE is a two-element vector
Pobj = logspace(0,3,121); % row variable
% Compute time allocation for the vehicle and drug conditions
a = 2; % median in vehicle condition was 1.84
Tsc1 = TAfun(a, FoptEquiv, FdaBend, FdaHMkDA(1), FdaRO, gDA, Pobj, …
     PobjE(1), PsubBend, PsubMin, RnormMax(1));
Tsc2 = TAfun(a, FoptEquiv, FdaBend, FdaHMkDA(2), FdaRO, gDA, Pobj, …
     PobjE(2), PsubBend, PsubMin, RnormMax(2));
title_str1 = strcat({‘Kda = ‘}, sprintf(‘%2.2f’, Kda(1)));
MTNkDA1sc = plot_MTN(Felec, Pobj, Tsc1, ‘off’, ‘MTNkDA1sc’, title_str1, …
    graphs2files, FigDir);
title_str2 = strcat({‘Kda = ‘}, sprintf(‘%2.2f’, Kda(2)));
MTNkDA2sc = plot_MTN(Felec, Pobj, Tsc2, ‘off’, ‘MTNkDA2sc’, title_str2, …
    graphs2files, FigDir);
MTNkDA1vskDA2sc = dual_subplot(MTNkDA1sc, MTNkDA2sc, ‘MTNkDA1vskDA2sc’,…
    graphs2files,FigDir);
if show_graphics
    MTNkDA1vskDA2sc.Visible = ‘on’;
end
~~~

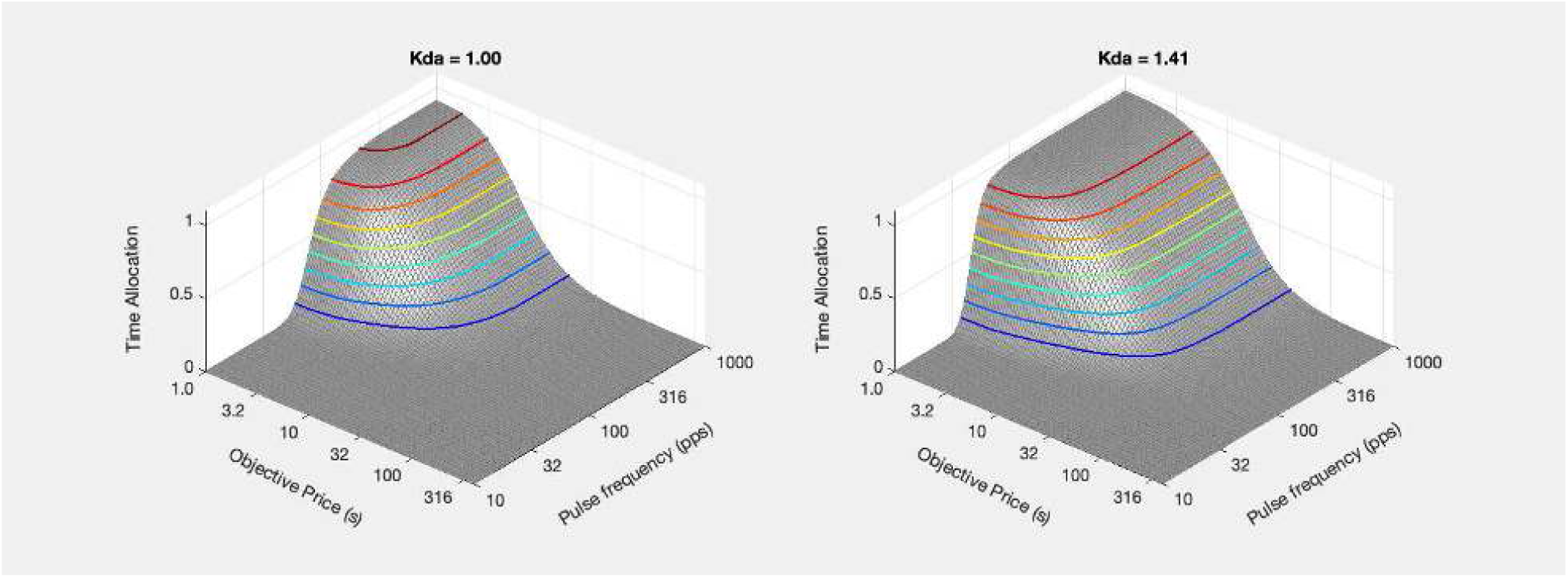

~~~
[fig_num, fig_tab] = add_fig(fig_num, fig_tab, “kDA1vskDA2_sc_mtns”, …
      “Effect of dopamine-transporter blockade on the reward mountain”);
~~~

`This is figure #43: (kDA1vskDA2_sc_mtns)`

~~~
FscHMstar = FilterFun(FdaHMkDA,FdaBend,FdaRO);
% Computed on the basis of frequency following in the dopamine neurons
PsubE = PsubFun(PobjE,PsubBend,PsubMin);
PsubEstar = PsubE ./ RnormMax;
PobjEstar = PsubBsFun(PsubEstar,PsubBend,PsubMin);
~~~

~~~
ContkDA1sc = plot_contour(Felec, Pobj, Tsc1, PobjE(1), FscHM(1), ‘off’, ‘ContkDA1sc’, title_str1, …
    strcat({‘Kda = ‘}, num2str(Kda(1))), graphs2files, FigDir);
ContkDA2sc = plot_contour(Felec, Pobj, Tsc2, PobjE(2), FscHM(2), ‘off’, ‘ContkDA2sc’, title_str2, …
    strcat({‘Kda = ‘}, num2str(Kda(2))), graphs2files, FigDir);
bg_kDA1vskDA2sc = plot_bgStar(FscHM(1), FscHM(2), FscHMstar(1), FscHMstar(2),…
    PobjE(1), PobjE(2), PobjEstar(1), PobjEstar(2),…
    ‘off’, ‘kDA1vskDA2sc_bg’, graphs2files, FigDir);
bg_root = ‘bg_kDA1vskDA2sc’;
quad_kDA1vskDA2sc = quad_subplot(ContkDA1sc, ContkDA2sc, bg_kDA1vskDA2sc, ‘quad_kDA1vskDA2sc’, bg_root, …
    graphs2files, FigDir);
if show_graphics
    quad_kDA1vskDA2sc.Visible = ‘on’;
end
~~~

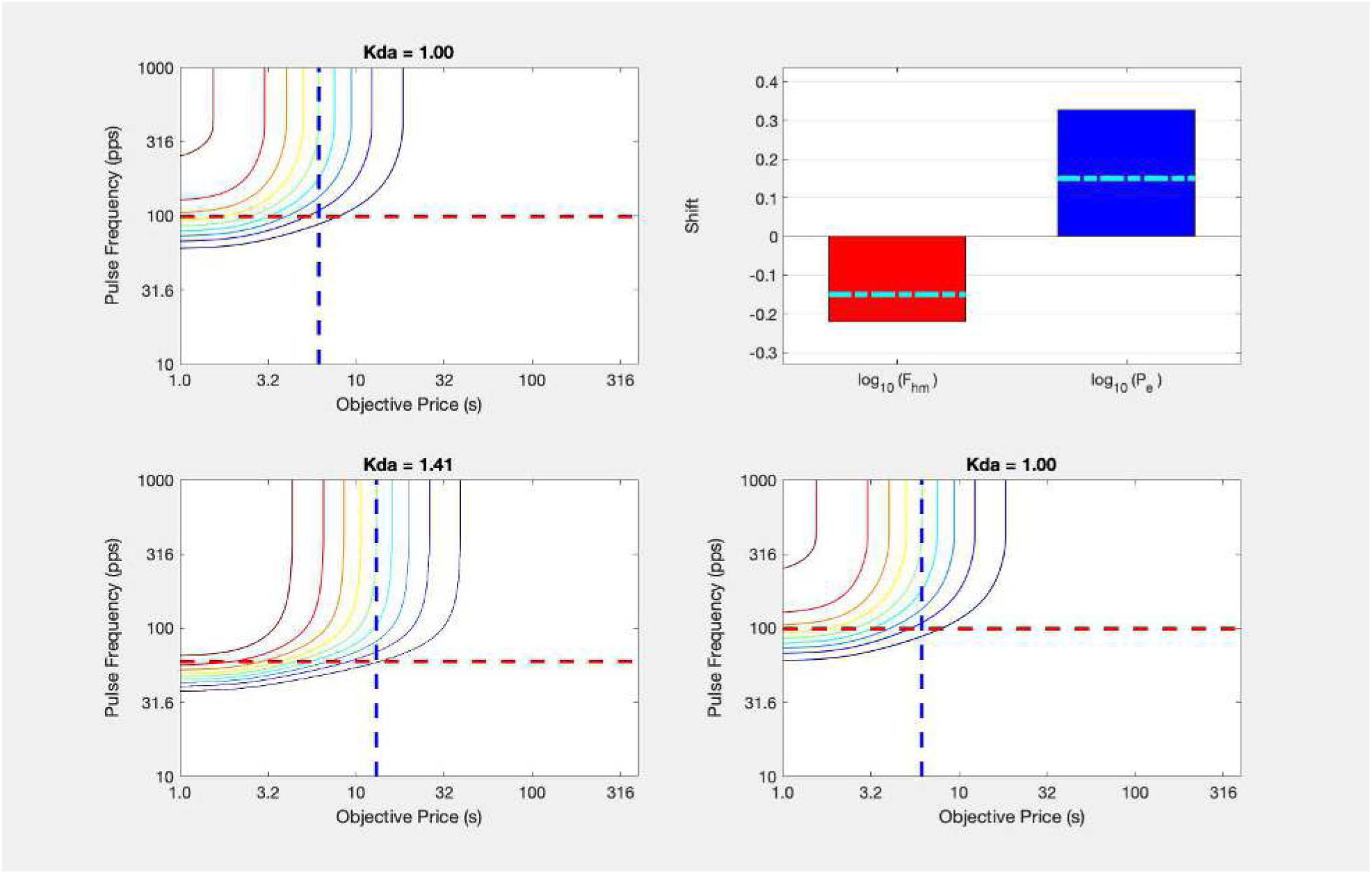

~~~
[fig_num, fig_tab] = add_fig(fig_num, fig_tab, “kDA1vskDA2_sc_quad”, …
     “Effect of dopamine-transporter blockade on the reward mountain”);
~~~

`This is figure #44: (kDA1vskDA2_sc_quad)`

~~~
toc
~~~

`Elapsed time is 9.650534 seconds.`

~~~
tic
clear -regexp ^Cont ^dual ^MTN ^quad;
~~~

The following two figures compare the shifts in the position of the reward mountain predicted by the series-circuit model and observed empirically when eICSS is challenged with GBR-12909, a dopamine-transporter blocker:

~~~
GBR_eICSS_summary = make_fig_from_png(fullfile(ImpFigDir,’GBR_eICSS_summary.png’), 100);
GBReICSSVSsimBG = dual_subplot(GBR_eICSS_summary, bg_kDA1vskDA2sc, ‘GBReICSSVSsimBG’,…
    graphs2files,FigDir);
% rescale the right panel to 60% width, 80% height & center in right panel
GBReICSSVSsimBGrs = adjust_right_panel(GBReICSSVSsimBG, 0.75, 1);
if show_graphics
    GBReICSSVSsimBGrs.Visible = ‘on’;
end
~~~

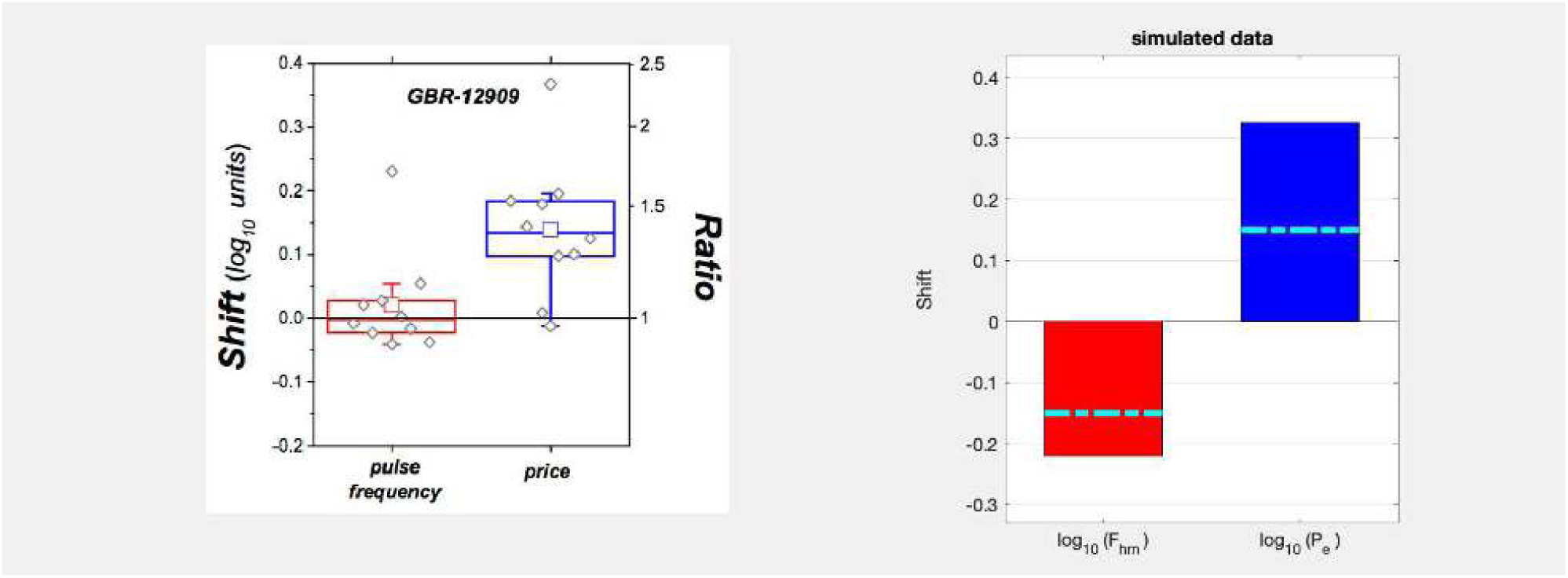

The predicted shift along the price axis is consistent qualitatively with the empirical results of the experiment in which eICSS was challenged with GBR-12909: the mountain shifts rightwards along the price axis. However, the predicted shift along the pulse-frequency axis is inconsistent with the empirical results. Whereas the series-circuit model predicts a leftward shift along the pulse-frequency axis, the mountain was not displaced systematically along the pulse-frequency axis in the empirical study. This discrepancy reveals a serious flaw in the series-circuit model.

The reason for the failure of the series-circuit model becomes clear upon inspection of

~~~
disp(string({strcat({‘Figure ‘},num2str(fig_tab.Number(fig_tab.Name==‘SeriesCircuit_eICSS_oICSS’)))}));
~~~

`Figure 42`

The change in dopaminergic neurotransmission produced by GBR-12909 rescales the input to the downstream reward-growth function (to the right of the dopamine neurons), changing its position parameter. In contrast to what is observed empirically in the eICSS studies, this causes the simulated mountain to shift along the pulse-frequency axis, regardless of whether the dopamine neurons are activated directly by optical stimulation or indirectly by synaptic input from the medial forebrain bundle neurons stimulated by an electrode.

An observer positioned at the input to the downstream reward-growth function cannot know whether optical or electrical stimulation was responsible for the phasic dopamine signal that constitutes the input to this function. Thus, the series-circuit model cannot readily generate differential predictions in response to optical and electrical inputs.

~~~
toc
~~~

`Elapsed time is 1.282094 seconds.`

### The qualitative predictions don’t depend meaningfully on the value of the parameter than scales the medial forebrain bundle drive on the dopamine neurons

~~~
tic;
clear -regexp ^GBR;
if show_graphics
    close all;
end
~~~

The next set of graphs shows that the predictions don’t change qualitatively when the maximum medial forebrain bundle drive on the dopamine neurons is reduced: The mountain continues to shift along both the price and pulse-frequency axes

~~~
KrgEqLo = 41; % Maximum MFB drive is reduced
% n.b. KrgEqLo must be sufficiently exceed FdaHMkDA so as to keep FscHMlo below the maximum firing rate
% Otherwise the back-solution function will generate an error message and return.
% Re-calculate the location-parameter values along the pulse-frequency axis for
% the series-circuit model as a whole. FFmax = FilterFun(FPmax, FmfbBend, FmfbRO);
if FFmax > LogistNormBsFun(gMFB,FmfbHM,max(FdaHMkDA)/KrgEqLo)
    for j=1:length(FdaHMkDA)
        FscHMlo(j) = FscHMbs(FdaHMkDA(j), FmfbBend, FmfbHM, FmfbRO, FPmax, gMFB, KrgEqLo) % result is a two-element vector
    end
else
    display(“FscHM could not be calculated because KrgEqLo is too low.”);
    display(“Set KrgEqLo such that FFmax > FilterFun(FPmax, Fbend, Fro).”);
    return
end
~~~

~~~
FscHMlo = 380.5649
FscHMlo = 1×2
  380.5649 103.3750
~~~

~~~
% Recalculate the drive on the DA neurons
FoptEquivLo = FoptEquivFun(Felec, FmfbBend, FmfbHM, FmfbRO, gMFB, KrgEqLo);
FPmaxEq = max(FoptEquivLo) % This is the highest equivalent optical pulse frequency that the DA neurons will “see”
~~~

`FPmaxEq = 37.7159`

~~~
RnormMax = fRbsrNorm(FPmaxEq, FdaBend, FdaHMkDA, FdaRO, gDA)
~~~

~~~
RnormMax = 1×2
    0.2640 0.7290
~~~

~~~
PobjE = PobjEfun(dotPhiObj, Kaa, Kec, Krg, dotRaa, pObj, PsubBend, PsubMin, RnormMax, KeffMod)
~~~

~~~
PobjE = 1×2
    2.5324 10.2974
~~~

~~~
% Compute time allocation for the vehicle and drug conditions
Tsc3 = TAfun(a, FoptEquivLo, FdaBend, FdaHMkDA(1), FdaRO, gDA, Pobj, …
    PobjE(1), PsubBend, PsubMin, RnormMax(1));
Tsc4 = TAfun(a, FoptEquivLo, FdaBend, FdaHMkDA(2), FdaRO, gDA, Pobj, …
    PobjE(2), PsubBend, PsubMin, RnormMax(2));
title_str1 = strcat({‘Kda = ‘}, sprintf(‘%2.2f’, Kda(1)));
MTNkDA1scKrgEqLo = plot_MTN(Felec, Pobj, Tsc3, ‘off’, ‘MTNkDA1scKrgEqLo’, title_str1, …
    graphs2files, FigDir);
title_str2 = strcat({‘Kda = ‘}, sprintf(‘%2.2f’, Kda(2)));
MTNkDA2scKrgEqLo = plot_MTN(Felec, Pobj, Tsc4, ‘off’, ‘MTNkDA2scKrgEqLo’, title_str2, …
    graphs2files, FigDir);
MTNkDA1vskDA2scKrgEqLo = dual_subplot(MTNkDA1scKrgEqLo, MTNkDA2scKrgEqLo, ‘MTNkDA1vskDA2scKrgEqLo’,…
    graphs2files,FigDir);
if show_graphics
    MTNkDA1vskDA2scKrgEqLo.Visible = ‘on’;
end
~~~

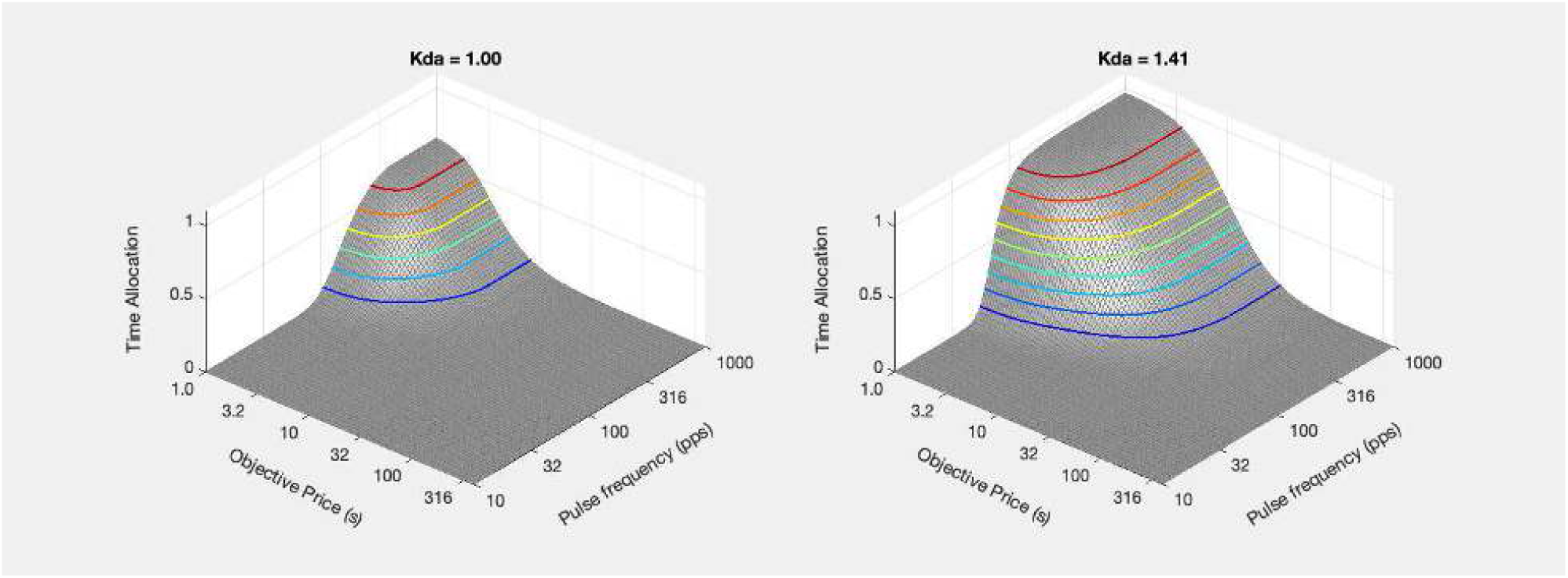

~~~
[fig_num, fig_tab] = add_fig(fig_num, fig_tab, “kDA1vskDA2_scKrgEqLo_mtns”, …
    “Effect of dopamine-transporter blockade on the reward mountain”);
~~~

`This is figure #45: (kDA1vskDA2_scKrgEqLo_mtns)`

~~~
FscHMstar = FilterFun(FdaHMkDA,FdaBend,FdaRO);
% Computed on the basis of frequency following in the dopamine neurons
PsubE = PsubFun(PobjE,PsubBend,PsubMin);
PsubEstar = PsubE ./ RnormMax;
PobjEstar = PsubBsFun(PsubEstar,PsubBend,PsubMin);
ContkDA1scKrgEqLo = plot_contour(Felec, Pobj, Tsc3, PobjE(1), FscHMlo(1), ‘off’, ‘ContkDA1scKrgEqLo’, title_str1, …
    strcat({‘Kda = ‘}, num2str(Kda(1))), graphs2files, FigDir);
ContkDA2scKrgEqLo = plot_contour(Felec, Pobj, Tsc4, PobjE(2), FscHMlo(2), ‘off’, ‘ContkDA2scKrgEqLo’, title_str2, …
    strcat({‘Kda = ‘}, num2str(Kda(2))), graphs2files, FigDir);
bg_kDA1vskDA2scKrgEqLo = plot_bgStar(FscHM(1), FscHM(2), FscHMstar(1), FscHMstar(2),…
    PobjE(1), PobjE(2), PobjEstar(1), PobjEstar(2),…
    ‘off’, ‘kDA1vskDA2scKrgEqLo_bg’, graphs2files, FigDir);
bg_root = ‘bg_kDA1vskDA2scKrgEqLo’;
quad_kDA1vskDA2scKrgEqLo = quad_subplot(ContkDA1scKrgEqLo, ContkDA2scKrgEqLo, bg_kDA1vskDA2scKrgEqLo, …
    ‘quad_kDA1vskDA2scKrgEqLo’, bg_root, graphs2files, FigDir);
if show_graphics
    quad_kDA1vskDA2scKrgEqLo.Visible = ‘on’;
end
~~~

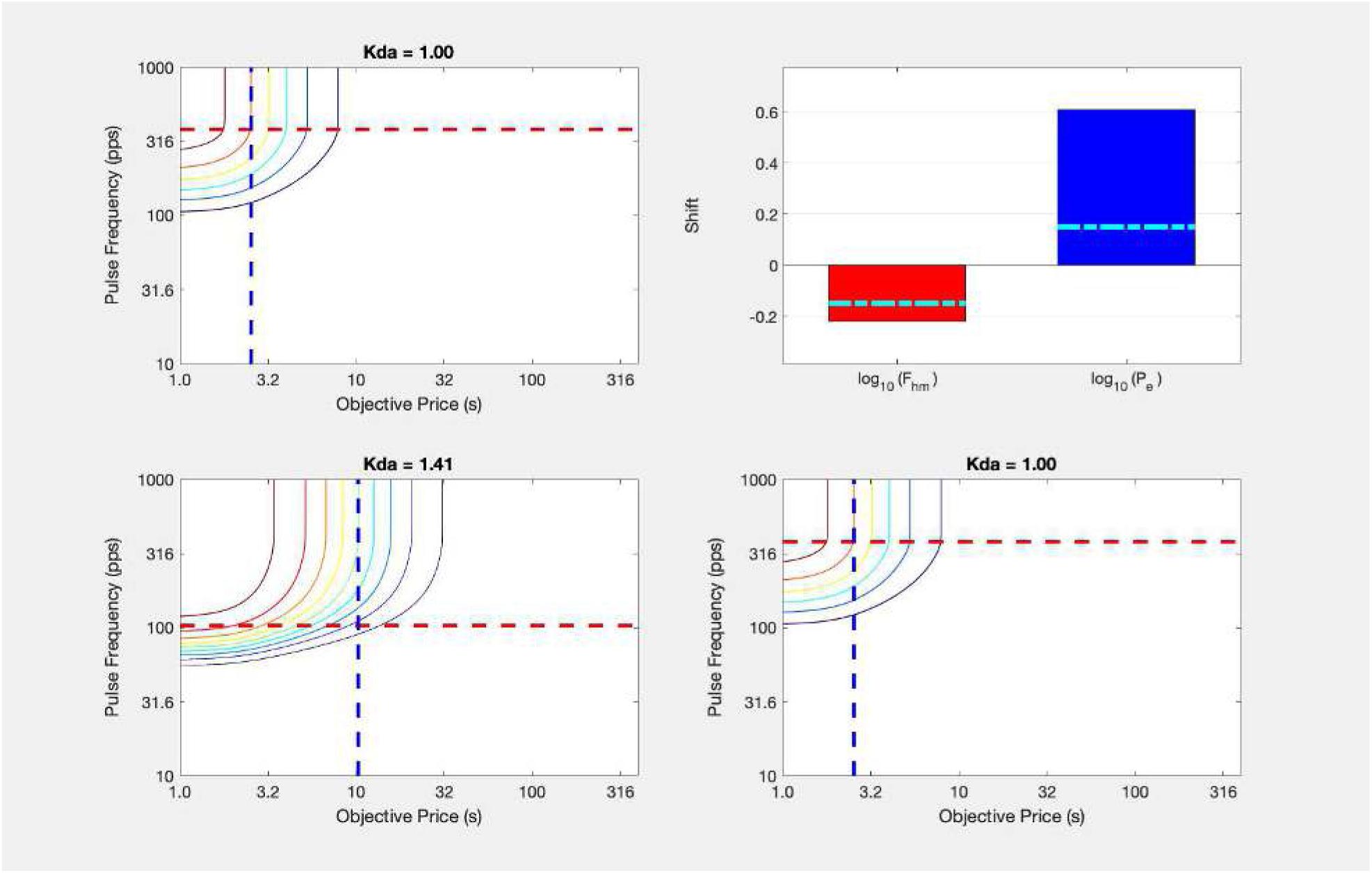

~~~
[fig_num, fig_tab] = add_fig(fig_num, fig_tab, “kDA1vskDA2_scKrgEqLo_quad”, …
    “Effect of dopamine-transporter blockade on the reward mountain”);
~~~

`This is figure #46: (kDA1vskDA2_scKrgEqLo_quad)`

~~~
toc
~~~

`Elapsed time is 9.818036 seconds.`

~~~
tic;
clear -regexp ^bg ^Cont ^dual ^MTN ^quad;
close all;
~~~

Next, we increase the medial forebrain bundle drive so that some of it is wasted. The firing rate of the dopamine neurons now asymptotes at sub-maximal levels of medial forebrain bundle drive. Again, the predictions of the series-circuit model do not change qualitatively: The mountain continues to shift along both the price and pulse-frequency axes.

~~~
KrgEqHi = 126; % Maximum MFB drive exceeds the DA frequency-following range
% Re-calculate the location-parameter values along the pulse-frequency axis for
% the series-circuit model as a whole.
FFmax = FilterFun(FPmax, FmfbBend, FmfbRO);
if FFmax > LogistNormBsFun(gMFB,FmfbHM,max(FdaHMkDA)/KrgEqHi)
    for j=1:length(FdaHMkDA)
        FscHMhi(j) = FscHMbs(FdaHMkDA(j), FmfbBend, FmfbHM, FmfbRO, FPmax, gMFB, KrgEqHi) % result is a two-element vector
    end
else
    display(“FscHM could not be calculated because KrgEqHi is too low.”);
    display(“Set KrgEqHi such that FFmax > FilterFun(FPmax, Fbend, Fro).”);
    return
end
~~~

~~~
FscHMhi = 44.9730
FscHMhi = 1×2
   44.9730 32.0551
~~~

~~~
% Recalculate the drive on the DA neurons
FoptEquivHi = FoptEquivFun(Felec, FmfbBend, FmfbHM, FmfbRO, gMFB, KrgEqHi);
FPmaxEq = max(FoptEquivHi) % This is the highest equivalent optical pulse frequency that the DA neurons will “see”
~~~

`FPmaxEq = 115.9075`

~~~
RnormMax = fRbsrNorm(FPmaxEq, FdaBend, FdaHMkDA, FdaRO, gDA)
~~~

~~~
RnormMax = 1×2
    0.8186 0.9713
~~~

~~~
PobjE = PobjEfun(dotPhiObj, Kaa, Kec, Krg, dotRaa, pObj, PsubBend, PsubMin, RnormMax, KeffMod)
~~~

~~~
PobjE = 1×2
    8.1862 13.7200
~~~

~~~
% Compute time allocation for the vehicle and drug conditions
Tsc5 = TAfun(a, FoptEquivHi, FdaBend, FdaHMkDA(1), FdaRO, gDA, Pobj, …
    PobjE(1), PsubBend, PsubMin, RnormMax(1));
Tsc6 = TAfun(a, FoptEquivHi, FdaBend, FdaHMkDA(2), FdaRO, gDA, Pobj, …
    PobjE(2), PsubBend, PsubMin, RnormMax(2));
title_str1 = strcat({‘Kda = ‘}, sprintf(‘%2.2f’, Kda(1)));
MTNkDA1scKrgEqHi = plot_MTN(Felec, Pobj, Tsc5, ‘off’, ‘MTNkDA1scKrgEqHi’, title_str1, …
     graphs2files, FigDir);
title_str2 = strcat({‘Kda = ‘}, sprintf(‘%2.2f’, Kda(2)));
MTNkDA2scKrgEqHi = plot_MTN(Felec, Pobj, Tsc6, ‘off’, ‘MTNkDA2scKrgEqHi’, title_str2, …
     graphs2files, FigDir);
MTNkDA1vskDA2scKrgEqHi = dual_subplot(MTNkDA1scKrgEqHi, MTNkDA2scKrgEqHi, ‘MTNkDA1vskDA2scKrgEqHi’,…
     graphs2files,FigDir);
if show_graphics
     MTNkDA1vskDA2scKrgEqHi.Visible = ‘on’;
end
~~~

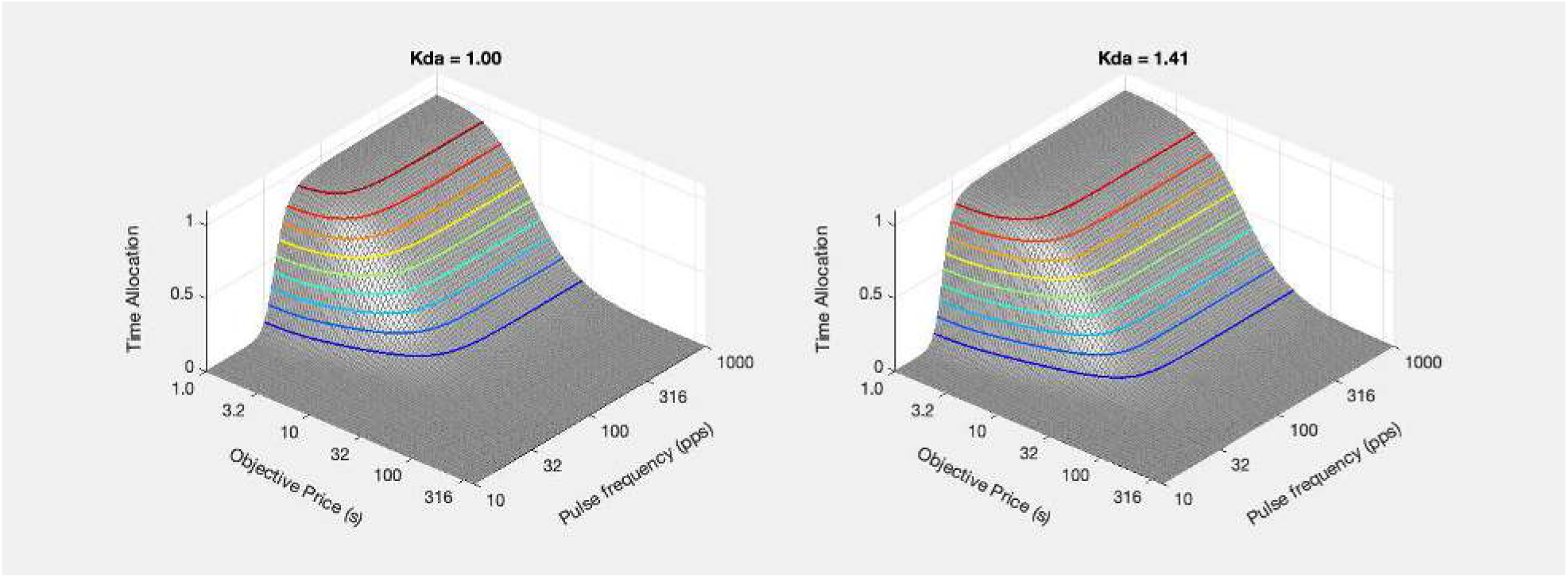

~~~
FscHMstar = FilterFun(FdaHMkDA,FdaBend,FdaRO);
% Computed on the basis of frequency following in the dopamine neurons
PsubE = PsubFun(PobjE,PsubBend,PsubMin);
PsubEstar = PsubE ./ RnormMax;
PobjEstar = PsubBsFun(PsubEstar,PsubBend,PsubMin);
ContkDA1scKrgEqHi = plot_contour(Felec, Pobj, Tsc5, PobjE(1), FscHMhi(1), ‘off’, ‘ContkDA1scKrgEqHi’, title_str1, …
    strcat({‘Kda = ‘}, num2str(Kda(1))), graphs2files, FigDir);
ContkDA2scKrgEqHi = plot_contour(Felec, Pobj, Tsc6, PobjE(2), FscHMhi(2), ‘off’, ‘ContkDA2scKrgEqHi’, title_str2, …
    strcat({‘Kda = ‘}, num2str(Kda(2))), graphs2files, FigDir);
bg_kDA1vskDA2scKrgEqHi = plot_bgStar(FscHM(1), FscHM(2), FscHMstar(1), FscHMstar(2),…
    PobjE(1), PobjE(2), PobjEstar(1), PobjEstar(2),…
    ‘off’, ‘kDA1vskDA2scKrgEqHi_bg’, graphs2files, FigDir);
bg_root = ‘bg_kDA1vskDA2scKrgEqHi’;
quad_kDA1vskDA2scKrgEqHi = quad_subplot(ContkDA1scKrgEqHi, ContkDA2scKrgEqHi, bg_kDA1vskDA2scKrgEqHi, …
    ‘quad_kDA1vskDA2scKrgEqHi’, bg_root, graphs2files, FigDir);
if show_graphics
    quad_kDA1vskDA2scKrgEqHi.Visible = ‘on’;
end
~~~

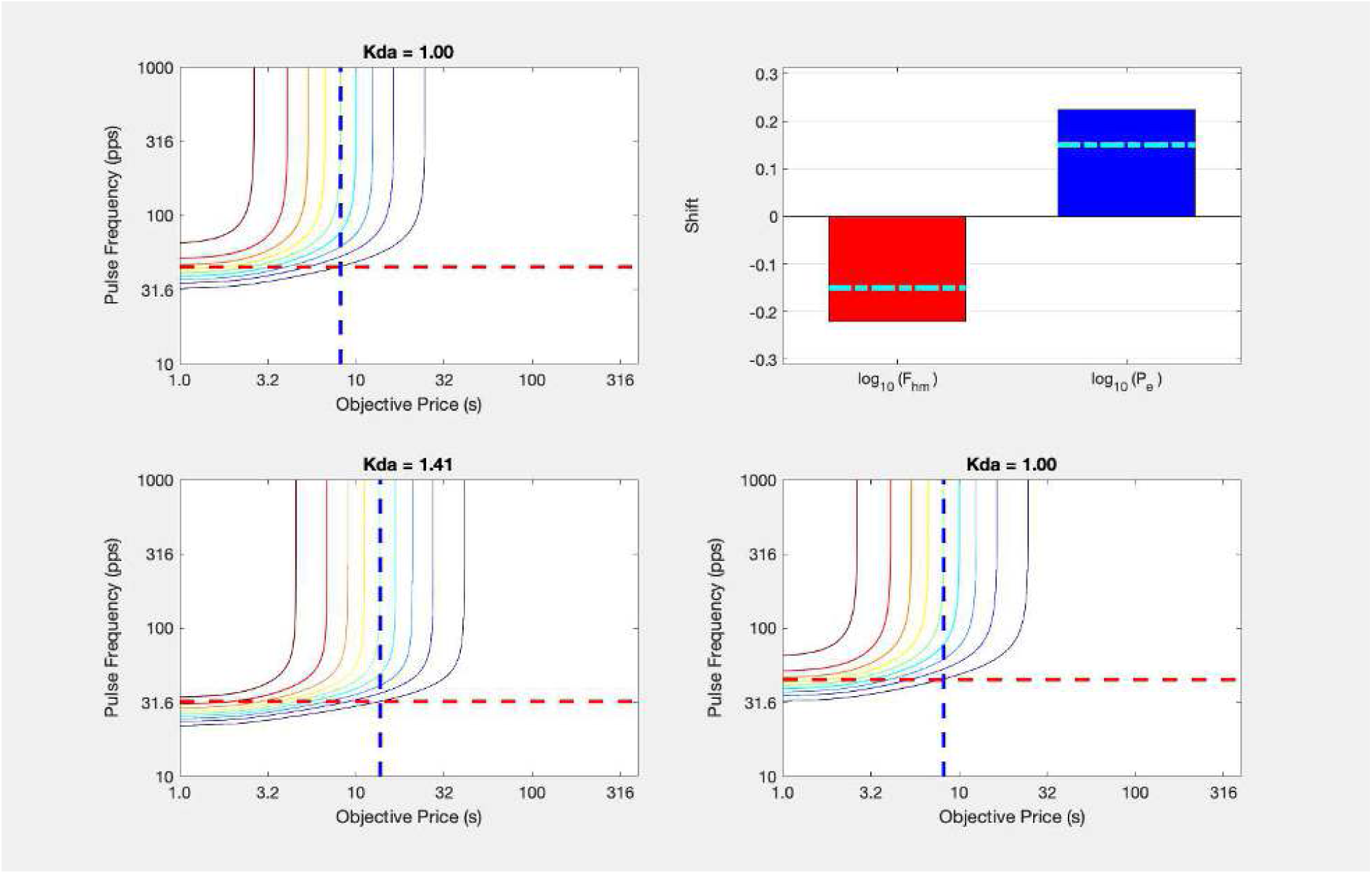

~~~
[fig_num, fig_tab] = add_fig(fig_num, fig_tab, “kDA1vskDA2_scKrgEqHi_quad”, …
    “Effect of dopamine-transporter blockade on the reward mountain”);
~~~

`This is figure #47: (kDA1vskDA2_scKrgEqHi_quad)`

This section demontrates that the inconsistency between the simulated and observed results is seen over a broad range of values of the parameter that scales the MFB drive on the dopamine neurons.

The problem with the series-circuit model is a fundamental one. It predicts shifts of the mountain along the pulse-frequency axis in response to perturbation of dopaminergic neurotransmission. In contrast, systematic, consistent shifts along the pulse-frequency axis are not seen in eICSS studies under the influence of the dopamine transporter blocker, GBR-12909 (Hernandez et al. 2012); the dopamine, norepinephrine, and serotonin blocker, cocaine (Hernandez et al., 2010); the D2, D3 and 5HT7 receptor blocker, pimozide (Trujillo-Pisanty et al., 2014); or the cannabinoid CB-1 blocker, AM-251 (Trujillo-Pisanty et al., 2011). (AM-251 inhibits dopamine release and attenuates the stimulation-induced increase in dopamine tone (Trujillo-Pisanty et al., 2011)). Thus, the series-circuit model fails to account for the eICSS data.

`Elapsed time is 8.968485 seconds.`

## The convergence model

~~~
tic;
close all;
~~~

The failure of the series-circuit model to account readily for the differential movement of the reward mountain in the eICSS and oICSS studies motivates the search for an alternative. We investigate a new model here. The challenge is to account for the stability of the reward mountain along the pulse-frequency axis under dopaminergic challenge in the eICSS data, the observed displacement along the pulse-frequency axis in the oICSS data, and the observed displacement along the price axis in both datasets.

~~~
if show_graphics
    show_imported_graphic(‘eICSS_oICSS_logistic+3Pe_noMasks_v1.png’,30,ImpFigDir)
end
~~~

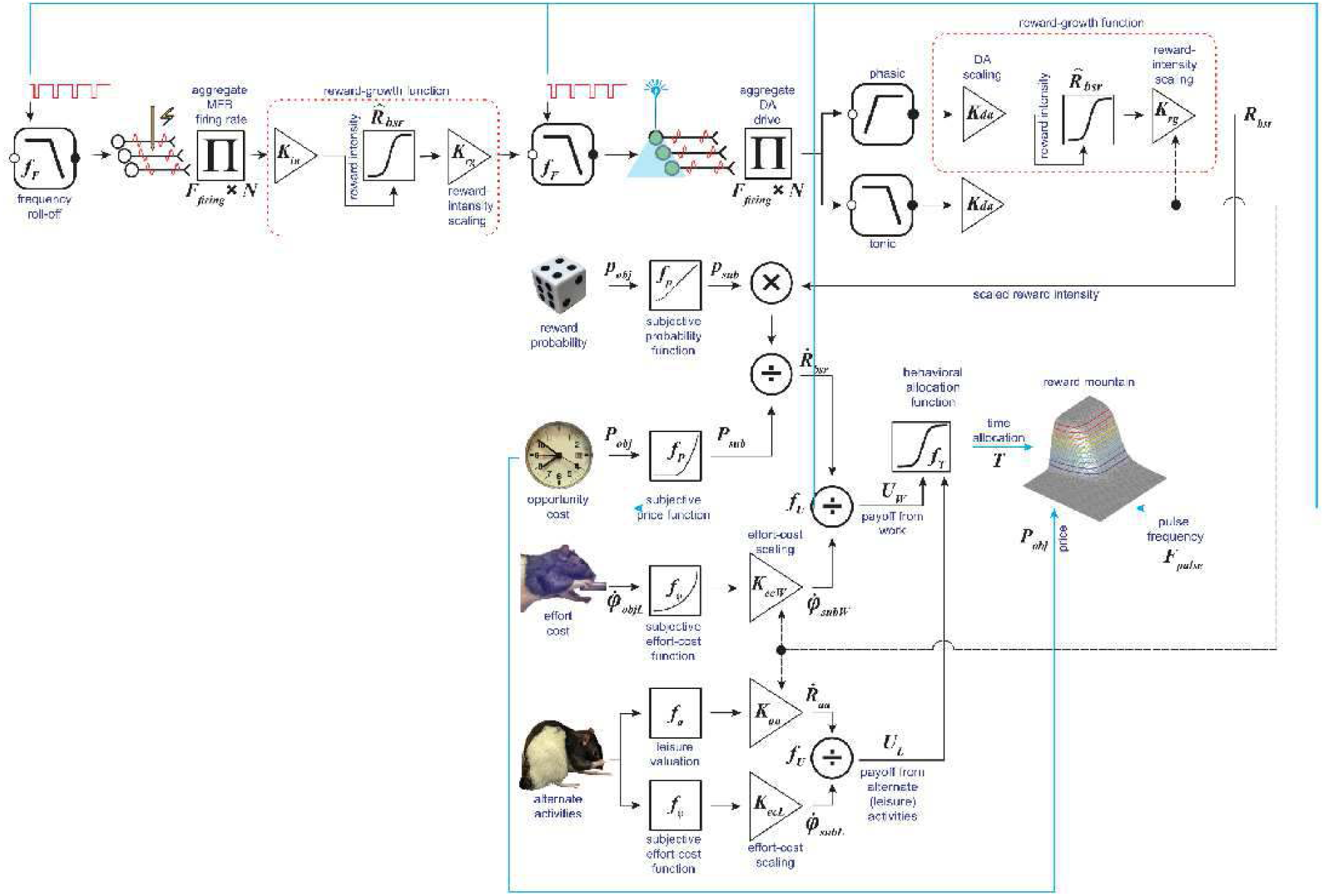

~~~
[fig_num, fig_tab] = add_fig(fig_num, fig_tab, “Convergence_model”, …
    “A model implementing converging pathways subserving oICSS and eICSS”);
~~~

`This is figure #48: (Convergence_model)`

In this model, the circuitry underlying eICSS of the medial forebrain bundle and oICSS of midbrain dopamine neurons includes parallel stages that are linked at two levels. The parallel limbs subserve oICSS and eICSS between the directly stimulated neurons and the scaled output of the reward-growth functions. The upstream link between the two limbs relays to midbrain dopamine neurons input from neurons activated by electrical stimulation of the medial forebrain bundle. The downsteam link combines the outputs of the two limbs so as to produce the signal representing the benefit of the experimenter-controlled reward.

### The input from the MFB to the midbrain dopamine neurons

It has long been known that electrical stimulation of the medial forebrain bundle provides trans-synaptic input to midbrain dopamine neurons (Maeda & Mogenson, 1980) and drives phasic dopamine release from their ventral striatal terminals (Gratton, Hoffer & Gerhardt, 1988; Yavich & Tiihonen, 2000a, 2000b; Wightman & Robinson, 2002; Yavitch & Tanila, 2007). Recently, Cossette, Conover & Shizgal (2016) demonstrated differences between the frequency-following characteristics of the MFB input to midbrain dopamine neurons that project to the medial shell of the nucleus accumbens and the frequency-following characteristics of the neurons subserving eICSS of the MFB. Whereas the neurons subserving the rewarding effect maintain high-fidelity frequency following up to ∼360 pulses *s*^−1^(Solomon et al., 2015), the input to the nucleus-accumbens projecting dopamine neurons fails to follow pulse frequencies greater than 130 pulses *s*^−1^. To accommodate this difference, a second low-pass filter is inserted below the link between the medial forebrain bundle fibers and the spike counter to the left of the dopamine neurons in

~~~
disp(string({strcat({‘Figure ‘},num2str(fig_tab.Number(fig_tab.Name==“Convergence_model”)))}));
~~~

`Figure 48`

At pulse frequencies below 130 pulses *s*^−1^, the amplitude of dopamine transients recorded in the nucleus accumbens in response to electrical stimulation of the medial forebrain bundle grows as function of both current and pulse frequency (Cossette, Conover & Shizgal, 2016). Increases in current can compensate for decreases in pulse frequency so as to hold constant the amplitude of the transient (and vice-versa). This suggests that with train duration held constant, the amplitude of the transient depends on the aggregate rate of firing in the medial forebrain bundle fibers that drive the activation of the dopamine neurons. That is the reason for the second spike counter.

As in the case of the series-circuit model, the trans-synaptic drive on the dopamine neurons is represented in terms of the optical pulse frequency that provides equivalent dopaminergic activation.

### Summation between the outputs of the two parallel limbs

The convergence model retains the idea that there is a final common path for neural signals that encode the predicted benefits of reward procurement. The two parallel circuit limbs, one subserving eICSS and the other oICSS, converge on this final common path. A simple way to implement this convergence is to add the outputs of the two limbs. This proposal faces a seemingly daunting challenge: electrical stimulation of the MFB activates midbrain dopamine neurons. If so, one would expect such stimulation to drive signaling in both of the hypothesized converging pathways. Wouldn’t this produce at least some displacement of the reward mountain along the pulse-frequency axis in response to dopamine-transporter blockade? We show below that this is not necessarily the case. Indeed, the simulations show that given reasonable assumptions and values drawn from the current data, a convergence model can replicate the eICSS findings.

The reward-intensity signal at the output of the eICSS limb of the circuit is computed as above, by applying

~~~
disp(string({strcat({‘Function ‘},num2str(fun_tab.Number(fun_tab.Name==‘fRbsrFull’)))}));
~~~

`Function 34`

The reward-intensity signal at the output of the oICSS limb of the circuit is computed in the same manner as when the input consists of optical stimulation pulses. However, additional steps are required to compute the output of this limb during eICSS, when electrically excited medial forebrain bundle neurons provide the input to the dopamine neurons. The electrical pulse frequency is translated in the input to the dopamine neurons (expressed in terms of the equivalent optical pulse frequency) by the function, *FmfbDAdriveFun*.

~~~
% [fun_num, fun_tab] = add_fun(fun_num, fun_tab, …
%     “FmfbDAdriveFun”, “FbendMFBda, Felec, FroMFBda, Kmfb, NnMFBs”,…
%     “Function that translates MFB drive on the DA neurons into an equivalent optical pulse frequency”);
[fun_num, fun_tab] = add_fun(fun_num, fun_tab, …
    “FFmfbDAbend”, “Felec, FFmfbDAmax, FoptEquivVec, FmfbDAro”,…
    “Function that translates MFB drive on the DA neurons into an equivalent optical pulse frequency”);
~~~

`This is function #41: (FFmfbDAbend)`

We now simulate the output of the model in response to electrical stimulation of the medial forebrain bundle.

~~~
% Reward intensity produced by the upper limb
C = 0.473; % median from Sonnenschein et al., 2003
Delec = 0.5;
logFelecBend = 1.3222; % from Solomon et al., 2015
FelecBend = 10^logFelecBend;
logFelecRO = 2.5587; % from Solomon et al., 2015
FelecRO = 10^logFelecRO;
NnMFB = 126;
RhoPi = 5000;
FPmax = 1000;
FhmUpper = FpulseHMfun(C, Delec, FelecBend, FPmax, FelecRO, NnMFB, RhoPi)
~~~

`FhmUpper = 77.2222`

~~~
numF = 121;
numParamVals = 6;
Felec = logspace(0,3,numF);
FelecMat = repmat(Felec,numParamVals,1);
gElec = 5;
KrgUpper = 1;
Rupper = fRbsr(Felec, FelecBend, FhmUpper, FelecRO, gElec, KrgUpper);
~~~

The position-parameter value for the upper limb (77 pulses *s*^−1^) is typical for the eICSS studies entailing measurement of the reward mountain.

~~~
disp(string({strcat({‘Figure ‘},num2str(fig_tab.Number(fig_tab.Name==‘RGfunsElecFhm’)))}));
~~~

Figure 3

~~~
TitleStrSemi = ‘Upper-limb reward intensity’;
pnam = “Fhm”;
fnam = “Fhm”;
% The data to be plotted must be in columns. Thus, FelecMat and RelecMat are transposed.
RG_MFBda_upper_semilog = plot_RG(Felec’,Rupper’,pnam,FhmUpper,fnam,TitleStrSemi,’lin’, ‘upper’);
axh = findall(RG_MFBda_upper_semilog,’Type’,’Axes’);
axh.XLabel.String = “Electrical pulse frequency”;
axh.YLabel.String = “Upper-limb reward intensity”;
if show_graphics
    RG_MFBda_upper_semilog.Visible = ‘on’;
end
~~~

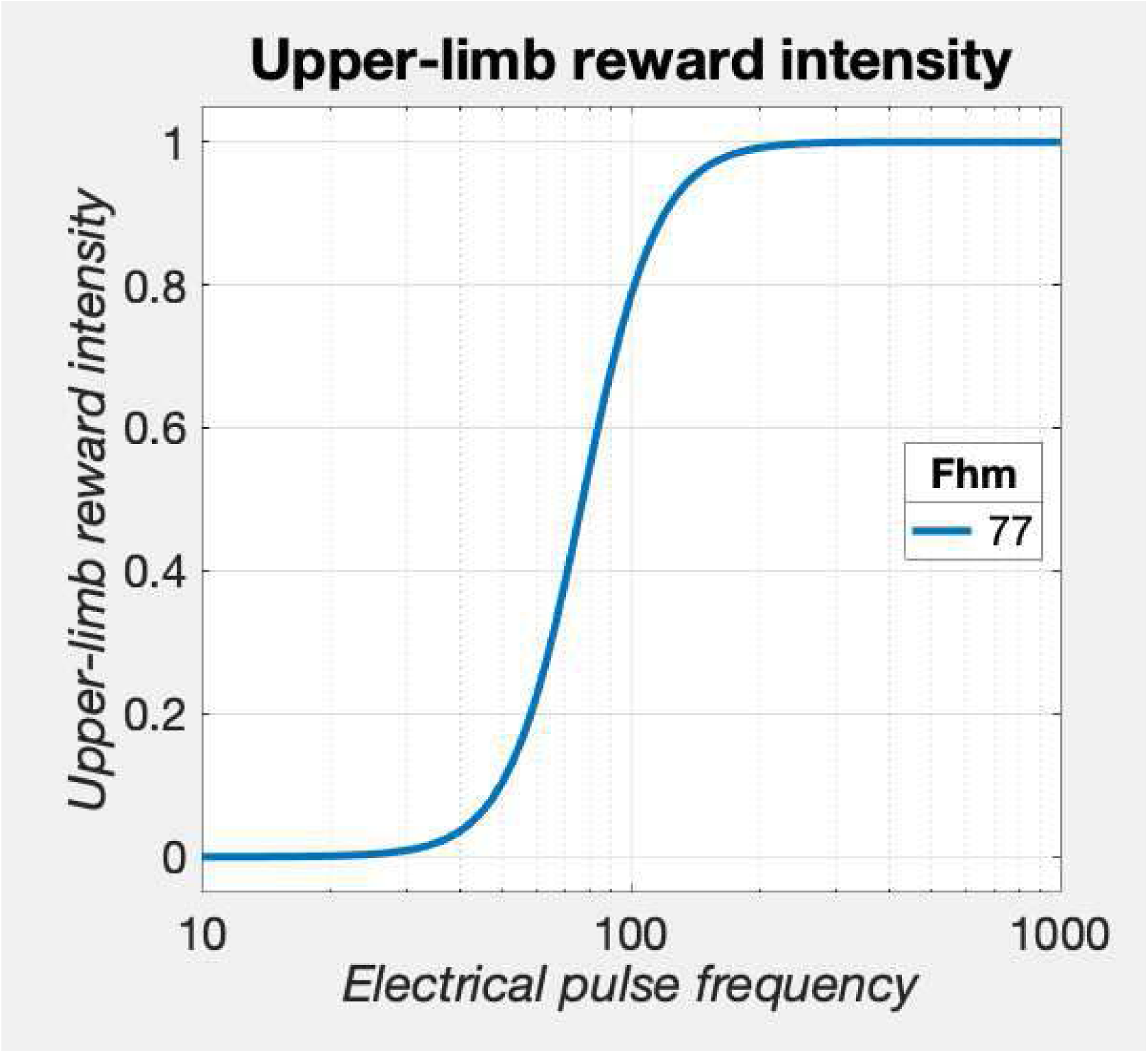

~~~
[fig_num, fig_tab] = add_fig(fig_num, fig_tab, “RG_upper_semilog”, …
“Growth of upper-limb reward intensity”);
~~~

`This is figure #49: (RG_upper_semilog)`

Reward-intensity growth in the lower limb depends on the strength of the MFB drive,

~~~
% Reward intensity produced by the lower limb
% First compute the drive on the DA neurons in terms of the equivalent optical pulse frequency
FmfbDAbend = 40; % Make this gradual to match data in Cossette et al., 2016
FmfbDAro = 125; % Frequency following as described by Cossette et al., 2016
FFmfbDAmax = FilterFun(FPmax,FmfbDAbend, FmfbDAro);
FoptEquivVec = [5;10;20;40;80;160];
FoptEquivMat = repmat(FoptEquivVec, 1, numF);
FmfbDAdriveMat = FmfbDAdriveFun(FmfbDAbend, Felec, FFmfbDAmax, FoptEquivVec, FmfbDAro);
~~~

which is shown here:

~~~
% Columnar data are required to plot multiple lines. Thus the input matrices have been transposed.
FF_graphMFBda = plot_freqFoll(FelecMat’,FmfbDAdriveMat’,’MFB_DA’,1,1000,1,250);
% modify graph
lgnd = legend(num2str(FoptEquivVec),’Location’,’best’);
lgnd.Title.String = “FoptEquiv”;
axh = findall(FF_graphMFBda,’Type’,’Axes’);
axh.XLabel.String = “Electrical pulse frequency”;
axh.YLabel.String = “Equivalent optical pulse frequency”;
if show_graphics
    FF_graphMFBda.Visible = ‘on’;
end
~~~

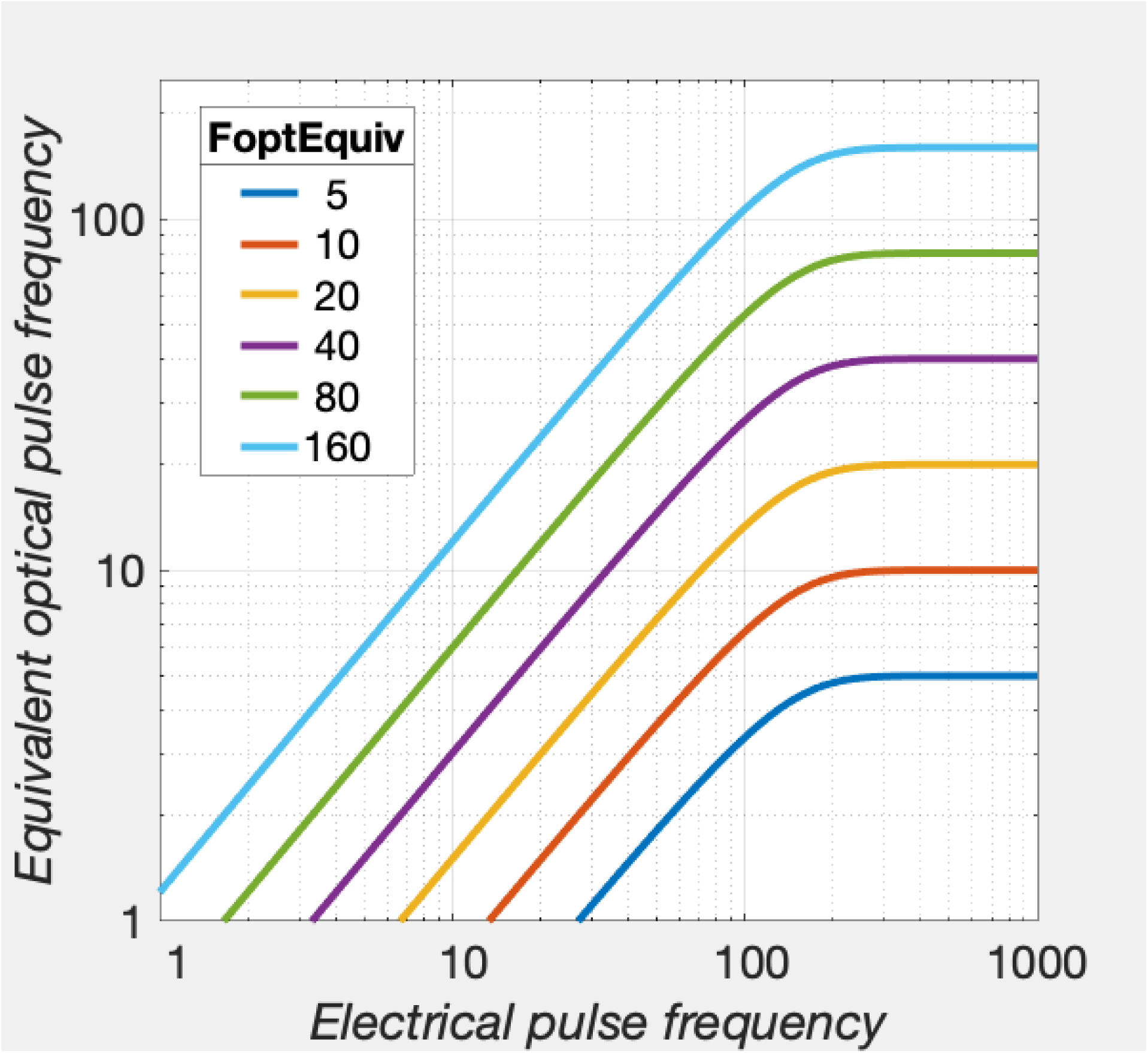

~~~
[fig_num, fig_tab] = add_fig(fig_num, fig_tab, “FreqFolMFBda”, …
    “Electrically induced firing frequency in MFB input to the dopamine neurons”);
~~~

`This is figure #50: (FreqFolMFBda)`

~~~
D = Delec;
FdaBend = 20;
FdaRO = 50;
gDA = 5;
KrgLower = 1;
NnDA = 100; % Same value as in the series-circuit section
RhoPiDA = 1572; % Same value as in the series-circuit section for D = 0.5
FPmax = 1000;
FdaHM = FpulseHMfun(C, D, FdaBend, FPmax, FdaRO, NnDA, RhoPiDA)
~~~

`FdaHM = 37.6342`

The FdaHM parameter positions the reward-growth curve for the dopamine neurons along the axis representing the dopamine firing frequency:

~~~
logFopt = 0:0.025:2.4;
Fopt = 10.^logFopt;
Rlower = fRbsr(Fopt, FdaBend, FdaHM, FdaRO, gDA, KrgLower);
TitleStrSemi = ‘Lower limb reward-intensity’;
pnam = “FdaHM”;
fnam = “FdaHM”;
% The data to be plotted must be in columns. Thus, FelecMat and RelecMat are transposed.
RG_DA_LowerEquiv_semilog = plot_RG(Fopt’,Rlower’,pnam,FdaHM,fnam,TitleStrSemi,’lin’, ‘LowerEquiv’);
RG_DA_LowerEquiv_semilog = modify_2D_graph(RG_DA_LowerEquiv_semilog, ‘Axes’, ‘XLim’, [1,250], …
   ‘RG_FdaHM_LowerEquiv_semilog’, graphs2files, FigDir);
axh = findall(RG_DA_LowerEquiv_semilog,’Type’,’Axes’);
axh.XLabel.String = “Equivalent optical pulse frequency”;
axh.YLabel.String = “Lower-limb reward intensity”;
if show_graphics
   RG_DA_LowerEquiv_semilog.Visible = ‘on’;
end
~~~

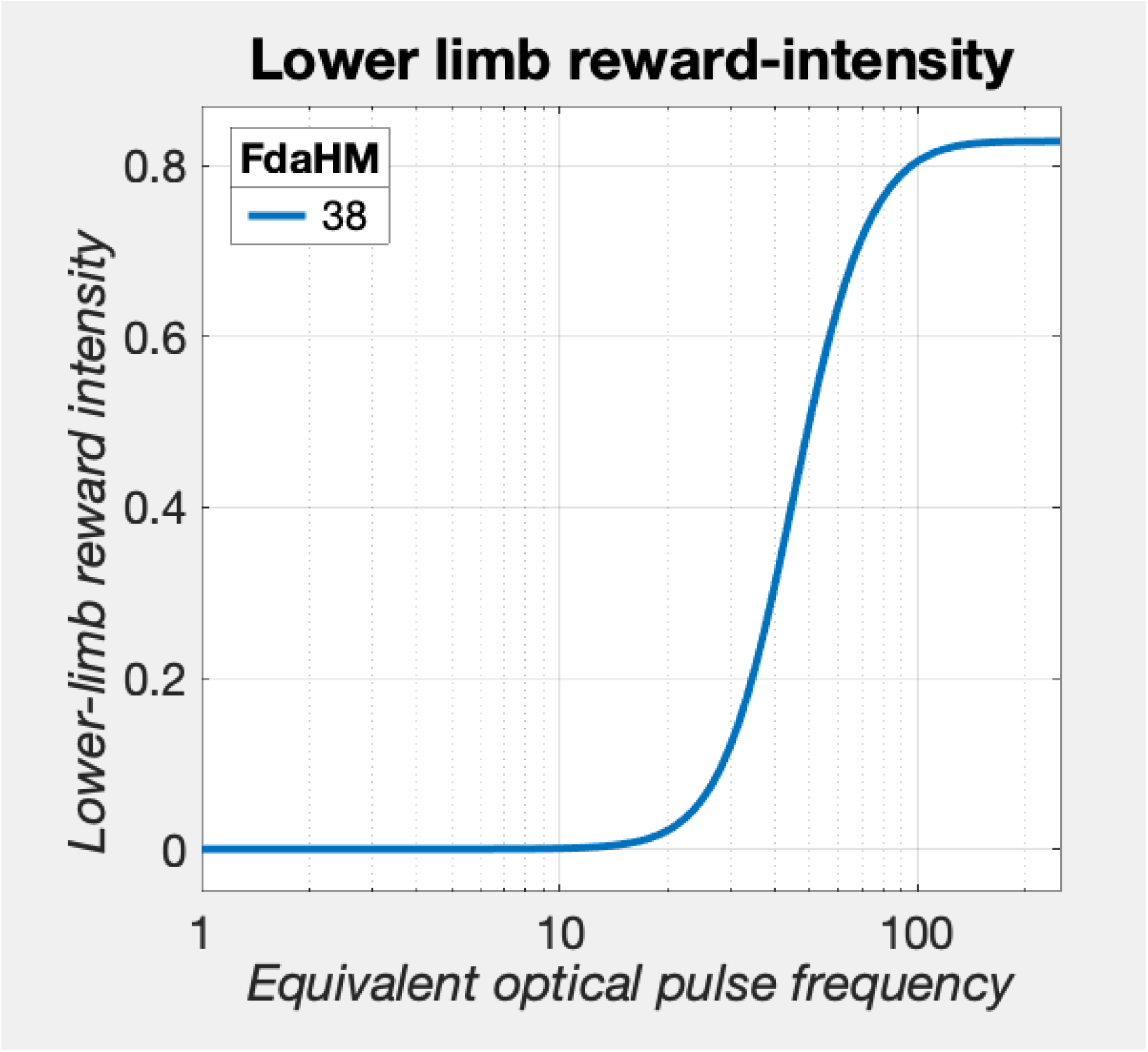

~~~
[fig_num, fig_tab] = add_fig(fig_num, fig_tab, “RG_DA_LowerEquiv_semilog”, …
    “Lower-limb reward intensity as a function of equivalent optical pulse frequencies”);
~~~

`This is figure #51: (RG_DA_LowerEquiv_semilog)`

In the following graph, we see how reward-intensity grows in the lower limb as a function of the ***electrical*** pulse frequency.

~~~
Rlower = fRbsr(FmfbDAdriveMat, FdaBend, FdaHM, FdaRO, gDA, KrgLower);
TitleStrSemi = ‘Lower-limb reward intensity’;
pnam = “optDAeq”;
fnam = “optDAeq”;
% The data to be plotted must be in columns. Thus, FelecMat and RelecMat are transposed.
RG_MFBda_lower_semilog = plot_RG(FelecMat’,Rlower’,pnam,FoptEquivVec,fnam,TitleStrSemi,’lin’, ‘lower’);
RG_MFBda_lower_semilog = modify_2D_graph(RG_MFBda_lower_semilog, ‘Axes’, ‘XLim’, [5,1000], …
     ‘RG_optDAeq_lower_semilog’, graphs2files, FigDir);
axh = findall(RG_MFBda_lower_semilog,’Type’,’Axes’);
axh.XLabel.String = “Electrical pulse frequency”;
axh.YLabel.String = “Lower-limb reward intensity”;
if show_graphics
    RG_MFBda_lower_semilog.Visible = ‘on’;
end
~~~

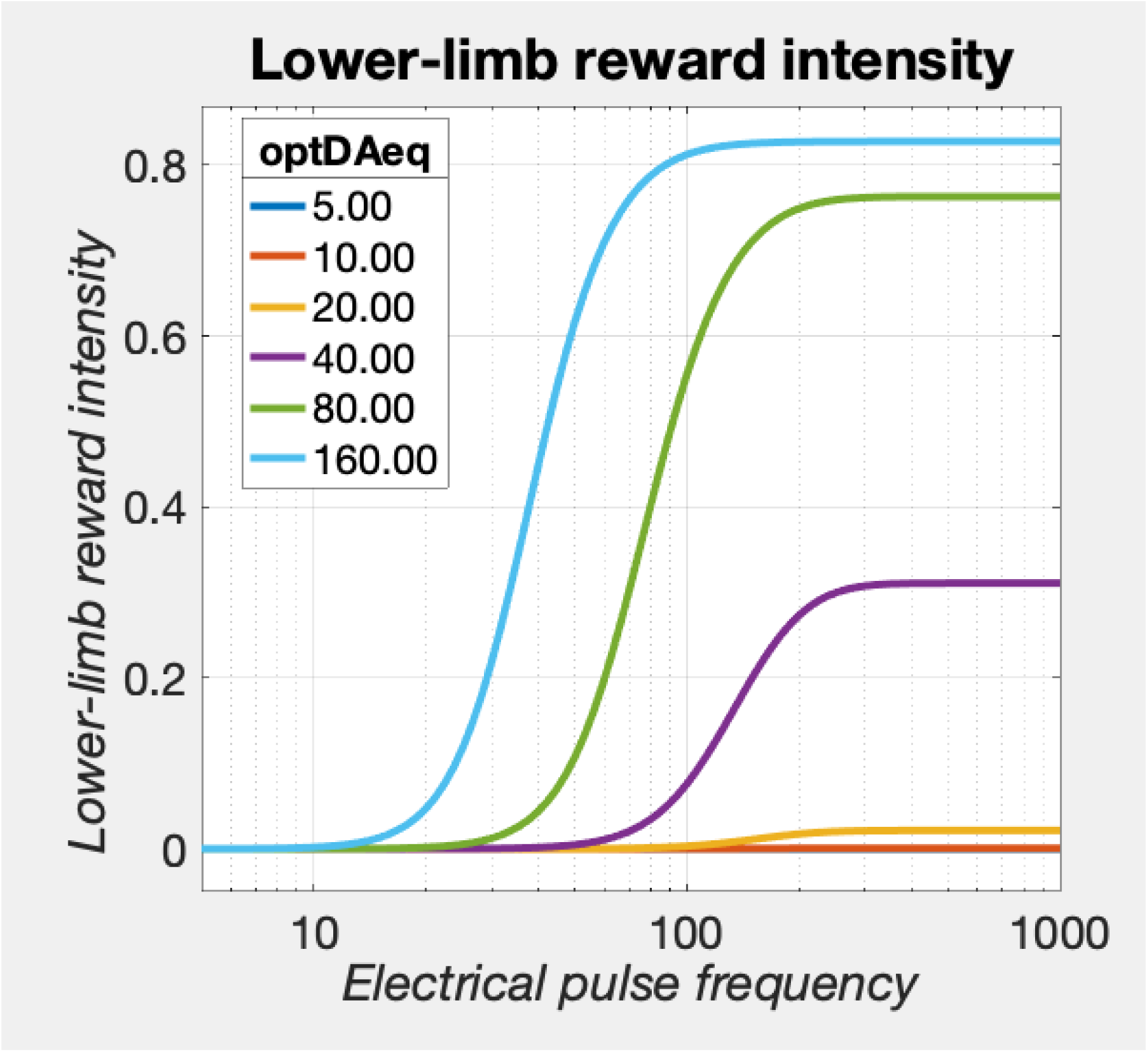

~~~
[fig_num, fig_tab] = add_fig(fig_num, fig_tab, “RG_lower_semilog”, …
    “Growth of lower-limb reward intensity at multiple values of Kmfb”);
~~~

`This is figure #52: (RG_lower_semilog)`

The *FmfbDAdrive* function expresses the MFB drive on the dopamine neurons in units of equivalent optical pulses, i.e., an FmfbDAdrive of 25 means that the effect of the trans-synaptic input from the MFB is equivalent to that produced by 25 pulses *s*^−1^ of direct optical stimulation.

The graph above shows that strong input is required to generate a substantial response from the lower limb. The reason for this is that the 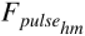 value is fairly close to the assumed frequency-following limit of the dopamine cells. The parameter values used in this simulation yield a 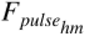 value close to the observed median in the current study when the train duration was 1 s. An adjustment is then made to predict model output for a train duration of 0.5 s, which was used in the eICSS work. This calculation are described above (see: *<u>The location parameter of the reward-growth function for BSR</u>*).

Growth curves for the summated reward intensity are shown here, at different values of the strength of the MFB drive on the dopamine neurons. The equivalent optical pulse frequencies (*FoptEquiv*) are the maximum values that the simulated MFB drive can achieve.

~~~
Rupper = repmat(Rupper,numParamVals,1); % add rows to Rupper so that it is the same size as Rlower
Rsum = Rupper + Rlower;
TitleStrSemi = ‘Summated reward intensity’;
pnam = “FoptEquiv”;
fnam = “MFBsum”;
% The data to be plotted must be in columns. Thus, FelecMat and RelecMat are transposed.
RG_MFBda_semilog = plot_RG(FelecMat’,Rsum’,pnam,FoptEquivVec,fnam,TitleStrSemi,’lin’);
RG_MFBda_semilog = modify_2D_graph(RG_MFBda_semilog, ‘Axes’, ‘XLim’, [5,1000], …
     ‘RG_MFBda_semilog’, graphs2files, FigDir);
axh = findall(RG_MFBda_semilog, ‘Type’, ‘Axes’);
axh.XLabel.String = “Electrical pulse frequency”;
axh.YLabel.String = “Upper + lower reward intensity”;
if show_graphics
    RG_MFBda_semilog.Visible = ‘on’;
end
~~~

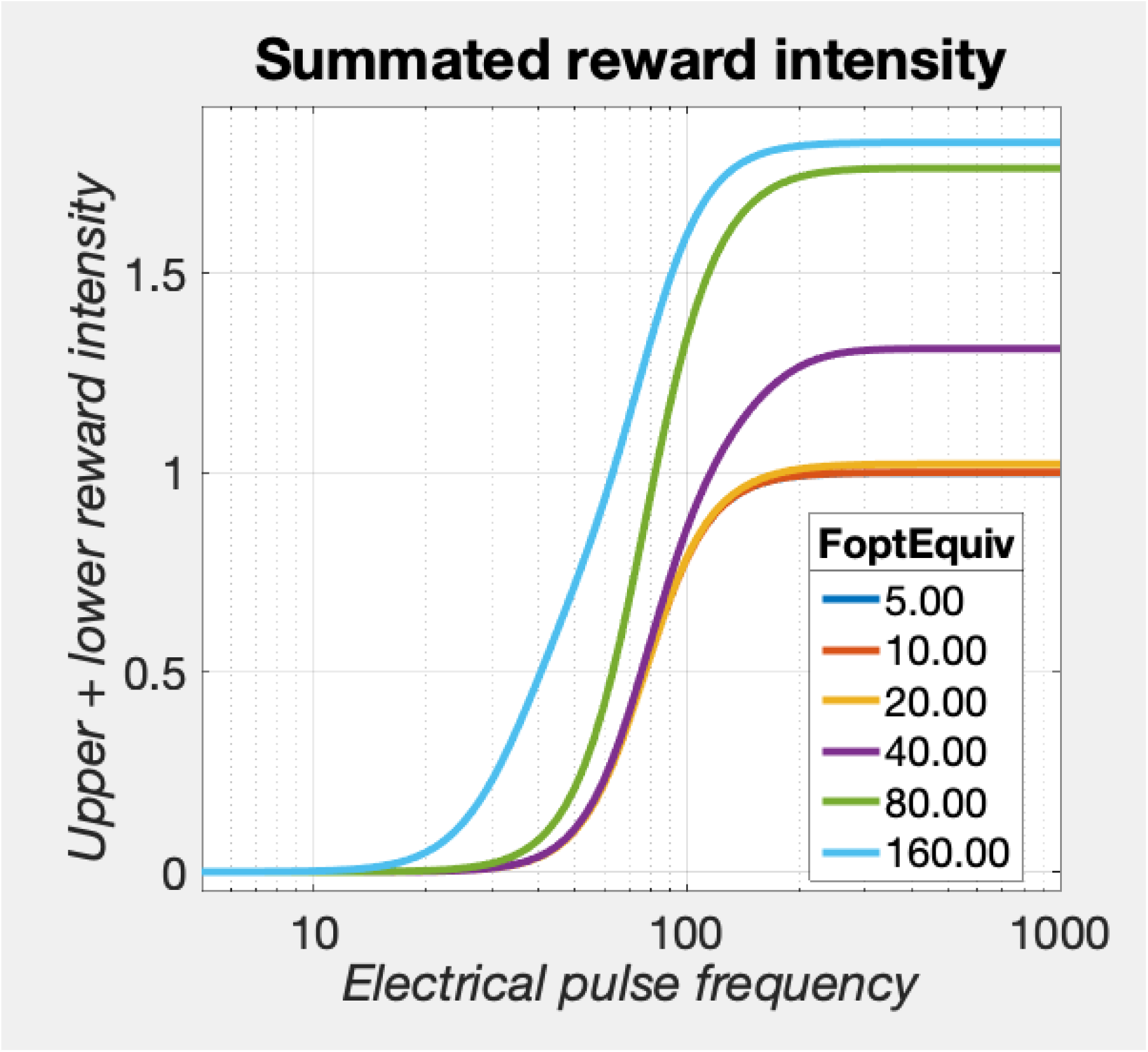

The reduced slope of the early portion of the leftmost curve is due to the fact that reward intensity in the lower limb begins rising at lower MFB pulse frequencies than reward intensity in the upper limb when the drive on the dopamine neurons is highest.

~~~
% Section break here to force display of RG_MFBda_semilog
~~~

~~~
close all;
[fig_num, fig_tab] = add_fig(fig_num, fig_tab, “RG_Kmfb_semilog”, …
    “Growth of summated reward intensity at at multiple values of Kmfb”);
~~~

`This is figure #53: (RG_Kmfb_semilog)`

To prepare for simulation of the mountain generated by the convergence model, we first derive the electrical pulse frequencies that produce half-maximal, summated reward intensity for each value of *FoptEquiv*

~~~
% Find FsumHM by means of interpolation
numF = 121;
Felec = logspace(0,3,numF);
[FsumHM, RsumMax] = find_FhmSum(Felec, Rsum);
~~~

We generate a mountain surface, scaling the maximum MFB drive on the dopamine neurons to be equivalent to an optical pulse frequency of 80 pulses *s*^−1^, a maximal or near-maximal value in the current study.

~~~
disp(string({strcat({‘Figure ‘},num2str(fig_tab.Number(fig_tab.Name==‘RG_Kmfb_semilog’)))}));
~~~

`Figure 53`

~~~
FoptEquiv = 80;
RupperFoptEquiv = Rupper(FoptEquivVec==FoptEquiv,:); % Selects the appropriate row from the matrix
RupperMax = max(RupperFoptEquiv);
RlowerFoptEquiv = Rlower(FoptEquivVec==FoptEquiv,:); % Selects the appropriate row from the matrix
RlowerMax = max(RlowerFoptEquiv);
Rsum = Rsum(FoptEquivVec==FoptEquiv,:);
FsumHM = FsumHM(FoptEquivVec==FoptEquiv);
RsumMax = RsumMax(FoptEquivVec==FoptEquiv);
FPmax = 1000;
RupperNormMax = fRbsrNorm(FPmax, FelecBend, FhmUpper, FelecRO, gElec);
RlowerNormMax = fRbsrNorm(FoptEquiv, FdaBend, FdaHM, FdaRO, gDA);
RupperRatio = RupperMax / (RlowerMax + RupperMax)
~~~

`RupperRatio = 0.5671`

~~~
RlowerRatio = RlowerMax / (RlowerMax + RupperMax)
~~~

`RlowerRatio = 0.4329`

~~~
RsumNormMax = (RupperRatio * RupperNormMax) + (RlowerRatio * RlowerNormMax); % Weighted average
dotPhiObj = 1;
Kaa = 1;
Kec = 1;
Krg = 1;
dotRaa = 0.1;
pObj = 1;
PsubBend = 0.5;
PsubMin = 1.82;
PobjE = PobjEfun(dotPhiObj, Kaa, Kec, Krg, dotRaa, pObj, PsubBend, PsubMin, RsumNormMax)
~~~

`PobjE = 8.9716`

~~~
a = 3;
numP = numF;
Pobj = logspace(0,3,numP); % row variable
Tsum = TAsumFun(a, Pobj, PobjE, PsubBend, PsubMin, Rsum’, RsumNormMax);
% n.b., numel(Rsumxx) = numel(Pobj). Rsumxx has been transposed. Thus, Tsumxx is a square matrix.
MTNsum = plot_MTN(Felec, Pobj, Tsum, ‘off’, strcat(‘MTNsum’,num2str(FoptEquiv)), ‘reward mountain’, …
    graphs2files, FigDir);
if show_graphics
    MTN.Visible = ‘on’;
end
~~~

We now simulate the effect of dopamine-transporter blockade.

`Elapsed time is 11.751399 seconds.`

~~~
tic;
close all;
~~~

### The effect of dopamine transporter blockade in the convergence model

The reward-intensity signal produced by the upper and lower limbs of the circuit shown in

~~~
disp(string({strcat({‘Figure ‘},num2str(fig_tab.Number(fig_tab.Name==“Convergence_model”)))}));
~~~

`Figure 48`

will first be computed separately and then combined additively, with two different values of ***K***_*da*_, one representing the vehicle condition, and the second representing the condition in which the dopamine-transporter blocker has been administered.

~~~
% Reward intensity produced by the upper limb
C = 0.473; % median from Sonnenschein et al., 2003
Delec = 0.5;
logFelecBend = 1.3222; % from Solomon et al., 2015
FelecBend = 10^logFelecBend;
logFelecRO = 2.5587; % from Solomon et al., 2015
FelecRO = 10^logFelecRO;
NnMFB = 126;
RhoPi = 5000;
FPmax = 1000;
FhmUpper = FpulseHMfun(C, Delec, FelecBend, FPmax, FelecRO, NnMFB, RhoPi)
~~~

`FhmUpper = 77.2222`

~~~
numF = 121;
numParamVals = 2; % for two values of Kda
Felec = logspace(0,3,numF);
FelecMat = repmat(Felec,numParamVals,1);
gElec = 5;
KrgUpper = 1;
Rupper = fRbsr(Felec, FelecBend, FhmUpper, FelecRO, gElec, KrgUpper);
% Reward intensity produced by the lower limb
% First compute the drive on the DA neurons in terms of the equivalent optical pulse frequency
FmfbDAbend = 40;
FmfbDAro = 125;
FFmfbDAmax = FilterFun(FPmax,FmfbDAbend, FmfbDAro);
FoptEquiv = 80; % maximal or near-maximal value in the current study
FmfbDAdrive = FmfbDAdriveFun(FmfbDAbend, Felec, FFmfbDAmax, FoptEquiv, FmfbDAro);
% Calculate FdaHM for the vehicle and drug conditions
C = 0.473; % median from Sonnenschein et al., 2003
D = Delec;
FdaBend = 20;
FdaRO = 50;
Kda = [1,10^0.15]; % Median for current oICSS study: 10^0.1446
gDA = 5; % Median in vehicle condition of current oICSS study: 4.61
KrgLower = 1;
NnDA = 100; % Make sure to use the same value as in the series-circuit section
RhoPiDA = 1572; % Same value as in the series-circuit section for D=0.5
FPmax = 1000;
FdaHMkDA = FpulseHMfun(C, D, FdaBend, FPmax, FdaRO, NnDA, RhoPiDA, Kda) % result is a two-element vector
~~~

~~~
FdaHMkDA = 1×2
    37.6342 25.1515
~~~

~~~
% Compute reward growth in the lower limb as a function of the equivalent optical pulse frequency
numFOpt = 97; % 0.025 log10 spacing from 0 to 2.4
numParamVals = 2; % 2 values of Kda Fopt = logspace(0,2.4,numFOpt);
FoptMat = repmat(Fopt,numParamVals,1); % number of rows = number of Kda vals
KdaMat = repmat(Kda’,1,numFOpt); % make matrix the same size as FoptMat
FdaHMmat = repmat(FdaHMkDA’,1,numFOpt); % make matrix the same size as FoptMat
Rlower = fRbsr(FoptMat, FdaBend, FdaHMmat, FdaRO, gDA, KrgLower);
TitleStrSemi = ‘Lower limb vs Fopt equivalent’;
pnam = “Kda”;
fnam = “Kda”;
% The data to be plotted must be in columns. Thus, FoptMat and Rlower are transposed.
RG_MFB_DA_FoptEquiv_Kda = plot_RG(FoptMat’,Rlower’,pnam,Kda,fnam,TitleStrSemi,’lin’, ‘FoptEquiv’);
RG_MFB_DA_FoptEquiv_Kda = modify_2D_graph(RG_MFB_DA_FoptEquiv_Kda, ‘Axes’, ‘XLim’, [1,250], …
    ‘RG_Kda_FoptEquiv_semilog’, graphs2files, FigDir);
axh = findall(RG_MFB_DA_FoptEquiv_Kda, ‘Type’,’Axes’);
axh.XLabel.String = “Equivalent optical pulse frequency”;
axh.YLabel.String = “Lower-limb reward intensity”;
if show_graphics
     RG_MFB_DA_FoptEquiv_Kda.Visible = ‘on’;
end
~~~

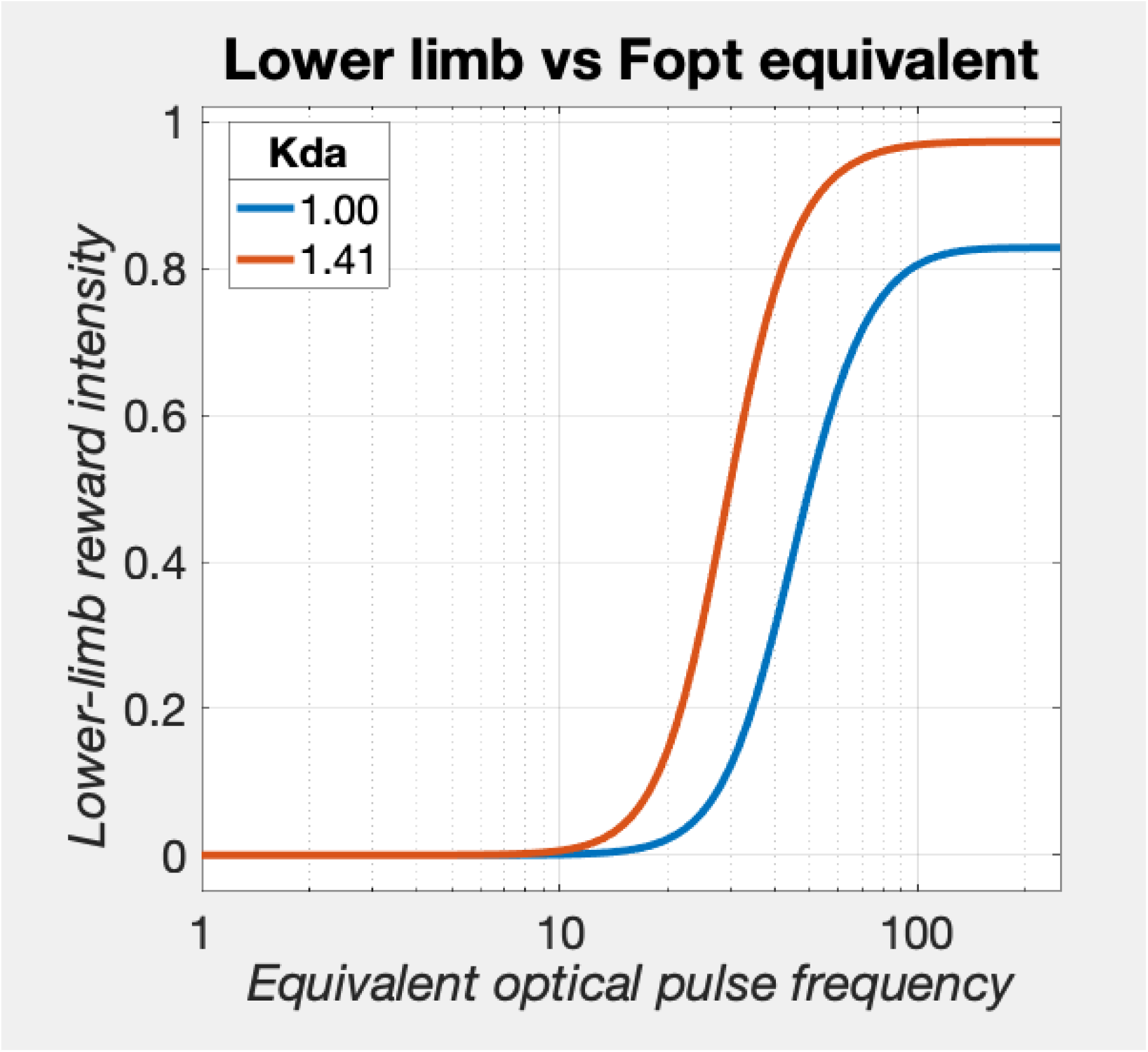

~~~
[fig_num, fig_tab] = add_fig(fig_num, fig_tab, “RG_MFB_DA_FoptEquiv_Kda”, …
    “Reward growth in lower limb as a function of equivalent optical pulse frequency”);
~~~

`This is figure #54: (RG_MFB_DA_FoptEquiv_Kda)`

The previous graph shows lower-limb reward intensity as a function of the **equivalent optical pulse frequency** in the presence and absence of dopamine-transporter blockade. The next graph will plot lower-limb reward intensity as a function of the **electrical MFB pulse frequency**, again in the presence and absence of dopamine transporter blockade.

~~~
KdaMat = repmat(Kda’,1,numF); % make matrix the same size as FelecMat
FdaHMmat = repmat(FdaHMkDA’,1,numF); % make matrix the same size as FelecMat
Rlower = fRbsr(FmfbDAdrive, FdaBend, FdaHMmat, FdaRO, gDA, KrgLower);
TitleStrSemi = ‘Lower limb vs Felec’;
pnam = “Kda”;
fnam = “Kda”;
% The data to be plotted must be in columns. Thus, FelecMat and RelecMat are transposed.
RG_MFB_DA_Felec_Kda = plot_RG(FelecMat’,Rlower’,pnam,Kda,fnam,TitleStrSemi,’lin’, ‘Felec’);
RG_MFB_DA_Felec_Kda = modify_2D_graph(RG_MFB_DA_Felec_Kda, ‘Axes’, ‘XLim’, [5,500], …
    ‘RG_Kda_Felec_semilog’, graphs2files, FigDir);
axh = findall(RG_MFB_DA_Felec_Kda, ‘Type’,’Axes’);
axh.XLabel.String = “Electrical pulse frequency”;
axh.YLabel.String = “Lower-limb reward intensity”;
if show_graphics
    RG_MFB_DA_Felec_Kda.Visible = ‘on’;
end
~~~

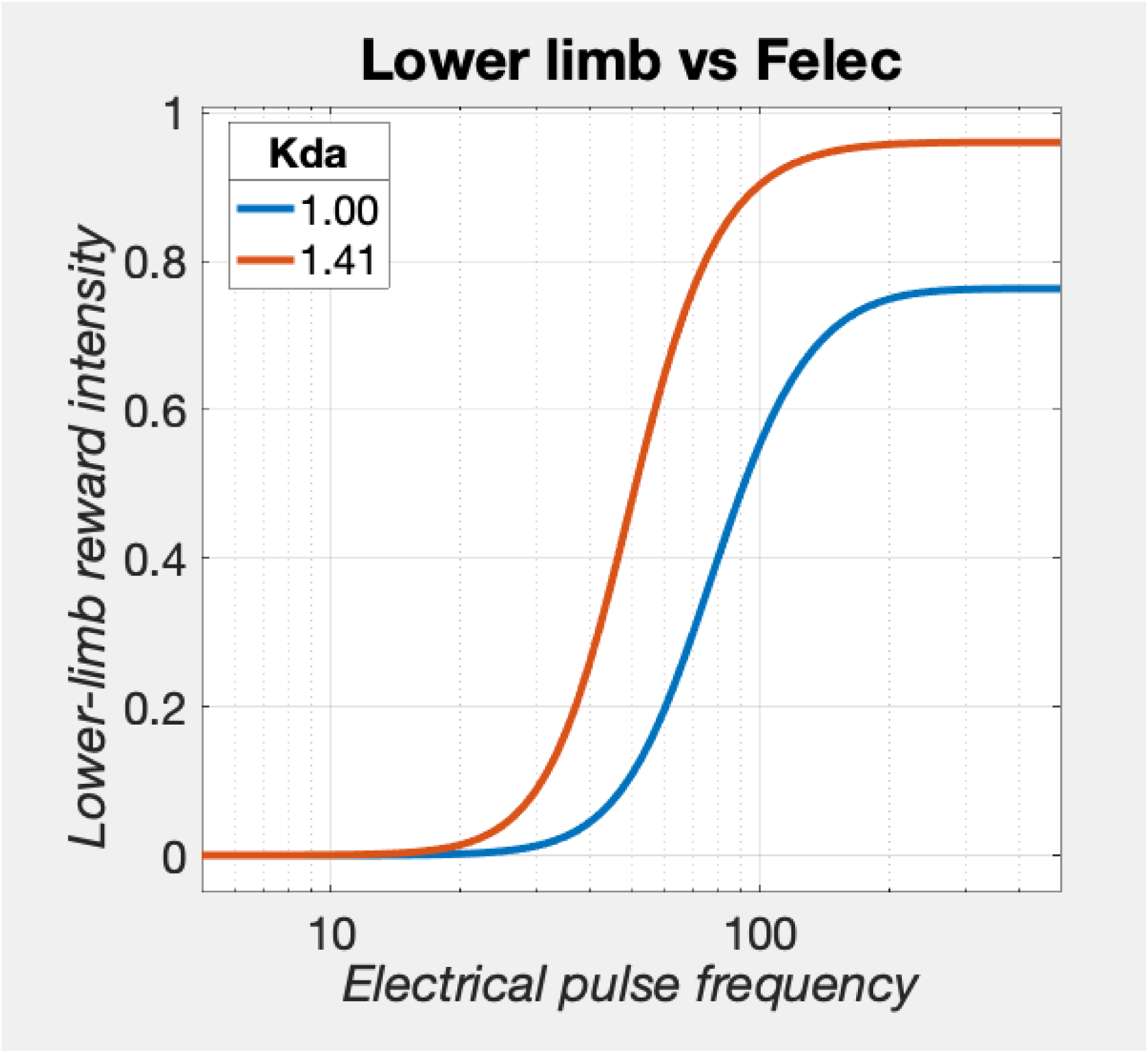

~~~
[fig_num, fig_tab] = add_fig(fig_num, fig_tab, “RG_MFB_DA_Felec_Kda”, …
“Reward growth in lower limb as a function of electrical pulse frequency”);
~~~

`This is figure #55: (RG_MFB_DA_Felec_Kda)`

In the convergence model, the reward-intensity signal in the upper limb is not changed by dopamine-transporter blockade. Thus, we can simply add the reward-intensity signal already computed to the results for the lower-limb reward-intensity signal to obtain the summated output of the two limbs. This corresponds to the ‘benefit’ signal at the right of

~~~
disp(string({strcat({‘Figure ‘},num2str(fig_tab.Number(fig_tab.Name==“Convergence_model”)))}));
~~~

`Figure 48`

~~~
Rupper = repmat(Rupper,numParamVals,1); % add rows to Rupper so that it is the same size as Rlower
Rsum = Rupper + Rlower;
~~~

The following graph shows the summated reward-intensity signal in response to MFB drive equivalent to an optical pulse frequency of

~~~
FoptEquiv
~~~

`FoptEquiv = 80`

pulses *s*^−1^

~~~
[FsumHM, RsumMax] = find_FhmSum(FelecMat, Rsum)
~~~

~~~
FsumHM = 2×1
   75.2675
   63.0957
RsumMax = 2×1
   1.7626
   1.9598
~~~

~~~
% plot Rupper, Rlower, and Rxor for the vehicle and drug conditions
% Three curves will be plotted per graph; the matrices must be dimensioned accordingly
FelecMatPlot = repmat(Felec, 3, 1);
RmatVeh = [Rupper(1,:);Rlower(1,:);Rsum(1,:)];
RmatDrg = [Rupper(2,:);Rlower(2,:);Rsum(2,:)];
pnam = “1:up 2:lo 3:sum”;
pVec = [1,2,3];
fnam = strcat(“Sum”,num2str(FoptEquiv),”Veh”);
TitleStr = strcat(“FoptEquiv=“,num2str(FoptEquiv),”; vehicle”);
RGsum_Rveh = plot_RG(FelecMatPlot’,RmatVeh’,pnam,pVec,fnam,TitleStr,’lin’);
axh = findall(RGsum_Rveh, ‘Type’, ‘Axes’);
axh.XLabel.String = ‘Electrical pulse frequency’;
lh = findall(RGsum_Rveh, ‘Type’, ‘Line’);
lcolor = {‘m’,’g’,’c’};
for j=1:3
    lh(j).Color = lcolor{j};
end
if show_graphics
    RGsum_Rveh.Visible = ‘on’;
end
~~~

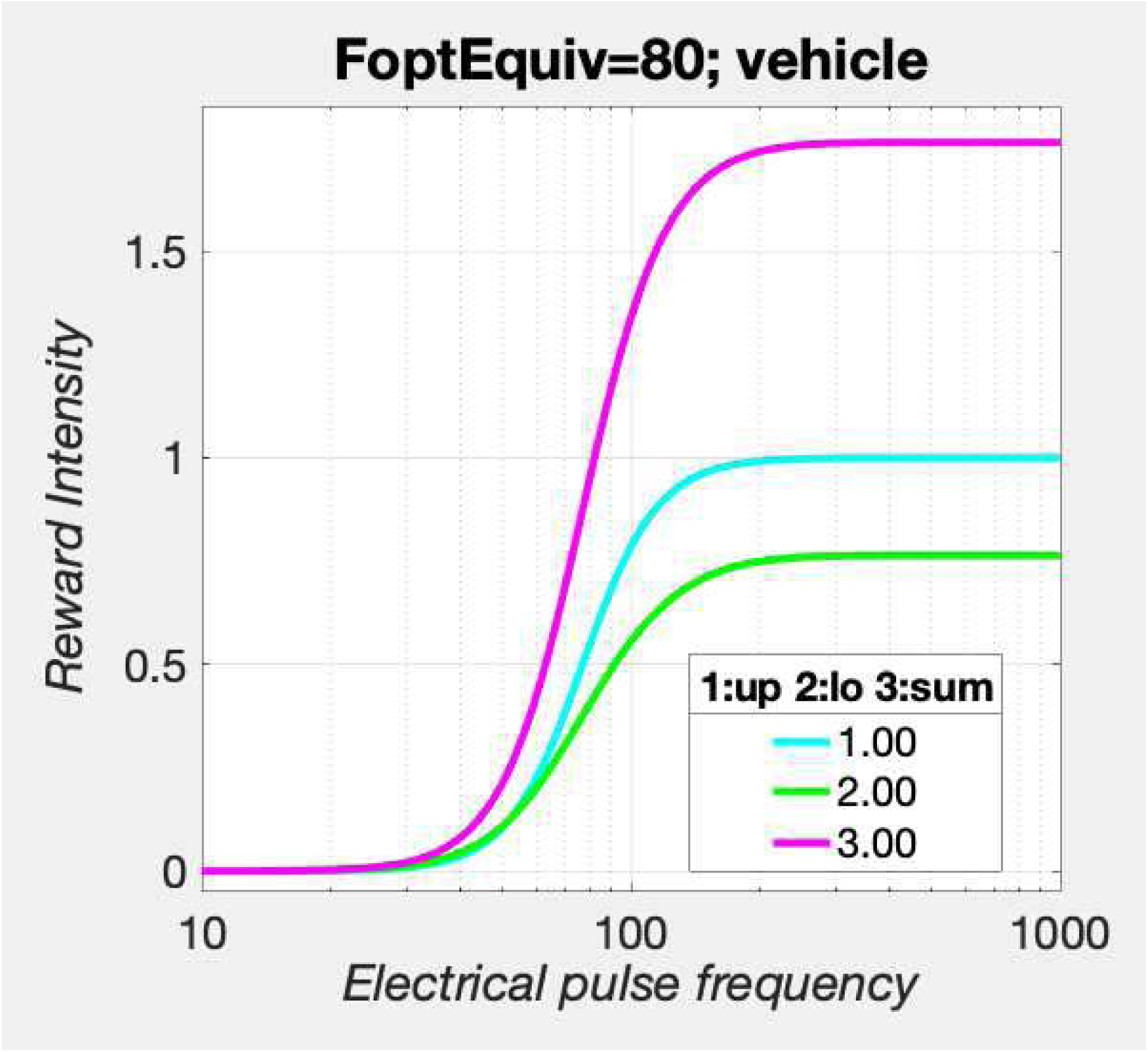

~~~
fig_nam = strcat(“RGsum_Veh_FoptEquiv”,num2str(FoptEquiv));
[fig_num, fig_tab] = add_fig(fig_num, fig_tab, fig_nam, “Reward summation in the convergence model”);
~~~

`This is figure #56: (RGsum_Veh_FoptEquiv80)`

~~~
fnam = strcat(“sum”,num2str(FoptEquiv),”Drg”);
TitleStr = strcat(“FoptEquiv=“,num2str(FoptEquiv),”; drug”);
RGsum_Rdrg = plot_RG(FelecMatPlot’,RmatDrg’,pnam,pVec,fnam,TitleStr,’lin’);
axh = findall(RGsum_Rdrg, ‘Type’, ‘Axes’);
axh.XLabel.String = ‘Electrical pulse frequency’;
lh = findall(RGsum_Rdrg, ‘Type’, ‘Line’);
lcolor = {‘m’,’g’,’c’};
for j=1:3
    lh(j).Color = lcolor{j};
    lh(j).LineStyle = ‘--’;
end
if show_graphics
    RG_RDrg.Visible = ‘on’;
end
fig_nam = strcat(“RGsum_Drg_FoptEquiv”,num2str(FoptEquiv));
[fig_num, fig_tab] = add_fig(fig_num, fig_tab, fig_nam, “Reward summation in the convergence model”);
~~~

`This is figure #57: (RGsum_Drg_FoptEquiv80)`

~~~
shg
~~~

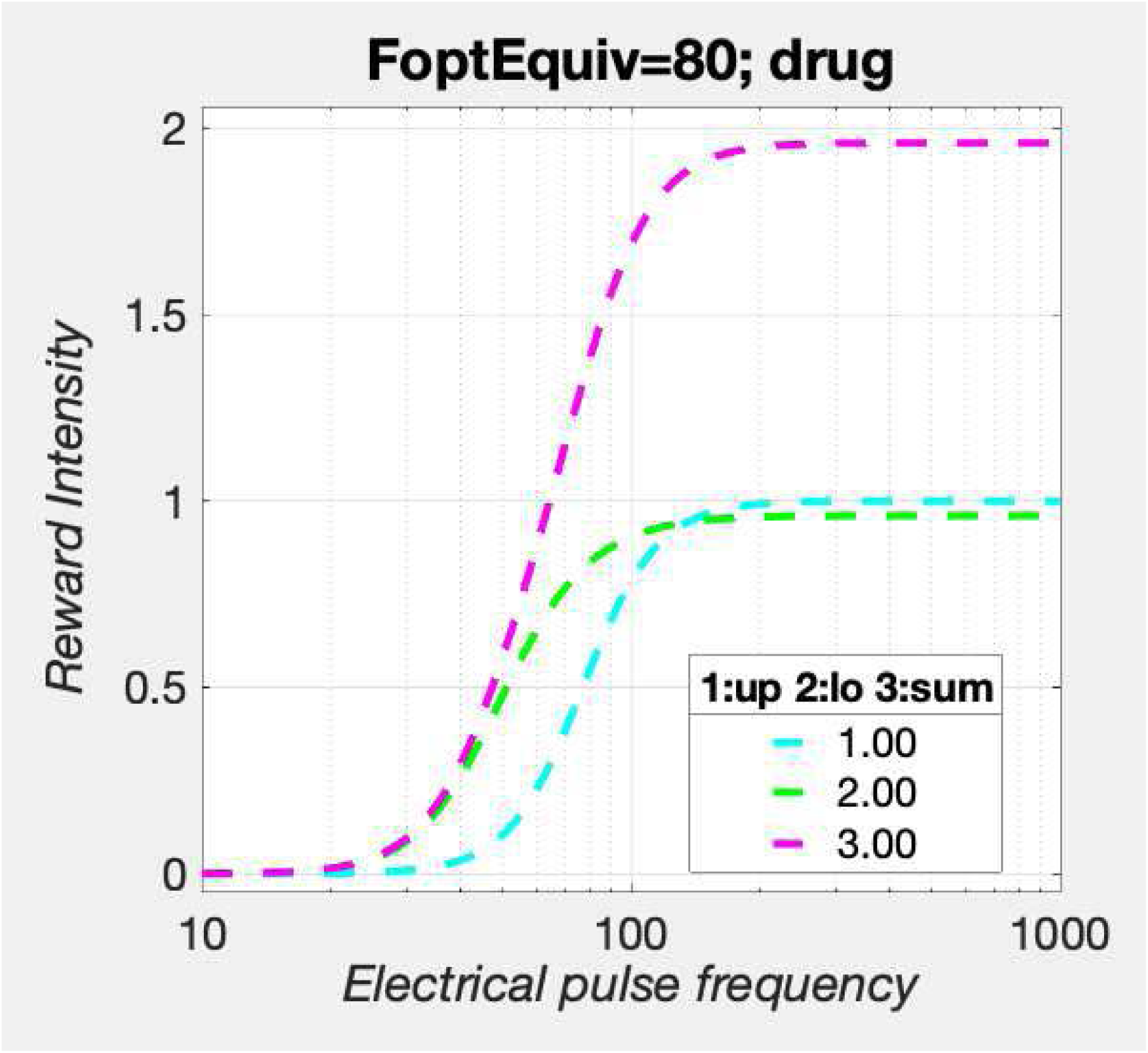

~~~
pnam = “Kda”;
pVec = [Kda(1),Kda(2)];
fnam = strcat(“Sum”,num2str(FoptEquiv),”VehDrg”);
TitleStr = strcat(“FoptEquiv=“,num2str(FoptEquiv),”; sum”);
RGsum_Rvehdrg = plot_RG(FelecMat’,Rsum’,pnam,pVec,fnam,TitleStr,’lin’);
axh = findall(RGsum_Rvehdrg, ‘Type’, ‘Axes’);
axh.XLabel.String = ‘Electrical pulse frequency’;
lh = findall(RGsum_Rvehdrg, ‘Type’, ‘Line’);
for j=1:2
    lh(j).Color = ‘m’;
end
lh(1).LineStyle = ‘--’;
lgndh = findall(RGsum_Rvehdrg,’Type’,’Legend’);
lgndh.Title.String = ‘condition’;
lgndh.String = {‘vehicle’,’drug’};
if show_graphics
    RGsum_Rvehdrg.Visible = ‘on’;
end
~~~

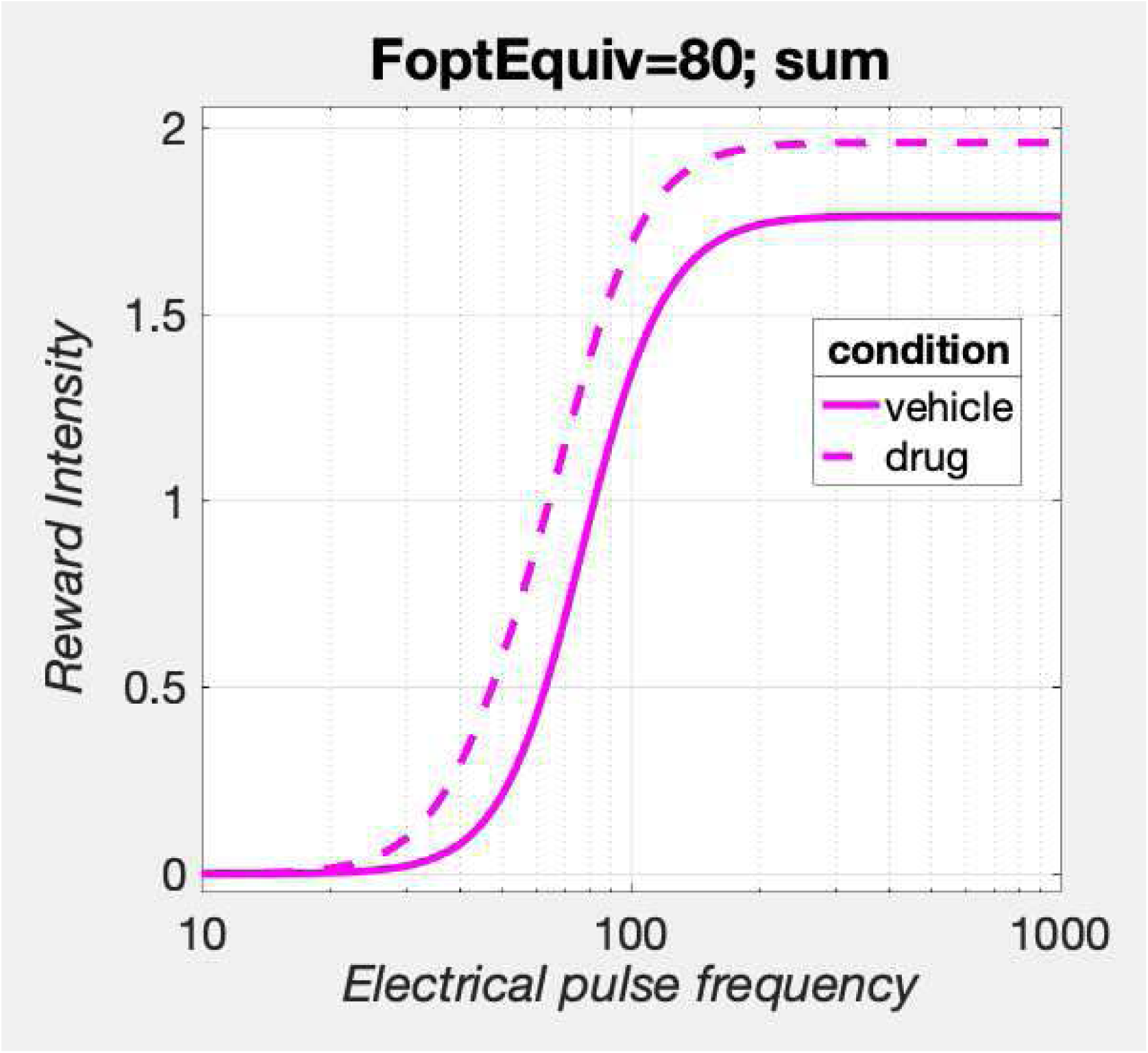

~~~
fig_nam = strcat(“RGsum_VehDrg_FoptEquiv”,num2str(FoptEquiv));
[fig_num, fig_tab] = add_fig(fig_num, fig_tab, fig_nam, “Drug modulation of reward summation in the convergence model”);
~~~

`This is figure #58: (RGsum_VehDrg_FoptEquiv80)`

At this high value of MFB drive, a small leftward curve shift is produced.

With the reward-intensity signal in hand, we can proceed to generate the mountains predicted in the presence and absence of dopamine-transporter blockade.

~~~
close all;
FPmax = 1000;
FhmUpperVec = repmat(FhmUpper,1,2); % Upper limb output is independent of Kda
RupperNormMax = fRbsrNorm(FPmax, FelecBend, FhmUpperVec, FelecRO, gElec)’;
RlowerNormMax = fRbsrNorm(FoptEquiv, FdaBend, FdaHMkDA, FdaRO, gDA)’;
RupperMax = max(Rupper, [], 2);
RlowerMax = max(Rlower, [], 2);
RupperRatio = RupperMax ./ (RlowerMax + RupperMax)
~~~

~~~
RupperRatio = 2×1
    0.5671
    0.5100
~~~

~~~
RlowerRatio = RlowerMax ./ (RlowerMax + RupperMax)
~~~

~~~
RlowerRatio = 2×1
    0.4329
    0.4900
~~~

~~~
RsumNormMax = (RupperRatio .* RupperNormMax) + (RlowerRatio .* RlowerNormMax); % Weighted average
dotPhiObj = 1;
Kaa = 1;
Kec = 1;
Krg = 1;
dotRaa = 0.1;
pObj = 1;
PsubBend = 0.5;
PsubMin = 1.82;
PobjE = PobjEfun(dotPhiObj, Kaa, Kec, Krg, dotRaa, pObj, PsubBend, PsubMin, RsumNormMax)
~~~

~~~
PobjE = 2×1
    8.9716
    9.8029
~~~

~~~
a = 3;
numP = numF;
Pobj = logspace(0,3,numP); % row variable
TsumKda1 = TAsumFun(a, Pobj, PobjE(1), PsubBend, PsubMin, Rsum(1,:)’, RsumNormMax(1));
TsumKda2 = TAsumFun(a, Pobj, PobjE(2), PsubBend, PsubMin, Rsum(2,:)’, RsumNormMax(2));
% n.b., numel(Rsum(x)) = numel(Pobj). Rsum(x) has been transposed. Thus, TsumKda(x) is a square matrix.
% Plot the individual mountains for the two values of Kda
title_str1 = strcat({‘Kda = ‘}, sprintf(‘%2.2f’, Kda(1)));
MTNsumKda1 = plot_MTN(Felec, Pobj, TsumKda1, ‘off’, ‘MTNsumKda1’, title_str1, …
     graphs2files, FigDir);
title_str2 = strcat({‘Kda = ‘}, sprintf(‘%2.2f’, Kda(2)));
MTNsumKda2 = plot_MTN(Felec, Pobj, TsumKda2, ‘off’, ‘MTNsumKda1’, title_str2, …
     graphs2files, FigDir);
% Plot the pair of mountains for the two values of Kda
dual_sum_plot = dual_subplot(MTNsumKda1, MTNsumKda2, ‘MTNsum_Kda1_Kda2’,…
     graphs2files,FigDir);
if show_graphics
     dual_sum_plot.Visible = ‘on’;
end
~~~

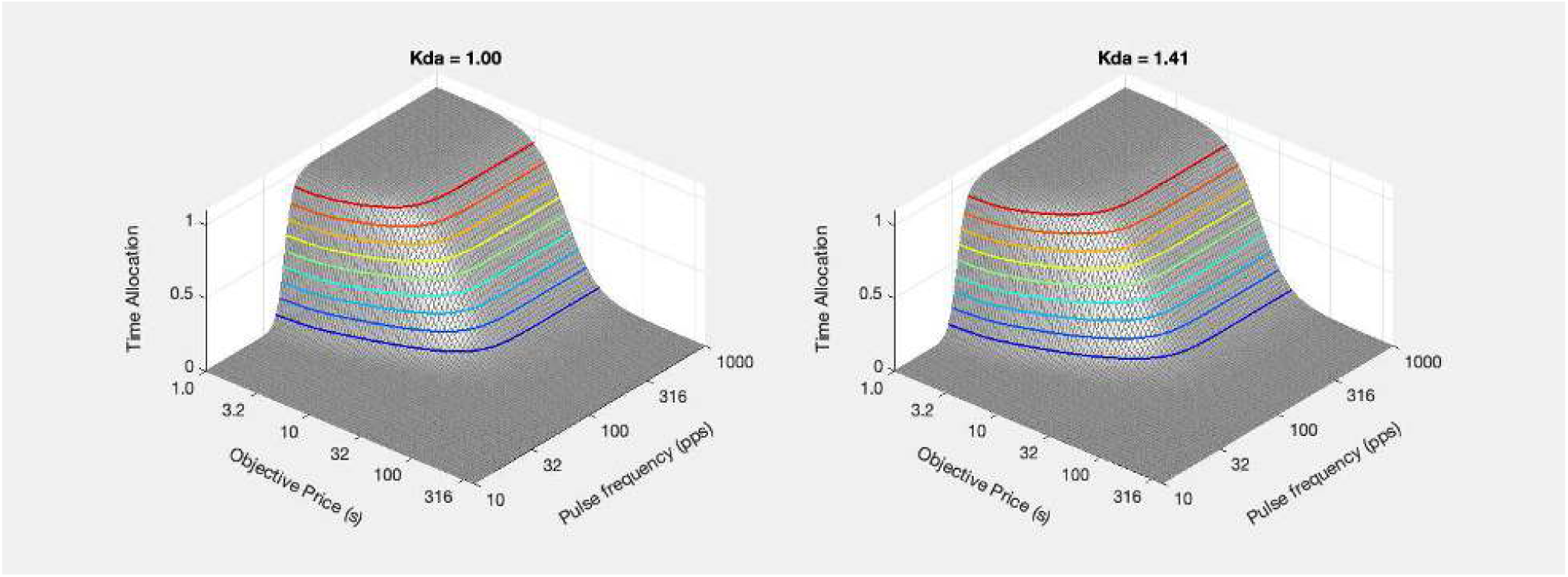

~~~
% Estimate the weighted average of the firing frequencies that produce half-maximal reward intensity in the
% upper and lower limbs
FhmUpperStar = FilterFun(FhmUpper,FelecBend,FelecRO);
FhmLowerDrive = FmfbDAdriveFun(FmfbDAbend, FsumHM, FFmfbDAmax, FoptEquiv, FmfbDAro);
FFhmLower = FilterFun(FhmLowerDrive,FdaBend, FdaRO);
FsumHMstar = FsumHM .* …
     ((RupperRatio .* (FhmUpperStar ./ FsumHM)) + (RlowerRatio .* (FFhmLower ./ FhmLowerDrive)));
% Weighted average of the frequency-following fidelity in the upper and lower limbs
PsubE = PsubFun(PobjE,PsubBend,PsubMin);
PsubEstar = PsubE ./ RsumNormMax;
PobjEstar = PsubBsFun(PsubEstar,PsubBend,PsubMin);
ContkDA1sum = plot_contour(Felec, Pobj, TsumKda1, PobjE(1), FsumHM(1), ‘off’, ‘ContkDA1sum’, title_str1, …
    strcat({‘Kda = ‘}, num2str(Kda(1))), graphs2files, FigDir);
ContkDA2sum = plot_contour(Felec, Pobj, TsumKda2, PobjE(2), FsumHM(2), ‘off’, ‘ContkDA2sum’, title_str2, …
    strcat({‘Kda = ‘}, num2str(Kda(2))), graphs2files, FigDir);
bg_kDA1vskDA2sum = plot_bgStar(FsumHM(1), FsumHM(2), FsumHMstar(1), FsumHMstar(2),…
    PobjE(1), PobjE(2), PobjEstar(1), PobjEstar(2),…
    ‘off’, ‘kDA1vskDA2sum_bg’, graphs2files, FigDir, −0.3, 0.3);
% Set a scale for the y-axis of this bar graph that ressembles the scales for the others
bg_root = ‘bg_kDA1vskDA2sum’;
quad_kDA1vskDA2sum = quad_subplot(ContkDA1sum, ContkDA2sum, bg_kDA1vskDA2sum, ‘quad_kDA1vskDA2sum’, bg_root, …
    graphs2files, FigDir);
if show_graphics
    quad_kDA1vskDA2sum.Visible = ‘on’;
end
~~~

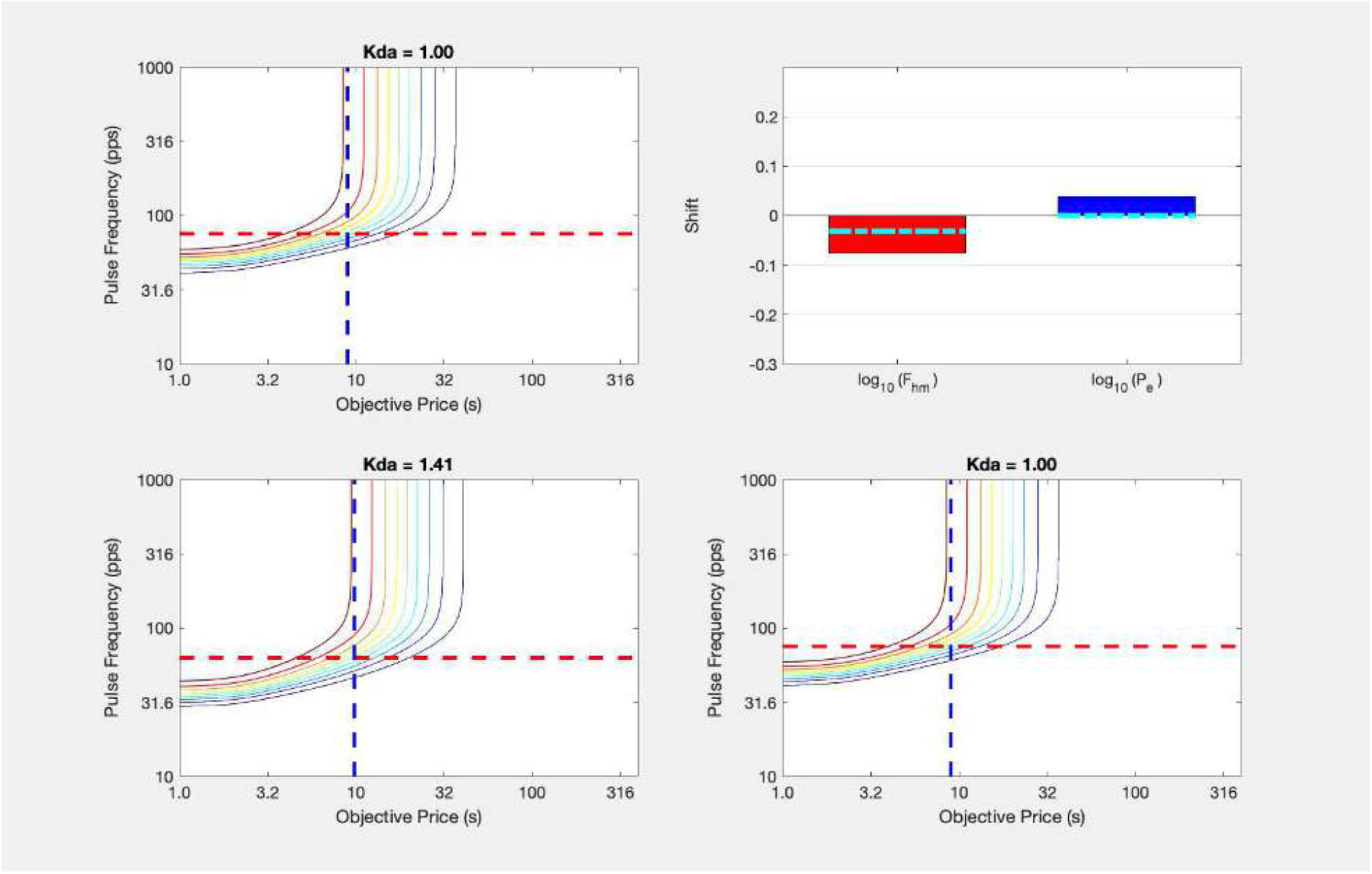

~~~
fig_nam = strcat(“quad_kDA1vskDA2sum”,num2str(FoptEquiv));
[fig_num, fig_tab] = add_fig(fig_num, fig_tab, …
    fig_nam, “Modest downward shift of the reward mountain induced by dopamine-transporter blockade”);
~~~

`This is figure #59: (quad_kDA1vskDA2sum80)`

~~~
% Insert section break to force display of the last graph
~~~

~~~
close all;
~~~

As expected from the small leftward displacement of the reward-intensity growth function by dopamine transporter blockade shown in

~~~
disp(string({strcat({‘Figure ‘},num2str(fig_tab.Number(fig_tab.Name==“RGsum_VehDrg_FoptEquiv80”)))}));
~~~

`Figure 58`

the mountain is shifted modestly along the pulse-frequency axis. None of the eight subjects in the Hernandez et al. (2012) eICSS study showed such a shift.

Adding dopaminergic modulation of effort cost (i.e., a drug-induced reduction in these costs) boosts the shift along the price axis. The same effect could be produced by boosting the output of the reward-growth function(s) (↑***K***_*rg*_) or decreasing the value of alternate activities.

With dopaminergic modulation of effort cost:

~~~
KeffMod = [1;10^0.15]; % Median shift in Pe in the current oICSS study: 10^0.1525
PobjE = PobjEfun(dotPhiObj, Kaa, Kec, Krg, dotRaa, pObj, PsubBend, PsubMin, RsumNormMax, KeffMod)
~~~

~~~
PobjE = 2×1
    8.9716
    13.8470
~~~

~~~
TsumEffModKda1 = TAsumFun(a, Pobj, PobjE(1), PsubBend, PsubMin, Rsum(1,:)’, RsumNormMax(1));
TsumEffModKda2 = TAsumFun(a, Pobj, PobjE(2), PsubBend, PsubMin, Rsum(2,:)’, RsumNormMax(2));
% n.b., numel(Rsum(x)) = numel(Pobj). Rsum(x) has been transposed. Thus, TsumKda(x) is a square matrix.
% Plot the individual mountains for the two values of Kda
title_str1 = strcat({‘Kda = ‘}, sprintf(‘%2.2f’, Kda(1)));
MTNsumEffModKda1 = plot_MTN(Felec, Pobj, TsumEffModKda1, ‘off’, ‘MTNsumEffModKda1’, title_str1, …
    graphs2files, FigDir);
title_str2 = strcat({‘Kda = ‘}, sprintf(‘%2.2f’, Kda(2)));
MTNsumEffModKda2 = plot_MTN(Felec, Pobj, TsumEffModKda2, ‘off’, ‘MTNsumEffModKda2’, title_str2, …
    graphs2files, FigDir);
% Plot the pair of mountains for the two values of Kda
dual_sum_plot = dual_subplot(MTNsumEffModKda1, MTNsumEffModKda2, ‘MTNsumEffMod_Kda1_Kda2’,…
    graphs2files,FigDir);
if show_graphics
    dual_sum_plot.Visible = ‘on’;
end
~~~

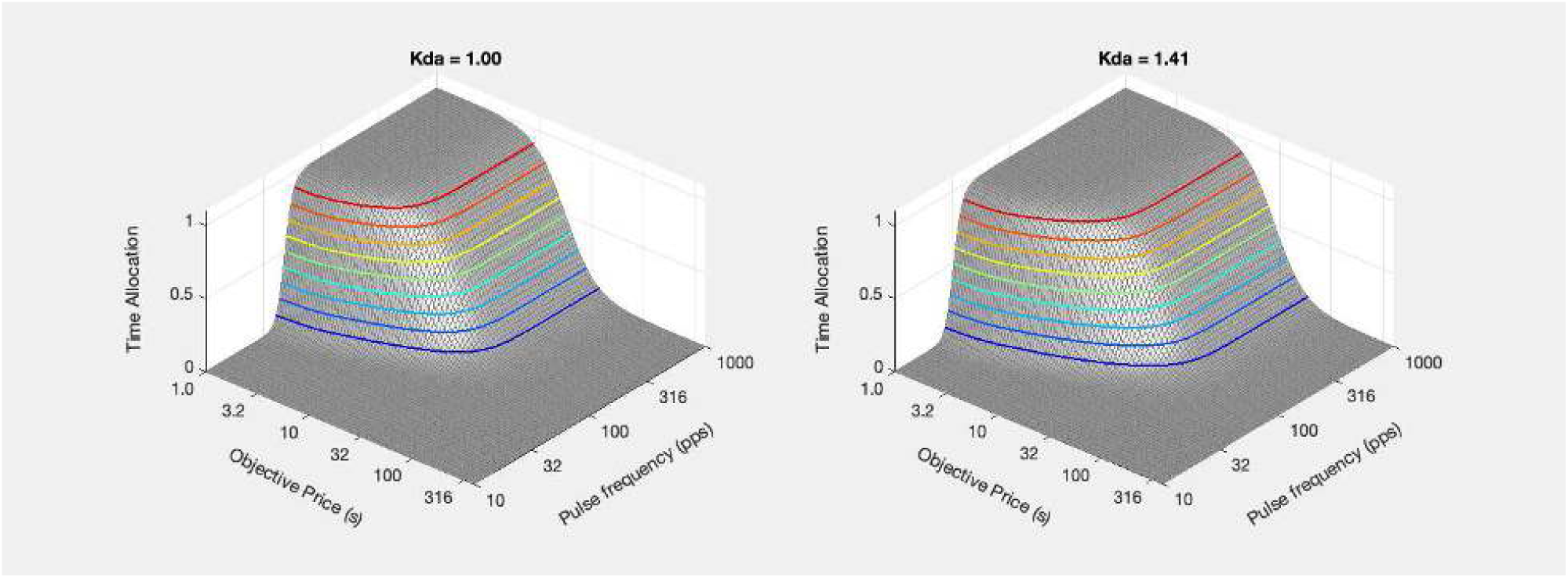

~~~
% Use value of FsumHMstar calculated above;
PsubE = PsubFun(PobjE,PsubBend,PsubMin);
PsubEstar = PsubE ./ RsumNormMax;
PobjEstar = PsubBsFun(PsubEstar,PsubBend,PsubMin);
ContkDA1sumEffMod = plot_contour(Felec, Pobj, TsumEffModKda1, PobjE(1), FsumHM(1), ‘off’, …
     ‘ContkDA1sumEffMod’, title_str1, …
     strcat({‘Kda = ‘}, num2str(Kda(1))), graphs2files, FigDir);
ContkDA2sumEffMod = plot_contour(Felec, Pobj, TsumEffModKda2, PobjE(2), FsumHM(2), ‘off’, …
     ‘ContkDA2sumEffMod’, title_str2, …
     strcat({‘Kda = ‘}, num2str(Kda(2))), graphs2files, FigDir);
bg_kDA1vskDA2sumEffMod = plot_bgStar(FsumHM(1), FsumHM(2), FsumHMstar(1), FsumHMstar(2),…
     PobjE(1), PobjE(2), PobjEstar(1), PobjEstar(2),…
     ‘off’, ‘kDA1vskDA2sumEffMod_bg’, graphs2files, FigDir, −0.3, 0.3);
% Set a scale for the y-axis of this bar graph that ressembles the scales for the others
bg_root = ‘bg_kDA1vskDA2sumEffMod’;
quad_kDA1vskDA2sumEffMod = quad_subplot(ContkDA1sumEffMod, ContkDA2sumEffMod, bg_kDA1vskDA2sumEffMod, …
     ‘quad_kDA1vskDA2sumEffMod’, bg_root, graphs2files, FigDir);
if show_graphics
     quad_kDA1vskDA2sumEffMod.Visible = ‘on’;
end
~~~

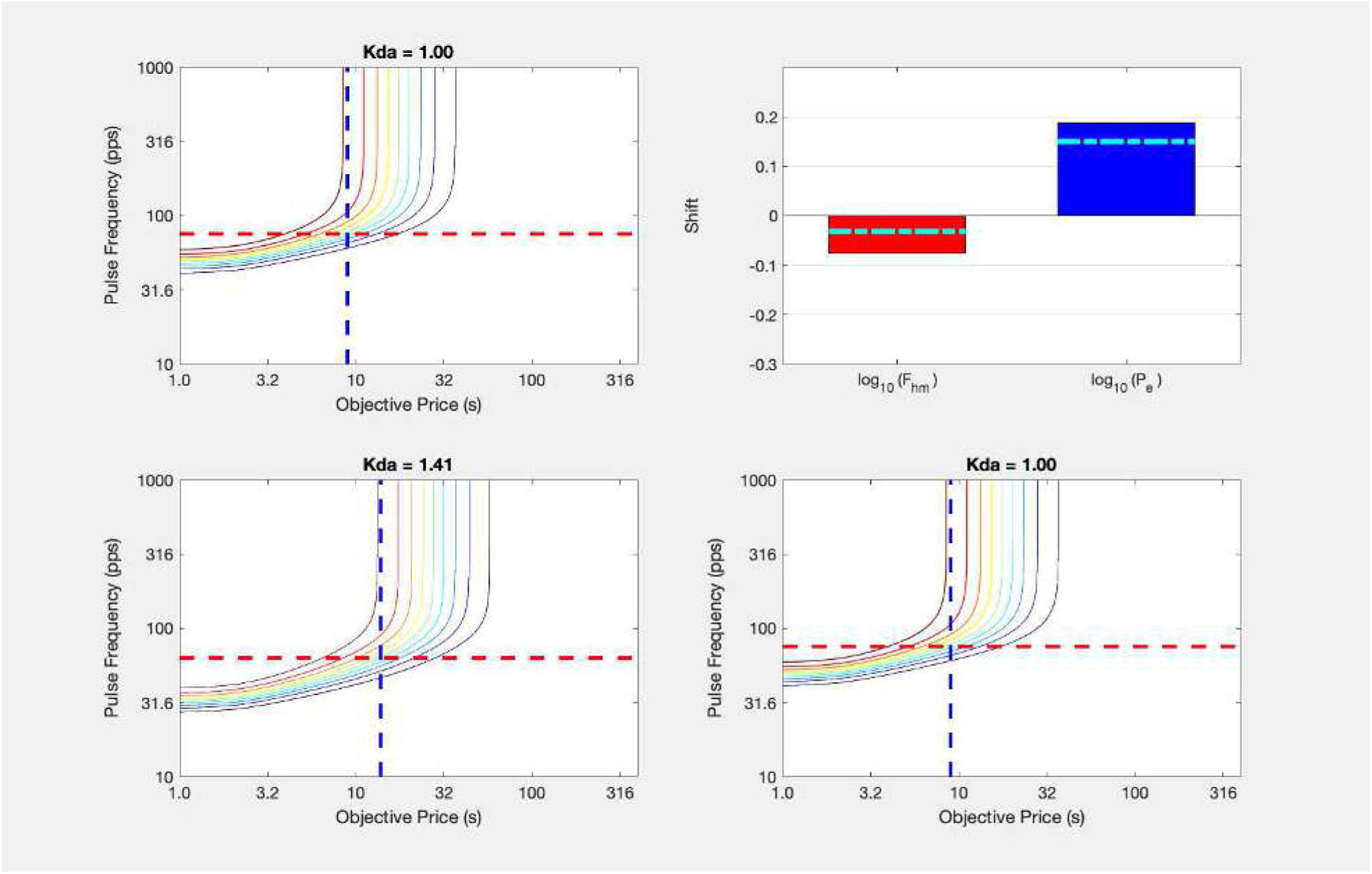

~~~
fig_nam = strcat(“quad_kDA1vskDA2sumEffMod”,num2str(FoptEquiv));
[fig_num, fig_tab] = add_fig(fig_num, fig_tab, …
    fig_nam, “Dual shifts of the reward mountain induced by dopamine-transporter blockade”);
~~~

`This is figure #60: (quad_kDA1vskDA2sumEffMod80)`

~~~
close all;
vars2save = who;
ws_name = fullfile(pwd, ‘WS_convergence_vars80.mat’);
save_ws;
~~~

~~~
Saving workspace to /Users/shizgal/Work/Research/papers/In_Progress/Opto_GBR2/Simulations/WS_convergence_vars80.mat
Copying backup to /Users/shizgal/Work/Research/papers/In_Progress/Opto_GBR2/Simulations/WS_convergence_vars80_bk191201.mat
~~~

The small shift along the pulse-frequency axis can be eliminated by reducing the MFB drive. We will illustrate this by means of reward-growth graphs.

~~~
% Rupper is unchanged - no need to regenerate it
% Regenerate Rlower using weaker MFB drive
FoptEquiv = 40; % Value was 80 in above simulations
FmfbDAdrive = FmfbDAdriveFun(FmfbDAbend, Felec, FFmfbDAmax, FoptEquiv, FmfbDAro);
FdaHMkDA = FpulseHMfun(C, D, FdaBend, FPmax, FdaRO, NnDA, RhoPiDA, Kda) % result is a two-element vector
~~~

~~~
FdaHMkDA = 1×2
    37.6342 25.1515
~~~

~~~
KdaMat = repmat(Kda’,1,numF);
FdaHMmat = repmat(FdaHMkDA’,1,numF);
Rlower = fRbsr(FmfbDAdrive, FdaBend, FdaHMmat, FdaRO, gDA, KrgLower);
Rsum = Rupper + Rlower;
[FsumHM, RsumMax] = find_FhmSum(FelecMat, Rsum)
~~~

~~~
FsumHM = 2×1
    84.1920
    84.2963
RsumMax = 2×1
    1.3105
    1.7715
~~~

~~~
RmatVeh = [Rupper(1,:);Rlower(1,:);Rsum(1,:)];
RmatDrg = [Rupper(2,:);Rlower(2,:);Rsum(2,:)];
pnam = “1:up 2:lo 3:sum”;
pVec = [1,2,3];
fnam = strcat(“Sum”,num2str(FoptEquiv),”Veh”);
TitleStr = strcat(“FoptEquiv=“,num2str(FoptEquiv),”; vehicle”);
RGsum_Rveh = plot_RG(FelecMatPlot’,RmatVeh’,pnam,pVec,fnam,TitleStr,’lin’);
axh = findall(RGsum_Rveh, ‘Type’, ‘Axes’);
axh.XLabel.String = ‘Electrical pulse frequency’;
lh = findall(RGsum_Rveh, ‘Type’, ‘Line’);
lcolor = {‘m’,’g’,’c’};
for j=1:3
    lh(j).Color = lcolor{j};
end
if show_graphics
    RGsum_Rveh.Visible = ‘on’;
end
~~~

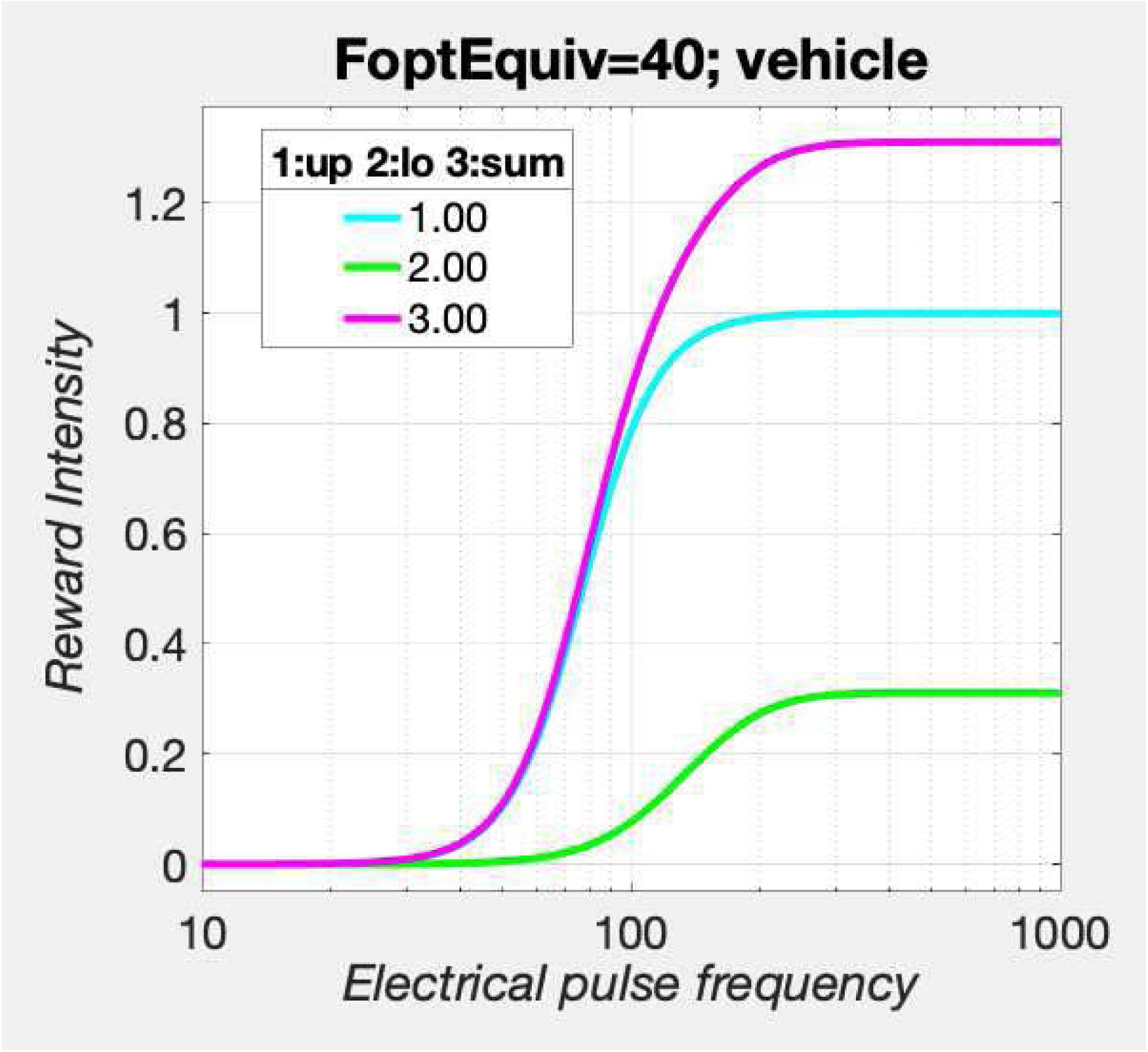

~~~
fig_nam = strcat(“RGsum_Veh_FoptEquiv”,num2str(FoptEquiv));
[fig_num, fig_tab] = add_fig(fig_num, fig_tab, fig_nam, “Reward summation in the convergence model”);
~~~

`This is figure #61: (RGsum_Veh_FoptEquiv40)`

~~~
pnam = “1:up 2:lo 3:sum”;
pVec = [1,2,3];
fnam = strcat(“Sum”,num2str(FoptEquiv),”Drg”);
TitleStr = strcat(“FoptEquiv=“,num2str(FoptEquiv),”; drug”);
RGsum_Rdrg = plot_RG(FelecMatPlot’,RmatDrg’,pnam,pVec,fnam,TitleStr,’lin’);
axh = findall(RGsum_Rdrg, ‘Type’, ‘Axes’);
axh.XLabel.String = ‘Electrical pulse frequency’;
lh = findall(RGsum_Rdrg, ‘Type’, ‘Line’);
lcolor = {‘m’,’g’,’c’};
for j=1:3
    lh(j).Color = lcolor{j};
    lh(j).LineStyle = ‘--’;
end
if show_graphics
    RGsum_Rdrg.Visible = ‘on’;
end
~~~

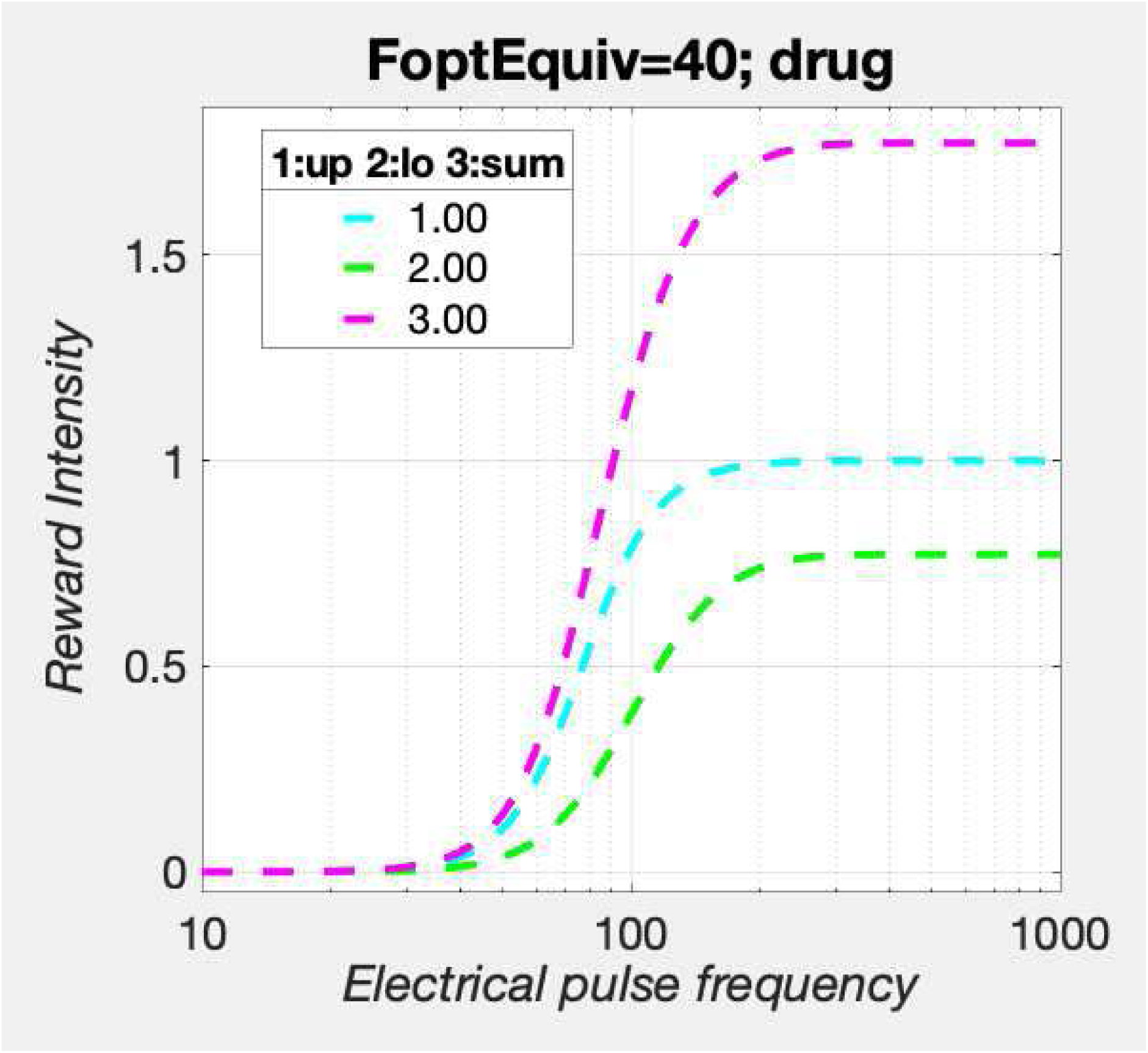

~~~
fig_nam = strcat(“RGsum_Drg_FoptEquiv”,num2str(FoptEquiv));
[fig_num, fig_tab] = add_fig(fig_num, fig_tab, fig_nam, “Reward summation in the convergence model”);
~~~

`This is figure #62: (RGsum_Drg_FoptEquiv40)`

~~~
pnam = “Kda”;
pVec = [Kda(1),Kda(2)];
fnam = strcat(“Sum”,num2str(FoptEquiv),”VehDrg”);
TitleStr = strcat(“FoptEquiv=“,num2str(FoptEquiv),”; sum”);
RGsum_Rvehdrg = plot_RG(FelecMat’,Rsum’,pnam,pVec,fnam,TitleStr,’lin’);
axh = findall(RGsum_Rvehdrg, ‘Type’, ‘Axes’);
axh.XLabel.String = ‘Electrical pulse frequency’;
lh = findall(RGsum_Rvehdrg, ‘Type’, ‘Line’);
lh(1).LineStyle = ‘--’;
for j=1:2
    lh(j).Color = ‘m’;
end
lgndh = findall(RGsum_Rvehdrg, ‘Type’, ‘Legend’);
lgndh.Title.String = “condition”;
lgndh.String = {‘vehicle’,’drug’};
if show_graphics
    RGsum_Rvehdrg.Visible = ‘on’;
end
~~~

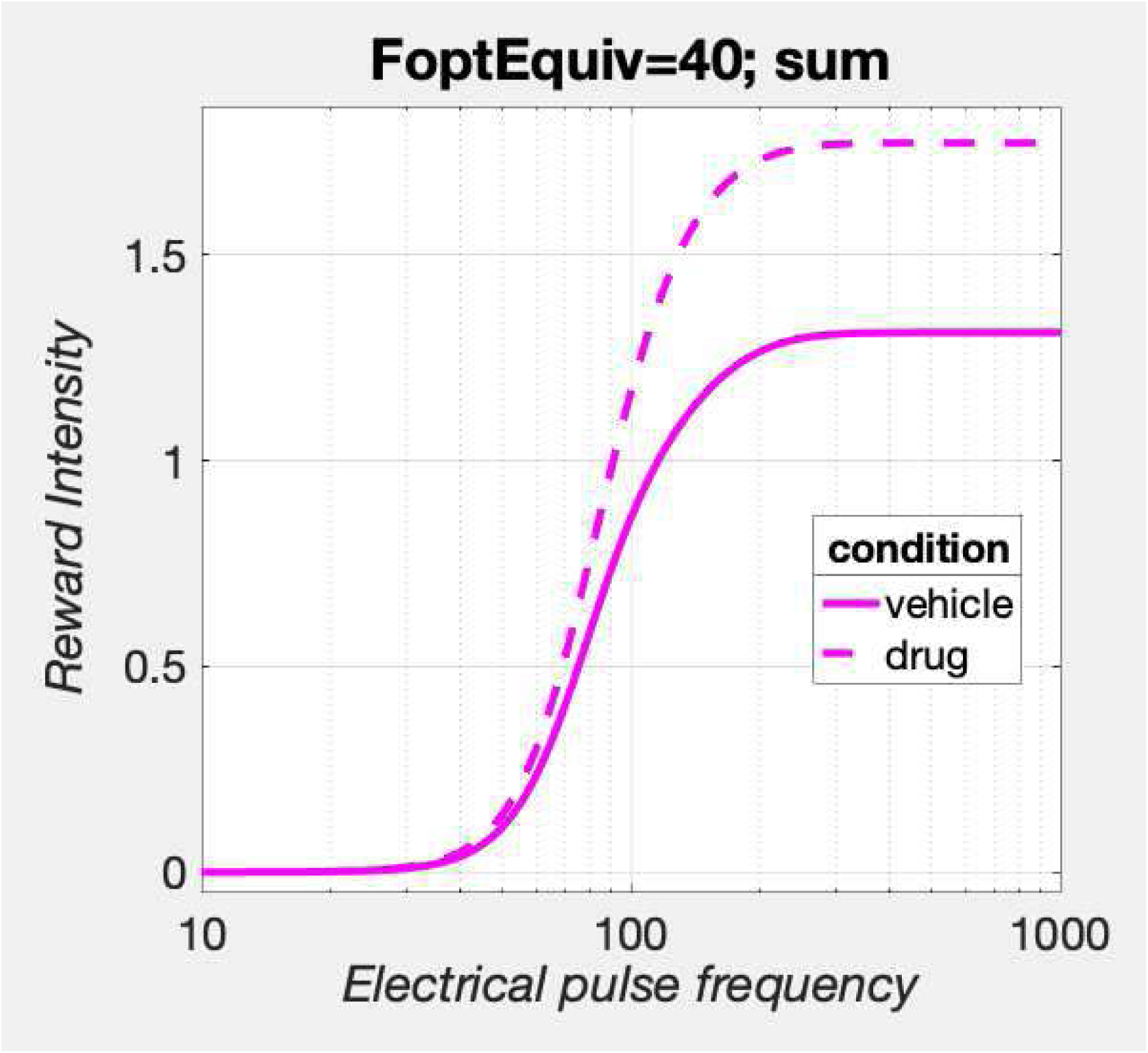

~~~
fig_nam = strcat(“RGsum_VehDrg_FoptEquiv”,num2str(FoptEquiv));
[fig_num, fig_tab] = add_fig(fig_num, fig_tab, fig_nam, “Drug modulation of reward summation in the convergence model”);
~~~

`This is figure #63: (RGsum_VehDrg_FoptEquiv40)`

At this weaker value of MFB drive, the reward-growth curve is shifted to the right. At this value of *K*_*da*_ (∼median for the current study), the drug cannot shift the reward-growth curve far enough to the left to displace the summated curve leftwards. Thus, we learn from this that shifts along the pulse-frequency axis in the convergence model depend on the strength of the MFB drive.

~~~
close all;
FPmax = 1000;
FhmUpperVec = repmat(FhmUpper,1,2); % Upper limb output is independent of Kda
RupperNormMax = fRbsrNorm(FPmax, FelecBend, FhmUpperVec, FelecRO, gElec)’;
RlowerNormMax = fRbsrNorm(FoptEquiv, FdaBend, FdaHMkDA, FdaRO, gDA)’;
RupperMax = max(Rupper, [], 2);
RlowerMax = max(Rlower, [], 2);
RupperRatio = RupperMax ./ (RlowerMax + RupperMax)
RupperRatio = 2×1
   0.7627
   0.5643
RlowerRatio = RlowerMax ./ (RlowerMax + RupperMax)
RlowerRatio = 2×1
   0.2373
   0.4357
RsumNormMax = (RupperRatio .* RupperNormMax) + (RlowerRatio .* RlowerNormMax); % Weighted average
dotPhiObj = 1;
Kaa = 1;
Kec = 1;
Krg = 1;
dotRaa = 0.1;
pObj = 1;
PsubBend = 0.5;
PsubMin = 1.82;
PobjE = PobjEfun(dotPhiObj, Kaa, Kec, Krg, dotRaa, pObj, PsubBend, PsubMin, RsumNormMax, KeffMod)
PobjE = 2×1
   8.3618
   12.7179
a = 3;
numP = numF;
Pobj = logspace(0,3,numP); % row variable
TsumEffModLoFoeqKda1 = TAsumFun(a, Pobj, PobjE(1), PsubBend, PsubMin, Rsum(1,:)’, RsumNormMax(1));
TsumEffModLoFoeqKda2 = TAsumFun(a, Pobj, PobjE(2), PsubBend, PsubMin, Rsum(2,:)’, RsumNormMax(2));
% n.b., numel(Rsum(x)) = numel(Pobj). Rsum(x) has been transposed. Thus, TsumKda(x) is a square matrix.
% Plot the individual mountains for the two values of Kda
title_str1 = strcat({‘Kda = ‘}, sprintf(‘%2.2f’, Kda(1)));
MTNsumEffModLoFoeqKda1 = plot_MTN(Felec, Pobj, TsumEffModLoFoeqKda1, ‘off’, ‘MTNsumEffModLoFoeqKda1’, title_str1, …
    graphs2files, FigDir);
title_str2 = strcat({‘Kda = ‘}, sprintf(‘%2.2f’, Kda(2)));
MTNsumEffModKLoFoeqda2 = plot_MTN(Felec, Pobj, TsumEffModLoFoeqKda2, ‘off’, ‘MTNsumEffModLoFoeqKda2’, title_str2, …
    graphs2files, FigDir);
% Plot the pair of mountains for the two values of Kda
dual_sum_plot = dual_subplot(MTNsumEffModLoFoeqKda1, MTNsumEffModKLoFoeqda2, ‘MTNsumEffModLoFoeq_Kda1_Kda2’,…
    graphs2files,FigDir);
if show_graphics
    dual_sum_plot.Visible = ‘on’;
end
~~~

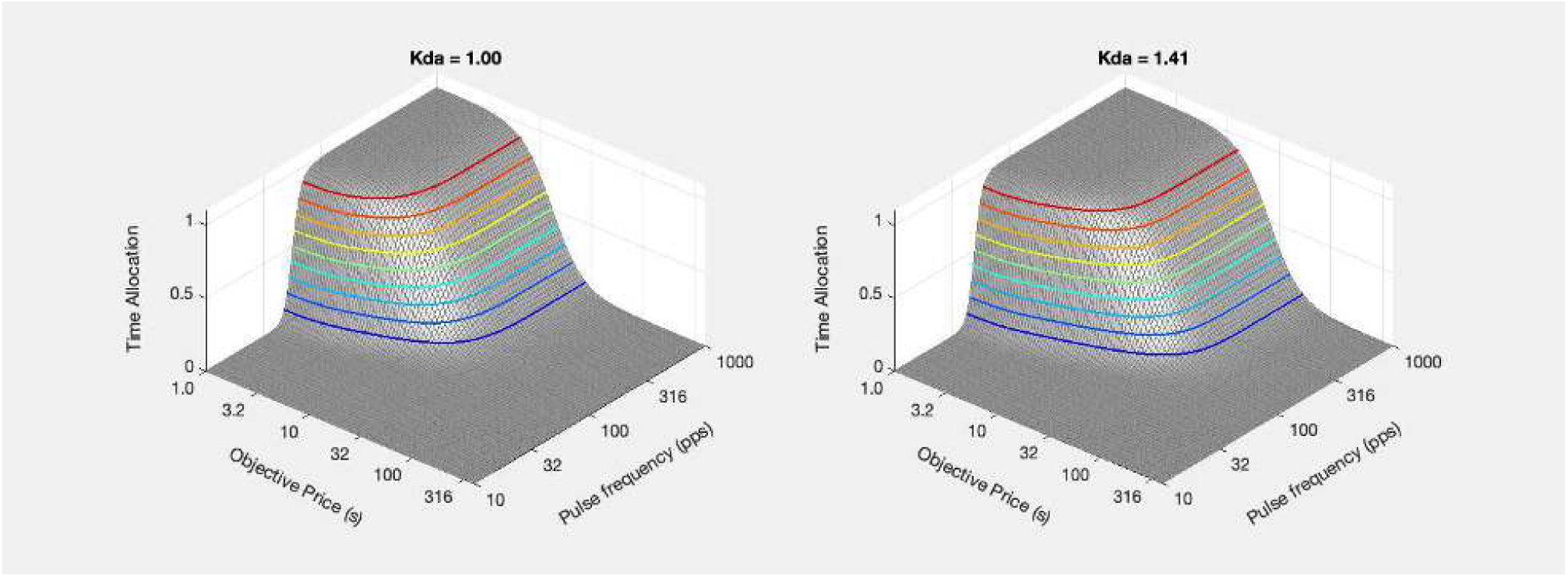

~~~
% Estimate the weighted average of the firing frequencies that produce half-maximal reward intensity in the
% upper and lower limbs
FhmUpperStar = FilterFun(FhmUpper,FelecBend,FelecRO);
FhmLowerDrive = FmfbDAdriveFun(FmfbDAbend, FsumHM, FFmfbDAmax, FoptEquiv, FmfbDAro);
FFhmLowerStar = FilterFun(FhmLowerDrive,FdaBend, FdaRO);
FsumHMstar = FsumHM .* …
     ((RupperRatio .* (FhmUpperStar ./ FsumHM)) + (RlowerRatio .* (FFhmLowerStar ./ FhmLowerDrive)));
PsubE = PsubFun(PobjE,PsubBend,PsubMin);
PsubEstar = PsubE ./ RsumNormMax;
PobjEstar = PsubBsFun(PsubEstar,PsubBend,PsubMin);
ContkDA1sumEffMod = plot_contour(Felec, Pobj, TsumEffModLoFoeqKda1, PobjE(1), FsumHM(1), ‘off’, …
     ‘ContkDA1sumEffModLoFoeq’, title_str1, …
     strcat({‘Kda = ‘}, num2str(Kda(1))), graphs2files, FigDir);
ContkDA2sumEffMod = plot_contour(Felec, Pobj, TsumEffModLoFoeqKda2, PobjE(2), FsumHM(2), ‘off’, …
     ‘ContkDA2sumEffModLoFoeq’, title_str2, …
     strcat({‘Kda = ‘}, num2str(Kda(2))), graphs2files, FigDir);
bg_kDA1vskDA2sumEffMod = plot_bgStar(FsumHM(1), FsumHM(2), FsumHMstar(1), FsumHMstar(2),…
    PobjE(1), PobjE(2), PobjEstar(1), PobjEstar(2),…
    ‘off’, ‘kDA1vskDA2sumEffMod_bg’, graphs2files, FigDir, −0.3, 0.3);
% Set a scale for the y-axis of this bar graph that ressembles the scales for the others
bg_root = ‘bg_kDA1vskDA2sumEffMod’;
quad_kDA1vskDA2sumEffMod = quad_subplot(ContkDA1sumEffMod, ContkDA2sumEffMod, bg_kDA1vskDA2sumEffMod, …
   ‘quad_kDA1vskDA2sumEffMod’, bg_root, graphs2files, FigDir);
if show_graphics
   quad_kDA1vskDA2sumEffMod.Visible = ‘on’;
end
~~~

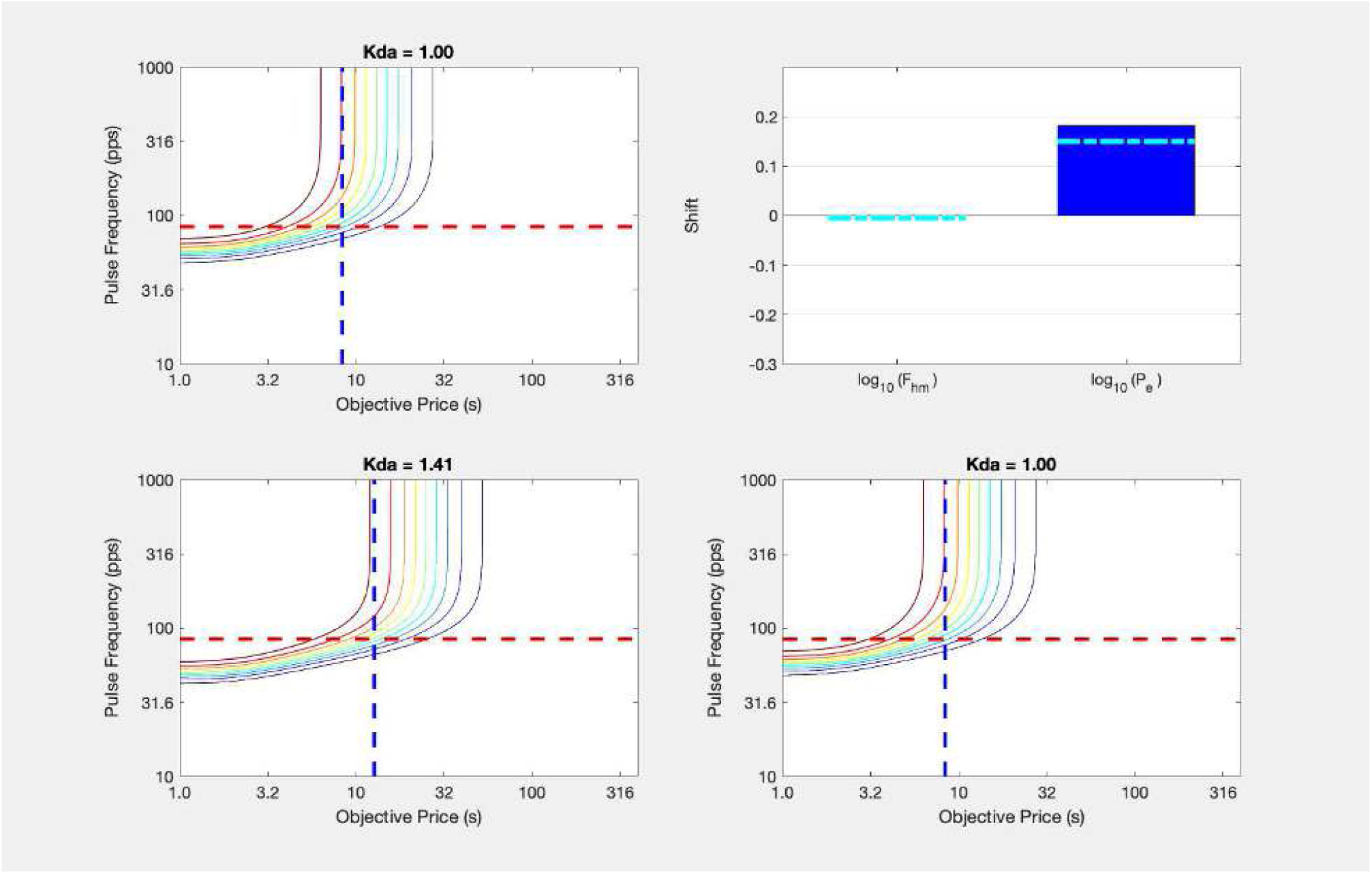

~~~
fig_nam = strcat(“quad_kDA1vskDA2sumEffMod”,num2str(FoptEquiv));
[fig_num, fig_tab] = add_fig(fig_num, fig_tab, …
    fig_nam, “Dual shifts of the reward mountain induced by dopamine-transporter blockade”);
~~~

`This is figure #64: (quad_kDA1vskDA2sumEffMod40)`

~~~
close all;
vars2save = who;
ws_name = fullfile(pwd, ‘WS_convergence_vars.mat’);
save_ws;
~~~

Saving workspace to /Users/shizgal/Work/Research/papers/In_Progress/Opto_GBR2/Simulations/WS_convergence_vars.mat Copying backup to /Users/shizgal/Work/Research/papers/In_Progress/Opto_GBR2/Simulations/WS_convergence_vars_bk191201.mat

`Elapsed time is 42.313713 seconds.`

~~~
tic;
~~~

### Remarks concerning the convergence model

The results in

~~~
disp(string({strcat({‘Figure ‘},num2str(fig_tab.Number(fig_tab.Name==‘quad_kDA1vskDA2sumEffMod’)))}));
Figure
~~~

above resemble those obtained in the eICSS study in which the reward mountain was measured under the influence of GBR-12901 (Hernandez et al., 2012). Thus, given moderate MFB drive and 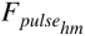 values compatible with those obtained in the present study (adjusted for a train duration of 0.5 s), the convergence model can generate outputs that match our earlier eICSS results.

The dependence of the output of the convergence model on experimental parameters is of interest. Although failure to observe shifts along the pulse-frequency axis was the most common result of the experiments in which the eICSS reward-mountain was measured under the influence of drugs that alter dopaminergic neurotransmission (27 of 32 cases reported by Hernandez et al., 2010, Trujillo-Pisanty et al., 2011, Hernandez et al., 2012, and Trujillo-Pisanty et al., 2014), it is not the only result. Reward mountains obtained from three subjects in the cocaine study (Hernandez et al., 2010) showed fairly substantial, reliable, shifts along the pulse-frequency axis. (The shift was marginally reliable in a fourth subject when tested initially but disappeared upon re-test.) Although no subject in the GBR-12909 (Hernandez et al., 2012) or pimozide (Trujillo-Pisanty et al., 2014) studies showed such shifts, one subject in the AM-251 study (Trujillo-Pisanty et al., 2011) did. Could variation in the strength of MFB drive on the dopamine neurons and in the individual 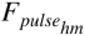 values for the dopamine pathway explain these findings?

Please see the main body of the manuscript for more detailed discussion.

## Sort tables and write to files

### Formatted equations

~~~
if tabdir
   writetable(sortrows(eqn_tab,2),…
     fullfile(tabdir,strcat(‘eqn_tab_v’,num2str(version),’.csv’)),…
     ‘WriteVariableNames’, true);
end
~~~

### Figures

~~~
if tabdir
   writetable(sortrows(fig_tab,2),…
   fullfile(tabdir,strcat(‘fig_tab_v’,num2str(version),’.csv’)),…
   ‘WriteVariableNames’, true);
end
~~~

### Functions

~~~
if tabdir
    writetable(sortrows(fun_tab,2),…
    fullfile(tabdir,strcat(‘fun_tab_v’,num2str(version),’.csv’)),…
    ‘WriteVariableNames’, true);
end
~~~

### Symbols

~~~
if tabdir
   writetable(sortrows(sym_tab,2),…
   fullfile(tabdir,strcat(‘sym_tab_v’,num2str(version),’.csv’)),…
   ‘WriteVariableNames’, true);
end
~~~

### Save workspace

~~~
if saveWS
   ws_name = fullfile(pwd, strcat(‘WS_GBR_eICSS_oICSS_v’,num2str(version),’.mat’));
   save_ws;
end
~~~

Saving workspace to /Users/shizgal/Work/Research/papers/In_Progress/Opto_GBR2/Simulations/WS_GBR_eICSS_oICSS_v10.mat Copying backup to /Users/shizgal/Work/Research/papers/In_Progress/Opto_GBR2/Simulations/WS_GBR_eICSS_oICSS_v10_bk191201.mat

Last revised on:

~~~
datetime(‘now’)
ans = *datetime*
   01-Dec-2019 18:52:43
toc % elapsed time for this section
~~~

`Elapsed time is 1.230854 seconds.`

~~~
toc(t_start) % elapsed time for the entire script
~~~

`Elapsed time is 226.200086 seconds.`

### Tools for this live script

~~~
function init_all
  global eqn_num eqn_tab fig_num fig_tab fun_num fun_tab sym_num sym_tab
  eqn_num = 0; % initialize counter
  eqn_nam = ““; % initialize equation-name string array
  eqn_desc = ““; % initialize equation-description string array;
  eqn_tab = table(eqn_num, eqn_nam, eqn_desc,’VariableNames’, {‘Number’; ‘Name’; ‘Description’});
  eqn_tab(1,:)=[]; % clear the table
  disp(“The equation table has been initialized.”);
  fig_num = 0; % initialize counter
  fig_nam = ““; % initialize figure-name string array
  fig_desc = ““; % initialize figure-description string array;
  fig_tab = table(fig_num, fig_nam, fig_desc,’VariableNames’, {‘Number’; ‘Name’; ‘Description’});
  fig_tab(1,:)=[]; % clear the table
  disp(“The figure table has been initialized.”);
  fun_num = 0; % initialize counter
  fun_nam = ““; % initialize function-name string array
  fun_args = ““; %initialize function-argument string array
  fun_desc = ““; % initialize function-description string array;
  fun_tab = table(fun_num, fun_nam, fun_args, fun_desc, …
      ‘VariableNames’, {‘Number’; ‘Name’; ‘Arguments’; ‘Description’});
  fun_tab(1,:)=[]; % clear the table
  disp(“The function table has been initialized.”);
  sym_num = 0; % initialize counter
  sym_nam = ““; % initialize symbol-name string array
  sym_desc = ““; % initialize symbol-description string array;
  sym_tab = table(sym_num, sym_nam, sym_desc,’VariableNames’, {‘Number’; ‘Name’; ‘Description’});
  sym_tab(1,:)=[]; % clear the table
  disp(“The symbol table has been initialized.”);
  [fun_num, fun_tab] = add_fun(fun_num, fun_tab, “init_all”, ““, “initialize tables”);
  [fun_num, fun_tab] = add_fun(fun_num, fun_tab, “add_eqn”, “eqn_num, eqn_tab, EqnNam, EqnDesc”,…
    “add equation to table”);
  [fun_num, fun_tab] = add_fun(fun_num, fun_tab, “add_fig”, “fig_num, fig_tab, FigNam, FigDesc”,…
    “add figure to table”);
  [fun_num, fun_tab] = add_fun(fun_num, fun_tab, “add_fun”, “fun_num, fun_tab, FunNam, FunArgs, FunDesc”,…
    “add local function to table”);
  [fun_num, fun_tab] = add_fun(fun_num, fun_tab, “add_sym”, “sym_num, sym_tab, SymNam, SymDesc”,…
    “add symbol to table”);
  [fun_num, fun_tab] = add_fun(fun_num, fun_tab, “FilterFun”, “F, Fbend, Fro”,…
    “Frequency-following function”);
  [fun_num, fun_tab] = add_fun(fun_num, fun_tab, “FilterFunBS”, “Fbend, Fro, Frate, FPmax”,…
    “Back-solution of frequency-following function”);
  [fun_num, fun_tab] = add_fun(fun_num, fun_tab, “LogistNormFun”, “exponent, input, location”,…
    “Rising logistic function”);
  [fun_num, fun_tab] = add_fun(fun_num, fun_tab, “LogistNormBsFun”, “exponent, location, output”,…
    “Back-solution of logistic function to return the input”);
  [fun_num, fun_tab] = add_fun(fun_num, fun_tab, “LogistNormBsLocFun”, “exponent, input, output”,…
    “Back-solution of logistic function to return the location parameter”);
  [fun_num, fun_tab] = add_fun(fun_num, fun_tab, “PsubFun”, “Pobj, PsubBend, PsubMin”,…
    “Subjective-price function”);
  [fun_num, fun_tab] = add_fun(fun_num, fun_tab, “PsubBsFun”, “Psub, PsubBend, PsubMin”,…
    “Back-solution of subjective-price function”);
  [fun_num, fun_tab] = add_fun(fun_num, fun_tab, “ScalarDivFun”, “dividend, divisor”,…
    “Scalar-division function”);
  [fun_num, fun_tab] = add_fun(fun_num, fun_tab, “ScalarDivBsFun”, “divisor, quotient”,…
    “Back-solution of scalar-division function”);
  [fun_num, fun_tab] = add_fun(fun_num, fun_tab, “ScalarMultFun”, “multiplicand, multiplier”,…
    “Scalar-multiplication function”);
  [fun_num, fun_tab] = add_fun(fun_num, fun_tab, “ScalarMultBsFun”, “multiplier, product”,…
    “Back-solution of scalar-multiplication function”);
  [fun_num, fun_tab] = add_fun(fun_num, fun_tab, “PsubEfun”, “Ceff, Rmax, Ue”,…
    “Function to compute subjective equivalent of the objective-price location parameter”);
  [fun_num, fun_tab] = add_fun(fun_num, fun_tab, “PobjEfun”, “Ceff, PsubBend, PsubMin, Rmax, Ue”,…
     “Function to compute objective-price location parameter”);
end % end of init_all function
function [eqn_num, eqn_tab] = add_eqn(eqn_num, eqn_tab, EqnNam, EqnDesc)
   if any(strcmp(EqnNam,eqn_tab.Name))
     disp(strcat({‘Equation ‘}, EqnNam, {‘has already been entered.’}));
     return;
     else
        eqn_num = eqn_num+1;
        eqn_tab(eqn_num,:) = {eqn_num, …
          string(EqnNam), …
          string(EqnDesc)};
      disp(strcat({‘This is equation #’}, num2str(eqn_tab.Number(height(eqn_tab))), …
     {‘: (‘}, eqn_tab.Name(height(eqn_tab)), {‘)’}));
   end
end
function [fig_num, fig_tab] = add_fig(fig_num, fig_tab, FigNam, FigDesc)
   if any(strcmp(FigNam,fig_tab.Name))
      disp(strcat({‘Figure ‘}, FigNam, {‘has already been entered.’}));
   return;
   else
      fig_num = fig_num+1;
      fig_tab(fig_num,:) = {fig_num, …
          string(FigNam), …
          string(FigDesc)};
      disp(strcat({‘This is figure #’}, num2str(fig_tab.Number(height(fig_tab))), …
        {‘: (‘}, fig_tab.Name(height(fig_tab)), {‘)’}));
     end
end
function [fun_num, fun_tab] = add_fun(fun_num, fun_tab, FunNam, FunArgs, FunDesc)
    if any(strcmp(FunNam,fun_tab.Name))
        disp(strcat({‘Function ‘}, FunNam, {‘has already been entered.’}));
    return;
  else
   fun_num = fun_num+1;
   fun_tab(fun_num,:) = {fun_num, …
      string(FunNam), …
      string(FunArgs), …
      string(FunDesc)};
   disp(strcat({‘This is function #’}, num2str(fun_tab.Number(height(fun_tab))), …
      {‘: (‘}, fun_tab.Name(height(fun_tab)), {‘)’}));
  end
end
function [sym_num, sym_tab] = add_sym(sym_num, sym_tab, SymNam, SymDesc)
    if any(strcmp(SymNam,sym_tab.Name))
    disp(strcat({‘Symbol ‘}, SymNam, {‘has already been entered.’}));
   return;
  else
    sym_num = sym_num+1;
    sym_tab(sym_num,:) = {sym_num, …
      string(SymNam), …
      string(SymDesc)};
%   disp(strcat({‘This is symbol #’}, num2str(sym_tab.Number(height(sym_tab))), …
%     {‘: (‘}, sym_tab.Name(height(sym_tab)), {‘)’}));
   end
end
function FigDir = set_figdir(figdirnam, graphs2files)
    global Figdir
    FigDir = fullfile(pwd,figdirnam);
    if graphs2files
          if ∼exist(FigDir, ‘dir’)
              mkdir(FigDir);
          end
     end
end
function ImpFigDir = set_impfigdir(impfigdirnam)
  ImpFigDir = fullfile(pwd,impfigdirnam);
  if ∼exist(ImpFigDir, ‘dir’)
    disp(strcat({‘Exiting because Imported-files directory ‘}, impfigdirnam, {‘does not exist.’}));
      return
   end
end
function tabdir = set_tabdir(tabdirnam, tabs2files)
     tabdir = fullfile(pwd,tabdirnam);
     if tabs2files
        if ∼exist(tabdir, ‘dir’)
            mkdir(tabdir);
           end
     end
end
~~~

### Functional building blocks for the simulations

~~~
function Frate = FilterFun(F, Fbend, Fro) % Frequency-following function
  Frate = Fbend.*(log(1+exp(Fro./Fbend))-log(1+exp((Fro-F)./Fbend)));
end
% see:
% Breton, Y.-A., Mullett, A., Conover, K., & Shizgal, P. (2013).
% Validation and extension of the reward-mountain model.
% Frontiers in Behavioral Neuroscience, 7, 125. https://doi.org/10.3389/fnbeh.2013.00125
% Solomon, R. B., Trujillo-Pisanty, I., Conover, K., & Shizgal, P. (2015).
% Psychophysical inference of frequency-following fidelity in the neural substrate for brain stimulation reward.
% Brain Research, 292, 327–341. https://doi.org/10.1016/j.bbr.2015.06.008
% Last revised by Peter Shizgal on 2019-09-05 09:57
function Fpulse = FilterFunBS(Fbend, Fro, FF, FPmax) % Back-solution of frequency-following function
   % Estimate the maximum achievable firing rate, given FPmax, Fbend, and Fro
   FFmax = FilterFun(FPmax, Fbend, Fro);
   FpulseRI = Fro - Fbend.*log(exp(-FF/Fbend).*(exp(Fro/Fbend) + 1) - 1);
   % FpulseRI is a complex number. When FF < FFmax, the imaginary part = zero. When Fpulse RI = FFmax, then Fpulse = Inf.
   % We retain the real component when FF < FFmax & >= 0. returning NaN when FF > FFmax | FF < 0 and Inf when FF = FFmax.
   sz = size(FF);
   Fpulse = zeros(sz); % pre-allocate
   neg = FF < 0;
   below = (FF < FFmax & FF >= 0);
   eq = FF == FFmax;
   above = FF > FFmax;
   Fpulse(neg) = NaN;
   Fpulse(below) = real(FpulseRI(below)); % Complex portion, if it exists, equals zero
   Fpulse(eq) = Inf;
   Fpulse(above) = NaN;
end
% This back-solution can be verified from the following code snippet, which requires the
% Matlab Symbolic-Math Toolbox. It is commented out here to allow this Live Script to run
% on systems that lack the Symbolic-Math toolbox.
% syms FdaRate FdaBend FdaRO Fopt positive;
% simplify(solve(FdaRate == FdaBend * (log(1+exp(FdaRO/FdaBend)) - (log(1+exp((FdaRO-Fopt)/FdaBend)))),Fopt))
% Fopt = FdaRO - FdaBend*log(exp(-FdaRate/FdaBend)*(exp(FdaRO/FdaBend) + 1) - 1)
%%
% This function returns imaginary numbers. The imaginary portion equals zero provided that Frate (the firing rate)
% is less than its maximum achievable value. That maximum is estimated from the forward solution, using FPmax
% as the input.
%%
% see:
% Breton, Y.-A., Mullett, A., Conover, K., & Shizgal, P. (2013).
% Validation and extension of the reward-mountain model.
% Frontiers in Behavioral Neuroscience, 7, 125. https://doi.org/10.3389/fnbeh.2013.00125
% Solomon, R. B., Trujillo-Pisanty, I., Conover, K., & Shizgal, P. (2015).
% Psychophysical inference of frequency-following fidelity in the neural substrate for brain stimulation reward.
% Behavioural Brain Research, 292, 327–341. https://doi.org/10.1016/j.bbr.2015.06.008
function output = LogistNormFun(exponent, input, location)
    output = …
       ((input .^ exponent) ./…
       ((input .^ exponent) + (location .^ exponent)));
end
% The symbol for the logistic function in the flow diagrams is a rising S-shaped curve in a box.
%
% Revised by Peter Shizgal on 2019-10-29 21:25
% The argument called “output” must be scalar
function input = LogistNormBsFun(exponent, location, output)
% Logistic function back-solved to return the input
   if (output >= 1) || (output <= 0)
     disp(“Illegal value. Call with 0 < output < 1”);
     return
  end
  input = location .* (output./(1-output)).^(1./exponent);
end
% Revised by Peter Shizgal on 2019-10-29 21:25
function location = LogistNormBsLocFun(exponent, input, output)
% The argument called “output” must be scalar
% Logistic function back-solved to return the location parameter
  if output >= 1 || output <= 0
   disp(“Illegal value. Call with 0 < output < 1”);
   return
  end
   location = input .* ((1 - output) ./output).^ (1./exponent);
end
% The symbol for the subjective-price function in the flow diagrams is a
% hockey-stick-shaped curve in a box.
%
% Solomon, R. B., Conover, K., & Shizgal, P. (2017).
% Valuation of opportunity costs by rats working for rewarding electrical brain stimulation.
% PLOS ONE, 12(8), e0182120. https://doi.org/10.1371/journal.pone.0182120
function Psub = PsubFun(Pobj, PsubBend, PsubMin) % Subjective-price function
    Psub = PsubMin + PsubBend .* (log(1 + exp((Pobj - PsubMin)./PsubBend)));
end
% The equation for PsubBsFun will return -Inf when Psub = PsubMin and a complex number when Psub includes one more values less
% than PsubMin. When Psub includes values between PsubMin and Pobj_0 (the value of Pobj at which Psub = 0),
% PsubBsFun returns negative values. The definition of PsubBsFun has thus been couched so as to return real values
% > 0 when Psub > Pobj_0, 0 when Psub = Pobj_0, and NaN otherwise. The reason that 0 is returned when obj_0, 0 when Psub = Pobj_0
% is to ensure consistency between the forward and backward solutions.
% Last revised by Peter Shizgal on 2019-09-08 15:06
function Pobj = PsubBsFun(Psub, PsubBend, PsubMin) % Back-solution of subjective-price function
   Psub_Pobj_0 = PsubMin + PsubBend .* log(1 + exp((-PsubMin)./PsubBend));
   above = Psub > Psub_Pobj_0;
   eq = Psub == Psub_Pobj_0;
   below = Psub < Psub_Pobj_0;
   Pobj = PsubMin + PsubBend .* log(−1 + exp((Psub(above) - PsubMin)./PsubBend));
   % Output will be real because Psub(above) > the value that drives Pobj to zero
   Pobj(eq) = 0;
   Pobj(below) = NaN;
end
% The next four functions are defined for consistency in formatting when composing more complex functions from the
% five building blocks (FilterFun, PsubFun, LogistNormFun, ScalarDivFun, ScalarMultFun) and their
% six back-solutions (FilterFunBS, PsubBsFun, LogistNormBsFun, LogistNormBsLocFun, ScalarDivBsFun, ScalarMultBsFun).
% Obviously, ScalarDiv function == “/”, ScalarDivBsFun == “*”, ScalarMultFun == “*”, and
% ScalarMultBsFun == “/”.
function quotient = ScalarDivFun(dividend, divisor)
  quotient = (dividend ./ divisor);
end
function dividend = ScalarDivBsFun(divisor, quotient)
  % Back solution of the scalar-division function (scalar multiplication)
  dividend = (quotient .* divisor);
end
function product = ScalarMultFun(multiplicand, multiplier)
  product = (multiplicand .* multiplier);
end
% Scalar multiplication is represented in the flow diagrams in three ways:
% 1) as a product operator (Pi notation) in a box (to compute the aggregate firing rate),
% 2) as a right-facing triangle (representing amplification), and
% 3) as a standard multiplication sign
function multiplicand = ScalarMultBsFun(multiplier, product)
  % Back solution of the scalar-multiplication function (scalar division)
  multiplicand = (product ./ multiplier);
end
~~~

### Functions composing the reward-mountain model

~~~
% Dummy function to relate the duration of the burst of increased firing in the directly stimulated neurons to
% the pulse frequency. We set the two to be equal, a reasonable assumption if frequency-following fidelity is
% high.
function Dburst = fD(Dtrain)
   Dburst = Dtrain;
end
% Function to compute the aggregate rate of firing required to produce a reward of half-maximal intensity
% see: Sonnenschein, B., Conover, K., & Shizgal, P. (2003).
% Growth of brain stimulation reward as a function of duration and stimulation strength.
% Behavioral Neuroscience, 117(5), 978–994. http://psycnet.apa.org/journals/bne/117/5/978
function FFaggHM = fH(C, D, RhoPi)
  FFaggHM = ScalarMultFun(…
       RhoPi,…
        (1 + ScalarDivFun(…
               C,…
               D…
              )…
       )…
      );
end
% Function to compute the average rate of firing required to produce a reward of half-maximal intensity
% see: Sonnenschein, B., Conover, K., & Shizgal, P. (2003).
% Growth of brain stimulation reward as a function of duration and stimulation strength.
% Behavioral Neuroscience, 117(5), 978–994. http://psycnet.apa.org/journals/bne/117/5/978
function FFhm = fHfiring(C, D, N, RhoPi, varargin)
  if size(varargin,2) == 0
    FFhm = ScalarDivFun(…
       fH(…
               C, …
               D, …
               RhoPi…
              ),…
        N…
      );
else
     Kin = varargin{1};
     FFhm = ScalarDivFun(…
            fH(…
               C, …
               D, …
               ScalarDivFun(…
                 RhoPi,…
                 Kin…
                  ),…
                )…
              N…
      );
  end
end
% Function to compute the average pulse frequency required to produce a reward of half-maximal intensity
% see: Sonnenschein, B., Conover, K., & Shizgal, P. (2003).
% Growth of brain stimulation reward as a function of duration and stimulation strength.
% Behavioral Neuroscience, 117(5), 978–994. http://psycnet.apa.org/journals/bne/117/5/978
function FpulseHM = FpulseHMfun(C, D, Fbend, FPmax, Fro, N, RhoPi, varargin)
  if size(varargin,2) == 0
    FpulseHM = FilterFunBS(…
         Fbend, …
         Fro, …
         fHfiring(C, …
            D, …
            N, …
            RhoPi),…
     FPmax…
    );
  else
    Kin = varargin{1};
    FpulseHM = FilterFunBS(…
         Fbend, …
         Fro, …
         fHfiring(C, …
           D, …
           N, …
           RhoPi,…
           Kin…
          ),…
      FPmax…
     );
  end
end
% Function to find the pulse frequency that produces a reward of half-maximal intensity given power growth of
% reward intensity
% FFmax, the maximal induced firing rate is obtained by setting the pulse frequency to 1000
function FPhmPG = find_FhmPG(FPmax, Fbend, Fro, g, Kin, Kout, RmaxPG)
  FFmax = FilterFun(FPmax,Fbend,Fro);
  FFhmPG = Kin .* ((0.5 .* RmaxPG) ./ Kout).^(1./g);
  FPhmPG = FilterFunBS(Fbend, Fro, FFhmPG, FPmax);
end
% Function to translate the effect of an electrical input to the directly stimulated mfb stage
% of the series circuit model into an equivalent optical input to the dopaminergic stage.
% Both inputs produce the same level of firing in the dopamine neurons.
function FoptEquiv = FoptEquivFun(Felec, FmfbBend, FmfbHM, FmfbRO, gMFB, KrgE)
   FoptEquiv = fRbsr(…
                Felec,…
                FmfbBend,…
                FmfbHM,…
                FmfbRO,…
                gMFB,…
                KrgE);
end
% Function to compute the equivalent optical drive on midbrain dopamine neurons
% due to trans-synaptic input from electrically excited MFB neurons
function FmfbDAdrive = FmfbDAdriveFun(FmfbDAbend, Felec, FFmfbDAmax, FoptEquiv, FmfbDAro)
   FmfbDAdrive = ScalarMultFun(…
          ScalarDivFun(…
                FilterFun(…
                    Felec,…
                    FmfbDAbend,…
                    FmfbDAro…
                   ),… % Filtered firing rate in MFB input
                FFmfbDAmax…
            ),… % Proportion of maximum firing rate achieved
           FoptEquiv…
          ); % Maximum equivalent optical pulse frequency applied to the dopamine neurons
end
% Find the electrical pulse frequency that produces half-maximal, summated reward intensity
% The solution is obtained by means of interpolation
%function [FsumHM, RsumMax] = find_FhmSum(Felec, Rsum)
function [FsumHM, RsumMax, CindAbove, CindBelow] = find_FhmSum(Felec, Rsum)
    RsumMax = max(Rsum, [], 2); % max along each row
    Rcrit = RsumMax./2;
    RsumHMlog = Rsum > Rcrit;
    RsumHMlogDiff = diff(RsumHMlog==1,1,2); % indices (offset by 1) marking transition from zero to one
% RsumHMlogDiff = diff(RsumHMlog); % indices (offset by 1) marking transition from zero to one
    siz = size(RsumHMlogDiff);
    IndDiffAbove = find(RsumHMlogDiff==1); % linear indices for first elements > Rcrit
    [Rind Cind] = ind2sub(siz,IndDiffAbove);
    % row & column indices for first elements > Rcrit offset by −1
    CindSort(Rind) = Cind; % column indices sorted by row number of Rsum matrix
    CindAbove = CindSort + 1; % row index of value just above Fh
    CindBelow = CindSort;
    % find exact Fhm by interpolation
    siz = size(Rsum); % one more column than diff matrix
    IndAbove = sub2ind(siz,1:length(Rind),CindAbove);
    IndBelow = IndAbove - siz(1);
    % The indices increment down the columns, so adjacent row indices are separated by the number of rows
    RsumAbove = Rsum(IndAbove);
    RsumBelow = Rsum(IndBelow);
    slp = (RsumAbove - RsumBelow) ./ (Felec(CindAbove) - Felec(CindBelow));
    FsumHM = (((Rcrit’ - RsumBelow) ./ slp) + Felec(1,CindBelow))’;
    % All rows of Felec are the same
    % Rcrit is a column vector but RsumBelow and Felec are row vectors
    % Return FsumHM as a column vector so that it has the same dimensions as RsumMax
end
% Function to compute the normalized value of the reward-growth function
% F is a pulse frequency, whereas FFhm is a firing frequency
function RbsrNorm = fRbsrNorm(F, Fbend, FFhm, Fro, g)
   RbsrNorm = LogistNormFun(…
      g,…
      FilterFun(…
         F,…
         Fbend,…
         Fro…
        ),…
     FFhm…
    );
end
% Function to compute the scaled value of the reward-growth function
% F is a pulse frequency, whereas Fhm is a firing frequency
function Rbsr = fRbsr(F, Fbend, Fhm, Fro, g, Krg)
   Rbsr = ScalarMultFun(…
          Krg,…
          LogistNormFun(…
              g,…
              FilterFun(…
                 F,…
                 Fbend,…
                 Fro…
                ),…
              Fhm…
             )…
         );
end
function Rbsr = fRbsrFull(C, D, F, Fbend, Fro, g, Krg, N, RhoPi, varargin)
   if size(varargin,2) == 0
      Rbsr = ScalarMultFun(…
              Krg,…
              LogistNormFun(…
                g,…
                FilterFun(…
                     F,…
                     Fbend,…
                     Fro…
                    ),…
                fHfiring(…
                     C, …
                     D, …
                     N, …
                     RhoPi…
                    )…
                )…
       );
   else
      Kin = varargin{1};
      Rbsr = ScalarMultFun(Krg,…
                     LogistNormFun(g,…
                         FilterFun(…
                             F,…
                             Fbend,…
                             Fro…
                            ),…
                    fHfiring(…
                           C, …
                           D, …
                           N, …
                           RhoPi,…
                           Kin…
                          )…
                   )…
            );
      end
end
% Back solution of the full reward-growth function for the series-circuit (sc) model
% KrgE translates the normalized output of the upstream reward-growth function into
% the equivalent of an optical pulse frequency applied to the dopamine neurons
function FscHM = FscHMbs(FhmDA, FmfbBend, FhmMFB, FmfbRO, FPmax, gMFB, KrgEq)
    FscHM = FilterFunBS(… % Electrical pulse frequency that produces HM reward intensity
             FmfbBend,…
             FmfbRO,…
             LogistNormBsFun(… % MFB firing-rate that produces HM reward intensity
                    gMFB,… % exponent of MFB RG function
                    FhmMFB,… % location parameter of MFB RG function
                    ScalarMultBsFun(… % normalized output of MFB RG function
                          KrgEq,…
                          FhmDA…
                         )… % normalized output of MFB RG function
                   ),… % MFB firing-rate that produces HM reward intensity
             FPmax… % pulse frequency sufficient to maximize firing
            ); % Electrical pulse frequency that produces HM reward intensity
end
% Power function to compute reward growth
% Kin scales the firings, and the product is raised to the power of g
function R = fRpg(F, Fbend, Fro, g, Kin, Kout)
        R = ScalarMultFun(…
             Kout,…
        ScalarDivFun(…
                          FilterFun(F,…
                               Fbend,…
                               Fro…
                              ),…
                          Kin…
                         )…
                          .^g…
       );
end
% Power function to compute reward growth, normalized to output values between 0 & 1
% A pulse frequency of 1000 is used to estimate Rmax
function Rnorm = RGpgNormFun(F, Fbend, Fro, g, Kin, Kout)
        Rnorm = ScalarDivFun(…
                         fRpg(F, Fbend, Fro, g, Kin, Kout),…
                         fRpg(1000, Fbend, Fro, g, Kin, Kout)…
                        );
end
% Dummy function to compute subjective probability (output = input)
% Breton, Y.-A., Conover, K., & Shizgal, P. (2014).
% The effect of probability discounting on reward seeking: a three-dimensional perspective.
% Frontiers in Behavioral Neuroscience, 8, 284. https://doi.org/10.3389/fnbeh.2014.00284
% found that subjective and objective reward probabilites in an eICSS experiment
% were indistiguishable when the objective probability was 0.5 or greater.
% Thus, subjective and objective probabilities are equated here.
% We do not know at what point this equivalence breaks down, so use of
% this function is invalid for objective probabilities <0.5.
function ProbSub = ProbSubFun(ProbObj)
     ProbSub = ProbObj;
end
% Function to compute subjective effort cost (output = input)
% This function is certainly not scalar! However, we haven’t measured it yet.
% We make the assumption that the rate of subjective exertion is held constant
% under the conditions of our experiment, and we thus impose a fixed value on the subjective rate of exertion.
% Due to factors such as fatigue, this can’t be entirely right. We hope
% that violations of this assumption aren’t serious.
% To accommodate the hypothesis that subjective effort costs are modulated by dopamine tone,
% the optional KeffMod parameter scales the subjective effort cost.
% We use this function to set the subjective rates of exertion for both work and leisure.
function dotPhiSubScaled = fphiSc(dotPhiSub, Kec, varargin)
     if size(varargin,2) == 0
        dotPhiSubScaled = ScalarMultFun(…
                         dotPhiSub,…
                         Kec…
                        );
    else
        KeffMod = varargin{1};
        dotPhiSubScaled = ScalarMultFun(…
                         dotPhiSub,…
                         ScalarDivFun(…
                             Kec,…
                             KeffMod…
                            )…
                        );
        end
end
% Dummy function to compute subjective value of alternate (“leisure”) activities (output = input)
% Again, we impose a fixed value, faute de mieux.
function RaaScaled = fRaa(Raa,Kaa)
    RaaScaled = ScalarMultFun(…
                      Raa,…
                      Kaa…
                     );
end
% Function to compute the subjective equivalent of the objective-price location parameter
function PsubE = PsubEfun(dotPhiObj, Kaa, Kec, Krg, Raa, pObj, RnormMax, varargin)
     if size(varargin,2) == 0
          PsubE = ScalarDivFun(…
                ScalarMultFun(…
                      ProbSubFun(pObj),…
                      ScalarMultFun(… % compute Rmax
                           Krg,…
                           RnormMax…
                          )…
                     ),…
                ScalarMultFun(…
                      fphiSc(dotPhiObj, Kec),…
                      fRaa(Raa, Kaa)…
                     )…
               );
   else
         KeffMod = varargin{1};
             PsubE = ScalarDivFun(…
                ScalarMultFun(…
                      ProbSubFun(pObj),…
                      ScalarMultFun(… % compute Rmax
                          Krg,…
                          RnormMax…
                         )…
                     ),…
                ScalarMultFun(…
                      fphiSc(dotPhiObj, Kec, KeffMod),…
                      fRaa(Raa, Kaa)…
                     )…
             );
      end
end
% Function to compute the subjective-price location parameter for the power-growth model
function PsubEpg = PsubEpgFun(dotPhiObj, Kaa, Kec, Krg, ObjAA, pObj, Rmax)
      PsubEpg = ScalarDivFun(…
                   ScalarMultFun(…
                         Rmax,…
                         ProbSubFun(pObj)…
                        ),…
                   ScalarMultFun(…
                         fphiSc(dotPhiObj, Kec),…
                         fRaa(ObjAA, Kaa)…
                        )…
              );
end
% Function to compute the objective-price location parameter
% This is the back-solution of the function that computes PsubE
function PobjE = PobjEfun(dotPhiObj, Kaa, Kec, Krg, ObjAA, pObj, PsubBend, PsubMin, RnormMax, varargin)
      if size(varargin,2) == 0
           PobjE = PsubBsFun(…
                   PsubEfun(dotPhiObj, Kaa, Kec, Krg, ObjAA, pObj, RnormMax), …
                   PsubBend, …
                   PsubMin…
                  );
   else
        KeffMod = varargin{1};
        PobjE = PsubBsFun(…
              PsubEfun(dotPhiObj, Kaa, Kec, Krg, ObjAA, pObj, RnormMax, KeffMod), …
              PsubBend, …
              PsubMin…
             );
     end
end
% Function to compute time allocation
% This function builds the mountain surface
% The values of the location parameters, FpulseHM & PobjE, are inputs
% These must be computed before this function is called
function T = TAfun(a, F, Fbend, FpulseHM, Fro, g, Pobj, PobjE, PsubBend, PsubMin, RnormMax)
     T = LogistNormFun(…
              a, …
              fRbsrNorm(…
                       F, …
                       Fbend, …
                       FpulseHM, …
                       Fro, …
                       g…
                      ), …
              ScalarDivFun(…
                         ScalarMultFun(…
                               PsubFun(Pobj, …
                                    PsubBend, …
                                    PsubMin),…
                                    RnormMax…
                              ),…
                         PsubFun(PobjE, …
                            PsubBend, …
                            PsubMin)…
                   )…
             );
end
% Function to compute time allocation using power growth of reward intensity
% This function builds the mountain surface
% The values of the location parameter, PsubEpg, is an input and must be computed before this function is called
function T = TApgFun(a, F, Fbend, Fro, g, Kin, Kout, Pobj, PsubBend, PsubEpg, PsubMin)
        T = LogistNormFun(…
                   a, …
                   RGpgNormFun(F, Fbend, Fro, g, Kin, Kout),…
                   ScalarDivFun(…
                          PsubFun(Pobj, …
                               PsubBend, …
                               PsubMin),…
                          PsubEpg…
                         )…
                );
end
% Function to compute time allocation for the convergence model
% The reward-growth function must be computed beforehand. Its output, RGvec, must be a column vector.
function T = TAsumFun(a, Pobj, PobjE, PsubBend, PsubMin, RGvec, RnormMax)
       T = LogistNormFun(…
                    a, …
                    RGvec, …
                    ScalarDivFun(…
                           ScalarMultFun(…
                                PsubFun(Pobj, …
                                      PsubBend, …
                                      PsubMin),…
                                      RnormMax…
                               ),…
                         PsubFun(PobjE, …
                             PsubBend, …
                             PsubMin)…
                )…
       );
end
~~~

### Functions that plot graphs and set attributes

~~~
% Plot of the frequency-following function
% function FF_graph = plot_freqFoll(F, FR, fnroot, graphs2files, figdir, varargin)
function FF_graph = plot_freqFoll(F, FR, fnroot, varargin)
   global graphs2files FigDir
   if size(varargin,2) == 0
      % To accommodate multiple plots based on input matrices, must compute {min, max} in 2 stages
      % Revise this section to operate in the log domain and then transform back to linear
      xmin = min(min(F)) - (0.1*(max(max(F))-min(min(F))));
      xmax = max(max(F)) + (0.1*(max(max(F))-min(min(F))));
      ymin = min(min(FR)) - (0.1*(max(max(FR))-min(min(FR))));
      ymax = max(max(FR)) + (0.1*(max(max(FR))-min(min(FR))));
   else
      xmin = varargin{1};
      xmax = varargin{2};
      ymin = varargin{3};
      ymax = varargin{4};
   end
      FF_graph = figure;
      FF_graph.Visible = ‘off’;
      FF_graph.Position = [0 0 600 600];
      gmfbfr = loglog(F,FR);
% gmfbfr.LineWidth = 4;
      arrayfun(@(x) set(x,’LineWidth’,4), gmfbfr); % Needed if plot includes multiple lines
      grid on;
      ax1 = gca;
      ax1.XLim = [xmin xmax];
      ax1.YLim = [ymin ymax];
      ax1.XLabel.String = ‘Pulse Frequency_{ }’;
      ax1.YLabel.String = ‘Firing Frequency_{ }’;
      ax1.FontSize = 24;
      ax1.XLabel.FontAngle = ‘italic’;
      ax1.YLabel.FontAngle = ‘italic’;
      arrayfun(@(x) set(ax1, ‘XTickLabel’, x), {num2str(ax1.XTick’)});
      arrayfun(@(x) set(ax1, ‘YTickLabel’, x), {num2str(ax1.YTick’)});
      if graphs2files
          saveas(gca,fullfile(FigDir,strcat(fnroot,’_freq_resp.fig’)));
          saveas(gca,fullfile(FigDir,strcat(fnroot,’_freq_resp.png’)));
    end
end
% Plot of the strength-duration function for trains
function SDT_graph = plot_StrDurTrains(Delec,FelecHM)
      global graphs2files FigDir
      SDT_graph = figure;
      SDT_graph.Visible = ‘off’;
      SDT_graph.Position = [0 0 600 600];
      gmfbfr = loglog(Delec,FelecHM);
      gmfbfr.LineWidth = 4;
      grid on;
      ax1 = gca;
      ymax = max(FelecHM)*10^0.1;
      ymin = min(FelecHM)*10^-0.1;
      ax1.YLim = [ymin ymax];
      ax1.XLabel.String = ‘Train Duration (s)_{ }’;
      ax1.YLabel.String = ‘F_{hm }’;
      ax1.FontSize = 24;
      ax1.XLabel.FontAngle = ‘italic’;
      ax1.YLabel.FontAngle = ‘italic’;
      ax1.XTickLabels = {‘0.1’, ‘1’, ‘10’, ‘100’};
      if graphs2files
          saveas(gca,fullfile(FigDir,’mfb_StrDurTrains.fig’));
          saveas(gca,fullfile(FigDir,’mfb_StrDurTrains.png’));
    end
end
% Plot one or more reward-growth functions
function out_graph = plot_RG(Fmat,RMat,pnam,pVec,fnam,TitleStr,linlog, varargin)
      global graphs2files FigDir
      fh = figure;
      fh.Position = [0 0 600 600];
      fh.Visible = ‘off’;
      plab = regexprep(regexprep(pnam,’\W’,’’),’_’,’’);
      switch linlog
           case ‘lin’ % linear y-axis, logarithmic x-axis
                gRG = semilogx(Fmat,RMat);
                ymin = −0.05;
                ymax = max(max(RMat)) * 1.05;
                if graphs2files
                if size(varargin,2) == 0
%                 figfile = fullfile(FigDir,char(strcat(‘RG_’,fnam’,’_’,’semilog.fig’)));
%                 pngfile = fullfile(FigDir,char(strcat(‘RG_’,fnam’,’_’,’semilog.png’)));
                figfile = fullfile(FigDir,char(strcat(‘RG_’,fnam,’_’,’semilog.fig’)));
                pngfile = fullfile(FigDir,char(strcat(‘RG_’,fnam,’_’,’semilog.png’)));
            else
                name_suffix = varargin{1};
                figfile = …
                   fullfile(FigDir,char(strcat(‘RG_’,fnam’,’_’,name_suffix,’_’,’semilog.fig’)));
                pngfile = …
                   fullfile(FigDir,char(strcat(‘RG_’,fnam’,’_’,name_suffix,’_’,’semilog.png’)));
           end
    end
case ‘log’ % logarithmic y-axis, logarithmic x-axis
     gRG = loglog(Fmat,RMat);
     ymin = 0.01;
     ymax = max(max(RMat)) * 10^0.1;
     if graphs2files
        if size(varargin,2) == 0
          figfile = fullfile(FigDir,char(strcat(‘RG_’,fnam’,’_’,’loglog.fig’)));
          pngfile = fullfile(FigDir,char(strcat(‘RG_’,fnam’,’_’,’loglog.png’)));
        else
          name_suffix = varargin{1};
          figfile = …
               fullfile(FigDir,char(strcat(‘RG_’,fnam’,’_’,name_suffix,’_’,’loglog.fig’)));
          pngfile = …
               fullfile(FigDir,char(strcat(‘RG_’,fnam’,’_’,name_suffix,’_’,’loglog.png’)));
            end
       end
  end
  for j = 1:length(gRG)
      gRG(j).LineWidth = 4;
  end
  grid on;
  ax1 = gca;
  ax1.Position = [0.175 0.15 0.75 0.75];
  ax1.XLim = [10 1000];
  ax1.YLim = [ymin ymax];
  ax1.XLabel.String = ‘Pulse Frequency_{ }’;
  ax1.YLabel.String = ‘Reward Intensity_{ }’;
  ax1.FontSize = 24;
  ax1.XLabel.FontAngle = ‘italic’;
  ax1.YLabel.FontAngle = ‘italic’;
  ax1.XTick = [1 10 100 1000];
  ax1.XTickLabels = {‘1’, ‘10’, ‘100’, ‘1000’};
  ax1.LineWidth = 1;
  switch linlog
       case ‘lin’
              XlabPos = ax1.XLabel.Position;
              XlabPos(2) = ax1.YLim(1) - ((ax1.YLim(2) - ax1.YLim(1)) * 0.075); % 7.5% below lower limit
              ax1.XLabel.Position = XlabPos;
       case ‘log’
              XlabPos = ax1.XLabel.Position;
              XlabPos(2) = 10^(log10(ax1.YLim(1)) - ((log10(ax1.YLim(2)) - log10(ax1.YLim(1)))*0.075)); % 7.5% below lower limit
              ax1.XLabel.Position = XlabPos;
  end
     YlabPos = ax1.YLabel.Position;
     YlabPos(1) = 10^(log10(ax1.XLim(1)) - ((log10(ax1.XLim(2)) - log10(ax1.XLim(1)))*0.1)); % 10% below lower limit
     ax1.YLabel.Position = YlabPos;
%   if min(pVec) < 0.01
%       formatstring = ‘%3.2e’;
%   else
%       formatstring = ‘%0.2f’;
%   end
     if min(pVec) < 10
          formatstring = ‘%3.2f’;
     else
          formatstring = ‘%3.0f’;
     end
% Make a cell array of the legend values and pass to the legend function
     lgnd = legend(arrayfun(@(x) sprintf(formatstring,x), pVec, ‘UniformOutput’, false),…
                 ‘Location’, ‘best’);
     lgnd.Title.String = strjoin(pnam,’\n’);
     title(ax1, TitleStr, ‘FontSize’, 30);
     out_graph = gcf;
     if graphs2files
        saveas(gca,figfile);
        saveas(gca,pngfile);
     end
end
% Plot of the subjective-price function
function SP_graph = plot_PsubFun(Pobj,Psub,graphs2files, figdir)
     SP_graph = figure;
     SP_graph.Visible = ‘off’;
     SP_graph.Position = [0 0 600 600];
     gPsub = loglog(Pobj,Psub);
     gPsub.LineWidth = 4;
     grid on;
     ax1 = gca;
     ax1.XLim = [0.1 100];
     ax1.YLim = [0.5 100];
     ax1.XTickLabels = {‘0.1’, ‘1’, ‘10’, ‘100’};
     ax1.YTickLabels = {‘1’, ‘10’, ‘100’};
     ax1.LineWidth = 1;
     ax1.XLabel.String = ‘Objective Price (s)_{ }’;
     ax1.YLabel.String = ‘Subjective Price (s)_{ }’;
     ax1.FontSize = 24;
     ax1.XLabel.FontAngle = ‘italic’;
     ax1.YLabel.FontAngle = ‘italic’;
     if graphs2files
        saveas(gca,fullfile(figdir,’Subj_price.fig’));
        saveas(gca,fullfile(figdir,’Subj_price.png’));
     end
end
% Function to plot a single mountain
% F is a logarithmically spaced vector of pulse-frequencies
% Pobj is a logarithmically spaced vector of prices
% T is a square matrix of time-allocation values computed with price as the row variable
% and pulse frequency as the column variable.
% Visible is a character vector (‘on’, or ‘off’) that determines whether or not the figure is displayed.
% The optional arguments are xmin, xmax, ymin, ymax.
function gh3d2 = plot_MTN(F, Pobj, T, Visible, mtn_root, title_str, …
     graphs2files, figdir, varargin)
     logF = log10(F);
     logP = log10(Pobj);
     [X,Y] = meshgrid(logP, logF);
     Z = T;
     if size(varargin,2) == 0
         xmin = 0;
         xmax = 2.6;
         ymin = 1;
         ymax = 3;
     else
         xmin = varargin{1};
         xmax = varargin{2};
         ymin = varargin{3};
         ymax = varargin{4};
     end
     gh3d = figure;
     gh3d.Visible = Visible;
     gh3d.Position = [0 0 600 600];
     surface(X, Y, Z, ‘CData’, Z,…
         ‘FaceLighting’,’gouraud’,…
         ‘FaceColor’,[0.5 0.5 0.5], ‘FaceAlpha’, 0.5,…
         ‘EdgeColor’, [0.5 0.5 0.5],…
         ‘SpecularColorReflectance’,0.5, ‘SpecularExponent’,4, ‘SpecularStrength’,1);
     light(‘Parent’,gca,’Position’,[10 −5 2]);
     view([40 30]);
     ax3d = findall(gca,’Type’,’Axes’);
     title(ax3d, title_str, ‘FontSize’, 36); ax3d.Position = [0.15, 0.15, 0.75, 0.75];
     ax3d.FontSize = 16;
     ax3d.XLabel.String = ‘Objective Price (s)’;
     ax3d.XLabel.FontSize = 22;
     ax3d.XLabel.Rotation = −27;
     ax3d.XLabel.Units = ‘Normalized’;
     ax3d.XLabel.Position = [0.325 −0.0625 0];
     ax3d.XAxis.Scale = ‘linear’;
     ax3d.XLim = [xmin,xmax];
     ax3d.XTick = [0, 0.5, 1, 1.5, 2, 2.5];
     ax3d.XTickLabel = {‘1.0’,’3.2’,’10’,’32’,’100’,’316’};
     ax3d.YLabel.String = ‘Pulse frequency (pps)’;
     ax3d.YLabel.FontSize = 22;
     ax3d.YLabel.Rotation = 32;
     ax3d.YLabel.Units = ‘Normalized’;
     ax3d.YLabel.Position = [0.85 0.05 0];
     ax3d.YAxis.Scale = ‘linear’;
     ax3d.YLim = [ymin,ymax];
     ax3d.YTick = [0, 0.5, 1, 1.5, 2, 2.5, 3];
     ax3d.YTickLabel = {‘1’,’3.2’,’10’,’32’, ‘100’, ‘316’, ‘1000’};
     ax3d.ZLim = [0,1.1];
     ax3d.ZLabel.String = ‘Time Allocation’;
     ax3d.ZLabel.FontSize = 22;
     ax3d.ZLabel.Rotation = 91;
     ax3d.ZLabel.Units = ‘Normalized’;
     ax3d.ZLabel.Position = [-0.125 0.475 0];
     ax3d.DataAspectRatio = [1 0.75 0.8];
     hold on;
     z_shim = 0.01;
     z_shim = Z + z_shim;
     contvec = 0.1:0.1:0.9;
     [∼, ch] = contour3(X, Y, z_shim, contvec);
     colormap(‘jet’);
     set(ch, ‘LineWidth’, 2);
     gh3d2 = figure(gcf);
     gh3d2.Visible = Visible;
     grid on;
     hold off;
     if graphs2files
        saveas(gca,fullfile(figdir,strcat(mtn_root,’.fig’)));
        saveas(gca,fullfile(figdir,strcat(mtn_root,’.png’)));
     end
end
% Function to plot a single contour graph
% F is a logarithmically spaced vector of pulse-frequencies.
% Pobj is a logarithmically spaced vector of prices.
% T is square matrix of time-allocation values computed with price as the row variable
% and pulse frequency as the column variable.
% Visible is a character vector (‘on’, or ‘off’) that determines whether or not the figure is displayed.
% The optional arguments are xmin, xmax, ymin, ymax.
function cont1 = plot_contour(F, Pobj, T, Pobj_e, Fhm, Visible, mtn_root, …
        title_str, annot_str, graphs2files, figdir, varargin)
     logF = log10(F);
     logP = log10(Pobj);
     [X,Y] = meshgrid(logP, logF);
     Z = T;
     if size(varargin,2) == 0
        xmin = 0;
        xmax = 2.6;
        ymin = 1;
        ymax = 3;
     else
        xmin = varargin{1};
        xmax = varargin{2};
        ymin = varargin{3};
        ymax = varargin{4};
     end
     contvec = 0.1:0.1:0.9;
     cont1 = figure;
     cont1.Visible = Visible;
     cont1.Position = [0 0 600 600];
     contour(X,Y,Z,contvec,’LineWidth’,1.25);
     colormap(‘jet’);
     hold on;
     axc1 = findall(gca,’Type’,’Axes’);
     title(axc1, title_str, ‘FontSize’, 36);
     axc1.Position = [0.15, 0.175, 0.825, 0.75];
     axc1.FontSize = 18;
     axc1.XLim = [xmin,xmax];
     axc1.XLabel.String = ‘Objective Price (s)’;
     axc1.XLabel.FontSize = 24;
     % Adjust position of X-axis label
     xlpos = axc1.XLabel.Position;
     xlpos(2) = ymin - ((ymax - ymin) * 0.1);
     axc1.XLabel.Position = xlpos;
     axc1.XTick = [−2, −1.5, −1, −0.5, 0, 0.5, 1, 1.5, 2, 2.5];
     axc1.XTickLabel = {‘‘,’’,’0.1’,’0.32’,’1.0’,’3.2’,’10’,’32’,’100’, ‘316’};
     axc1.YLim = [ymin,ymax];
     axc1.YLabel.String = ‘Pulse Frequency (pps)’;
     axc1.YLabel.FontSize = 24;
     % Adjust position of Y-axis label
     ylpos = axc1.YLabel.Position;
     ylpos(1) = xmin - ((xmax - xmin) * 0.125);
     axc1.YLabel.Position = ylpos;
     axc1.YTick = [0. 0.5, 1, 1.5, 2, 2.5, 3];
     axc1.YTickLabel = {‘1’, ‘3.2’, ‘10’, ‘31.6’, ‘100’, ‘316’, ‘1000’};
     FhmLine1 = line(‘XData’, axc1.XLim,’YData’, [log10(Fhm), log10(Fhm)]);
     FhmLine1.Color = ‘r’;
     FhmLine1.LineWidth = 4;
     FhmLine1.LineStyle = ‘--’;
     PeLine1 = line(‘XData’, [log10(Pobj_e), log10(Pobj_e)],’YData’, axc1.YLim);
     PeLine1.Color = ‘b’;
     PeLine1.LineWidth = 4;
     PeLine1.LineStyle = ‘--’;
     tb2 = annotation(‘textbox’);
     tb2.String = annot_str;
     tb2.FontSize = 18;
     tb2.Position = [0.7, 0.25, 0.15, 0.075];
     hold off;
     legend([FhmLine1, PeLine1], [{‘F_{hm }’}, {‘P_e ‘}], ‘Location’,’northwest’);
     if graphs2files
        saveas(gca,fullfile(figdir,strcat(mtn_root,’.fig’)));
        saveas(gca,fullfile(figdir,strcat(mtn_root,’.png’)));
     end
end
% Function to produce a bargraph showing changes in the location parameters
function bg = plot_bg(Fhm1, Fhm2, Pobj_e1, Pobj_e2, Visible, bg_root,…
        graphs2files, figdir, varargin)
     logFhmShift = log10(Fhm2) - log10(Fhm1);
     logPobjShift = log10(Pobj_e2) - log10(Pobj_e1);
     if size(varargin,2) == 0
        span = max(logFhmShift, logPobjShift) - min(logFhmShift, logPobjShift);
        ymax = max(logFhmShift, logPobjShift) + 0.2 * span;
        ymin = min(logFhmShift, logPobjShift) - 0.2 * span;
     else
        ymin = varargin{1};
        ymax = varargin{2};
     end
     bg = figure;
     bg.Visible = Visible;
     bg.Position = [0 0 600 800];
     bar(1,logFhmShift, ‘r’);
     hold on;
     bar(2,logPobjShift, ‘b’);
     hold off;
     ax = findall(gca,’Type’,’Axes’);
     ax.Position = [0.2 0.15 0.6 0.8]; %[left bottom width height]
     ax.XLim = [0.5,2.5];
     ax.YLim = [ymin, ymax];
     ax.YLabel.String = ‘Shift’;
     ylabpos = ax.YLabel.Position;
     ylabpos(1) = 0.15;
     ax.YLabel.Position = ylabpos;
     ax.XAxis.FontSize = 22;
     ax.YAxis.FontSize = 22;
     ax.YGrid = ‘on’;
     ax.XTick = [1,2];
     ax.XTickLabel = {‘log_{10 }(F_{hm })’, ‘log_{10 }(P_e)’};
     bh = findall(gcf,’Type’,’Bar’);
     bh(1).BarWidth = 0.6;
     bh(2).BarWidth = 0.6;
     if graphs2files
        saveas(gca,fullfile(figdir,strcat(bg_root,’.fig’)));
        saveas(gca,fullfile(figdir,strcat(bg_root,’.png’)));
     end
end
% Function to produce a bargraph showing changes in the location parameters
function bg = plot_bgStar(Fhm1, Fhm2, FhmStar1, FhmStar2, Pobj_e1, Pobj_e2, PobjEstar1, PobjEstar2, …
     Visible, bg_root, graphs2files, figdir, varargin)
     logFhmShift = log10(Fhm2) - log10(Fhm1);
     logPobjShift = log10(Pobj_e2) - log10(Pobj_e1);
     logFhmStarShift = log10(FhmStar2) - log10(FhmStar1); % These are the corrected values
     logPobjStarShift = log10(PobjEstar2) - log10(PobjEstar1); % These are the corrected values
     if size(varargin,2) == 0
        span = max(logFhmShift, logPobjShift) - min(logFhmShift, logPobjShift);
        ymax = max(logFhmShift, logPobjShift) + 0.2 * span;
        ymin = min(logFhmShift, logPobjShift) - 0.2 * span;
     else
        ymin = varargin{1};
        ymax = varargin{2};
     end
     bwidth = 0.6; % width of bars
     FhmStarX = [1-(bwidth/2), 1+(bwidth/2)];
     FhmStarY = [logFhmStarShift, logFhmStarShift];
     PeStarX = [2-(bwidth/2), 2+(bwidth/2)];
     PeStarY = [logPobjStarShift, logPobjStarShift];
     bg = figure;
     bg.Visible = Visible;
     bg.Position = [0 0 600 800];
     bar(1,logFhmShift, ‘r’);
     hold on;
     bar(2,logPobjShift, ‘b’);
     bh = findall(gcf,’Type’,’Bar’);
     bh(1).BarWidth = bwidth;
     bh(2).BarWidth = bwidth;
     % Add lines showing corrected location parameters
     plot(FhmStarX, FhmStarY, ‘Color’,’c’, ‘LineStyle’, ‘-.’, ‘LineWidth’,6);
     plot(PeStarX, PeStarY, ‘Color’,’c’, ‘LineStyle’, ‘-.’, ‘LineWidth’,6);
     hold off;
     ax = findall(gca,’Type’,’Axes’);
     ax.Position = [0.2 0.15 0.6 0.8]; %[left bottom width height]
     ax.XLim = [0.5,2.5];
     ax.YLim = [ymin, ymax];
     ax.YLabel.String = ‘Shift’;
     ylabpos = ax.YLabel.Position;
     ylabpos(1) = 0.15;
     ax.YLabel.Position = ylabpos;
     ax.XAxis.FontSize = 22;
     ax.YAxis.FontSize = 22;
     ax.YGrid = ‘on’;
     ax.XTick = [1,2];
     ax.XTickLabel = {‘log_{10 }(F_{hm })’, ‘log_{10 }(P_e)’};
     bh = findall(gcf,’Type’,’Bar’);
     bh(1).BarWidth = 0.6;
     bh(2).BarWidth = 0.6;
     if graphs2files
        saveas(gca,fullfile(figdir,strcat(bg_root,’.fig’)));
        saveas(gca,fullfile(figdir,strcat(bg_root,’.png’)));
     end
end
% Modify attributes of a 2D graph. handle_attribute and attribute_value can be cell arrays,
% in which case attribute value must have the same number of rows as the length of the handle
% and the same number of columns as the number on handle_attributes.
% The restrictions of the set command apply. For example, this function cannot modify legend attributes.
function out_graph = modify_2D_graph(in_graph, handle_type, handle_attribute, attribute_value, …
     gFileNam, graphs2files, figdir)
     gh = findall(in_graph, ‘Type’, handle_type);
     set(gh, handle_attribute, attribute_value);
     if strcmp(handle_attribute,’XLim’) || strcmp(handle_attribute,’YLim’)
        ax = findall(gcf, ‘Type’, ‘Axes’);
        XlabPos = ax.XLabel.Position;
        YlabPos = ax.YLabel.Position;
        LinLogX = ax.XAxis.Scale;
        switch LinLogX
           case ‘linear’
               XlabPos(1) = ax.XLim(1) + ((ax.XLim(2) - ax.XLim(1)) / 2);
               YlabPos(1) = ax.Xlim(1) - ((ax.XLim(2) - ax.XLim(1)) * 0.1); % 10% below lower limit
           case ‘log’
               XlabPos(1) = 10^(log10(ax.XLim(1)) + ((log10(ax.XLim(2)) - log10(ax.XLim(1))) / 2));
               YlabPos(1) = 10^(log10(ax.XLim(1)) - ((log10(ax.XLim(2)) - log10(ax.XLim(1)))*0.1)); % 10% below lower limit
        end
        ax.XLabel.Position = XlabPos;
        LinLogY = ax.YAxis.Scale;
        switch LinLogY
           case ‘linear’
               XlabPos(2) = ax.YLim(1) - ((ax.YLim(2) - ax.YLim(1)) * 0.0/075); % 7.5% below lower limit;
               YlabPos(2) = ax.YLim(1) + (ax.YLim(1) + ((ax.YLim(2) - ax.YLim(1)) / 2));
           case ‘log’
               XlabPos(2) = 10^(log10(ax.YLim(1)) - ((log10(ax.YLim(2)) - log10(ax.YLim(1)))*0.075)); % 7.5% below lower limit
               YlabPos(2) = 10^(log10(ax.YLim(1)) + ((log10(ax.YLim(2)) - log10(ax.YLim(1))) / 2));
        end
        ax.YLabel.Position = YlabPos;
     end
     out_graph = gcf;
     if graphs2files
        figfile = fullfile(figdir, strcat(gFileNam, ‘.fig’));
        saveas(gcf,figfile);
        pngfile = regexprep(figfile, ‘.fig’,’.png’);
        saveas(gcf,pngfile);
     end
end
~~~

### Functions that display multi-panel Matlab figures

~~~
function dual_sub = dual_subplot(g1, g2, dual_sub_out, graphs2files, figdir)
     dual_sub = figure;
     dual_sub.Visible = ‘off’;
     dual_sub.Units = ‘pixels’;
     dual_sub.Position = [0 0 1500 600];
     colormap ‘jet’;
     ax(1) = findall(g1, ‘Type’, ‘Axes’);
     ax(2) = findall(g2, ‘Type’, ‘Axes’);
     % Collect legend properties if a legend exists in the input figures
     for j = 1:2
        lg(j) = false;
        if ∼isempty(ax(j).Legend) lg(j) = true;
            lgh = ax(j).Legend;
            lgstr(j).str = lgh.String;
            lgstr(j).tstr = lgh.Title.String;
        end
     end
     for j = 1:2
        ax_copy(j) = copyobj(ax(j), dual_sub);
        sph(j) = subplot(1,2,j,ax_copy(j));
%             if ∼isempty(ax(j).Legend)
          if lg(j)
%               splg(j) = legend(sph(j),lgstr(j).str,’Location’,’southeast’);
              % This position is rigid, but it should work in this specific case.
              % The reward-growth graphs have legends; the mountain plots do not.
              splg(j) = legend(sph(j),lgstr(j).str,’Location’,’best’);
              splg(j).Title.String = lgstr(j).tstr;
            end
     end
     if graphs2files
            saveas(dual_sub,fullfile(figdir,strcat(dual_sub_out,’.fig’)));
            saveas(dual_sub,fullfile(figdir,strcat(dual_sub_out,’.png’)));
     end
end
% function to plot the quad-panel display of the contour and bar graphs
% cont1 is plotted twice, once in the upper left and once in the lower right
function quad_sub = quad_subplot(cont1, cont2, bg, quad_sub_out, bg_root, …
          graphs2files, figdir)
     quad_sub = figure; quad_sub.Visible = ‘off’;
     quad_sub.Units = ‘pixels’;
     quad_sub.Position = [0 0 1500 1500];
     colormap ‘jet’;
     fh(1) = cont1;
     fh(2) = bg;
     fh(3) = cont2;
     fh(4) = cont1;
     for j = 1:4
        ax(j) = findall(fh(j), ‘Type’, ‘Axes’);
     end
     % Collect textbox properties if a textbox exists in the input figure
     for j = 1:4
     tbh = findall(fh(j),’Type’,’TextBox’);
        tb(j) = false;
        if ∼isempty(tbh)
           tb(j) = true;
           tbstr(j) = tbh.String;
        end
     end
     bg_str = findall(ax(2), ‘-Property’, ‘FontSize’);
     bg_str = findall(bg_str(:),’-Property’,’FontSize’);
     for j = 1:length(bg_str)
        bg_str(j).FontSize = bg_str(j).FontSize * 1.75;
     end
     for j = 1:4
        ax_copy(j) = copyobj(ax(j), quad_sub);
        sph(j) = subplot(2,2,j,ax_copy(j));
        if tb(j)
           axpos = sph(j).Position;
           x_offset = axpos(3) * 0.7;
           y_offset = axpos(4) * −0.1;
           tbdim = [(axpos(1) + x_offset) (axpos(2) + y_offset) 0.1 0.1];
           ann(j) = annotation(‘textbox’, tbdim, ‘String’, tbstr(j),’FitBoxToText’,’on’);
           ann(j).FontSize = 16;
        end
     end
     if graphs2files
        saveas(quad_sub,fullfile(figdir,strcat(quad_sub_out,’.fig’)));
        saveas(quad_sub,fullfile(figdir,strcat(quad_sub_out,’.png’)));
     end
end
function out_fig = adjust_right_panel(dual_sub, h, v)
     out_fig = dual_sub;
     out_fig.Visible = ‘off’;
     ax = findall(out_fig,’Type’,’Axes’);
     ax1_pos = ax(1).Position;
     ax1_ip = ax(1).InnerPosition;
     ax1_op = ax(1).OuterPosition;
     ax(1).InnerPosition = ax(1).OuterPosition; % tight borders
     lgutter = ax1_op(1) - 0.5;
     panel_width = 0.5 - lgutter;
     panel_height = 1;
     new_ax_width = ax1_pos(3) * h; % rescale height
     new_ax_height = ax1_pos(4) * v; % rescale width
     new_hmargin = (panel_width - new_ax_width) / 2;
     new_vmargin = (panel_height - new_ax_height) / 2;
     ax1_newxpos = 0.5 + lgutter + new_hmargin;
     ax1_newpos(1)= ax1_newxpos;
     ax1_newvpos = new_vmargin; % unnecessary, but included for consistency
     ax1_newpos(2) = ax1_newvpos;
     ax1_newpos(3) = new_ax_width;
     ax1_newpos(4) = new_ax_height;
     ax(1).Title.String = ‘simulated data’;
     ax(1).Position = ax1_newpos;
end
~~~

### Functions that create & display images from stored files

~~~
function show_imported_graphic(gnam, mag, ImpFigDir)
     imshow(fullfile(ImpFigDir,gnam),’Border’,’tight’,’InitialMagnification’,mag);
end
function F = make_fig_from_png(png_file, mag)
     img = imread(png_file, ‘png’);
     sz = size(img);
     F = figure; image(img);
     pos = F.Position; % the conventional x and y dimensions are reversed pos(3) = sz(2) * mag;
     pos(4) = sz(1) * mag;
     F.Position = pos; axis tight;
     ah = findall(F,’Type’,’Axes’);
     ah.Visible = ‘off’;
     F.Visible = ‘off’;
end
~~~

